## Supplementary material for "CCL5 promotes breast cancer recurrence through macrophage recruitment in residual tumors": Figure 3 Source Data

Source Data Figure 3 - pg/mL protein expression of cytokines in primary and residual tumors

| Identifier | 1507-4L | 1507-4R | 1510-4L | 1510-4R | 1503-4L | 1503-4R |
|---|---|---|---|---|---|---|
| **Cytokine** | **Primary** | **Primary** | **Primary** | **Primary** | **Residual** | **Residual** |
| AR | 32.0607815 | 32.7198676 | 28.1006253 | 28.9267303 | 3.95134791 | 3.66448537 |
| Axl | 651.80807 | 986.443981 | 601.225216 | 925.582985 | 898.065698 | 366.801966 |
| CD27L | 79.5177683 | 17.0191643 | 98.0888993 | 20.4330608 | 76.7328845 | 15.0814708 |
| CD30 | 14.2977581 | 13.1497234 | 9.44225342 | 5.3235828 | 14.6036829 | 12.971881 |
| CD40 | 322.786106 | 249.568429 | 646.749837 | 275.246132 | 566.273079 | 429.967822 |
| CXCL16 | 18.7539786 | 16.4137751 | 125.784645 | 13.596758 | 15.5412176 | 12.1708182 |
| E-selectin | 31.3971661 | 27.3857538 | 34.9365191 | 18.2981481 | 30.9842764 | 30.5023802 |
| EGF | 14.795709 | 31.1298043 | 5.45217765 | 9.4970828 | 6.15343769 | 12.8284811 |
| Fractalkine | 406.9508 | 623.573942 | 1477.22806 | 503.508706 | 468.857894 | 182.807476 |
| GITR | 61.7591175 | 47.3421301 | 37.4193553 | 25.2288045 | 119.866628 | 255.188953 |
| GM-CSF | 1.63713866 | 0.946845 | 7.49597173 | 0.96718127 | 1.81029096 | NA |
| HGF | 100.772686 | 76.1834585 | 150.644942 | 75.7352335 | 170.487956 | 120.319408 |
| I-TAC | 2.16562815 | 11.9376755 | 15.6828001 | 12.0079814 | 18.0940353 | 20.4527728 |
| IFNg | 0.67944465 | 0.14100793 | 21.5989837 | 14.4818065 | 18.4138487 | NA |
| IGF-1 | 10363.7082 | 1501.8345 | 842.130439 | 144.634109 | 157219.857 | 277154.506 |
| IGFBP-2 | 217.075793 | 527.671294 | 338.307533 | 177.033838 | 212.402876 | 266.779861 |
| IGFBP-3 | 434.755123 | 4787.34452 | 8534.7905 | 10786.6545 | 1370.27892 | 409.908616 |
| IGFBP-5 | 289.098396 | 107.866513 | 329.635564 | 58.9965571 | 270.094941 | 78.4062881 |
| IGFBP-6 | 47.7724167 | 58.1726153 | 116.715403 | 33.8258224 | 584.732712 | 250.214845 |
| IL-10 | 6.7445882 | 13.526799 | 10.7346921 | 4.00925937 | 4.84715446 | NA |
| IL-12A | 6.94391438 | 10.9741066 | 9.28929983 | 4.77830753 | 6.8785694 | NA |
| IL-12p70 | 11.4581869 | 11.0898348 | 34.1694764 | 9.34088122 | 8.45076009 | 6.95953195 |
| IL-13 | 95.3370115 | 249.834405 | 202.977575 | 156.145114 | 255.734073 | NA |
| IL-17 | 1.1996594 | 1.91609719 | 1.72365145 | 1.92687411 | 1.76607188 | NA |
| IL-17E | 90.9623857 | 110.280058 | 265.431421 | 124.128867 | 68.1137084 | 101.483981 |
| IL-17F | 15.6260898 | 15.1294384 | 19.8504626 | 5.65549978 | 13.4990358 | 15.0863081 |
| IL-1a | 18.5047385 | 36.158668 | 30.1942851 | 20.2943938 | 17.0687951 | NA |
| IL-1b | 1.53521393 | 1.81453154 | 1.62415771 | 1.66299469 | 3.68843028 | NA |
| IL-1ra | 3524.43916 | 3199.28802 | 1516.96686 | 2897.41973 | 3901.63382 | 3274.32302 |
| IL-2 | 1.09506618 | 2.84342665 | 3.66234158 | 0.97296916 | 2.15649977 | NA |
| IL-2 Ra | 29.1177346 | 26.6521759 | 42.929386 | 17.3910152 | 8.61614884 | 26.6377603 |
| IL-20 | 18.6840261 | 30.4487896 | 43.8826189 | 17.5599691 | 18.5235125 | 20.5179384 |
| IL-23 | 112.002421 | 77.2238439 | 206.274931 | 78.852885 | 110.938765 | 71.449357 |
| IL-28 | 21.795572 | 29.5402579 | 42.4719079 | 27.8055098 | 14.3176347 | 21.0873137 |
| IL-3 | 0.14297744 | 0.23454826 | 0.21043605 | 0.22611133 | 0.32442175 | NA |
| IL-4 | 0.22667143 | 0.23468003 | 0.35659813 | 0.19871333 | 0.25868031 | NA |
| IL-5 | 1.04604055 | 0.76144347 | 4.25549934 | 3.18442345 | 1.65928368 | NA |
| IL-6 | 4.9039758 | 15.2670101 | 70.6117238 | 2.74435632 | 4.52779527 | NA |
| IL-9 | 60.1803765 | 109.910522 | 105.400757 | 78.5291642 | 113.364082 | NA |
| KC | 7.62172892 | 15.7248842 | 14.1472184 | 7.73185848 | 10.3036907 | NA |
| M-CSF | 3.72456512 | 7.41949182 | 9.02083734 | 2.73581689 | 2.98908685 | NA |
| MCP-1 | 3.72456512 | 7.41949182 | 9.02083734 | 2.73581689 | 2.98908685 | NA |

| | | | | | | |
|---|---|---|---|---|---|---|
| MDC | 190.191804 | 80.1197593 | 260.027802 | 59.7545446 | 175.40664 | 300.72815 |
| MIP-2 | 6.54521232 | 27.9710404 | 27.1895637 | 3.10032733 | 8.10001011 | 5.87990977 |
| MIP-3a | 1.90361843 | 4.33169046 | 2.17765786 | 2.02683282 | 2.19452883 | 1.61239217 |
| OPG | 128.080125 | 64.0728827 | 93.2827461 | 104.368944 | 1948.71399 | 3490.26021 |
| OPN | 2742.13604 | 3759.92268 | 7908.55218 | 5233.53861 | 2144.3341 | 1750.87307 |
| P-selectin | 1251.59265 | 1191.45227 | 572.51044 | 221.576626 | 2267.27288 | 1231.10403 |
| Pro-MMP-9 | 6785.78047 | 7966.85567 | 424.668879 | 4375.69802 | 1073.0975 | 517.161353 |
| Prolactin | 31.8530103 | 22.843987 | 38.3101504 | 25.6061844 | 48.7975766 | 30.482195 |
| RANTES | 2.95564727 | 4.89584212 | 2.27977934 | 2.0413161 | 3.04394723 | NA |
| Resistin | 1517.99686 | 710.983589 | 51.7068015 | 19.0246794 | 1275.42211 | 308.037327 |
| SCF | 20.2772296 | 25.6138473 | 80.7402194 | 34.4496073 | 24.9094333 | 31.3375025 |
| SDF-1a | 35.5398386 | 50.3107358 | 36.3283935 | 37.6400276 | 90.7204121 | 42.9792637 |
| TNFa | 0.16520154 | 0.20655112 | 0.21849986 | 0.37586856 | 0.42850112 | NA |
| TPO | 108.64762 | 93.1257735 | 120.926013 | 82.4712092 | 129.817735 | 74.1104201 |
| VCAM-1 | 1919.11784 | 1625.46398 | 1465.37134 | 491.935574 | 6664.93947 | 4590.42781 |
| VEGF | 15.1571793 | 22.878486 | 106.847725 | 34.2661129 | 9.77993099 | 10.1720341 |
| VEGF-D | 5.56992926 | 20.2067539 | 32.520791 | 25.7092522 | 7.29773927 | 14.3323805 |

| 1505-4L | 1505-4R | 1511-4L | 1511-4R | 1502-4L | 1502-4R | 1504-4L |
|---|---|---|---|---|---|---|
| **Residual** | **Residual** | **Residual** | **Residual** | **Residual** | **Residual** | **Residual** |
| 4.78056437 | 4.37798062 | 4.04670314 | 10.7317451 | 2.71857135 | 1.98154478 | 3.44973239 |
| 587.121743 | 429.630107 | 726.276889 | 558.753207 | 641.652169 | 382.109284 | 582.133924 |
| 86.9984319 | 72.7440852 | 42.0264028 | 76.3989339 | 54.4149074 | 14.0982759 | 80.8672972 |
| 11.2402331 | 9.24511191 | 15.1904523 | 9.39574022 | 13.9152453 | 10.1238605 | 17.7766808 |
| 307.760843 | 132.675889 | 421.270528 | 194.698277 | 358.78277 | 156.748833 | 218.923733 |
| 27.4184564 | 13.1865982 | 3.90179611 | 14.1175669 | 18.1306606 | 8.60806029 | 16.9467937 |
| 41.0610331 | 53.1811157 | 34.1081307 | 45.3913636 | 31.3881822 | 28.5090755 | 61.3174635 |
| 20.4632508 | 20.2971288 | 13.0677916 | 60.1676132 | 3.08748058 | 14.8940873 | 23.7972661 |
| 925.089578 | 1159.06337 | 571.475558 | 1765.25251 | 422.528999 | 353.98797 | 574.931885 |
| 46.1607972 | 36.3591945 | 123.791469 | 121.735869 | 61.5342072 | 32.6924274 | 114.073643 |
| 0.09855255 | 7.64057571 | 3.56744667 | NA | NA | NA | NA |
| 463.328059 | 248.620178 | 219.745516 | 332.887143 | 240.25832 | 121.959186 | 328.029798 |
| 16.6658939 | 3.35635417 | 8.15028138 | 61.8861854 | 85.9150887 | 23.0471427 | 43.7549505 |
| 3.5453726 | NA | 12.3353731 | NA | NA | NA | NA |
| 353581.795 | 187972.056 | 3397153.66 | 879282.421 | 80279.6323 | 55962.2038 | 2175731.48 |
| 247.777998 | 236.139964 | 191.017468 | 238.790498 | 129.50811 | 169.336552 | 119.910442 |
| 683.625735 | 3023.15166 | 424.539756 | 354.29747 | 393.321644 | 333.575385 | 492.871505 |
| 561.346376 | 361.067911 | 119.773431 | 308.642796 | 345.943833 | 255.556948 | 143.480304 |
| 666.515465 | 684.422887 | 427.959557 | 758.056041 | 472.346604 | 370.206369 | 810.805302 |
| 9.98399716 | 13.3466789 | 9.21157957 | NA | NA | NA | NA |
| 9.67539214 | 13.8330953 | 3.94750005 | NA | NA | NA | NA |
| 20.6599547 | 16.7962384 | 15.4526389 | 14.675028 | 10.3288157 | 9.10970339 | 6.64457305 |
| 516.757432 | 683.197018 | 274.95815 | NA | NA | NA | NA |
| 1.22805266 | 3.0177997 | 2.23908462 | NA | NA | NA | NA |
| 246.33341 | 129.234885 | 71.4674587 | 266.030902 | 83.0974095 | 80.3990325 | 290.440176 |
| 18.3242145 | 20.86456 | 19.0710905 | 51.6477458 | 16.7965649 | 21.2775532 | 48.4694557 |
| 45.7354819 | 55.4927199 | 33.1893106 | NA | NA | NA | NA |
| 0.73923568 | 2.40170448 | 2.29973835 | NA | NA | NA | NA |
| 2350.56842 | 1189.49376 | 5924.53863 | 3767.87706 | 711.441483 | 158.793871 | 5961.33035 |
| 2.27845964 | 0.57157449 | 3.72508635 | NA | NA | NA | NA |
| 16.5644824 | 16.2021805 | 10.1771486 | 34.7698991 | 21.1901594 | 8.0534159 | 25.3395066 |
| 19.4930916 | 11.5801687 | 25.7671579 | 27.4406436 | 17.0674466 | 20.5886167 | 49.0624812 |
| 244.561934 | 220.872991 | 95.0739131 | 220.458653 | 128.935 | 104.149541 | 97.4125157 |
| 29.4395834 | 25.9935013 | 18.6287646 | 16.0267003 | 20.8139995 | 29.015606 | 14.8148619 |
| 0.35355251 | 0.81594434 | 0.27086241 | NA | NA | NA | NA |
| 0.44894378 | 0.26344623 | 0.51379938 | NA | NA | NA | NA |
| 2.08632747 | 0.37441792 | 1.50386298 | NA | NA | NA | NA |
| NA | 0.43952708 | 4.6209603 | NA | NA | NA | NA |
| 124.590197 | 165.500292 | 86.9382231 | NA | NA | NA | NA |
| 30.3177256 | 43.4973439 | 17.0832739 | NA | NA | NA | NA |
| 9.56190334 | 10.3252348 | 4.77959955 | NA | NA | NA | NA |
| 9.56190334 | 10.3252348 | 4.77959955 | NA | NA | NA | NA |

| | | | | | | |
|---|---|---|---|---|---|---|
| 40.7866643 | 22.9170725 | 26.6952221 | 18.9328239 | 105.483926 | 86.206069 | 160.600207 |
| 31.0397088 | 12.2607642 | 19.3103753 | 78.7083571 | 32.3369466 | 27.9168863 | 28.515306 |
| 1.67014564 | 2.28956199 | 1.11119794 | 2.69637313 | 10.9844712 | 6.16980199 | 8.32522098 |
| 1741.73367 | 2570.05248 | 1249.97967 | 148.017565 | 4153.28925 | 4119.95321 | 3753.92213 |
| 3445.66194 | 2927.11037 | 2673.34036 | 1731.42659 | 3230.01554 | 2106.06401 | 3808.16684 |
| 3451.10209 | 3915.4621 | 3451.83267 | 3234.65256 | 2284.77095 | 1574.75716 | 2269.64251 |
| 1420.14853 | 867.359353 | 1272.74291 | 586.122775 | 663.311755 | 218.97699 | 62792.105 |
| 54.2841742 | 32.024257 | 38.3413074 | 135.531683 | 38.03023 | 45.3633271 | 40.6132687 |
| 8.95548277 | 10.2780852 | 4.96768239 | NA | NA | NA | NA |
| 3213.10609 | 4263.38832 | 3490.23049 | 3900.74318 | 2130.60733 | 2783.95971 | 2170.03009 |
| 55.0458701 | 66.3332312 | 28.3841324 | 58.3538668 | 23.7306746 | 24.7391485 | 60.1646832 |
| 27.8467348 | 36.1499348 | 66.344283 | 82.1056468 | 16.6322638 | 11.7912041 | 110.996815 |
| 0.02172824 | 2.4426465 | 4.26380224 | NA | NA | NA | NA |
| 69.5814305 | 65.1111063 | 90.5705873 | 124.833028 | 172.705877 | 122.42397 | 81.4819085 |
| 3409.56119 | 3868.20237 | 4189.22618 | 3221.53436 | 6432.65083 | 2568.50985 | 8781.40217 |
| 77.8265016 | 40.0173971 | 19.6548591 | 140.711933 | 67.2194224 | 60.2839433 | 46.3700812 |
| 25.5062717 | 11.1582347 | 29.1901301 | 48.3919213 | 12.4572647 | 12.0464456 | 33.9788706 |

| 1504-4R Residual | 1509-4L Residual | 1509-4R Residual | Fold Change | p-value | log2(Fold Change) | -LOG10(p-value) |
|---|---|---|---|---|---|---|
| 3.20826947 | 4.38335174 | 4.02553686 | 0.14043914 | 7.0835E-12 | -2.831983062 | 11.1497532 |
| 373.851437 | 624.025784 | 328.611632 | 0.68445604 | 0.02797175 | -0.546970205 | 1.553280312 |
| 10.5223752 | 6.78636068 | 21.8586095 | 0.86570091 | 0.71700685 | -0.208059426 | 0.144476694 |
| 11.2414152 | 11.3644763 | 13.8625775 | 1.19181469 | 0.25692407 | 0.253159938 | 0.590195201 |
| 180.452562 | 258.261331 | 188.817373 | 0.76167607 | 0.30994898 | -0.392750532 | 0.508709784 |
| 18.4746836 | 10.0233828 | 9.71441864 | 0.32127426 | 0.06784612 | -1.6381227 | 1.168474966 |
| 42.8103575 | 28.4922637 | 54.5308944 | 1.43512148 | 0.06658043 | 0.521172863 | 1.176653388 |
| 26.5388369 | 2.37451985 | 23.1436335 | 1.24196781 | 0.66770324 | 0.312627778 | 0.175416518 |
| 264.82661 | 417.640763 | 497.503425 | 0.84172542 | 0.65862382 | -0.248578416 | 0.181362569 |
| 64.5307009 | 29.1900619 | 86.6525871 | 2.11893315 | 0.16368426 | 1.083338071 | 0.785993079 |
| NA | NA | NA | 1.18735436 | 0.82689689 | 0.247750566 | 0.082548643 |
| 281.687758 | 142.225208 | 147.45126 | 2.32808176 | 0.02678651 | 1.219141724 | 1.572083888 |
| 18.0499955 | 19.5697449 | 21.1190382 | 2.71219784 | 0.16857258 | 1.43946242 | 0.773213057 |
| NA | NA | NA | 1.23914867 | 0.77254826 | 0.309349287 | 0.112074381 |
| 60703.0478 | 424466.338 | 1462538.97 | 246.701385 | 0.16348944 | 7.946622005 | 0.786510301 |
| 228.664119 | 158.598858 | 275.457938 | 0.65455317 | 0.04647101 | -0.611417706 | 1.3328179 |
| 1364.1843 | 1102.23912 | 405.272894 | 0.14066519 | 0.0010215 | -2.829662756 | 2.990762531 |
| 112.656507 | 203.866278 | 183.833434 | 1.24943999 | 0.54220201 | 0.321281615 | 0.265838875 |
| 346.218348 | 835.378477 | 513.33207 | 8.7336566 | 0.00019724 | 3.126585807 | 3.704995671 |
| NA | NA | NA | 1.0678009 | 0.83553701 | 0.094642669 | 0.07803431 |
| NA | NA | NA | 1.073437 | 0.82227906 | 0.102237521 | 0.084980769 |
| 7.48078236 | 10.0330558 | 8.9336987 | 0.68386369 | 0.20265493 | -0.548219304 | 0.693242832 |
| NA | NA | NA | 2.45727837 | 0.05452437 | 1.297061302 | 1.263409321 |
| NA | NA | NA | 1.21943021 | 0.40661745 | 0.286207191 | 0.390813983 |
| 60.0882686 | 97.5247447 | 68.5800831 | 0.88173484 | 0.72471822 | -0.181583226 | 0.139830818 |
| 15.7367441 | 15.8458704 | 16.4276308 | 1.61772449 | 0.22411421 | 0.693965928 | 0.649530606 |
| NA | NA | NA | 1.44064007 | 0.25902332 | 0.526709936 | 0.586661138 |
| NA | NA | NA | 1.3755084 | 0.34387542 | 0.459964954 | 0.463598871 |
| 4617.90421 | 1000.20881 | 1709.56284 | 1.03451617 | 0.92832989 | 0.04895619 | 0.032297666 |
| NA | NA | NA | 1.01840685 | 0.96733526 | 0.026314023 | 0.014422982 |
| 13.3181981 | 6.90452845 | 16.5975483 | 0.58681606 | 0.03747247 | -0.769019752 | 1.426287666 |
| 20.0642952 | 20.6515061 | 22.5659313 | 0.82394089 | 0.41075619 | -0.279387252 | 0.386415878 |
| 94.0643721 | 92.6652464 | 111.805629 | 1.11898684 | 0.68951251 | 0.162193067 | 0.161457852 |
| 24.7537227 | 19.6600753 | 20.2688787 | 0.69844541 | 0.02041793 | -0.517780744 | 1.689988281 |
| NA | NA | NA | 2.16784102 | 0.1122318 | 1.116258961 | 0.949884086 |
| NA | NA | NA | 1.46053296 | 0.16380825 | 0.546494918 | 0.785664231 |
| NA | NA | NA | 0.60815882 | 0.36258153 | -0.717479954 | 0.440594325 |
| NA | NA | NA | 0.13669173 | 0.33529782 | -2.871002088 | 0.474569274 |
| NA | NA | NA | 1.38520891 | 0.14074161 | 0.470103575 | 0.851577483 |
| NA | NA | NA | 2.23771123 | 0.1171879 | 1.162023871 | 0.93111724 |
| NA | NA | NA | 1.20764043 | 0.62846958 | 0.272190956 | 0.201715738 |
| NA | NA | NA | 1.20764043 | 0.62846958 | 0.272190956 | 0.201715738 |

| | | | | | | |
|---|---|---|---|---|---|---|
| 141.03981 | 75.2378447 | 35.6469467 | 0.67202941 | 0.34833502 | -0.573403734 | 0.458002865 |
| 25.3977085 | 11.9697003 | 25.2939311 | 1.5776776 | 0.38283083 | 0.65780242 | 0.416993098 |
| 15.7139694 | 2.07554556 | 2.2254689 | 1.82215113 | 0.38528841 | 0.865642623 | 0.414214052 |
| 2190.85927 | 3269.87214 | 1677.24605 | 25.9222971 | 0.00198994 | 4.696121665 | 2.701160348 |
| 1962.6842 | 1498.4415 | 1695.30297 | 0.49163784 | 0.00376029 | -1.024332145 | 2.424778467 |
| 2340.97796 | 2400.28082 | 2996.69703 | 3.23522519 | 0.00090931 | 1.693866134 | 3.041289731 |
| 10243.2791 | 1311.68732 | 9950.6607 | 1.54991799 | 0.77270663 | 0.632191877 | 0.111985363 |
| 34.8794979 | 29.1829029 | 32.697147 | 1.57438054 | 0.27402909 | 0.654784297 | 0.562203326 |
| NA | NA | NA | 2.23824258 | 0.08228521 | 1.162366401 | 1.084678201 |
| 2739.74059 | 2467.70133 | 3997.9215 | 4.74565062 | 0.00389954 | 2.24660589 | 2.408986247 |
| 30.0531129 | 22.2280171 | 24.5111286 | 0.93077617 | 0.81045378 | -0.103493814 | 0.091271747 |
| 88.0609358 | 14.6491045 | 30.8718351 | 1.29135345 | 0.52145104 | 0.368883926 | 0.28278646 |
| NA | NA | NA | 7.40764102 | 0.16578228 | 2.889014186 | 0.780461894 |
| 117.990475 | 65.2357625 | 80.0441062 | 0.98222523 | 0.92092554 | -0.025874213 | 0.035775481 |
| 4692.14284 | 6018.66248 | 4622.98455 | 3.57817996 | 0.00168381 | 1.839225949 | 2.773706321 |
| 40.6093307 | 39.9873839 | 31.8777388 | 1.08756569 | 0.85803876 | 0.121102541 | 0.066493093 |
| 25.7310865 | 4.02210181 | 29.8503646 | 1.00770824 | 0.98274501 | 0.011078 | 0.007559152 |
