## Supplementary material for "CCL5 promotes breast cancer recurrence through macrophage recruitment in residual tumors": Figure 6 Source Data 1

| Entrez | Symbol | baseMean | log2FoldChai | lfcSE | stat | pvalue |
|---|---|---|---|---|---|---|
| 12550 | CDH1 | 4861.07602 | -9.5604968 | 0.22971772 | -41.618455 | 0 |
| 192897 | ITGB4 | 3887.37157 | -9.5575724 | 0.23007633 | -41.540876 | 0 |
| 19268 | PTPRF | 5146.98838 | -8.7192055 | 0.15473271 | -56.350112 | 0 |
| 12822 | COL18A1 | 7932.88747 | -7.1977722 | 0.1276078 | -56.405426 | 0 |
| 11501 | ADAM8 | 1804.88923 | -6.9035043 | 0.14499333 | -47.612565 | 0 |
| 27428 | SHROOM3 | 4354.6161 | -5.202511 | 0.09120314 | -57.043113 | 0 |
| 56615 | MGST1 | 1260.82559 | -4.6865999 | 0.09288435 | -50.456293 | 0 |
| 16480 | JUP | 5623.38192 | -3.3685862 | 0.07466912 | -45.113509 | 0 |
| 68682 | SLC44A2 | 4098.37135 | -2.9186268 | 0.05818952 | -50.157262 | 0 |
| 19272 | PTPRK | 4117.81288 | -2.6243152 | 0.05589871 | -46.947689 | 0 |
| 235505 | CD109 | 6642.81817 | 2.85042173 | 0.06206195 | 45.928653 | 0 |
| 13823 | EPB41L3 | 3402.27131 | 2.86322068 | 0.05276115 | 54.267595 | 0 |
| 66180 | P3H4 | 1456.83432 | 5.63114881 | 0.13270337 | 42.4341068 | 0 |
| 66854 | TRIM35 | 2920.65802 | 7.08021153 | 0.16024049 | 44.1849106 | 0 |
| 14178 | FGF7 | 4368.70621 | 7.75119464 | 0.19204675 | 40.3609784 | 0 |
| 16949 | LOXL1 | 3446.11354 | 8.19439124 | 0.20991286 | 39.0371086 | 0 |
| 78560 | ADGRA2 | 2801.95719 | 8.47452002 | 0.22018335 | 38.4884696 | 0 |
| 17387 | MMP14 | 7457.87184 | 11.0461502 | 0.2860003 | 38.6228621 | 0 |
| 12843 | COL1A2 | 15551.1413 | 11.4138114 | 0.21982662 | 51.9218802 | 0 |
| 17082 | IL1RL1 | 47941.3721 | 11.773132 | 0.14839128 | 79.3384383 | 0 |
| 12835 | COL6A3 | 14314.1977 | 12.0938573 | 0.30768717 | 39.3056922 | 0 |
| 54403 | SLC4A4 | 3452.72686 | -2.9039342 | 0.08114806 | -35.785627 | 1.85E-280 |
| 108735 | SFT2D2 | 1968.57366 | -4.3390261 | 0.12151841 | -35.706739 | 3.11E-279 |
| 67313 | 5730559C18I | 623.476967 | -4.0764402 | 0.11546867 | -35.30343 | 5.20E-273 |
| 53322 | NUCB2 | 1124.18669 | -2.3225723 | 0.06649247 | -34.929857 | 2.62E-267 |
| 22329 | VCAM1 | 796.766244 | 4.65638018 | 0.13649059 | 34.1150274 | 4.42E-255 |
| 319944 | TAF2 | 3269.95485 | 1.4538521 | 0.0427352 | 34.0200113 | 1.13E-253 |
| 71704 | ARHGEF3 | 1383.01955 | -3.7259875 | 0.10964093 | -33.983544 | 3.90E-253 |
| 213742 | Xist | 16880.0823 | -13.03547 | 0.39247549 | -33.213463 | 6.88E-242 |
| 269356 | SLC4A11 | 982.902251 | -4.6057563 | 0.13951563 | -33.012475 | 5.38E-239 |
| 12737 | CLDN1 | 3663.03804 | -7.7584842 | 0.235222 | -32.983667 | 1.39E-238 |
| 16478 | jund | 3475.35848 | -2.2185391 | 0.06817405 | -32.542282 | 2.69E-232 |
| 70737 | CGN | 1455.82314 | -8.3251244 | 0.25686306 | -32.41075 | 1.94E-230 |
| 142980 | TLR3 | 2005.52388 | -4.587316 | 0.14404388 | -31.846658 | 1.46E-222 |
| 21857 | TIMP1 | 12060.8743 | 1.98506732 | 0.06251713 | 31.7523743 | 2.94E-221 |
| 16456 | F11R | 2309.75009 | -6.6728962 | 0.21192418 | -31.487187 | 1.30E-217 |
| 107765 | ANKRD1 | 3895.44496 | -5.5241939 | 0.17815728 | -31.007399 | 4.28E-211 |
| 20198 | S100A4 | 12366.7232 | 3.23248821 | 0.10426913 | 31.0013924 | 5.16E-211 |
| 110310 | KRT7 | 10128.6738 | -12.490927 | 0.40954841 | -30.499269 | 2.66E-204 |
| 103963 | RPN1 | 10936.2668 | 0.79380476 | 0.02626196 | 30.2264071 | 1.07E-200 |
| 12443 | CCND1 | 41605.3531 | -1.1018228 | 0.03650411 | -30.183525 | 3.90E-200 |
| 56431 | DSTN | 12235.7135 | -1.8294208 | 0.06090359 | -30.037978 | 3.13E-198 |
| 20204 | PRRX2 | 1351.01998 | 7.86458293 | 0.26280439 | 29.9256147 | 9.14E-197 |

| | | | | | | |
|---|---|---|---|---|---|---|
| 13052 | CXADR | 1826.4733 | -2.9595397 | 0.0990993 | -29.864385 | 5.71E-196 |
| 105732 | fam83h | 1526.75914 | -2.9312148 | 0.1000561 | -29.295712 | 1.18E-188 |
| 11529 | ADH7 | 1806.26995 | 2.90571691 | 0.09982851 | 29.1070835 | 2.92E-186 |
| 243548 | PRICKLE2 | 1321.72377 | 8.3178865 | 0.28867131 | 28.8143858 | 1.42E-182 |
| 230752 | EVA1B | 588.941546 | 4.38675814 | 0.15273876 | 28.7206609 | 2.11E-181 |
| 69581 | RHOU | 2332.98937 | -2.4068093 | 0.08397127 | -28.662297 | 1.13E-180 |
| 353499 | TMC4 | 1000.31025 | -6.181134 | 0.21699486 | -28.485163 | 1.79E-178 |
| 13649 | EGFR | 864.615996 | -6.0598525 | 0.21288488 | -28.465397 | 3.14E-178 |
| 14314 | FSTL1 | 10751.9428 | 12.8515381 | 0.45279195 | 28.3828765 | 3.29E-177 |
| 14302 | FRK | 1341.59902 | -1.603136 | 0.05693184 | -28.158866 | 1.87E-174 |
| 234593 | NDRG4 | 1425.3777 | -2.8636482 | 0.10220655 | -28.018245 | 9.74E-173 |
| 26434 | PRND | 627.84185 | -6.0016637 | 0.2146307 | -27.962746 | 4.61E-172 |
| 16819 | LCN2 | 34254.0328 | -7.6309919 | 0.27460171 | -27.789309 | 5.84E-170 |
| 20843 | STAG2 | 2094.48985 | -3.6130607 | 0.1302063 | -27.74874 | 1.80E-169 |
| 109222 | RARRES1 | 647.658032 | 7.19911106 | 0.25964554 | 27.726689 | 3.33E-169 |
| 72599 | PDIA5 | 633.707669 | 2.57372969 | 0.09297743 | 27.681232 | 1.17E-168 |
| 17069 | LY6E | 17709.9497 | -3.4248091 | 0.12401527 | -27.616027 | 7.14E-168 |
| 64143 | RALB | 3883.15835 | -1.2472871 | 0.04549972 | -27.413074 | 1.92E-165 |
| 54161 | COPG1 | 9057.1017 | 0.82259684 | 0.03005712 | 27.3677895 | 6.63E-165 |
| 11981 | ATP9A | 1767.85834 | -9.6910136 | 0.3541573 | -27.363585 | 7.44E-165 |
| 242620 | DMRTA2 | 563.717502 | 6.78405791 | 0.24817892 | 27.3353511 | 1.61E-164 |
| 12035 | BCAT1 | 4564.14837 | 6.3077898 | 0.23132452 | 27.2681419 | 1.01E-163 |
| 19088 | PRKAR2B | 3287.31731 | 2.5672346 | 0.09505329 | 27.0083717 | 1.18E-160 |
| 14230 | FKBP10 | 3649.85142 | 11.2925948 | 0.41926392 | 26.9343349 | 8.70E-160 |
| 233765 | PLEKHA7 | 511.20641 | -5.0608334 | 0.18816739 | -26.89538 | 2.49E-159 |
| 100342 | FAM46B | 926.188373 | -5.0511815 | 0.18824196 | -26.833451 | 1.32E-158 |
| 14605 | TSC22D3 | 908.524288 | -2.9610501 | 0.11084213 | -26.714122 | 3.23E-157 |
| 20354 | SEMA4D | 617.935976 | -7.1233543 | 0.26665809 | -26.713438 | 3.29E-157 |
| 12490 | CD34 | 3666.31369 | 7.38913588 | 0.27668027 | 26.7064074 | 3.97E-157 |
| 56401 | P3H1 | 1200.51345 | 2.4650194 | 0.09237173 | 26.685863 | 6.87E-157 |
| 226352 | EPB41L5 | 576.616612 | -3.3223286 | 0.12455958 | -26.672606 | 9.79E-157 |
| 72507 | DZIP1L | 1068.23108 | 8.77872271 | 0.33015746 | 26.5895026 | 8.98E-156 |
| 12808 | COBL | 837.636101 | -6.6322646 | 0.24966847 | -26.564286 | 1.76E-155 |
| 19414 | RASA3 | 3496.6973 | 1.51151565 | 0.05740756 | 26.3295573 | 8.80E-153 |
| 246727 | OAS3 | 331.340138 | -3.6954205 | 0.14068652 | -26.267055 | 4.56E-152 |
| 20334 | SEC23A | 3726.22228 | 1.47699111 | 0.05649747 | 26.1426068 | 1.20E-150 |
| 67701 | WFDC2 | 5475.30741 | -8.3135988 | 0.31834236 | -26.115276 | 2.45E-150 |
| 16156 | IL11 | 1382.85965 | 7.67219178 | 0.29519008 | 25.9906831 | 6.31E-149 |
| 11651 | AKT1 | 14240.7647 | 0.58254352 | 0.0224615 | 25.9352007 | 2.67E-148 |
| 19041 | PPL | 4581.50062 | -6.7005296 | 0.25891621 | -25.879144 | 1.14E-147 |
| 65963 | TMEM176B | 3838.40711 | -3.4917276 | 0.13518192 | -25.829842 | 4.10E-147 |
| 13661 | EHF | 1704.0765 | -7.443851 | 0.2883471 | -25.815592 | 5.93E-147 |
| 21356 | TAPBP | 3924.33264 | -1.6488067 | 0.06389097 | -25.806566 | 7.48E-147 |
| 102098 | ARHGEF18 | 1783.26711 | -1.2365813 | 0.0481759 | -25.668046 | 2.66E-145 |
| 12988 | CSK | 3128.1329 | 1.16042705 | 0.04538468 | 25.5686954 | 3.40E-144 |

| | | | | | | |
|---|---|---|---|---|---|---|
| 78255 | RALGPS2 | 665.878752 | -4.2574836 | 0.16687758 | -25.512617 | 1.43E-143 |
| 226419 | DYRK3 | 741.026901 | -2.1291048 | 0.08462402 | -25.15958 | 1.11E-139 |
| 68671 | PCYT2 | 763.257961 | -7.236016 | 0.29028658 | -24.927147 | 3.78E-137 |
| 68628 | FBXW9 | 857.552522 | -2.06323 | 0.08285647 | -24.901255 | 7.21E-137 |
| 13866 | ERBB2 | 8968.3762 | -3.3475017 | 0.13482964 | -24.82764 | 4.51E-136 |
| 18822 | PLOD1 | 1602.41328 | 1.5984434 | 0.06457915 | 24.7516933 | 2.97E-135 |
| 23876 | FBLN5 | 2397.38925 | 4.43035202 | 0.17900347 | 24.75009 | 3.09E-135 |
| 18104 | NQO1 | 7086.87019 | 5.51462342 | 0.22334541 | 24.6910083 | 1.34E-134 |
| 17158 | MAN2A1 | 9062.27386 | 1.80659906 | 0.07322894 | 24.6705612 | 2.21E-134 |
| 71461 | PTK7 | 2255.40571 | -4.3246915 | 0.17560701 | -24.6271 | 6.48E-134 |
| 18024 | NFE2L2 | 5035.66982 | 0.9917139 | 0.04124185 | 24.0462989 | 9.13E-128 |
| 71913 | TMEM79 | 682.583267 | -5.491701 | 0.22882148 | -23.999937 | 2.79E-127 |
| 19285 | PTRF | 12085.3187 | 4.69238181 | 0.19638227 | 23.8941212 | 3.53E-126 |
| 12725 | CLCN3 | 1749.89901 | -2.2575657 | 0.09463277 | -23.856066 | 8.76E-126 |
| 239554 | FOXRED2 | 1803.8527 | -5.6323846 | 0.23636029 | -23.829657 | 1.65E-125 |
| 109050 | FAM212B | 544.166292 | -3.1545288 | 0.13243414 | -23.819604 | 2.09E-125 |
| 26415 | mapk13 | 721.655502 | -8.2345501 | 0.34662211 | -23.756563 | 9.40E-125 |
| 12905 | CRADD | 304.746777 | -2.5925582 | 0.10924695 | -23.731173 | 1.72E-124 |
| 110532 | ADARB1 | 539.920178 | -2.0243364 | 0.08533704 | -23.72166 | 2.16E-124 |
| 216148 | SHC2 | 504.354974 | 7.36539021 | 0.31064721 | 23.7098228 | 2.86E-124 |
| 18414 | OSMR | 1742.61278 | 2.29944452 | 0.09740832 | 23.6062425 | 3.33E-123 |
| 83397 | AKAP12 | 9504.52208 | -4.8152906 | 0.20476065 | -23.51668 | 2.75E-122 |
| 12288 | CACNA1C | 795.940471 | 7.90977984 | 0.3370753 | 23.4659136 | 9.09E-122 |
| 67169 | Nradd | 498.737175 | 2.24139448 | 0.09572329 | 23.4153523 | 2.98E-121 |
| 12695 | PATJ | 847.698939 | -8.4660043 | 0.36184678 | -23.396655 | 4.62E-121 |
| 18693 | PICK1 | 510.813955 | -1.9896955 | 0.08554451 | -23.259184 | 1.15E-119 |
| 97440 | B3GNT9 | 245.251186 | 4.67990213 | 0.20168534 | 23.2039779 | 4.15E-119 |
| 211770 | TRIB1 | 2180.2119 | 1.92849138 | 0.08363076 | 23.0595933 | 1.18E-117 |
| 66629 | GOLPH3 | 6192.76774 | 0.83419599 | 0.03628222 | 22.9918671 | 5.62E-117 |
| 235633 | ALS2CL | 1221.34638 | -9.5433598 | 0.41719882 | -22.874848 | 8.27E-116 |
| 347722 | AGAP1 | 2413.04399 | -1.408315 | 0.06176533 | -22.801059 | 4.48E-115 |
| 21804 | TGFB1I1 | 671.193939 | 6.15705614 | 0.27006426 | 22.798486 | 4.75E-115 |
| 20732 | SPINT1 | 799.114895 | -7.6247387 | 0.33509519 | -22.753949 | 1.31E-114 |
| 83675 | BICC1 | 509.357856 | 7.70565322 | 0.33950256 | 22.6968926 | 4.81E-114 |
| 212285 | ARAP2 | 1170.217 | -3.1741263 | 0.13998315 | -22.67506 | 7.90E-114 |
| 54710 | HS3ST3B1 | 441.414663 | -6.9111431 | 0.3048402 | -22.671364 | 8.59E-114 |
| 106766 | STAP2 | 377.276891 | -5.22139 | 0.2308959 | -22.613611 | 3.18E-113 |
| 252973 | GRHL2 | 930.094638 | -6.8219585 | 0.30186672 | -22.599241 | 4.41E-113 |
| 12931 | CRLF1 | 3541.65722 | 4.82409841 | 0.21352532 | 22.5926292 | 5.12E-113 |
| 18611 | Pea15a | 5349.26167 | -1.6585759 | 0.07341826 | -22.590783 | 5.34E-113 |
| 13383 | DLG1 | 4934.32835 | -1.6257629 | 0.07205325 | -22.563352 | 9.93E-113 |
| 109332 | CDCP1 | 914.110998 | -8.7350505 | 0.38771053 | -22.529826 | 2.12E-112 |
| 228413 | PRRG4 | 2963.21463 | -5.7772851 | 0.25686343 | -22.491661 | 5.01E-112 |
| 245857 | SSH3 | 722.490836 | -5.5408322 | 0.24694568 | -22.437453 | 1.70E-111 |
| 99470 | MAGI3 | 1000.07533 | -1.9560011 | 0.08731418 | -22.401871 | 3.77E-111 |

| | | | | | | |
|---|---|---|---|---|---|---|
| 213948 | ATG9B | 1688.34278 | -5.9512475 | 0.26618145 | -22.35786 | 1.01E-110 |
| 227358 | FAM132B | 1023.9063 | -6.4864792 | 0.29056575 | -22.323619 | 2.18E-110 |
| 243219 | 2900026A02 | 2610.16231 | -2.2655264 | 0.10176487 | -22.262362 | 8.56E-110 |
| 319352 | PIANP | 415.278515 | 6.46181331 | 0.29068578 | 22.2295472 | 1.78E-109 |
| 22152 | TUBB3 | 355.423219 | 5.79787401 | 0.26094803 | 22.2185007 | 2.28E-109 |
| 17196 | MBP | 630.351016 | -7.5877065 | 0.34262192 | -22.146004 | 1.14E-108 |
| 18514 | PBX1 | 5560.18339 | -1.1228415 | 0.05070614 | -22.144092 | 1.19E-108 |
| 242785 | KLHL21 | 4588.78716 | 1.31326999 | 0.05954243 | 22.0560368 | 8.36E-108 |
| 80285 | PARP9 | 675.697718 | -1.7925077 | 0.08133129 | -22.039584 | 1.20E-107 |
| 330938 | DIXDC1 | 602.264938 | 2.68260063 | 0.12172688 | 22.0378649 | 1.25E-107 |
| 330814 | ADGRL1 | 1423.8838 | 3.38447399 | 0.15382833 | 22.0016292 | 2.78E-107 |
| 277414 | Trp53i11 | 472.286932 | -6.3279371 | 0.28822073 | -21.955177 | 7.73E-107 |
| 16476 | JUN | 3921.02421 | 1.819649 | 0.08308203 | 21.9018366 | 2.50E-106 |
| 211945 | PLEKHH1 | 254.919315 | -3.7353739 | 0.17097339 | -21.847692 | 8.18E-106 |
| 72754 | ARHGEF10L | 776.113537 | -3.7639945 | 0.17244632 | -21.82705 | 1.28E-105 |
| 70417 | MEGF10 | 1723.66153 | -8.8236614 | 0.40506162 | -21.783504 | 3.33E-105 |
| 208943 | MYO5C | 998.15261 | -9.2507741 | 0.42535794 | -21.74821 | 7.18E-105 |
| 22439 | XK | 242.873645 | -4.2555561 | 0.19598173 | -21.714045 | 1.51E-104 |
| 17937 | nab2 | 2436.42567 | 2.06781534 | 0.09532708 | 21.6917939 | 2.45E-104 |
| 207683 | igsf11 | 996.126823 | -8.087177 | 0.37326868 | -21.665833 | 4.31E-104 |
| 117606 | BOC | 372.265935 | 2.1646708 | 0.10015932 | 21.6122744 | 1.38E-103 |
| 216233 | SOCS2 | 1661.22264 | -4.5664338 | 0.21266345 | -21.472584 | 2.81E-102 |
| 629378 | DACT3 | 306.87503 | 4.11106838 | 0.19237577 | 21.3699909 | 2.54E-101 |
| 66058 | TMEM176A | 1828.79839 | -3.9005379 | 0.1830508 | -21.3085 | 9.47E-101 |
| 97031 | TPRN | 1103.52408 | -1.3681826 | 0.06440646 | -21.242942 | 3.83E-100 |
| 22352 | VIM | 86453.4352 | 1.4315901 | 0.06758567 | 21.1818588 | 1.40E-99 |
| 12266 | C3 | 7835.67587 | -3.0516315 | 0.14423159 | -21.157858 | 2.34E-99 |
| 21802 | TGFA | 1081.28659 | -8.7466008 | 0.41410937 | -21.121475 | 5.05E-99 |
| 13024 | Ctla2a | 783.450582 | -6.7342684 | 0.32003019 | -21.042604 | 2.67E-98 |
| 228876 | Zfp334 | 534.950271 | 2.13476522 | 0.10179482 | 20.971256 | 1.20E-97 |
| 13196 | ASAP1 | 9857.54332 | 1.63899096 | 0.07836088 | 20.9159338 | 3.83E-97 |
| 76509 | PLET1 | 354.796771 | -7.205255 | 0.34476993 | -20.898734 | 5.50E-97 |
| 71960 | MYH14 | 831.152546 | -9.2296781 | 0.44177843 | -20.892097 | 6.32E-97 |
| 75841 | RNF139 | 1737.63562 | 0.9882879 | 0.04737179 | 20.8623729 | 1.18E-96 |
| 330662 | DOCK1 | 2448.40376 | 2.33542267 | 0.11197912 | 20.8558766 | 1.35E-96 |
| 229474 | FHDC1 | 1312.06921 | -1.9894021 | 0.09573482 | -20.780341 | 6.52E-96 |
| 224833 | AI661453 | 611.442031 | -6.3159048 | 0.30448164 | -20.743138 | 1.41E-95 |
| 94346 | TMEM40 | 251.077728 | -6.3038295 | 0.30396016 | -20.738999 | 1.54E-95 |
| 223978 | CPPED1 | 1451.53354 | -1.1504397 | 0.05553806 | -20.714439 | 2.57E-95 |
| 102626 | MAPKAPK3 | 261.861285 | -3.3126579 | 0.16034649 | -20.659373 | 8.04E-95 |
| 21366 | SLC6A6 | 16623.7015 | 1.08599341 | 0.05258814 | 20.6509176 | 9.58E-95 |
| 14583 | GFPT1 | 3776.40041 | 0.97335495 | 0.04734157 | 20.5602581 | 6.23E-94 |
| 70472 | ATAD2 | 9239.99108 | 1.11723964 | 0.05442598 | 20.5276891 | 1.22E-93 |
| 14026 | EVL | 2539.32878 | 1.3768974 | 0.06719751 | 20.490304 | 2.63E-93 |
| 54648 | CCDC120 | 494.579463 | -1.9699646 | 0.09615341 | -20.487724 | 2.77E-93 |

| | | | | | | |
|---|---|---|---|---|---|---|
| 13433 | DNMT1 | 10488.995 | 0.78553383 | 0.03835048 | 20.4830243 | 3.05E-93 |
| 230779 | SERINC2 | 1294.80303 | -8.7342224 | 0.42728069 | -20.441416 | 7.16E-93 |
| 13392 | DLX2 | 761.704611 | 9.15555205 | 0.44874133 | 20.4027388 | 1.58E-92 |
| 18195 | NSF | 2352.26216 | 0.83833812 | 0.04133102 | 20.2835105 | 1.80E-91 |
| 59125 | NEK7 | 5248.4455 | -1.0706942 | 0.05282662 | -20.268082 | 2.46E-91 |
| 81535 | sgpp1 | 2407.51699 | 1.49515834 | 0.07385688 | 20.2439941 | 4.01E-91 |
| 53868 | RAB25 | 297.394327 | -6.6699402 | 0.32961284 | -20.235681 | 4.75E-91 |
| 71973 | RBPMS2 | 702.537491 | 4.65581955 | 0.23165234 | 20.0983059 | 7.64E-90 |
| 100303738 | D030025P21 | 577.039606 | 1.5452671 | 0.07693346 | 20.0857625 | 9.83E-90 |
| 320827 | C530008M17 | 457.097574 | 7.34179939 | 0.36554872 | 20.084325 | 1.01E-89 |
| 81840 | SORCS2 | 282.321305 | -4.5628971 | 0.22834382 | -19.982573 | 7.81E-89 |
| 12390 | CAV2 | 2871.35754 | -1.51334 | 0.07581608 | -19.960673 | 1.21E-88 |
| 12286 | CACNA1A | 234.367351 | 3.06523033 | 0.15356477 | 19.9605047 | 1.21E-88 |
| 234214 | SORBS2 | 9480.42577 | -10.119405 | 0.50829297 | -19.908607 | 3.43E-88 |
| 11518 | ADD1 | 5006.8303 | -1.006918 | 0.050657 | -19.877172 | 6.41E-88 |
| 232943 | KLC3 | 372.568983 | -3.7790403 | 0.190764 | -19.810028 | 2.44E-87 |
| 17075 | EPCAM | 2620.70612 | -11.347593 | 0.57398525 | -19.769833 | 5.42E-87 |
| 69698 | SLC52A3 | 183.444242 | -5.2650802 | 0.26737401 | -19.691818 | 2.53E-86 |
| 67980 | GNPDA2 | 454.980632 | -1.9555588 | 0.09957567 | -19.638921 | 7.19E-86 |
| 240752 | PIK3C2B | 191.452092 | -3.4678058 | 0.17668658 | -19.626877 | 9.11E-86 |
| 13849 | EPHX1 | 3092.38751 | -2.466031 | 0.12569502 | -19.619163 | 1.06E-85 |
| 73341 | ARHGEF6 | 194.820678 | -5.4786189 | 0.27990077 | -19.573433 | 2.61E-85 |
| 18640 | PFKFB2 | 1962.83455 | -2.0322182 | 0.1039541 | -19.549187 | 4.19E-85 |
| 65113 | NDFIP1 | 2337.11868 | -0.8470231 | 0.04338014 | -19.525599 | 6.65E-85 |
| 20743 | SPTBN2 | 522.86443 | -5.7808043 | 0.29639693 | -19.50359 | 1.02E-84 |
| 69697 | CAMSAP3 | 240.230655 | -5.4765071 | 0.28108952 | -19.483143 | 1.53E-84 |
| 93672 | IL24 | 315.574704 | -6.517129 | 0.33459243 | -19.477814 | 1.69E-84 |
| 12520 | CD81 | 5805.97248 | -1.16407 | 0.05992336 | -19.42598 | 4.65E-84 |
| 79555 | BC005537 | 8107.86592 | 0.738134 | 0.03802024 | 19.4142404 | 5.85E-84 |
| 16865 | EIF2D | 1703.0216 | -1.088172 | 0.05608645 | -19.40169 | 7.47E-84 |
| 68794 | FLNC | 10505.8388 | 2.23135462 | 0.11524025 | 19.3626324 | 1.60E-83 |
| 18645 | PFN2 | 851.35898 | 3.4663686 | 0.17959464 | 19.3010697 | 5.26E-83 |
| 213573 | cracr2b | 269.773806 | -6.6623352 | 0.34631321 | -19.237889 | 1.78E-82 |
| 20733 | SPINT2 | 2627.8478 | -11.763942 | 0.61186247 | -19.226448 | 2.22E-82 |
| 14007 | CELF2 | 1655.24432 | -1.4539921 | 0.07579328 | -19.183655 | 5.07E-82 |
| 14924 | MAGI1 | 1126.87041 | -1.5339538 | 0.07998926 | -19.176997 | 5.76E-82 |
| 240832 | TOR1AIP2 | 6052.32514 | -0.5720145 | 0.02983659 | -19.171578 | 6.39E-82 |
| 13732 | EMP3 | 3924.3761 | 3.20534981 | 0.16790527 | 19.0902281 | 3.04E-81 |
| 227960 | GCA | 226.503101 | -6.2764321 | 0.32894347 | -19.08058 | 3.66E-81 |
| 21873 | TJP2 | 3473.67439 | -2.3232828 | 0.12185238 | -19.066372 | 4.81E-81 |
| 245945 | RBM47 | 498.573316 | -8.24908 | 0.43336385 | -19.034998 | 8.75E-81 |
| 20442 | ST3GAL1 | 2176.46889 | 1.61412653 | 0.08495668 | 18.999406 | 1.72E-80 |
| 18557 | CDK18 | 346.900172 | -2.260908 | 0.11905504 | -18.990443 | 2.05E-80 |
| 67226 | TMEM19 | 1386.21223 | -1.2782969 | 0.06732009 | -18.988343 | 2.13E-80 |
| 72297 | B3GNT3 | 287.576595 | -4.4179797 | 0.23290173 | -18.969287 | 3.06E-80 |

| | | | | | | |
|---|---|---|---|---|---|---|
| 13867 | ERBB3 | 549.456798 | -8.1895153 | 0.43244749 | -18.937595 | 5.59E-80 |
| 216549 | AFTPH | 1809.97905 | -0.8242471 | 0.04360423 | -18.902914 | 1.08E-79 |
| 77411 | ESRP2 | 318.475693 | -7.4928184 | 0.39680537 | -18.882855 | 1.58E-79 |
| 76073 | PCGF5 | 480.816556 | -2.0350467 | 0.10826061 | -18.797665 | 7.89E-79 |
| 27205 | PODXL | 935.568229 | -6.7842433 | 0.36226268 | -18.727414 | 2.96E-78 |
| 216225 | SLC5A8 | 243.589083 | -5.889736 | 0.3151279 | -18.689986 | 5.97E-78 |
| 100340 | SMPDL3B | 183.24965 | -2.6575625 | 0.14229084 | -18.676975 | 7.62E-78 |
| 13591 | EBF1 | 470.107476 | 5.24049016 | 0.28154162 | 18.6135542 | 2.50E-77 |
| 23983 | PCBP1 | 14347.4054 | 0.95030448 | 0.05107999 | 18.6042419 | 2.97E-77 |
| 14797 | AES | 7464.48606 | -1.0714085 | 0.05773737 | -18.556588 | 7.21E-77 |
| 435684 | SHF | 877.730019 | -1.9216951 | 0.10356233 | -18.555929 | 7.30E-77 |
| 22409 | WNT10A | 780.636123 | -6.4375953 | 0.34705038 | -18.549454 | 8.24E-77 |
| 69815 | KRTCAP3 | 175.045701 | -5.0105131 | 0.2711296 | -18.48014 | 2.98E-76 |
| 213326 | SCYL2 | 1602.30074 | -0.9404128 | 0.05096659 | -18.451554 | 5.07E-76 |
| 67154 | MTDH | 9744.43865 | 0.84533496 | 0.04592084 | 18.408525 | 1.12E-75 |
| 68859 | SMIM1 | 226.328906 | -4.3339172 | 0.23547466 | -18.405026 | 1.20E-75 |
| 12609 | CEBPD | 1670.89262 | 2.87671093 | 0.15645585 | 18.386727 | 1.68E-75 |
| 20346 | SEMA3A | 1343.23327 | 4.06049172 | 0.22127846 | 18.3501441 | 3.29E-75 |
| 50798 | GNE | 1121.91456 | -1.7188385 | 0.09381396 | -18.321777 | 5.55E-75 |
| 21813 | TGFBR2 | 6010.78334 | -0.7389903 | 0.04043415 | -18.276388 | 1.28E-74 |
| 12816 | COL12A1 | 12375.1401 | 3.02955128 | 0.16598288 | 18.2521915 | 1.99E-74 |
| 66952 | 2310030G06 | 273.198188 | -5.0084165 | 0.2751147 | -18.204831 | 4.73E-74 |
| 20514 | slc1a5 | 2598.60978 | 3.06426622 | 0.16843741 | 18.1923134 | 5.94E-74 |
| 66102 | CXCL16 | 3110.61849 | -7.3020865 | 0.40243404 | -18.144803 | 1.41E-73 |
| 11877 | arvcf | 291.55601 | -2.5852161 | 0.14257032 | -18.13292 | 1.75E-73 |
| 108037 | SHMT2 | 7845.35379 | 0.84752303 | 0.04677214 | 18.120255 | 2.21E-73 |
| 12227 | BTG2 | 410.166107 | -2.6828596 | 0.1483537 | -18.084211 | 4.24E-73 |
| 12450 | CCNG1 | 12694.0492 | -1.5356653 | 0.0849587 | -18.075432 | 4.98E-73 |
| 320487 | HEATR5A | 3431.69318 | 1.44652382 | 0.0802536 | 18.0244112 | 1.25E-72 |
| 93842 | IGSF9 | 746.857594 | -7.7195354 | 0.42929573 | -17.981859 | 2.70E-72 |
| 223499 | DCAF13 | 2889.98995 | 0.95025215 | 0.05300886 | 17.9262892 | 7.35E-72 |
| 17974 | NCK2 | 672.816289 | -1.8718978 | 0.10447834 | -17.916611 | 8.75E-72 |
| 50490 | NOX4 | 849.504419 | -4.4128744 | 0.24731168 | -17.843373 | 3.25E-71 |
| 11745 | ANXA3 | 16571.2932 | -1.0550837 | 0.0591879 | -17.826003 | 4.44E-71 |
| 13382 | DLD | 4740.96321 | 0.57385672 | 0.03222518 | 17.8077119 | 6.16E-71 |
| 110796 | TSHZ1 | 3703.99287 | 1.81967151 | 0.10220197 | 17.8046612 | 6.50E-71 |
| 17207 | MCF2L | 168.546357 | 2.79674921 | 0.15718482 | 17.7927442 | 8.04E-71 |
| 14027 | EVPL | 150.293271 | -5.1389752 | 0.28930593 | -17.763117 | 1.36E-70 |
| 353211 | PRUNE2 | 3769.12128 | -5.2993375 | 0.29861192 | -17.746571 | 1.83E-70 |
| 235611 | PLXNB1 | 952.474065 | -4.9101052 | 0.27685113 | -17.735543 | 2.23E-70 |
| 106014 | FAM19A5 | 583.015674 | -7.9154426 | 0.44730337 | -17.695915 | 4.51E-70 |
| 67451 | PKP2 | 754.077657 | -2.4010063 | 0.13591688 | -17.665255 | 7.77E-70 |
| 56047 | MSLN | 8683.33444 | -8.749898 | 0.49567376 | -17.652534 | 9.73E-70 |
| 14339 | AKTIP | 825.361688 | -1.4500483 | 0.08216558 | -17.64788 | 1.06E-69 |
| 12337 | CAPN5 | 748.130362 | -3.8129115 | 0.21626289 | -17.63091 | 1.43E-69 |

| | | | | | | |
|---|---|---|---|---|---|---|
| 75605 | KDM5B | 1354.95963 | -1.3323306 | 0.0758891 | -17.556285 | 5.32E-69 |
| 270084 | LPCAT2 | 199.529035 | -6.6215258 | 0.37763935 | -17.533993 | 7.88E-69 |
| 320817 | ATAD2B | 1198.7169 | 1.06337868 | 0.06078012 | 17.4955023 | 1.55E-68 |
| 66195 | Lce1g | 1345.16238 | -8.0792948 | 0.46260672 | -17.464716 | 2.66E-68 |
| 319530 | Zfp750 | 275.688186 | -7.2812269 | 0.41777307 | -17.428665 | 5.00E-68 |
| 20657 | SOD3 | 3077.26204 | 3.07023184 | 0.17681553 | 17.3640399 | 1.54E-67 |
| 240505 | CDC42BPG | 2765.36761 | -5.4844056 | 0.31591124 | -17.36059 | 1.64E-67 |
| 72736 | TMX1 | 4146.56067 | 0.78924792 | 0.0455323 | 17.3338029 | 2.61E-67 |
| 52521 | Zfp622 | 2598.01087 | 0.9816782 | 0.05665451 | 17.3274511 | 2.92E-67 |
| 29816 | HIP1R | 2827.44673 | -1.241966 | 0.07189721 | -17.274189 | 7.36E-67 |
| 55960 | EBAG9 | 745.891769 | 1.04236984 | 0.06064718 | 17.1874414 | 3.30E-66 |
| 12236 | BUB1B | 4520.56517 | 1.04592768 | 0.0608772 | 17.1809418 | 3.69E-66 |
| 74105 | GGA2 | 1596.64888 | 1.01866131 | 0.05931074 | 17.1749887 | 4.09E-66 |
| 12111 | BGN | 6053.61834 | 13.6000746 | 0.79212527 | 17.1690958 | 4.52E-66 |
| 18704 | PIK3C2A | 9447.16482 | -1.4886089 | 0.08671456 | -17.16677 | 4.71E-66 |
| 71740 | NECTIN4 | 172.764124 | -6.0741294 | 0.35402982 | -17.157112 | 5.56E-66 |
| 104884 | TDP1 | 766.623048 | 1.20106084 | 0.07011195 | 17.1306155 | 8.77E-66 |
| 224860 | PLCL2 | 248.297159 | 6.25655198 | 0.36527855 | 17.128167 | 9.15E-66 |
| 329540 | NOL4L | 446.613843 | -5.7134429 | 0.33407463 | -17.102295 | 1.43E-65 |
| 547253 | PARP14 | 475.138035 | -1.981079 | 0.11584544 | -17.101053 | 1.46E-65 |
| 245128 | AU018091 | 151.45356 | -5.5058374 | 0.32199233 | -17.099281 | 1.50E-65 |
| 16373 | IRX3 | 1737.75372 | -2.0764913 | 0.12148613 | -17.092414 | 1.69E-65 |
| 17390 | MMP2 | 1052.97987 | 7.16713803 | 0.41964197 | 17.0791734 | 2.12E-65 |
| 55925 | syt8 | 497.874893 | -7.1817866 | 0.4207927 | -17.06728 | 2.60E-65 |
| 73182 | PEAR1 | 812.324107 | 7.05588929 | 0.41358377 | 17.0603632 | 2.93E-65 |
| 53885 | nphp1 | 541.341731 | -1.4283308 | 0.08383461 | -17.037485 | 4.33E-65 |
| 105844 | CARD10 | 632.614675 | -5.2155988 | 0.3061789 | -17.034481 | 4.56E-65 |
| 104886 | RAB15 | 2067.90032 | -4.0976488 | 0.24061349 | -17.030004 | 4.92E-65 |
| 18772 | pkp1 | 3576.49298 | -12.473523 | 0.7326442 | -17.025349 | 5.33E-65 |
| 211652 | WWC1 | 1893.14873 | -4.0295516 | 0.23672355 | -17.022183 | 5.62E-65 |
| 18717 | PIP5K1C | 5818.01597 | -1.2801554 | 0.075403 | -16.977512 | 1.20E-64 |
| 404634 | H2AFY2 | 629.619088 | 1.91760871 | 0.1129654 | 16.9751859 | 1.25E-64 |
| 53886 | CDKL2 | 646.143439 | -2.1296804 | 0.12574384 | -16.936658 | 2.41E-64 |
| 71889 | EPN3 | 355.357184 | -8.2898947 | 0.49005977 | -16.916089 | 3.42E-64 |
| 320706 | SOGA1 | 2845.39023 | 1.33739519 | 0.07908036 | 16.9118508 | 3.68E-64 |
| 100862012 | Gm21399 | 3567.86733 | 1.71637731 | 0.101524 | 16.9061235 | 4.06E-64 |
| 66260 | TMEM54 | 275.368546 | -7.5071019 | 0.4440927 | -16.904358 | 4.18E-64 |
| 12575 | CDKN1A | 5630.17347 | -1.4998043 | 0.08874512 | -16.900134 | 4.49E-64 |
| 16782 | LAMC2 | 5234.81416 | -2.3358612 | 0.13831228 | -16.888314 | 5.48E-64 |
| 27401 | SKP2 | 1767.49317 | 2.0333222 | 0.12059495 | 16.8607578 | 8.75E-64 |
| 13506 | DSC2 | 405.136283 | -7.9400714 | 0.47104296 | -16.856363 | 9.42E-64 |
| 27280 | PHLDA3 | 4323.8307 | -0.8520614 | 0.05056503 | -16.850805 | 1.03E-63 |
| 70110 | IFI35 | 458.427315 | -1.8561582 | 0.11021631 | -16.841048 | 1.22E-63 |
| 212439 | AA986860 | 249.664125 | -3.6754736 | 0.21833937 | -16.833765 | 1.38E-63 |
| 70564 | FAM213A | 502.20596 | -5.6167554 | 0.33491871 | -16.770504 | 4.01E-63 |

| | | | | | | |
|---|---|---|---|---|---|---|
| 15275 | HK1 | 7514.9451 | -1.6561905 | 0.09880205 | -16.762714 | 4.57E-63 |
| 12654 | Chil1 | 292.314966 | -5.9513352 | 0.35539913 | -16.745497 | 6.11E-63 |
| 18016 | NF2 | 4008.00173 | -0.7340575 | 0.0438921 | -16.724137 | 8.74E-63 |
| 72565 | UACA | 6295.81411 | 2.1833829 | 0.13060774 | 16.7171021 | 9.84E-63 |
| 73690 | GLIPR1 | 999.568758 | 6.70815124 | 0.4015535 | 16.7054982 | 1.20E-62 |
| 16012 | IGFBP6 | 773.285804 | 9.83565438 | 0.58933341 | 16.6894566 | 1.56E-62 |
| 16780 | LAMB3 | 2443.30575 | -5.6618046 | 0.33931355 | -16.686055 | 1.66E-62 |
| 69551 | 2310022B05 | 6062.4516 | 1.48355731 | 0.08893519 | 16.6813316 | 1.79E-62 |
| 20502 | SLC16A2 | 854.329646 | -9.9276397 | 0.59518784 | -16.679843 | 1.84E-62 |
| 77963 | HOOK1 | 457.002442 | -8.6570237 | 0.52047821 | -16.632826 | 4.03E-62 |
| 18647 | CDK14 | 1468.24243 | 1.02101493 | 0.06146808 | 16.6104901 | 5.85E-62 |
| 66898 | BAIAP2L1 | 712.130029 | -9.0045802 | 0.54216262 | -16.608633 | 6.04E-62 |
| 100737 | DCUN1D4 | 745.379956 | -1.1465345 | 0.06930792 | -16.54262 | 1.81E-61 |
| 105005 | FAM84A | 228.236238 | -6.226942 | 0.37675162 | -16.527977 | 2.31E-61 |
| 212980 | SLC45A3 | 180.49425 | -3.3612683 | 0.20382571 | -16.490895 | 4.27E-61 |
| 320924 | CCBE1 | 6017.5316 | -6.4562671 | 0.39177142 | -16.479679 | 5.14E-61 |
| 83965 | ENPP5 | 463.095032 | -8.0283498 | 0.48721083 | -16.478184 | 5.26E-61 |
| 108099 | PRKAG2 | 387.489912 | -2.5552615 | 0.15519875 | -16.464446 | 6.61E-61 |
| 21384 | TBX15 | 354.15515 | 8.5068825 | 0.51716756 | 16.4489872 | 8.53E-61 |
| 20454 | ST3GAL5 | 1779.03137 | -1.2476419 | 0.07599881 | -16.416598 | 1.45E-60 |
| 353342 | Peg13 | 625.167968 | 1.86676103 | 0.11377908 | 16.4068912 | 1.71E-60 |
| 19106 | eif2ak2 | 1553.87178 | -0.8094408 | 0.04937309 | -16.394372 | 2.10E-60 |
| 76041 | CCDC125 | 117.193683 | -4.9651161 | 0.30302703 | -16.38506 | 2.45E-60 |
| 76295 | ATP11B | 2065.7785 | -0.8361431 | 0.05113144 | -16.352818 | 4.15E-60 |
| 215113 | SLC43A2 | 501.420209 | -2.9012183 | 0.17786399 | -16.311443 | 8.18E-60 |
| 105787 | prkaa1 | 4426.19871 | 1.15958723 | 0.07111534 | 16.3057258 | 8.99E-60 |
| 14257 | FLT4 | 174.258824 | -3.6709979 | 0.22539751 | -16.286773 | 1.23E-59 |
| 14251 | FLOT1 | 1098.48627 | -2.2146961 | 0.13634152 | -16.243739 | 2.47E-59 |
| 14972 | H2-K1 | 7310.29979 | -1.4757355 | 0.0909306 | -16.22925 | 3.13E-59 |
| 97775 | D930048N14 | 116.47808 | -3.0811277 | 0.19020139 | -16.199292 | 5.10E-59 |
| 26563 | ROR1 | 482.548629 | -2.6335674 | 0.16273109 | -16.183554 | 6.59E-59 |
| 54637 | PRAF2 | 645.612561 | 1.1176734 | 0.06906742 | 16.1823523 | 6.72E-59 |
| 68734 | SMEK1 | 4207.54954 | 0.80678169 | 0.04986196 | 16.1803057 | 6.94E-59 |
| 19271 | PTPRJ | 822.497638 | -2.1204383 | 0.1312486 | -16.155893 | 1.03E-58 |
| 16878 | LIF | 2062.30939 | -2.0507497 | 0.12698687 | -16.149306 | 1.15E-58 |
| 19707 | REPS1 | 2629.90021 | 0.74926992 | 0.04643514 | 16.135839 | 1.43E-58 |
| 103534 | MGAT4B | 5049.60033 | -1.031643 | 0.06395761 | -16.130107 | 1.57E-58 |
| 22249 | UNC13B | 275.710172 | -6.5831185 | 0.40872814 | -16.10635 | 2.30E-58 |
| 12484 | Cd24a | 9536.77934 | -6.6771093 | 0.4147834 | -16.097822 | 2.64E-58 |
| 20350 | SEMA3F | 1145.68735 | -2.0348569 | 0.12640824 | -16.097502 | 2.66E-58 |
| 108705 | PTTG1IP | 7473.98941 | -1.8248097 | 0.11359493 | -16.064184 | 4.55E-58 |
| 54324 | ARHGEF5 | 3005.55655 | -3.9419335 | 0.24612366 | -16.016069 | 9.87E-58 |
| 108837 | IBTK | 2187.72324 | 0.98522716 | 0.06158367 | 15.9981896 | 1.32E-57 |
| 16668 | KRT18 | 9523.75959 | -16.647388 | 1.04137128 | -15.986026 | 1.60E-57 |
| 360013 | MYO18A | 2720.63357 | -1.4538226 | 0.09098996 | -15.977836 | 1.82E-57 |

| | | | | | | |
|---|---|---|---|---|---|---|
| 27410 | ABCA3 | 2058.13372 | -1.4178506 | 0.08876955 | -15.972264 | 1.99E-57 |
| 15170 | PTPN6 | 146.611597 | -5.291143 | 0.33156761 | -15.957961 | 2.51E-57 |
| 216961 | CORO6 | 226.110411 | -3.5958567 | 0.22535155 | -15.956654 | 2.56E-57 |
| 67622 | MXRA7 | 322.350074 | 8.20221817 | 0.51453875 | 15.9409144 | 3.29E-57 |
| 13139 | DGKA | 428.770666 | -2.693359 | 0.16925627 | -15.912906 | 5.16E-57 |
| 11911 | ATF4 | 7407.67951 | 0.66829946 | 0.04205027 | 15.8928695 | 7.10E-57 |
| 105387 | Akr1c14 | 282.827317 | 4.45304811 | 0.28046402 | 15.8774307 | 9.08E-57 |
| 83814 | NEDD4L | 2556.27633 | 1.12821053 | 0.07109004 | 15.8701637 | 1.02E-56 |
| 214230 | PAK6 | 284.713012 | -7.818729 | 0.49305005 | -15.857881 | 1.24E-56 |
| 28035 | USP39 | 3286.81326 | 0.78030848 | 0.04920869 | 15.8571272 | 1.26E-56 |
| 16196 | IL7 | 123.893498 | -5.3921251 | 0.34045634 | -15.837934 | 1.70E-56 |
| 18605 | ENPP1 | 941.271851 | -3.6658109 | 0.23147839 | -15.836515 | 1.74E-56 |
| 30963 | HACD1 | 254.247226 | 2.17636315 | 0.13750097 | 15.8279843 | 2.00E-56 |
| 59093 | PCBP3 | 236.644482 | -7.4085679 | 0.46882656 | -15.802364 | 3.00E-56 |
| 231207 | CPEB2 | 1127.07905 | -1.2639517 | 0.08010324 | -15.779034 | 4.34E-56 |
| 56468 | SOCS5 | 3707.50495 | 0.61310468 | 0.03896601 | 15.7343479 | 8.80E-56 |
| 107371 | EXOC6 | 459.563915 | -1.818662 | 0.1157368 | -15.713776 | 1.22E-55 |
| 17260 | MEF2C | 406.874103 | 3.98508082 | 0.25399971 | 15.6893127 | 1.79E-55 |
| 545975 | CERS3 | 302.834496 | -8.0629954 | 0.5141044 | -15.683576 | 1.96E-55 |
| 64817 | SVEP1 | 521.06673 | 9.0734206 | 0.57860694 | 15.6814929 | 2.02E-55 |
| 12399 | RUNX3 | 368.039703 | 7.63770032 | 0.48766848 | 15.6616649 | 2.77E-55 |
| 319482 | 9530053A07 | 110.189458 | -4.5373095 | 0.29002045 | -15.644792 | 3.61E-55 |
| 12475 | CD14 | 957.399643 | -3.3814366 | 0.21636279 | -15.628549 | 4.65E-55 |
| 14000 | DROSHA | 4841.66442 | 1.10582099 | 0.07080611 | 15.6175932 | 5.53E-55 |
| 26564 | ROR2 | 490.059192 | 9.39238908 | 0.60145134 | 15.6162078 | 5.65E-55 |
| 24055 | SH3BP2 | 1057.75301 | -3.550064 | 0.22736522 | -15.613927 | 5.85E-55 |
| 53421 | SEC61A1 | 12219.7407 | 0.78640642 | 0.05040806 | 15.600806 | 7.19E-55 |
| 108153 | ADAMTS7 | 751.993937 | 2.84586357 | 0.18242675 | 15.6000342 | 7.28E-55 |
| 13010 | CST3 | 6522.96437 | 1.14249469 | 0.07326622 | 15.5937432 | 8.03E-55 |
| 22240 | DPYSL3 | 1190.71077 | 6.54936059 | 0.42063389 | 15.570216 | 1.16E-54 |
| 24110 | USP18 | 133.393167 | -4.6030659 | 0.29612214 | -15.544484 | 1.73E-54 |
| 71145 | SCARA5 | 820.166138 | -5.0724564 | 0.32646255 | -15.537637 | 1.93E-54 |
| 233977 | ppfia1 | 3574.22432 | -0.6696783 | 0.04316314 | -15.51505 | 2.74E-54 |
| 17300 | FOXC1 | 2326.48537 | -0.8115158 | 0.05246857 | -15.466705 | 5.82E-54 |
| 13844 | EPHB2 | 863.973206 | 4.31471723 | 0.27951449 | 15.4364709 | 9.31E-54 |
| 50493 | TXNRD1 | 29396.7423 | 1.29422446 | 0.08384566 | 15.4357953 | 9.40E-54 |
| 26951 | ZW10 | 1362.91701 | 0.82695497 | 0.05358646 | 15.4321629 | 9.95E-54 |
| 14241 | FOXL1 | 646.889585 | 10.0591895 | 0.65271342 | 15.4113417 | 1.37E-53 |
| 12842 | COL1A1 | 9088.13435 | 8.57999137 | 0.5568263 | 15.4087394 | 1.43E-53 |
| 14600 | GHR | 1416.24114 | 1.77250754 | 0.11528616 | 15.3748504 | 2.41E-53 |
| 16691 | KRT8 | 6141.34879 | -13.989124 | 0.91046249 | -15.364855 | 2.82E-53 |
| 96957 | TMEM62 | 286.571052 | -2.753023 | 0.1793191 | -15.352648 | 3.40E-53 |
| 192678 | RASSF3 | 2081.95732 | -2.5650001 | 0.16716982 | -15.34368 | 3.90E-53 |
| 382620 | TMED8 | 1844.49532 | 1.22077296 | 0.07956411 | 15.343261 | 3.93E-53 |
| 244654 | MTSS1L | 389.492131 | 1.42532471 | 0.09293275 | 15.3371629 | 4.32E-53 |

| | | | | | | |
|---|---|---|---|---|---|---|
| 215280 | WIPF1 | 1644.09733 | 3.12869433 | 0.20400456 | 15.3363939 | 4.37E-53 |
| 15488 | HSD17B4 | 2362.81722 | -0.92613 | 0.06040896 | -15.331005 | 4.75E-53 |
| 224024 | SCARF2 | 433.682221 | 8.34619286 | 0.54469867 | 15.3225871 | 5.40E-53 |
| 16182 | IL18R1 | 522.615172 | -7.3037659 | 0.47672345 | -15.320761 | 5.56E-53 |
| 27008 | MICALL1 | 3846.7457 | -0.7897269 | 0.05157795 | -15.311327 | 6.42E-53 |
| 320705 | BEND6 | 196.663851 | 2.64213737 | 0.17262852 | 15.3053351 | 7.04E-53 |
| 66724 | TAB3 | 958.650997 | -1.1712084 | 0.07659963 | -15.29 | 8.91E-53 |
| 108148 | GALNT2 | 5550.47916 | -0.9814263 | 0.06427768 | -15.268539 | 1.24E-52 |
| 54598 | CALCRL | 161.390655 | 2.69837455 | 0.17675559 | 15.2661345 | 1.29E-52 |
| 71175 | NIPBL | 6915.56336 | 1.28250052 | 0.08414596 | 15.2413802 | 1.88E-52 |
| 269941 | CHSY1 | 2351.31373 | 1.89473704 | 0.12442402 | 15.228065 | 2.30E-52 |
| 232341 | WNK1 | 27526.0747 | -1.3064695 | 0.08589156 | -15.210686 | 3.00E-52 |
| 14132 | FCGRT | 142.976944 | 6.08984726 | 0.40142466 | 15.1705858 | 5.54E-52 |
| 192231 | HEXIM1 | 1536.03261 | 0.71679178 | 0.04729486 | 15.1558086 | 6.94E-52 |
| 237353 | SH3RF3 | 307.289956 | 3.85536357 | 0.2545311 | 15.1469253 | 7.94E-52 |
| 12876 | CPE | 16256.0979 | -1.3379489 | 0.08834498 | -15.144595 | 8.23E-52 |
| 17153 | MAL | 1719.83572 | -10.930305 | 0.72225157 | -15.133654 | 9.71E-52 |
| 15245 | HHIP | 132.56634 | -3.7348053 | 0.24706017 | -15.116987 | 1.25E-51 |
| 20024 | SUB1 | 3687.3724 | 1.45624153 | 0.09638091 | 15.1092314 | 1.41E-51 |
| 21452 | TCN2 | 674.086538 | -5.9745978 | 0.3966456 | -15.062811 | 2.84E-51 |
| 214305 | hhipl1 | 851.671501 | 2.04419082 | 0.13572504 | 15.0612647 | 2.91E-51 |
| 108075 | LTBP4 | 1182.56682 | 5.09359505 | 0.33821185 | 15.0603685 | 2.95E-51 |
| 170720 | CARD14 | 413.960659 | -7.8770336 | 0.52347879 | -15.047474 | 3.59E-51 |
| 76976 | Arxes2 | 169.62453 | 2.45472634 | 0.16313999 | 15.0467479 | 3.63E-51 |
| 57778 | FMNL1 | 704.6218 | -5.6039883 | 0.37252279 | -15.043343 | 3.82E-51 |
| 66522 | PGPEP1 | 939.849547 | -1.5807253 | 0.10509728 | -15.040592 | 3.98E-51 |
| 68519 | EML1 | 2155.92848 | 1.6397781 | 0.10915783 | 15.0220845 | 5.26E-51 |
| 12231 | BTN1A1 | 374.471693 | -5.1272514 | 0.342107 | -14.987274 | 8.89E-51 |
| 93737 | PARD6G | 436.124318 | 2.03844213 | 0.13607863 | 14.9798847 | 9.94E-51 |
| 223921 | AAAS | 1598.73989 | -0.939911 | 0.06275339 | -14.977854 | 1.02E-50 |
| 20698 | SPHK1 | 996.445709 | -2.5615162 | 0.17151598 | -14.934563 | 1.96E-50 |
| 100503676 | Gm38426 | 134.166288 | -3.9094222 | 0.26213177 | -14.913958 | 2.67E-50 |
| 195209 | Gm22 | 1025.44659 | -3.0632529 | 0.2054179 | -14.912298 | 2.74E-50 |
| 59035 | CARM1 | 3389.52417 | 0.95692128 | 0.06425767 | 14.8919386 | 3.72E-50 |
| 94216 | COL4A6 | 966.400287 | -10.578487 | 0.71064844 | -14.885683 | 4.08E-50 |
| 67880 | DCXR | 280.805046 | -1.7335682 | 0.1165567 | -14.873175 | 4.92E-50 |
| 106877 | AFAP1L1 | 386.791275 | -8.5865004 | 0.57741284 | -14.870643 | 5.11E-50 |
| 66853 | PNPLA2 | 1330.44954 | -1.3497212 | 0.09093031 | -14.843469 | 7.67E-50 |
| 19294 | NECTIN2 | 998.170926 | -2.634622 | 0.17754513 | -14.839168 | 8.18E-50 |
| 66086 | FOPNL | 1376.27406 | -0.669393 | 0.04522982 | -14.799816 | 1.47E-49 |
| 80976 | SYT13 | 1986.63144 | -6.101928 | 0.41381186 | -14.745658 | 3.28E-49 |
| 22673 | Zfp185 | 268.564205 | -7.7333002 | 0.52475695 | -14.736918 | 3.73E-49 |
| 218699 | PXK | 1447.96212 | 0.97942243 | 0.06649028 | 14.7303102 | 4.12E-49 |
| 19143 | ST14 | 1984.48465 | -12.362555 | 0.84005795 | -14.716312 | 5.07E-49 |
| 13038 | CTSK | 1064.02543 | 10.7743695 | 0.73309022 | 14.6971945 | 6.72E-49 |

| | | | | | | |
|---|---|---|---|---|---|---|
| 26941 | SLC9A3R1 | 1265.12361 | -3.39788 | 0.23148894 | -14.678369 | 8.87E-49 |
| 328258 | SLC25A48 | 2633.49791 | -12.771814 | 0.87050596 | -14.671714 | 9.78E-49 |
| 11421 | ACE | 197.60499 | -7.4425244 | 0.5074667 | -14.666035 | 1.06E-48 |
| 12740 | CLDN4 | 4557.06842 | -15.584024 | 1.06316416 | -14.658154 | 1.19E-48 |
| 16803 | LBP | 206.23998 | -6.3450705 | 0.43352418 | -14.636024 | 1.65E-48 |
| 207798 | GRAMD1C | 90.9547459 | -4.5790318 | 0.31306329 | -14.626537 | 1.90E-48 |
| 22223 | UCHL1 | 488.107092 | 3.80235044 | 0.26015968 | 14.6154486 | 2.24E-48 |
| 319742 | MPZL3 | 251.887182 | -5.9002026 | 0.40373176 | -14.614165 | 2.28E-48 |
| 234683 | ELMO3 | 170.409659 | -3.0742789 | 0.21047985 | -14.606049 | 2.57E-48 |
| 15980 | IFNGR2 | 1025.70431 | -0.9550214 | 0.06549795 | -14.580935 | 3.71E-48 |
| 22422 | WNT7B | 5112.73007 | -15.02854 | 1.03114929 | -14.574553 | 4.08E-48 |
| 19400 | RAPSN | 101.414067 | -5.0239638 | 0.345099 | -14.558036 | 5.19E-48 |
| 20168 | RTN3 | 6894.65671 | -0.7274996 | 0.04999576 | -14.551226 | 5.74E-48 |
| 171508 | creld1 | 288.691066 | -5.1312629 | 0.35308737 | -14.532559 | 7.54E-48 |
| 20720 | SERPINE2 | 4425.00281 | -7.2435465 | 0.49876252 | -14.523037 | 8.66E-48 |
| 229488 | FAM160A1 | 672.585072 | -1.131496 | 0.07796342 | -14.513164 | 1.00E-47 |
| 66381 | RNF113A2 | 387.192096 | 1.66568696 | 0.11478498 | 14.5113664 | 1.03E-47 |
| 30935 | TOR3A | 1905.95218 | -1.3067787 | 0.09011019 | -14.502008 | 1.18E-47 |
| 242050 | IGSF10 | 366.072081 | 5.18554619 | 0.35785758 | 14.4905306 | 1.39E-47 |
| 244373 | ERLIN2 | 2207.80388 | -1.0436572 | 0.07202821 | -14.489563 | 1.41E-47 |
| 72043 | SULF2 | 5011.05102 | -3.6294011 | 0.2508068 | -14.470904 | 1.85E-47 |
| 11630 | AIM1 | 5192.05551 | -3.0730814 | 0.21264196 | -14.451905 | 2.44E-47 |
| 85031 | PLA1A | 184.047673 | -5.6792915 | 0.39437299 | -14.400813 | 5.11E-47 |
| 13244 | DEGS1 | 8248.44686 | -0.4579427 | 0.03183749 | -14.383755 | 6.54E-47 |
| 235180 | FEZ1 | 110.989656 | 4.66909085 | 0.32477571 | 14.376355 | 7.28E-47 |
| 14269 | FNBP1 | 1425.12685 | 1.86671701 | 0.12994262 | 14.3657026 | 8.49E-47 |
| 109929 | ZBTB25 | 452.604659 | 1.29518537 | 0.09026389 | 14.3488762 | 1.08E-46 |
| 170772 | GLCCI1 | 483.030703 | 2.23042546 | 0.15554839 | 14.3391101 | 1.25E-46 |
| 259277 | KLK8 | 279.018498 | -3.5569134 | 0.24854425 | -14.310987 | 1.87E-46 |
| 226251 | ABLIM1 | 1274.60886 | -2.3504201 | 0.16439297 | -14.297571 | 2.27E-46 |
| 64058 | PERP | 6641.14557 | -4.1249174 | 0.28851434 | -14.297097 | 2.28E-46 |
| 54391 | RFK | 1755.56882 | -0.7283444 | 0.05095088 | -14.295029 | 2.35E-46 |
| 14537 | GCNT1 | 292.240274 | -6.8464433 | 0.47979957 | -14.269382 | 3.40E-46 |
| 20947 | SWAP70 | 3174.00304 | 1.59490692 | 0.11178262 | 14.2679335 | 3.47E-46 |
| 54525 | SYT7 | 335.298438 | 4.98716222 | 0.34960861 | 14.264987 | 3.62E-46 |
| 12223 | BTC | 191.647468 | -3.3275787 | 0.23345984 | -14.253324 | 4.27E-46 |
| 328949 | MCC | 3052.7032 | 1.3608579 | 0.0956621 | 14.2256745 | 6.35E-46 |
| 21415 | TCF7L1 | 460.405857 | 1.67863872 | 0.1180206 | 14.2232683 | 6.57E-46 |
| 67213 | CMTM6 | 1009.03994 | -0.7881283 | 0.05550286 | -14.199778 | 9.19E-46 |
| 215351 | SENP6 | 2912.11264 | 0.71842819 | 0.05062203 | 14.1920064 | 1.03E-45 |
| 16568 | KIF3A | 1076.24278 | -0.8412641 | 0.05928156 | -14.190992 | 1.04E-45 |
| 223775 | PIM3 | 1331.20619 | -1.9078251 | 0.13449617 | -14.184978 | 1.14E-45 |
| 78757 | RICTOR | 2667.69513 | 1.04053807 | 0.07336087 | 14.1838287 | 1.15E-45 |
| 15530 | HSPG2 | 15201.1649 | 1.75826949 | 0.12405852 | 14.1729043 | 1.35E-45 |
| 320051 | EXPH5 | 209.432617 | -7.8912293 | 0.55683262 | -14.171636 | 1.37E-45 |

| | | | | | | |
|---|---|---|---|---|---|---|
| 89867 | SEC16B | 179.539294 | 7.3578095 | 0.51920074 | 14.1714155 | 1.38E-45 |
| 56742 | PSRC1 | 855.849593 | -1.5554031 | 0.10996084 | -14.145064 | 2.00E-45 |
| 20466 | SIN3A | 2408.12069 | 0.57364172 | 0.0406269 | 14.11975 | 2.87E-45 |
| 52626 | CDKN2AIPNL | 2222.92814 | -1.4298031 | 0.10130252 | -14.114191 | 3.11E-45 |
| 58194 | SH3KBP1 | 4543.83838 | 1.56981081 | 0.11127649 | 14.1073002 | 3.42E-45 |
| 20310 | CXCL2 | 261.505258 | -3.9300847 | 0.27871555 | -14.100701 | 3.76E-45 |
| 105837 | MTBP | 1742.07817 | 1.31292894 | 0.09327985 | 14.0751606 | 5.40E-45 |
| 52428 | RHPN2 | 216.13895 | -3.764721 | 0.26767282 | -14.064637 | 6.26E-45 |
| 93703 | PCDHGB6 | 177.305888 | -7.4566222 | 0.53033976 | -14.060085 | 6.68E-45 |
| 381175 | CCDC68 | 168.223279 | -7.3784435 | 0.52595857 | -14.028564 | 1.04E-44 |
| 14042 | EXT1 | 6859.91987 | 1.19980018 | 0.08566073 | 14.0064199 | 1.42E-44 |
| 80884 | MAGED2 | 1413.82673 | -1.8098421 | 0.12931412 | -13.995703 | 1.66E-44 |
| 21929 | TNFAIP3 | 2359.46456 | -4.0149048 | 0.28691845 | -13.993192 | 1.72E-44 |
| 13860 | EPS8 | 11247.2905 | 2.55706998 | 0.18280485 | 13.9879766 | 1.85E-44 |
| 103172 | CHCHD10 | 353.911429 | -8.2804233 | 0.59276011 | -13.969265 | 2.40E-44 |
| 225870 | RIN1 | 2692.12955 | 0.91069703 | 0.06524018 | 13.9591443 | 2.77E-44 |
| 100502825 | Rpl37rt | 1899.64798 | 0.9424032 | 0.06755307 | 13.9505609 | 3.12E-44 |
| 170706 | TMEM37 | 308.562799 | -2.2559595 | 0.16181411 | -13.941674 | 3.54E-44 |
| 67241 | SMC6 | 5212.799 | 0.64389185 | 0.04623695 | 13.9259157 | 4.41E-44 |
| 236899 | PCYT1B | 1466.06692 | -2.1968329 | 0.15794427 | -13.908912 | 5.59E-44 |
| 70082 | LYSMD2 | 345.837026 | 2.22400388 | 0.15992532 | 13.9065147 | 5.78E-44 |
| 333654 | PPP1R13L | 1584.1024 | -1.5137031 | 0.10889586 | -13.900466 | 6.29E-44 |
| 21429 | UBTF | 5930.7899 | 0.55373051 | 0.03984512 | 13.8970718 | 6.60E-44 |
| 72284 | ORAOV1 | 546.970172 | -1.0278332 | 0.07404522 | -13.881156 | 8.24E-44 |
| 15939 | IER5 | 3680.09205 | 1.44558893 | 0.1041542 | 13.8793147 | 8.45E-44 |
| 54135 | LSR | 1192.95368 | -11.627578 | 0.83797718 | -13.875769 | 8.88E-44 |
| 67905 | PPM1M | 434.713861 | 1.28829595 | 0.09299996 | 13.8526506 | 1.23E-43 |
| 207920 | ESRP1 | 2161.36382 | -14.507734 | 1.04730177 | -13.852487 | 1.23E-43 |
| 14104 | FASN | 31457.6296 | 0.71114683 | 0.05136162 | 13.845881 | 1.35E-43 |
| 13845 | EPHB3 | 600.591075 | -1.4857412 | 0.10732276 | -13.843673 | 1.39E-43 |
| 217410 | TRIB2 | 2203.44312 | 2.43644166 | 0.17612301 | 13.8337495 | 1.59E-43 |
| 241639 | FERMT1 | 633.40316 | -10.296869 | 0.74487732 | -13.823577 | 1.84E-43 |
| 17532 | MRAS | 1067.94548 | 2.11017062 | 0.15267006 | 13.8217711 | 1.88E-43 |
| 140780 | BMP2K | 1427.0815 | 0.74516092 | 0.05392378 | 13.8187802 | 1.96E-43 |
| 100502698 | rubcn | 2249.64807 | -0.9908407 | 0.07177368 | -13.80507 | 2.38E-43 |
| 18719 | PIP5K1B | 256.36283 | -7.9851067 | 0.57889071 | -13.793807 | 2.78E-43 |
| 80877 | LRBA | 1743.531 | -1.8173491 | 0.13179518 | -13.789192 | 2.96E-43 |
| 75541 | Nat8f4 | 102.373635 | -4.9238332 | 0.35754452 | -13.771245 | 3.80E-43 |
| 16404 | itga7 | 146.615077 | -2.656691 | 0.19306427 | -13.760656 | 4.40E-43 |
| 68646 | NADK2 | 866.77412 | 2.25705714 | 0.16405402 | 13.7580118 | 4.56E-43 |
| 211401 | MTSS1 | 1557.46605 | -1.157757 | 0.08415968 | -13.756671 | 4.64E-43 |
| 30795 | FKBP3 | 2666.88765 | 0.98850411 | 0.07189443 | 13.749384 | 5.14E-43 |
| 69718 | IPMK | 1582.79906 | 1.27528469 | 0.09282788 | 13.7381652 | 6.00E-43 |
| 54120 | GIPC2 | 169.425922 | -5.3861677 | 0.39213734 | -13.735411 | 6.23E-43 |
| 12454 | CCNK | 2330.94395 | 0.8711918 | 0.06349439 | 13.7207673 | 7.63E-43 |

| | | | | | | |
|---|---|---|---|---|---|---|
| 13835 | EPHA1 | 472.382723 | -9.5504268 | 0.69725391 | -13.697201 | 1.06E-42 |
| 56018 | STARD10 | 726.86509 | -4.3009389 | 0.31415918 | -13.690318 | 1.16E-42 |
| 83603 | ELOVL4 | 459.386622 | 2.06407833 | 0.1507831 | 13.689056 | 1.18E-42 |
| 380967 | TMEM106C | 615.782608 | -1.1756114 | 0.08597622 | -13.673681 | 1.46E-42 |
| 20499 | SLC12A7 | 1402.79995 | -1.8584337 | 0.13610426 | -13.654485 | 1.90E-42 |
| 18260 | OCLN | 153.133908 | -7.2421207 | 0.53082768 | -13.643073 | 2.22E-42 |
| 67338 | RFFL | 744.94758 | 1.54750913 | 0.1135393 | 13.6297224 | 2.67E-42 |
| 93699 | PCDHGB1 | 141.772039 | 7.00381329 | 0.51441455 | 13.6151151 | 3.26E-42 |
| 15015 | H2-Q4 | 1877.17171 | -1.830738 | 0.13455526 | -13.605845 | 3.70E-42 |
| 18553 | PCSK6 | 1031.56662 | -8.1216899 | 0.59735734 | -13.596033 | 4.23E-42 |
| 16400 | ITGA3 | 15639.8056 | -3.7353429 | 0.27478952 | -13.593469 | 4.38E-42 |
| 11487 | ADAM10 | 6702.96909 | -0.8179451 | 0.06020073 | -13.586962 | 4.79E-42 |
| 55948 | SFN | 363.597229 | -5.655091 | 0.41679589 | -13.568011 | 6.20E-42 |
| 218865 | CHDH | 179.942398 | -6.2900119 | 0.46361919 | -13.567195 | 6.27E-42 |
| 14369 | FZD7 | 3205.11769 | 1.77907291 | 0.13114556 | 13.5656359 | 6.40E-42 |
| 72434 | LYPD3 | 203.894397 | -7.8468362 | 0.5792442 | -13.546681 | 8.29E-42 |
| 211389 | SUOX | 516.237127 | -2.6231351 | 0.19382153 | -13.533765 | 9.88E-42 |
| 22145 | tuba4a | 6266.4687 | -0.6542044 | 0.04834753 | -13.531291 | 1.02E-41 |
| 107065 | LRRTM2 | 131.774963 | 4.93934661 | 0.36503915 | 13.5310051 | 1.03E-41 |
| 16922 | PHYH | 183.817197 | -1.8813557 | 0.13915475 | -13.519881 | 1.19E-41 |
| 15586 | HYAL1 | 310.14691 | -2.4101329 | 0.17830231 | -13.517116 | 1.24E-41 |
| 70790 | UBR5 | 16672.4598 | 1.43261689 | 0.10612104 | 13.4998382 | 1.57E-41 |
| 224613 | FLYWCH1 | 1215.0211 | 1.18982576 | 0.08817205 | 13.4943643 | 1.69E-41 |
| 217030 | SYNRG | 859.551215 | -1.0595611 | 0.07852841 | -13.49271 | 1.73E-41 |
| 269999 | ORAI3 | 547.545769 | 1.4756896 | 0.10941911 | 13.4865806 | 1.88E-41 |
| 69219 | DDAH1 | 2172.57202 | -2.2753976 | 0.16892002 | -13.470266 | 2.34E-41 |
| 20195 | S100A11 | 2897.10004 | -1.0258332 | 0.07621634 | -13.459492 | 2.71E-41 |
| 109620 | DSP | 4258.41567 | -14.044447 | 1.0435265 | -13.45864 | 2.74E-41 |
| 83679 | PDE4DIP | 3258.43674 | -1.0809976 | 0.08038465 | -13.447811 | 3.17E-41 |
| 18826 | LCP1 | 2066.97174 | -12.010028 | 0.89346903 | -13.442019 | 3.43E-41 |
| 104721 | DDX1 | 7416.08757 | 0.60911609 | 0.04531747 | 13.4410868 | 3.47E-41 |
| 73246 | RASSF6 | 411.429908 | -3.8851929 | 0.28913454 | -13.437319 | 3.65E-41 |
| 229541 | DENND4B | 439.033517 | 2.33618573 | 0.1741738 | 13.4129575 | 5.08E-41 |
| 330406 | B4GALNT3 | 227.798077 | -8.2336315 | 0.6145306 | -13.398245 | 6.19E-41 |
| 225115 | svil | 2996.45652 | -1.5939563 | 0.11899135 | -13.395564 | 6.42E-41 |
| 67138 | HERC6 | 433.997159 | -1.3925474 | 0.10397388 | -13.393242 | 6.62E-41 |
| 20336 | EXOC4 | 2513.19237 | -1.20638 | 0.09018763 | -13.376336 | 8.31E-41 |
| 218518 | MARVELD2 | 203.026426 | -8.0673701 | 0.60343896 | -13.368991 | 9.18E-41 |
| 67815 | SEC14L2 | 184.077627 | -3.7717493 | 0.28222809 | -13.364188 | 9.79E-41 |
| 16975 | LRP8 | 3453.27176 | 1.02482574 | 0.07678808 | 13.3461559 | 1.25E-40 |
| 71766 | RAVER1 | 4708.91795 | 0.51145714 | 0.03836591 | 13.3310328 | 1.53E-40 |
| 666704 | SAMD1 | 1187.64531 | 1.1464522 | 0.08601505 | 13.3285077 | 1.58E-40 |
| 18214 | DDR2 | 5522.96991 | 6.40344818 | 0.4813072 | 13.3042849 | 2.19E-40 |
| 14425 | GALNT3 | 875.080479 | -11.180034 | 0.84095365 | -13.294472 | 2.49E-40 |
| 60599 | Trp53inp1 | 2895.72267 | -3.3473807 | 0.25193714 | -13.286571 | 2.77E-40 |

| | | | | | | |
|---|---|---|---|---|---|---|
| 20973 | SYNGR2 | 2833.71741 | -0.9878318 | 0.07437854 | -13.28114 | 2.98E-40 |
| 223527 | ENY2 | 1829.71182 | 1.33917166 | 0.10083737 | 13.2805101 | 3.00E-40 |
| 14814 | GRIN2D | 83.1069151 | 4.39846416 | 0.3312031 | 13.2802627 | 3.01E-40 |
| 16403 | ITGA6 | 3708.39428 | -2.2110044 | 0.16652756 | -13.277108 | 3.14E-40 |
| 16525 | KCNK1 | 1139.35259 | -13.583911 | 1.02452937 | -13.258683 | 4.02E-40 |
| 68292 | STT3B | 8177.42935 | 0.56535188 | 0.04267049 | 13.2492463 | 4.56E-40 |
| 207181 | RBMS3 | 250.408424 | 2.81952738 | 0.21297931 | 13.2385036 | 5.26E-40 |
| 100040608 | Fancf | 295.578069 | 1.53481631 | 0.11604892 | 13.2255975 | 6.24E-40 |
| 207278 | FCHSD2 | 876.356745 | 0.98369839 | 0.07438925 | 13.2236632 | 6.41E-40 |
| 12217 | BSN | 196.798707 | 4.38869607 | 0.33204742 | 13.2170763 | 6.99E-40 |
| 12739 | CLDN3 | 2637.27455 | -13.355104 | 1.01068124 | -13.213962 | 7.29E-40 |
| 66471 | ANP32E | 6334.92428 | 0.54646736 | 0.04136669 | 13.2103246 | 7.65E-40 |
| 11910 | ATF3 | 256.505831 | -3.8408471 | 0.2910605 | -13.196044 | 9.25E-40 |
| 56274 | stk3 | 2032.15038 | 1.00005125 | 0.0758399 | 13.1863464 | 1.05E-39 |
| 72461 | PRCP | 1473.91986 | 1.50534222 | 0.1141654 | 13.1856251 | 1.06E-39 |
| 192216 | TMEM47 | 1122.88261 | -13.563398 | 1.02926462 | -13.177756 | 1.18E-39 |
| 12349 | Car2 | 475.495249 | -9.8788901 | 0.74996553 | -13.172459 | 1.26E-39 |
| 382571 | KCNF1 | 311.184468 | -3.0766387 | 0.23358885 | -13.171171 | 1.29E-39 |
| 13008 | CSRP2 | 954.032128 | 4.1307338 | 0.31418064 | 13.1476397 | 1.76E-39 |
| 105349 | Akr1c18 | 497.802911 | 6.42890041 | 0.48953193 | 13.1327499 | 2.14E-39 |
| 59003 | MAEA | 4639.35606 | -0.5390683 | 0.04109264 | -13.118365 | 2.58E-39 |
| 232164 | PAIP2B | 1365.63739 | -0.8569867 | 0.06537087 | -13.109611 | 2.90E-39 |
| 68775 | ATP6V1C2 | 98.2995507 | -6.1309658 | 0.46785586 | -13.10439 | 3.11E-39 |
| 232934 | MYPOP | 362.034857 | 1.92474193 | 0.14716271 | 13.0790058 | 4.34E-39 |
| 70673 | PRDM16 | 173.650627 | 7.8928932 | 0.60353054 | 13.0778688 | 4.41E-39 |
| 110109 | NOP2 | 5069.83486 | 0.55816641 | 0.04268537 | 13.0762917 | 4.50E-39 |
| 432940 | OTULIN | 1113.12822 | 0.84528737 | 0.06467449 | 13.0698739 | 4.89E-39 |
| 70097 | SASH1 | 3661.77229 | 1.28562601 | 0.09841664 | 13.0630961 | 5.35E-39 |
| 69540 | KLK10 | 536.768244 | 1.58896532 | 0.12171622 | 13.0546715 | 5.98E-39 |
| 13014 | CSTB | 3832.19576 | -1.2145752 | 0.09308103 | -13.04858 | 6.47E-39 |
| 231668 | vsig10 | 320.652497 | -1.941462 | 0.14884188 | -13.043788 | 6.89E-39 |
| 12053 | BCL6 | 928.74085 | -1.0354304 | 0.07942003 | -13.037395 | 7.50E-39 |
| 229011 | SAMD10 | 285.3645 | -2.1313736 | 0.16349659 | -13.036196 | 7.62E-39 |
| 77579 | MYH10 | 2761.06677 | -14.13964 | 1.08499201 | -13.032022 | 8.04E-39 |
| 26943 | SERINC3 | 15086.9653 | -2.6750134 | 0.20558217 | -13.011894 | 1.05E-38 |
| 100502897 | Gm12854 | 1303.76578 | -1.0745404 | 0.08258706 | -13.011002 | 1.06E-38 |
| 327826 | FRS2 | 2873.88996 | 1.12369999 | 0.08638479 | 13.0080763 | 1.10E-38 |
| 380921 | DGKH | 204.077027 | 3.34247819 | 0.25701045 | 13.0052231 | 1.14E-38 |
| 20377 | SFRP1 | 1145.61911 | -13.592189 | 1.04651426 | -12.988059 | 1.43E-38 |
| 59092 | PCBP4 | 2794.87553 | 1.89312208 | 0.14600526 | 12.9661225 | 1.90E-38 |
| 26408 | MAP3K5 | 674.365519 | -2.6285313 | 0.20290268 | -12.954641 | 2.21E-38 |
| 16210 | IMPACT | 4189.33055 | 1.31287064 | 0.10134456 | 12.9545244 | 2.22E-38 |
| 229445 | CTSO | 163.341438 | -1.7943475 | 0.13866101 | -12.940534 | 2.66E-38 |
| 20878 | AURKA | 2865.27819 | 0.57249791 | 0.04424074 | 12.940513 | 2.66E-38 |
| 100862305 | #N/A | 165.418361 | 7.81974542 | 0.60449062 | 12.9360907 | 2.82E-38 |

| | | | | | | |
|---|---|---|---|---|---|---|
| 104806 | fancm | 1738.56631 | 0.73825868 | 0.05711213 | 12.926477 | 3.19E-38 |
| 93732 | ACOX2 | 137.76502 | -7.2805756 | 0.5635889 | -12.918238 | 3.55E-38 |
| 27055 | FKBP9 | 5234.2927 | 0.87019473 | 0.067407 | 12.9095601 | 3.98E-38 |
| 74137 | NUAK2 | 449.259048 | -2.5134404 | 0.19511301 | -12.881973 | 5.69E-38 |
| 214627 | PAPD5 | 2689.14058 | 0.71105118 | 0.05521105 | 12.878783 | 5.93E-38 |
| 11839 | AREG | 708.04565 | -4.8856192 | 0.37974883 | -12.865396 | 7.05E-38 |
| 74006 | DNM1L | 3653.97213 | -0.4722067 | 0.03671264 | -12.862236 | 7.34E-38 |
| 53624 | CLDN7 | 844.988088 | -13.153229 | 1.02331438 | -12.853556 | 8.21E-38 |
| 14404 | GABRE | 101.160862 | -6.3123059 | 0.49134588 | -12.84697 | 8.94E-38 |
| 99167 | SSX2IP | 1040.96086 | -1.1017997 | 0.08585926 | -12.832625 | 1.08E-37 |
| 56460 | PKP3 | 996.788229 | -4.6109026 | 0.35945514 | -12.827477 | 1.15E-37 |
| 13198 | DDIT3 | 519.003858 | -1.4088104 | 0.10984136 | -12.825864 | 1.17E-37 |
| 21423 | TCF3 | 3660.69809 | 0.65222308 | 0.05088658 | 12.8171929 | 1.31E-37 |
| 16402 | ITGA5 | 5362.04575 | 1.02569337 | 0.08010186 | 12.8048626 | 1.54E-37 |
| 21930 | TNFAIP6 | 105.376322 | 2.5958382 | 0.20276122 | 12.8024392 | 1.59E-37 |
| 13025 | Ctla2b | 192.993143 | -1.9510123 | 0.15247813 | -12.795358 | 1.74E-37 |
| 13437 | DNPEP | 1105.98635 | -0.7799356 | 0.06095772 | -12.794698 | 1.76E-37 |
| 27273 | PDK4 | 356.680343 | -8.8739068 | 0.69429168 | -12.781237 | 2.09E-37 |
| 13476 | REEP5 | 4947.95059 | -0.9725771 | 0.07614393 | -12.772877 | 2.32E-37 |
| 21934 | TNFRSF11A | 80.8848666 | -4.4712825 | 0.35050563 | -12.756664 | 2.86E-37 |
| 54608 | ABHD2 | 7428.07783 | -1.4916197 | 0.11699863 | -12.749035 | 3.16E-37 |
| 269437 | PLCH1 | 109.182515 | -2.5782991 | 0.20233013 | -12.743031 | 3.41E-37 |
| 17035 | LXN | 1807.37334 | 1.84137046 | 0.14451095 | 12.7420829 | 3.45E-37 |
| 14630 | GCLM | 8064.64792 | 1.46763081 | 0.11519327 | 12.7405948 | 3.52E-37 |
| 223650 | EPPK1 | 162.907162 | -7.5199419 | 0.59045315 | -12.735882 | 3.74E-37 |
| 14299 | NCS1 | 2736.07104 | 1.80694899 | 0.14212398 | 12.7138927 | 4.95E-37 |
| 17754 | MAP1A | 2424.72168 | 6.54841184 | 0.51514777 | 12.7117154 | 5.09E-37 |
| 16009 | IGFBP3 | 141.825205 | -6.438996 | 0.50672737 | -12.707022 | 5.41E-37 |
| 22381 | WBP5 | 3127.30232 | -0.9667001 | 0.07608249 | -12.705946 | 5.48E-37 |
| 22631 | YWHAZ | 9188.44143 | 0.84282471 | 0.06634716 | 12.703252 | 5.67E-37 |
| 109095 | RBM15B | 1779.30656 | 0.80568644 | 0.0634452 | 12.6989344 | 5.99E-37 |
| 66985 | RASSF7 | 587.393872 | -1.7186963 | 0.13539955 | -12.693515 | 6.42E-37 |
| 338372 | MAP3K9 | 583.824666 | -7.9179862 | 0.62427489 | -12.683493 | 7.30E-37 |
| 26457 | SLC27A1 | 2349.13829 | 2.23484668 | 0.17645904 | 12.6649599 | 9.25E-37 |
| 16763 | LAD1 | 3754.56556 | -9.1751129 | 0.72505384 | -12.654388 | 1.06E-36 |
| 27260 | PLEK2 | 83.4825644 | -3.2869207 | 0.25988915 | -12.647395 | 1.16E-36 |
| 13511 | DSG2 | 836.318603 | -13.137777 | 1.038987 | -12.644794 | 1.20E-36 |
| 19200 | PSTPIP1 | 331.466633 | -2.6998866 | 0.21352395 | -12.644421 | 1.20E-36 |
| 52055 | rab11fip5 | 5329.45791 | -1.309692 | 0.10360133 | -12.641652 | 1.24E-36 |
| 101772 | ANO1 | 4451.50715 | -6.5412495 | 0.51798052 | -12.62837 | 1.47E-36 |
| 73737 | 1110008P14I | 1082.22872 | -1.8683281 | 0.14797055 | -12.626351 | 1.51E-36 |
| 17191 | MBD2 | 2675.32912 | -0.5680307 | 0.04499366 | -12.624684 | 1.54E-36 |
| 78244 | DNAJC21 | 1271.29181 | 1.25159338 | 0.09916266 | 12.6216194 | 1.60E-36 |
| 58220 | PARD6B | 546.08799 | -3.3602378 | 0.26630848 | -12.61784 | 1.68E-36 |
| 12444 | CCND2 | 4727.03784 | 6.18010355 | 0.4902137 | 12.606958 | 1.93E-36 |

| | | | | | | |
|---|---|---|---|---|---|---|
| 231858 | RADIL | 112.455255 | 2.60802415 | 0.20687223 | 12.6069322 | 1.93E-36 |
| 217869 | EIF5 | 12725.0096 | 0.60342685 | 0.0478899 | 12.6002936 | 2.10E-36 |
| 24063 | SPRY1 | 234.516666 | -3.3766947 | 0.26808303 | -12.595705 | 2.23E-36 |
| 257632 | NOD2 | 101.890568 | -4.9960831 | 0.39675387 | -12.592399 | 2.32E-36 |
| 67832 | BRIX1 | 3211.33594 | 1.05026189 | 0.08341976 | 12.5900859 | 2.39E-36 |
| 54563 | NUP210 | 438.995139 | -6.4132055 | 0.50939865 | -12.589758 | 2.40E-36 |
| 18080 | NIN | 3167.64099 | 1.50112715 | 0.11928915 | 12.5839371 | 2.59E-36 |
| 208898 | UNC13C | 124.719248 | -5.3927235 | 0.4286845 | -12.579703 | 2.73E-36 |
| 223601 | FAM49B | 3808.8549 | 1.07946052 | 0.08586606 | 12.5714457 | 3.03E-36 |
| 109246 | TSPAN9 | 3422.70928 | 1.24274602 | 0.09890765 | 12.5647104 | 3.30E-36 |
| 14538 | gcnt2 | 695.886204 | -3.1664163 | 0.25221072 | -12.554646 | 3.75E-36 |
| 14806 | GRIK2 | 105.759143 | -6.7016277 | 0.53380265 | -12.554504 | 3.76E-36 |
| 16420 | ITGB6 | 797.774425 | -13.070184 | 1.04127161 | -12.552137 | 3.87E-36 |
| 19355 | RAD1 | 605.900928 | 1.28936657 | 0.10280463 | 12.5419117 | 4.40E-36 |
| 94040 | CLMN | 279.444038 | -6.8640016 | 0.54736252 | -12.540138 | 4.50E-36 |
| 20317 | SERPINF1 | 4589.94562 | 9.06517918 | 0.7236849 | 12.5264175 | 5.35E-36 |
| 18132 | NOTCH4 | 943.952527 | -5.2074473 | 0.41589149 | -12.521168 | 5.72E-36 |
| 12833 | COL6A1 | 3610.90632 | 4.02047067 | 0.32128813 | 12.5135985 | 6.29E-36 |
| 243771 | PARP12 | 882.567775 | -1.1948178 | 0.09548979 | -12.512518 | 6.38E-36 |
| 69064 | FUOM | 156.190635 | -3.1404499 | 0.25110914 | -12.506315 | 6.89E-36 |
| 18519 | KAT2B | 1214.19641 | -1.6735628 | 0.13388642 | -12.499869 | 7.48E-36 |
| 54670 | ATP8B1 | 211.974642 | -8.122672 | 0.65066885 | -12.483573 | 9.18E-36 |
| 54139 | IRF6 | 650.957524 | -12.776687 | 1.02380514 | -12.479608 | 9.65E-36 |
| 209760 | TMC7 | 266.874688 | -7.2212081 | 0.57889879 | -12.474042 | 1.03E-35 |
| 101943 | SF3B3 | 10847.4724 | 0.55180787 | 0.04424227 | 12.4724144 | 1.06E-35 |
| 230073 | DDX58 | 1083.31761 | -1.9936821 | 0.15986144 | -12.471313 | 1.07E-35 |
| 246703 | apoa1bp | 782.912089 | -1.1158233 | 0.08947352 | -12.470989 | 1.07E-35 |
| 50772 | MAPK6 | 14726.5414 | 1.36961171 | 0.10986414 | 12.4664131 | 1.14E-35 |
| 230594 | ZCCHC11 | 948.630854 | 1.74714151 | 0.14024534 | 12.4577512 | 1.27E-35 |
| 66848 | FUCA2 | 1914.1463 | -2.5114777 | 0.20161878 | -12.456566 | 1.29E-35 |
| 68016 | MURC | 158.692674 | 7.77058577 | 0.62396378 | 12.4535847 | 1.34E-35 |
| 223455 | 41703 | 5621.65879 | 0.84556247 | 0.06791248 | 12.4507679 | 1.39E-35 |
| 70369 | BAG5 | 1117.63691 | 0.91033152 | 0.07315365 | 12.4441026 | 1.51E-35 |
| 100198 | H6PD | 2387.89844 | -0.6811981 | 0.05474155 | -12.443895 | 1.51E-35 |
| 269704 | Zfp664 | 3909.23815 | -0.5629035 | 0.04523533 | -12.44389 | 1.51E-35 |
| 16450 | JAG2 | 84.3950682 | -3.7090808 | 0.29810749 | -12.442092 | 1.54E-35 |
| 17909 | MYO10 | 11761.2232 | 1.18730318 | 0.09546375 | 12.4372145 | 1.64E-35 |
| 22367 | VRK1 | 1534.42491 | 0.78270949 | 0.0629528 | 12.4332752 | 1.72E-35 |
| 69608 | SEC24D | 4371.35548 | 0.90921534 | 0.07313166 | 12.4325824 | 1.74E-35 |
| 329251 | PPP1R12B | 240.325435 | 2.192844 | 0.17651025 | 12.4233237 | 1.95E-35 |
| 171486 | CD99L2 | 1804.47824 | -0.9364326 | 0.07539604 | -12.420182 | 2.03E-35 |
| 26360 | angptl2 | 14433.579 | 2.07744963 | 0.16737922 | 12.411634 | 2.26E-35 |
| 18082 | NIPSNAP1 | 540.825418 | -5.1591978 | 0.41577131 | -12.408739 | 2.34E-35 |
| 234797 | 6430548M08 | 291.167993 | -3.1327956 | 0.25252119 | -12.40607 | 2.42E-35 |
| 235584 | DUSP7 | 2809.13577 | -1.2015526 | 0.0968772 | -12.402842 | 2.52E-35 |

| | | | | | | |
|---|---|---|---|---|---|---|
| 21817 | TGM2 | 3034.3422 | -4.4286182 | 0.35786466 | -12.37512 | 3.56E-35 |
| 12370 | casp8 | 1270.5214 | -1.003031 | 0.08110149 | -12.367603 | 3.91E-35 |
| 433256 | ACSL5 | 6717.31108 | -0.9329603 | 0.07546759 | -12.362397 | 4.18E-35 |
| 14234 | FOXC2 | 948.48614 | 12.648382 | 1.02379504 | 12.3544084 | 4.61E-35 |
| 12006 | AXIN2 | 208.42842 | -5.6851067 | 0.46027622 | -12.351511 | 4.78E-35 |
| 330260 | PON2 | 1397.53408 | -1.6186057 | 0.13114617 | -12.341998 | 5.38E-35 |
| 18795 | PLCB1 | 240.576622 | 1.74941356 | 0.14197677 | 12.3218298 | 6.91E-35 |
| 70237 | BHLHB9 | 969.655008 | 1.57346785 | 0.12770413 | 12.3211982 | 6.97E-35 |
| 67432 | HOGA1 | 128.151371 | -3.3688723 | 0.27355075 | -12.315347 | 7.49E-35 |
| 21872 | TJP1 | 6176.04592 | -0.9019311 | 0.07331721 | -12.301765 | 8.86E-35 |
| 20613 | SNAI1 | 1238.76827 | 3.5244195 | 0.28668843 | 12.2935533 | 9.81E-35 |
| 97130 | C77080 | 3840.30464 | -1.6012195 | 0.13045736 | -12.273891 | 1.25E-34 |
| 78658 | NCAPD3 | 1742.0846 | 0.68262007 | 0.05563021 | 12.270673 | 1.30E-34 |
| 56356 | GLTP | 2015.63745 | -1.0479649 | 0.0854773 | -12.260154 | 1.48E-34 |
| 68498 | TSPAN11 | 946.652724 | 11.3403892 | 0.92521452 | 12.2570376 | 1.54E-34 |
| 64339 | FNDC4 | 131.276295 | -3.5416796 | 0.28911068 | -12.250255 | 1.67E-34 |
| 239217 | KCTD12 | 1028.62091 | 3.74081232 | 0.30545493 | 12.2466915 | 1.75E-34 |
| 17256 | MEA1 | 919.702258 | -0.7015724 | 0.05730615 | -12.242532 | 1.84E-34 |
| 14368 | FZD6 | 1684.53722 | -2.8690107 | 0.23448338 | -12.235454 | 2.01E-34 |
| 11443 | chrnb1 | 623.095649 | -3.7899497 | 0.30989167 | -12.229918 | 2.15E-34 |
| 19201 | PSTPIP2 | 85.5424141 | -3.4164226 | 0.27935847 | -12.229529 | 2.16E-34 |
| 56532 | ripk3 | 1380.503 | 12.4734763 | 1.02010656 | 12.2276209 | 2.21E-34 |
| 14325 | Ftl1 | 15855.7109 | 1.43762696 | 0.11772542 | 12.2116951 | 2.69E-34 |
| 75705 | EIF4B | 10409.7985 | -0.7581167 | 0.06211114 | -12.20581 | 2.89E-34 |
| 24059 | SLCO2A1 | 6288.59973 | -5.5789709 | 0.45713744 | -12.204144 | 2.95E-34 |
| 67092 | GATM | 716.42807 | 3.16497429 | 0.25936483 | 12.202789 | 3.00E-34 |
| 20187 | RYK | 4248.73028 | 1.22429098 | 0.10042184 | 12.1914811 | 3.45E-34 |
| 100017 | LDLRAP1 | 989.780296 | 0.75350325 | 0.06183244 | 12.1862129 | 3.68E-34 |
| 100042150 | NRG2 | 107.930585 | -5.3091016 | 0.43609121 | -12.174292 | 4.26E-34 |
| 353187 | NR1D2 | 1109.14574 | 1.06112313 | 0.08722204 | 12.1657689 | 4.73E-34 |
| 239393 | LRP12 | 1528.46473 | 1.6980213 | 0.13963415 | 12.160502 | 5.04E-34 |
| 14012 | MPZL2 | 872.368326 | -13.198739 | 1.08785622 | -12.132797 | 7.08E-34 |
| 217653 | MIS18BP1 | 1555.63831 | 0.80919561 | 0.06674865 | 12.1230262 | 7.98E-34 |
| 18767 | PKIA | 517.294771 | -12.444541 | 1.02705574 | -12.116714 | 8.61E-34 |
| 27276 | PLEKHB1 | 270.929565 | -6.3352955 | 0.52305724 | -12.11205 | 9.12E-34 |
| 140743 | REM2 | 130.77309 | -6.8356583 | 0.56457854 | -12.107542 | 9.63E-34 |
| 93686 | RBFOX2 | 5663.44342 | 0.59566542 | 0.04922023 | 12.102044 | 1.03E-33 |
| 67211 | ARMC10 | 598.164715 | -1.0223969 | 0.08452509 | -12.09578 | 1.11E-33 |
| 66864 | CLEC14A | 585.321891 | 12.6738265 | 1.04795914 | 12.0938174 | 1.14E-33 |
| 140570 | PLXNB2 | 14219.1294 | -0.8773955 | 0.07257941 | -12.088766 | 1.21E-33 |
| 67016 | TBC1D2B | 1936.23788 | 2.50560062 | 0.20731655 | 12.0858686 | 1.25E-33 |
| 72831 | DHX30 | 2399.13089 | 0.62996254 | 0.05214452 | 12.0810878 | 1.33E-33 |
| 224703 | 41699 | 598.13293 | -1.1176792 | 0.09252207 | -12.080136 | 1.34E-33 |
| 22410 | WNT10B | 116.069415 | 7.31137398 | 0.60527174 | 12.0794901 | 1.36E-33 |
| 68176 | FAM212A | 93.4683857 | 3.80290167 | 0.31499439 | 12.0729189 | 1.47E-33 |

| | | | | | | |
|---|---|---|---|---|---|---|
| 20969 | sdc1 | 7692.55755 | -0.8953276 | 0.07420606 | -12.065425 | 1.61E-33 |
| 21983 | TPBG | 3793.25724 | 1.85270827 | 0.15365033 | 12.0579518 | 1.76E-33 |
| 217304 | CD300LB | 548.488329 | 5.07749687 | 0.42118605 | 12.0552352 | 1.82E-33 |
| 72469 | PLCD3 | 714.577973 | -1.8885635 | 0.15681581 | -12.043196 | 2.11E-33 |
| 64138 | CTSZ | 2497.53612 | 0.79060655 | 0.06565675 | 12.0415128 | 2.15E-33 |
| 16870 | LHX2 | 97.1873739 | 6.16709635 | 0.51253547 | 12.0325259 | 2.40E-33 |
| 93696 | CHRAC1 | 795.237078 | 0.975504 | 0.08123027 | 12.0091199 | 3.18E-33 |
| 71678 | BROX | 1183.61983 | -0.68354 | 0.05692544 | -12.007637 | 3.24E-33 |
| 22032 | TRAF4 | 2265.57368 | -0.8199867 | 0.06829566 | -12.006426 | 3.29E-33 |
| 76686 | CLIP3 | 165.903649 | 7.59702385 | 0.63289209 | 12.0036638 | 3.40E-33 |
| 195733 | GRHL1 | 70.6190153 | -3.4502615 | 0.28747308 | -12.002033 | 3.47E-33 |
| 65116 | prrg2 | 254.675454 | -1.5126907 | 0.12607527 | -11.998314 | 3.63E-33 |
| 228366 | LARGE2 | 154.413984 | -7.9386045 | 0.6618112 | -11.995271 | 3.76E-33 |
| 209200 | DTX3L | 743.351449 | -1.5600146 | 0.13005633 | -11.994915 | 3.78E-33 |
| 70028 | DOPEY2 | 417.631656 | -1.0238645 | 0.08546106 | -11.98048 | 4.50E-33 |
| 74559 | ELOVL7 | 469.159916 | -4.4848023 | 0.37436986 | -11.979603 | 4.54E-33 |
| 80889 | mesdc1 | 2229.14678 | 1.44780158 | 0.12087956 | 11.977224 | 4.68E-33 |
| 71957 | CPSF3L | 1567.03354 | 0.76153053 | 0.06360372 | 11.9730505 | 4.92E-33 |
| 56612 | PFDN5 | 712.693058 | -1.0016231 | 0.08369763 | -11.967162 | 5.28E-33 |
| 216795 | WNT9A | 153.540887 | -3.7401982 | 0.31276736 | -11.958403 | 5.87E-33 |
| 234515 | INPP4B | 441.757138 | 2.096344 | 0.17580168 | 11.9244819 | 8.82E-33 |
| 277010 | MARVELD1 | 1356.82147 | 2.22166251 | 0.18631256 | 11.9243837 | 8.83E-33 |
| 30928 | zbtb18 | 891.366567 | -2.0583571 | 0.1729776 | -11.899559 | 1.19E-32 |
| 57752 | TACC2 | 4455.31996 | 1.25677311 | 0.10562587 | 11.8983452 | 1.21E-32 |
| 98432 | PHLPP1 | 1050.73486 | -1.1340281 | 0.09556056 | -11.867115 | 1.75E-32 |
| 53312 | NUB1 | 2490.17433 | -0.6547682 | 0.05518493 | -11.864982 | 1.80E-32 |
| 66270 | fam134b | 2214.14213 | -0.4698564 | 0.03965888 | -11.847446 | 2.22E-32 |
| 18584 | PDE8A | 216.778906 | 4.83621454 | 0.40826841 | 11.845674 | 2.27E-32 |
| 18458 | PABPC1 | 52704.6267 | 0.9581117 | 0.08095815 | 11.8346541 | 2.58E-32 |
| 83602 | GTF2A1 | 1377.22344 | 0.77480692 | 0.06549607 | 11.8298224 | 2.74E-32 |
| 73699 | PPP2R1B | 2478.95716 | 0.67351942 | 0.05694031 | 11.8285167 | 2.78E-32 |
| 76773 | WDYHV1 | 573.954336 | 1.07607185 | 0.09101239 | 11.8233553 | 2.96E-32 |
| 56217 | MPP5 | 1466.27224 | -0.7163235 | 0.06062957 | -11.814755 | 3.28E-32 |
| 20540 | SLC7A7 | 199.974509 | -5.9035422 | 0.50002805 | -11.806422 | 3.62E-32 |
| 66743 | RNF220 | 1381.83338 | 0.93332789 | 0.07917589 | 11.7880315 | 4.50E-32 |
| 17954 | NAP1L2 | 133.614765 | 7.07636722 | 0.60031693 | 11.787719 | 4.52E-32 |
| 26562 | NCDN | 1769.13236 | 1.17940642 | 0.10011424 | 11.780606 | 4.91E-32 |
| 630499 | H2-K2 | 186.577139 | -1.615673 | 0.13720695 | -11.775446 | 5.22E-32 |
| 104184 | BLMH | 1797.32149 | 0.5516642 | 0.04686063 | 11.772445 | 5.41E-32 |
| 52662 | LDLRAD4 | 410.337964 | -1.674378 | 0.14230731 | -11.765932 | 5.85E-32 |
| 56325 | ABCB9 | 183.12434 | -4.826969 | 0.41038438 | -11.762068 | 6.12E-32 |
| 71586 | IFIH1 | 334.3018 | -2.6624882 | 0.22651171 | -11.754307 | 6.71E-32 |
| 213438 | A630033H20 | 131.63234 | 7.26519248 | 0.61883572 | 11.7400987 | 7.94E-32 |
| 223272 | ITGBL1 | 427.313794 | 12.219775 | 1.0412784 | 11.7353582 | 8.40E-32 |
| 104445 | CDC42EP1 | 1765.58599 | -1.466929 | 0.12525894 | -11.711172 | 1.12E-31 |

| | | | | | | |
|---|---|---|---|---|---|---|
| 69399 | 1700025G04 | 1692.22196 | -2.2173077 | 0.18947233 | -11.702541 | 1.24E-31 |
| 22021 | TPST1 | 725.083026 | 1.33447619 | 0.11419617 | 11.6858226 | 1.51E-31 |
| 21956 | TNNT2 | 1767.62211 | -2.8854144 | 0.24714051 | -11.675198 | 1.71E-31 |
| 17919 | MYO5B | 333.666323 | -9.7877663 | 0.83840023 | -11.674336 | 1.72E-31 |
| 60441 | MRPL38 | 1305.37209 | -0.6789514 | 0.05821463 | -11.662901 | 1.97E-31 |
| 110208 | PGD | 12311.0788 | 0.8609118 | 0.07383942 | 11.6592432 | 2.06E-31 |
| 107684 | CORO2A | 484.024719 | -3.5456419 | 0.30422971 | -11.654489 | 2.18E-31 |
| 223473 | NIPAL2 | 608.723694 | -11.957592 | 1.02603552 | -11.65417 | 2.19E-31 |
| 24017 | RNF13 | 1728.59242 | -0.536979 | 0.04609168 | -11.650235 | 2.29E-31 |
| 80909 | GATSL2 | 514.279 | -1.6162183 | 0.13875364 | -11.648115 | 2.35E-31 |
| 66844 | ORMDL2 | 718.118968 | -1.2564601 | 0.10795621 | -11.638609 | 2.62E-31 |
| 16188 | IL3RA | 106.147373 | -4.0525043 | 0.34819877 | -11.638479 | 2.63E-31 |
| 20842 | STAG1 | 3859.04236 | 0.7155997 | 0.06152391 | 11.6312446 | 2.86E-31 |
| 21453 | TCOF1 | 4880.37064 | 1.08935716 | 0.09366452 | 11.630414 | 2.89E-31 |
| 58887 | REPIN1 | 766.071202 | -0.9235321 | 0.0794632 | -11.622136 | 3.18E-31 |
| 269181 | MGAT4A | 765.636977 | -5.0164144 | 0.4318806 | -11.615281 | 3.45E-31 |
| 105785 | KDELR3 | 497.792954 | 2.14634289 | 0.18480399 | 11.6141585 | 3.49E-31 |
| 12159 | BMP4 | 119.406819 | -5.9436961 | 0.51178831 | -11.613583 | 3.52E-31 |
| 228061 | AGPS | 4987.99007 | -0.5219027 | 0.04494939 | -11.610896 | 3.63E-31 |
| 29861 | dpf1 | 129.274295 | 2.51876959 | 0.21701982 | 11.6061732 | 3.83E-31 |
| 102462 | IMP3 | 988.111907 | 0.74703122 | 0.06438438 | 11.6026783 | 3.99E-31 |
| 16889 | lipA | 1295.63073 | 0.70223084 | 0.06060066 | 11.5878422 | 4.75E-31 |
| 57890 | IL17RE | 346.594074 | -11.867235 | 1.0243806 | -11.584791 | 4.92E-31 |
| 72999 | INSIG2 | 1168.25316 | -1.0968713 | 0.09471098 | -11.581248 | 5.13E-31 |
| 110058 | syt17 | 276.472741 | -5.619729 | 0.48552807 | -11.574468 | 5.55E-31 |
| 14055 | EZH1 | 576.689101 | -1.0117345 | 0.08745415 | -11.568742 | 5.93E-31 |
| 207521 | DTX4 | 1792.17631 | -2.3344274 | 0.20184065 | -11.565695 | 6.15E-31 |
| 12182 | BST1 | 451.673674 | -7.9743955 | 0.68954235 | -11.564765 | 6.22E-31 |
| 108156 | MTHFD1 | 5390.19399 | 0.69274368 | 0.05990239 | 11.5645423 | 6.23E-31 |
| 213068 | TMEM71 | 57.6086128 | -5.0927182 | 0.44068352 | -11.556407 | 6.85E-31 |
| 100503884 | CCDC149 | 134.966474 | -7.4837877 | 0.64818164 | -11.545819 | 7.75E-31 |
| 210998 | FAM91A1 | 9038.27041 | 0.55790112 | 0.0483278 | 11.5441023 | 7.91E-31 |
| 76408 | ABCC3 | 152.408232 | -6.518711 | 0.56485357 | -11.540533 | 8.24E-31 |
| 78339 | TTYH3 | 1610.43687 | -0.6098057 | 0.05286557 | -11.535026 | 8.79E-31 |
| 12064 | BDNF | 1928.34693 | 3.68671372 | 0.31965353 | 11.533468 | 8.95E-31 |
| 15258 | HIPK2 | 1495.63514 | -1.1974399 | 0.1038284 | -11.532875 | 9.01E-31 |
| 74055 | PLCE1 | 64.0687007 | -4.6883885 | 0.40684117 | -11.523879 | 1.00E-30 |
| 21985 | TPD52 | 3405.86636 | 0.6134511 | 0.05331453 | 11.506265 | 1.23E-30 |
| 226751 | CDC42BPA | 5565.60269 | -1.1078717 | 0.09631936 | -11.502066 | 1.29E-30 |
| 72388 | RIPK4 | 411.030149 | -12.113578 | 1.05357241 | -11.497622 | 1.36E-30 |
| 22402 | WISP1 | 1170.29971 | 2.69958603 | 0.2349354 | 11.4907587 | 1.47E-30 |
| 15248 | HIC1 | 245.796984 | 4.69919147 | 0.40946526 | 11.4764108 | 1.73E-30 |
| 70835 | PRSS22 | 152.484328 | -6.6574073 | 0.58016542 | -11.475016 | 1.76E-30 |
| 66913 | KDELR2 | 4554.24942 | 0.70326954 | 0.06131981 | 11.4688808 | 1.89E-30 |
| 26570 | SLC7A11 | 12248.2108 | 1.77687188 | 0.15494395 | 11.4678366 | 1.91E-30 |

| | | | | | | |
|---|---|---|---|---|---|---|
| 110960 | TARS | 7580.97967 | 1.24665608 | 0.1087384 | 11.464727 | 1.98E-30 |
| 67860 | S100A16 | 1562.37514 | -10.097293 | 0.88120469 | -11.45851 | 2.13E-30 |
| 13808 | ENO3 | 950.388462 | 3.82422603 | 0.33399572 | 11.4499254 | 2.35E-30 |
| 56873 | LMBR1 | 399.989449 | -1.8438591 | 0.16105303 | -11.44877 | 2.39E-30 |
| 19252 | DUSP1 | 660.024523 | 2.296123 | 0.2011508 | 11.4149334 | 3.52E-30 |
| 22183 | Zrsr1 | 179.381883 | -8.4772353 | 0.74300602 | -11.409376 | 3.75E-30 |
| 216892 | SPNS2 | 1888.62268 | -4.0331268 | 0.35367698 | -11.403419 | 4.02E-30 |
| 23936 | LYNX1 | 427.869297 | -2.2959719 | 0.20143956 | -11.39782 | 4.29E-30 |
| 52377 | RCN3 | 890.80784 | 11.8383089 | 1.03898205 | 11.394142 | 4.47E-30 |
| 74552 | NIPAL3 | 153.360579 | -3.0073017 | 0.26415112 | -11.384777 | 4.98E-30 |
| 19731 | RGL1 | 1045.5604 | 3.01921896 | 0.26538954 | 11.3765561 | 5.47E-30 |
| 18626 | PER1 | 1604.48842 | -1.3908368 | 0.12227103 | -11.375031 | 5.57E-30 |
| 74103 | NEBL | 248.817338 | -4.211111 | 0.37035893 | -11.370351 | 5.88E-30 |
| 54672 | ADGRG3 | 85.575944 | -4.7013823 | 0.41359027 | -11.367246 | 6.09E-30 |
| 65079 | RTN4R | 96.0571648 | -2.965007 | 0.26103843 | -11.358508 | 6.73E-30 |
| 338349 | CNTLN | 434.159648 | 1.37189044 | 0.12078173 | 11.3584268 | 6.73E-30 |
| 23801 | ALOXE3 | 220.70246 | -4.9412089 | 0.4350486 | -11.357832 | 6.78E-30 |
| 106347 | ILDR1 | 378.686659 | -11.994521 | 1.05663246 | -11.351649 | 7.28E-30 |
| 238455 | MACC1 | 1047.92198 | -4.8333123 | 0.42625608 | -11.338987 | 8.41E-30 |
| 21976 | TOP3B | 1543.13065 | -0.7828138 | 0.06910206 | -11.328372 | 9.50E-30 |
| 386454 | RNF39 | 82.1527623 | -3.9699179 | 0.35053677 | -11.325254 | 9.84E-30 |
| 268564 | ZBTB1 | 1629.26032 | 0.68626827 | 0.06067428 | 11.3106953 | 1.16E-29 |
| 18011 | Neurl1a | 543.719076 | -0.9724376 | 0.0861057 | -11.293533 | 1.41E-29 |
| 21961 | TNS1 | 1923.60604 | -2.8273468 | 0.25037935 | -11.292253 | 1.43E-29 |
| 72657 | 2700094K13I | 1852.53463 | 0.88446355 | 0.0783966 | 11.2819125 | 1.61E-29 |
| 16155 | IL10RB | 626.576693 | -1.5917451 | 0.1412181 | -11.271537 | 1.81E-29 |
| 68790 | Fendrr | 66.7038609 | 5.14028969 | 0.45635696 | 11.2637477 | 1.98E-29 |
| 75146 | MFSD13A | 102.511037 | -2.9838405 | 0.26529602 | -11.247212 | 2.39E-29 |
| 76867 | RHBDD1 | 1576.36301 | -1.423497 | 0.12662329 | -11.241984 | 2.54E-29 |
| 16874 | LHX6 | 252.458351 | 2.2025508 | 0.19603308 | 11.2356079 | 2.73E-29 |
| 66371 | CHMP4C | 448.266061 | -11.516172 | 1.02533957 | -11.231569 | 2.85E-29 |
| 16177 | IL1R1 | 489.696697 | 3.38354294 | 0.30141019 | 11.2257086 | 3.05E-29 |
| 20361 | SEMA7A | 492.194567 | 3.87431186 | 0.3453813 | 11.2174916 | 3.35E-29 |
| 223453 | DAP | 10833.4411 | 1.18088429 | 0.10532175 | 11.2121602 | 3.55E-29 |
| 66111 | TMED3 | 1392.26768 | 1.19085799 | 0.10622066 | 11.2111708 | 3.59E-29 |
| 16592 | FABP5 | 358.045393 | -1.0465035 | 0.09339172 | -11.205527 | 3.83E-29 |
| 242721 | KLHDC7A | 288.514305 | -9.5789738 | 0.85511655 | -11.201951 | 3.99E-29 |
| 76974 | urah | 250.340398 | -8.9629149 | 0.80031317 | -11.19926 | 4.11E-29 |
| 56275 | RBM14 | 2526.29859 | 0.84492633 | 0.075517 | 11.1885581 | 4.64E-29 |
| 68832 | LDAH | 1747.63095 | 0.61610547 | 0.05510139 | 11.1813064 | 5.03E-29 |
| 98952 | FAM102A | 1120.01382 | -2.7394562 | 0.24539915 | -11.163267 | 6.17E-29 |
| 110074 | DUT | 2207.74886 | 0.65505421 | 0.05868337 | 11.1625181 | 6.22E-29 |
| 235036 | PPAN | 1769.75465 | 0.75087728 | 0.06727191 | 11.1618257 | 6.27E-29 |
| 69693 | POF1B | 112.703524 | -6.9894044 | 0.62639654 | -11.158114 | 6.54E-29 |
| 229003 | HELZ2 | 959.595268 | -1.5548054 | 0.13955806 | -11.140922 | 7.93E-29 |

| | | | | | | |
|---|---|---|---|---|---|---|
| 217325 | LLGL2 | 745.121385 | -3.7581861 | 0.33736842 | -11.139709 | 8.04E-29 |
| 14173 | FGF2 | 107.553818 | 2.55443759 | 0.22971952 | 11.1198108 | 1.00E-28 |
| 69590 | GPX8 | 1362.36228 | 1.38021839 | 0.12413841 | 11.1183831 | 1.02E-28 |
| 68135 | EIF3H | 5547.00701 | 0.79740546 | 0.07172911 | 11.1169014 | 1.04E-28 |
| 54396 | Irgm2 | 254.544469 | -3.2696363 | 0.29444384 | -11.104448 | 1.19E-28 |
| 224903 | SAFB | 2904.6615 | 0.52406402 | 0.04720003 | 11.1030449 | 1.21E-28 |
| 72787 | NDC1 | 3309.14735 | 1.32070604 | 0.11895544 | 11.1025276 | 1.22E-28 |
| 69906 | SLC25A32 | 1209.98034 | 1.60773653 | 0.14512434 | 11.078338 | 1.60E-28 |
| 20668 | SOX13 | 111.762813 | -7.4692302 | 0.67440616 | -11.07527 | 1.65E-28 |
| 11492 | ADAM19 | 2564.38082 | 5.11308311 | 0.46188063 | 11.0701397 | 1.75E-28 |
| 18477 | PRDX1 | 13260.9035 | 1.7742068 | 0.16036068 | 11.0638519 | 1.88E-28 |
| 19672 | RCN1 | 2369.40373 | 1.17766952 | 0.10646948 | 11.0610993 | 1.94E-28 |
| 97761 | SGSM2 | 605.426663 | -1.8106466 | 0.1637156 | -11.059708 | 1.97E-28 |
| 13110 | Cyp2j6 | 151.722081 | -3.1723358 | 0.28686501 | -11.058636 | 1.99E-28 |
| 171095 | IL17RC | 1011.30357 | -0.9004713 | 0.08143342 | -11.057761 | 2.01E-28 |
| 56811 | DKK2 | 643.583115 | 12.0891182 | 1.09344274 | 11.0560139 | 2.05E-28 |
| 67864 | YIPF4 | 1412.14616 | -0.5455553 | 0.04936985 | -11.050372 | 2.18E-28 |
| 74868 | tmem65 | 3087.10678 | 1.2222403 | 0.11063468 | 11.0475337 | 2.25E-28 |
| 217140 | SCRN2 | 198.651991 | -1.8248802 | 0.16529704 | -11.040006 | 2.45E-28 |
| 13018 | CTCF | 3532.65035 | 0.5154902 | 0.04669661 | 11.0391364 | 2.47E-28 |
| 213827 | ARCN1 | 4889.18653 | 0.48617328 | 0.04404727 | 11.0375356 | 2.52E-28 |
| 80914 | UCK2 | 2790.26434 | 1.22133239 | 0.11067889 | 11.0349177 | 2.59E-28 |
| 27103 | EIF2AK4 | 1162.547 | 0.89875711 | 0.08145801 | 11.033379 | 2.64E-28 |
| 22339 | VEGFA | 3376.19457 | 1.72788557 | 0.15663635 | 11.0311916 | 2.70E-28 |
| 219148 | FAM167A | 238.394801 | -11.327959 | 1.02691571 | -11.031051 | 2.71E-28 |
| 17850 | MUT | 787.769171 | -0.7459516 | 0.06765638 | -11.02559 | 2.88E-28 |
| 20591 | KDM5C | 4532.70486 | -0.4424964 | 0.04015368 | -11.020068 | 3.06E-28 |
| 71963 | CDCA4 | 3136.23707 | 0.96869955 | 0.08794336 | 11.0150392 | 3.23E-28 |
| 20724 | SERPINB5 | 875.886561 | -11.773283 | 1.07064435 | -10.996446 | 3.97E-28 |
| 192212 | PROM2 | 504.819497 | -12.409933 | 1.12898801 | -10.992086 | 4.17E-28 |
| 94112 | MED15 | 2059.77467 | -0.7114849 | 0.06474135 | -10.989651 | 4.29E-28 |
| 67155 | SMARCA2 | 1008.80705 | 1.34873647 | 0.12279757 | 10.9834133 | 4.59E-28 |
| 213603 | SLC44A3 | 375.585499 | -11.261971 | 1.02545728 | -10.982389 | 4.64E-28 |
| 15040 | H2-T23 | 109.063228 | -2.4086571 | 0.21934579 | -10.981095 | 4.71E-28 |
| 18602 | PADI4 | 730.073536 | -11.499641 | 1.04732768 | -10.979984 | 4.77E-28 |
| 381259 | TMEM237 | 904.524369 | -1.019661 | 0.09287979 | -10.978287 | 4.86E-28 |
| 75580 | ZBTB4 | 1065.26452 | -1.2406625 | 0.1130253 | -10.976857 | 4.94E-28 |
| 53331 | STX7 | 1550.74368 | -1.04918 | 0.0955999 | -10.974698 | 5.06E-28 |
| 76788 | KLHDC10 | 2625.88215 | -0.5764913 | 0.0525339 | -10.973701 | 5.11E-28 |
| 69833 | POLR2F | 1391.5369 | -0.817702 | 0.07462309 | -10.957761 | 6.10E-28 |
| 102623 | #N/A | 146.484458 | -8.1780361 | 0.74634051 | -10.957513 | 6.12E-28 |
| 12033 | BCAP29 | 1162.00093 | 0.72676942 | 0.06633625 | 10.9558415 | 6.23E-28 |
| 230872 | CROCC | 239.240898 | 1.18902074 | 0.10854374 | 10.9543004 | 6.34E-28 |
| 74123 | FOXP4 | 2578.70079 | -0.646755 | 0.05907378 | -10.948258 | 6.77E-28 |
| 80297 | SPTBN4 | 50.6927873 | 3.2042704 | 0.29295603 | 10.9377177 | 7.61E-28 |

| | | | | | | |
|---|---|---|---|---|---|---|
| 11732 | ANK | 25880.5409 | 2.76466483 | 0.25276948 | 10.9374945 | 7.63E-28 |
| 77045 | BCL7A | 683.956593 | 0.92089932 | 0.08420682 | 10.9361614 | 7.74E-28 |
| 212307 | MAPRE2 | 2125.07007 | 0.57058174 | 0.05218728 | 10.9333489 | 7.98E-28 |
| 12890 | CPLX2 | 327.672021 | 2.2316375 | 0.20420741 | 10.9282883 | 8.44E-28 |
| 269831 | TSPAN12 | 237.270522 | -11.319698 | 1.03738086 | -10.911805 | 1.01E-27 |
| 28193 | REEP3 | 4804.43431 | -0.7968113 | 0.07304454 | -10.908568 | 1.05E-27 |
| 236930 | ercc6l | 1883.50185 | 0.62518116 | 0.05732309 | 10.9062717 | 1.08E-27 |
| 244049 | MCTP2 | 90.9429687 | -5.2866796 | 0.48507341 | -10.898721 | 1.17E-27 |
| 227545 | PROSER2 | 364.495888 | -1.997491 | 0.18330131 | -10.897309 | 1.19E-27 |
| 380705 | TMEM102 | 214.751517 | -11.176078 | 1.02592974 | -10.89361 | 1.24E-27 |
| 17311 | Kitl | 3908.46304 | 1.61033881 | 0.14798775 | 10.8815686 | 1.41E-27 |
| 17472 | GBP4 | 71.2733998 | -5.1033528 | 0.4689954 | -10.881456 | 1.41E-27 |
| 170753 | Zfp704 | 831.062555 | 1.36574854 | 0.12552483 | 10.8803057 | 1.43E-27 |
| 14745 | LPAR1 | 1600.89785 | 2.62606915 | 0.24150067 | 10.873962 | 1.53E-27 |
| 18764 | PKD2 | 3663.32129 | 1.04149685 | 0.09583132 | 10.8680217 | 1.64E-27 |
| 99526 | USP53 | 481.842179 | -0.8289097 | 0.07629006 | -10.865239 | 1.69E-27 |
| 207474 | Kctd12b | 280.430428 | -5.7724134 | 0.53157355 | -10.859106 | 1.81E-27 |
| 22772 | ZIC2 | 200.940269 | 11.1316803 | 1.0258846 | 10.8508114 | 1.98E-27 |
| 14615 | GJC1 | 864.332914 | 6.11243356 | 0.56419356 | 10.8339301 | 2.38E-27 |
| 67099 | METTL21A | 483.981774 | 0.91255207 | 0.08428794 | 10.8266026 | 2.58E-27 |
| 232223 | TXNRD3 | 586.570851 | 1.39778322 | 0.1293435 | 10.8067526 | 3.20E-27 |
| 70717 | MEDAG | 106.544246 | 5.27728399 | 0.48866251 | 10.7994452 | 3.46E-27 |
| 70310 | PLSCR3 | 1790.10932 | -0.7283434 | 0.06744942 | -10.798365 | 3.50E-27 |
| 11544 | ADPRH | 1104.6102 | 1.04777979 | 0.09710496 | 10.790178 | 3.83E-27 |
| 107358 | TM9SF3 | 11846.608 | -0.3570125 | 0.03309012 | -10.789096 | 3.88E-27 |
| 18685 | PHTF1 | 1434.54558 | -0.6376321 | 0.05912534 | -10.784412 | 4.08E-27 |
| 22634 | PLAGL1 | 221.048786 | 2.09865958 | 0.19462417 | 10.7831393 | 4.14E-27 |
| 232286 | TMF1 | 2940.91043 | 0.7349467 | 0.06821066 | 10.7746603 | 4.53E-27 |
| 69674 | MIF4GD | 312.334852 | -1.2783934 | 0.11866952 | -10.772719 | 4.63E-27 |
| 16423 | CD47 | 4035.63969 | -2.2079823 | 0.20497414 | -10.772004 | 4.67E-27 |
| 14238 | FOXF2 | 290.894103 | 3.32031573 | 0.30835976 | 10.7676686 | 4.89E-27 |
| 66938 | SH3D21 | 59.6649805 | -3.3656059 | 0.31265573 | -10.764574 | 5.06E-27 |
| 76800 | USP42 | 835.474598 | 0.7424122 | 0.06899754 | 10.7599812 | 5.32E-27 |
| 545886 | Gm5886 | 191.224534 | 11.0598384 | 1.02848436 | 10.7535309 | 5.70E-27 |
| 225791 | ZADH2 | 3015.15194 | 0.88230204 | 0.08204781 | 10.7535112 | 5.70E-27 |
| 16206 | LRIG1 | 9545.00157 | 0.49709977 | 0.04625936 | 10.7459288 | 6.19E-27 |
| 26441 | PSMA4 | 1492.15881 | 0.62610348 | 0.05831188 | 10.7371513 | 6.81E-27 |
| 231642 | ALKBH2 | 109.309797 | -2.9910116 | 0.27867446 | -10.732995 | 7.12E-27 |
| 227580 | C1QL3 | 165.643842 | -6.0914398 | 0.56761612 | -10.731619 | 7.23E-27 |
| 74895 | CCDC181 | 232.28985 | -1.5431298 | 0.14384188 | -10.727959 | 7.52E-27 |
| 21885 | TLE1 | 1158.8998 | 1.14837079 | 0.10714559 | 10.7178538 | 8.39E-27 |
| 67281 | RPL37 | 1550.89498 | 1.06988447 | 0.09986346 | 10.7134724 | 8.80E-27 |
| 224022 | SLC7A4 | 96.1994821 | -4.0101803 | 0.3744042 | -10.710832 | 9.05E-27 |
| 58894 | Zfp862-ps | 741.787967 | 0.95004959 | 0.08873073 | 10.7071085 | 9.43E-27 |
| 100043468 | Zfp955b | 390.461699 | 1.21406755 | 0.11341562 | 10.7045886 | 9.69E-27 |

| | | | | | | |
|---|---|---|---|---|---|---|
| 110958 | m1ap | 181.66839 | -3.8416277 | 0.35891856 | -10.703341 | 9.82E-27 |
| 20660 | SORL1 | 444.893286 | -10.78815 | 1.00805042 | -10.701994 | 9.96E-27 |
| 67452 | PNPLA8 | 1440.21473 | 0.76857665 | 0.07183952 | 10.6985219 | 1.03E-26 |
| 18023 | NFE2L1 | 10838.2521 | -0.3389917 | 0.03173955 | -10.680419 | 1.26E-26 |
| 171170 | MBNL3 | 1512.5832 | -1.7659155 | 0.16541231 | -10.675841 | 1.32E-26 |
| 20564 | SLIT3 | 176.584897 | 6.21878786 | 0.58252204 | 10.6756268 | 1.32E-26 |
| 50868 | KEAP1 | 1961.90306 | 0.48435534 | 0.04538556 | 10.6720152 | 1.38E-26 |
| 21946 | PGLYRP1 | 749.808212 | -12.258791 | 1.14884379 | -10.670546 | 1.40E-26 |
| 235442 | RAB8B | 1703.33238 | 2.06398269 | 0.19344198 | 10.6697767 | 1.41E-26 |
| 11746 | ANXA4 | 5269.95202 | -0.9239251 | 0.08660184 | -10.668653 | 1.43E-26 |
| 625249 | GPX4 | 494.492454 | -0.9346229 | 0.0877471 | -10.651324 | 1.72E-26 |
| 102414 | CLK3 | 1312.94109 | 0.66049432 | 0.0620314 | 10.6477414 | 1.79E-26 |
| 68731 | RBFA | 616.058807 | -0.7683685 | 0.0721654 | -10.647325 | 1.79E-26 |
| 230972 | ARHGEF16 | 206.589869 | -9.0993424 | 0.85578606 | -10.63273 | 2.10E-26 |
| 71774 | SHROOM1 | 388.181243 | -1.6408992 | 0.15438263 | -10.628781 | 2.19E-26 |
| 13032 | CTSC | 340.62039 | -11.841827 | 1.11421647 | -10.627941 | 2.21E-26 |
| 219140 | SPATA13 | 317.559148 | 1.89290658 | 0.17812423 | 10.6268897 | 2.23E-26 |
| 18019 | NFATC2 | 214.285592 | -7.5902197 | 0.71466573 | -10.620657 | 2.39E-26 |
| 116870 | MTA1 | 4479.51759 | 0.97581318 | 0.09191858 | 10.6160605 | 2.51E-26 |
| 22042 | TFRC | 14323.1012 | -0.9364806 | 0.08825955 | -10.61053 | 2.66E-26 |
| 23943 | ESYT1 | 1859.87967 | -1.9644115 | 0.18522493 | -10.605546 | 2.81E-26 |
| 239528 | AGO2 | 6154.25825 | 1.03555806 | 0.09769305 | 10.6001202 | 2.98E-26 |
| 17761 | MAP7 | 284.274785 | -10.859916 | 1.02520015 | -10.592971 | 3.21E-26 |
| 14451 | GAS1 | 1214.83202 | 4.80305249 | 0.4541257 | 10.5764825 | 3.83E-26 |
| 69226 | SNX24 | 344.509761 | -1.1133893 | 0.10529879 | -10.573619 | 3.95E-26 |
| 11768 | AP1M2 | 139.745975 | -8.1115396 | 0.76759679 | -10.567449 | 4.22E-26 |
| 18025 | nfe2l3 | 148.735539 | -7.4066964 | 0.70115536 | -10.56356 | 4.40E-26 |
| 106052 | FBXO4 | 758.196838 | 1.06096715 | 0.10049349 | 10.5575714 | 4.69E-26 |
| 17772 | MTM1 | 294.798454 | -1.8023042 | 0.17085393 | -10.548802 | 5.14E-26 |
| 66060 | CYSTM1 | 416.447429 | -3.8155826 | 0.36171722 | -10.548524 | 5.16E-26 |
| 226525 | RASAL2 | 2247.52072 | -1.0965299 | 0.10402749 | -10.540771 | 5.60E-26 |
| 22592 | ERCC5 | 1024.78962 | -0.884023 | 0.08390594 | -10.535881 | 5.90E-26 |
| 12394 | RUNX1 | 3178.07195 | -0.9988301 | 0.0948658 | -10.528875 | 6.36E-26 |
| 20465 | SIM2 | 58.5382333 | -4.3311147 | 0.41168135 | -10.520551 | 6.95E-26 |
| 623230 | TMEM200B | 272.764866 | 9.55555329 | 0.90836545 | 10.5195032 | 7.02E-26 |
| 12364 | CASP12 | 452.458031 | -1.2849569 | 0.12217749 | -10.517133 | 7.20E-26 |
| 57278 | bcam | 57.5763986 | -5.8246246 | 0.5543894 | -10.506378 | 8.07E-26 |
| 83701 | SRRT | 5628.89777 | 0.36581814 | 0.03482667 | 10.5039661 | 8.28E-26 |
| 13197 | GADD45A | 527.524288 | -1.2130045 | 0.11554645 | -10.497981 | 8.82E-26 |
| 14062 | F2R | 4236.19177 | 1.5827607 | 0.15084647 | 10.4925275 | 9.35E-26 |
| 64704 | HTRA2 | 2331.76832 | 0.73735537 | 0.07030459 | 10.4880123 | 9.81E-26 |
| 56338 | TXNIP | 1102.67364 | -2.6431281 | 0.25209685 | -10.484574 | 1.02E-25 |
| 19357 | RAD21 | 16034.783 | 1.02584818 | 0.09785899 | 10.482922 | 1.03E-25 |
| 231832 | TMEM184A | 135.793233 | -8.0753108 | 0.77034735 | -10.482688 | 1.04E-25 |
| 21419 | TFAP2B | 398.132832 | -12.066968 | 1.15203919 | -10.474443 | 1.13E-25 |

| | | | | | | |
|---|---|---|---|---|---|---|
| 13056 | CYB561 | 584.368428 | -5.7403703 | 0.5481965 | -10.471373 | 1.17E-25 |
| 194590 | REPS2 | 107.869431 | -4.4815192 | 0.4280359 | -10.469961 | 1.19E-25 |
| 12985 | CSF3 | 829.209017 | -2.5877721 | 0.24751121 | -10.455171 | 1.39E-25 |
| 26919 | Zfp346 | 1031.24246 | 0.87277092 | 0.08349958 | 10.4523987 | 1.43E-25 |
| 76448 | PPP1R18 | 3786.37444 | 0.445191 | 0.04260798 | 10.4485352 | 1.49E-25 |
| 69717 | Gm10499 | 471.808428 | -1.9457562 | 0.18631291 | -10.443486 | 1.57E-25 |
| 216438 | 41706 | 264.008438 | -2.6418907 | 0.25299464 | -10.442477 | 1.59E-25 |
| 19885 | rorc | 167.786334 | -10.821469 | 1.03645423 | -10.440855 | 1.61E-25 |
| 231712 | TRAFD1 | 646.01238 | -0.9213309 | 0.08828185 | -10.436244 | 1.69E-25 |
| 66659 | ACP6 | 514.920817 | -1.2078371 | 0.11575881 | -10.434083 | 1.73E-25 |
| 73296 | RHOBTB3 | 3587.53452 | 1.28303032 | 0.12296698 | 10.4339414 | 1.74E-25 |
| 21912 | TSPAN7 | 559.014833 | -12.556704 | 1.20362283 | -10.432424 | 1.76E-25 |
| 68070 | PDZD2 | 121.073861 | 4.76648873 | 0.4568983 | 10.4322749 | 1.77E-25 |
| 17536 | MEIS2 | 488.244583 | 2.0954069 | 0.20088668 | 10.4307907 | 1.79E-25 |
| 17281 | FYCO1 | 891.434059 | 0.7395936 | 0.07095741 | 10.4230636 | 1.95E-25 |
| 52538 | ACAA2 | 891.120378 | -0.7161342 | 0.06872999 | -10.41953 | 2.02E-25 |
| 109264 | ME3 | 107.209623 | -4.6733956 | 0.44853178 | -10.419319 | 2.02E-25 |
| 214459 | FNBP1L | 4334.63527 | -1.3264941 | 0.12735805 | -10.415471 | 2.11E-25 |
| 242505 | RASEF | 170.191089 | -10.841861 | 1.04140793 | -10.410772 | 2.21E-25 |
| 16572 | KIF5A | 80.9709249 | -2.6635555 | 0.25597246 | -10.405633 | 2.34E-25 |
| 77305 | WDR82 | 3690.43921 | 0.57955635 | 0.05569641 | 10.4056317 | 2.34E-25 |
| 20527 | SLC2A3 | 97.787099 | 5.73277834 | 0.55102159 | 10.4039087 | 2.38E-25 |
| 54427 | DNMT3L | 69.8143905 | 6.56788348 | 0.63154663 | 10.3996809 | 2.49E-25 |
| 170762 | NUP155 | 6044.1824 | 0.85185684 | 0.08191862 | 10.3988188 | 2.51E-25 |
| 23947 | MID2 | 1120.20877 | -2.5563347 | 0.24583646 | -10.398518 | 2.52E-25 |
| 105245 | TXNDC5 | 9588.08462 | 1.13487462 | 0.1091642 | 10.3960329 | 2.58E-25 |
| 76560 | PRSS8 | 156.08696 | -10.71709 | 1.03103343 | -10.394512 | 2.63E-25 |
| 29858 | PMM1 | 1049.09561 | -1.3684218 | 0.13165262 | -10.394186 | 2.64E-25 |
| 213056 | FAM126B | 809.458024 | -1.7639161 | 0.16973252 | -10.392328 | 2.69E-25 |
| 17121 | MXD3 | 269.430157 | 1.21374965 | 0.11680633 | 10.3911287 | 2.72E-25 |
| 142682 | ZCCHC14 | 4227.13704 | 1.27696667 | 0.12295707 | 10.3854679 | 2.89E-25 |
| 11433 | ACP5 | 265.727153 | -6.7022929 | 0.6456494 | -10.3807 | 3.04E-25 |
| 74762 | MDGA1 | 65.382956 | 3.31255988 | 0.3191212 | 10.3802564 | 3.05E-25 |
| 234740 | TMEM231 | 241.076574 | -1.069458 | 0.10305448 | -10.377599 | 3.14E-25 |
| 100041680 | Gm3470 | 59.3785934 | 4.41623341 | 0.42590378 | 10.3690872 | 3.43E-25 |
| 54200 | SULT2B1 | 167.953361 | -4.2100945 | 0.40605725 | -10.368229 | 3.46E-25 |
| 70675 | VCPIP1 | 2360.70033 | -0.5074049 | 0.04896296 | -10.363035 | 3.65E-25 |
| 58809 | RNASE4 | 267.30634 | 1.62076448 | 0.15642616 | 10.3612112 | 3.72E-25 |
| 242585 | SLC35D1 | 562.072595 | 1.03275914 | 0.09969323 | 10.3593704 | 3.79E-25 |
| 56417 | ADAR | 1756.14512 | -0.9134459 | 0.08824122 | -10.351691 | 4.11E-25 |
| 74048 | Vsir | 156.765592 | -4.6560458 | 0.4498981 | -10.349112 | 4.22E-25 |
| 116871 | mta3 | 988.820275 | -0.9811594 | 0.0948537 | -10.343923 | 4.46E-25 |
| 230967 | CEP104 | 1098.69678 | 0.79406786 | 0.07677481 | 10.3428176 | 4.51E-25 |
| 75553 | ZC3H14 | 3957.39769 | 0.56560759 | 0.05470008 | 10.3401595 | 4.64E-25 |
| 68964 | CTC1 | 1578.18548 | -0.5992387 | 0.05795954 | -10.338913 | 4.70E-25 |

| | | | | | | |
|---|---|---|---|---|---|---|
| 231293 | CWH43 | 225.469387 | -11.246364 | 1.08812798 | -10.335516 | 4.87E-25 |
| 320405 | CADPS2 | 309.316615 | -10.98063 | 1.06298136 | -10.33003 | 5.15E-25 |
| 72778 | DNAJC22 | 203.679411 | -4.9126719 | 0.47576757 | -10.325781 | 5.39E-25 |
| 67515 | TTC33 | 800.122695 | 1.23751255 | 0.11986764 | 10.3239922 | 5.49E-25 |
| 117592 | B3GALT6 | 485.312476 | 1.6438521 | 0.15933329 | 10.3170659 | 5.90E-25 |
| 71529 | KAZN | 223.767882 | -11.235436 | 1.08914908 | -10.315793 | 5.98E-25 |
| 75991 | SLAIN2 | 2956.99394 | 0.4451351 | 0.04316529 | 10.31234 | 6.20E-25 |
| 11459 | ACTA1 | 444.54448 | -6.2306135 | 0.6044203 | -10.308412 | 6.46E-25 |
| 18018 | NFATC1 | 976.304024 | -0.9705633 | 0.09418331 | -10.305045 | 6.69E-25 |
| 107242 | AI837181 | 1607.72247 | 0.67869558 | 0.06588093 | 10.3018512 | 6.91E-25 |
| 70445 | CD248 | 612.537128 | 6.98254148 | 0.67814901 | 10.2964709 | 7.31E-25 |
| 208884 | ZDHHC9 | 1149.53861 | -0.8128032 | 0.07894287 | -10.296095 | 7.34E-25 |
| 100861925 | #N/A | 340.495209 | -10.402299 | 1.01097274 | -10.289396 | 7.87E-25 |
| 11856 | ARHGAP6 | 524.677809 | -4.1594917 | 0.40425436 | -10.289293 | 7.88E-25 |
| 544817 | ARHGAP27 | 1095.78329 | -4.5259141 | 0.43993699 | -10.287642 | 8.01E-25 |
| 234311 | DDX60 | 97.0946381 | -2.5679118 | 0.24968662 | -10.284539 | 8.27E-25 |
| 12363 | CASP4 | 132.719955 | -3.0209189 | 0.29380926 | -10.281905 | 8.50E-25 |
| 20707 | Serpinb9c | 60.2458261 | -5.8825289 | 0.57247191 | -10.275664 | 9.07E-25 |
| 73442 | HSPA12A | 232.384231 | 2.92702062 | 0.28497434 | 10.2711724 | 9.50E-25 |
| 170833 | HOOK2 | 786.243827 | -2.0833983 | 0.20287848 | -10.269193 | 9.70E-25 |
| 114301 | PALMD | 146.201344 | -10.621345 | 1.03429637 | -10.269151 | 9.71E-25 |
| 242700 | IFNLR1 | 140.398388 | -10.564308 | 1.02904456 | -10.266133 | 1.00E-24 |
| 100504231 | Gm15708 | 110.962967 | 3.90996491 | 0.3809186 | 10.264568 | 1.02E-24 |
| 14284 | FOSL2 | 6756.04726 | 1.48566779 | 0.14485803 | 10.2560265 | 1.11E-24 |
| 67425 | EPS8L1 | 137.92925 | -10.537679 | 1.02773156 | -10.253338 | 1.14E-24 |
| 69574 | CMBL | 45.2776841 | -4.1746785 | 0.40720904 | -10.25193 | 1.16E-24 |
| 69719 | CAD | 9087.85231 | 0.70377883 | 0.06873332 | 10.2392671 | 1.32E-24 |
| 67122 | NRARP | 59.5513502 | -3.9896201 | 0.38977998 | -10.23557 | 1.37E-24 |
| 216835 | USP43 | 88.4893192 | -6.4504408 | 0.63026739 | -10.234451 | 1.39E-24 |
| 20744 | STRBP | 1160.79878 | -1.2971904 | 0.12679788 | -10.230379 | 1.45E-24 |
| 107581 | COL16A1 | 3809.7902 | 2.33646333 | 0.22845134 | 10.2274004 | 1.49E-24 |
| 20509 | SLC19A1 | 1468.98772 | 1.02206867 | 0.09995267 | 10.2255264 | 1.52E-24 |
| 70415 | STK26 | 142.863951 | -10.589321 | 1.03571024 | -10.224212 | 1.54E-24 |
| 76252 | ATP6V0E2 | 357.715986 | -3.4738841 | 0.33990582 | -10.220137 | 1.61E-24 |
| 57377 | MOGS | 2291.29905 | 1.55333251 | 0.15206771 | 10.2147425 | 1.70E-24 |
| 18974 | POLE2 | 856.568737 | 0.95687796 | 0.09373084 | 10.2087842 | 1.81E-24 |
| 276770 | EIF5A | 19091.0531 | 0.38699496 | 0.03793818 | 10.2006727 | 1.97E-24 |
| 108012 | AP1S2 | 623.847119 | -1.0768523 | 0.10563187 | -10.194388 | 2.10E-24 |
| 56248 | AK3 | 1130.05378 | -3.1141608 | 0.30551436 | -10.193173 | 2.13E-24 |
| 16150 | IKBKB | 1469.78275 | -0.6983235 | 0.0685653 | -10.184794 | 2.32E-24 |
| 67846 | TMEM39A | 780.84186 | 0.84495454 | 0.08299742 | 10.1804911 | 2.42E-24 |
| 231287 | ATP10D | 1380.71497 | 1.74683913 | 0.1715952 | 10.1799997 | 2.44E-24 |
| 12759 | CLU | 3709.93987 | -3.4402891 | 0.33809567 | -10.17549 | 2.55E-24 |
| 56447 | COPZ1 | 2118.85659 | -1.0709549 | 0.10525118 | -10.175229 | 2.56E-24 |
| 22027 | HSP90B1 | 31091.9258 | 0.36488755 | 0.03586677 | 10.1734141 | 2.61E-24 |

| | | | | | | |
|---|---|---|---|---|---|---|
| 68667 | TRPM4 | 486.999733 | -1.7046579 | 0.16758149 | -10.172113 | 2.64E-24 |
| 12903 | CRABP1 | 1864.62236 | 13.6239937 | 1.33958226 | 10.17033 | 2.69E-24 |
| 114774 | PAWR | 1244.52755 | -1.7309924 | 0.17026238 | -10.166617 | 2.79E-24 |
| 72789 | VEPH1 | 96.9067042 | -3.0436345 | 0.29940061 | -10.165759 | 2.82E-24 |
| 215015 | FAM20B | 2932.79465 | -0.5252751 | 0.05169085 | -10.161858 | 2.93E-24 |
| 20349 | SEMA3E | 1017.04973 | -3.1585841 | 0.31088717 | -10.159905 | 2.99E-24 |
| 235320 | ZBTB16 | 303.727903 | -10.23527 | 1.0075141 | -10.158935 | 3.02E-24 |
| 57783 | TNIP1 | 1128.54055 | -0.800435 | 0.07879436 | -10.158532 | 3.04E-24 |
| 81879 | TFCP2L1 | 381.307391 | -8.9843872 | 0.88452441 | -10.157308 | 3.07E-24 |
| 103266 | TMEM263 | 1500.16217 | 0.95826524 | 0.09437028 | 10.154312 | 3.17E-24 |
| 232430 | CREBL2 | 261.499451 | -1.752291 | 0.17265669 | -10.14899 | 3.35E-24 |
| 70380 | MOSPD1 | 327.328775 | -1.0795613 | 0.10638465 | -10.147717 | 3.39E-24 |
| 11676 | ALDOC | 117.630578 | -4.7238912 | 0.46568827 | -10.143891 | 3.53E-24 |
| 319859 | E030011O05 | 238.61137 | -1.7969821 | 0.17723383 | -10.139047 | 3.71E-24 |
| 107029 | ME2 | 2687.31443 | 0.69397877 | 0.06845894 | 10.1371537 | 3.78E-24 |
| 13537 | DUSP2 | 156.448542 | 7.33316215 | 0.72348029 | 10.1359529 | 3.83E-24 |
| 53791 | TLR5 | 134.577465 | -10.501082 | 1.03616285 | -10.134586 | 3.88E-24 |
| 14228 | FKBP4 | 6718.69304 | 0.75928069 | 0.07495462 | 10.1298717 | 4.07E-24 |
| 21810 | TGFBI | 217.446587 | 4.52429818 | 0.44667977 | 10.1287287 | 4.12E-24 |
| 246730 | Oas1a | 134.507971 | -3.0378686 | 0.29999595 | -10.126365 | 4.22E-24 |
| 52815 | LDHD | 74.6771816 | -6.8750727 | 0.67899422 | -10.125377 | 4.26E-24 |
| 243374 | GIMAP8 | 132.740641 | -10.483259 | 1.03570124 | -10.121895 | 4.42E-24 |
| 56736 | RNF14 | 2628.61054 | -0.7144079 | 0.07058589 | -10.121115 | 4.45E-24 |
| 22401 | ZMAT3 | 4298.47619 | -0.9178907 | 0.09072353 | -10.11745 | 4.62E-24 |
| 14462 | GATA3 | 2065.9535 | -1.0460037 | 0.10341297 | -10.114821 | 4.75E-24 |
| 230766 | FAM167B | 149.067021 | 10.700757 | 1.05850364 | 10.1093247 | 5.02E-24 |
| 69065 | CHAC1 | 762.80673 | 0.87187229 | 0.08624876 | 10.1088096 | 5.05E-24 |
| 69554 | KLHDC2 | 2881.97317 | 0.65875441 | 0.0651721 | 10.1079205 | 5.10E-24 |
| 329260 | DENND1B | 915.835503 | -1.1906193 | 0.11779555 | -10.107507 | 5.12E-24 |
| 214895 | LMAN2L | 325.80948 | -1.2576216 | 0.12448506 | -10.10259 | 5.38E-24 |
| 52502 | CARHSP1 | 2939.58238 | -0.6820809 | 0.0675416 | -10.098679 | 5.60E-24 |
| 67729 | mansc1 | 190.677409 | -11.005651 | 1.08993915 | -10.097491 | 5.67E-24 |
| 12297 | CACNB3 | 1377.20207 | -0.7343522 | 0.07278668 | -10.089101 | 6.17E-24 |
| 12554 | CDH13 | 153.934686 | -10.695315 | 1.06022301 | -10.087798 | 6.26E-24 |
| 103012 | Firre | 670.477276 | -2.2040245 | 0.21878121 | -10.074103 | 7.19E-24 |
| 55942 | SERTAD1 | 1171.27961 | 0.67034588 | 0.06655788 | 10.0716224 | 7.38E-24 |
| 414084 | TNIP3 | 229.833875 | -11.274113 | 1.11961436 | -10.069639 | 7.53E-24 |
| 317758 | Gimap9 | 135.406993 | -8.486525 | 0.8429264 | -10.067931 | 7.66E-24 |
| 215436 | SLC35E3 | 758.674085 | -1.1577217 | 0.11500158 | -10.067007 | 7.73E-24 |
| 80880 | KANK3 | 87.3712471 | -2.2454252 | 0.22309854 | -10.064724 | 7.91E-24 |
| 106582 | NRM | 395.11904 | 1.19086053 | 0.11839139 | 10.0586751 | 8.41E-24 |
| 320982 | ARL4C | 3864.84804 | -2.5722171 | 0.2557961 | -10.055732 | 8.67E-24 |
| 52348 | VPS37A | 1854.80015 | -0.8900037 | 0.08851601 | -10.05472 | 8.76E-24 |
| 16776 | LAMA5 | 8274.48075 | -5.0372891 | 0.50118 | -10.050858 | 9.11E-24 |
| 98660 | ATP1A2 | 212.773987 | -11.162812 | 1.11099511 | -10.04758 | 9.42E-24 |

| | | | | | | |
|---|---|---|---|---|---|---|
| 217882 | CEP170B | 10156.174 | -0.9790447 | 0.09744841 | -10.0468 | 9.49E-24 |
| 381314 | iars2 | 3444.70245 | -0.4320725 | 0.04302218 | -10.043018 | 9.86E-24 |
| 18188 | NRTN | 43.0553445 | -3.8631822 | 0.38470203 | -10.042012 | 9.96E-24 |
| 76952 | NT5C2 | 1505.20769 | -1.4189306 | 0.14136398 | -10.037427 | 1.04E-23 |
| 22761 | ZFPM1 | 731.806751 | -1.5672726 | 0.15621511 | -10.032785 | 1.09E-23 |
| 55963 | SLC1A4 | 2167.33419 | 1.10229028 | 0.10993062 | 10.0271453 | 1.16E-23 |
| 20271 | SCN5A | 140.178539 | -10.560392 | 1.05324448 | -10.026535 | 1.17E-23 |
| 11363 | ACADL | 4335.28242 | -1.3421303 | 0.13396599 | -10.018441 | 1.26E-23 |
| 16391 | IRF9 | 323.664006 | -1.7249559 | 0.17218185 | -10.018221 | 1.27E-23 |
| 71653 | SHTN1 | 843.680098 | -1.8280893 | 0.18254903 | -10.014237 | 1.32E-23 |
| 232288 | FRMD4B | 284.918763 | -3.6158979 | 0.36129452 | -10.008172 | 1.40E-23 |
| 215632 | PSD4 | 286.336893 | -10.152106 | 1.01548098 | -9.9973377 | 1.57E-23 |
| 236312 | Ifi209 | 54.5223707 | -3.9711529 | 0.39769943 | -9.9853121 | 1.77E-23 |
| 21682 | TEC | 248.663229 | -2.1427136 | 0.21468075 | -9.9809303 | 1.85E-23 |
| 66274 | LYRM9 | 210.671976 | -1.1735574 | 0.11760158 | -9.9790959 | 1.88E-23 |
| 14635 | GALK1 | 1633.78614 | 0.9731959 | 0.09754293 | 9.97710324 | 1.92E-23 |
| 12837 | COL8A1 | 8048.2415 | 1.24715323 | 0.12505604 | 9.97275513 | 2.01E-23 |
| 217169 | TNS4 | 1962.48768 | -4.7544081 | 0.47676017 | -9.9723264 | 2.01E-23 |
| 69073 | KDF1 | 116.496646 | -10.292836 | 1.03238159 | -9.9699916 | 2.06E-23 |
| 13797 | EMX2 | 81.4923304 | 7.37972745 | 0.74050281 | 9.96583314 | 2.15E-23 |
| 16859 | lgals9 | 1728.23365 | -2.0142608 | 0.20217643 | -9.9628863 | 2.22E-23 |
| 80794 | CBLC | 183.623168 | -10.22931 | 1.02795465 | -9.9511292 | 2.49E-23 |
| 23999 | TWF2 | 643.154697 | 0.9432132 | 0.09482897 | 9.94646643 | 2.61E-23 |
| 14964 | H2-D1 | 16322.9757 | -1.5892819 | 0.15979401 | -9.9458165 | 2.63E-23 |
| 24051 | SGCB | 626.877676 | -1.4079681 | 0.14160559 | -9.9428844 | 2.71E-23 |
| 70450 | UNC13D | 770.778478 | -5.0005629 | 0.50331763 | -9.935203 | 2.93E-23 |
| 140792 | COLEC12 | 2198.94239 | 2.15427273 | 0.21689843 | 9.93217301 | 3.02E-23 |
| 74153 | uba7 | 51.1216668 | -3.6472998 | 0.36725366 | -9.9312822 | 3.04E-23 |
| 18034 | NFKB2 | 2852.37535 | -0.7313175 | 0.07367566 | -9.9261748 | 3.20E-23 |
| 71458 | BCOR | 1315.145 | 1.07011723 | 0.10782419 | 9.92464894 | 3.25E-23 |
| 19108 | PRKX | 1640.11513 | -0.9448711 | 0.09521556 | -9.9234945 | 3.29E-23 |
| 243659 | STYK1 | 263.821764 | -2.0512216 | 0.20695807 | -9.911291 | 3.72E-23 |
| 100043172 | Gm10080 | 809.078309 | -0.7163852 | 0.07237619 | -9.8980777 | 4.24E-23 |
| 54373 | PRSS16 | 63.4210485 | -3.2799094 | 0.33144016 | -9.8959321 | 4.34E-23 |
| 29876 | CLIC4 | 9138.01314 | 1.50569499 | 0.15238191 | 9.88106161 | 5.03E-23 |
| 116848 | BAZ2A | 2862.55771 | -0.6302329 | 0.06380375 | -9.8776782 | 5.20E-23 |
| 17246 | MDM2 | 7173.81037 | -1.3194954 | 0.13358989 | -9.87721 | 5.23E-23 |
| 20588 | SMARCC1 | 6008.05492 | 0.57182879 | 0.05791028 | 9.87439194 | 5.38E-23 |
| 96979 | PTGES2 | 797.849025 | -0.7814743 | 0.0791515 | -9.8731459 | 5.44E-23 |
| 17768 | mthfd2 | 3628.90443 | 1.56497879 | 0.15857445 | 9.8690477 | 5.67E-23 |
| 99887 | TMEM56 | 72.3935055 | -3.1080395 | 0.31494662 | -9.8684643 | 5.70E-23 |
| 17283 | MEN1 | 1622.6558 | 0.77446454 | 0.07850936 | 9.86461451 | 5.93E-23 |
| 13710 | ELF3 | 106.60753 | -10.164992 | 1.03194712 | -9.8503031 | 6.83E-23 |
| 170721 | PAPLN | 65.3173568 | -5.6550557 | 0.57414033 | -9.8496053 | 6.88E-23 |
| 211949 | SPSB4 | 87.5791739 | -2.6747091 | 0.27155756 | -9.8495109 | 6.89E-23 |

| | | | | | | |
|---|---|---|---|---|---|---|
| 192197 | BCAS3 | 389.572599 | -1.0026887 | 0.10181753 | -9.8478989 | 7.00E-23 |
| 105439 | SLAIN1 | 173.492291 | -10.869464 | 1.10382913 | -9.8470529 | 7.06E-23 |
| 432530 | ADCY1 | 278.350195 | -10.829208 | 1.09992979 | -9.8453627 | 7.18E-23 |
| 53857 | TUBA8 | 155.988333 | -10.714699 | 1.08936321 | -9.8357453 | 7.90E-23 |
| 12864 | COX6C | 1629.47643 | 0.7131077 | 0.07260519 | 9.82171855 | 9.08E-23 |
| 17112 | TM4SF1 | 14125.6009 | 1.35407006 | 0.13789076 | 9.81987549 | 9.25E-23 |
| 52840 | DBNDD2 | 604.666255 | -0.9841675 | 0.10029144 | -9.8130755 | 9.89E-23 |
| 231440 | PARM1 | 248.39502 | -10.663973 | 1.08688534 | -9.8114977 | 1.00E-22 |
| 71365 | PDSS2 | 478.716211 | 1.00987713 | 0.10292911 | 9.81138504 | 1.01E-22 |
| 68023 | PDF | 588.124052 | 0.97394422 | 0.09927072 | 9.81099165 | 1.01E-22 |
| 232313 | GXYLT2 | 543.478573 | 1.95874549 | 0.19973004 | 9.8069651 | 1.05E-22 |
| 329628 | FAT4 | 4140.0256 | 2.40077676 | 0.24482154 | 9.80623189 | 1.06E-22 |
| 236573 | Gbp9 | 298.313496 | -4.4167307 | 0.45049641 | -9.8041418 | 1.08E-22 |
| 227541 | CAMK1D | 1014.99455 | 2.49702701 | 0.25472566 | 9.80280906 | 1.09E-22 |
| 17002 | LTF | 269.593819 | -10.782272 | 1.10008494 | -9.8013084 | 1.11E-22 |
| 73608 | MARVELD3 | 185.640865 | -10.245241 | 1.04531204 | -9.8011316 | 1.11E-22 |
| 105298 | EPDR1 | 745.335601 | 2.51144841 | 0.25624448 | 9.80098559 | 1.11E-22 |
| 71675 | 0610010F05I | 1372.79289 | -0.8677411 | 0.08856336 | -9.7979704 | 1.15E-22 |
| 105014 | rdh14 | 467.260905 | 0.96168384 | 0.09815507 | 9.79759689 | 1.15E-22 |
| 333433 | GPD1L | 1182.64581 | -0.9833558 | 0.10037674 | -9.7966501 | 1.16E-22 |
| 54651 | USP27X | 321.742024 | -1.6501551 | 0.16851357 | -9.7924163 | 1.21E-22 |
| 239570 | TTC38 | 113.377493 | -5.7979163 | 0.59214896 | -9.7913138 | 1.23E-22 |
| 380863 | TMEM171 | 84.1637098 | -3.9523967 | 0.40372041 | -9.7899354 | 1.24E-22 |
| 26932 | PPP2R5E | 2802.75869 | 0.38663424 | 0.03950333 | 9.78738369 | 1.28E-22 |
| 68299 | VPS53 | 1246.5858 | -0.6548776 | 0.06691922 | -9.786092 | 1.29E-22 |
| 15903 | ID3 | 3420.21732 | 1.49969928 | 0.1532849 | 9.78373787 | 1.32E-22 |
| 16341 | EIF3E | 3535.48035 | 1.18096681 | 0.12072717 | 9.78211258 | 1.34E-22 |
| 17268 | MEIS1 | 134.000701 | 1.89117426 | 0.19336597 | 9.78028491 | 1.37E-22 |
| 70297 | GCC2 | 1441.56147 | -0.5964814 | 0.06099473 | -9.7792285 | 1.38E-22 |
| 327959 | XAF1 | 138.543709 | -3.1211221 | 0.3191772 | -9.7786498 | 1.39E-22 |
| 14365 | FZD3 | 105.704187 | -10.155289 | 1.0399186 | -9.7654656 | 1.58E-22 |
| 235086 | IGSF9B | 101.345935 | -10.093013 | 1.03426479 | -9.7586355 | 1.69E-22 |
| 12465 | CCT5 | 20414.3503 | 0.99215481 | 0.10172278 | 9.75351639 | 1.78E-22 |
| 20568 | SLPI | 4059.94111 | -2.0165846 | 0.20677871 | -9.7523804 | 1.80E-22 |
| 208659 | FAM20A | 76.8748233 | -4.2586504 | 0.43688871 | -9.7476777 | 1.89E-22 |
| 21871 | ATP6V0A2 | 2590.49127 | -0.6683124 | 0.06859055 | -9.7435057 | 1.97E-22 |
| 74747 | DDIT4 | 899.156287 | -1.3687851 | 0.140555 | -9.738431 | 2.07E-22 |
| 269180 | INPP4A | 409.02265 | -1.1137577 | 0.11440829 | -9.7349392 | 2.14E-22 |
| 29810 | BAG3 | 3826.1362 | 1.09192689 | 0.11218748 | 9.73305471 | 2.18E-22 |
| 228608 | SMOX | 1117.43215 | -2.1384734 | 0.21977825 | -9.730141 | 2.24E-22 |
| 101602 | AI467606 | 40.8848522 | 4.14865262 | 0.42647614 | 9.72774842 | 2.30E-22 |
| 12488 | CD2AP | 3150.29264 | -0.53544 | 0.05504723 | -9.7269204 | 2.31E-22 |
| 72042 | COTL1 | 3489.71902 | -3.4867848 | 0.35853465 | -9.7250985 | 2.36E-22 |
| 12029 | BCL6B | 218.744577 | 10.5319814 | 1.08357028 | 9.71970309 | 2.49E-22 |
| 15372 | HMX2 | 74.014245 | 7.24630136 | 0.74569068 | 9.71756992 | 2.54E-22 |

| | | | | | | |
|---|---|---|---|---|---|---|
| 20848 | STAT3 | 3450.5599 | -0.4375842 | 0.04503446 | -9.7166527 | 2.56E-22 |
| 217737 | AHSA1 | 4746.01124 | 0.77338757 | 0.079602 | 9.71567958 | 2.59E-22 |
| 19345 | RAB5C | 4998.90872 | -0.3791973 | 0.03903347 | -9.7146703 | 2.61E-22 |
| 213081 | WDR19 | 304.678023 | -1.1934446 | 0.12289831 | -9.7108299 | 2.71E-22 |
| 18053 | NGFR | 168.611932 | -10.828195 | 1.11594466 | -9.7031646 | 2.92E-22 |
| 319468 | PPM1H | 550.918445 | -2.4472873 | 0.25225946 | -9.7014691 | 2.97E-22 |
| 171212 | GALNT10 | 3006.99926 | -1.1088246 | 0.11430254 | -9.7007871 | 2.99E-22 |
| 320679 | SAMD12 | 44.1749289 | -2.9241358 | 0.30164533 | -9.6939536 | 3.20E-22 |
| 74488 | LRRC15 | 47.8321662 | 4.2487368 | 0.43832614 | 9.69309474 | 3.23E-22 |
| 269589 | SYTL1 | 78.4495162 | -4.3144125 | 0.44515008 | -9.6920403 | 3.26E-22 |
| 17231 | Mcpt8 | 96.5401415 | -10.024256 | 1.03437683 | -9.6911066 | 3.29E-22 |
| 51786 | CPSF2 | 4921.22767 | 0.7348802 | 0.07584602 | 9.68910648 | 3.35E-22 |
| 474332 | DNM3OS | 139.666041 | 10.6065531 | 1.09572171 | 9.67996984 | 3.67E-22 |
| 381463 | nr1h5 | 696.769516 | -1.363242 | 0.14089856 | -9.6753432 | 3.84E-22 |
| 18674 | SLC25A3 | 14608.6555 | -0.3779899 | 0.03907182 | -9.674234 | 3.88E-22 |
| 319713 | ABLIM3 | 210.082953 | -9.1137419 | 0.94220089 | -9.6728224 | 3.93E-22 |
| 20556 | Slfn2 | 290.263329 | -10.168375 | 1.05159609 | -9.6694682 | 4.06E-22 |
| 98402 | SH3BP4 | 2091.43034 | -1.3318285 | 0.13784032 | -9.6621108 | 4.37E-22 |
| 102162 | TAF5L | 1348.66631 | 0.58950434 | 0.06101975 | 9.66087698 | 4.42E-22 |
| 107272 | PSAT1 | 9386.26114 | 0.81445046 | 0.08430721 | 9.66050786 | 4.44E-22 |
| 667977 | Gm8909 | 524.156968 | -1.6095033 | 0.16661073 | -9.6602621 | 4.45E-22 |
| 19946 | RPL30 | 173.991055 | 1.29394533 | 0.13395009 | 9.65990652 | 4.46E-22 |
| 72000 | LMNTD2 | 37.5567884 | -3.9648976 | 0.41067492 | -9.6545892 | 4.70E-22 |
| 17126 | SMAD2 | 2286.91998 | -0.4280825 | 0.04433997 | -9.654552 | 4.70E-22 |
| 78593 | NRIP3 | 48.4663677 | -5.5554881 | 0.57586149 | -9.6472645 | 5.05E-22 |
| 76901 | JADE2 | 2700.14244 | -2.1343046 | 0.2212972 | -9.6445169 | 5.19E-22 |
| 50528 | TMPRSS2 | 92.3981279 | -9.9611711 | 1.03283793 | -9.6444669 | 5.19E-22 |
| 30056 | TIMM9 | 788.840262 | 0.82803862 | 0.08595144 | 9.63379523 | 5.76E-22 |
| 76740 | EFR3A | 7227.09786 | 1.09587937 | 0.1138492 | 9.62570967 | 6.23E-22 |
| 67703 | KIRREL3 | 152.524764 | -9.9592796 | 1.03573054 | -9.6157052 | 6.86E-22 |
| 104776 | ALDH6A1 | 167.233155 | -1.2748726 | 0.13265649 | -9.6103297 | 7.23E-22 |
| 234356 | CSGALNACT1 | 550.248161 | 2.50424563 | 0.26068387 | 9.60644638 | 7.51E-22 |
| 93715 | PCDHGA7 | 110.750539 | -8.1898824 | 0.85272143 | -9.6044054 | 7.66E-22 |
| 278279 | TMTC2 | 99.4074116 | -10.064059 | 1.04843143 | -9.5991582 | 8.06E-22 |
| 22750 | Zfp9 | 522.335841 | 1.47454557 | 0.15364195 | 9.59728518 | 8.21E-22 |
| 54610 | TBC1D8 | 143.66449 | -4.6071485 | 0.48013285 | -9.5955704 | 8.35E-22 |
| 53381 | PRDX4 | 1900.11556 | 0.72780438 | 0.07591105 | 9.58759483 | 9.02E-22 |
| 12539 | CDC37 | 6246.68098 | 0.41592575 | 0.04338713 | 9.58638489 | 9.12E-22 |
| 14786 | GRB7 | 145.306332 | -9.8916317 | 1.03198932 | -9.5850136 | 9.24E-22 |
| 68036 | Zfp706 | 3682.884 | 0.62382271 | 0.06508577 | 9.58462554 | 9.28E-22 |
| 78088 | SOWAHB | 116.370801 | -10.293634 | 1.07435521 | -9.5812199 | 9.59E-22 |
| 71729 | RGS12 | 1349.71204 | -1.7387915 | 0.18153775 | -9.5781263 | 9.88E-22 |
| 20439 | SIAH2 | 695.999908 | 0.90844949 | 0.09485606 | 9.57713759 | 9.98E-22 |
| 224079 | ATP13A4 | 963.599968 | -5.1105393 | 0.53386689 | -9.5726843 | 1.04E-21 |
| 54394 | CRLF3 | 1226.92939 | 0.8821536 | 0.0921695 | 9.57099239 | 1.06E-21 |

| | | | | | | |
|---|---|---|---|---|---|---|
| 67573 | LOXL4 | 2252.21718 | -1.3838567 | 0.14460757 | -9.5697389 | 1.07E-21 |
| 74012 | RAP2B | 1809.67461 | -0.6571651 | 0.06870892 | -9.5644808 | 1.13E-21 |
| 66912 | BZW2 | 3724.49274 | 1.58205 | 0.16541974 | 9.5638525 | 1.13E-21 |
| 18559 | PCTP | 137.361206 | -1.4653765 | 0.15332245 | -9.557482 | 1.21E-21 |
| 70348 | Ube2cbp | 233.11511 | 1.07538788 | 0.11257136 | 9.55294368 | 1.26E-21 |
| 68943 | PINK1 | 431.24583 | -0.8580658 | 0.08984379 | -9.5506415 | 1.29E-21 |
| 223435 | TRIO | 7542.65674 | 0.84603931 | 0.08859497 | 9.54951803 | 1.30E-21 |
| 68020 | APOPT1 | 520.522795 | 0.72627125 | 0.07607605 | 9.54664813 | 1.34E-21 |
| 16443 | ITSN1 | 2244.17854 | 0.88510834 | 0.09273651 | 9.54433551 | 1.37E-21 |
| 69550 | BST2 | 1375.17497 | -3.1871092 | 0.33393931 | -9.5439775 | 1.37E-21 |
| 57295 | ICMT | 2766.80325 | 0.62146736 | 0.06512018 | 9.54339149 | 1.38E-21 |
| 19302 | PEX2 | 1159.70306 | -0.5238423 | 0.05504026 | -9.5174396 | 1.77E-21 |
| 98682 | MFSD6 | 2846.92694 | -2.792302 | 0.29352149 | -9.5131095 | 1.85E-21 |
| 21803 | TGFB1 | 1663.02477 | 0.90555196 | 0.09519535 | 9.51256469 | 1.86E-21 |
| 68616 | GDPD3 | 62.031409 | -3.7989132 | 0.3994709 | -9.509862 | 1.91E-21 |
| 67039 | RBM25 | 5111.16795 | 0.61765586 | 0.0649645 | 9.50758981 | 1.95E-21 |
| 20911 | STXBP2 | 1701.77914 | -1.1421253 | 0.12013546 | -9.5069789 | 1.96E-21 |
| 83813 | TNK1 | 198.363316 | -9.6205414 | 1.01196719 | -9.5067721 | 1.97E-21 |
| 327655 | PPIP5K1 | 701.935806 | -0.7502196 | 0.07893101 | -9.5047509 | 2.01E-21 |
| 66196 | MYO19 | 1286.80693 | 0.79084058 | 0.08321156 | 9.50397466 | 2.02E-21 |
| 239833 | LMLN | 231.133512 | -1.4961138 | 0.15746746 | -9.5010985 | 2.08E-21 |
| 17117 | AMACR | 307.314386 | 1.71261137 | 0.1802593 | 9.50082102 | 2.08E-21 |
| 20425 | SHMT1 | 1369.83282 | 0.48069764 | 0.05061074 | 9.49793761 | 2.14E-21 |
| 242100 | PGLYRP3 | 82.9009882 | -9.8039325 | 1.03233313 | -9.496869 | 2.16E-21 |
| 70892 | TTLL7 | 1623.29779 | -1.0321024 | 0.10873813 | -9.4916325 | 2.27E-21 |
| 14963 | H2-Bl | 211.76839 | -1.7127572 | 0.1805246 | -9.4876667 | 2.36E-21 |
| 19283 | PTPRZ1 | 94.8283122 | -9.9982672 | 1.05385315 | -9.4873439 | 2.37E-21 |
| 224912 | CRB3 | 148.917493 | -10.649117 | 1.12369512 | -9.4768737 | 2.62E-21 |
| 56531 | YLPM1 | 3275.88895 | 0.45735437 | 0.04828042 | 9.4728748 | 2.72E-21 |
| 16542 | KDR | 316.686193 | -8.1127258 | 0.85646289 | -9.4723611 | 2.74E-21 |
| 212999 | TNPO2 | 6057.97747 | -0.626898 | 0.06621291 | -9.4679124 | 2.85E-21 |
| 68477 | RMND5A | 1805.9149 | 0.58220336 | 0.06151961 | 9.46370304 | 2.97E-21 |
| 16590 | KIT | 114.243459 | -10.265033 | 1.08521229 | -9.4590087 | 3.11E-21 |
| 12614 | CELSR1 | 251.398041 | -6.6261161 | 0.70095044 | -9.4530451 | 3.29E-21 |
| 20847 | STAT2 | 523.143194 | -1.172775 | 0.12406655 | -9.4527892 | 3.30E-21 |
| 68097 | DYNLL2 | 4737.08902 | -0.8769789 | 0.09279123 | -9.451097 | 3.35E-21 |
| 14782 | GSR | 4542.85359 | 0.92050671 | 0.09747615 | 9.44340424 | 3.61E-21 |
| 22678 | ZFP2 | 81.2775804 | -9.7742642 | 1.03605349 | -9.4341309 | 3.94E-21 |
| 72050 | KDELC1 | 1184.69012 | 0.89660737 | 0.09505125 | 9.43288358 | 3.99E-21 |
| 23794 | ADAMTS5 | 245.849387 | 3.63100403 | 0.38497692 | 9.43174461 | 4.03E-21 |
| 230726 | RHBDL2 | 97.1724721 | -8.0043833 | 0.84909322 | -9.4269783 | 4.22E-21 |
| 14366 | FZD4 | 110.369891 | 2.45550608 | 0.26051084 | 9.42573486 | 4.27E-21 |
| 218121 | MBOAT1 | 879.283624 | -2.4294677 | 0.25779082 | -9.4241822 | 4.33E-21 |
| 53618 | FUT8 | 1428.11814 | 1.13264969 | 0.12024566 | 9.41946408 | 4.53E-21 |
| 75826 | SENP2 | 2328.81743 | -0.416633 | 0.04426062 | -9.4131754 | 4.81E-21 |

| | | | | | | |
|---|---|---|---|---|---|---|
| 27418 | MKLN1 | 1588.89689 | -0.5261225 | 0.05589528 | -9.4126474 | 4.84E-21 |
| 100503499 | Gm16136 | 86.9809278 | -9.8709651 | 1.04909899 | -9.4089931 | 5.01E-21 |
| 76527 | IL34 | 1048.22121 | -1.3591078 | 0.1444686 | -9.4076343 | 5.07E-21 |
| 13345 | TWIST2 | 1278.97836 | 4.85378524 | 0.51601348 | 9.40631476 | 5.14E-21 |
| 14873 | GSTO1 | 11134.889 | 2.08863812 | 0.2220964 | 9.40419607 | 5.24E-21 |
| 67728 | DPH2 | 662.388494 | 1.01528151 | 0.10799133 | 9.40150926 | 5.38E-21 |
| 15529 | SDC2 | 5885.39919 | 0.95811774 | 0.10192978 | 9.39978249 | 5.47E-21 |
| 106957 | SLC39A6 | 5777.66662 | 0.99749446 | 0.10612619 | 9.39913608 | 5.50E-21 |
| 53611 | VTI1A | 911.162657 | -0.9317659 | 0.0991455 | -9.3979655 | 5.56E-21 |
| 26879 | B3GALNT1 | 208.969889 | 9.7491503 | 1.03749808 | 9.39678878 | 5.63E-21 |
| 20259 | SCIN | 143.179722 | -1.5309043 | 0.16293604 | -9.395738 | 5.68E-21 |
| 20661 | SORT1 | 431.26416 | -4.5592139 | 0.48541757 | -9.3923546 | 5.87E-21 |
| 21789 | TFPI2 | 139.121973 | 9.8784513 | 1.05184063 | 9.39158557 | 5.91E-21 |
| 60533 | CD274 | 75.099447 | -4.0670509 | 0.4334451 | -9.3830819 | 6.41E-21 |
| 212712 | SATB2 | 1226.39439 | 1.96348266 | 0.20931647 | 9.3804499 | 6.57E-21 |
| 194744 | SLC25A43 | 134.044336 | -1.4169917 | 0.15108797 | -9.3785871 | 6.69E-21 |
| 72077 | GCNT3 | 207.614497 | -8.6637373 | 0.92412004 | -9.3751211 | 6.91E-21 |
| 231327 | PPAT | 2432.19603 | 0.68191087 | 0.07274582 | 9.37388413 | 6.99E-21 |
| 67286 | IFT22 | 326.930929 | -1.2102019 | 0.12912456 | -9.3723605 | 7.09E-21 |
| 68817 | DDI2 | 738.094186 | 1.10078331 | 0.11755732 | 9.36380098 | 7.69E-21 |
| 269400 | RTEL1 | 1980.5727 | 0.46825358 | 0.05002185 | 9.36098087 | 7.90E-21 |
| 11980 | ATP8A1 | 144.248686 | -10.601926 | 1.13305707 | -9.3569211 | 8.21E-21 |
| 72278 | CCPG1 | 530.602515 | -0.9142907 | 0.09776466 | -9.3519547 | 8.60E-21 |
| 236266 | ALMS1 | 1050.66734 | 0.79818991 | 0.08541937 | 9.34436651 | 9.24E-21 |
| 16658 | MAFB | 105.490418 | -10.149997 | 1.08656156 | -9.3413912 | 9.51E-21 |
| 18109 | MYCN | 45.2562409 | 5.32931393 | 0.57051663 | 9.34120693 | 9.52E-21 |
| 19725 | RFX2 | 109.46288 | -1.8350897 | 0.19661624 | -9.3333579 | 1.03E-20 |
| 19159 | CYTH3 | 1629.12847 | 1.44668243 | 0.15505699 | 9.33000457 | 1.06E-20 |
| 19358 | RAD23A | 1948.71418 | -0.4803466 | 0.05149177 | -9.3286085 | 1.07E-20 |
| 218215 | RNF144B | 149.63741 | -6.2157971 | 0.66634429 | -9.3282064 | 1.08E-20 |
| 16330 | INPP5B | 877.469167 | 0.84065628 | 0.09014868 | 9.32522018 | 1.11E-20 |
| 20775 | SQLE | 10393.0069 | 0.73306575 | 0.07861672 | 9.32455325 | 1.11E-20 |
| 209497 | TMEM164 | 1612.80517 | 0.72778379 | 0.07812553 | 9.31556967 | 1.21E-20 |
| 103573 | XPO1 | 10916.003 | 0.43702374 | 0.04692164 | 9.31390507 | 1.23E-20 |
| 14236 | FOXN2 | 1572.70521 | 0.92543299 | 0.09937298 | 9.31272257 | 1.25E-20 |
| 99326 | GARNL3 | 156.664391 | -2.9748729 | 0.31954492 | -9.3097173 | 1.28E-20 |
| 16331 | INPP5D | 49.0013838 | -2.6758148 | 0.28745989 | -9.3084807 | 1.30E-20 |
| 70762 | DCLK2 | 164.431333 | 2.16007129 | 0.23206903 | 9.30788264 | 1.30E-20 |
| 73062 | PPP1R16A | 747.14155 | -0.6230787 | 0.0669455 | -9.3072516 | 1.31E-20 |
| 71207 | NUDT4 | 6029.18357 | -1.0311928 | 0.11080603 | -9.3062879 | 1.32E-20 |
| 58801 | PMAIP1 | 297.751175 | -5.9454833 | 0.63914508 | -9.3022438 | 1.38E-20 |
| 23918 | IMPDH2 | 435.58062 | 0.97564789 | 0.10489003 | 9.30162636 | 1.38E-20 |
| 27375 | TJP3 | 101.131482 | -10.0913 | 1.08525302 | -9.2985693 | 1.42E-20 |
| 68196 | HSBP1 | 1805.91242 | -0.5345929 | 0.05749909 | -9.2974156 | 1.44E-20 |
| 18176 | NRAS | 5118.61173 | -0.4485956 | 0.04826499 | -9.2944317 | 1.48E-20 |

| | | | | | | |
|---|---|---|---|---|---|---|
| 18738 | pitpna | 6741.96168 | -0.5458682 | 0.05873159 | -9.2942856 | 1.48E-20 |
| 74413 | TC2N | 296.414669 | -10.205013 | 1.09800576 | -9.2941338 | 1.48E-20 |
| 54004 | DIAPH2 | 1153.35244 | -0.8942823 | 0.09624077 | -9.2921354 | 1.51E-20 |
| 76967 | 2700049A03 | 896.422703 | 0.7790193 | 0.08383752 | 9.29201273 | 1.51E-20 |
| 76044 | NCAPG2 | 4869.49309 | 0.60094267 | 0.06467734 | 9.29139383 | 1.52E-20 |
| 215819 | NHSL1 | 3943.26265 | -1.9897841 | 0.21415816 | -9.2911896 | 1.53E-20 |
| 15476 | HS3ST1 | 99.316953 | -10.063212 | 1.08359501 | -9.2868758 | 1.59E-20 |
| 70155 | OGFRL1 | 569.511635 | 0.86290063 | 0.09294913 | 9.28357946 | 1.64E-20 |
| 14049 | EYA2 | 72.0281212 | -9.601484 | 1.03442923 | -9.2819148 | 1.66E-20 |
| 67894 | FAM45A | 834.786489 | -0.6738959 | 0.0726271 | -9.2788492 | 1.71E-20 |
| 111241 | Hmga1-rs1 | 6400.77692 | 1.34910402 | 0.14544381 | 9.27577487 | 1.76E-20 |
| 237769 | Gm12258 | 78.4617647 | -1.9732846 | 0.2128677 | -9.2700049 | 1.86E-20 |
| 211064 | ALKBH1 | 509.389444 | 0.75595071 | 0.08155496 | 9.26921853 | 1.88E-20 |
| 67916 | PLPP3 | 1884.23886 | 3.13702674 | 0.33844322 | 9.26898982 | 1.88E-20 |
| 114128 | LAPTM4B | 2444.03161 | 0.61758902 | 0.06666495 | 9.2640738 | 1.97E-20 |
| 24088 | TLR2 | 1147.00189 | -1.7310065 | 0.18696758 | -9.258324 | 2.08E-20 |
| 19229 | PTK2B | 581.179819 | -5.7693489 | 0.62344142 | -9.2540353 | 2.16E-20 |
| 16392 | ISL1 | 74.3293732 | 9.69544488 | 1.04789509 | 9.25230494 | 2.20E-20 |
| 78943 | ERN1 | 536.089436 | -1.2321514 | 0.13323428 | -9.248006 | 2.29E-20 |
| 70355 | GPRC5C | 252.631885 | -9.4018247 | 1.01682798 | -9.2462294 | 2.33E-20 |
| 107986 | ddb2 | 182.638366 | -3.1949443 | 0.34565591 | -9.243135 | 2.39E-20 |
| 73720 | CST6 | 619.803038 | -4.3087135 | 0.46641639 | -9.237912 | 2.51E-20 |
| 13848 | EPHB6 | 819.965623 | -2.1437492 | 0.23213431 | -9.234952 | 2.58E-20 |
| 231003 | KLHL17 | 307.61892 | 0.88350656 | 0.09568022 | 9.23395159 | 2.61E-20 |
| 75939 | 4930579G24 | 838.820092 | 0.73169433 | 0.07924496 | 9.23332363 | 2.62E-20 |
| 53860 | 41890 | 6974.8427 | -0.6132706 | 0.06645935 | -9.2277543 | 2.76E-20 |
| 71709 | SYDE1 | 1644.3781 | 0.96563574 | 0.10466064 | 9.22635029 | 2.80E-20 |
| 233979 | TPCN2 | 393.353451 | 0.77037205 | 0.08351489 | 9.22436774 | 2.85E-20 |
| 230770 | TMEM39B | 307.485797 | 1.56110446 | 0.16929531 | 9.2211914 | 2.94E-20 |
| 320772 | MDGA2 | 80.2485337 | -9.7580901 | 1.05837953 | -9.2198402 | 2.98E-20 |
| 58235 | NECTIN1 | 3327.44257 | -2.2455884 | 0.24356707 | -9.2195896 | 2.98E-20 |
| 11993 | AUP1 | 3406.07667 | 0.46040833 | 0.04996616 | 9.21440353 | 3.13E-20 |
| 20893 | BHLHE40 | 3207.49516 | -1.3646925 | 0.14810739 | -9.2142092 | 3.14E-20 |
| 67465 | SF3A1 | 4688.52967 | 0.40423898 | 0.04390396 | 9.20734702 | 3.34E-20 |
| 67306 | ZC2HC1A | 1138.14062 | -1.1417407 | 0.12407134 | -9.2022923 | 3.50E-20 |
| 14696 | GNB4 | 463.153926 | 2.83804803 | 0.30841904 | 9.20192219 | 3.52E-20 |
| 75415 | ARHGAP12 | 1409.7389 | -1.0156162 | 0.11044463 | -9.1957046 | 3.73E-20 |
| 93706 | PCDHGC3 | 6361.9055 | 1.59218773 | 0.17315188 | 9.19532473 | 3.74E-20 |
| 545551 | BC021767 | 55.8566627 | -6.4491379 | 0.70155206 | -9.1926719 | 3.83E-20 |
| 76055 | MGEA5 | 4628.56715 | -0.649249 | 0.07064825 | -9.1898818 | 3.93E-20 |
| 16774 | LAMA3 | 331.781963 | -4.7348676 | 0.51526931 | -9.1891124 | 3.96E-20 |
| 79202 | Tnfrsf22 | 899.630066 | -0.6308736 | 0.06865838 | -9.1885893 | 3.98E-20 |
| 381290 | ATP2B4 | 354.340625 | -4.3620802 | 0.47479313 | -9.1873279 | 4.03E-20 |
| 93734 | MPV17L | 120.520398 | -2.2079052 | 0.2403859 | -9.1848368 | 4.12E-20 |
| 70294 | rnf126 | 2653.78511 | 0.436309 | 0.04750454 | 9.18457389 | 4.13E-20 |

| | | | | | | |
|---|---|---|---|---|---|---|
| 75717 | CUL5 | 3024.1812 | 0.69144966 | 0.07528438 | 9.18450404 | 4.13E-20 |
| 18476 | PAFAH1B3 | 444.638629 | -1.0405167 | 0.11329571 | -9.1840779 | 4.15E-20 |
| 107723 | SLC12A6 | 1311.10156 | -0.6568955 | 0.07156033 | -9.1796042 | 4.33E-20 |
| 14154 | FEM1A | 2037.4586 | 0.73806643 | 0.08046553 | 9.17245511 | 4.62E-20 |
| 17756 | MAP2 | 1321.51889 | -4.4646509 | 0.48682546 | -9.1709477 | 4.69E-20 |
| 80750 | N4BP1 | 2326.60329 | -0.9075799 | 0.09897409 | -9.1698744 | 4.74E-20 |
| 11475 | ACTA2 | 503.482433 | 4.93452072 | 0.53823092 | 9.16803646 | 4.82E-20 |
| 20909 | Stx4a | 1341.58386 | 0.63706293 | 0.06952941 | 9.16249651 | 5.07E-20 |
| 30806 | ADAMTS8 | 82.0097786 | -7.3418739 | 0.80169839 | -9.1579003 | 5.29E-20 |
| 98758 | HNRNPF | 1877.33316 | 0.58499935 | 0.06402095 | 9.13762411 | 6.38E-20 |
| 73723 | SH3BGRL3 | 1343.28955 | 0.94823164 | 0.10381206 | 9.13411848 | 6.59E-20 |
| 217166 | NR1D1 | 762.727058 | 1.4919882 | 0.16337231 | 9.1324422 | 6.70E-20 |
| 18771 | PKNOX1 | 714.334323 | 0.77752209 | 0.08514172 | 9.13209315 | 6.72E-20 |
| 14629 | GCLC | 4846.37142 | 1.16159059 | 0.12732933 | 9.12272629 | 7.33E-20 |
| 11928 | ATP1A1 | 29955.0743 | -0.5572531 | 0.06109565 | -9.1209942 | 7.44E-20 |
| 78920 | DLST | 6530.78475 | 0.69429654 | 0.07613877 | 9.11883085 | 7.59E-20 |
| 214763 | MB21D1 | 650.988493 | 0.86070873 | 0.09444197 | 9.11362519 | 7.97E-20 |
| 75216 | CEP128 | 1437.30294 | 0.63475668 | 0.06965491 | 9.1128779 | 8.02E-20 |
| 70691 | 3830403N18 | 99.7949175 | 10.1214637 | 1.11221054 | 9.1003127 | 9.01E-20 |
| 225659 | CEP76 | 1159.58802 | 0.77388645 | 0.08508466 | 9.09548712 | 9.42E-20 |
| 72401 | SLC43A1 | 70.9538241 | 6.56997389 | 0.72260426 | 9.09207744 | 9.72E-20 |
| 320528 | VPS13C | 2635.4308 | 1.1605795 | 0.12765562 | 9.09148792 | 9.77E-20 |
| 66686 | DCBLD1 | 517.65673 | -0.7809531 | 0.08590435 | -9.0909615 | 9.82E-20 |
| 433182 | eno1b | 1251.23345 | 0.89811389 | 0.09880191 | 9.09004547 | 9.90E-20 |
| 13405 | DMD | 242.144402 | -11.349537 | 1.25041871 | -9.076589 | 1.12E-19 |
| 17873 | GADD45B | 1180.65165 | -1.4242186 | 0.15695658 | -9.0739656 | 1.15E-19 |
| 319880 | TMCC3 | 2254.68964 | -1.1351501 | 0.12517147 | -9.0687606 | 1.20E-19 |
| 68659 | FAM198B | 1470.91244 | 3.5867843 | 0.39555579 | 9.06770771 | 1.22E-19 |
| 17702 | MSX2 | 63.1130797 | -9.408501 | 1.03761675 | -9.0674143 | 1.22E-19 |
| 30944 | Zfp354c | 156.213415 | 1.56750858 | 0.17297524 | 9.06203998 | 1.28E-19 |
| 18755 | PRKCH | 61.1129297 | -3.5581074 | 0.3927049 | -9.0605117 | 1.30E-19 |
| 69101 | YDJC | 166.551073 | 2.88278168 | 0.31859487 | 9.04842459 | 1.45E-19 |
| 76441 | DAAM2 | 110.756393 | -6.3062021 | 0.69708343 | -9.0465528 | 1.48E-19 |
| 73469 | RNF38 | 1288.60952 | -0.9497704 | 0.10499843 | -9.045568 | 1.49E-19 |
| 74185 | GBE1 | 3180.03994 | 0.92794663 | 0.10259669 | 9.04460618 | 1.50E-19 |
| 67374 | JAM2 | 1850.89164 | 2.01165318 | 0.22244782 | 9.04325851 | 1.52E-19 |
| 20479 | VPS4B | 2435.79261 | -0.4781972 | 0.05288737 | -9.0418032 | 1.54E-19 |
| 110877 | SLC18A1 | 61.115979 | -4.6274179 | 0.5122376 | -9.0337333 | 1.66E-19 |
| 23805 | APC2 | 173.62405 | -3.9839415 | 0.44117073 | -9.0303849 | 1.71E-19 |
| 228839 | TGIF2 | 80.0683742 | 3.1321565 | 0.34704631 | 9.02518321 | 1.79E-19 |
| 56454 | ALDH18A1 | 3076.98983 | 0.78070096 | 0.08652475 | 9.02286258 | 1.83E-19 |
| 208994 | FAM83B | 62.9644503 | -9.405089 | 1.04312229 | -9.016286 | 1.95E-19 |
| 66166 | S100A14 | 109.66313 | -9.4829569 | 1.05239343 | -9.0108477 | 2.04E-19 |
| 14567 | GDI1 | 1509.78016 | -0.7817192 | 0.08675349 | -9.0108106 | 2.05E-19 |
| 67484 | EEPD1 | 97.1365576 | -4.5257436 | 0.5023485 | -9.0091711 | 2.08E-19 |

| | | | | | | |
|---|---|---|---|---|---|---|
| 12530 | CDC25A | 1367.89394 | 0.73882347 | 0.08202928 | 9.00682624 | 2.12E-19 |
| 20877 | AURKB | 2449.24761 | 0.4582678 | 0.05090924 | 9.00166261 | 2.22E-19 |
| 12408 | CBR1 | 353.673745 | -1.2722518 | 0.14138361 | -8.9985809 | 2.29E-19 |
| 12054 | BCL7B | 757.918637 | 0.6685301 | 0.07429486 | 8.99833606 | 2.29E-19 |
| 83925 | TRPS1 | 4615.65334 | 0.917639 | 0.10199262 | 8.99711163 | 2.32E-19 |
| 23961 | Oas1b | 51.0042506 | -3.1391138 | 0.34938249 | -8.9847484 | 2.59E-19 |
| 217684 | SUSD6 | 4407.91588 | -0.4268995 | 0.04757755 | -8.9727091 | 2.89E-19 |
| 13033 | CTSD | 30798.514 | -1.6758056 | 0.18684074 | -8.9691661 | 2.99E-19 |
| 208111 | Zfp976 | 84.7267979 | -4.174652 | 0.46545762 | -8.9689197 | 2.99E-19 |
| 68169 | NDNF | 71.4188849 | -9.5869693 | 1.06932189 | -8.9654662 | 3.09E-19 |
| 66713 | ACTR2 | 11406.3192 | -0.4791341 | 0.0534702 | -8.9607681 | 3.22E-19 |
| 108013 | CELF4 | 121.684973 | -3.5225348 | 0.3933905 | -8.9542957 | 3.42E-19 |
| 380714 | RPH3AL | 46.8789762 | -5.7135139 | 0.63807562 | -8.9542896 | 3.42E-19 |
| 243084 | TMPRSS11E | 177.790989 | -8.8727332 | 0.9914554 | -8.9492005 | 3.58E-19 |
| 14229 | FKBP5 | 3963.53536 | -1.8374491 | 0.20538674 | -8.9462891 | 3.68E-19 |
| 15361 | HMGA1 | 1039.57822 | 1.64368396 | 0.18378028 | 8.94374502 | 3.76E-19 |
| 68789 | trmt61b | 227.397878 | -1.2749679 | 0.14255672 | -8.9435834 | 3.77E-19 |
| 68262 | AGPAT4 | 375.442324 | 1.42496262 | 0.15934487 | 8.94263239 | 3.80E-19 |
| 66403 | ASF1A | 837.667565 | 0.7551275 | 0.0844938 | 8.93707606 | 4.00E-19 |
| 319262 | FCHSD1 | 603.300715 | -2.2279684 | 0.24939454 | -8.9335094 | 4.13E-19 |
| 109108 | SLC30A9 | 2282.50663 | -0.3893225 | 0.04359723 | -8.9299837 | 4.26E-19 |
| 234729 | VAC14 | 1718.76502 | -0.74468 | 0.08342275 | -8.9265819 | 4.39E-19 |
| 93888 | Pcdhb17 | 146.138631 | 2.08696072 | 0.23383926 | 8.92476604 | 4.47E-19 |
| 223918 | spryd3 | 1224.01527 | -1.169879 | 0.13116443 | -8.9191788 | 4.70E-19 |
| 235044 | PLPPR2 | 201.440704 | 2.38266108 | 0.2671721 | 8.91807599 | 4.74E-19 |
| 71198 | OTUD1 | 602.793175 | 1.35193655 | 0.15160247 | 8.91764209 | 4.76E-19 |
| 27047 | OMD | 78.0359986 | 7.01665299 | 0.78684465 | 8.91745653 | 4.77E-19 |
| 329650 | MED12L | 135.834246 | 1.5520495 | 0.17411323 | 8.9140238 | 4.92E-19 |
| 330671 | b4galnt4 | 81.5390474 | 3.7952189 | 0.42575926 | 8.91400208 | 4.92E-19 |
| 75572 | ACYP2 | 264.555534 | 1.43942759 | 0.16150057 | 8.91283271 | 4.97E-19 |
| 29819 | STAU2 | 460.221227 | -1.9809966 | 0.22230573 | -8.9111361 | 5.05E-19 |
| 19155 | NPEPPS | 3967.37374 | -0.5731034 | 0.06432794 | -8.9090904 | 5.15E-19 |
| 12825 | COL3A1 | 1085.00627 | 12.8427495 | 1.44165817 | 8.90831803 | 5.18E-19 |
| 12861 | Cox6a1 | 2836.58531 | -0.6671895 | 0.07495304 | -8.9014335 | 5.51E-19 |
| 66218 | NDUFB9 | 2286.28314 | 0.48058721 | 0.05400458 | 8.8990088 | 5.63E-19 |
| 68799 | RGMB | 803.543973 | 1.96321059 | 0.22078608 | 8.89191293 | 6.01E-19 |
| 17391 | MMP24 | 248.73175 | -9.3561086 | 1.05222864 | -8.8917068 | 6.02E-19 |
| 74718 | SNX16 | 380.838576 | -1.0852666 | 0.12209124 | -8.8889799 | 6.17E-19 |
| 230848 | ZBTB40 | 734.410007 | 0.8657912 | 0.09741104 | 8.88801971 | 6.22E-19 |
| 13639 | EFNA4 | 180.590777 | -1.1740001 | 0.13211531 | -8.886178 | 6.32E-19 |
| 320213 | SENP5 | 3652.52911 | -0.5089137 | 0.05733976 | -8.8754059 | 6.97E-19 |
| 208266 | DOT1L | 2765.38676 | 0.53072344 | 0.05981757 | 8.87236678 | 7.16E-19 |
| 74147 | ehhadh | 80.3561995 | -6.4924859 | 0.73198573 | -8.8696891 | 7.34E-19 |
| 22441 | Xlr | 130.168007 | 8.0446277 | 0.9070663 | 8.86884199 | 7.39E-19 |
| 242125 | MAB21L3 | 41.1493527 | -3.9344737 | 0.44377597 | -8.8659007 | 7.59E-19 |

| | | | | | | |
|---|---|---|---|---|---|---|
| 12795 | PLK3 | 1189.3051 | -0.9606695 | 0.10836738 | -8.8649323 | 7.66E-19 |
| 328329 | MAST4 | 4538.13317 | 0.89919021 | 0.10143598 | 8.86460782 | 7.68E-19 |
| 271813 | AGBL2 | 85.6644591 | -6.2130772 | 0.7015436 | -8.8562951 | 8.27E-19 |
| 434341 | NLRC5 | 88.0754135 | -2.3096421 | 0.26079257 | -8.8562421 | 8.28E-19 |
| 19362 | RAD51AP1 | 783.148697 | 0.78807981 | 0.08900018 | 8.85481153 | 8.38E-19 |
| 228421 | KIF18A | 955.281159 | 0.67671008 | 0.0764391 | 8.85293129 | 8.53E-19 |
| 64685 | NMI | 177.627628 | -1.349134 | 0.15250442 | -8.846524 | 9.03E-19 |
| 77531 | ANKS1B | 42.1586137 | 3.82729028 | 0.43278104 | 8.84347953 | 9.28E-19 |
| 384309 | TRIM56 | 1015.62821 | -0.6500819 | 0.0735136 | -8.843015 | 9.32E-19 |
| 71722 | CIC | 5763.30243 | 0.90972133 | 0.10298209 | 8.83378205 | 1.01E-18 |
| 13627 | EEF1A1 | 45810.2988 | 0.22278981 | 0.02522563 | 8.83188168 | 1.03E-18 |
| 623781 | Gm14137 | 369.682983 | 1.29670061 | 0.14682146 | 8.8318191 | 1.03E-18 |
| 17125 | SMAD1 | 1126.14423 | 0.60228613 | 0.06822202 | 8.82832507 | 1.06E-18 |
| 21413 | TCF4 | 2517.57652 | 1.45611115 | 0.16495659 | 8.82723847 | 1.07E-18 |
| 622434 | ARHGEF26 | 127.689131 | -2.4782153 | 0.28098082 | -8.8198737 | 1.15E-18 |
| 12521 | CD82 | 1332.47445 | -2.2650446 | 0.25694837 | -8.8151743 | 1.19E-18 |
| 329727 | DENND2C | 429.487928 | -1.6589778 | 0.18829254 | -8.81064 | 1.24E-18 |
| 319586 | CELF5 | 133.672121 | -4.9248788 | 0.55902485 | -8.8097672 | 1.25E-18 |
| 241556 | TSPAN18 | 121.73002 | -8.9175429 | 1.01241908 | -8.8081538 | 1.27E-18 |
| 69090 | ASCC1 | 446.325828 | -0.9216245 | 0.10465728 | -8.8061193 | 1.30E-18 |
| 16201 | ILF3 | 4172.12697 | 0.64558547 | 0.07334587 | 8.80193378 | 1.34E-18 |
| 65960 | TWSG1 | 3963.95043 | -0.5173206 | 0.0587836 | -8.800424 | 1.36E-18 |
| 100503318 | Gm15853 | 68.1912523 | 2.2726975 | 0.25841621 | 8.79471719 | 1.43E-18 |
| 27660 | 1700088E04I | 54.5572509 | -2.123457 | 0.24166649 | -8.786725 | 1.54E-18 |
| 22004 | TPM2 | 1501.55837 | 4.22022534 | 0.48038628 | 8.78506637 | 1.56E-18 |
| 19889 | RP2 | 1060.60059 | -0.6288793 | 0.07159499 | -8.7838455 | 1.58E-18 |
| 13527 | DTNA | 383.512954 | -2.4682863 | 0.28100932 | -8.7836457 | 1.58E-18 |
| 320939 | 5930403L14I | 54.0396163 | 9.2370431 | 1.05201685 | 8.78031859 | 1.63E-18 |
| 240283 | DMXL1 | 6343.81976 | -2.5877505 | 0.29477391 | -8.7787639 | 1.65E-18 |
| 237988 | CDR2L | 1269.93279 | 2.55959253 | 0.29159605 | 8.77787104 | 1.67E-18 |
| 16880 | LIFR | 298.630473 | 4.03209363 | 0.45945807 | 8.77575971 | 1.70E-18 |
| 74032 | SDR42E1 | 134.924643 | -1.8276551 | 0.20827847 | -8.7750549 | 1.71E-18 |
| 231093 | AGBL5 | 177.943916 | -1.1315436 | 0.12900311 | -8.7714444 | 1.76E-18 |
| 71853 | PDIA6 | 10648.3513 | 0.85771619 | 0.09781921 | 8.76838188 | 1.81E-18 |
| 12419 | CBX5 | 9148.82463 | -0.5030974 | 0.05738273 | -8.7674021 | 1.83E-18 |
| 107587 | osr2 | 50.0854193 | 9.12732207 | 1.04111232 | 8.76689472 | 1.84E-18 |
| 69675 | PXDN | 4472.19671 | 1.04639469 | 0.11942847 | 8.76168552 | 1.92E-18 |
| 223227 | SOX21 | 41.7184045 | -5.7615789 | 0.65771177 | -8.7600361 | 1.95E-18 |
| 65962 | slc9a3r2 | 264.999644 | -1.1481061 | 0.13108147 | -8.7587214 | 1.97E-18 |
| 217578 | BAZ1A | 3320.22795 | 0.68404571 | 0.0781122 | 8.7572203 | 2.00E-18 |
| 223753 | CERK | 608.424024 | 0.61130433 | 0.06981678 | 8.7558367 | 2.03E-18 |
| 244871 | ZC3H12C | 992.033264 | -0.8670679 | 0.09905384 | -8.7535013 | 2.07E-18 |
| 94089 | TRIM7 | 110.954615 | -2.2681544 | 0.25935585 | -8.7453372 | 2.22E-18 |
| 72027 | SLC39A4 | 182.793778 | -4.1199824 | 0.47130564 | -8.7416361 | 2.30E-18 |
| 21826 | THBS2 | 1953.6596 | 7.33777648 | 0.8395037 | 8.74061244 | 2.32E-18 |

| | | | | | | |
|---|---|---|---|---|---|---|
| 28135 | CEP63 | 542.315279 | 0.73098654 | 0.08368736 | 8.7347304 | 2.44E-18 |
| 233552 | GDPD5 | 638.294141 | -3.5503469 | 0.40696275 | -8.7240096 | 2.69E-18 |
| 11800 | API5 | 4391.6884 | 0.36939109 | 0.04235569 | 8.72116772 | 2.75E-18 |
| 77590 | CHST15 | 122.069045 | 6.94346185 | 0.79619023 | 8.72085789 | 2.76E-18 |
| 338320 | mia2 | 105.547854 | -1.9213803 | 0.22044578 | -8.7158858 | 2.88E-18 |
| 27206 | NRK | 57.3610641 | -9.2698721 | 1.06384317 | -8.7135702 | 2.94E-18 |
| 15013 | H2-Q2 | 519.98305 | -1.6724043 | 0.19195524 | -8.7124703 | 2.97E-18 |
| 230696 | AU022252 | 294.90434 | 1.10824866 | 0.12724572 | 8.70951655 | 3.05E-18 |
| 217558 | G2E3 | 2430.26988 | 0.7119263 | 0.08175364 | 8.70819126 | 3.09E-18 |
| 66175 | mustn1 | 67.0377998 | 5.07340298 | 0.58324458 | 8.69858567 | 3.36E-18 |
| 194126 | MTMR11 | 694.127257 | 2.05498152 | 0.23629954 | 8.69651095 | 3.42E-18 |
| 14555 | GPD1 | 38.6162609 | -3.6163603 | 0.41607375 | -8.6916331 | 3.57E-18 |
| 16172 | IL17RA | 1752.66546 | 1.00632499 | 0.11584864 | 8.68654968 | 3.74E-18 |
| 76499 | CLASP2 | 2050.27499 | 0.68998909 | 0.07944099 | 8.68555522 | 3.77E-18 |
| 73836 | SLC35B2 | 1539.6466 | 0.55549809 | 0.0640497 | 8.67292332 | 4.21E-18 |
| 14252 | FLOT2 | 2000.47667 | -0.9045998 | 0.10431946 | -8.6714387 | 4.27E-18 |
| 16890 | LIPE | 154.887455 | -1.7576115 | 0.2027134 | -8.670426 | 4.31E-18 |
| 12468 | CCT7 | 13518.8658 | 0.74288086 | 0.08575607 | 8.66272088 | 4.61E-18 |
| 78388 | MVP | 1836.00109 | -1.3593137 | 0.15700457 | -8.6577966 | 4.81E-18 |
| 244864 | LAYN | 2023.33051 | 4.10829829 | 0.47482893 | 8.652165 | 5.05E-18 |
| 666532 | Zfp991 | 41.6193581 | -5.7859173 | 0.66873846 | -8.6519882 | 5.06E-18 |
| 72121 | DENND2D | 56.3650972 | -9.2447697 | 1.06913141 | -8.6469911 | 5.29E-18 |
| 50876 | TMOD2 | 177.851388 | 3.66275489 | 0.42372778 | 8.64412264 | 5.42E-18 |
| 574428 | ZMYND15 | 54.1021282 | -5.3813046 | 0.62259162 | -8.6433939 | 5.46E-18 |
| 11576 | AFP | 172.120875 | 3.07838149 | 0.35619198 | 8.64247837 | 5.50E-18 |
| 67102 | D16Ertd472e | 688.997017 | 1.16069369 | 0.13438923 | 8.6368056 | 5.78E-18 |
| 319839 | A530020G20 | 118.299311 | -8.8736451 | 1.02762652 | -8.6350876 | 5.87E-18 |
| 56318 | ACPP | 317.015618 | -4.7786437 | 0.55360406 | -8.6318798 | 6.04E-18 |
| 58185 | RSAD2 | 69.0695072 | -2.8261526 | 0.32742704 | -8.6313964 | 6.06E-18 |
| 216618 | CFAP36 | 942.394751 | -0.6605983 | 0.07656172 | -8.6283114 | 6.23E-18 |
| 60315 | myg1 | 791.592625 | -0.9927764 | 0.11511091 | -8.6245204 | 6.44E-18 |
| 67216 | MBOAT2 | 131.045524 | -2.5672446 | 0.2976684 | -8.6245116 | 6.44E-18 |
| 20130 | RRAS | 677.131707 | -1.1461739 | 0.13289956 | -8.6243615 | 6.45E-18 |
| 15234 | HGF | 47.9379157 | 6.60281267 | 0.76573508 | 8.62284205 | 6.53E-18 |
| 216363 | RAB3IP | 1504.25747 | -0.5544629 | 0.06433449 | -8.6184394 | 6.79E-18 |
| 22724 | ZBTB7B | 1944.85012 | -0.9940086 | 0.11536447 | -8.6162452 | 6.92E-18 |
| 235610 | ATRIP | 564.985666 | 0.64855144 | 0.07529679 | 8.61326852 | 7.10E-18 |
| 242960 | FBXL5 | 1850.38295 | -0.3878362 | 0.04503262 | -8.6123385 | 7.16E-18 |
| 71900 | TMEM106B | 7126.65682 | -0.861218 | 0.10002876 | -8.6097034 | 7.32E-18 |
| 71874 | 2310007B03 | 161.816356 | -5.3630654 | 0.62291117 | -8.6096793 | 7.33E-18 |
| 268783 | MTMR12 | 1592.40603 | 0.78338269 | 0.090999 | 8.60869513 | 7.39E-18 |
| 26874 | ABCD2 | 33.5741564 | 3.16670404 | 0.36787156 | 8.60817848 | 7.42E-18 |
| 12038 | BCHE | 194.976646 | -11.036944 | 1.28401828 | -8.5956282 | 8.28E-18 |
| 16161 | IL12RB1 | 74.7097354 | 3.81521493 | 0.44387471 | 8.59525192 | 8.31E-18 |
| 234889 | GUCY1A2 | 60.7732844 | -5.5460936 | 0.6452816 | -8.5948423 | 8.34E-18 |

| | | | | | | |
|---|---|---|---|---|---|---|
| 100502841 | EPG5 | 1428.3964 | -0.8785477 | 0.1022658 | -8.5908263 | 8.63E-18 |
| 56375 | B4GALT4 | 267.82829 | -2.3450144 | 0.27298554 | -8.5902516 | 8.68E-18 |
| 243725 | PPP1R9A | 51.1208107 | -4.9196063 | 0.57296534 | -8.5862197 | 8.99E-18 |
| 433904 | OCIAD2 | 382.027534 | -2.1029573 | 0.24497314 | -8.5844405 | 9.13E-18 |
| 11731 | Ang2 | 274.063094 | 3.32483488 | 0.38734064 | 8.58374911 | 9.18E-18 |
| 52024 | ANKRD22 | 46.9592045 | -8.9813733 | 1.04799863 | -8.5700239 | 1.03E-17 |
| 67263 | ZSWIM6 | 710.904319 | 1.42608534 | 0.16662906 | 8.55844321 | 1.14E-17 |
| 76454 | FBXO31 | 888.032275 | 1.3407216 | 0.15665654 | 8.55835091 | 1.14E-17 |
| 109019 | NABP1 | 2256.51452 | -1.3282948 | 0.1552789 | -8.5542519 | 1.19E-17 |
| 67788 | SFR1 | 4070.07474 | 0.5730699 | 0.06700179 | 8.55305307 | 1.20E-17 |
| 98258 | TXNDC9 | 1857.3802 | -0.4375066 | 0.05118103 | -8.5482163 | 1.25E-17 |
| 66859 | SLC16A9 | 129.134243 | 5.28478265 | 0.6182615 | 8.5478113 | 1.25E-17 |
| 54561 | NAP1L3 | 74.36365 | -8.9216453 | 1.04373935 | -8.5477713 | 1.25E-17 |
| 20184 | UIMC1 | 797.497412 | 0.93741413 | 0.10969483 | 8.54565444 | 1.28E-17 |
| 216011 | LRRC20 | 383.976499 | -1.251458 | 0.14648146 | -8.543457 | 1.30E-17 |
| 170571 | CNTNAP4 | 83.5818155 | 9.14201799 | 1.07039951 | 8.54075313 | 1.33E-17 |
| 67398 | SRPR | 2307.78402 | 0.67005382 | 0.07847251 | 8.53870709 | 1.36E-17 |
| 12070 | BEX3 | 2792.51799 | -0.4772284 | 0.05589813 | -8.5374659 | 1.37E-17 |
| 66665 | MSANTD3 | 752.538211 | 0.73606851 | 0.08621771 | 8.53732364 | 1.37E-17 |
| 83383 | TFAP4 | 576.889736 | 0.91419537 | 0.1071385 | 8.53283732 | 1.43E-17 |
| 380614 | intu | 211.966275 | -0.9935613 | 0.11647025 | -8.5306017 | 1.46E-17 |
| 18426 | OVOL1 | 33.4912669 | -4.8320024 | 0.56667213 | -8.5269809 | 1.50E-17 |
| 66616 | SNX9 | 3463.94676 | 1.23258793 | 0.14457774 | 8.52543381 | 1.52E-17 |
| 239319 | CARD6 | 717.755171 | 1.03202326 | 0.12107922 | 8.52353712 | 1.55E-17 |
| 16431 | ITM2A | 123.892387 | -8.9374814 | 1.04908564 | -8.5193058 | 1.61E-17 |
| 621837 | Gm6265 | 614.437629 | 0.70990868 | 0.08336352 | 8.51581964 | 1.65E-17 |
| 224093 | FAM43A | 188.9747 | 1.75942988 | 0.20668662 | 8.51254864 | 1.70E-17 |
| 19654 | RBM6 | 1378.99797 | 0.4204051 | 0.04938863 | 8.51218359 | 1.71E-17 |
| 68404 | NRN1 | 254.837011 | 5.09256443 | 0.59857703 | 8.50778461 | 1.77E-17 |
| 67469 | ABHD5 | 2541.33454 | 1.32060487 | 0.15522395 | 8.50773915 | 1.77E-17 |
| 77853 | MSL2 | 1592.85944 | 0.52569747 | 0.06181854 | 8.50388061 | 1.83E-17 |
| 237175 | ADGRG2 | 1518.60217 | -2.5219692 | 0.29692247 | -8.4936962 | 2.00E-17 |
| 192120 | BSPRY | 89.2882859 | -7.8846794 | 0.92836069 | -8.4931207 | 2.01E-17 |
| 245007 | ZBTB38 | 2219.86439 | -1.1207954 | 0.13196518 | -8.4931148 | 2.01E-17 |
| 66603 | GEMIN2 | 452.803987 | 0.67149286 | 0.07907618 | 8.49172077 | 2.04E-17 |
| 671878 | Gm14680 | 364.614806 | 1.17781902 | 0.1387803 | 8.4869326 | 2.12E-17 |
| 11790 | SPEG | 60.9016066 | -2.5046882 | 0.29542111 | -8.4783655 | 2.28E-17 |
| 14667 | GM2A | 1285.41254 | -2.3679996 | 0.2793018 | -8.4782828 | 2.29E-17 |
| 330122 | CXCL3 | 247.948044 | -9.9489913 | 1.17350816 | -8.4779907 | 2.29E-17 |
| 15116 | HAS1 | 128.779417 | 9.76656455 | 1.15263401 | 8.47325731 | 2.39E-17 |
| 237459 | CDK17 | 1501.18463 | -0.5476499 | 0.06463529 | -8.4729249 | 2.39E-17 |
| 19219 | PTGER4 | 566.512022 | 2.59414986 | 0.30639013 | 8.46681924 | 2.52E-17 |
| 223827 | GXYLT1 | 2160.49641 | -0.6731103 | 0.07950461 | -8.4663051 | 2.53E-17 |
| 18596 | PDGFRB | 2855.734 | 8.03696441 | 0.94934917 | 8.46576226 | 2.54E-17 |
| 56442 | SERINC1 | 8577.90031 | -0.3543899 | 0.04186348 | -8.4653703 | 2.55E-17 |

| | | | | | | |
|---|---|---|---|---|---|---|
| 20405 | SH3GL1 | 1940.93272 | 0.6097208 | 0.07203214 | 8.46456652 | 2.57E-17 |
| 15220 | FOXQ1 | 129.3496 | -2.6993004 | 0.31894438 | -8.4632323 | 2.60E-17 |
| 18789 | PAPOLA | 9651.34324 | 0.53074957 | 0.06272563 | 8.46144598 | 2.64E-17 |
| 28169 | AGPAT3 | 2586.37458 | -0.8744597 | 0.10335282 | -8.460918 | 2.65E-17 |
| 20818 | SRPRB | 2288.47703 | 0.56311402 | 0.06655933 | 8.4603313 | 2.67E-17 |
| 12492 | SCARB2 | 2039.12532 | -1.2212617 | 0.14443297 | -8.4555604 | 2.78E-17 |
| 20710 | Serpinb9e | 51.4113806 | -4.3463503 | 0.51403508 | -8.4553573 | 2.78E-17 |
| 22041 | trf | 230.323245 | -8.0232347 | 0.94899693 | -8.454437 | 2.80E-17 |
| 18762 | PRKCZ | 42.9197825 | -8.8529812 | 1.04718565 | -8.4540704 | 2.81E-17 |
| 67865 | RGS10 | 147.38133 | 2.27827932 | 0.26951152 | 8.45336538 | 2.83E-17 |
| 226841 | VASH2 | 816.078022 | 1.97901513 | 0.23411929 | 8.4530202 | 2.84E-17 |
| 22288 | UTRN | 3902.40806 | -1.0068315 | 0.11914051 | -8.4507902 | 2.89E-17 |
| 20908 | STX3 | 901.673103 | -1.0609228 | 0.12556391 | -8.4492658 | 2.93E-17 |
| 240025 | DACT2 | 264.694491 | -10.028953 | 1.18696777 | -8.449221 | 2.93E-17 |
| 18807 | PLD3 | 1893.76649 | -1.0560279 | 0.12502412 | -8.4465929 | 3.00E-17 |
| 100043314 | Tigit | 400.075867 | -5.2529523 | 0.62201296 | -8.4450849 | 3.04E-17 |
| 68992 | Zfp580 | 97.7847452 | 2.74983663 | 0.32561724 | 8.44499705 | 3.04E-17 |
| 97884 | B3GALNT2 | 1650.07897 | -0.6730382 | 0.07972599 | -8.441892 | 3.12E-17 |
| 11836 | ARAF | 1677.5545 | -0.505986 | 0.05993906 | -8.4416743 | 3.13E-17 |
| 22057 | TOB1 | 1374.01674 | -1.6702073 | 0.19792449 | -8.4386085 | 3.21E-17 |
| 22632 | YY1 | 3599.13468 | 0.59332701 | 0.07035578 | 8.43323759 | 3.36E-17 |
| 270163 | MYO9A | 2464.27648 | 0.82736545 | 0.09813262 | 8.43109544 | 3.42E-17 |
| 11830 | AQP5 | 517.418751 | -5.9221393 | 0.70251548 | -8.4299058 | 3.46E-17 |
| 14528 | GCH1 | 390.327268 | -2.634739 | 0.31272965 | -8.4249733 | 3.61E-17 |
| 77552 | SHISA4 | 608.283385 | -2.0348459 | 0.24162521 | -8.4214966 | 3.72E-17 |
| 15051 | H2-T9 | 217.466072 | -1.4853313 | 0.17642121 | -8.4192328 | 3.79E-17 |
| 71409 | FMNL2 | 2984.40831 | -1.6302646 | 0.19371541 | -8.4157716 | 3.90E-17 |
| 12010 | B2M | 14725.3789 | -1.4087543 | 0.16754407 | -8.408261 | 4.16E-17 |
| 14872 | GstT2 | 53.7795572 | -3.9935529 | 0.47504601 | -8.4066656 | 4.22E-17 |
| 77318 | ankrd55 | 69.4841695 | 2.62056891 | 0.31196475 | 8.40020837 | 4.46E-17 |
| 14709 | GNG8 | 35.3241665 | 5.84092023 | 0.69560263 | 8.39692081 | 4.58E-17 |
| 234967 | SLC36A4 | 281.3684 | 0.99231521 | 0.11819731 | 8.39541287 | 4.64E-17 |
| 114715 | SPRED1 | 2948.91083 | 1.10528253 | 0.13176764 | 8.38811821 | 4.94E-17 |
| 14588 | GFRA4 | 32.0062299 | 3.70075473 | 0.44121234 | 8.38769546 | 4.96E-17 |
| 330256 | Gm20559 | 76.6921827 | -1.9031209 | 0.22699064 | -8.3841382 | 5.11E-17 |
| 18779 | PLA2R1 | 138.079323 | -4.5334365 | 0.54081839 | -8.3825486 | 5.18E-17 |
| 74011 | SLC25A27 | 55.7902324 | -3.7709322 | 0.45000773 | -8.3797054 | 5.31E-17 |
| 213649 | ARHGEF19 | 337.621087 | -1.8612009 | 0.2221778 | -8.377079 | 5.43E-17 |
| 69771 | 1810019D21 | 33.1930453 | -4.9821534 | 0.59482581 | -8.375819 | 5.48E-17 |
| 74256 | CYLD | 2464.67209 | -0.6899934 | 0.08239714 | -8.373997 | 5.57E-17 |
| 67759 | PLGRKT | 503.231883 | -0.8848891 | 0.10568229 | -8.3731068 | 5.61E-17 |
| 17921 | MYO7A | 265.336443 | -6.321798 | 0.75517705 | -8.3712793 | 5.70E-17 |
| 140500 | acap3 | 1765.00429 | 1.19017163 | 0.14219874 | 8.36977592 | 5.77E-17 |
| 20481 | SKI | 4492.65662 | 0.70278834 | 0.08406027 | 8.36052939 | 6.24E-17 |
| 208285 | Cyp4f17 | 186.963782 | -1.8403515 | 0.22017011 | -8.3587705 | 6.34E-17 |

| | | | | | | |
|---|---|---|---|---|---|---|
| 69894 | 2010107G23 | 75.1843961 | -9.6634327 | 1.15628164 | -8.3573347 | 6.42E-17 |
| 69875 | NDUFA11 | 814.050591 | -0.5342201 | 0.06394421 | -8.3544721 | 6.57E-17 |
| 16973 | LRP5 | 3183.35835 | -1.0462254 | 0.12523092 | -8.35437 | 6.58E-17 |
| 98878 | EHD4 | 3124.10499 | -2.0171289 | 0.24146284 | -8.3537861 | 6.61E-17 |
| 56213 | HTRA1 | 6914.07527 | -1.8700234 | 0.22398023 | -8.3490559 | 6.88E-17 |
| 319875 | TMPRSS11B | 46.1526002 | -2.505556 | 0.30010656 | -8.3488877 | 6.89E-17 |
| 66925 | SDHD | 3006.86412 | 0.57147241 | 0.06846329 | 8.34713644 | 6.99E-17 |
| 17392 | MMP3 | 1307.72453 | 3.87925157 | 0.46481219 | 8.34584737 | 7.07E-17 |
| 60440 | iigp1 | 69.6495328 | -4.9182596 | 0.58937422 | -8.3448841 | 7.13E-17 |
| 67433 | CCDC127 | 1995.5204 | 0.43943739 | 0.05266443 | 8.34410306 | 7.18E-17 |
| 259302 | SRGAP3 | 1830.00079 | -1.000542 | 0.1201671 | -8.3262556 | 8.34E-17 |
| 74043 | PEX26 | 566.199793 | -1.1037004 | 0.13260415 | -8.3232719 | 8.56E-17 |
| 13682 | EIF4A2 | 3634.91091 | -0.5156192 | 0.06195916 | -8.3219214 | 8.65E-17 |
| 16593 | KLC1 | 4595.37393 | 0.88974117 | 0.10695194 | 8.3190749 | 8.87E-17 |
| 24052 | SGCD | 67.1781334 | 8.82728424 | 1.06174004 | 8.31397887 | 9.25E-17 |
| 230784 | SESN2 | 831.615149 | -0.7540306 | 0.09072834 | -8.3108608 | 9.50E-17 |
| 74610 | ABCB8 | 689.533783 | -0.7729513 | 0.09303574 | -8.3081112 | 9.72E-17 |
| 54448 | Il1f6 | 58.5018863 | -7.2688991 | 0.87597448 | -8.2980718 | 1.06E-16 |
| 17769 | MTHFR | 999.326338 | 1.25836604 | 0.15167991 | 8.29619482 | 1.07E-16 |
| 380785 | BEGAIN | 33.1358984 | 4.56579459 | 0.55035268 | 8.29612489 | 1.08E-16 |
| 71908 | CLDN23 | 47.6995695 | -9.0084439 | 1.08590335 | -8.2958063 | 1.08E-16 |
| 15374 | HN1 | 3372.73346 | -0.6331976 | 0.07634494 | -8.2939029 | 1.10E-16 |
| 66569 | GDPD1 | 790.471555 | -2.2847646 | 0.27554939 | -8.29167 | 1.12E-16 |
| 57905 | ISY1 | 1564.04408 | 1.1852849 | 0.14295095 | 8.29154988 | 1.12E-16 |
| 140494 | ATP6V0A4 | 84.9509897 | 1.74237295 | 0.21016676 | 8.29043069 | 1.13E-16 |
| 231380 | UBA6 | 2177.35236 | -0.4177297 | 0.05040042 | -8.2882193 | 1.15E-16 |
| 260315 | NAV3 | 1936.23136 | 0.90041043 | 0.10864951 | 8.28729381 | 1.16E-16 |
| 21844 | TIAM1 | 830.102454 | -3.3126494 | 0.39972749 | -8.2872696 | 1.16E-16 |
| 245827 | FAT2 | 180.745629 | -7.8804259 | 0.95179355 | -8.2795538 | 1.24E-16 |
| 56316 | GGCX | 829.986191 | 0.68715206 | 0.08301907 | 8.27703873 | 1.26E-16 |
| 319758 | RFX7 | 2463.43131 | 0.41354054 | 0.04999401 | 8.27180243 | 1.32E-16 |
| 105841 | DENND3 | 311.2144 | -2.3285574 | 0.28169862 | -8.2661301 | 1.38E-16 |
| 230157 | TMEFF1 | 358.471874 | 2.96428407 | 0.35862204 | 8.26576102 | 1.39E-16 |
| 67460 | DECR1 | 634.385575 | -1.7988829 | 0.21764966 | -8.2650391 | 1.40E-16 |
| 72543 | MVB12B | 1241.91375 | -1.3725582 | 0.16618599 | -8.2591692 | 1.47E-16 |
| 56358 | COPZ2 | 255.567829 | 1.13657787 | 0.13763692 | 8.25779788 | 1.48E-16 |
| 66994 | CEP19 | 346.407228 | -0.828101 | 0.10033026 | -8.2537508 | 1.53E-16 |
| 108888 | ATAD3A | 1442.43617 | 0.72012243 | 0.08727515 | 8.25117342 | 1.57E-16 |
| 105853 | MAL2 | 160.015523 | -10.029233 | 1.21550121 | -8.2511091 | 1.57E-16 |
| 217615 | #N/A | 2766.33852 | 0.51207337 | 0.06217066 | 8.23657602 | 1.77E-16 |
| 18131 | NOTCH3 | 93.1175465 | -3.7955781 | 0.46150112 | -8.224418 | 1.96E-16 |
| 52850 | SGSM1 | 335.747294 | -1.9168682 | 0.23339141 | -8.2131051 | 2.16E-16 |
| 19015 | PPARD | 2570.44762 | 1.36862322 | 0.16665135 | 8.21249387 | 2.17E-16 |
| 226976 | KANSL3 | 1946.70398 | -0.7611823 | 0.09269217 | -8.2119371 | 2.18E-16 |
| 67429 | NUDCD1 | 2217.16479 | 1.30544481 | 0.15903859 | 8.20835257 | 2.24E-16 |

| | | | | | | |
|---|---|---|---|---|---|---|
| 12317 | CALR | 25783.6527 | 0.46384556 | 0.05653129 | 8.20511185 | 2.30E-16 |
| 320100 | RELT | 368.84885 | 1.38293398 | 0.16884431 | 8.19058684 | 2.60E-16 |
| 67071 | RPS6KA6 | 215.605428 | -11.182111 | 1.36551733 | -8.1889195 | 2.64E-16 |
| 20639 | SNRPB2 | 1290.67111 | 0.60402383 | 0.07380897 | 8.18361018 | 2.75E-16 |
| 17240 | MDFI | 84.8275298 | -4.6634509 | 0.5699535 | -8.1821603 | 2.79E-16 |
| 228432 | ANO3 | 82.585361 | -7.768672 | 0.94989064 | -8.1784909 | 2.87E-16 |
| 67873 | mri1 | 551.826872 | 0.8133069 | 0.09945724 | 8.17745329 | 2.90E-16 |
| 74682 | WDR35 | 1430.18111 | -0.7525761 | 0.09203214 | -8.1773184 | 2.90E-16 |
| 21888 | TLE4 | 54.8806178 | -9.206984 | 1.12621067 | -8.1751881 | 2.95E-16 |
| 68839 | ANKRD46 | 1732.91147 | 0.56838228 | 0.06953498 | 8.17404884 | 2.98E-16 |
| 105859 | CSDC2 | 43.393883 | -5.4153316 | 0.66266935 | -8.1719965 | 3.03E-16 |
| 71648 | OPTN | 800.509019 | -1.610232 | 0.1971515 | -8.1674855 | 3.15E-16 |
| 241520 | FAM171B | 379.279134 | 0.93804703 | 0.11493092 | 8.16183321 | 3.30E-16 |
| 109658 | TXLNA | 3929.39134 | 1.12048517 | 0.13729923 | 8.16089925 | 3.33E-16 |
| 12785 | CNBP | 17326.9707 | 0.88370452 | 0.10840902 | 8.15157707 | 3.59E-16 |
| 272636 | ESYT3 | 186.45044 | -8.5224673 | 1.0456252 | -8.1505948 | 3.62E-16 |
| 66443 | TNFAIP8L1 | 70.6742688 | -2.0762295 | 0.25475223 | -8.1499953 | 3.64E-16 |
| 230657 | TMEM69 | 443.758586 | 0.89573623 | 0.10992705 | 8.14846075 | 3.69E-16 |
| 241075 | PLEKHM3 | 660.744277 | -1.2894178 | 0.15826172 | -8.1473762 | 3.72E-16 |
| 56209 | GDE1 | 2022.14194 | -0.5350775 | 0.06569172 | -8.145281 | 3.78E-16 |
| 68126 | FAHD2A | 34.7651104 | -8.5489076 | 1.05008562 | -8.141153 | 3.92E-16 |
| 117591 | SLC2A9 | 27.4935639 | -3.4001195 | 0.41767426 | -8.1406011 | 3.93E-16 |
| 231646 | MYO1H | 71.2949308 | 2.20048564 | 0.2703382 | 8.13975088 | 3.96E-16 |
| 15114 | HAP1 | 88.6560329 | -1.6323256 | 0.20055996 | -8.1388406 | 3.99E-16 |
| 54325 | ELOVL1 | 2249.9739 | 0.97684769 | 0.12003596 | 8.13795884 | 4.02E-16 |
| 19222 | PTGIR | 34.131133 | 8.57299098 | 1.05361024 | 8.13677646 | 4.06E-16 |
| 74766 | YIPF2 | 294.92848 | -1.2506576 | 0.15379183 | -8.1321457 | 4.22E-16 |
| 12395 | RUNX1T1 | 54.515656 | 8.52304273 | 1.0484578 | 8.12912329 | 4.32E-16 |
| 22259 | NR1H3 | 52.0206298 | -6.3598076 | 0.78282903 | -8.1241335 | 4.51E-16 |
| 19042 | PPM1A | 3413.98768 | 0.40061512 | 0.04932991 | 8.12113974 | 4.62E-16 |
| 83398 | NDST3 | 38.3130685 | -2.6905173 | 0.33143967 | -8.1176684 | 4.75E-16 |
| 210530 | P3H2 | 85.80529 | 1.79990587 | 0.22172696 | 8.11766806 | 4.75E-16 |
| 14048 | EYA1 | 77.485144 | 5.56085876 | 0.68506911 | 8.11722303 | 4.77E-16 |
| 12925 | CRIP1 | 958.368933 | 1.68675617 | 0.20780335 | 8.11707891 | 4.78E-16 |
| 17192 | MBD3 | 4263.16608 | 0.44572659 | 0.05492071 | 8.11581935 | 4.83E-16 |
| 140486 | IGF2BP1 | 499.824646 | -3.7663692 | 0.46418355 | -8.1139654 | 4.90E-16 |
| 12904 | CRABP2 | 368.629613 | 2.89911591 | 0.35738484 | 8.11202812 | 4.98E-16 |
| 68612 | UBE2C | 3581.49562 | 0.39604673 | 0.04882762 | 8.11112056 | 5.02E-16 |
| 12305 | ddr1 | 2446.01243 | -1.0371337 | 0.12787979 | -8.1102243 | 5.05E-16 |
| 52480 | SNHG14 | 53.6150848 | -9.1732732 | 1.1314035 | -8.1078706 | 5.15E-16 |
| 14469 | GBP2 | 310.253417 | -1.1949966 | 0.14747367 | -8.1031184 | 5.36E-16 |
| 235534 | PXYLP1 | 687.041327 | -1.3478406 | 0.16637287 | -8.1013242 | 5.44E-16 |
| 97827 | EXD2 | 730.734907 | 0.69875992 | 0.086257 | 8.10090639 | 5.46E-16 |
| 74596 | cds1 | 51.280424 | -6.31325 | 0.77942731 | -8.0998574 | 5.50E-16 |
| 20135 | RRM2 | 10643.1192 | 0.65650367 | 0.08105193 | 8.09979027 | 5.51E-16 |

| | | | | | | |
|---|---|---|---|---|---|---|
| 67043 | SYAP1 | 1434.21023 | -0.5206118 | 0.06435385 | -8.0898317 | 5.97E-16 |
| 97863 | FAM8A1 | 603.985568 | -0.7505378 | 0.09279498 | -8.0881294 | 6.06E-16 |
| 217218 | ATXN7L3 | 2595.25244 | 0.33729361 | 0.04170513 | 8.08758049 | 6.09E-16 |
| 71982 | SNX10 | 917.482469 | -0.5765971 | 0.07130354 | -8.0865145 | 6.14E-16 |
| 74376 | MYO18B | 107.574686 | -3.3002855 | 0.40815744 | -8.085815 | 6.17E-16 |
| 22213 | UBE2G2 | 1930.06032 | -0.5596939 | 0.06924645 | -8.0826371 | 6.34E-16 |
| 78038 | MCCC2 | 321.265818 | -0.9373668 | 0.11600667 | -8.0802836 | 6.46E-16 |
| 23989 | MED24 | 1657.28364 | 0.40977084 | 0.05071471 | 8.07992108 | 6.48E-16 |
| 18616 | PEG3 | 1639.12246 | -14.10878 | 1.74711757 | -8.0754611 | 6.72E-16 |
| 13113 | Cyp3a13 | 55.2916471 | -8.4959913 | 1.05227295 | -8.0739425 | 6.81E-16 |
| 77877 | 6030458C11I | 1158.24353 | 0.81363852 | 0.10078053 | 8.07337016 | 6.84E-16 |
| 12151 | BMI1 | 2671.62181 | 0.56621174 | 0.07018561 | 8.06734757 | 7.18E-16 |
| 77574 | TCAF1 | 992.707887 | -0.6619726 | 0.08206113 | -8.066823 | 7.22E-16 |
| 20019 | POLR1A | 6564.54864 | 0.9710808 | 0.12039717 | 8.065645 | 7.29E-16 |
| 73095 | SLC25A42 | 133.194297 | -1.5982828 | 0.1981817 | -8.0647349 | 7.34E-16 |
| 242915 | GAREM2 | 126.953403 | 2.65262867 | 0.32892009 | 8.06466002 | 7.34E-16 |
| 74143 | OPA1 | 3167.41506 | -0.5376788 | 0.06668517 | -8.0629434 | 7.45E-16 |
| 18933 | PRRX1 | 8837.01789 | 4.30883455 | 0.5346042 | 8.0598591 | 7.64E-16 |
| 107586 | OVOL2 | 36.1432343 | -6.1465319 | 0.76281017 | -8.0577477 | 7.77E-16 |
| 18392 | ORC1 | 975.845013 | 1.22672562 | 0.15251806 | 8.04314988 | 8.76E-16 |
| 22156 | TUFT1 | 301.585659 | -2.6870925 | 0.33445628 | -8.0342116 | 9.42E-16 |
| 70021 | NT5DC2 | 1201.1799 | 3.99872812 | 0.4980433 | 8.02887646 | 9.84E-16 |
| 237082 | NXT2 | 718.338449 | -0.5581565 | 0.06952969 | -8.0275993 | 9.94E-16 |
| 17132 | MAF | 267.60028 | 3.07915155 | 0.38371529 | 8.02457342 | 1.02E-15 |
| 53416 | STK39 | 2700.75602 | -1.7432837 | 0.21724779 | -8.0244023 | 1.02E-15 |
| 69047 | ATP2C2 | 34.5280271 | -5.7486447 | 0.71643673 | -8.0239391 | 1.02E-15 |
| 215789 | PHACTR2 | 80.2767128 | -6.5028847 | 0.81090606 | -8.0192825 | 1.06E-15 |
| 105522 | ANKRD28 | 5792.83498 | 1.51336282 | 0.18883815 | 8.01407354 | 1.11E-15 |
| 94062 | MRPL3 | 1355.94886 | 0.48533115 | 0.06056196 | 8.01379467 | 1.11E-15 |
| 67951 | TUBB6 | 11053.533 | -0.263067 | 0.03282768 | -8.0135741 | 1.11E-15 |
| 234725 | Zfp612 | 42.5740941 | -8.8400912 | 1.10359859 | -8.0102415 | 1.14E-15 |
| 101631 | PWWP2B | 1032.59958 | -1.0237174 | 0.12780915 | -8.0097346 | 1.15E-15 |
| 13123 | CYP7B1 | 36.7807456 | -8.6293508 | 1.07753817 | -8.0083945 | 1.16E-15 |
| 11595 | ACAN | 96.4327361 | -9.2993996 | 1.1612591 | -8.0080316 | 1.17E-15 |
| 105734 | TIGD5 | 152.290173 | 1.15106125 | 0.14375995 | 8.00682811 | 1.18E-15 |
| 16562 | KIF1C | 13949.0191 | -0.8239707 | 0.10294363 | -8.0040958 | 1.20E-15 |
| 72580 | ZUFSP | 228.790973 | -0.8840444 | 0.11050866 | -7.9997749 | 1.25E-15 |
| 72649 | TMEM209 | 2171.55571 | -0.568547 | 0.07111261 | -7.9950247 | 1.30E-15 |
| 12589 | IFT81 | 344.251019 | -0.9342028 | 0.11685854 | -7.994305 | 1.30E-15 |
| 19024 | PPFIBP2 | 420.110001 | -5.5773458 | 0.697693 | -7.9939827 | 1.31E-15 |
| 94093 | TRIM33 | 2065.50195 | -0.4533884 | 0.05671789 | -7.9937465 | 1.31E-15 |
| 212168 | ZSWIM4 | 1157.4909 | -0.7945251 | 0.09949444 | -7.9856224 | 1.40E-15 |
| 67455 | KLHL13 | 35.4992771 | -8.577599 | 1.07414204 | -7.9855351 | 1.40E-15 |
| 99890 | PRMT6 | 807.445305 | 0.59848235 | 0.07497454 | 7.98247452 | 1.43E-15 |
| 50934 | SLC7A8 | 167.949257 | 7.58520945 | 0.95151211 | 7.9717424 | 1.56E-15 |

| | | | | | | |
|---|---|---|---|---|---|---|
| 67887 | SARAF | 2230.8869 | -0.3473703 | 0.04357777 | -7.9712734 | 1.57E-15 |
| 19179 | PSMC1 | 3540.77797 | 0.58545161 | 0.0734682 | 7.96877572 | 1.60E-15 |
| 228765 | SDCBP2 | 107.599726 | -4.072753 | 0.51126489 | -7.966033 | 1.64E-15 |
| 232035 | CCSER1 | 48.7714818 | -8.31239 | 1.04385052 | -7.9631995 | 1.68E-15 |
| 76178 | COA5 | 3242.70971 | -0.5846355 | 0.07345547 | -7.9590466 | 1.73E-15 |
| 100952 | EMILIN1 | 198.04206 | 4.75154052 | 0.59700172 | 7.95900644 | 1.73E-15 |
| 14874 | GSTZ1 | 623.313471 | 0.65273174 | 0.08202749 | 7.95747524 | 1.76E-15 |
| 234959 | MED17 | 1300.80326 | 0.56971995 | 0.07162098 | 7.95465191 | 1.80E-15 |
| 232187 | SMYD5 | 2085.26524 | 0.96224467 | 0.12098626 | 7.9533387 | 1.82E-15 |
| 22320 | VAMP8 | 1536.53901 | -1.0193657 | 0.1282069 | -7.9509425 | 1.85E-15 |
| 230603 | TTC39A | 203.134477 | -3.2325461 | 0.40705355 | -7.941329 | 2.00E-15 |
| 13136 | CD55 | 146.823369 | -6.2589505 | 0.78819589 | -7.9408566 | 2.01E-15 |
| 71901 | FAM219A | 307.339489 | 1.35864973 | 0.17114892 | 7.93840677 | 2.05E-15 |
| 11370 | ACADVL | 1384.57957 | -0.7031131 | 0.08858403 | -7.9372441 | 2.07E-15 |
| 72114 | ZBED3 | 854.977915 | 1.04531329 | 0.13169987 | 7.93708679 | 2.07E-15 |
| 224807 | TMEM63B | 3363.00049 | -0.9225114 | 0.11622807 | -7.9370793 | 2.07E-15 |
| 21841 | TIA1 | 2022.48615 | 0.50185444 | 0.06324031 | 7.9356732 | 2.09E-15 |
| 192657 | ELL2 | 2431.8523 | 0.84389797 | 0.10634972 | 7.93512172 | 2.10E-15 |
| 19248 | PTPN12 | 2690.65532 | -0.4049313 | 0.05104081 | -7.9334803 | 2.13E-15 |
| 14457 | GAS7 | 866.192633 | 1.3343317 | 0.16819703 | 7.93314656 | 2.14E-15 |
| 27883 | TANGO2 | 248.317197 | -0.8755086 | 0.11040401 | -7.9300432 | 2.19E-15 |
| 235559 | TOPBP1 | 7790.64097 | 0.48854849 | 0.06164993 | 7.92455853 | 2.29E-15 |
| 14254 | FLT1 | 886.535794 | -2.4962483 | 0.31514917 | -7.9208469 | 2.36E-15 |
| 69225 | NAXD | 880.789739 | -0.4791102 | 0.06049293 | -7.9201014 | 2.37E-15 |
| 73998 | herc3 | 1453.7909 | -0.6953468 | 0.08784993 | -7.9151661 | 2.47E-15 |
| 19087 | PRKAR2A | 4239.66066 | 1.0333429 | 0.13055286 | 7.91513032 | 2.47E-15 |
| 654470 | tctn1 | 211.689504 | -1.1291025 | 0.14265641 | -7.9148391 | 2.48E-15 |
| 269514 | FBXL4 | 429.189365 | -1.5723415 | 0.19868253 | -7.9138387 | 2.50E-15 |
| 233571 | P2RY6 | 27.950871 | -4.7340427 | 0.59823794 | -7.9133107 | 2.51E-15 |
| 67781 | ILF2 | 3130.55551 | 0.40989841 | 0.05180201 | 7.91278896 | 2.52E-15 |
| 211577 | MRGPRF | 92.9899406 | 9.2962825 | 1.17494281 | 7.91211491 | 2.53E-15 |
| 67117 | DYNLT3 | 2728.11112 | -1.5040858 | 0.19026672 | -7.9051441 | 2.68E-15 |
| 66483 | RPL36AL | 530.226108 | 0.55743343 | 0.07051851 | 7.90478131 | 2.68E-15 |
| 408022 | PRIMPOL | 263.683382 | -0.9997968 | 0.12651315 | -7.9027106 | 2.73E-15 |
| 73828 | DCAF4 | 581.995334 | -0.7708063 | 0.0975919 | -7.8982608 | 2.83E-15 |
| 17777 | MTTP | 177.314508 | -2.3794072 | 0.30140137 | -7.8944805 | 2.92E-15 |
| 56403 | SYNCRIP | 9286.27856 | 0.79002705 | 0.10009823 | 7.89251806 | 2.96E-15 |
| 68537 | MRPL13 | 1820.1582 | 0.73777481 | 0.0934789 | 7.89242069 | 2.96E-15 |
| 209011 | SIRT7 | 682.519466 | -0.9049897 | 0.11466736 | -7.8923043 | 2.97E-15 |
| 224088 | ATP13A3 | 9646.21929 | -0.4494992 | 0.05695592 | -7.8920544 | 2.97E-15 |
| 18749 | PRKACB | 5938.40226 | -1.7023151 | 0.21571494 | -7.8915033 | 2.99E-15 |
| 319269 | A130040M12 | 1258.84478 | 3.82065092 | 0.48416 | 7.89129815 | 2.99E-15 |
| 66168 | GRINA | 3866.62369 | -1.4666407 | 0.18601476 | -7.8845392 | 3.16E-15 |
| 75568 | CAPSL | 30.024265 | -8.3383705 | 1.0578001 | -7.8827469 | 3.20E-15 |
| 94220 | CNNM4 | 1019.87826 | -0.7997599 | 0.1014591 | -7.8825847 | 3.21E-15 |

| | | | | | | |
|---|---|---|---|---|---|---|
| 226162 | DPCD | 363.263924 | -0.7964172 | 0.10106604 | -7.8801658 | 3.27E-15 |
| 75273 | pelp1 | 2313.33028 | 0.39227008 | 0.04978105 | 7.87990824 | 3.28E-15 |
| 110639 | PRPS2 | 1941.63058 | -0.7003876 | 0.08893676 | -7.8751194 | 3.40E-15 |
| 17169 | MARK3 | 3636.55189 | 0.58915042 | 0.07487942 | 7.86798853 | 3.60E-15 |
| 14859 | GSTA3 | 43.5084629 | -3.7475896 | 0.47643805 | -7.8658487 | 3.67E-15 |
| 18551 | PCSK4 | 108.506047 | 1.57718713 | 0.20060684 | 7.86208041 | 3.78E-15 |
| 19070 | MOB4 | 2265.37444 | -0.6069688 | 0.07720366 | -7.8619176 | 3.78E-15 |
| 73916 | IFT57 | 542.807736 | -0.8459643 | 0.10763039 | -7.8599013 | 3.84E-15 |
| 229900 | gbp7 | 238.527731 | -2.4616697 | 0.31336605 | -7.8555723 | 3.98E-15 |
| 80281 | CTTNBP2NL | 2674.03344 | -0.2976418 | 0.03789276 | -7.8548441 | 4.00E-15 |
| 207742 | RNF43 | 57.7682472 | -8.5599828 | 1.08988565 | -7.8540192 | 4.03E-15 |
| 58521 | EID1 | 1947.07733 | 0.7908643 | 0.10070717 | 7.85310812 | 4.06E-15 |
| 73288 | VPS50 | 1490.09951 | -0.7525171 | 0.09583148 | -7.852504 | 4.08E-15 |
| 69274 | CTDSPL | 967.894963 | -0.63517 | 0.08095167 | -7.8462858 | 4.29E-15 |
| 69709 | PTRHD1 | 433.480054 | 0.719429 | 0.09169936 | 7.84551858 | 4.31E-15 |
| 108937 | RNF169 | 1037.05232 | -0.7893998 | 0.10065242 | -7.8428294 | 4.41E-15 |
| 106565 | DLK2 | 360.136425 | -2.035758 | 0.25959187 | -7.8421483 | 4.43E-15 |
| 244144 | USP35 | 107.286949 | -1.342738 | 0.17124803 | -7.840896 | 4.47E-15 |
| 208748 | PRRG3 | 142.671882 | -10.586315 | 1.35067741 | -7.8377819 | 4.59E-15 |
| 19878 | ROCK2 | 13350.3271 | 0.73931962 | 0.09435401 | 7.83559318 | 4.67E-15 |
| 320207 | PIK3R5 | 560.986383 | -4.5224168 | 0.57727591 | -7.8340647 | 4.72E-15 |
| 319636 | fsd1l | 127.072601 | -1.3495086 | 0.1722643 | -7.8339423 | 4.73E-15 |
| 12416 | CBX2 | 519.473247 | -1.4344283 | 0.18315737 | -7.8316711 | 4.81E-15 |
| 321006 | vprbp | 1821.29832 | 0.5417441 | 0.0691788 | 7.83107156 | 4.84E-15 |
| 70549 | TLN2 | 1329.63763 | 2.13495052 | 0.27266235 | 7.83001572 | 4.88E-15 |
| 270066 | SLC35E1 | 3280.40468 | 0.30709112 | 0.03922393 | 7.82917865 | 4.91E-15 |
| 107513 | SSR1 | 10962.8702 | 0.86104766 | 0.11000049 | 7.82767118 | 4.97E-15 |
| 68038 | CHID1 | 642.357348 | -0.9296512 | 0.11885192 | -7.8219283 | 5.20E-15 |
| 140740 | SEC63 | 4829.11161 | -0.3982747 | 0.05091888 | -7.8217484 | 5.21E-15 |
| 12293 | CACNA2D1 | 727.426677 | 2.58482098 | 0.33062313 | 7.81802823 | 5.37E-15 |
| 74202 | FBLIM1 | 2534.87212 | -1.7204691 | 0.22008684 | -7.8172283 | 5.40E-15 |
| 232157 | mob1a | 3601.19927 | 0.66881404 | 0.08556941 | 7.81604103 | 5.45E-15 |
| 269855 | SSC5D | 103.32095 | 3.83962077 | 0.4914253 | 7.81323374 | 5.57E-15 |
| 233315 | MTMR10 | 2568.19989 | -0.3577272 | 0.04579367 | -7.8117171 | 5.64E-15 |
| 233875 | INO80E | 1218.74131 | 1.10612111 | 0.14160178 | 7.81149132 | 5.65E-15 |
| 69882 | VWA9 | 817.479241 | 0.49858356 | 0.06384353 | 7.80946067 | 5.74E-15 |
| 19819 | RNASEH1 | 579.916534 | 0.8533081 | 0.10930552 | 7.80663342 | 5.87E-15 |
| 30938 | FGD3 | 660.098879 | -3.0398323 | 0.38940578 | -7.8063359 | 5.89E-15 |
| 21953 | TNNI2 | 88.5616579 | -7.8717881 | 1.00839805 | -7.806231 | 5.89E-15 |
| 73379 | DCBLD2 | 5779.66347 | -1.0627514 | 0.13614443 | -7.8060581 | 5.90E-15 |
| 80385 | TUSC2 | 309.212671 | -0.7082005 | 0.09074583 | -7.8042205 | 5.99E-15 |
| 227753 | GSN | 9140.22001 | -0.8350221 | 0.10703472 | -7.8014128 | 6.12E-15 |
| 17999 | NEDD4 | 35071.0363 | 0.37014229 | 0.04744689 | 7.80119124 | 6.13E-15 |
| 16834 | COG1 | 1301.27999 | 0.48341092 | 0.0620151 | 7.79505225 | 6.44E-15 |
| 78177 | NINL | 704.003801 | 0.7259843 | 0.09318407 | 7.7908626 | 6.66E-15 |

| | | | | | | |
|---|---|---|---|---|---|---|
| 68736 | TYW5 | 338.335102 | -0.6698342 | 0.08598285 | -7.7903231 | 6.68E-15 |
| 240087 | MDC1 | 2898.43124 | 0.51096194 | 0.06559872 | 7.78920571 | 6.74E-15 |
| 65100 | ZIC5 | 30.2787331 | 8.40007796 | 1.07861895 | 7.78780863 | 6.82E-15 |
| 213550 | DIS3L | 955.571978 | 0.65269987 | 0.0838661 | 7.78264289 | 7.10E-15 |
| 212390 | klhl32 | 40.0566185 | -8.7560323 | 1.12563168 | -7.7787721 | 7.32E-15 |
| 56473 | FADS2 | 3387.72969 | 0.56764152 | 0.07298431 | 7.77758304 | 7.39E-15 |
| 94185 | TNFRSF21 | 613.75393 | -3.1830936 | 0.40929405 | -7.7770336 | 7.42E-15 |
| 225997 | TRPM6 | 77.8375195 | -4.4030426 | 0.56647281 | -7.7727343 | 7.68E-15 |
| 67157 | 2610301B20 | 519.594034 | -1.184036 | 0.15234423 | -7.7721093 | 7.72E-15 |
| 59027 | NAMPT | 4357.63841 | 0.8964638 | 0.11535123 | 7.77160177 | 7.75E-15 |
| 30060 | mfi2 | 102.111017 | -9.3815739 | 1.20722565 | -7.7711851 | 7.78E-15 |
| 72183 | SNX6 | 2710.80631 | 0.44607304 | 0.05740938 | 7.77003809 | 7.85E-15 |
| 66488 | FAM136A | 798.34693 | 0.85824554 | 0.11053523 | 7.76445278 | 8.20E-15 |
| 27878 | TADA1 | 984.71558 | 1.13001514 | 0.14557753 | 7.76229103 | 8.34E-15 |
| 271564 | VPS13A | 3186.22462 | -0.777148 | 0.10014982 | -7.7598543 | 8.50E-15 |
| 217344 | RHBDF2 | 924.986165 | -4.3567437 | 0.56148543 | -7.7593175 | 8.54E-15 |
| 320184 | LRRC58 | 8586.67312 | 0.36740825 | 0.04735502 | 7.75859173 | 8.59E-15 |
| 100503043 | ARMCX4 | 1934.43646 | 1.3570701 | 0.17495198 | 7.75681483 | 8.71E-15 |
| 225363 | ETF1 | 13716.5748 | 0.22798492 | 0.02939569 | 7.75572546 | 8.78E-15 |
| 22137 | TTK | 1510.34457 | 0.54364523 | 0.07010025 | 7.75525415 | 8.82E-15 |
| 11826 | AQP1 | 286.595462 | 3.01540343 | 0.38905475 | 7.75058898 | 9.15E-15 |
| 27078 | B9D1 | 280.201162 | -1.0135081 | 0.13077773 | -7.7498523 | 9.20E-15 |
| 17386 | MMP13 | 1443.75464 | 4.10839679 | 0.53015837 | 7.74937652 | 9.23E-15 |
| 72205 | EML2 | 633.043028 | -2.4786912 | 0.3198886 | -7.7486076 | 9.29E-15 |
| 244550 | PODNL1 | 59.5335098 | 4.86515045 | 0.62794155 | 7.74777603 | 9.35E-15 |
| 72141 | ADPGK | 1704.23826 | 1.10018335 | 0.14208173 | 7.74331312 | 9.69E-15 |
| 18491 | PAPPA | 148.140179 | -7.1766215 | 0.92697711 | -7.7419619 | 9.79E-15 |
| 64899 | LPIN3 | 170.567045 | -2.0193797 | 0.26090242 | -7.7399806 | 9.94E-15 |
| 12662 | CHM | 1188.2283 | -0.5783332 | 0.07473766 | -7.7381774 | 1.01E-14 |
| 353190 | EDC3 | 930.52419 | 0.53213282 | 0.06878052 | 7.73667939 | 1.02E-14 |
| 60321 | WBP11 | 3933.42076 | 0.72772256 | 0.09412278 | 7.7316304 | 1.06E-14 |
| 20512 | SLC1A3 | 36.0637853 | 4.019954 | 0.51998765 | 7.73086443 | 1.07E-14 |
| 20603 | SMS | 407.965362 | 0.74229188 | 0.09604589 | 7.72851257 | 1.09E-14 |
| 109077 | INTS5 | 2091.188 | 0.50906352 | 0.06587422 | 7.72781105 | 1.09E-14 |
| 319974 | AUTS2 | 56.3213018 | -9.2446327 | 1.19635447 | -7.7273358 | 1.10E-14 |
| 67724 | POP1 | 2521.8024 | 1.52656138 | 0.1975662 | 7.72683467 | 1.10E-14 |
| 71345 | ANO9 | 38.7360157 | -8.7076732 | 1.12785783 | -7.7205415 | 1.16E-14 |
| 27015 | POLK | 1085.80143 | -0.7066602 | 0.09154657 | -7.7191333 | 1.17E-14 |
| 67488 | CALCOCO1 | 500.955662 | -1.5839639 | 0.20532782 | -7.7143169 | 1.22E-14 |
| 100042069 | Gm15210 | 1997.23934 | 0.8790847 | 0.11396065 | 7.71393212 | 1.22E-14 |
| 70103 | ZNHIT1 | 741.737894 | -0.5289971 | 0.06857828 | -7.713771 | 1.22E-14 |
| 101994 | CHAMP1 | 2065.71884 | 0.4701416 | 0.06097408 | 7.71051592 | 1.25E-14 |
| 382090 | CEP162 | 442.546693 | -0.6980872 | 0.09055608 | -7.7088935 | 1.27E-14 |
| 52858 | CDIPT | 2960.42428 | -0.5897176 | 0.07650407 | -7.7083158 | 1.27E-14 |
| 72747 | TTC39C | 438.964746 | 0.74580328 | 0.09676921 | 7.70703065 | 1.29E-14 |

| | | | | | | |
|---|---|---|---|---|---|---|
| 57339 | JPH1 | 77.4881502 | -9.7052198 | 1.25947395 | -7.7057725 | 1.30E-14 |
| 69743 | CASZ1 | 83.5519186 | -3.3153971 | 0.43029112 | -7.7050093 | 1.31E-14 |
| 16563 | KIF2A | 3022.85391 | 1.20675252 | 0.15664355 | 7.70381247 | 1.32E-14 |
| 64658 | MRPS25 | 1111.46935 | 0.63106431 | 0.08192387 | 7.70305822 | 1.33E-14 |
| 76650 | SRXN1 | 12510.9159 | 1.93913529 | 0.2518985 | 7.69808195 | 1.38E-14 |
| 230587 | GLIS1 | 77.1902117 | 2.67057064 | 0.34696975 | 7.69684006 | 1.39E-14 |
| 99681 | TCHH | 53.8328899 | -5.7174505 | 0.74296057 | -7.6954965 | 1.41E-14 |
| 73167 | ARHGAP8 | 84.6709986 | -7.05682 | 0.91700769 | -7.6954862 | 1.41E-14 |
| 98496 | PID1 | 241.119141 | -3.2458908 | 0.42185917 | -7.6942522 | 1.42E-14 |
| 75590 | DUSP9 | 1370.86959 | 2.11633847 | 0.27546854 | 7.6826866 | 1.56E-14 |
| 21973 | TOP2A | 19940.1602 | 0.45689621 | 0.05947163 | 7.68259066 | 1.56E-14 |
| 269016 | SH3RF2 | 31.2166552 | -8.3961633 | 1.09313417 | -7.6808168 | 1.58E-14 |
| 22040 | TREX1 | 186.528063 | 1.30507797 | 0.16993923 | 7.67967454 | 1.59E-14 |
| 15006 | H2-Q1 | 163.100376 | -1.2837211 | 0.16731104 | -7.6726621 | 1.68E-14 |
| 67857 | PPP6C | 1292.44809 | 0.49756374 | 0.06485548 | 7.67188473 | 1.69E-14 |
| 66736 | EMC2 | 1222.92084 | 1.2256175 | 0.15980663 | 7.66937826 | 1.73E-14 |
| 67993 | NUDT12 | 251.764836 | -1.5422212 | 0.20113577 | -7.6675632 | 1.75E-14 |
| 22629 | YWHAH | 3609.2774 | -0.3398981 | 0.04435458 | -7.6632019 | 1.81E-14 |
| 67629 | SPC24 | 648.018901 | 0.84512881 | 0.11028782 | 7.66293849 | 1.82E-14 |
| 14373 | G0S2 | 85.2548586 | -9.1214358 | 1.19055092 | -7.6615252 | 1.84E-14 |
| 171395 | PKD1L1 | 35.8156607 | -6.1362762 | 0.80119526 | -7.6589023 | 1.88E-14 |
| 664883 | NOVA1 | 121.819224 | 4.44784164 | 0.5810565 | 7.654749 | 1.94E-14 |
| 235627 | NBEAL2 | 615.721645 | -3.0370167 | 0.39697982 | -7.650305 | 2.01E-14 |
| 13728 | MARK2 | 2863.77694 | -0.365146 | 0.04774089 | -7.6484955 | 2.03E-14 |
| 14707 | GNG5 | 436.067656 | -0.6955856 | 0.0909472 | -7.6482359 | 2.04E-14 |
| 269328 | MUC15 | 39.8756074 | -8.0204953 | 1.04883968 | -7.6470174 | 2.06E-14 |
| 545554 | ANKRD34A | 89.5080916 | 3.6791851 | 0.48127258 | 7.64470135 | 2.09E-14 |
| 21770 | PPP2R5D | 2690.13342 | -0.6535256 | 0.08551115 | -7.6425778 | 2.13E-14 |
| 382038 | URB2 | 2888.26563 | 0.3629517 | 0.04750689 | 7.63998007 | 2.17E-14 |
| 215446 | entpd3 | 26.990793 | -8.1827812 | 1.07123983 | -7.6386081 | 2.20E-14 |
| 20355 | SEMA4F | 1279.21049 | -6.4952655 | 0.85043054 | -7.6376203 | 2.21E-14 |
| 66366 | ERGIC3 | 2041.64643 | -0.7934822 | 0.10401238 | -7.6287284 | 2.37E-14 |
| 224674 | SLC37A1 | 39.861122 | -6.7191025 | 0.88089251 | -7.6276077 | 2.39E-14 |
| 102371 | MYZAP | 138.103027 | -2.6135779 | 0.34269526 | -7.6265363 | 2.41E-14 |
| 14263 | FMO5 | 33.5476144 | -2.6255833 | 0.34433072 | -7.6251788 | 2.44E-14 |
| 12660 | CHKA | 1844.46704 | -1.5017117 | 0.19694607 | -7.6249896 | 2.44E-14 |
| 72102 | DUSP11 | 4215.73477 | 0.69995955 | 0.09180009 | 7.62482399 | 2.44E-14 |
| 22770 | ZHX1 | 1504.39365 | 1.35556532 | 0.17778962 | 7.62454706 | 2.45E-14 |
| 19329 | RAB17 | 67.2508735 | -8.7786135 | 1.1515989 | -7.6229783 | 2.48E-14 |
| 53328 | PGRMC1 | 5773.47966 | 0.56020422 | 0.07349371 | 7.62247823 | 2.49E-14 |
| 94088 | TRIM6 | 66.2737684 | -1.9952905 | 0.26184453 | -7.6201345 | 2.53E-14 |
| 22688 | Zfp26 | 1164.55663 | 0.51714616 | 0.06789576 | 7.61676671 | 2.60E-14 |
| 68260 | TRMT12 | 339.98557 | 0.86210706 | 0.11318932 | 7.61650512 | 2.61E-14 |
| 320299 | IQCB1 | 324.239727 | -0.6382523 | 0.08380125 | -7.6162625 | 2.61E-14 |
| 320508 | CACHD1 | 721.17799 | 3.21223553 | 0.42206 | 7.61085041 | 2.72E-14 |

| | | | | | | |
|---|---|---|---|---|---|---|
| 96935 | SUSD4 | 87.2002058 | -8.4379791 | 1.10978622 | -7.6032473 | 2.89E-14 |
| 12810 | COCH | 55.1068646 | 7.22380219 | 0.9502334 | 7.60213457 | 2.91E-14 |
| 271887 | RBBP8NL | 26.4269062 | -8.1570863 | 1.07346102 | -7.5988659 | 2.99E-14 |
| 22321 | VARS | 7133.39592 | 0.43969949 | 0.05786434 | 7.59879944 | 2.99E-14 |
| 74127 | KRT80 | 341.296872 | -2.2498975 | 0.29617158 | -7.5966015 | 3.04E-14 |
| 18938 | PPP1R14B | 1507.81869 | 0.46754944 | 0.061582 | 7.5923074 | 3.14E-14 |
| 68047 | MPND | 950.068047 | -0.550005 | 0.07249876 | -7.5864055 | 3.29E-14 |
| 226777 | C130074G19 | 70.9996946 | -4.7883667 | 0.63125568 | -7.5854632 | 3.31E-14 |
| 74330 | DNAJC14 | 2362.89283 | -0.3205653 | 0.04226219 | -7.585155 | 3.32E-14 |
| 20586 | SMARCA4 | 7978.36365 | 0.48115857 | 0.0634553 | 7.5826382 | 3.39E-14 |
| 230648 | EFCAB14 | 1373.3247 | 0.58501626 | 0.07716669 | 7.58120206 | 3.42E-14 |
| 74841 | USP38 | 1860.39846 | 0.43270651 | 0.05707786 | 7.58098652 | 3.43E-14 |
| 71780 | ISYNA1 | 720.600237 | -4.435414 | 0.58532941 | -7.5776373 | 3.52E-14 |
| 18223 | NUMBL | 1356.76114 | 1.29901677 | 0.17142791 | 7.57762701 | 3.52E-14 |
| 21808 | TGFB2 | 687.260286 | -4.73832 | 0.62555833 | -7.5745454 | 3.60E-14 |
| 22272 | Uqcrq | 2156.70442 | -0.4495814 | 0.059362 | -7.5735558 | 3.63E-14 |
| 100041231 | Gm3219 | 131.939771 | 1.18566192 | 0.15655506 | 7.5734499 | 3.63E-14 |
| 213499 | FBXO42 | 1414.60017 | 0.7970249 | 0.10528902 | 7.56987633 | 3.74E-14 |
| 22421 | WNT7A | 1301.28561 | -13.775792 | 1.81985941 | -7.5697013 | 3.74E-14 |
| 109711 | ACTN1 | 30752.2549 | -0.3405961 | 0.04500495 | -7.5679691 | 3.79E-14 |
| 57435 | PLIN4 | 192.362425 | 6.02853437 | 0.79661927 | 7.56764819 | 3.80E-14 |
| 16412 | ITGB1 | 44994.535 | -0.8535331 | 0.11283217 | -7.5646257 | 3.89E-14 |
| 83410 | CSTF2T | 858.934728 | -0.4521003 | 0.0597679 | -7.5642662 | 3.90E-14 |
| 75015 | 4930503B20 | 25.1731392 | -8.0817016 | 1.06871171 | -7.562097 | 3.97E-14 |
| 74035 | NOL9 | 2791.73547 | 1.09037003 | 0.14419517 | 7.56176507 | 3.98E-14 |
| 18591 | PDGFB | 117.984043 | -8.3054941 | 1.09850767 | -7.5607066 | 4.01E-14 |
| 53890 | SART3 | 2471.4074 | 0.32952064 | 0.04358519 | 7.56038168 | 4.02E-14 |
| 22210 | UBE2B | 2234.38719 | -0.690564 | 0.09140522 | -7.5549735 | 4.19E-14 |
| 278240 | Spin2c | 25.0002419 | -8.0726017 | 1.06864502 | -7.5540536 | 4.22E-14 |
| 234699 | EDC4 | 2701.31723 | 0.37881548 | 0.05014733 | 7.55405034 | 4.22E-14 |
| 320165 | TACC1 | 1227.78732 | 3.28939791 | 0.43560628 | 7.55130972 | 4.31E-14 |
| 216131 | TRAPPC10 | 2760.80275 | -0.4328491 | 0.05736241 | -7.5458663 | 4.49E-14 |
| 11733 | ANK1 | 35.1620295 | -3.232474 | 0.42864564 | -7.5411335 | 4.66E-14 |
| 232714 | MGAM | 191.053733 | -4.4769749 | 0.59386098 | -7.5387592 | 4.74E-14 |
| 229055 | ZBTB10 | 490.000375 | 0.91600934 | 0.12152457 | 7.53764746 | 4.79E-14 |
| 613262 | BC029722 | 801.897001 | -0.9477269 | 0.12574414 | -7.5369473 | 4.81E-14 |
| 229600 | BC028528 | 37.9774171 | 8.00284471 | 1.06184995 | 7.53670017 | 4.82E-14 |
| 79362 | BHLHE41 | 252.814784 | -2.6244061 | 0.34831933 | -7.5344831 | 4.90E-14 |
| 74025 | NPHP3 | 203.05598 | 1.11492637 | 0.14798476 | 7.53406192 | 4.92E-14 |
| 407819 | BC031181 | 707.282574 | -0.6694895 | 0.08887905 | -7.5325899 | 4.97E-14 |
| 109168 | ATL3 | 8133.52595 | -0.813142 | 0.10802754 | -7.5271731 | 5.19E-14 |
| 78581 | UTP23 | 740.091612 | 1.01181796 | 0.13450613 | 7.52246717 | 5.38E-14 |
| 243983 | ZDHHC13 | 866.611189 | -0.7571466 | 0.10065549 | -7.5221587 | 5.39E-14 |
| 99512 | WDR47 | 544.938453 | -0.6452489 | 0.08578136 | -7.5220178 | 5.39E-14 |
| 19273 | PTPRU | 32.1474456 | 2.87263733 | 0.38206646 | 7.51868495 | 5.53E-14 |

| | | | | | | |
|---|---|---|---|---|---|---|
| 105827 | AMIGO2 | 589.204812 | 4.83755179 | 0.64391104 | 7.51276415 | 5.79E-14 |
| 18158 | nppb | 150.973266 | 2.32341875 | 0.30932828 | 7.5111747 | 5.86E-14 |
| 241308 | RALGPS1 | 72.0553457 | -3.1790121 | 0.42333041 | -7.5095294 | 5.93E-14 |
| 64451 | DIP2A | 2157.48927 | -0.5592969 | 0.07448442 | -7.5089111 | 5.96E-14 |
| 17698 | MSN | 47334.338 | -0.3103664 | 0.04133479 | -7.5085992 | 5.98E-14 |
| 76380 | CEP112 | 65.3683381 | -1.7892622 | 0.23836995 | -7.5062408 | 6.08E-14 |
| 11535 | ADM | 1555.95628 | 3.89870319 | 0.51972277 | 7.5015055 | 6.31E-14 |
| 333424 | A4GNT | 23.065058 | 8.00916921 | 1.06768404 | 7.50144134 | 6.31E-14 |
| 226418 | YOD1 | 433.195617 | -0.8366853 | 0.11154582 | -7.5008218 | 6.34E-14 |
| 20666 | SOX11 | 372.20052 | -11.969942 | 1.59731723 | -7.4937786 | 6.69E-14 |
| 319888 | oacyl | 30.1715743 | -4.5354762 | 0.60530241 | -7.4929095 | 6.74E-14 |
| 226250 | AFAP1L2 | 143.991319 | -2.6197117 | 0.34962649 | -7.4928865 | 6.74E-14 |
| 665113 | TNIK | 98.8710091 | -8.6061545 | 1.14880819 | -7.4913764 | 6.82E-14 |
| 110446 | ACAT1 | 1264.38457 | 0.86688436 | 0.11573817 | 7.4900474 | 6.88E-14 |
| 14421 | B4GALNT1 | 1927.79987 | -0.6221491 | 0.08307001 | -7.4894552 | 6.92E-14 |
| 71726 | SMUG1 | 492.243148 | -1.0478456 | 0.13994524 | -7.4875405 | 7.02E-14 |
| 17454 | MOV10 | 590.778784 | -1.6051068 | 0.2144784 | -7.4837687 | 7.22E-14 |
| 234582 | CCDC102A | 531.3858 | 1.61096001 | 0.21543815 | 7.47759864 | 7.57E-14 |
| 104001 | RTN1 | 42.1603963 | -6.7974973 | 0.90927445 | -7.4757377 | 7.68E-14 |
| 13202 | DDT | 301.94368 | -0.8595857 | 0.11503041 | -7.4726823 | 7.86E-14 |
| 74412 | GLE1 | 1526.81531 | 0.44139495 | 0.0590679 | 7.47267088 | 7.86E-14 |
| 269608 | PLEKHG5 | 489.053844 | 1.01210047 | 0.13545458 | 7.47188075 | 7.91E-14 |
| 16468 | JARID2 | 391.609823 | 1.06915636 | 0.14309519 | 7.47164444 | 7.92E-14 |
| 233912 | ARMC5 | 1182.78915 | 1.31188464 | 0.17558906 | 7.47133472 | 7.94E-14 |
| 257871 | Olfr1372-ps1 | 38.7509339 | -5.2547311 | 0.70337502 | -7.4707389 | 7.97E-14 |
| 16907 | LMNB2 | 1610.79037 | 0.61218488 | 0.08206722 | 7.45955421 | 8.68E-14 |
| 22368 | TRPV2 | 336.345829 | 3.762745 | 0.50446782 | 7.45884051 | 8.73E-14 |
| 18074 | NID2 | 24.8044134 | 5.0457588 | 0.67657902 | 7.45775236 | 8.80E-14 |
| 12572 | CDK7 | 1297.6361 | 0.36575984 | 0.04904433 | 7.45773883 | 8.80E-14 |
| 67678 | LSM3 | 1150.16174 | 0.9642068 | 0.1292969 | 7.45730818 | 8.83E-14 |
| 14367 | FZD5 | 1288.91802 | -1.137558 | 0.15257997 | -7.4554871 | 8.95E-14 |
| 106369 | YPEL1 | 36.0916793 | -2.8953597 | 0.38859049 | -7.4509276 | 9.27E-14 |
| 56392 | SHOC2 | 2630.07209 | -0.5247036 | 0.07042287 | -7.4507563 | 9.28E-14 |
| 12927 | BCAR1 | 5036.17636 | -0.6822286 | 0.09157194 | -7.4501926 | 9.32E-14 |
| 235661 | DYNC1LI1 | 2185.74995 | 0.45763673 | 0.06144867 | 7.44746348 | 9.52E-14 |
| 11931 | ATP1B1 | 1678.11339 | -4.4552015 | 0.59840783 | -7.4450923 | 9.69E-14 |
| 74111 | RBM19 | 2330.17717 | 0.48023532 | 0.06451572 | 7.44369419 | 9.79E-14 |
| 19663 | RBPMS | 786.773764 | -1.6916217 | 0.22728411 | -7.4427624 | 9.86E-14 |
| 11764 | AP1B1 | 2619.99472 | -0.5298945 | 0.07122522 | -7.4397034 | 1.01E-13 |
| 319740 | ZFYVE27 | 875.377475 | -0.8845763 | 0.11891855 | -7.4385053 | 1.02E-13 |
| 70435 | INF2 | 10741.1486 | 1.03610632 | 0.13929223 | 7.43836418 | 1.02E-13 |
| 237823 | PFAS | 3572.82329 | 0.60550472 | 0.08140455 | 7.43821746 | 1.02E-13 |
| 71893 | NOXO1 | 150.337924 | -4.2800973 | 0.57549884 | -7.4371954 | 1.03E-13 |
| 233016 | BLVRB | 887.202022 | 1.70430656 | 0.2291978 | 7.43596403 | 1.04E-13 |
| 22685 | Zfp239 | 315.894266 | 0.72298033 | 0.09726751 | 7.43290641 | 1.06E-13 |

| | | | | | | |
|---|---|---|---|---|---|---|
| 104080 | NXPH4 | 26.130473 | 3.21248063 | 0.43242548 | 7.42898092 | 1.09E-13 |
| 100087 | KTI12 | 1052.8512 | 1.12156315 | 0.15098112 | 7.42849923 | 1.10E-13 |
| 75599 | PCDH1 | 1232.92342 | -1.3773697 | 0.18542767 | -7.4280699 | 1.10E-13 |
| 65255 | ASB4 | 22.2911379 | -7.9070759 | 1.06449831 | -7.4279835 | 1.10E-13 |
| 100049161 | Gm13032 | 23.8428022 | -3.3382696 | 0.45014056 | -7.4160603 | 1.21E-13 |
| 53603 | TSLP | 36.3560144 | -5.5495782 | 0.74888916 | -7.4104133 | 1.26E-13 |
| 239796 | MB21D2 | 757.364688 | 1.07276962 | 0.14478588 | 7.40935252 | 1.27E-13 |
| 434484 | SP140 | 59.9367951 | -2.1207274 | 0.28626996 | -7.4081383 | 1.28E-13 |
| 20390 | SFTPD | 30.1312129 | -8.3458799 | 1.1268597 | -7.4063168 | 1.30E-13 |
| 110279 | BCR | 2593.25146 | -0.4995494 | 0.06745633 | -7.4055229 | 1.31E-13 |
| 74477 | 4933427D14 | 568.186321 | -0.5768103 | 0.07790896 | -7.4036459 | 1.32E-13 |
| 218232 | PTPDC1 | 530.37677 | 1.06491613 | 0.14391338 | 7.39970215 | 1.36E-13 |
| 55994 | SMAD9 | 524.231295 | 2.92238948 | 0.39511753 | 7.39625374 | 1.40E-13 |
| 66083 | SETD6 | 457.769976 | -0.651527 | 0.08809332 | -7.3958725 | 1.40E-13 |
| 381979 | BRSK1 | 191.624584 | 2.77523264 | 0.3753772 | 7.3931839 | 1.43E-13 |
| 20741 | SPTB | 30.7057536 | -5.9059249 | 0.79898844 | -7.3917527 | 1.45E-13 |
| 433926 | LRRC8B | 737.966643 | -0.9566916 | 0.12943269 | -7.3914215 | 1.45E-13 |
| 70122 | MLLT3 | 1059.95027 | 1.11700623 | 0.15113059 | 7.39100031 | 1.46E-13 |
| 216831 | ARHGAP44 | 750.232576 | -2.0977989 | 0.28385162 | -7.390477 | 1.46E-13 |
| 71361 | AIFM2 | 446.58519 | -1.1164889 | 0.15107973 | -7.3900645 | 1.47E-13 |
| 223922 | ATF7 | 499.03078 | -0.8325464 | 0.11271227 | -7.3864757 | 1.51E-13 |
| 232970 | PHLDB3 | 189.679171 | -0.9748361 | 0.13209163 | -7.379999 | 1.58E-13 |
| 66549 | AGGF1 | 1908.06255 | 0.59128447 | 0.08011992 | 7.3799931 | 1.58E-13 |
| 100042856 | Gm4070 | 21.5308247 | -7.8572202 | 1.06528078 | -7.375727 | 1.63E-13 |
| 17758 | MAP4 | 10015.3256 | 0.7117831 | 0.09651065 | 7.3751765 | 1.64E-13 |
| 67718 | Lce1h | 28.7181342 | -8.2717197 | 1.12212359 | -7.3714872 | 1.69E-13 |
| 66291 | SMIM8 | 122.172258 | -1.7299201 | 0.23468386 | -7.3712786 | 1.69E-13 |
| 53881 | SLC5A3 | 6293.97626 | 0.54839542 | 0.07442485 | 7.36844497 | 1.73E-13 |
| 20289 | SCX | 360.568974 | -2.7889387 | 0.3785371 | -7.3676761 | 1.74E-13 |
| 13549 | DYRK1B | 130.834642 | -1.4532912 | 0.19726759 | -7.3671057 | 1.74E-13 |
| 17161 | MAOA | 4768.15414 | -1.587121 | 0.21546081 | -7.3661699 | 1.76E-13 |
| 235402 | LINGO1 | 100.531049 | 3.39368651 | 0.46084987 | 7.36397412 | 1.79E-13 |
| 68149 | OTUB2 | 160.758518 | 1.25237855 | 0.17019064 | 7.3586805 | 1.86E-13 |
| 108672 | ZDHHC15 | 36.3388897 | -8.6154426 | 1.17081126 | -7.3585238 | 1.86E-13 |
| 65112 | PMEPA1 | 7673.39116 | -1.3402535 | 0.18222854 | -7.3547946 | 1.91E-13 |
| 64933 | AP3M2 | 222.445626 | -1.4721973 | 0.20016962 | -7.3547489 | 1.91E-13 |
| 16635 | Klra4 | 704.097492 | -2.0129763 | 0.27370472 | -7.3545545 | 1.92E-13 |
| 94221 | GOPC | 2265.49432 | -0.6374 | 0.08667577 | -7.3538424 | 1.93E-13 |
| 72324 | PLXDC1 | 125.660732 | 3.72911624 | 0.50718875 | 7.35252162 | 1.95E-13 |
| 21892 | TLL1 | 33.0528129 | -4.8128416 | 0.65461008 | -7.3522265 | 1.95E-13 |
| 100504230 | AU020206 | 566.271757 | 1.14004345 | 0.15506447 | 7.35206083 | 1.95E-13 |
| 231452 | SDAD1 | 2188.81629 | 0.43661264 | 0.05941399 | 7.34865059 | 2.00E-13 |
| 15468 | PRMT2 | 696.70853 | -1.0985666 | 0.149509 | -7.347829 | 2.01E-13 |
| 24102 | TREX2 | 63.8349611 | -9.4271917 | 1.28310334 | -7.3471804 | 2.02E-13 |
| 73094 | SGIP1 | 43.0820839 | 6.0925752 | 0.82961792 | 7.34383266 | 2.08E-13 |

| | | | | | | |
|---|---|---|---|---|---|---|
| 245631 | MUM1L1 | 696.661534 | 1.57877208 | 0.21498093 | 7.34377723 | 2.08E-13 |
| 77652 | Zfp955a | 543.015443 | 1.08774846 | 0.14813845 | 7.34278276 | 2.09E-13 |
| 27225 | DDX24 | 4872.1912 | 0.54551205 | 0.07432834 | 7.33921966 | 2.15E-13 |
| 381148 | PROB1 | 84.1487075 | -2.2229274 | 0.30290989 | -7.3385767 | 2.16E-13 |
| 73945 | OTUD4 | 9956.78752 | 0.55859214 | 0.07614349 | 7.33604609 | 2.20E-13 |
| 18015 | NF1 | 2364.97556 | -1.2089811 | 0.16482714 | -7.3348422 | 2.22E-13 |
| 83486 | RBM5 | 1997.92565 | 0.3944781 | 0.05378387 | 7.3345051 | 2.23E-13 |
| 100503546 | Gm15958 | 21.6145166 | -7.8650494 | 1.07308489 | -7.3293823 | 2.31E-13 |
| 20471 | SIX1 | 715.916144 | 2.86507537 | 0.39095387 | 7.32842315 | 2.33E-13 |
| 69071 | TMEM97 | 1121.32274 | 0.77202714 | 0.10535365 | 7.32795853 | 2.34E-13 |
| 18037 | NFKBIE | 402.004844 | -1.4558965 | 0.19872105 | -7.3263327 | 2.37E-13 |
| 238386 | BTBD7 | 2235.53875 | 0.66958222 | 0.09141292 | 7.32480968 | 2.39E-13 |
| 69632 | ARHGEF12 | 5624.56818 | 0.59210496 | 0.08083723 | 7.32465668 | 2.40E-13 |
| 102545 | CMTM7 | 736.189074 | 0.87763389 | 0.11986667 | 7.32175079 | 2.45E-13 |
| 380928 | LMO7 | 1034.58454 | 1.86495018 | 0.25495942 | 7.31469402 | 2.58E-13 |
| 17703 | Msx3 | 40.6187009 | 5.50965102 | 0.75327419 | 7.31427025 | 2.59E-13 |
| 56384 | LETM1 | 2313.45953 | -0.519348 | 0.071013 | -7.3134212 | 2.60E-13 |
| 230734 | YRDC | 1311.51729 | 1.00291656 | 0.13714687 | 7.31271928 | 2.62E-13 |
| 11920 | ATM | 2735.21354 | 0.71811753 | 0.09822851 | 7.31068337 | 2.66E-13 |
| 74320 | WDR33 | 2863.08899 | 0.34644014 | 0.04742574 | 7.3048973 | 2.77E-13 |
| 22319 | VAMP3 | 1896.16362 | 0.98849769 | 0.1353399 | 7.30381594 | 2.80E-13 |
| 74102 | SLC35A5 | 452.318901 | -0.7544753 | 0.10334063 | -7.300858 | 2.86E-13 |
| 210274 | SHANK2 | 28.2306552 | -4.4445134 | 0.60886867 | -7.2996257 | 2.89E-13 |
| 108903 | TBCD | 2805.66047 | -0.4391436 | 0.06018203 | -7.2969222 | 2.94E-13 |
| 319954 | Egfros | 32.193518 | -7.7107222 | 1.05699449 | -7.2949502 | 2.99E-13 |
| 237887 | Slfn10-ps | 69.357366 | -7.0636506 | 0.96833812 | -7.2946118 | 3.00E-13 |
| 16923 | SH2B3 | 1482.33799 | 1.28665061 | 0.17639246 | 7.29424947 | 3.00E-13 |
| 234875 | TTC13 | 2300.54185 | -0.6716415 | 0.09212416 | -7.2906121 | 3.09E-13 |
| 56441 | NAT6 | 415.734421 | -0.7447145 | 0.10218722 | -7.2877461 | 3.15E-13 |
| 18810 | PLEC | 42545.299 | 0.50025842 | 0.06864512 | 7.28760315 | 3.16E-13 |
| 237860 | SSH2 | 1210.50078 | -0.7648593 | 0.10498612 | -7.285338 | 3.21E-13 |
| 19713 | RET | 30.7290555 | -6.3284533 | 0.86873169 | -7.2847041 | 3.22E-13 |
| 13388 | DLL1 | 116.245814 | -4.5089219 | 0.61901182 | -7.2840642 | 3.24E-13 |
| 21376 | TBRG1 | 3341.49967 | 0.38332856 | 0.05266857 | 7.27812785 | 3.38E-13 |
| 378462 | MORN2 | 64.1624804 | -2.4485025 | 0.33646247 | -7.2771936 | 3.41E-13 |
| 14789 | P3H3 | 1136.8737 | 3.61168244 | 0.49639 | 7.27589687 | 3.44E-13 |
| 15461 | HRAS | 2345.56195 | -0.4576213 | 0.0629073 | -7.2745345 | 3.48E-13 |
| 17978 | NCOA2 | 1955.09496 | -0.5469372 | 0.07522416 | -7.2707654 | 3.57E-13 |
| 80290 | GPR146 | 187.021441 | -4.4004963 | 0.60529315 | -7.270025 | 3.59E-13 |
| 56055 | GTPBP2 | 871.930335 | -0.7859145 | 0.10811996 | -7.2689118 | 3.62E-13 |
| 68026 | 2810417H13 | 1845.04682 | 0.91602011 | 0.12602467 | 7.26857777 | 3.63E-13 |
| 78885 | CORO7 | 524.687491 | -1.3837034 | 0.19037379 | -7.2683504 | 3.64E-13 |
| 71943 | TOM1L1 | 668.953552 | -1.8734071 | 0.25783081 | -7.2660326 | 3.70E-13 |
| 224997 | DLGAP1 | 24.8612374 | -3.9484844 | 0.5435087 | -7.2648044 | 3.74E-13 |
| 93886 | PCDHB15 | 30.3339405 | 4.07249552 | 0.56117267 | 7.25711668 | 3.95E-13 |

| | | | | | | |
|---|---|---|---|---|---|---|
| 245049 | MYRIP | 67.5470765 | -9.5071821 | 1.31024415 | -7.2560386 | 3.99E-13 |
| 80286 | TUSC3 | 1531.49684 | -0.5721421 | 0.07886023 | -7.2551404 | 4.01E-13 |
| 15925 | IDE | 5184.73175 | -0.5264307 | 0.0725619 | -7.2549191 | 4.02E-13 |
| 67891 | RPL4 | 23631.4098 | 0.36181621 | 0.04989 | 7.25227886 | 4.10E-13 |
| 100732 | MAPRE3 | 404.860717 | -1.6321232 | 0.22511235 | -7.2502607 | 4.16E-13 |
| 214639 | 4930486L24I | 607.870356 | -4.0451479 | 0.55815911 | -7.2473024 | 4.25E-13 |
| 13663 | EI24 | 2599.95546 | -0.4434133 | 0.06119076 | -7.2464083 | 4.28E-13 |
| 228913 | Zfp217 | 928.79719 | -0.4878146 | 0.0673186 | -7.2463559 | 4.28E-13 |
| 110959 | NUDT19 | 539.475154 | 1.09351887 | 0.15091522 | 7.24591501 | 4.30E-13 |
| 545471 | Zfp345 | 22.0940797 | 7.94655528 | 1.09742837 | 7.24106964 | 4.45E-13 |
| 70727 | RASGEF1A | 29.6746959 | -4.6645932 | 0.64427562 | -7.2400584 | 4.48E-13 |
| 108812 | ALS2CR12 | 31.177691 | -3.0976382 | 0.42787951 | -7.2395104 | 4.50E-13 |
| 110173 | MANBA | 574.880521 | -1.5396234 | 0.21272549 | -7.2376062 | 4.57E-13 |
| 70484 | SLC35D2 | 138.450942 | -1.1861107 | 0.16388719 | -7.2373611 | 4.57E-13 |
| 69884 | 2010300F17I | 80.3090626 | -3.4712872 | 0.47977886 | -7.2351818 | 4.65E-13 |
| 230767 | IQCC | 172.851455 | 1.22389872 | 0.16919986 | 7.23344985 | 4.71E-13 |
| 70546 | ZDHHC2 | 592.57787 | -1.8132495 | 0.25078052 | -7.2304238 | 4.81E-13 |
| 11737 | ANP32A | 2069.24877 | 0.796101 | 0.1101074 | 7.23022221 | 4.82E-13 |
| 71007 | D7Ertd443e | 35.54302 | -8.5800483 | 1.18697084 | -7.2285249 | 4.88E-13 |
| 19649 | ROBO3 | 24.8038194 | -5.6072596 | 0.77571421 | -7.2285122 | 4.88E-13 |
| 215335 | SLC36A1 | 2122.1688 | -1.5813277 | 0.21886667 | -7.2250731 | 5.01E-13 |
| 78287 | RBSN | 1830.70365 | 0.83854988 | 0.11611494 | 7.22172285 | 5.13E-13 |
| 213696 | DUOXA1 | 94.0124245 | -3.6951118 | 0.51178742 | -7.220013 | 5.20E-13 |
| 12801 | CNR1 | 666.246677 | -4.6864012 | 0.64914368 | -7.2193589 | 5.22E-13 |
| 20917 | SUCLG2 | 881.165313 | 0.93481145 | 0.12949647 | 7.21881803 | 5.24E-13 |
| 20587 | SMARCB1 | 2419.56114 | 0.39515249 | 0.05477506 | 7.21409564 | 5.43E-13 |
| 67812 | UBXN4 | 4004.22541 | -0.3400692 | 0.04716749 | -7.2098212 | 5.60E-13 |
| 230761 | Zfp362 | 578.274254 | 0.79406025 | 0.11013652 | 7.20978124 | 5.60E-13 |
| 76365 | TBX18 | 725.576972 | 7.53212495 | 1.04488199 | 7.20858909 | 5.65E-13 |
| 77996 | Cutal | 58.02454 | -8.5654641 | 1.18911726 | -7.2032124 | 5.88E-13 |
| 22025 | NR2C1 | 711.054218 | -0.7695732 | 0.10685501 | -7.2020317 | 5.93E-13 |
| 67922 | FAM32A | 1677.1488 | -0.3670205 | 0.05096931 | -7.2008134 | 5.99E-13 |
| 71738 | MAMDC2 | 56.3280119 | -6.7959617 | 0.9439651 | -7.1993781 | 6.05E-13 |
| 16917 | LMX1B | 114.954747 | -8.2240525 | 1.14257394 | -7.1978296 | 6.12E-13 |
| 320554 | TCP11L1 | 896.981292 | -3.4346604 | 0.47719489 | -7.1976051 | 6.13E-13 |
| 11520 | PLIN2 | 5170.66752 | -1.182789 | 0.1643736 | -7.1957355 | 6.21E-13 |
| 83674 | CNNM1 | 46.133745 | 5.16134998 | 0.71756829 | 7.19283457 | 6.35E-13 |
| 19349 | Rab7 | 1922.42403 | 0.42951138 | 0.05973564 | 7.19020277 | 6.47E-13 |
| 74016 | PHF19 | 690.107511 | 2.70556042 | 0.37661541 | 7.18388145 | 6.78E-13 |
| 80837 | RHOJ | 845.733302 | 4.52601158 | 0.63004269 | 7.18365863 | 6.79E-13 |
| 13806 | ENO1 | 2962.86137 | 0.99285405 | 0.13822284 | 7.18299554 | 6.82E-13 |
| 66953 | CDCA7 | 846.477753 | 0.67695478 | 0.09429826 | 7.17886824 | 7.03E-13 |
| 56505 | RUVBL1 | 476.763962 | 0.88043245 | 0.12264286 | 7.17883148 | 7.03E-13 |
| 11538 | ADNP | 4616.27312 | -0.3459003 | 0.04818433 | -7.1786888 | 7.04E-13 |
| 16549 | KHSRP | 4510.90809 | -0.4649832 | 0.06480881 | -7.1746918 | 7.25E-13 |

| | | | | | | |
|---|---|---|---|---|---|---|
| 60527 | FADS3 | 3990.15948 | -1.0758939 | 0.1499639 | -7.1743527 | 7.27E-13 |
| 23885 | GMCL1 | 888.679452 | 0.50225421 | 0.07002039 | 7.17297102 | 7.34E-13 |
| 20779 | SRC | 3037.87331 | -1.1475661 | 0.16000928 | -7.1718721 | 7.40E-13 |
| 671535 | PARP10 | 744.808545 | -1.7533596 | 0.24463746 | -7.1671754 | 7.66E-13 |
| 68501 | NSMCE2 | 982.802812 | 0.57304055 | 0.07995488 | 7.16704902 | 7.66E-13 |
| 235323 | USP28 | 632.262581 | 0.82641883 | 0.11530851 | 7.16702394 | 7.66E-13 |
| 69181 | DYRK2 | 662.437984 | 0.82478572 | 0.11510588 | 7.16545222 | 7.75E-13 |
| 18718 | PIP4K2A | 900.103672 | 0.68916701 | 0.09618827 | 7.16477174 | 7.79E-13 |
| 229357 | GPR149 | 671.238584 | 7.31759539 | 1.02160314 | 7.16285526 | 7.90E-13 |
| 12224 | KLF5 | 1176.00744 | -1.3492661 | 0.18845293 | -7.1596981 | 8.09E-13 |
| 72058 | IGSF5 | 29.0051436 | -8.2865476 | 1.15813399 | -7.1550854 | 8.36E-13 |
| 27061 | BCAP31 | 3302.4951 | -0.4739916 | 0.06625327 | -7.1542365 | 8.41E-13 |
| 69094 | TMEM160 | 458.166605 | 0.85054054 | 0.11888661 | 7.15421662 | 8.42E-13 |
| 14148 | FDX1 | 572.694794 | -0.9237129 | 0.12913541 | -7.1530565 | 8.49E-13 |
| 67075 | MAGT1 | 2740.48066 | -0.2768021 | 0.03871472 | -7.1497897 | 8.69E-13 |
| 215051 | BUD13 | 512.151448 | 0.78136279 | 0.10929453 | 7.14914845 | 8.73E-13 |
| 15547 | TRMT2A | 1328.72769 | -0.366863 | 0.05132103 | -7.1483947 | 8.78E-13 |
| 67273 | NDUFA10 | 2378.3813 | -0.4757027 | 0.06656857 | -7.1460554 | 8.93E-13 |
| 74104 | ABCB6 | 592.937152 | 1.05912609 | 0.14821453 | 7.14589935 | 8.94E-13 |
| 67890 | UFM1 | 1391.71058 | -0.4396427 | 0.06152812 | -7.1453953 | 8.97E-13 |
| 19744 | RHEB | 3196.18415 | -0.3431027 | 0.04802088 | -7.1448649 | 9.01E-13 |
| 100088 | RCC1 | 3576.42968 | 1.53306961 | 0.21464962 | 7.14219578 | 9.19E-13 |
| 67040 | DDX17 | 10645.3279 | -0.8838012 | 0.12377773 | -7.1402278 | 9.32E-13 |
| 107771 | Bmyc | 22.7806865 | -7.9428519 | 1.11305797 | -7.1360631 | 9.60E-13 |
| 73835 | ifitm5 | 22.9080772 | 4.1337595 | 0.579302 | 7.13575903 | 9.63E-13 |
| 23984 | PDE10A | 89.0091121 | 4.8759927 | 0.68343894 | 7.13449642 | 9.71E-13 |
| 72500 | IER5L | 1747.89229 | 1.03555924 | 0.14519141 | 7.1323725 | 9.87E-13 |
| 69091 | VPS26B | 2217.75515 | 0.43210274 | 0.0605927 | 7.13126679 | 9.94E-13 |
| 74580 | PYROXD2 | 26.6290177 | -3.5265192 | 0.4946857 | -7.1288077 | 1.01E-12 |
| 105278 | CDK20 | 215.269202 | 1.07868244 | 0.15141289 | 7.12411243 | 1.05E-12 |
| 67205 | UTP11L | 1032.73309 | 0.83804743 | 0.11763797 | 7.12395365 | 1.05E-12 |
| 216766 | GEMIN5 | 3124.88164 | 0.58600555 | 0.08226876 | 7.1230629 | 1.06E-12 |
| 68423 | ANKRD13D | 79.4222789 | -2.3315103 | 0.32741295 | -7.1210083 | 1.07E-12 |
| 18806 | PLD2 | 897.896309 | -0.7520974 | 0.10563925 | -7.119489 | 1.08E-12 |
| 15381 | HNRNPC | 4455.56844 | 0.42378142 | 0.05952614 | 7.1192496 | 1.09E-12 |
| 217431 | NOL10 | 3033.54653 | 0.70490364 | 0.09910085 | 7.11299264 | 1.14E-12 |
| 320354 | Grin1os | 23.3134444 | -7.975096 | 1.1217491 | -7.1095186 | 1.16E-12 |
| 70257 | 2010107E04I | 1164.15166 | 0.51321916 | 0.07219178 | 7.10910778 | 1.17E-12 |
| 60322 | CHST7 | 258.24116 | 1.0702729 | 0.15060852 | 7.10632364 | 1.19E-12 |
| 76943 | PSAPL1 | 19.9157116 | -7.7426439 | 1.08976872 | -7.1048506 | 1.20E-12 |
| 76089 | RAPGEF2 | 1675.65735 | 0.49995483 | 0.07036853 | 7.10480671 | 1.20E-12 |
| 21816 | TGM1 | 54.7889353 | -4.3072862 | 0.6063619 | -7.1034907 | 1.22E-12 |
| 14664 | SLC6A9 | 1065.46673 | 1.27675117 | 0.17975533 | 7.10271641 | 1.22E-12 |
| 16650 | KPNA6 | 4690.56425 | 0.81623701 | 0.11494597 | 7.10104961 | 1.24E-12 |
| 320506 | LMBRD2 | 1400.27586 | 0.63874174 | 0.08995154 | 7.10095392 | 1.24E-12 |

| | | | | | | |
|---|---|---|---|---|---|---|
| 76561 | SNX7 | 1273.64007 | 1.34917677 | 0.19004608 | 7.09920876 | 1.25E-12 |
| 114304 | SLC28A3 | 48.842932 | -9.0390713 | 1.27334823 | -7.098664 | 1.26E-12 |
| 16886 | LIMK2 | 1712.59 | -1.3207598 | 0.18606056 | -7.0985477 | 1.26E-12 |
| 75744 | SVIP | 46.5674298 | -8.9701235 | 1.26378808 | -7.0978066 | 1.27E-12 |
| 14778 | GPX3 | 124.936482 | -9.672676 | 1.36288325 | -7.0972154 | 1.27E-12 |
| 16924 | LNX1 | 61.1890822 | -7.9262189 | 1.11691487 | -7.0965291 | 1.28E-12 |
| 53413 | EXOC7 | 1031.89573 | -0.5216313 | 0.07352073 | -7.0950238 | 1.29E-12 |
| 319581 | XKR5 | 229.234013 | 4.93437215 | 0.69553342 | 7.09437108 | 1.30E-12 |
| 16956 | LPL | 47.7548173 | -4.6775249 | 0.65939044 | -7.0937105 | 1.31E-12 |
| 236727 | SLC9A7 | 37.9772137 | -6.6449963 | 0.93684538 | -7.0929488 | 1.31E-12 |
| 242122 | VTCN1 | 96.7644578 | -8.5725399 | 1.2090725 | -7.0901786 | 1.34E-12 |
| 74411 | plpp6 | 381.618657 | -1.3987515 | 0.1972957 | -7.0896202 | 1.34E-12 |
| 75758 | 9130401M01 | 1449.09563 | 0.74333763 | 0.10489139 | 7.08673603 | 1.37E-12 |
| 109205 | SOBP | 316.796293 | 4.70831708 | 0.66449801 | 7.08552469 | 1.39E-12 |
| 106618 | WDR90 | 795.137599 | 1.23624805 | 0.17455055 | 7.08246432 | 1.42E-12 |
| 103551 | E130012A19 | 600.41812 | 2.00362167 | 0.28295752 | 7.08099818 | 1.43E-12 |
| 214150 | AGO3 | 596.412654 | 1.25140132 | 0.17675015 | 7.08005819 | 1.44E-12 |
| 72107 | DSCC1 | 305.224397 | 1.00153155 | 0.14145979 | 7.0799733 | 1.44E-12 |
| 213491 | SZRD1 | 4253.81508 | 1.07049964 | 0.15123496 | 7.07838747 | 1.46E-12 |
| 11671 | ALDH3A2 | 2381.19647 | -0.9674244 | 0.1367345 | -7.0752034 | 1.49E-12 |
| 15242 | HHEX | 26.6839245 | 6.17627195 | 0.8731788 | 7.07331869 | 1.51E-12 |
| 78895 | PUS7L | 344.041705 | 0.98900526 | 0.13998077 | 7.06529369 | 1.60E-12 |
| 66841 | ETFDH | 1112.47751 | -0.8542078 | 0.12091796 | -7.0643585 | 1.61E-12 |
| 64898 | LPIN2 | 1605.09174 | 0.3337986 | 0.04725983 | 7.0630508 | 1.63E-12 |
| 67778 | Zfp639 | 664.560667 | 0.45643356 | 0.06465047 | 7.0600191 | 1.66E-12 |
| 50790 | ACSL4 | 10803.1483 | -1.0548023 | 0.14944406 | -7.0581746 | 1.69E-12 |
| 56774 | SLC6A14 | 31.2640593 | -8.3986345 | 1.18998048 | -7.0577919 | 1.69E-12 |
| 69327 | 1700007K13I | 46.8703942 | -5.0616864 | 0.71719922 | -7.0575738 | 1.69E-12 |
| 68728 | Trp53inp2 | 1234.37423 | -1.0996581 | 0.15589594 | -7.0537959 | 1.74E-12 |
| 12870 | CP | 10255.9126 | -1.8357715 | 0.26050768 | -7.0468994 | 1.83E-12 |
| 12981 | CSF2 | 17.6119115 | -7.5679411 | 1.07403977 | -7.0462392 | 1.84E-12 |
| 67414 | MFN1 | 1878.20302 | -0.4520946 | 0.06416577 | -7.0457294 | 1.84E-12 |
| 67199 | PFDN1 | 1630.19045 | -0.8422384 | 0.11958773 | -7.0428498 | 1.88E-12 |
| 107522 | ECE2 | 413.59903 | 0.80648319 | 0.11456204 | 7.03970668 | 1.93E-12 |
| 11519 | ADD2 | 335.755694 | -11.099484 | 1.57723241 | -7.0373165 | 1.96E-12 |
| 23959 | NT5E | 2300.64927 | -3.4356971 | 0.48823563 | -7.036965 | 1.96E-12 |
| 56807 | SCAMP5 | 429.153481 | -3.0332189 | 0.43119383 | -7.0344673 | 2.00E-12 |
| 20856 | STC2 | 269.491107 | 2.16271588 | 0.30746303 | 7.03406812 | 2.01E-12 |
| 14768 | LANCL1 | 703.194416 | -0.8395729 | 0.11944781 | -7.028784 | 2.08E-12 |
| 18712 | PIM1 | 420.279096 | -1.5377414 | 0.21884751 | -7.0265427 | 2.12E-12 |
| 66055 | SF3B6 | 1681.26887 | 0.62093161 | 0.08837086 | 7.02642937 | 2.12E-12 |
| 65106 | ARL6IP5 | 5687.23439 | 1.16210186 | 0.16540786 | 7.0256751 | 2.13E-12 |
| 238161 | akap6 | 164.941928 | -2.6477421 | 0.3768807 | -7.0254118 | 2.13E-12 |
| 56030 | TMEM131 | 5536.62892 | -0.364035 | 0.05181764 | -7.0253116 | 2.14E-12 |
| 654812 | ANGPTL7 | 78.3859129 | -9.7231659 | 1.3846629 | -7.0220455 | 2.19E-12 |

| | | | | | | |
|---|---|---|---|---|---|---|
| 16971 | LRP1 | 26893.7172 | 2.65685844 | 0.37839097 | 7.02146361 | 2.20E-12 |
| 13640 | EFNA5 | 2346.5783 | -1.07822 | 0.15372716 | -7.0138547 | 2.32E-12 |
| 66887 | LONP2 | 1085.44857 | -0.6422526 | 0.09163389 | -7.0088985 | 2.40E-12 |
| 69232 | QRICH1 | 3593.76219 | 0.46700446 | 0.06663854 | 7.0080238 | 2.42E-12 |
| 666938 | BEND4 | 100.084771 | -6.3266902 | 0.90292175 | -7.0069086 | 2.44E-12 |
| 80901 | Cxcr6 | 30.381525 | 7.6783623 | 1.09600473 | 7.00577478 | 2.46E-12 |
| 70686 | DUSP16 | 1379.0707 | -0.9975822 | 0.14239492 | -7.0057433 | 2.46E-12 |
| 269344 | ELL3 | 19.686307 | -7.7321912 | 1.103708 | -7.0056493 | 2.46E-12 |
| 76282 | GPT | 109.401708 | -1.2817162 | 0.18295996 | -7.0054464 | 2.46E-12 |
| 22031 | TRAF3 | 3272.71333 | 0.46551313 | 0.06646599 | 7.00377936 | 2.49E-12 |
| 382252 | A830080D01 | 559.252057 | -0.7086912 | 0.10119898 | -7.0029484 | 2.51E-12 |
| 57914 | CRLF2 | 705.272531 | -0.9515456 | 0.13587864 | -7.0029079 | 2.51E-12 |
| 237336 | TBPL1 | 971.041677 | 0.47692954 | 0.06810847 | 7.00249988 | 2.51E-12 |
| 246154 | VASN | 4704.99473 | 1.60355942 | 0.22905379 | 7.0007985 | 2.55E-12 |
| 14456 | GAS6 | 141.868598 | -9.8554219 | 1.40822346 | -6.9984787 | 2.59E-12 |
| 399566 | BTBD6 | 479.663218 | 0.88789448 | 0.12688298 | 6.99774302 | 2.60E-12 |
| 11820 | APP | 18702.3652 | -0.9735146 | 0.13913181 | -6.9970667 | 2.61E-12 |
| 242523 | DMRTA1 | 80.0556803 | -5.6794057 | 0.81173915 | -6.9965896 | 2.62E-12 |
| 107528 | MAGEE1 | 627.943474 | 1.12259707 | 0.16046125 | 6.99606341 | 2.63E-12 |
| 384783 | IRS2 | 406.822666 | -2.2754285 | 0.32524913 | -6.9959557 | 2.63E-12 |
| 16569 | KIF3B | 1972.05979 | -0.7041538 | 0.10066627 | -6.9949335 | 2.65E-12 |
| 66734 | MAP1LC3A | 454.690154 | -1.1153512 | 0.15946959 | -6.9941309 | 2.67E-12 |
| 18583 | PDE7A | 1154.47315 | -1.0420031 | 0.14900243 | -6.9931953 | 2.69E-12 |
| 63873 | TRPV4 | 1218.07119 | -1.9714745 | 0.28191727 | -6.9930961 | 2.69E-12 |
| 12125 | BCL2L11 | 644.248888 | -1.8005621 | 0.25749761 | -6.9925392 | 2.70E-12 |
| 66818 | SMIM7 | 1764.622 | -0.5366757 | 0.07675337 | -6.992211 | 2.71E-12 |
| 72518 | 2610307P16I | 43.2159306 | -4.8149185 | 0.68887202 | -6.9895689 | 2.76E-12 |
| 108100 | BAIAP2 | 2373.96115 | -1.1607303 | 0.16607298 | -6.9892781 | 2.76E-12 |
| 18227 | NR4A2 | 225.860871 | 1.20831052 | 0.17307152 | 6.98156761 | 2.92E-12 |
| 12021 | BARD1 | 1105.90618 | -0.7096501 | 0.10165998 | -6.9806248 | 2.94E-12 |
| 18405 | ORM1 | 28.5176617 | -8.2660778 | 1.18491771 | -6.9760775 | 3.04E-12 |
| 214359 | TMEM51 | 23.4132774 | -7.9765264 | 1.14349284 | -6.9755805 | 3.05E-12 |
| 67943 | MESDC2 | 2530.0989 | 0.6134971 | 0.08797325 | 6.97367768 | 3.09E-12 |
| 66622 | UBR7 | 2106.96696 | 0.80015737 | 0.11474313 | 6.97346661 | 3.09E-12 |
| 24069 | sufu | 701.730336 | 0.56851732 | 0.08153195 | 6.97293912 | 3.10E-12 |
| 22122 | TSTA3 | 1125.54355 | -0.6610029 | 0.0948131 | -6.9716413 | 3.13E-12 |
| 72480 | TSPYL4 | 145.855978 | -1.1856062 | 0.17013604 | -6.9685779 | 3.20E-12 |
| 320411 | A730089K16 | 16.5817003 | 7.53240087 | 1.08111993 | 6.96722043 | 3.23E-12 |
| 107999 | GTPBP6 | 724.772627 | -0.624592 | 0.08970113 | -6.9630343 | 3.33E-12 |
| 12332 | CAPG | 11395.5885 | 0.42145105 | 0.06053502 | 6.96210285 | 3.35E-12 |
| 17995 | NDUFV1 | 2426.40523 | -0.2761874 | 0.03967127 | -6.9618987 | 3.36E-12 |
| 13855 | EPN2 | 1971.62628 | -1.044376 | 0.15002394 | -6.9613958 | 3.37E-12 |
| 270151 | NLRX1 | 162.310144 | -1.7466188 | 0.2509375 | -6.9603739 | 3.39E-12 |
| 67568 | MRFAP1 | 4271.61113 | 0.46520745 | 0.06684004 | 6.96001185 | 3.40E-12 |
| 76464 | KNL1 | 2839.18906 | 0.52700419 | 0.07577223 | 6.9551098 | 3.52E-12 |

| | | | | | | |
|---|---|---|---|---|---|---|
| 71791 | CPA4 | 27.282554 | -7.468501 | 1.07449327 | -6.9507193 | 3.63E-12 |
| 26358 | Aldh1a7 | 94.5264108 | -6.376227 | 0.91743108 | -6.9500883 | 3.65E-12 |
| 13555 | E2F1 | 2020.77409 | 0.36975068 | 0.05320361 | 6.94972937 | 3.66E-12 |
| 105445 | DOCK9 | 2728.39809 | -2.3739332 | 0.3416595 | -6.948243 | 3.70E-12 |
| 14620 | GJB3 | 475.395475 | -5.5045196 | 0.79249317 | -6.9458259 | 3.76E-12 |
| 11740 | SLC25A5 | 5292.23836 | 0.3371588 | 0.04854242 | 6.94565351 | 3.77E-12 |
| 208618 | ETL4 | 1321.37443 | -1.8800611 | 0.27094376 | -6.9389349 | 3.95E-12 |
| 78600 | PDE6H | 25.2733265 | 5.34070957 | 0.76999602 | 6.93602232 | 4.03E-12 |
| 217716 | MLH3 | 780.244082 | -0.522733 | 0.07545709 | -6.9275535 | 4.28E-12 |
| 18805 | PLD1 | 404.747755 | -1.1352117 | 0.16388468 | -6.9268932 | 4.30E-12 |
| 226090 | ERMP1 | 3314.54311 | 1.47301796 | 0.21270368 | 6.9252116 | 4.35E-12 |
| 235469 | Zfp280d | 733.791291 | 0.66414769 | 0.0959035 | 6.92516602 | 4.35E-12 |
| 16337 | INSR | 1421.36965 | -0.9645707 | 0.13929007 | -6.9249063 | 4.36E-12 |
| 211712 | pcdh9 | 16.4377134 | -7.4698253 | 1.07928122 | -6.9211112 | 4.48E-12 |
| 13870 | ercc1 | 1670.45874 | -0.4665547 | 0.06742385 | -6.9197285 | 4.53E-12 |
| 50880 | SCLY | 705.550503 | -2.0577248 | 0.29739854 | -6.9190816 | 4.55E-12 |
| 57370 | B4GALT3 | 605.820159 | -0.8240463 | 0.119161 | -6.915402 | 4.67E-12 |
| 227292 | CTDSP1 | 1945.46137 | 0.33910353 | 0.04905966 | 6.91206364 | 4.78E-12 |
| 213311 | FBXL21 | 31.6931264 | -5.3610252 | 0.77603512 | -6.908225 | 4.91E-12 |
| 216177 | AU041133 | 121.560875 | -1.4596697 | 0.21140311 | -6.9046746 | 5.03E-12 |
| 113868 | Acaa1a | 431.677125 | -0.7632206 | 0.11054115 | -6.9044025 | 5.04E-12 |
| 225348 | WDR36 | 4608.2014 | -0.3585135 | 0.05198162 | -6.8969277 | 5.31E-12 |
| 108115 | SLCO4A1 | 50.7998757 | -7.6459177 | 1.10865367 | -6.8965791 | 5.33E-12 |
| 69665 | UPK3BL | 59.5157614 | -7.2705414 | 1.05428858 | -6.8961588 | 5.34E-12 |
| 217216 | BC030867 | 775.122097 | 0.63812101 | 0.09258058 | 6.8926015 | 5.48E-12 |
| 57376 | SMARCE1 | 1092.46098 | 0.4910219 | 0.07124746 | 6.89178071 | 5.51E-12 |
| 78912 | SP2 | 795.378229 | 0.60963688 | 0.08847869 | 6.89021122 | 5.57E-12 |
| 116748 | LSM10 | 238.133745 | 0.83412272 | 0.12109875 | 6.88795498 | 5.66E-12 |
| 68178 | CGNL1 | 8506.85436 | -1.9401915 | 0.28172316 | -6.8868725 | 5.70E-12 |
| 22123 | PSMD3 | 6799.68295 | 0.2994015 | 0.04348241 | 6.88557768 | 5.76E-12 |
| 67903 | GIPC1 | 1863.06317 | -0.8572192 | 0.12456978 | -6.8814379 | 5.93E-12 |
| 102644 | OAF | 4915.16399 | 3.75964177 | 0.54641675 | 6.88053903 | 5.96E-12 |
| 19303 | PXN | 4595.9217 | -0.5995493 | 0.08715747 | -6.878921 | 6.03E-12 |
| 106512 | GPSM3 | 72.9528495 | -5.4201137 | 0.78793537 | -6.878881 | 6.03E-12 |
| 14802 | GRIA4 | 25.5149381 | -8.1013099 | 1.17775311 | -6.8786147 | 6.04E-12 |
| 70031 | CMTM8 | 32.5189603 | -7.7281761 | 1.12354063 | -6.8784127 | 6.05E-12 |
| 26448 | MOK | 47.810139 | -1.9035524 | 0.27703931 | -6.871055 | 6.37E-12 |
| 76263 | GSTK1 | 17.6217771 | -7.5672224 | 1.10133215 | -6.8709721 | 6.38E-12 |
| 51797 | Ctps | 4869.21117 | 1.16742095 | 0.17001144 | 6.86672004 | 6.57E-12 |
| 212733 | CCDC64B | 27.8144034 | -7.4974215 | 1.09207746 | -6.8652836 | 6.64E-12 |
| 70575 | GFOD2 | 362.009058 | -0.7543564 | 0.10989865 | -6.8641101 | 6.69E-12 |
| 68001 | 1110004E09I | 665.908036 | -0.609338 | 0.08877607 | -6.8637642 | 6.71E-12 |
| 20544 | SLC9A1 | 2187.02272 | 0.71985082 | 0.10488178 | 6.86345 | 6.72E-12 |
| 55936 | CTPS2 | 1242.33453 | -0.6582672 | 0.09593055 | -6.8619143 | 6.79E-12 |
| 14904 | GTPBP1 | 2302.16575 | 0.35412087 | 0.05164674 | 6.85659706 | 7.05E-12 |

| | | | | | | |
|---|---|---|---|---|---|---|
| 13665 | EIF2S1 | 4253.94542 | 0.50413272 | 0.07353083 | 6.85607287 | 7.08E-12 |
| 215378 | BRINP3 | 1752.92532 | -6.2266595 | 0.90835401 | -6.854882 | 7.14E-12 |
| 70747 | TSPAN2 | 384.622377 | -4.6732455 | 0.6818159 | -6.8541163 | 7.18E-12 |
| 56753 | TACSTD2 | 477.179569 | -12.328411 | 1.79907073 | -6.8526549 | 7.25E-12 |
| 56451 | SUCLG1 | 2173.71701 | 0.44249938 | 0.0645833 | 6.85160657 | 7.30E-12 |
| 238317 | ELMSAN1 | 2196.71243 | 0.83507446 | 0.12189909 | 6.85053916 | 7.36E-12 |
| 19775 | XPR1 | 2505.81487 | -0.3821295 | 0.05582119 | -6.8455989 | 7.62E-12 |
| 77644 | C330007P06I | 872.695002 | -0.701069 | 0.10246922 | -6.8417518 | 7.82E-12 |
| 338364 | TRIM65 | 257.086386 | 0.94111515 | 0.13757627 | 6.8406794 | 7.88E-12 |
| 15227 | FOXF1 | 277.274711 | 4.36624167 | 0.63840438 | 6.83930407 | 7.96E-12 |
| 215449 | RAP1B | 7378.1533 | -0.4590256 | 0.06712144 | -6.8387335 | 7.99E-12 |
| 14231 | FKBP7 | 215.041749 | 1.50901078 | 0.2207627 | 6.83544273 | 8.18E-12 |
| 68250 | FAM96A | 1376.01509 | 0.70963481 | 0.10385293 | 6.83307466 | 8.31E-12 |
| 68427 | SLC39A13 | 1997.99944 | 0.47879869 | 0.07010054 | 6.83017139 | 8.48E-12 |
| 69368 | WDFY1 | 1814.59028 | -0.7984121 | 0.11691634 | -6.8289179 | 8.56E-12 |
| 74375 | GCC1 | 726.474471 | -0.6541516 | 0.09582257 | -6.8266973 | 8.69E-12 |
| 71721 | FAM13C | 97.1817706 | 4.11023964 | 0.60209003 | 6.82661962 | 8.69E-12 |
| 12385 | CTNNA1 | 15720.5671 | -0.4071039 | 0.05964264 | -6.8257198 | 8.75E-12 |
| 14281 | FOS | 69.8502212 | -1.9989999 | 0.29293213 | -6.8241058 | 8.85E-12 |
| 76267 | FADS1 | 5003.89666 | 1.39298828 | 0.20416017 | 6.82301699 | 8.91E-12 |
| 18654 | PGF | 89.5288775 | -6.4487784 | 0.94534417 | -6.8216198 | 9.00E-12 |
| 18577 | PDE4A | 516.512105 | 3.71099867 | 0.54401444 | 6.82150771 | 9.01E-12 |
| 14038 | WFDC18 | 81.1972881 | -9.7729326 | 1.4338262 | -6.8159813 | 9.36E-12 |
| 13548 | DYRK1A | 2576.66723 | -0.6649116 | 0.09761749 | -6.8113985 | 9.67E-12 |
| 55943 | STX8 | 415.13722 | -0.606123 | 0.08901517 | -6.8092101 | 9.81E-12 |
| 246293 | KLHL8 | 169.829999 | -0.8846484 | 0.12995648 | -6.8072669 | 9.95E-12 |
| 226861 | HHAT | 210.353598 | 0.75901276 | 0.11153792 | 6.80497514 | 1.01E-11 |
| 242297 | FAM110B | 134.257664 | -1.030286 | 0.15140537 | -6.804818 | 1.01E-11 |
| 67800 | DGAT2 | 122.164888 | 1.62175858 | 0.23833936 | 6.80440955 | 1.01E-11 |
| 73178 | WASL | 2714.21941 | -0.5404309 | 0.07946609 | -6.8007736 | 1.04E-11 |
| 21354 | TAP1 | 292.202381 | -1.5714243 | 0.23115743 | -6.7980695 | 1.06E-11 |
| 14621 | GJB4 | 605.576389 | -1.950793 | 0.28698896 | -6.7974495 | 1.06E-11 |
| 231070 | INSIG1 | 2545.99016 | 0.42467563 | 0.06248669 | 6.79625706 | 1.07E-11 |
| 53945 | SLC40A1 | 75.0300076 | -2.698743 | 0.39726671 | -6.7932775 | 1.10E-11 |
| 59012 | MOXD1 | 101.068507 | -5.3414109 | 0.78639099 | -6.7923093 | 1.10E-11 |
| 67477 | ABHD15 | 49.7721174 | 2.44136958 | 0.35946025 | 6.79176508 | 1.11E-11 |
| 20833 | SSRP1 | 9057.39 | 0.36832727 | 0.05423188 | 6.7917112 | 1.11E-11 |
| 68875 | TMCC2 | 579.597344 | 1.87281975 | 0.27578399 | 6.79089359 | 1.11E-11 |
| 106869 | TNFAIP8 | 586.982306 | -1.4950915 | 0.22020192 | -6.7896389 | 1.12E-11 |
| 381085 | TBC1D22B | 824.16091 | -0.616792 | 0.09084883 | -6.7892127 | 1.13E-11 |
| 14706 | GNG4 | 25.1495959 | 5.07854192 | 0.74818525 | 6.78781349 | 1.14E-11 |
| 544696 | TBC1D32 | 300.028392 | -0.8725735 | 0.12855417 | -6.7875939 | 1.14E-11 |
| 56086 | SET | 4230.93827 | 0.45736956 | 0.06738326 | 6.78758424 | 1.14E-11 |
| 80748 | BC004004 | 1343.49092 | -0.5470933 | 0.0806023 | -6.7875646 | 1.14E-11 |
| 15983 | IFRD2 | 1653.52616 | 0.8817282 | 0.12991563 | 6.78692942 | 1.15E-11 |

| | | | | | | |
|---|---|---|---|---|---|---|
| 331026 | GMPPB | 542.367588 | 0.63312035 | 0.0932997 | 6.78587795 | 1.15E-11 |
| 74120 | Zfp263 | 723.900796 | -0.7134032 | 0.10519719 | -6.7815807 | 1.19E-11 |
| 242736 | PRAMEF8 | 605.240846 | 0.56293201 | 0.08302389 | 6.78036171 | 1.20E-11 |
| 108123 | NAPG | 1080.43846 | -0.3879049 | 0.05723681 | -6.7771941 | 1.23E-11 |
| 20617 | snca | 45.9308243 | -8.9505087 | 1.32097985 | -6.7756588 | 1.24E-11 |
| 75646 | RAI14 | 6934.10083 | 1.15215628 | 0.17012994 | 6.77221363 | 1.27E-11 |
| 80292 | ZXDC | 1446.01733 | 0.47447144 | 0.07007975 | 6.77044946 | 1.28E-11 |
| 77134 | HNRNPA0 | 12415.979 | 0.87164757 | 0.12878178 | 6.76840776 | 1.30E-11 |
| 11668 | ALDH1A1 | 364.994935 | -9.53367 | 1.40893809 | -6.7665642 | 1.32E-11 |
| 234814 | MTHFSD | 575.048907 | -0.6265753 | 0.09262548 | -6.7646102 | 1.34E-11 |
| 58203 | ZBP1 | 63.6397761 | -4.267466 | 0.63095513 | -6.7635016 | 1.35E-11 |
| 69876 | THAP3 | 187.929978 | 0.86970296 | 0.12861396 | 6.76211964 | 1.36E-11 |
| 29805 | ZNHIT2 | 584.023007 | 0.50224529 | 0.07428403 | 6.76114801 | 1.37E-11 |
| 15958 | IFIT2 | 1462.23038 | 0.81838909 | 0.12111209 | 6.75728665 | 1.41E-11 |
| 213522 | PLEKHG6 | 113.063442 | -3.5785887 | 0.52960198 | -6.7571287 | 1.41E-11 |
| 71999 | FBXO22 | 1203.5187 | 0.39819266 | 0.0589765 | 6.75171725 | 1.46E-11 |
| 30956 | AASS | 17.5369473 | -7.5663889 | 1.12166894 | -6.7456525 | 1.52E-11 |
| 52639 | WIPI1 | 1337.16491 | -0.9553734 | 0.14165107 | -6.744555 | 1.53E-11 |
| 217864 | RCOR1 | 1702.11112 | 0.52034212 | 0.07715724 | 6.74391796 | 1.54E-11 |
| 20717 | Serpina3m | 110.178119 | -1.959593 | 0.29061252 | -6.7429753 | 1.55E-11 |
| 66817 | tmem170 | 129.81124 | -1.0367975 | 0.1537788 | -6.7421353 | 1.56E-11 |
| 217262 | ABCA9 | 26.4801654 | 2.99955536 | 0.44529016 | 6.7361816 | 1.63E-11 |
| 23871 | ETS1 | 4159.04414 | 0.97665793 | 0.14498816 | 6.73612197 | 1.63E-11 |
| 11987 | SLC7A1 | 7279.82013 | 0.5086028 | 0.0755242 | 6.73430259 | 1.65E-11 |
| 72097 | 2010300C02l | 24.0768813 | -5.542137 | 0.82308732 | -6.7333524 | 1.66E-11 |
| 16425 | ITIH2 | 204.608765 | -2.1361886 | 0.31727379 | -6.7329501 | 1.66E-11 |
| 99138 | STARD7 | 2687.44226 | 0.41022018 | 0.06092823 | 6.73284316 | 1.66E-11 |
| 56273 | PEX14 | 1020.50912 | 0.58541911 | 0.0869509 | 6.73275493 | 1.66E-11 |
| 74735 | TRIM14 | 48.2233867 | -8.2977343 | 1.23296918 | -6.7298798 | 1.70E-11 |
| 231510 | AGPAT9 | 388.703363 | -1.003521 | 0.14918348 | -6.7267568 | 1.73E-11 |
| 217335 | FBF1 | 1829.3043 | -0.5551642 | 0.08254655 | -6.7254679 | 1.75E-11 |
| 211488 | ADO | 2231.84017 | 0.6843945 | 0.10177484 | 6.72459445 | 1.76E-11 |
| 20677 | SOX4 | 4408.34521 | 0.47905613 | 0.07126703 | 6.72198791 | 1.79E-11 |
| 66972 | SLC25A23 | 127.414711 | -7.9836684 | 1.18770828 | -6.7219102 | 1.79E-11 |
| 66181 | NOP10 | 789.695099 | 0.56773429 | 0.08447279 | 6.72091288 | 1.81E-11 |
| 230098 | ARHGEF39 | 221.758398 | 0.8857225 | 0.13182692 | 6.71882865 | 1.83E-11 |
| 225341 | LIMS2 | 331.530888 | -3.3781388 | 0.50291613 | -6.7171017 | 1.85E-11 |
| 239337 | adamts12 | 265.134558 | 2.97331622 | 0.44269402 | 6.71641381 | 1.86E-11 |
| 93714 | PCDHGA6 | 32.0209121 | -4.3176875 | 0.64294545 | -6.7154802 | 1.87E-11 |
| 93806 | Serpinb9g | 38.4137853 | -5.9255283 | 0.88240027 | -6.7152386 | 1.88E-11 |
| 72672 | Zfp518a | 420.442847 | -1.0487894 | 0.15625293 | -6.7121264 | 1.92E-11 |
| 19157 | CYTH1 | 489.411959 | -0.5941853 | 0.08852583 | -6.7119998 | 1.92E-11 |
| 232232 | HDAC11 | 102.421738 | -3.0305646 | 0.45151552 | -6.7119831 | 1.92E-11 |
| 100710 | PDS5B | 2989.07076 | 0.50136425 | 0.07472773 | 6.70921332 | 1.96E-11 |
| 236285 | LANCL3 | 1254.01561 | 1.92755796 | 0.28764105 | 6.70126166 | 2.07E-11 |

| | | | | | | |
|---|---|---|---|---|---|---|
| 18301 | FXYD5 | 2382.88687 | 1.61145668 | 0.24049184 | 6.70067098 | 2.07E-11 |
| 675363 | Gm9640 | 20.6378346 | -4.9736786 | 0.74240623 | -6.6994031 | 2.09E-11 |
| 67581 | TBC1D23 | 1639.38861 | 0.66059907 | 0.09864917 | 6.696448 | 2.14E-11 |
| 227102 | ORMDL1 | 453.602487 | -0.9779851 | 0.14607423 | -6.6951244 | 2.15E-11 |
| 208967 | THNSL1 | 214.261899 | -1.9235984 | 0.28732023 | -6.6949632 | 2.16E-11 |
| 73192 | XPOT | 4174.25917 | 0.43074837 | 0.06435741 | 6.69306532 | 2.19E-11 |
| 11491 | ADAM17 | 3777.29732 | 0.64468856 | 0.09632518 | 6.69283542 | 2.19E-11 |
| 109218 | TMEM139 | 16.4304003 | -7.4660167 | 1.11568698 | -6.6918561 | 2.20E-11 |
| 67828 | LCE1F | 17.5057207 | -7.5616224 | 1.12997732 | -6.6918355 | 2.20E-11 |
| 223669 | Zfp7 | 546.699371 | -0.9908785 | 0.1480828 | -6.6913812 | 2.21E-11 |
| 104248 | CABIN1 | 2586.05451 | -0.3371816 | 0.05039106 | -6.6912993 | 2.21E-11 |
| 12846 | COMT | 2229.24014 | -0.620546 | 0.09281176 | -6.6860702 | 2.29E-11 |
| 21417 | ZEB1 | 1477.6829 | 1.31352956 | 0.19646383 | 6.68585943 | 2.30E-11 |
| 110750 | CSE1L | 7246.93367 | 0.31346125 | 0.04690683 | 6.68263521 | 2.35E-11 |
| 73137 | PRRC1 | 1426.76739 | 0.46795741 | 0.0700438 | 6.68092568 | 2.37E-11 |
| 11770 | FABP4 | 199.333244 | -4.8387052 | 0.72460696 | -6.6776963 | 2.43E-11 |
| 66618 | SNRNP27 | 699.144913 | 0.68642161 | 0.10280884 | 6.67667874 | 2.44E-11 |
| 11720 | MAT1A | 23.3872658 | -7.2516464 | 1.08650772 | -6.6742705 | 2.48E-11 |
| 52150 | KCNK6 | 43.8068278 | 3.3741107 | 0.50570716 | 6.67206433 | 2.52E-11 |
| 71169 | NBAS | 1466.96615 | 0.38320242 | 0.05745592 | 6.66950239 | 2.57E-11 |
| 104360 | ISL2 | 25.7787723 | 6.11254874 | 0.91650129 | 6.66943825 | 2.57E-11 |
| 71985 | ACAD10 | 79.7149577 | -1.6312336 | 0.24464825 | -6.6676691 | 2.60E-11 |
| 218793 | UBE2E2 | 721.616933 | -0.5800322 | 0.08699809 | -6.6671832 | 2.61E-11 |
| 407786 | TAF9B | 207.163916 | 0.69462874 | 0.10418753 | 6.66710076 | 2.61E-11 |
| 209446 | TFE3 | 2237.46158 | 0.38288467 | 0.05744166 | 6.66562698 | 2.64E-11 |
| 78521 | B230219D22 | 4372.34117 | 0.40293322 | 0.06050898 | 6.65906517 | 2.76E-11 |
| 103743 | TMEM98 | 551.013695 | -1.1851896 | 0.17801368 | -6.6578568 | 2.78E-11 |
| 11908 | ATF1 | 1428.52928 | 0.49536338 | 0.07440598 | 6.65757522 | 2.78E-11 |
| 51812 | MCRS1 | 369.043038 | 0.5953787 | 0.08943739 | 6.65693271 | 2.80E-11 |
| 385643 | Kng2 | 60.0937905 | 5.41460149 | 0.81380487 | 6.65343957 | 2.86E-11 |
| 71445 | 5530601H04 | 141.20701 | -1.2978437 | 0.1951289 | -6.6512122 | 2.91E-11 |
| 380601 | FASTKD5 | 822.239021 | 0.50389408 | 0.07576177 | 6.65103317 | 2.91E-11 |
| 18114 | RRP1 | 4964.44072 | -0.3322083 | 0.04994882 | -6.6509739 | 2.91E-11 |
| 22375 | WARS | 3547.73207 | 0.22679679 | 0.03410647 | 6.64967037 | 2.94E-11 |
| 319604 | FAM168A | 2302.44166 | -0.470262 | 0.07072644 | -6.6490276 | 2.95E-11 |
| 17083 | TMED1 | 994.890833 | 0.90805001 | 0.136607 | 6.64717057 | 2.99E-11 |
| 244745 | DPY19L1 | 7360.68704 | 0.86036413 | 0.12947286 | 6.64513096 | 3.03E-11 |
| 330836 | SLC7A6 | 1877.38342 | 0.77904171 | 0.11724519 | 6.64455132 | 3.04E-11 |
| 67826 | SNAP47 | 840.063465 | -0.9378149 | 0.14115368 | -6.6439281 | 3.05E-11 |
| 16981 | LRRN3 | 51.9531855 | 3.28949554 | 0.49524083 | 6.64221387 | 3.09E-11 |
| 269614 | PANK4 | 579.151025 | 0.57033444 | 0.08588063 | 6.64101387 | 3.12E-11 |
| 230597 | ZFYVE9 | 478.959315 | 0.70811057 | 0.10663201 | 6.6406945 | 3.12E-11 |
| 78834 | Zfp623 | 433.825568 | -0.5725701 | 0.08625596 | -6.6380353 | 3.18E-11 |
| 320878 | MICAL2 | 1706.8943 | 1.04363315 | 0.15724151 | 6.63713503 | 3.20E-11 |
| 97064 | WWTR1 | 5659.7422 | 0.66219078 | 0.09979929 | 6.63522567 | 3.24E-11 |

| | | | | | | |
|---|---|---|---|---|---|---|
| 22160 | TWIST1 | 2215.914 | 3.13826336 | 0.47297496 | 6.63515755 | 3.24E-11 |
| 235072 | 41888 | 8277.71085 | 0.43384981 | 0.06539859 | 6.63393244 | 3.27E-11 |
| 104725 | SPTSSA | 1604.31889 | 0.50650608 | 0.07635924 | 6.63320001 | 3.28E-11 |
| 24058 | SIGIRR | 172.170198 | 1.95841951 | 0.29530171 | 6.63192748 | 3.31E-11 |
| 53761 | PRRC2A | 23147.1266 | 0.21778182 | 0.03283923 | 6.63175786 | 3.32E-11 |
| 74142 | LONP1 | 4512.36809 | -0.5064385 | 0.07636953 | -6.6314215 | 3.32E-11 |
| 70779 | PRDM5 | 73.9978593 | -9.6400334 | 1.45401715 | -6.629931 | 3.36E-11 |
| 19274 | PTPRM | 1120.15788 | 1.38617642 | 0.20909673 | 6.6293548 | 3.37E-11 |
| 22601 | YAP1 | 5220.614 | 0.51971317 | 0.07839698 | 6.62924968 | 3.37E-11 |
| 78795 | armc9 | 411.709728 | -1.1746314 | 0.17726528 | -6.6264042 | 3.44E-11 |
| 213895 | BMS1 | 4273.38944 | 0.57544426 | 0.08688441 | 6.62310158 | 3.52E-11 |
| 14225 | FKBP1A | 7742.88614 | 0.39245277 | 0.05925526 | 6.62308706 | 3.52E-11 |
| 74558 | GVIN1 | 13.8635482 | -7.2248908 | 1.09089196 | -6.6229206 | 3.52E-11 |
| 67489 | AP4B1 | 461.77688 | -0.523565 | 0.07905647 | -6.6226703 | 3.53E-11 |
| 399603 | FAM84B | 1178.47451 | -1.1618612 | 0.17551863 | -6.619589 | 3.60E-11 |
| 320595 | PHF8 | 1348.73546 | -0.6427113 | 0.09713703 | -6.6165426 | 3.68E-11 |
| 22642 | ZBTB17 | 806.253441 | 0.92233826 | 0.13944827 | 6.61419632 | 3.74E-11 |
| 329160 | 9130024F11I | 76.2679233 | 2.145007 | 0.32443497 | 6.61151595 | 3.80E-11 |
| 66960 | FAM188A | 3932.88058 | -2.003135 | 0.30304329 | -6.6100621 | 3.84E-11 |
| 67921 | UBE2F | 749.294599 | -0.6471862 | 0.09802817 | -6.6020426 | 4.06E-11 |
| 21763 | TEX2 | 1457.45479 | -0.7646418 | 0.1158557 | -6.5999497 | 4.11E-11 |
| 67131 | ACBD4 | 355.048601 | -0.9117599 | 0.13814968 | -6.5997974 | 4.12E-11 |
| 17064 | CD93 | 310.140969 | -3.5652722 | 0.540371 | -6.5978229 | 4.17E-11 |
| 66660 | SLTM | 2929.23689 | 0.26490781 | 0.0401574 | 6.59673767 | 4.20E-11 |
| 215627 | ZBTB8B | 20.2708531 | -7.0417816 | 1.06766612 | -6.5954903 | 4.24E-11 |
| 72119 | TPX2 | 6345.4251 | 0.34251831 | 0.05193796 | 6.59475891 | 4.26E-11 |
| 20444 | ST3GAL2 | 2499.49377 | 1.13555783 | 0.17220732 | 6.59413208 | 4.28E-11 |
| 74287 | KCMF1 | 4003.64484 | 0.73855365 | 0.11200173 | 6.59412699 | 4.28E-11 |
| 26443 | PSMA6 | 4545.54142 | 0.391774 | 0.05942313 | 6.59295414 | 4.31E-11 |
| 13850 | EPHX2 | 38.6538151 | -4.9017012 | 0.74349016 | -6.5928259 | 4.32E-11 |
| 56229 | THSD1 | 50.9709475 | -2.2835273 | 0.34639303 | -6.5923016 | 4.33E-11 |
| 68421 | LMBRD1 | 1193.08481 | -1.1588997 | 0.17580414 | -6.591993 | 4.34E-11 |
| 320795 | PKN1 | 1558.9468 | -0.5115504 | 0.07764128 | -6.5886393 | 4.44E-11 |
| 12813 | COL10A1 | 38.5369637 | 5.52742824 | 0.83895256 | 6.58848723 | 4.44E-11 |
| 54445 | unc93b1 | 126.18617 | -3.4080479 | 0.51731278 | -6.5879833 | 4.46E-11 |
| 17864 | MYBL1 | 655.61528 | -1.1753902 | 0.17847957 | -6.5855728 | 4.53E-11 |
| 237411 | Zfp938 | 216.812439 | -0.8909117 | 0.13529149 | -6.5851276 | 4.54E-11 |
| 67217 | L3HYPDH | 831.874761 | 1.48818209 | 0.22601653 | 6.58439477 | 4.57E-11 |
| 66849 | PPP1R2 | 1418.00076 | -0.8033604 | 0.12201422 | -6.5841534 | 4.57E-11 |
| 70661 | SIK3 | 2105.61082 | 0.55594728 | 0.08446828 | 6.58172857 | 4.65E-11 |
| 27049 | ETV3 | 998.696369 | -0.537493 | 0.08167458 | -6.5809088 | 4.68E-11 |
| 18189 | NRXN1 | 46.5780073 | 4.81367606 | 0.73161758 | 6.57949754 | 4.72E-11 |
| 54124 | CKS1B | 2059.54053 | 0.41041113 | 0.06237925 | 6.57928896 | 4.73E-11 |
| 229521 | SYT11 | 113.258493 | -1.3897367 | 0.21125184 | -6.5785778 | 4.75E-11 |
| 240354 | MALT1 | 528.598357 | -0.7891564 | 0.11996298 | -6.578333 | 4.76E-11 |

| | | | | | | |
|---|---|---|---|---|---|---|
| 434156 | EID2B | 115.361993 | 0.95815481 | 0.14566176 | 6.57794342 | 4.77E-11 |
| 215821 | ARFGEF3 | 66.9258658 | -8.0584395 | 1.22508392 | -6.5778673 | 4.77E-11 |
| 69724 | RNASEH2A | 763.212631 | -0.6230811 | 0.09475453 | -6.5757392 | 4.84E-11 |
| 320311 | RNF152 | 41.094302 | -7.3553542 | 1.11881289 | -6.5742487 | 4.89E-11 |
| 56443 | ARPC1A | 1973.53798 | -0.4489241 | 0.06830506 | -6.5723404 | 4.95E-11 |
| 79201 | Tnfrsf23 | 2569.17725 | -1.203965 | 0.18323378 | -6.5706501 | 5.01E-11 |
| 381802 | TSEN2 | 331.518206 | 0.88980735 | 0.135465 | 6.56854053 | 5.08E-11 |
| 16559 | KIF17 | 34.8583681 | -3.1696311 | 0.48256133 | -6.5683486 | 5.09E-11 |
| 69216 | SVBP | 249.431849 | 0.80225614 | 0.12214911 | 6.56784242 | 5.10E-11 |
| 59287 | NCSTN | 3538.43397 | -0.4725748 | 0.0720105 | -6.5625816 | 5.29E-11 |
| 83430 | IL23A | 22.7977686 | -7.9386201 | 1.20984831 | -6.5616657 | 5.32E-11 |
| 215615 | RNPEP | 1907.13022 | -0.8093922 | 0.12348779 | -6.554431 | 5.59E-11 |
| 66822 | FBXO25 | 555.059149 | -0.6410976 | 0.0978363 | -6.5527577 | 5.65E-11 |
| 225724 | MAPK4 | 35.7662734 | -8.5893252 | 1.31088793 | -6.5522956 | 5.67E-11 |
| 19331 | RAB19 | 17.1722884 | -7.5334508 | 1.15020343 | -6.5496681 | 5.77E-11 |
| 68142 | INO80 | 1579.73809 | 0.51717098 | 0.07896496 | 6.54937299 | 5.78E-11 |
| 67963 | NPC2 | 11856.7097 | 0.85514269 | 0.13057872 | 6.54886727 | 5.80E-11 |
| 268709 | FAM107A | 20.3980044 | -7.7835096 | 1.18858285 | -6.548563 | 5.81E-11 |
| 20615 | SNAPIN | 1085.7211 | -0.4391238 | 0.06705781 | -6.5484358 | 5.81E-11 |
| 14319 | FTH1 | 114638.114 | -1.7683596 | 0.27012765 | -6.546385 | 5.89E-11 |
| 56812 | DNAJB2 | 996.537203 | -0.696658 | 0.10642951 | -6.5457223 | 5.92E-11 |
| 12014 | BACH2 | 31.0014354 | -3.2147484 | 0.49121415 | -6.5444946 | 5.97E-11 |
| 109065 | DNAAF2 | 838.820488 | 0.77882357 | 0.11901198 | 6.54407723 | 5.99E-11 |
| 66824 | pycard | 24.650627 | -8.0514495 | 1.23045518 | -6.5434724 | 6.01E-11 |
| 74769 | pik3cb | 1722.40961 | -0.7508332 | 0.11476337 | -6.5424462 | 6.05E-11 |
| 103098 | SLC6A15 | 109.644666 | -10.206473 | 1.56011076 | -6.5421461 | 6.06E-11 |
| 22166 | Txn1 | 9988.79529 | 0.78595575 | 0.12016852 | 6.54044606 | 6.13E-11 |
| 68975 | MED27 | 412.497862 | 0.52025711 | 0.07965357 | 6.53149763 | 6.51E-11 |
| 19212 | PTER | 514.499053 | -1.14377 | 0.17515338 | -6.5301053 | 6.57E-11 |
| 11758 | PRDX6 | 9919.87004 | 0.98863642 | 0.15142228 | 6.52900216 | 6.62E-11 |
| 229706 | SLC6A17 | 223.772794 | -11.236054 | 1.72181793 | -6.5256923 | 6.77E-11 |
| 13557 | E2F3 | 1618.39831 | 0.67717589 | 0.1037772 | 6.5252856 | 6.79E-11 |
| 16784 | LAMP2 | 11171.3497 | -0.661617 | 0.10139867 | -6.5249086 | 6.80E-11 |
| 232087 | MAT2A | 7883.55694 | 0.3960216 | 0.06072515 | 6.52154223 | 6.96E-11 |
| 243653 | CLEC1A | 69.0902014 | 7.15668718 | 1.09742224 | 6.5213615 | 6.97E-11 |
| 75219 | DUSP18 | 1378.53449 | -0.4472374 | 0.06859159 | -6.5202947 | 7.02E-11 |
| 100039691 | Lncenc1 | 27.6542014 | -2.3428434 | 0.35942056 | -6.51839 | 7.11E-11 |
| 17384 | MMP10 | 230.605487 | 2.86683703 | 0.43991505 | 6.51679684 | 7.18E-11 |
| 68895 | RASL11A | 47.3607252 | 2.5535578 | 0.39203229 | 6.51364143 | 7.34E-11 |
| 13819 | EPAS1 | 322.573368 | -1.4820643 | 0.22764049 | -6.5105478 | 7.49E-11 |
| 240168 | RASGRP3 | 267.924956 | 4.61388187 | 0.70902875 | 6.50732692 | 7.65E-11 |
| 14114 | FBLN1 | 51.2299273 | -9.1097715 | 1.40016455 | -6.5062149 | 7.71E-11 |
| 70024 | MCM10 | 1601.68724 | 0.56557631 | 0.08697359 | 6.50285144 | 7.88E-11 |
| 75533 | NME5 | 25.0492148 | -3.7400945 | 0.57542121 | -6.4997508 | 8.05E-11 |
| 216858 | KCTD11 | 504.492209 | -1.0747638 | 0.16544309 | -6.4962749 | 8.23E-11 |

| | | | | | | |
|---|---|---|---|---|---|---|
| 18793 | PLAUR | 5989.55178 | 0.78754758 | 0.1214195 | 6.48617044 | 8.80E-11 |
| 15212 | HEXB | 1245.72293 | -0.607669 | 0.09370072 | -6.4852123 | 8.86E-11 |
| 73338 | ITPRIPL1 | 741.813498 | -0.7673495 | 0.11834467 | -6.4840227 | 8.93E-11 |
| 72258 | KCNK10 | 24.6829614 | 5.3046228 | 0.81815642 | 6.48362911 | 8.95E-11 |
| 68744 | Zfp740 | 1711.84048 | -0.6282869 | 0.09690555 | -6.4834979 | 8.96E-11 |
| 12368 | CASP6 | 366.52246 | -0.5845983 | 0.090168 | -6.483434 | 8.97E-11 |
| 94091 | TRIM11 | 1021.57927 | -0.7351727 | 0.11345363 | -6.4799402 | 9.18E-11 |
| 54123 | IRF7 | 221.635105 | -2.7782257 | 0.42879919 | -6.4790834 | 9.23E-11 |
| 230099 | Car9 | 45.8469025 | -3.9101351 | 0.60355194 | -6.4785394 | 9.26E-11 |
| 84111 | GPR87 | 14.6362247 | -7.3006048 | 1.12706424 | -6.477541 | 9.32E-11 |
| 15161 | HCFC1 | 9808.97452 | 0.43596081 | 0.0673054 | 6.47735287 | 9.33E-11 |
| 67870 | ENOPH1 | 1239.38253 | 0.38056378 | 0.05876562 | 6.47595951 | 9.42E-11 |
| 27784 | COMMD8 | 1476.79353 | -0.3414857 | 0.05275632 | -6.4728884 | 9.61E-11 |
| 69162 | SEC31A | 8861.33295 | 0.29810002 | 0.04606666 | 6.47105755 | 9.73E-11 |
| 66646 | RPE | 1657.36433 | -0.6320109 | 0.09767906 | -6.4702801 | 9.78E-11 |
| 213439 | GPR174 | 36.2916888 | -7.1581715 | 1.10664295 | -6.4683659 | 9.91E-11 |
| 21778 | TEX9 | 246.315285 | -0.7121913 | 0.11010937 | -6.468035 | 9.93E-11 |
| 237926 | RSAD1 | 324.026063 | 0.93370186 | 0.14438636 | 6.46669013 | 1.00E-10 |
| 269209 | STK36 | 99.5887496 | -1.175296 | 0.18179502 | -6.4649513 | 1.01E-10 |
| 18676 | PHF2 | 2077.61257 | 0.62442907 | 0.09659913 | 6.46412745 | 1.02E-10 |
| 14218 | SH3PXD2A | 9127.93994 | 0.83971519 | 0.12996039 | 6.46131624 | 1.04E-10 |
| 21987 | TPD52L1 | 47.6901397 | -3.3466057 | 0.51800268 | -6.4605952 | 1.04E-10 |
| 225010 | LCLAT1 | 1742.31472 | 0.46997995 | 0.0727681 | 6.45859839 | 1.06E-10 |
| 432572 | SPECC1 | 6589.05693 | 0.5076828 | 0.07863638 | 6.45608088 | 1.07E-10 |
| 545056 | Gm5801 | 338.741439 | 0.60778923 | 0.09416461 | 6.45453997 | 1.09E-10 |
| 27399 | IP6K1 | 2611.51173 | 0.4451109 | 0.06896445 | 6.45420792 | 1.09E-10 |
| 22110 | TSPYL1 | 2039.85304 | -0.4704748 | 0.07291786 | -6.4521201 | 1.10E-10 |
| 209131 | SNX30 | 1541.33411 | 2.12338926 | 0.32919615 | 6.45022517 | 1.12E-10 |
| 19656 | RBMXL1 | 428.877008 | 0.55181397 | 0.0855741 | 6.44837604 | 1.13E-10 |
| 231876 | LMTK2 | 2611.64743 | -0.8775374 | 0.13615388 | -6.4451885 | 1.15E-10 |
| 235130 | ADAMTS15 | 20.1278339 | -7.0290494 | 1.09061281 | -6.4450457 | 1.16E-10 |
| 93892 | Pcdhb21 | 89.6413432 | 2.98277492 | 0.46287929 | 6.44395842 | 1.16E-10 |
| 71952 | 2410016O06 | 879.489396 | 1.11569361 | 0.17317485 | 6.44258455 | 1.17E-10 |
| 20226 | SARS | 7124.47145 | -0.3208989 | 0.04981793 | -6.4414337 | 1.18E-10 |
| 236732 | RBM10 | 676.897486 | 0.6203525 | 0.09632032 | 6.44051559 | 1.19E-10 |
| 234878 | BC021891 | 83.2374066 | -9.8088383 | 1.52302908 | -6.4403486 | 1.19E-10 |
| 11666 | ABCD1 | 314.4505 | 0.74934593 | 0.11635761 | 6.44002483 | 1.19E-10 |
| 21402 | Skp1a | 3861.5938 | -0.2172419 | 0.03373416 | -6.4398198 | 1.20E-10 |
| 52065 | MFHAS1 | 1580.50003 | 1.05794623 | 0.16446414 | 6.43268649 | 1.25E-10 |
| 17476 | MPEG1 | 25.5679481 | -5.6581008 | 0.88023445 | -6.4279475 | 1.29E-10 |
| 70560 | WARS2 | 475.152572 | -0.6403522 | 0.09963516 | -6.42697 | 1.30E-10 |
| 70611 | FBXO33 | 1105.0239 | 0.88102011 | 0.13711326 | 6.42549144 | 1.31E-10 |
| 59048 | C1GALT1C1 | 606.133914 | -0.9912078 | 0.15426843 | -6.425215 | 1.32E-10 |
| 68947 | CHST8 | 12.8803811 | -7.1140639 | 1.10729135 | -6.4247444 | 1.32E-10 |
| 94253 | HECW1 | 44.0111035 | -8.8908048 | 1.38396001 | -6.4241775 | 1.33E-10 |

| | | | | | | |
|---|---|---|---|---|---|---|
| 217340 | RNF157 | 409.75452 | 1.06543657 | 0.16595932 | 6.41986602 | 1.36E-10 |
| 78249 | ADGRF4 | 189.912456 | -4.8415385 | 0.75462626 | -6.4158097 | 1.40E-10 |
| 98910 | USP6NL | 2592.95399 | -0.6716674 | 0.10471999 | -6.4139365 | 1.42E-10 |
| 100039781 | Hrct1 | 129.289842 | -2.6936452 | 0.41999094 | -6.4135794 | 1.42E-10 |
| 21969 | TOP1 | 4364.68796 | 0.42318565 | 0.06598354 | 6.4135033 | 1.42E-10 |
| 74551 | PCK2 | 1700.90982 | 1.15419259 | 0.17998208 | 6.41281964 | 1.43E-10 |
| 24000 | PTPN21 | 3905.04738 | 0.59010495 | 0.09202177 | 6.41266676 | 1.43E-10 |
| 72844 | KCTD17 | 1602.37539 | 0.8491375 | 0.132422 | 6.41235968 | 1.43E-10 |
| 14371 | FZD9 | 34.7786909 | 2.54543858 | 0.39700776 | 6.41155868 | 1.44E-10 |
| 102060 | GADD45GIP1 | 944.014017 | 0.62949883 | 0.09818593 | 6.4112939 | 1.44E-10 |
| 77300 | RAPH1 | 8184.7878 | -1.0228289 | 0.15955264 | -6.4106047 | 1.45E-10 |
| 68259 | IFT80 | 762.264862 | -0.7509233 | 0.11720486 | -6.4069292 | 1.48E-10 |
| 18408 | SLC25A15 | 301.07087 | 0.9081757 | 0.14184559 | 6.40256556 | 1.53E-10 |
| 69790 | MED30 | 1001.62315 | 0.83593099 | 0.13060537 | 6.40043362 | 1.55E-10 |
| 71371 | ARID5B | 2232.4 | 0.90554469 | 0.14152245 | 6.39859389 | 1.57E-10 |
| 12812 | COIL | 520.677637 | 0.69567418 | 0.1087269 | 6.39836313 | 1.57E-10 |
| 17527 | MPV17 | 839.968073 | -0.528251 | 0.08257683 | -6.3970855 | 1.58E-10 |
| 226255 | ATRNL1 | 1609.39048 | -0.8866154 | 0.13861175 | -6.3963942 | 1.59E-10 |
| 22262 | UOX | 12.3412808 | -7.0561305 | 1.10321456 | -6.395973 | 1.60E-10 |
| 11854 | RHOD | 904.39444 | -0.9591217 | 0.14996476 | -6.3956468 | 1.60E-10 |
| 240753 | PLEKHA6 | 20.1960311 | -4.7121735 | 0.73687744 | -6.394786 | 1.61E-10 |
| 268566 | GPHN | 1116.88531 | 0.59922 | 0.09377539 | 6.38994969 | 1.66E-10 |
| 74388 | DPP8 | 4074.90953 | 0.24281471 | 0.0380056 | 6.38891869 | 1.67E-10 |
| 14772 | GRK4 | 28.9591141 | -2.2147758 | 0.3466701 | -6.3887131 | 1.67E-10 |
| 215194 | KRI1 | 1253.78974 | 0.45510042 | 0.07124838 | 6.38751966 | 1.69E-10 |
| 56405 | DUSP14 | 415.650704 | -1.0107304 | 0.15824984 | -6.3869283 | 1.69E-10 |
| 57813 | TK2 | 373.443463 | -0.7572221 | 0.11856806 | -6.3863916 | 1.70E-10 |
| 17151 | CCNDBP1 | 651.953751 | -0.6649342 | 0.10413344 | -6.3854052 | 1.71E-10 |
| 71481 | alpk1 | 107.840092 | -1.3481291 | 0.21117295 | -6.3840049 | 1.73E-10 |
| 26372 | CLCN6 | 400.860657 | 0.75800217 | 0.11873662 | 6.38389564 | 1.73E-10 |
| 226351 | TMEM185B | 805.515823 | -0.6018553 | 0.09429166 | -6.3829109 | 1.74E-10 |
| 59007 | NGLY1 | 535.689484 | 0.49458082 | 0.07749168 | 6.38237319 | 1.74E-10 |
| 72338 | WDR89 | 207.642162 | 1.09304076 | 0.17129992 | 6.38085969 | 1.76E-10 |
| 56096 | PLAC1 | 22.1648337 | -4.5741844 | 0.71690993 | -6.3804171 | 1.77E-10 |
| 230393 | FOCAD | 844.400589 | 1.03113219 | 0.16166701 | 6.37812354 | 1.79E-10 |
| 24004 | RAI2 | 36.468106 | 3.13868996 | 0.49214024 | 6.37763325 | 1.80E-10 |
| 27494 | AMOT | 3167.78358 | 0.54460598 | 0.08540156 | 6.3770028 | 1.81E-10 |
| 17216 | MCM2 | 7459.98844 | 0.75947471 | 0.11911775 | 6.37583147 | 1.82E-10 |
| 100101484 | Trac | 17.1618714 | -4.718078 | 0.74001079 | -6.3756881 | 1.82E-10 |
| 319613 | SYBU | 115.808692 | 1.58074917 | 0.24804749 | 6.3727683 | 1.86E-10 |
| 545085 | WDR70 | 1061.1296 | 0.90505558 | 0.14204885 | 6.37143902 | 1.87E-10 |
| 12616 | CENPB | 8768.18893 | 0.94444012 | 0.1482735 | 6.36958134 | 1.90E-10 |
| 240057 | SYNGAP1 | 121.822507 | -1.1703466 | 0.18374032 | -6.3695688 | 1.90E-10 |
| 100040294 | Gm2694 | 20.0693358 | 7.07981038 | 1.11160964 | 6.36897172 | 1.90E-10 |
| 13363 | dhh | 57.9755403 | -6.2586672 | 0.98289369 | -6.3675932 | 1.92E-10 |

| | | | | | | |
|---|---|---|---|---|---|---|
| 66278 | SMIM20 | 307.80215 | -0.7932509 | 0.12460353 | -6.3661988 | 1.94E-10 |
| 72395 | 2610028E06I | 12.2817919 | 7.10242621 | 1.11572856 | 6.36572949 | 1.94E-10 |
| 22117 | TST | 187.885839 | -1.6148982 | 0.25368836 | -6.3656773 | 1.94E-10 |
| 54201 | Zfp316 | 449.540598 | -0.7699616 | 0.12098318 | -6.3642034 | 1.96E-10 |
| 263406 | PLEKHG3 | 2911.21051 | -0.3420634 | 0.05374887 | -6.3641047 | 1.96E-10 |
| 77777 | ULBP1 | 472.586169 | -1.7541728 | 0.27566913 | -6.3633269 | 1.97E-10 |
| 69480 | TTC9 | 390.817402 | -8.1154205 | 1.27549374 | -6.3625718 | 1.98E-10 |
| 67526 | ATG12 | 1409.71762 | -0.7099237 | 0.11161357 | -6.36055 | 2.01E-10 |
| 108657 | RNPEPL1 | 1251.98719 | -1.1782539 | 0.18533645 | -6.3573778 | 2.05E-10 |
| 14766 | ADGRG1 | 291.372285 | 2.1861385 | 0.34401393 | 6.35479646 | 2.09E-10 |
| 20846 | STAT1 | 1160.11823 | -0.8443205 | 0.13288236 | -6.353895 | 2.10E-10 |
| 22169 | CMPK2 | 52.8447099 | -2.5468754 | 0.40090884 | -6.3527544 | 2.11E-10 |
| 227290 | AAMP | 3036.68789 | -0.4283335 | 0.06747496 | -6.3480363 | 2.18E-10 |
| 21771 | CIRH1A | 3361.13467 | 0.5775661 | 0.09103848 | 6.34419728 | 2.24E-10 |
| 215705 | ARRDC1 | 542.073405 | -2.8242519 | 0.4454445 | -6.3403003 | 2.29E-10 |
| 56149 | GRASP | 440.21043 | 2.32910785 | 0.36736457 | 6.34004483 | 2.30E-10 |
| 105372 | UTP15 | 2088.54707 | 0.50393893 | 0.07949038 | 6.33962128 | 2.30E-10 |
| 12323 | CAMK2B | 17.127812 | -3.8220907 | 0.60331775 | -6.3351206 | 2.37E-10 |
| 434540 | Psg20 | 25.9464958 | -8.1259186 | 1.28279109 | -6.3345611 | 2.38E-10 |
| 668158 | CCDC85C | 546.993855 | 0.80329506 | 0.12684025 | 6.33312433 | 2.40E-10 |
| 20912 | STXBP3 | 1024.3743 | -0.726222 | 0.1146771 | -6.3327552 | 2.41E-10 |
| 53883 | CELSR2 | 30.1536195 | -5.8950201 | 0.93095641 | -6.3322192 | 2.42E-10 |
| 16145 | Igtp | 186.847972 | -2.6041136 | 0.41128291 | -6.3316844 | 2.42E-10 |
| 14739 | S1PR2 | 1787.40192 | 0.64583501 | 0.1020248 | 6.33017677 | 2.45E-10 |
| 140499 | UBE2J2 | 700.28059 | 0.72912186 | 0.11518602 | 6.3299509 | 2.45E-10 |
| 67958 | U2SURP | 3465.39828 | 0.43545503 | 0.06879674 | 6.32958857 | 2.46E-10 |
| 27386 | NPAS3 | 20.3983325 | -2.8300097 | 0.44734814 | -6.3261908 | 2.51E-10 |
| 54611 | PDE3A | 34.8233067 | 8.60347767 | 1.3601824 | 6.32523821 | 2.53E-10 |
| 68810 | NEXN | 169.948052 | 1.68704975 | 0.26672469 | 6.32506024 | 2.53E-10 |
| 69902 | MRTO4 | 616.912251 | 0.84981347 | 0.13438598 | 6.32367646 | 2.55E-10 |
| 237360 | ADAMTS14 | 394.887402 | 2.9735825 | 0.47034859 | 6.32208233 | 2.58E-10 |
| 17913 | MYO1C | 18502.4196 | -0.5049904 | 0.07989038 | -6.3210417 | 2.60E-10 |
| 244556 | Zfp791 | 28.2047041 | -2.6336256 | 0.41667792 | -6.3205309 | 2.61E-10 |
| 18181 | NRF1 | 1353.87582 | 0.37246799 | 0.05894277 | 6.31914584 | 2.63E-10 |
| 19049 | PPP1R1B | 19.0099786 | -7.6766171 | 1.21490758 | -6.318684 | 2.64E-10 |
| 72267 | LRRC8E | 83.8966553 | -2.192983 | 0.34715835 | -6.3169532 | 2.67E-10 |
| 380930 | 9330188P03I | 471.408939 | 1.59898423 | 0.25314861 | 6.31638558 | 2.68E-10 |
| 209737 | KIF15 | 1647.12362 | 0.68459916 | 0.10839096 | 6.31601699 | 2.68E-10 |
| 18174 | SLC11A2 | 3541.87412 | -0.4443797 | 0.07037667 | -6.3143045 | 2.71E-10 |
| 216197 | CKAP4 | 7438.58229 | 3.18732724 | 0.50483236 | 6.31363501 | 2.73E-10 |
| 12700 | CISH | 41.6247191 | -2.338509 | 0.37062293 | -6.3096718 | 2.80E-10 |
| 193043 | ZFP3 | 99.188788 | -1.5174623 | 0.24057159 | -6.3077369 | 2.83E-10 |
| 55988 | SNX12 | 2257.23499 | -0.4579705 | 0.07261136 | -6.3071461 | 2.84E-10 |
| 11479 | ACVR1B | 431.233148 | -0.8300855 | 0.13166727 | -6.3044177 | 2.89E-10 |
| 77125 | IL33 | 86.535861 | -2.9940444 | 0.47514322 | -6.3013515 | 2.95E-10 |

| | | | | | | |
|---|---|---|---|---|---|---|
| 240726 | SLCO5A1 | 27.4059005 | -7.4801201 | 1.18710775 | -6.3011299 | 2.95E-10 |
| 241062 | PGAP1 | 361.703825 | -0.7768825 | 0.12330257 | -6.300619 | 2.96E-10 |
| 65967 | EEFSEC | 733.251319 | 0.37034375 | 0.05878499 | 6.29997168 | 2.98E-10 |
| 170439 | ELOVL6 | 3574.95857 | 0.88429233 | 0.1404449 | 6.29636497 | 3.05E-10 |
| 232210 | HMCES | 766.536573 | 0.60713358 | 0.09642649 | 6.2963362 | 3.05E-10 |
| 71566 | CLMP | 1260.98074 | 8.09270229 | 1.28540317 | 6.29584748 | 3.06E-10 |
| 72039 | MCCC1 | 350.81043 | -0.7792282 | 0.12379614 | -6.2944466 | 3.08E-10 |
| 69129 | PEX11G | 59.4462109 | -1.6182184 | 0.25728819 | -6.2895167 | 3.18E-10 |
| 57257 | VAV3 | 590.810304 | -1.5052726 | 0.23938421 | -6.2881032 | 3.21E-10 |
| 11639 | AK4 | 505.725163 | 1.5684888 | 0.24950324 | 6.28644668 | 3.25E-10 |
| 100504191 | Gm16576 | 116.206798 | -0.8645442 | 0.1376124 | -6.2824587 | 3.33E-10 |
| 56284 | MRPL19 | 1248.75227 | 0.74440671 | 0.11851015 | 6.28137532 | 3.36E-10 |
| 210293 | dock10 | 53.0338505 | -7.738098 | 1.23204424 | -6.2806982 | 3.37E-10 |
| 69051 | PYCR2 | 1617.2789 | 0.3707395 | 0.05905462 | 6.27790836 | 3.43E-10 |
| 71934 | Car13 | 1789.00339 | -0.6086307 | 0.09698545 | -6.2754851 | 3.49E-10 |
| 14063 | F2RL1 | 82.9429277 | -3.2844786 | 0.52364605 | -6.2723258 | 3.56E-10 |
| 59038 | PXMP4 | 98.4200164 | -1.1834183 | 0.18867404 | -6.2722897 | 3.56E-10 |
| 77622 | APEX2 | 352.290337 | 0.56345533 | 0.08983254 | 6.27228518 | 3.56E-10 |
| 73694 | NDUFAF7 | 430.144644 | -0.5826451 | 0.09289682 | -6.2719598 | 3.57E-10 |
| 238377 | GPR68 | 11.1527402 | -6.9094239 | 1.10185308 | -6.2707307 | 3.59E-10 |
| 78512 | 3300005D01 | 314.051846 | 2.42448598 | 0.38688219 | 6.26672928 | 3.69E-10 |
| 81703 | JDP2 | 931.45199 | 1.11767356 | 0.17849226 | 6.26174821 | 3.81E-10 |
| 72429 | DNAJC25 | 534.727657 | 0.72618159 | 0.11601007 | 6.25964272 | 3.86E-10 |
| 213575 | DYNC2LI1 | 236.950154 | -1.415818 | 0.22620221 | -6.2590812 | 3.87E-10 |
| 17930 | MYOM2 | 22.5165956 | -5.4377781 | 0.86882011 | -6.2588078 | 3.88E-10 |
| 14170 | Fgf15 | 46.8420465 | -7.5414335 | 1.20541143 | -6.2563148 | 3.94E-10 |
| 17178 | FXYD3 | 27.0537409 | -8.1864192 | 1.30886542 | -6.254592 | 3.99E-10 |
| 319942 | A530016L24I | 24.4212091 | -2.743045 | 0.43876678 | -6.2517154 | 4.06E-10 |
| 11804 | APLP2 | 12818.0757 | -0.716833 | 0.11472115 | -6.2484816 | 4.14E-10 |
| 73732 | MUC16 | 11.3414258 | -6.9319567 | 1.10951888 | -6.247714 | 4.17E-10 |
| 66961 | NEAT1 | 6843.66116 | -2.147397 | 0.34379255 | -6.2462001 | 4.21E-10 |
| 14073 | FAAH | 122.528444 | -3.4244616 | 0.5483676 | -6.2448284 | 4.24E-10 |
| 69136 | TUSC1 | 59.5798881 | -7.8663095 | 1.25972126 | -6.2444843 | 4.25E-10 |
| 66163 | MRPL4 | 711.590458 | 0.67432192 | 0.10799233 | 6.24416489 | 4.26E-10 |
| 72149 | STRADA | 510.505994 | 0.60304265 | 0.0966389 | 6.24016445 | 4.37E-10 |
| 231801 | AGFG2 | 1073.16112 | -1.2474973 | 0.1999344 | -6.239533 | 4.39E-10 |
| 227618 | LRRC26 | 11.033674 | -6.893944 | 1.10534164 | -6.236935 | 4.46E-10 |
| 110454 | Ly6a | 1377.91228 | -3.5025563 | 0.56158652 | -6.2368953 | 4.46E-10 |
| 226122 | UBTD1 | 1466.26042 | 1.18133115 | 0.18943768 | 6.23598844 | 4.49E-10 |
| 52686 | Mettl2 | 653.753326 | -0.4703604 | 0.07542998 | -6.2357217 | 4.50E-10 |
| 19305 | PEX5 | 2392.37769 | 0.32841223 | 0.0526755 | 6.23462993 | 4.53E-10 |
| 217342 | UBE2O | 1894.08338 | 0.5884958 | 0.09440851 | 6.23350345 | 4.56E-10 |
| 107271 | YARS | 3457.91359 | 0.93130506 | 0.1494653 | 6.23091155 | 4.64E-10 |
| 73181 | NFATC4 | 482.794047 | 2.89191011 | 0.46420474 | 6.22981604 | 4.67E-10 |
| 30841 | KDM2B | 1025.87481 | 0.63654419 | 0.10217795 | 6.22976107 | 4.67E-10 |

| | | | | | | |
|---|---|---|---|---|---|---|
| 54353 | SKAP2 | 1227.95766 | -1.0839341 | 0.17401791 | -6.2288654 | 4.70E-10 |
| 68212 | TMBIM4 | 1394.35815 | -0.8999653 | 0.14452906 | -6.2268815 | 4.76E-10 |
| 23967 | OSR1 | 1419.46689 | 1.72448468 | 0.2770194 | 6.22514048 | 4.81E-10 |
| 22214 | UBE2H | 975.618052 | -1.0690174 | 0.17175359 | -6.2241344 | 4.84E-10 |
| 12894 | CPT1A | 3972.84419 | -0.8195608 | 0.13174057 | -6.2210207 | 4.94E-10 |
| 226123 | MORN4 | 437.585768 | -1.1194541 | 0.1799876 | -6.219618 | 4.98E-10 |
| 11782 | AP4S1 | 393.737743 | 1.21072683 | 0.19467208 | 6.21931431 | 4.99E-10 |
| 54451 | CPSF3 | 2188.04141 | 0.41801766 | 0.06722469 | 6.21821577 | 5.03E-10 |
| 17708 | COX1 | 65702.0454 | -0.7456176 | 0.11990864 | -6.2182142 | 5.03E-10 |
| 110557 | H2-Q6 | 18.7630725 | -3.6615871 | 0.58898526 | -6.2167721 | 5.07E-10 |
| 216150 | CDC34 | 2818.57383 | -0.6414185 | 0.10319743 | -6.2154501 | 5.12E-10 |
| 237775 | Zfp867 | 284.607847 | -0.7779176 | 0.12516479 | -6.2151468 | 5.13E-10 |
| 233893 | Zfp764 | 267.901501 | 0.81087721 | 0.13047951 | 6.21459418 | 5.15E-10 |
| 17904 | MYL6 | 1176.2726 | -1.0192496 | 0.16401713 | -6.2142876 | 5.16E-10 |
| 224023 | KLHL22 | 872.783936 | -2.3493903 | 0.37830286 | -6.2103426 | 5.29E-10 |
| 78787 | USP54 | 1010.85152 | -0.4312719 | 0.06949815 | -6.2055155 | 5.45E-10 |
| 171285 | HAVCR2 | 860.614641 | -4.2073234 | 0.6780603 | -6.2049399 | 5.47E-10 |
| 52318 | D1Ertd448e | 548.100803 | -0.9078715 | 0.14632213 | -6.2046084 | 5.48E-10 |
| 66446 | EXOSC7 | 552.683756 | 0.67355489 | 0.10856651 | 6.20407607 | 5.50E-10 |
| 407812 | Zfp941 | 135.801316 | 1.4871071 | 0.23973897 | 6.20302619 | 5.54E-10 |
| 226856 | LPGAT1 | 6348.20874 | 0.41452618 | 0.06683028 | 6.20266983 | 5.55E-10 |
| 114714 | RAD51C | 266.708835 | 0.67152899 | 0.10828257 | 6.20163534 | 5.59E-10 |
| 76863 | DCUN1D5 | 2149.10096 | 0.47533675 | 0.07666279 | 6.200358 | 5.63E-10 |
| 319670 | EML5 | 533.083657 | 2.23489763 | 0.36046382 | 6.2000609 | 5.64E-10 |
| 67549 | gpr89 | 996.810987 | -0.4893878 | 0.07893425 | -6.1999431 | 5.65E-10 |
| 66813 | BCL2L14 | 16.4400931 | -6.7393494 | 1.0875454 | -6.1968442 | 5.76E-10 |
| 13837 | EPHA3 | 38.4907171 | 5.67737675 | 0.91627668 | 6.19613801 | 5.79E-10 |
| 213211 | RNF26 | 352.555768 | 0.70708709 | 0.11418421 | 6.19251189 | 5.92E-10 |
| 12048 | BCL2L1 | 4824.84429 | -1.6110755 | 0.26019443 | -6.1918137 | 5.95E-10 |
| 231002 | PLEKHN1 | 208.679838 | -1.138682 | 0.18395324 | -6.1900621 | 6.01E-10 |
| 66244 | NEMF | 2135.76008 | 0.44752458 | 0.0723054 | 6.18936615 | 6.04E-10 |
| 242691 | GPATCH3 | 315.486659 | 0.61120742 | 0.09875818 | 6.1889294 | 6.06E-10 |
| 19704 | UPF1 | 4601.08459 | 0.2685 | 0.04339456 | 6.18741129 | 6.12E-10 |
| 100038541 | Gm10554 | 59.1211525 | 1.93148344 | 0.31218671 | 6.18694968 | 6.13E-10 |
| 73124 | GOLIM4 | 8847.19836 | -0.5065259 | 0.08187663 | -6.1864528 | 6.15E-10 |
| 105975 | AU022754 | 22.4960531 | -7.1900875 | 1.16233301 | -6.1859101 | 6.17E-10 |
| 380654 | CFAP54 | 18.4893453 | -3.2653272 | 0.52787591 | -6.1857856 | 6.18E-10 |
| 14634 | GLI3 | 2271.6408 | 1.08830584 | 0.17593858 | 6.18571439 | 6.18E-10 |
| 326623 | TNFSF15 | 17.2540689 | -6.8025806 | 1.09978462 | -6.1853752 | 6.20E-10 |
| 70757 | HACD2 | 474.220333 | -0.5143038 | 0.08315824 | -6.1846408 | 6.22E-10 |
| 74646 | SPSB1 | 648.447632 | -0.7381796 | 0.11938959 | -6.182948 | 6.29E-10 |
| 213084 | CDKL3 | 119.150718 | -1.3717074 | 0.22186017 | -6.1827566 | 6.30E-10 |
| 71473 | 8430423G03 | 12.5162414 | 7.12356125 | 1.15237714 | 6.18162319 | 6.34E-10 |
| 328274 | Zfp459 | 82.6975117 | -1.6092015 | 0.26044304 | -6.178708 | 6.46E-10 |
| 107250 | KAZALD1 | 94.3918211 | 1.82534463 | 0.2954856 | 6.17744031 | 6.51E-10 |

| | | | | | | |
|---|---|---|---|---|---|---|
| 29864 | RNF11 | 5776.87923 | -0.6678231 | 0.10810799 | -6.1773703 | 6.52E-10 |
| 320586 | A630089N07 | 71.8566883 | -1.8036811 | 0.29203932 | -6.1761586 | 6.57E-10 |
| 209012 | ULK4 | 94.4830988 | -1.1757636 | 0.19048695 | -6.17241 | 6.73E-10 |
| 67998 | FAM134C | 1490.46875 | -0.5378437 | 0.0871521 | -6.1713225 | 6.77E-10 |
| 217203 | TMEM106A | 222.298373 | -1.0095811 | 0.16360598 | -6.170808 | 6.79E-10 |
| 107373 | FAM111A | 5468.33653 | 0.50715029 | 0.08219648 | 6.16997563 | 6.83E-10 |
| 78334 | CDK19 | 1581.03479 | -1.1881978 | 0.19273374 | -6.1649705 | 7.05E-10 |
| 241289 | PPP1R26 | 40.5214059 | -3.5180776 | 0.57075372 | -6.1639155 | 7.10E-10 |
| 442799 | 9330154J02F | 58.5649041 | 1.30737484 | 0.21213112 | 6.1630507 | 7.14E-10 |
| 71790 | ANXA9 | 78.3271579 | -1.2231757 | 0.19853249 | -6.161086 | 7.22E-10 |
| 20348 | SEMA3C | 5384.0401 | -1.0513803 | 0.17065855 | -6.1607242 | 7.24E-10 |
| 21750 | TERF2 | 844.288943 | 0.35169671 | 0.05708711 | 6.16070258 | 7.24E-10 |
| 12716 | Ckmt1 | 11.3972585 | -6.9430577 | 1.12707268 | -6.1602573 | 7.26E-10 |
| 20873 | PLK4 | 1965.37551 | 0.45376085 | 0.07366514 | 6.1597773 | 7.28E-10 |
| 22350 | EZR | 5621.00487 | -1.4643249 | 0.23776846 | -6.158617 | 7.34E-10 |
| 68187 | FAM135A | 1707.52788 | -0.4415152 | 0.07170001 | -6.1578124 | 7.38E-10 |
| 69539 | TRNP1 | 40.1406796 | -1.7279707 | 0.28062613 | -6.1575545 | 7.39E-10 |
| 59016 | THAP11 | 910.171007 | 0.64961509 | 0.10550146 | 6.15740359 | 7.39E-10 |
| 238692 | Zfp874a | 102.681167 | -1.0604915 | 0.17229495 | -6.1550932 | 7.50E-10 |
| 15270 | H2AFX | 2420.57579 | 1.04069955 | 0.16914825 | 6.15258833 | 7.62E-10 |
| 94245 | DTNBP1 | 1308.3032 | 0.60182439 | 0.09782848 | 6.15183198 | 7.66E-10 |
| 214812 | Zfp609 | 2470.06748 | 0.49577634 | 0.08059273 | 6.15162573 | 7.67E-10 |
| 57028 | PDXP | 112.972651 | 1.45509144 | 0.23658485 | 6.1503999 | 7.73E-10 |
| 226849 | PPP2R5A | 1494.61241 | 0.48094568 | 0.07821665 | 6.14889104 | 7.80E-10 |
| 226844 | Mfsd7b | 1680.05247 | 0.32884889 | 0.0534813 | 6.14885734 | 7.80E-10 |
| 69893 | COA7 | 360.729778 | 0.53989588 | 0.08780609 | 6.14872959 | 7.81E-10 |
| 544864 | Gm5785 | 82.0244957 | 1.26568635 | 0.20586668 | 6.14808731 | 7.84E-10 |
| 72572 | SPATS2 | 1305.88677 | 0.71275656 | 0.11593772 | 6.14775369 | 7.86E-10 |
| 12343 | CAPZA2 | 6805.14703 | -0.2931531 | 0.04770017 | -6.1457469 | 7.96E-10 |
| 234463 | TMEM184C | 1569.71451 | -0.4211005 | 0.06853494 | -6.1443191 | 8.03E-10 |
| 97159 | A430005L14I | 323.784901 | 0.76371718 | 0.1243796 | 6.14021241 | 8.24E-10 |
| 13614 | EDN1 | 224.455968 | -5.4528717 | 0.88828265 | -6.1386674 | 8.32E-10 |
| 71514 | SFPQ | 11115.8824 | 0.7810585 | 0.12730428 | 6.13536699 | 8.50E-10 |
| 53978 | LPAR2 | 693.789536 | -1.239494 | 0.20204982 | -6.134596 | 8.54E-10 |
| 29871 | SCMH1 | 1098.61588 | 0.92327371 | 0.15052091 | 6.13385684 | 8.58E-10 |
| 72269 | CDA | 19.255086 | -6.9648121 | 1.1364886 | -6.1283607 | 8.88E-10 |
| 16430 | STT3A | 8692.14866 | 0.45756697 | 0.07469007 | 6.1262092 | 9.00E-10 |
| 67010 | RBM7 | 1657.94849 | 0.57163695 | 0.0933266 | 6.12512355 | 9.06E-10 |
| 217356 | tmc8 | 27.6726948 | -2.1231713 | 0.34677178 | -6.1226761 | 9.20E-10 |
| 20411 | sorbs1 | 1011.34195 | -1.6306457 | 0.26644043 | -6.1201136 | 9.35E-10 |
| 18073 | NID1 | 61847.2113 | 4.10931123 | 0.67145546 | 6.12000569 | 9.36E-10 |
| 18599 | PADI1 | 36.1760609 | -5.3383619 | 0.8729297 | -6.1154545 | 9.63E-10 |
| 67938 | MYL12B | 1701.84959 | -0.9321521 | 0.15245029 | -6.1144658 | 9.69E-10 |
| 68591 | MOCOS | 305.714868 | -2.7110576 | 0.4435397 | -6.1123224 | 9.82E-10 |
| 14312 | BRD2 | 10306.946 | 0.37239587 | 0.06093507 | 6.11135514 | 9.88E-10 |

| | | | | | | |
|---|---|---|---|---|---|---|
| 242253 | WDR63 | 16.64473 | 5.48898564 | 0.89817798 | 6.11124497 | 9.89E-10 |
| 76936 | HNRNPM | 10975.4616 | 0.3803559 | 0.06225767 | 6.10938196 | 1.00E-09 |
| 56277 | TMEM45A | 208.360389 | 5.46270409 | 0.89419762 | 6.10905686 | 1.00E-09 |
| 17181 | MATN2 | 412.034007 | 1.74928238 | 0.28646175 | 6.1065129 | 1.02E-09 |
| 234779 | PLCG2 | 1295.46493 | -1.8512564 | 0.30321179 | -6.1054894 | 1.02E-09 |
| 19144 | KLK6 | 101.12481 | -6.0137527 | 0.98506864 | -6.1049073 | 1.03E-09 |
| 57373 | AKIP1 | 259.188743 | -1.5166264 | 0.24843431 | -6.1047382 | 1.03E-09 |
| 68969 | EIF1B | 678.31582 | 0.63871353 | 0.10463058 | 6.10446315 | 1.03E-09 |
| 214112 | NIPAL4 | 26.8694234 | -3.3785697 | 0.55346533 | -6.1043927 | 1.03E-09 |
| 19092 | PRKG2 | 3273.86112 | 0.99861609 | 0.16361597 | 6.10341463 | 1.04E-09 |
| 69309 | SLC16A13 | 17.4001331 | 7.60362406 | 1.2462538 | 6.10118426 | 1.05E-09 |
| 56077 | DGKE | 507.72194 | -0.7978384 | 0.13090481 | -6.0947985 | 1.10E-09 |
| 27215 | AZI2 | 1172.47893 | 0.58942416 | 0.09679701 | 6.08928069 | 1.13E-09 |
| 76626 | MSI2 | 1256.80617 | -0.6433625 | 0.10565613 | -6.0892111 | 1.13E-09 |
| 213233 | TAPBPL | 342.773748 | -1.477396 | 0.2426909 | -6.0875625 | 1.15E-09 |
| 74002 | PSD2 | 51.6312862 | -7.1072045 | 1.16765901 | -6.0867123 | 1.15E-09 |
| 14119 | FBN2 | 16.5544144 | 3.67400479 | 0.60375357 | 6.08527215 | 1.16E-09 |
| 108067 | EIF2B3 | 880.967266 | 0.84921769 | 0.13955756 | 6.08507135 | 1.16E-09 |
| 26442 | PSMA5 | 552.905516 | 0.41883079 | 0.06884001 | 6.08411906 | 1.17E-09 |
| 19253 | PTPN18 | 15.8583686 | -6.6820432 | 1.09833514 | -6.0837926 | 1.17E-09 |
| 69668 | CCDC115 | 414.094556 | 0.51590935 | 0.08485084 | 6.08019161 | 1.20E-09 |
| 16876 | LHX9 | 20.8071501 | 3.56833869 | 0.58688402 | 6.08014291 | 1.20E-09 |
| 12371 | CASP9 | 461.794714 | 0.48349368 | 0.07952991 | 6.07939443 | 1.21E-09 |
| 76415 | FAM187B | 10.8834547 | -6.8776315 | 1.13131205 | -6.0793408 | 1.21E-09 |
| 72281 | SH2D4A | 117.289106 | -2.7862291 | 0.45834343 | -6.0789114 | 1.21E-09 |
| 94282 | SFXN5 | 156.372211 | 0.90879291 | 0.14954248 | 6.07715559 | 1.22E-09 |
| 338351 | AKAP17B | 183.329721 | -1.0953613 | 0.18028693 | -6.0756558 | 1.23E-09 |
| 13214 | DEFB1 | 14.1134113 | -7.247365 | 1.19310921 | -6.0743518 | 1.24E-09 |
| 381546 | CCDC24 | 77.0901429 | 3.11910827 | 0.51378952 | 6.07078992 | 1.27E-09 |
| 100862030 | #N/A | 10.3562079 | -6.8009 | 1.1203866 | -6.0701368 | 1.28E-09 |
| 67486 | polr3g | 494.122915 | 1.00770685 | 0.166012 | 6.07008428 | 1.28E-09 |
| 20501 | SLC16A1 | 6043.93892 | 0.58542957 | 0.09646803 | 6.06863816 | 1.29E-09 |
| 56386 | B4GALT6 | 1298.42898 | -0.9256453 | 0.15253384 | -6.0684588 | 1.29E-09 |
| 269023 | Zfp608 | 635.958014 | -0.534302 | 0.08811778 | -6.0634981 | 1.33E-09 |
| 68758 | ABHD11 | 617.616023 | -0.4824885 | 0.07958115 | -6.0628492 | 1.34E-09 |
| 66929 | ASF1B | 1440.00635 | 0.34834535 | 0.05746628 | 6.06173431 | 1.35E-09 |
| 51886 | FUBP1 | 6191.62909 | 1.13745747 | 0.18770463 | 6.05982636 | 1.36E-09 |
| 77048 | cep83 | 1230.76317 | 0.32000856 | 0.05281478 | 6.05907168 | 1.37E-09 |
| 77044 | ARID2 | 2504.42457 | 0.5583625 | 0.09219699 | 6.05618982 | 1.39E-09 |
| 16324 | INHBB | 273.843703 | -10.103578 | 1.66861189 | -6.0550799 | 1.40E-09 |
| 57425 | U90926 | 138.504077 | 4.99192382 | 0.82447477 | 6.05467143 | 1.41E-09 |
| 12062 | BDKRB2 | 223.386036 | -1.8407226 | 0.30403467 | -6.0543181 | 1.41E-09 |
| 77422 | C330018D20 | 225.829776 | -0.9638295 | 0.1592303 | -6.0530533 | 1.42E-09 |
| 69577 | fastkd3 | 595.427804 | 0.5637481 | 0.09313901 | 6.05276029 | 1.42E-09 |
| 18187 | NRP2 | 12678.1038 | 1.6149117 | 0.26681158 | 6.05262977 | 1.43E-09 |

| | | | | | | |
|---|---|---|---|---|---|---|
| 382639 | ZBTB42 | 175.117134 | -1.3863774 | 0.22912516 | -6.0507427 | 1.44E-09 |
| 66867 | HMG20A | 805.368632 | 0.49755431 | 0.08226463 | 6.04821673 | 1.46E-09 |
| 22289 | KDM6A | 1321.35197 | -0.6592585 | 0.10905453 | -6.0452182 | 1.49E-09 |
| 69325 | 1700012B09 | 14.1291731 | 6.56486935 | 1.08601544 | 6.04491343 | 1.49E-09 |
| 56516 | RBMS2 | 2778.97887 | -0.5128901 | 0.08490178 | -6.0409818 | 1.53E-09 |
| 18858 | PMP22 | 1322.09329 | -0.3603561 | 0.05966306 | -6.0398527 | 1.54E-09 |
| 67795 | RNLS | 13.097785 | -7.1373679 | 1.18204497 | -6.0381526 | 1.56E-09 |
| 104383 | rcor2 | 396.607261 | 3.17581342 | 0.52598086 | 6.0378878 | 1.56E-09 |
| 238123 | COG5 | 619.18148 | 0.50239756 | 0.08326424 | 6.03377384 | 1.60E-09 |
| 84036 | KCNN1 | 47.6028537 | -3.5410098 | 0.58712116 | -6.0311397 | 1.63E-09 |
| 17155 | Man1a | 1736.15418 | 0.86656411 | 0.14370175 | 6.03029623 | 1.64E-09 |
| 18600 | PADI2 | 1063.26648 | -3.2922993 | 0.54606072 | -6.0291817 | 1.65E-09 |
| 75901 | DCP1A | 1243.7492 | 0.34778031 | 0.05775048 | 6.02212021 | 1.72E-09 |
| 109331 | RNF20 | 2418.3673 | 0.6577256 | 0.1092487 | 6.0204432 | 1.74E-09 |
| 117198 | IVNS1ABP | 5949.20951 | -0.6763381 | 0.11234343 | -6.0202726 | 1.74E-09 |
| 28106 | MYDGF | 1763.60944 | 0.36275087 | 0.06026122 | 6.01964058 | 1.75E-09 |
| 84004 | MCAM | 96.5773126 | -2.1943636 | 0.36455899 | -6.0192278 | 1.75E-09 |
| 121021 | CSPG4 | 5180.87598 | 9.0929479 | 1.51140689 | 6.01621442 | 1.79E-09 |
| 353172 | GARS | 9641.25352 | 0.39044554 | 0.06489975 | 6.01613287 | 1.79E-09 |
| 71862 | gpr160 | 14.2347106 | -6.5300892 | 1.08546945 | -6.0159125 | 1.79E-09 |
| 78748 | RASSF10 | 178.096067 | -7.4540118 | 1.23909952 | -6.0156684 | 1.79E-09 |
| 17524 | MPP1 | 3285.90136 | -0.99105 | 0.16478957 | -6.0140335 | 1.81E-09 |
| 67495 | TMEM167B | 1180.09376 | -0.5446118 | 0.09061092 | -6.0104439 | 1.85E-09 |
| 74008 | ARSG | 47.6628691 | -1.4895814 | 0.24786122 | -6.0097396 | 1.86E-09 |
| 13874 | EREG | 5441.93615 | 7.0330502 | 1.17104408 | 6.0057946 | 1.90E-09 |
| 72462 | RRP1B | 1598.16222 | 0.39436313 | 0.06566877 | 6.00533726 | 1.91E-09 |
| 100503607 | BB319198 | 14.7504749 | 6.62932376 | 1.1045657 | 6.00174691 | 1.95E-09 |
| 16666 | KRT16 | 97.5919971 | -1.8084535 | 0.30134438 | -6.0012852 | 1.96E-09 |
| 69716 | TRIP13 | 1459.79226 | 0.87297636 | 0.14547859 | 6.00072065 | 1.96E-09 |
| 11749 | ANXA6 | 4980.43422 | -2.251503 | 0.37528656 | -5.9994235 | 1.98E-09 |
| 228491 | Zfp770 | 677.906046 | 0.49387422 | 0.0823369 | 5.99821234 | 2.00E-09 |
| 52163 | CAMK1 | 1102.44197 | -1.0304948 | 0.17181497 | -5.9977009 | 2.00E-09 |
| 18521 | PCBP2 | 10151.288 | -0.6130607 | 0.1022303 | -5.9968587 | 2.01E-09 |
| 230678 | tmem125 | 15.1171264 | -6.6113358 | 1.1027926 | -5.9950853 | 2.03E-09 |
| 11950 | ATP5F1 | 2433.51967 | -0.326889 | 0.05455775 | -5.9916139 | 2.08E-09 |
| 18406 | ORM2 | 43.5540093 | -8.875772 | 1.48172426 | -5.9901644 | 2.10E-09 |
| 69163 | MRPL44 | 633.118006 | -0.4697344 | 0.07844635 | -5.9879702 | 2.12E-09 |
| 56412 | NOA1 | 1115.49891 | 0.39084959 | 0.06527325 | 5.98789795 | 2.13E-09 |
| 414108 | 9230114K14l | 166.307938 | -1.7220729 | 0.28763571 | -5.9869929 | 2.14E-09 |
| 14057 | SFXN1 | 3099.5263 | 0.77273283 | 0.12908292 | 5.9863291 | 2.15E-09 |
| 245877 | MAP7D1 | 7441.84237 | 0.98843476 | 0.16514816 | 5.9851395 | 2.16E-09 |
| 243923 | RGS9BP | 15.4667152 | 3.1344336 | 0.52397409 | 5.98203933 | 2.20E-09 |
| 233902 | FBXL19 | 1297.86362 | 0.93607006 | 0.156486 | 5.98181359 | 2.21E-09 |
| 223649 | NRBP2 | 134.103559 | -1.114284 | 0.18628149 | -5.9817213 | 2.21E-09 |
| 67457 | FRMD8 | 3556.04134 | -1.3560204 | 0.22679151 | -5.9791495 | 2.24E-09 |

| | | | | | | |
|---|---|---|---|---|---|---|
| 56724 | CRIPT | 809.784653 | -0.4102466 | 0.06861593 | -5.978882 | 2.25E-09 |
| 18675 | PHEX | 298.75913 | 2.10569652 | 0.35226432 | 5.97760368 | 2.26E-09 |
| 69367 | GLRX2 | 392.615852 | -0.5272345 | 0.08822842 | -5.9757898 | 2.29E-09 |
| 109294 | PREX2 | 52.447756 | -7.7050558 | 1.28940284 | -5.9756777 | 2.29E-09 |
| 19225 | PTGS2 | 20990.4807 | 1.6377352 | 0.27407152 | 5.97557595 | 2.29E-09 |
| 20931 | SURF2 | 805.771477 | 0.4965818 | 0.08313585 | 5.97313705 | 2.33E-09 |
| 320253 | 41700 | 48.9230706 | -1.8456043 | 0.30905588 | -5.9717493 | 2.35E-09 |
| 75564 | RSPH9 | 10.0412304 | -6.7554503 | 1.13134062 | -5.9711905 | 2.36E-09 |
| 225912 | CYB561A3 | 720.421458 | -2.0766227 | 0.34781348 | -5.9705068 | 2.37E-09 |
| 67103 | PTGR1 | 2574.47046 | 2.18980665 | 0.36688029 | 5.96872254 | 2.39E-09 |
| 18400 | SLC22A18 | 24.9072452 | -2.9207467 | 0.4893788 | -5.9682738 | 2.40E-09 |
| 71883 | COQ2 | 349.317485 | -0.5805812 | 0.09728924 | -5.9675782 | 2.41E-09 |
| 74549 | MAU2 | 1662.1285 | -0.4080155 | 0.06837213 | -5.9675704 | 2.41E-09 |
| 74764 | KLC4 | 226.994516 | -1.031356 | 0.17283159 | -5.9674041 | 2.41E-09 |
| 211673 | ARFGEF1 | 5198.25804 | -0.5775978 | 0.09679524 | -5.9672124 | 2.41E-09 |
| 68166 | SPIRE1 | 1747.88869 | 1.21272883 | 0.20324318 | 5.96688569 | 2.42E-09 |
| 100972 | RAB28 | 945.249028 | -0.4347696 | 0.07287459 | -5.9659969 | 2.43E-09 |
| 15505 | HSPH1 | 4574.87256 | 0.96401504 | 0.16159081 | 5.96577903 | 2.43E-09 |
| 239606 | SLC2A13 | 126.725098 | 3.40832688 | 0.57131949 | 5.96571081 | 2.44E-09 |
| 67006 | CISD2 | 1979.01429 | 0.41691529 | 0.06989267 | 5.96507933 | 2.45E-09 |
| 629777 | Zfp335os | 55.502328 | 1.31840527 | 0.22107496 | 5.96361201 | 2.47E-09 |
| 29812 | NDRG3 | 877.799192 | 0.59315849 | 0.09948777 | 5.96212461 | 2.49E-09 |
| 107976 | BRE | 631.206153 | -0.6409512 | 0.10751834 | -5.9613196 | 2.50E-09 |
| 22325 | VAV2 | 705.694772 | -0.5360269 | 0.08992037 | -5.9611286 | 2.51E-09 |
| 66469 | FAM213B | 12.4559892 | -7.0733705 | 1.18667522 | -5.9606625 | 2.51E-09 |
| 11834 | AQR | 2781.89597 | 0.32095957 | 0.0538828 | 5.95662434 | 2.58E-09 |
| 17474 | CLEC4D | 269.161282 | 3.28622278 | 0.5517116 | 5.95641413 | 2.58E-09 |
| 69786 | TPRKB | 1449.4069 | 0.64978558 | 0.10913374 | 5.95403022 | 2.62E-09 |
| 20688 | SP4 | 370.518235 | 1.50764814 | 0.2533341 | 5.95122456 | 2.66E-09 |
| 80708 | PACSIN3 | 1044.63151 | -1.3649729 | 0.22939378 | -5.9503483 | 2.68E-09 |
| 12367 | CASP3 | 2291.08286 | -0.8826618 | 0.1483577 | -5.9495517 | 2.69E-09 |
| 212167 | GSAP | 182.638768 | -1.2097579 | 0.20336552 | -5.9486874 | 2.70E-09 |
| 20229 | SAT1 | 1714.75541 | -1.0015494 | 0.1683967 | -5.9475597 | 2.72E-09 |
| 66916 | NDUFB7 | 1040.902 | -0.4386482 | 0.07376287 | -5.946735 | 2.74E-09 |
| 11886 | ASAH1 | 5452.68696 | -1.0862417 | 0.18268441 | -5.9460008 | 2.75E-09 |
| 56016 | HEBP2 | 412.915482 | -0.8289669 | 0.13944357 | -5.9448195 | 2.77E-09 |
| 68743 | ANLN | 17401.0988 | 0.70939582 | 0.11934712 | 5.94397112 | 2.78E-09 |
| 68703 | RERE | 3704.00823 | 0.65155085 | 0.10964547 | 5.94234173 | 2.81E-09 |
| 23821 | BACE1 | 2092.2974 | -0.5284498 | 0.08894493 | -5.9413147 | 2.83E-09 |
| 68066 | SLC25A39 | 4141.48128 | -0.239199 | 0.04026701 | -5.9403218 | 2.84E-09 |
| 66090 | YPEL3 | 147.126473 | -1.1167127 | 0.1880172 | -5.939418 | 2.86E-09 |
| 98970 | FIBCD1 | 62.3212602 | 3.94564541 | 0.66439325 | 5.93871988 | 2.87E-09 |
| 241688 | DZANK1 | 18.325573 | -3.6190119 | 0.6094139 | -5.9385123 | 2.88E-09 |
| 21981 | PPP1R13B | 357.132024 | -0.9813687 | 0.16530276 | -5.9367955 | 2.91E-09 |
| 55932 | GBP3 | 147.737451 | -2.0439895 | 0.34440812 | -5.9347891 | 2.94E-09 |

| | | | | | | |
|---|---|---|---|---|---|---|
| 18472 | PAFAH1B1 | 7261.60462 | -0.2333738 | 0.03932584 | -5.9343618 | 2.95E-09 |
| 56013 | SRCIN1 | 56.3210275 | -7.811231 | 1.31659671 | -5.9328957 | 2.98E-09 |
| 15904 | ID4 | 34.0657278 | -4.085713 | 0.68880245 | -5.931618 | 3.00E-09 |
| 320174 | A830082K12 | 103.091667 | 1.27747405 | 0.21540135 | 5.93066872 | 3.02E-09 |
| 16498 | KCNAB2 | 3579.98266 | 7.59772779 | 1.28113863 | 5.93044938 | 3.02E-09 |
| 56397 | MORF4L2 | 5129.90934 | 0.46398242 | 0.07828951 | 5.92649507 | 3.09E-09 |
| 320469 | 9930014A18 | 258.194586 | -0.8725896 | 0.14724839 | -5.9259705 | 3.10E-09 |
| 110948 | HLCS | 417.977229 | -0.6214377 | 0.10486689 | -5.9259663 | 3.10E-09 |
| 239556 | CACNA1I | 18.2001921 | -7.6134237 | 1.28526554 | -5.9236192 | 3.15E-09 |
| 75735 | PANK1 | 617.85788 | 0.62527286 | 0.10557113 | 5.92276363 | 3.17E-09 |
| 53599 | CD164 | 9780.75969 | -0.1542037 | 0.02603733 | -5.922407 | 3.17E-09 |
| 22026 | NR2C2 | 2729.13026 | 0.54583484 | 0.09217865 | 5.92148906 | 3.19E-09 |
| 75178 | MEIOB | 13.480736 | -6.4503998 | 1.08942873 | -5.9209012 | 3.20E-09 |
| 15894 | ICAM1 | 33.0689303 | -6.0465812 | 1.02149385 | -5.9193515 | 3.23E-09 |
| 18769 | PKIG | 455.961781 | -0.5856197 | 0.0990199 | -5.9141619 | 3.34E-09 |
| 16414 | ITGB2 | 113.257516 | -3.5668049 | 0.60330998 | -5.9120602 | 3.38E-09 |
| 18854 | PML | 1617.02823 | -0.8786599 | 0.14866559 | -5.910311 | 3.41E-09 |
| 13512 | DSG3 | 19.4991743 | -7.7138149 | 1.30526202 | -5.9097827 | 3.43E-09 |
| 69917 | NABP2 | 1241.3475 | -0.3256366 | 0.05510836 | -5.9090241 | 3.44E-09 |
| 112407 | EGLN3 | 1567.62198 | -2.2033303 | 0.3728872 | -5.9088388 | 3.45E-09 |
| 20111 | RPS6KA1 | 834.605406 | -1.3301069 | 0.22522085 | -5.9057895 | 3.51E-09 |
| 332175 | ZDHHC23 | 13.2384975 | -6.4198691 | 1.08704733 | -5.9057862 | 3.51E-09 |
| 66904 | PCCB | 551.496802 | 0.64311873 | 0.10890009 | 5.90558514 | 3.51E-09 |
| 68011 | SNRPG | 316.79266 | 0.85471791 | 0.14473141 | 5.90554533 | 3.51E-09 |
| 22218 | SUMO1 | 379.983603 | -0.7644123 | 0.12945197 | -5.9049874 | 3.53E-09 |
| 74091 | NPL | 37.7132083 | 2.1071965 | 0.35699976 | 5.90251525 | 3.58E-09 |
| 81000 | RAD54L2 | 2083.95093 | 0.65258583 | 0.11059078 | 5.90090613 | 3.62E-09 |
| 76850 | AGO4 | 55.6508371 | -1.2033908 | 0.20399822 | -5.8990258 | 3.66E-09 |
| 14561 | GDF11 | 580.846494 | 1.39371324 | 0.23626637 | 5.89890656 | 3.66E-09 |
| 69834 | RAB43 | 1975.78831 | 0.84916927 | 0.14400779 | 5.89668972 | 3.71E-09 |
| 22235 | UGDH | 6053.75512 | 1.00088945 | 0.16979507 | 5.89469095 | 3.75E-09 |
| 27056 | IRF5 | 99.728133 | 2.42536576 | 0.41146055 | 5.89452809 | 3.76E-09 |
| 80861 | DHX58 | 110.486206 | -1.1418495 | 0.19371922 | -5.8943534 | 3.76E-09 |
| 12124 | BIK | 24.2846503 | -5.5509639 | 0.94221402 | -5.8914045 | 3.83E-09 |
| 56635 | Prl2a1 | 19.1354288 | -6.9601558 | 1.18150876 | -5.8909049 | 3.84E-09 |
| 108907 | NUSAP1 | 1984.50228 | -0.2677213 | 0.04545901 | -5.8892892 | 3.88E-09 |
| 22652 | MKRN3 | 14.0965726 | -6.510555 | 1.10578131 | -5.887742 | 3.92E-09 |
| 56489 | IKBKE | 600.074302 | -2.0798396 | 0.35325279 | -5.8876804 | 3.92E-09 |
| 194231 | CNKSR1 | 256.936109 | -1.9172935 | 0.32569817 | -5.8867187 | 3.94E-09 |
| 13390 | DLX1 | 178.185781 | 2.083861 | 0.35399677 | 5.88666679 | 3.94E-09 |
| 27392 | PIGN | 3614.22453 | -0.4239246 | 0.07204058 | -5.8845254 | 3.99E-09 |
| 14791 | EMG1 | 2177.42109 | 0.58705917 | 0.09979099 | 5.88288758 | 4.03E-09 |
| 15399 | HOXA2 | 46.9340538 | -2.715405 | 0.4616114 | -5.8824479 | 4.04E-09 |
| 71833 | DCAF7 | 5013.12294 | -0.3799157 | 0.06460907 | -5.880223 | 4.10E-09 |
| 67789 | DALRD3 | 632.219393 | -0.6758174 | 0.11497491 | -5.877956 | 4.15E-09 |

| | | | | | | |
|---|---|---|---|---|---|---|
| 18407 | Orm3 | 10.2382811 | -6.7897717 | 1.15530895 | -5.8770181 | 4.18E-09 |
| 70497 | ARHGAP17 | 1321.10522 | 0.6032133 | 0.10269966 | 5.87356669 | 4.27E-09 |
| 329152 | HECW2 | 35.132227 | -3.0204779 | 0.51437147 | -5.8721723 | 4.30E-09 |
| 234852 | CHMP1A | 2173.19788 | -0.5440535 | 0.09268389 | -5.8699901 | 4.36E-09 |
| 269695 | RNFT2 | 65.5811114 | -1.3370625 | 0.2278712 | -5.8676242 | 4.42E-09 |
| 67765 | 5830432E09I | 18.6980055 | 6.24273828 | 1.06427264 | 5.8657322 | 4.47E-09 |
| 71972 | dnmbp | 920.496794 | 0.88208141 | 0.1503854 | 5.86547248 | 4.48E-09 |
| 210719 | MKX | 280.68415 | 0.91967461 | 0.15680761 | 5.86498698 | 4.49E-09 |
| 23886 | GDF15 | 32.0976276 | -2.2938025 | 0.39111892 | -5.8647189 | 4.50E-09 |
| 216441 | SLC26A10 | 19.5675848 | -5.2690273 | 0.89854363 | -5.8639638 | 4.52E-09 |
| 280411 | LIX1L | 1637.74863 | 1.19953874 | 0.20466441 | 5.86100298 | 4.60E-09 |
| 104943 | FAM110C | 1215.00616 | 1.09673476 | 0.18715964 | 5.85988929 | 4.63E-09 |
| 12814 | COL11A1 | 585.497395 | -10.553434 | 1.80132269 | -5.8587139 | 4.66E-09 |
| 13036 | CTSH | 1367.68856 | -1.9387331 | 0.33094619 | -5.8581522 | 4.68E-09 |
| 26412 | MAP4K2 | 295.891664 | -1.2582761 | 0.21479997 | -5.8578969 | 4.69E-09 |
| 11772 | AP2A2 | 5740.30333 | 0.78021523 | 0.13323854 | 5.85577747 | 4.75E-09 |
| 22187 | UBB | 2366.49175 | -0.5709055 | 0.09752661 | -5.8538432 | 4.80E-09 |
| 15558 | HTR2A | 63.3737236 | 5.35602018 | 0.91525538 | 5.8519407 | 4.86E-09 |
| 17918 | MYO5A | 7831.38557 | 0.82157366 | 0.14051602 | 5.84683262 | 5.01E-09 |
| 227737 | FAM129B | 17327.1852 | -1.0586705 | 0.18107409 | -5.8466148 | 5.02E-09 |
| 66174 | NUDT14 | 178.091396 | -1.7781215 | 0.30428569 | -5.8435923 | 5.11E-09 |
| 545062 | Gm5802 | 14.3282269 | -4.4399105 | 0.75979848 | -5.8435369 | 5.11E-09 |
| 23872 | ETS2 | 3005.66845 | -1.1973302 | 0.20489968 | -5.8434948 | 5.11E-09 |
| 67978 | TCTN2 | 290.039717 | -1.1237685 | 0.19233125 | -5.8428804 | 5.13E-09 |
| 13685 | EIF4EBP1 | 2585.02035 | 0.36188092 | 0.06195175 | 5.8413346 | 5.18E-09 |
| 13838 | EPHA4 | 101.359453 | -3.3124452 | 0.56707118 | -5.8413217 | 5.18E-09 |
| 17301 | FOXD2 | 37.4767284 | 1.60278137 | 0.27441576 | 5.84070456 | 5.20E-09 |
| 14726 | PDPN | 4536.42519 | 8.06661198 | 1.38111795 | 5.84063944 | 5.20E-09 |
| 408066 | BC067074 | 47.0931619 | -3.7358468 | 0.63967642 | -5.8402133 | 5.21E-09 |
| 72823 | PARD3B | 30.9110105 | -2.6745911 | 0.45807355 | -5.8387809 | 5.26E-09 |
| 18046 | NFYC | 294.911597 | 1.11768747 | 0.19147066 | 5.83738259 | 5.30E-09 |
| 105083 | PELO | 936.682614 | 0.76686695 | 0.13143677 | 5.83449343 | 5.40E-09 |
| 68346 | SIRT5 | 83.053394 | -1.3670553 | 0.2344723 | -5.8303486 | 5.53E-09 |
| 75578 | FGGY | 170.869925 | -2.9784229 | 0.51103969 | -5.8281637 | 5.60E-09 |
| 13528 | DTNB | 1585.6744 | -0.5155769 | 0.08847341 | -5.8274783 | 5.63E-09 |
| 14205 | VEGFD | 450.244098 | 2.7497767 | 0.4720069 | 5.82571294 | 5.69E-09 |
| 18701 | PIGF | 427.465629 | -0.7363849 | 0.12642638 | -5.8246139 | 5.72E-09 |
| 15007 | H2-Q10 | 211.844171 | -1.3705609 | 0.23531294 | -5.8244178 | 5.73E-09 |
| 320064 | D130017N08 | 19.3197056 | -5.2119953 | 0.89506362 | -5.8230445 | 5.78E-09 |
| 72230 | Zfp558 | 50.8699225 | -1.8598341 | 0.31963875 | -5.8185501 | 5.94E-09 |
| 19288 | PTX3 | 833.608437 | 7.30706432 | 1.25591566 | 5.81811703 | 5.95E-09 |
| 52331 | STBD1 | 533.660686 | -2.3771577 | 0.40860124 | -5.8177935 | 5.96E-09 |
| 330277 | FAM71F1 | 34.8256741 | 1.555814 | 0.2674408 | 5.81741457 | 5.98E-09 |
| 19044 | PPOX | 307.885549 | -0.829306 | 0.14261296 | -5.8150815 | 6.06E-09 |
| 435145 | SHISA8 | 16.072787 | 7.48884922 | 1.28814745 | 5.81365836 | 6.11E-09 |

| | | | | | | |
|---|---|---|---|---|---|---|
| 108699 | CHN1 | 42.0133302 | 1.40054263 | 0.2411316 | 5.80820868 | 6.31E-09 |
| 18710 | PIK3R3 | 591.88661 | -1.0295191 | 0.17725893 | -5.8079953 | 6.32E-09 |
| 216825 | USP22 | 3228.8589 | -0.6161348 | 0.10610747 | -5.8067052 | 6.37E-09 |
| 71795 | PITPNC1 | 536.736559 | -1.2654193 | 0.21793332 | -5.8064517 | 6.38E-09 |
| 320187 | E230001N04 | 15.4407401 | -4.5472809 | 0.78314526 | -5.8064336 | 6.38E-09 |
| 52589 | NCALD | 16.7949188 | -5.0085995 | 0.86276618 | -5.8052803 | 6.43E-09 |
| 12268 | c4b | 13.6330351 | 3.5331385 | 0.60894673 | 5.80204853 | 6.55E-09 |
| 320129 | GRK3 | 15.2146208 | -6.6276133 | 1.14282235 | -5.7993382 | 6.66E-09 |
| 20482 | SKIL | 4849.92847 | -0.6659416 | 0.11485652 | -5.7980311 | 6.71E-09 |
| 14118 | FBN1 | 2498.36202 | 4.77771078 | 0.82405084 | 5.79783499 | 6.72E-09 |
| 228807 | Zfp341 | 195.335097 | 0.831744 | 0.1435114 | 5.79566519 | 6.81E-09 |
| 101604 | E430018J23F | 149.366796 | 0.98364528 | 0.16972995 | 5.79535494 | 6.82E-09 |
| 66895 | PXDC1 | 1226.46752 | 0.92788398 | 0.16012023 | 5.79492032 | 6.84E-09 |
| 68165 | FDX1L | 564.951659 | 0.5775627 | 0.09969169 | 5.79348871 | 6.89E-09 |
| 75062 | SF3A3 | 2034.03887 | 0.9463938 | 0.16338518 | 5.79240909 | 6.94E-09 |
| 14871 | GstT1 | 19.9453381 | -5.6777165 | 0.98041619 | -5.7911289 | 6.99E-09 |
| 56378 | ARPC3 | 3098.72615 | -0.4293936 | 0.07414731 | -5.7910878 | 6.99E-09 |
| 67621 | BEND5 | 25.3384995 | -3.3746294 | 0.58291569 | -5.7892238 | 7.07E-09 |
| 321022 | CDV3 | 4062.13949 | 0.4689769 | 0.08101269 | 5.78893157 | 7.08E-09 |
| 114863 | PROSC | 2121.85321 | -0.4854971 | 0.08387004 | -5.7886837 | 7.09E-09 |
| 69768 | 1600019K03I | 18.467364 | 6.22108423 | 1.07478661 | 5.78820405 | 7.11E-09 |
| 68276 | TOE1 | 759.625227 | 0.82040323 | 0.1417812 | 5.78640353 | 7.19E-09 |
| 72584 | CUL4B | 4327.03536 | -0.2802191 | 0.04843169 | -5.7858625 | 7.21E-09 |
| 238130 | DOCK4 | 956.679159 | -2.0997065 | 0.36293098 | -5.7854154 | 7.23E-09 |
| 76787 | PPFIA3 | 55.1785282 | -2.826673 | 0.48922434 | -5.7778666 | 7.57E-09 |
| 93884 | PCDHB13 | 20.1133519 | 3.84440725 | 0.66549142 | 5.77679463 | 7.61E-09 |
| 69077 | PSMD11 | 2236.68594 | 0.51444608 | 0.08907175 | 5.77563702 | 7.67E-09 |
| 12161 | BMP6 | 30.7358959 | 2.55660491 | 0.44271354 | 5.77485138 | 7.70E-09 |
| 72433 | RAB38 | 21.6901982 | -5.7989965 | 1.00418696 | -5.7748176 | 7.70E-09 |
| 70356 | ST13 | 7153.74584 | -0.189613 | 0.03283798 | -5.7741987 | 7.73E-09 |
| 74145 | F13A1 | 31.3072116 | 2.94091041 | 0.50937662 | 5.77354807 | 7.76E-09 |
| 56535 | PEX3 | 575.390004 | -0.4412046 | 0.0764353 | -5.7722616 | 7.82E-09 |
| 75767 | RAB11FIP1 | 1399.99666 | -1.1124098 | 0.19275836 | -5.7710067 | 7.88E-09 |
| 56095 | FTSJ3 | 5137.14601 | 0.27327435 | 0.04737179 | 5.76871525 | 7.99E-09 |
| 22785 | SLC30A4 | 9354.02417 | 1.29625499 | 0.22470497 | 5.7686975 | 7.99E-09 |
| 67492 | ZFAND4 | 205.107359 | 1.64726276 | 0.28567941 | 5.76612348 | 8.11E-09 |
| 11541 | ADORA2B | 34.8433204 | 2.57140173 | 0.44601963 | 5.765221 | 8.16E-09 |
| 74015 | FCHO1 | 13.9959751 | -7.2398944 | 1.25637249 | -5.7625381 | 8.29E-09 |
| 100336 | PPP1R8 | 1810.08565 | 0.70908851 | 0.12309741 | 5.76038511 | 8.39E-09 |
| 93711 | PCDHGA3 | 40.7654672 | -2.6099735 | 0.45322033 | -5.75873 | 8.47E-09 |
| 72103 | APLF | 486.802424 | 0.54106618 | 0.09395985 | 5.75848287 | 8.49E-09 |
| 23792 | ADAM23 | 40.592397 | 5.06344141 | 0.87940666 | 5.75779291 | 8.52E-09 |
| 103711 | PNPO | 301.730367 | -0.9335312 | 0.16218172 | -5.7560818 | 8.61E-09 |
| 72630 | HSPA12B | 19.2982666 | 4.43238266 | 0.77005655 | 5.75591842 | 8.62E-09 |
| 209086 | SAMD9L | 856.166656 | -1.5374651 | 0.26728491 | -5.7521584 | 8.81E-09 |

| | | | | | | |
|---|---|---|---|---|---|---|
| 227696 | PHYHD1 | 51.3299688 | -2.9244491 | 0.50849133 | -5.751227 | 8.86E-09 |
| 414872 | ZYG11B | 2180.59823 | 0.59001779 | 0.1026198 | 5.74955103 | 8.95E-09 |
| 68192 | LEPROTL1 | 1703.87299 | -0.5578016 | 0.09704056 | -5.7481281 | 9.02E-09 |
| 12982 | CSF2RA | 39.6716677 | -6.7456244 | 1.17371821 | -5.7472265 | 9.07E-09 |
| 83797 | SMARCD1 | 1287.55836 | 0.39839288 | 0.06936771 | 5.74320349 | 9.29E-09 |
| 74166 | TMEM38A | 246.676457 | -0.934587 | 0.16272997 | -5.7431765 | 9.29E-09 |
| 236539 | PHGDH | 1872.08755 | -0.3202408 | 0.05576842 | -5.7423316 | 9.34E-09 |
| 19697 | RELA | 2807.29711 | -0.4782056 | 0.08327949 | -5.7421778 | 9.35E-09 |
| 78830 | SLC25A12 | 1885.50301 | 0.22448786 | 0.03910025 | 5.74134017 | 9.39E-09 |
| 17762 | MAPT | 32.7853273 | -4.652493 | 0.81060265 | -5.7395482 | 9.49E-09 |
| 72056 | 1810055G02 | 1362.81773 | 0.65609657 | 0.11432674 | 5.73878467 | 9.54E-09 |
| 330171 | KCTD10 | 7115.06794 | 0.59830312 | 0.10428978 | 5.73692971 | 9.64E-09 |
| 56041 | USO1 | 4307.20421 | 0.32018084 | 0.05581327 | 5.73664339 | 9.66E-09 |
| 64050 | YEATS4 | 1126.70576 | 0.45276101 | 0.07892681 | 5.73646645 | 9.67E-09 |
| 338523 | KDM7A | 559.812104 | 0.80137016 | 0.13971423 | 5.73578059 | 9.71E-09 |
| 69046 | ISCA1 | 322.624669 | 0.90293608 | 0.15744767 | 5.73483294 | 9.76E-09 |
| 26377 | DAPP1 | 304.530627 | -0.7660979 | 0.13358754 | -5.7348004 | 9.76E-09 |
| 74493 | TNKS2 | 6610.04943 | -0.3626386 | 0.06325015 | -5.7334026 | 9.84E-09 |
| 78408 | FAM131A | 21.627272 | -4.3580764 | 0.76014369 | -5.7332271 | 9.85E-09 |
| 20358 | SEMA6A | 60.4419562 | -4.4755144 | 0.78069224 | -5.7327512 | 9.88E-09 |
| 74777 | SEPN1 | 2303.59926 | 0.42157687 | 0.07360602 | 5.72747792 | 1.02E-08 |
| 73293 | CCDC103 | 27.9460935 | -2.037643 | 0.35579379 | -5.7270337 | 1.02E-08 |
| 328190 | A530046M15 | 17.3827031 | -2.489606 | 0.4348318 | -5.7254461 | 1.03E-08 |
| 382406 | POC1B | 618.317142 | 0.42964977 | 0.07504368 | 5.72532892 | 1.03E-08 |
| 232236 | CCDC174 | 407.924574 | 0.67661589 | 0.11819377 | 5.72463239 | 1.04E-08 |
| 12912 | CREB1 | 1817.47626 | 0.39592549 | 0.06916515 | 5.72434905 | 1.04E-08 |
| 19361 | RAD51 | 1604.62475 | -0.3676244 | 0.06422738 | -5.7237951 | 1.04E-08 |
| 230828 | IL22RA1 | 10.6103111 | -6.832785 | 1.19401705 | -5.7225188 | 1.05E-08 |
| 667742 | PIEZO2 | 82.6917931 | -7.373486 | 1.28905746 | -5.72006 | 1.06E-08 |
| 54524 | SYT6 | 22.8804226 | -7.9475103 | 1.38992875 | -5.7179264 | 1.08E-08 |
| 338535 | E030013I19R | 14.5427219 | -7.2895706 | 1.27585588 | -5.713475 | 1.11E-08 |
| 50794 | KLF13 | 3777.49056 | -0.4831779 | 0.08457067 | -5.7133033 | 1.11E-08 |
| 171580 | MICAL1 | 53.7453553 | 4.40664621 | 0.77144462 | 5.71220031 | 1.12E-08 |
| 270076 | GCDH | 377.194226 | -0.9243156 | 0.16182162 | -5.7119415 | 1.12E-08 |
| 19126 | PROM1 | 231.454557 | -10.562355 | 1.84932405 | -5.711468 | 1.12E-08 |
| 67760 | SLC38A2 | 13462.2868 | 0.52120811 | 0.09128947 | 5.70939989 | 1.13E-08 |
| 50767 | PNPLA6 | 1088.522 | -0.716881 | 0.12561323 | -5.70705 | 1.15E-08 |
| 72194 | FBXL20 | 721.270258 | -1.3490386 | 0.23638907 | -5.7068571 | 1.15E-08 |
| 12857 | Cox4I1 | 5881.93383 | -0.5339742 | 0.09358531 | -5.7057477 | 1.16E-08 |
| 72946 | LRRC47 | 2558.05257 | 0.60453153 | 0.10597183 | 5.70464368 | 1.17E-08 |
| 75828 | HORMAD2 | 18.5983845 | -4.1237737 | 0.72291546 | -5.704365 | 1.17E-08 |
| 12728 | CLCN5 | 1566.28466 | -1.1867361 | 0.2080527 | -5.704017 | 1.17E-08 |
| 76130 | LAS1L | 1891.20265 | -0.5922114 | 0.10385438 | -5.7023244 | 1.18E-08 |
| 52504 | CENPO | 463.025012 | 0.78183661 | 0.13713338 | 5.70128609 | 1.19E-08 |
| 20528 | SLC2A4 | 15.0794893 | -5.2863878 | 0.92749005 | -5.6996707 | 1.20E-08 |

| | | | | | | |
|---|---|---|---|---|---|---|
| 63958 | UBE4B | 4169.19117 | 0.68958271 | 0.12101716 | 5.69822257 | 1.21E-08 |
| 21420 | TFAP2C | 80.2320421 | -6.4729234 | 1.13678908 | -5.6940408 | 1.24E-08 |
| 14248 | FLII | 4793.49621 | -0.3753731 | 0.06593205 | -5.6933319 | 1.25E-08 |
| 211612 | PTCHD1 | 134.291865 | 2.04841736 | 0.35991233 | 5.69143426 | 1.26E-08 |
| 19746 | rhd | 29.4750852 | 4.58000047 | 0.80484909 | 5.69050836 | 1.27E-08 |
| 23962 | Oasl2 | 165.833967 | -2.8930025 | 0.50858358 | -5.6883521 | 1.28E-08 |
| 106861 | ABHD3 | 52.2943752 | -2.1784553 | 0.38306037 | -5.6869763 | 1.29E-08 |
| 15478 | HS3ST3A1 | 20.1314289 | -7.7634848 | 1.36609898 | -5.6829592 | 1.32E-08 |
| 55950 | bri3 | 1271.16233 | -0.6756797 | 0.11890992 | -5.6822821 | 1.33E-08 |
| 20524 | SLC25A17 | 1365.84259 | -0.4344487 | 0.07649376 | -5.6795319 | 1.35E-08 |
| 67088 | CAND2 | 767.251862 | 1.65333333 | 0.29111007 | 5.67940969 | 1.35E-08 |
| 12173 | BNC1 | 877.276896 | 2.39290783 | 0.42144965 | 5.67780236 | 1.36E-08 |
| 78610 | UVRAG | 1884.81801 | 0.58992663 | 0.10390761 | 5.67741518 | 1.37E-08 |
| 268451 | RAB11FIP4 | 188.357528 | -3.834312 | 0.67547039 | -5.6765064 | 1.37E-08 |
| 67861 | AKR1B10 | 179.799926 | -0.685579 | 0.12078391 | -5.676079 | 1.38E-08 |
| 16600 | KLF4 | 3150.74476 | 1.93756318 | 0.34138364 | 5.6756181 | 1.38E-08 |
| 52668 | IFI27 | 975.737921 | -1.3012877 | 0.22931216 | -5.6747435 | 1.39E-08 |
| 18720 | PIP5K1A | 2768.22181 | -0.7516828 | 0.13246779 | -5.6744568 | 1.39E-08 |
| 107526 | GIMAP4 | 9.58158183 | -6.6950472 | 1.17991592 | -5.6741731 | 1.39E-08 |
| 503610 | ZDHHC18 | 2010.77226 | 0.89636855 | 0.15799913 | 5.67324978 | 1.40E-08 |
| 70420 | ARPIN | 1151.87797 | 0.53436561 | 0.09420564 | 5.67233114 | 1.41E-08 |
| 14465 | GATA6 | 249.941948 | -2.541464 | 0.44823677 | -5.6699142 | 1.43E-08 |
| 21340 | TAF1B | 452.084106 | 0.50317501 | 0.0887552 | 5.6692453 | 1.43E-08 |
| 12545 | CDC7 | 806.3629 | 0.55837101 | 0.09852007 | 5.66758669 | 1.45E-08 |
| 232816 | Zfp628 | 780.955884 | 0.78089205 | 0.13780084 | 5.66681619 | 1.45E-08 |
| 64656 | MRPS23 | 1604.77246 | 0.56327372 | 0.0994061 | 5.66639001 | 1.46E-08 |
| 103199 | FIG4 | 1088.75407 | -0.3280433 | 0.05791865 | -5.663863 | 1.48E-08 |
| 216829 | Mmgt2 | 561.795147 | -0.7789506 | 0.13755791 | -5.6627103 | 1.49E-08 |
| 244418 | D8Ertd82e | 634.540256 | -1.1088582 | 0.1958848 | -5.6607669 | 1.51E-08 |
| 67048 | VMA21 | 1022.24905 | -0.5037292 | 0.08905696 | -5.6562586 | 1.55E-08 |
| 15944 | Irgm1 | 952.714497 | -1.1522156 | 0.20371366 | -5.6560546 | 1.55E-08 |
| 217333 | TRIM47 | 2837.39115 | 1.88011802 | 0.33258077 | 5.65311706 | 1.58E-08 |
| 214240 | DISP2 | 96.7044455 | -6.7636408 | 1.19651085 | -5.6528036 | 1.58E-08 |
| 217830 | 9030617O03 | 605.691009 | -1.3346206 | 0.23611343 | -5.6524552 | 1.58E-08 |
| 317717 | SEC22A | 423.044795 | -0.6923826 | 0.12254193 | -5.6501691 | 1.60E-08 |
| 104457 | 0610010K14I | 520.834829 | 0.54425491 | 0.09633089 | 5.649848 | 1.61E-08 |
| 50996 | PDCD7 | 414.450334 | 0.59566242 | 0.10543666 | 5.6494809 | 1.61E-08 |
| 12549 | ARHGAP31 | 4260.16064 | -0.6294522 | 0.11143054 | -5.6488304 | 1.62E-08 |
| 66223 | MRPL35 | 1056.41585 | 0.57588035 | 0.10198263 | 5.64684753 | 1.63E-08 |
| 70231 | GORASP2 | 4856.79356 | 0.26798036 | 0.04746572 | 5.64576646 | 1.64E-08 |
| 110253 | TRIOBP | 1938.67828 | -0.5458394 | 0.09669904 | -5.6447242 | 1.65E-08 |
| 19300 | ABCD4 | 1423.31823 | 1.44067349 | 0.25531008 | 5.64283827 | 1.67E-08 |
| 83703 | DBR1 | 975.314149 | 0.43930372 | 0.07790093 | 5.63926119 | 1.71E-08 |
| 78286 | NAV2 | 4073.5301 | 0.7682141 | 0.13623757 | 5.63878317 | 1.71E-08 |
| 68644 | ABHD14A | 220.735162 | 0.98249795 | 0.17427714 | 5.63756054 | 1.72E-08 |

| | | | | | | |
|---|---|---|---|---|---|---|
| 66131 | TIPIN | 1392.33802 | 0.58216912 | 0.10327297 | 5.63718774 | 1.73E-08 |
| 217692 | SIPA1L1 | 7561.82982 | 0.50558732 | 0.08968839 | 5.6371546 | 1.73E-08 |
| 209294 | Csta1 | 8.13502011 | -6.4537012 | 1.14499516 | -5.636444 | 1.74E-08 |
| 67885 | 1500011K16I | 203.421232 | -0.7448221 | 0.13219055 | -5.6344583 | 1.76E-08 |
| 69993 | CHN2 | 178.163492 | -10.907217 | 1.93646429 | -5.6325424 | 1.78E-08 |
| 11435 | CHRNA1 | 14.473337 | 4.50625275 | 0.80029563 | 5.63073514 | 1.79E-08 |
| 68861 | 1190002N15 | 724.209801 | 1.06604895 | 0.18953275 | 5.62461618 | 1.86E-08 |
| 269424 | JADE1 | 2158.184 | -0.5701252 | 0.10136616 | -5.6244137 | 1.86E-08 |
| 14784 | GRB2 | 1997.61613 | -0.2470728 | 0.04393392 | -5.6237368 | 1.87E-08 |
| 225339 | AMMECR1L | 2380.65155 | -0.3586248 | 0.06378486 | -5.6224122 | 1.88E-08 |
| 27364 | SRR | 186.482671 | -1.3083108 | 0.23285543 | -5.6185541 | 1.93E-08 |
| 67249 | TBC1D19 | 632.610841 | -0.8821246 | 0.15701734 | -5.6180071 | 1.93E-08 |
| 21752 | TERT | 106.733827 | -1.942847 | 0.34594715 | -5.6160226 | 1.95E-08 |
| 70579 | ZC3H11A | 3889.97639 | -0.2790937 | 0.04970936 | -5.6145107 | 1.97E-08 |
| 12418 | CBX4 | 869.474078 | -0.3583624 | 0.06382804 | -5.6144987 | 1.97E-08 |
| 73122 | TGFBRAP1 | 1633.79525 | -0.3773579 | 0.06721202 | -5.6144407 | 1.97E-08 |
| 233987 | Zfp958 | 473.561161 | -0.7642473 | 0.13613088 | -5.6140631 | 1.98E-08 |
| 239405 | RSPO2 | 33.8277747 | 6.52180804 | 1.1617599 | 5.6137314 | 1.98E-08 |
| 14829 | GRPR | 25.2242527 | -5.5976767 | 0.99723449 | -5.6132001 | 1.99E-08 |
| 66454 | NMNAT1 | 136.510055 | 0.86093753 | 0.15339817 | 5.61243658 | 1.99E-08 |
| 381280 | HJURP | 569.278873 | -0.8874395 | 0.15819935 | -5.6096276 | 2.03E-08 |
| 72313 | FRYL | 1708.40735 | -0.8454781 | 0.15076828 | -5.6077984 | 2.05E-08 |
| 18606 | ENPP2 | 457.843311 | 2.63244975 | 0.46976509 | 5.60375774 | 2.10E-08 |
| 12750 | CLK4 | 819.088723 | -0.6294703 | 0.11234302 | -5.6031104 | 2.11E-08 |
| 216134 | PDXK | 1938.30086 | 1.26013896 | 0.22501311 | 5.6002913 | 2.14E-08 |
| 17540 | MRVI1 | 13.5907117 | -3.470101 | 0.61963665 | -5.6002191 | 2.14E-08 |
| 624582 | Morf4l1-ps1 | 121.202566 | 0.91695029 | 0.16374181 | 5.59997649 | 2.14E-08 |
| 321003 | XPNPEP3 | 512.927758 | -0.5462529 | 0.09755 | -5.5997221 | 2.15E-08 |
| 71206 | KATNAL2 | 63.1061788 | -1.4755492 | 0.2636009 | -5.5976639 | 2.17E-08 |
| 103694 | TMED4 | 1411.22774 | -0.5990384 | 0.10703038 | -5.5969009 | 2.18E-08 |
| 319719 | SIMC1 | 834.769944 | 0.45748758 | 0.08176757 | 5.59497579 | 2.21E-08 |
| 73047 | CAMK2N2 | 173.883348 | 1.08754443 | 0.19448658 | 5.59187397 | 2.25E-08 |
| 19116 | PRLR | 392.093287 | -11.323256 | 2.02569395 | -5.5898158 | 2.27E-08 |
| 435626 | RUFY4 | 40.1532258 | -8.7582458 | 1.56725386 | -5.5882751 | 2.29E-08 |
| 14232 | FKBP8 | 3836.51848 | -0.3009015 | 0.05385133 | -5.5876326 | 2.30E-08 |
| 67665 | DCTN4 | 3317.96857 | -0.4103376 | 0.07347284 | -5.5848875 | 2.34E-08 |
| 93673 | Nat8f2 | 8.53866601 | -6.5291642 | 1.16911051 | -5.584728 | 2.34E-08 |
| 71276 | CCDC57 | 147.199486 | -0.9096951 | 0.16299175 | -5.5812341 | 2.39E-08 |
| 53601 | PCDH12 | 8.17606851 | -6.4635895 | 1.15824589 | -5.5804985 | 2.40E-08 |
| 13595 | EBP | 1340.93089 | -0.7098366 | 0.12724834 | -5.5783562 | 2.43E-08 |
| 80913 | PUM2 | 9741.41269 | 0.28405845 | 0.05092281 | 5.5782168 | 2.43E-08 |
| 71093 | ATOH8 | 701.470534 | 2.42157958 | 0.43411583 | 5.57818763 | 2.43E-08 |
| 106564 | PPCS | 242.615299 | 0.74055217 | 0.13276452 | 5.57793743 | 2.43E-08 |
| 74019 | TRAF3IP1 | 614.141772 | -0.5742235 | 0.10296809 | -5.5767128 | 2.45E-08 |
| 218460 | WDR41 | 874.099392 | 0.5046097 | 0.09049162 | 5.57631416 | 2.46E-08 |

| | | | | | | |
|---|---|---|---|---|---|---|
| 54199 | CCRL2 | 22.2680073 | -7.174983 | 1.28711114 | -5.5744859 | 2.48E-08 |
| 231532 | arhgap24 | 309.174275 | 2.78175764 | 0.49905792 | 5.57401765 | 2.49E-08 |
| 235184 | MSANTD2 | 328.945826 | 0.59754197 | 0.10724876 | 5.57155112 | 2.52E-08 |
| 56809 | GMEB1 | 927.892565 | 0.68472978 | 0.12294016 | 5.56961839 | 2.55E-08 |
| 14809 | GRIK5 | 86.4247532 | -3.6700062 | 0.65902179 | -5.5688693 | 2.56E-08 |
| 12321 | CALU | 20214.4969 | 0.29083172 | 0.05222589 | 5.56872732 | 2.57E-08 |
| 20535 | SLC4A2 | 4017.00628 | -0.3618924 | 0.06499133 | -5.5683176 | 2.57E-08 |
| 70551 | TMTC4 | 387.830212 | -0.7425246 | 0.13334882 | -5.5682876 | 2.57E-08 |
| 28126 | NOP16 | 1131.09191 | 0.85373334 | 0.15333577 | 5.56773782 | 2.58E-08 |
| 83435 | PLEKHA3 | 1637.5935 | 0.68527061 | 0.12308885 | 5.56728437 | 2.59E-08 |
| 436188 | Gm5751 | 7.88950376 | 6.45711212 | 1.16037578 | 5.56467331 | 2.63E-08 |
| 232670 | TSPAN33 | 21.1125073 | -3.3356266 | 0.59954145 | -5.5636297 | 2.64E-08 |
| 68792 | SRPX2 | 211.210065 | 4.92267934 | 0.88542224 | 5.55969695 | 2.70E-08 |
| 432995 | SMIM22 | 22.0807457 | -7.8935207 | 1.41979053 | -5.5596375 | 2.70E-08 |
| 14489 | MTPN | 12871.7879 | -0.3743445 | 0.06735638 | -5.5576694 | 2.73E-08 |
| 234365 | YJEFN3 | 33.248234 | -2.1628562 | 0.38921048 | -5.5570349 | 2.74E-08 |
| 319582 | 6430573F11I | 9.77776105 | -6.7204193 | 1.20951413 | -5.5562966 | 2.76E-08 |
| 66890 | LMAN2 | 5142.17677 | 0.33177091 | 0.05971191 | 5.55619312 | 2.76E-08 |
| 114643 | Oas1c | 54.6928975 | -2.2115303 | 0.39809089 | -5.5553402 | 2.77E-08 |
| 70047 | TRNT1 | 2165.33096 | 0.94894199 | 0.17089459 | 5.55279119 | 2.81E-08 |
| 68310 | ZMYM1 | 740.308434 | 0.92488931 | 0.16660876 | 5.55126475 | 2.84E-08 |
| 237847 | RTN4RL1 | 45.182252 | 4.9149398 | 0.88538479 | 5.55119068 | 2.84E-08 |
| 11769 | AP1S1 | 3704.20517 | -0.4920868 | 0.08864695 | -5.5510858 | 2.84E-08 |
| 14381 | G6pdx | 4478.36063 | -0.1979989 | 0.03569881 | -5.5463732 | 2.92E-08 |
| 19344 | RAB5B | 823.641148 | -0.4158696 | 0.07498759 | -5.5458456 | 2.93E-08 |
| 242521 | KLHL9 | 3351.75448 | 0.69753709 | 0.12579724 | 5.54493162 | 2.94E-08 |
| 29809 | RABGAP1L | 749.139665 | -0.7791401 | 0.14051946 | -5.5447132 | 2.94E-08 |
| 17463 | PSMD7 | 4963.79475 | 0.47570491 | 0.08579517 | 5.54465835 | 2.95E-08 |
| 100042480 | NHSL2 | 31.3882179 | 3.36223406 | 0.60690058 | 5.54000796 | 3.02E-08 |
| 64655 | MRPS22 | 673.260356 | 0.38627417 | 0.06972707 | 5.5398024 | 3.03E-08 |
| 67804 | SNX2 | 2246.83046 | -0.469219 | 0.08470023 | -5.5397615 | 3.03E-08 |
| 19249 | PTPN13 | 1494.16185 | -1.1784338 | 0.21286648 | -5.5360234 | 3.09E-08 |
| 75292 | PRKD3 | 2651.4577 | 0.48198157 | 0.08707556 | 5.53521091 | 3.11E-08 |
| 78893 | CNOT10 | 1483.24884 | 0.46881838 | 0.08470082 | 5.53499248 | 3.11E-08 |
| 59036 | DACT1 | 56.9408589 | 3.89632004 | 0.70397797 | 5.5347187 | 3.12E-08 |
| 338370 | NALCN | 67.6903355 | -9.5105751 | 1.71907849 | -5.5323682 | 3.16E-08 |
| 231803 | MEPCE | 1827.42625 | 0.32288601 | 0.05837414 | 5.53131886 | 3.18E-08 |
| 230596 | PRPF38A | 909.79512 | 0.78272002 | 0.14153684 | 5.53015028 | 3.20E-08 |
| 83669 | wdr6 | 3905.82032 | 0.38043128 | 0.06880069 | 5.52946895 | 3.21E-08 |
| 207728 | PDE2A | 140.164338 | -1.5546026 | 0.28119396 | -5.5285774 | 3.23E-08 |
| 211135 | D130040H23 | 7.59102559 | -6.3562567 | 1.14985206 | -5.5278909 | 3.24E-08 |
| 217588 | MBIP | 380.196803 | 0.54437907 | 0.09849401 | 5.52702689 | 3.26E-08 |
| 21345 | TAGLN | 82.4859639 | 3.99052624 | 0.7220774 | 5.52645218 | 3.27E-08 |
| 66049 | ROGDI | 251.026638 | -0.814474 | 0.14737981 | -5.5263605 | 3.27E-08 |
| 654432 | Gm7334 | 335.101686 | 1.62380873 | 0.29395993 | 5.52391178 | 3.32E-08 |

| | | | | | | |
|---|---|---|---|---|---|---|
| 15979 | IFNGR1 | 1154.90814 | -0.9153879 | 0.16576509 | -5.5221994 | 3.35E-08 |
| 209224 | ENOX2 | 201.939596 | -0.6814086 | 0.12340862 | -5.5215643 | 3.36E-08 |
| 15166 | HCN2 | 102.222476 | -2.3365015 | 0.42337556 | -5.5187444 | 3.41E-08 |
| 16504 | KCNC3 | 14.1341065 | -6.5127053 | 1.18014125 | -5.5185812 | 3.42E-08 |
| 237868 | SARM1 | 22.0183409 | -3.0312129 | 0.54930471 | -5.5182723 | 3.42E-08 |
| 27373 | csnk1e | 1516.11639 | -0.5867764 | 0.10636525 | -5.5166175 | 3.46E-08 |
| 13385 | dlg4 | 405.245508 | 2.8260162 | 0.51247618 | 5.51443426 | 3.50E-08 |
| 17005 | LTK | 41.8676509 | -8.0927556 | 1.46801336 | -5.5127261 | 3.53E-08 |
| 268469 | Zfp652 | 545.801703 | -0.4840301 | 0.08781939 | -5.5116542 | 3.55E-08 |
| 225913 | TKFC | 549.628927 | -0.6039654 | 0.10958211 | -5.5115334 | 3.56E-08 |
| 102657 | CD276 | 1448.02437 | -0.3363287 | 0.06102461 | -5.5113615 | 3.56E-08 |
| 67619 | NOB1 | 1814.59371 | 0.67737709 | 0.12290632 | 5.51132831 | 3.56E-08 |
| 71336 | RBKS | 31.2096308 | -3.2846225 | 0.59602061 | -5.510921 | 3.57E-08 |
| 70456 | MPC2 | 655.047231 | -0.5760591 | 0.1045441 | -5.5102019 | 3.58E-08 |
| 66752 | 4933404O12 | 679.483012 | 1.04018021 | 0.18878077 | 5.50999037 | 3.59E-08 |
| 56374 | TMEM59 | 3791.66096 | 0.5026578 | 0.09124795 | 5.50870223 | 3.61E-08 |
| 11855 | ARHGAP5 | 3880.77116 | 0.46078724 | 0.0836666 | 5.50742192 | 3.64E-08 |
| 76793 | SNIP1 | 724.831621 | 0.72269223 | 0.13122646 | 5.50721427 | 3.65E-08 |
| 279561 | WNK3 | 37.6255805 | -1.8473617 | 0.33545067 | -5.5071038 | 3.65E-08 |
| 54208 | ARL6IP1 | 5027.07227 | -1.0624521 | 0.19294326 | -5.5065523 | 3.66E-08 |
| 66416 | NDUFA7 | 867.409719 | -0.6489317 | 0.11785472 | -5.5062 | 3.67E-08 |
| 66214 | RGCC | 14.6943172 | 3.51486647 | 0.63836241 | 5.50606744 | 3.67E-08 |
| 235682 | Zfp445 | 2610.7651 | 0.37538586 | 0.06817972 | 5.50582867 | 3.67E-08 |
| 20708 | Serpinb6b | 48.2548068 | -3.7731166 | 0.68539443 | -5.5050296 | 3.69E-08 |
| 93695 | GPNMB | 7974.91158 | 2.38860957 | 0.43410926 | 5.50232342 | 3.75E-08 |
| 74176 | TGM5 | 12.0544811 | -6.2816568 | 1.14163731 | -5.5023226 | 3.75E-08 |
| 68323 | NUDT22 | 241.162826 | -0.7092362 | 0.12903873 | -5.4963049 | 3.88E-08 |
| 13429 | DNM1 | 2153.93509 | 4.23819716 | 0.77114536 | 5.49597702 | 3.89E-08 |
| 216820 | DHRS7B | 685.975942 | -0.5808832 | 0.10570242 | -5.4954581 | 3.90E-08 |
| 11603 | AGRN | 5684.92331 | -1.8946094 | 0.34480435 | -5.4947377 | 3.91E-08 |
| 240725 | SULF1 | 156.025258 | -10.715591 | 1.95019225 | -5.494633 | 3.92E-08 |
| 19056 | PPP3CB | 2902.17392 | 0.339329 | 0.06176882 | 5.49353258 | 3.94E-08 |
| 668894 | Gm14025 | 7.56188484 | -6.3516065 | 1.15670977 | -5.4910978 | 3.99E-08 |
| 16500 | KCNB1 | 21.1802534 | -5.4038487 | 0.98444146 | -5.4892535 | 4.04E-08 |
| 16832 | LDHB | 227.478477 | -2.5632216 | 0.46705391 | -5.4880638 | 4.06E-08 |
| 212706 | N4BP3 | 329.383716 | -1.3886519 | 0.25305096 | -5.4876372 | 4.07E-08 |
| 69934 | TRMT10B | 142.673014 | -1.0764403 | 0.19624238 | -5.4852592 | 4.13E-08 |
| 72709 | C1QTNF6 | 465.279405 | 4.36383224 | 0.79557032 | 5.48516218 | 4.13E-08 |
| 18218 | DUSP8 | 508.282429 | -1.121074 | 0.20440295 | -5.4846273 | 4.14E-08 |
| 66361 | ZFAND1 | 65.981174 | -1.0908489 | 0.19891734 | -5.4839307 | 4.16E-08 |
| 56070 | TCERG1 | 4275.55869 | 0.31194424 | 0.05688684 | 5.4835926 | 4.17E-08 |
| 66146 | TMEM57 | 1582.33626 | 0.58616952 | 0.10698086 | 5.47920001 | 4.27E-08 |
| 12499 | ENTPD5 | 997.846174 | 0.58402547 | 0.10662075 | 5.47759649 | 4.31E-08 |
| 67072 | CDC37L1 | 1179.25099 | -0.4123526 | 0.0752816 | -5.4774679 | 4.31E-08 |
| 12841 | COL9A3 | 46.9765912 | -1.8474207 | 0.3372773 | -5.4774535 | 4.31E-08 |

| | | | | | | |
|---|---|---|---|---|---|---|
| 218630 | CCNO | 22.9620963 | 3.88625557 | 0.70980305 | 5.47511816 | 4.37E-08 |
| 11842 | ARF3 | 7197.27616 | -0.5596071 | 0.10227043 | -5.4718362 | 4.45E-08 |
| 52028 | BBS1 | 154.650409 | -1.1453589 | 0.20933641 | -5.4713796 | 4.47E-08 |
| 18828 | PLSCR2 | 1816.27175 | -1.7376621 | 0.31775516 | -5.4685566 | 4.54E-08 |
| 216161 | SBNO2 | 2128.34745 | -0.5776021 | 0.10566128 | -5.466545 | 4.59E-08 |
| 72084 | PIGX | 435.955277 | -0.4349257 | 0.07956691 | -5.4661627 | 4.60E-08 |
| 56527 | MAST1 | 11.8801372 | -6.2657432 | 1.14632118 | -5.4659579 | 4.60E-08 |
| 73205 | 3110043O21 | 649.133062 | -1.3962172 | 0.2555844 | -5.4628421 | 4.69E-08 |
| 19217 | PTGER2 | 55.6130718 | 3.58103449 | 0.65553342 | 5.46277941 | 4.69E-08 |
| 229504 | ISG20L2 | 1994.6624 | 0.70644605 | 0.12936898 | 5.46070675 | 4.74E-08 |
| 13655 | EGR3 | 99.336702 | 3.6223708 | 0.66365616 | 5.45820412 | 4.81E-08 |
| 74158 | JOSD1 | 2192.13232 | 0.45717 | 0.08377326 | 5.45723041 | 4.84E-08 |
| 76156 | FAM131B | 236.424802 | 3.56617811 | 0.65370402 | 5.45534068 | 4.89E-08 |
| 74190 | EXOC3L4 | 169.130032 | 3.70232531 | 0.67884279 | 5.45387735 | 4.93E-08 |
| 20441 | ST3GAL3 | 305.412513 | 0.69793853 | 0.12797569 | 5.45368051 | 4.93E-08 |
| 71562 | AFMID | 192.631093 | -1.1238692 | 0.20611532 | -5.4526234 | 4.96E-08 |
| 225187 | ANKRD29 | 264.993676 | -1.2950269 | 0.23751916 | -5.4523048 | 4.97E-08 |
| 103140 | GstT3 | 199.411932 | -1.7138007 | 0.31436902 | -5.4515571 | 4.99E-08 |
| 66628 | THG1L | 422.050972 | 0.75916629 | 0.13935282 | 5.44779989 | 5.10E-08 |
| 18008 | NES | 4814.0584 | -1.8443449 | 0.33855685 | -5.4476667 | 5.10E-08 |
| 246198 | MLLT6 | 1883.0143 | 0.31603722 | 0.05805293 | 5.44394988 | 5.21E-08 |
| 102566 | ANO10 | 1366.85276 | -0.7203832 | 0.13235071 | -5.4429875 | 5.24E-08 |
| 330361 | GCFC2 | 474.130321 | 0.58641573 | 0.1077436 | 5.44269658 | 5.25E-08 |
| 100561 | SLC15A4 | 1097.60975 | -0.414899 | 0.07623281 | -5.4425259 | 5.25E-08 |
| 103677 | SMG6 | 1308.81802 | 0.54017907 | 0.09926781 | 5.44163393 | 5.28E-08 |
| 11568 | AEBP1 | 23428.4978 | 1.19297729 | 0.21923246 | 5.44160874 | 5.28E-08 |
| 70617 | 5730508B09 | 107.609627 | -0.9346893 | 0.17192293 | -5.4366761 | 5.43E-08 |
| 22129 | TTC3 | 3445.38387 | 0.47726479 | 0.08779847 | 5.43591217 | 5.45E-08 |
| 17420 | MNAT1 | 983.178576 | 0.29744899 | 0.05472133 | 5.43570479 | 5.46E-08 |
| 240880 | SCYL3 | 935.06156 | -0.3148505 | 0.05792522 | -5.4354641 | 5.47E-08 |
| 27224 | TCEB3 | 2776.92293 | 0.63372009 | 0.11659717 | 5.43512403 | 5.48E-08 |
| 634731 | SUSD1 | 61.8241419 | -4.7045492 | 0.86565644 | -5.4346609 | 5.49E-08 |
| 76422 | Mroh3 | 8.02957948 | -6.4346691 | 1.18407917 | -5.4343233 | 5.50E-08 |
| 20665 | SOX10 | 206.648704 | -11.121021 | 2.04661412 | -5.433863 | 5.51E-08 |
| 386655 | EID2 | 267.431951 | 1.62032086 | 0.29819027 | 5.43384885 | 5.52E-08 |
| 68487 | TMEM140 | 135.71451 | -2.5188945 | 0.46356817 | -5.4337089 | 5.52E-08 |
| 226551 | SUCO | 2332.0022 | -0.5766469 | 0.10612869 | -5.4334682 | 5.53E-08 |
| 238252 | GPR135 | 43.1627648 | 1.31899222 | 0.24281524 | 5.43208161 | 5.57E-08 |
| 83436 | PLEKHA2 | 2118.03613 | -1.3505001 | 0.24867272 | -5.4308334 | 5.61E-08 |
| 12406 | SERPINH1 | 15007.9861 | 1.09985938 | 0.20277347 | 5.42407928 | 5.83E-08 |
| 245841 | POLR2H | 497.720811 | 0.5074186 | 0.09358769 | 5.42185221 | 5.90E-08 |
| 67561 | WDR48 | 1758.10763 | 0.38791113 | 0.07154948 | 5.42157895 | 5.91E-08 |
| 16497 | KCNAB1 | 60.5269235 | -3.4730257 | 0.64059747 | -5.4215415 | 5.91E-08 |
| 228357 | LRP4 | 149.091231 | 1.55246499 | 0.28635769 | 5.42141886 | 5.91E-08 |
| 102448 | XYLB | 389.635015 | 0.88656485 | 0.16359676 | 5.4192079 | 5.99E-08 |

| | | | | | | |
|---|---|---|---|---|---|---|
| 75612 | GNS | 9646.43756 | -1.1159962 | 0.20597936 | -5.4180002 | 6.03E-08 |
| 68151 | WLS | 7133.3636 | -0.4039096 | 0.07458385 | -5.4155098 | 6.11E-08 |
| 68770 | PHTF2 | 647.646423 | 0.45739981 | 0.08446436 | 5.41529924 | 6.12E-08 |
| 381101 | DNPH1 | 577.842625 | 0.52268131 | 0.0965362 | 5.41435577 | 6.15E-08 |
| 17237 | MGRN1 | 2196.02415 | -0.5301682 | 0.09797439 | -5.4112935 | 6.26E-08 |
| 12830 | COL4A5 | 2210.69119 | -0.7068517 | 0.13064293 | -5.4105621 | 6.28E-08 |
| 68002 | SDHAF4 | 189.328749 | -0.7272639 | 0.13443615 | -5.4097344 | 6.31E-08 |
| 100503791 | Gm16201 | 18.4466206 | -3.9466052 | 0.72955491 | -5.4096069 | 6.32E-08 |
| 108897 | AIF1L | 2134.47483 | -4.1786342 | 0.77244815 | -5.4095983 | 6.32E-08 |
| 320560 | DENND5B | 342.424561 | 1.45350297 | 0.26870749 | 5.40923871 | 6.33E-08 |
| 27045 | NIT1 | 414.742823 | -0.4883468 | 0.0902812 | -5.4091747 | 6.33E-08 |
| 216859 | ACAP1 | 15.4386618 | -5.3197618 | 0.98347027 | -5.4091739 | 6.33E-08 |
| 328968 | Carmn | 12.5218588 | 5.0438382 | 0.93294401 | 5.40636753 | 6.43E-08 |
| 330177 | TAOK3 | 747.742032 | -0.8141554 | 0.15061425 | -5.4055666 | 6.46E-08 |
| 15108 | HSD17B10 | 1019.60204 | -0.3556892 | 0.06580153 | -5.4054857 | 6.46E-08 |
| 15018 | H2-Q7 | 34.4027994 | -1.9397149 | 0.35885077 | -5.4053524 | 6.47E-08 |
| 69257 | ELF2 | 1907.05717 | 0.39970103 | 0.07396236 | 5.40411375 | 6.51E-08 |
| 103468 | NUP107 | 3047.97984 | 0.38451855 | 0.0711705 | 5.40277962 | 6.56E-08 |
| 73327 | PRADC1 | 306.743689 | 0.8152882 | 0.15091767 | 5.40220491 | 6.58E-08 |
| 94186 | STRN3 | 4685.24471 | 0.54642018 | 0.10116842 | 5.40109425 | 6.62E-08 |
| 20537 | SLC5A1 | 37.1294003 | -4.2209772 | 0.78157339 | -5.4006152 | 6.64E-08 |
| 15507 | HSPB1 | 71.7240415 | -3.9985018 | 0.74065517 | -5.3986011 | 6.72E-08 |
| 76877 | RAB36 | 41.5112812 | -1.5885943 | 0.29432544 | -5.3974072 | 6.76E-08 |
| 268396 | SH3PXD2B | 4887.54918 | 1.33037367 | 0.24650471 | 5.39695025 | 6.78E-08 |
| 74637 | shpk | 83.5522951 | -1.3030904 | 0.24145525 | -5.3968195 | 6.78E-08 |
| 231580 | GAK | 3836.11967 | -0.4178995 | 0.07744437 | -5.3961251 | 6.81E-08 |
| 207818 | SMAGP | 32.8713356 | -5.6690792 | 1.05066908 | -5.3956848 | 6.83E-08 |
| 68904 | ABHD13 | 1495.0226 | -0.3274676 | 0.06069504 | -5.3952943 | 6.84E-08 |
| 67091 | TRAPPC6A | 346.3611 | -1.1378804 | 0.2109595 | -5.3938332 | 6.90E-08 |
| 117600 | SRGAP1 | 562.261019 | -1.6534238 | 0.30655676 | -5.3935325 | 6.91E-08 |
| 22379 | FMNL3 | 3087.7439 | 1.47264256 | 0.27306858 | 5.39294041 | 6.93E-08 |
| 67775 | RTP4 | 151.887378 | -2.2192908 | 0.41154739 | -5.3925523 | 6.95E-08 |
| 56546 | sec1 | 30.8018034 | 3.00402105 | 0.55713473 | 5.39191128 | 6.97E-08 |
| 93694 | CLEC2D | 192.706845 | -2.3215468 | 0.43060397 | -5.3913734 | 6.99E-08 |
| 56440 | SNX1 | 1595.33901 | 0.27199014 | 0.05045367 | 5.39088949 | 7.01E-08 |
| 11994 | PCDH15 | 14.7910716 | -5.2332039 | 0.97150214 | -5.3867137 | 7.18E-08 |
| 18746 | PKM | 41131.8831 | 0.41746638 | 0.07755152 | 5.38308429 | 7.32E-08 |
| 226691 | Ifi207 | 155.360056 | -1.0090281 | 0.18751529 | -5.3810441 | 7.41E-08 |
| 22411 | WNT11 | 54.3847984 | -1.8369157 | 0.34136793 | -5.3810434 | 7.41E-08 |
| 11798 | XIAP | 2855.23831 | -0.3171298 | 0.05897859 | -5.3770321 | 7.57E-08 |
| 13841 | EPHA7 | 27.5302844 | -8.2118815 | 1.52759953 | -5.3756769 | 7.63E-08 |
| 68153 | GTF2E2 | 792.424786 | 0.70403008 | 0.1309724 | 5.37540788 | 7.64E-08 |
| 102294 | Cyp4v3 | 151.990501 | -1.4300937 | 0.26614983 | -5.3732655 | 7.73E-08 |
| 78892 | CRISPLD2 | 80.5278442 | -5.4273571 | 1.01010247 | -5.3730758 | 7.74E-08 |
| 330959 | SNAPC5 | 247.047296 | -0.6982431 | 0.12995511 | -5.372956 | 7.75E-08 |

| | | | | | | |
|---|---|---|---|---|---|---|
| 75452 | ASCC2 | 1616.72206 | -0.4581688 | 0.08529174 | -5.3717845 | 7.80E-08 |
| 231214 | CC2D2A | 603.562463 | -0.3780624 | 0.07040046 | -5.3701694 | 7.87E-08 |
| 21927 | TNFAIP1 | 4662.637 | -0.1941835 | 0.03617302 | -5.3681859 | 7.95E-08 |
| 100210 | GPN2 | 476.500955 | 1.021276 | 0.1903461 | 5.36536348 | 8.08E-08 |
| 11302 | AATK | 117.844102 | -5.2743191 | 0.98317704 | -5.3645669 | 8.11E-08 |
| 76261 | 0610040J01F | 47.7124097 | -9.0058779 | 1.67881908 | -5.3644124 | 8.12E-08 |
| 234730 | FUK | 477.088538 | -0.597093 | 0.11134992 | -5.3623118 | 8.22E-08 |
| 78245 | ACBD7 | 11.139026 | 4.4604259 | 0.83192017 | 5.3616033 | 8.25E-08 |
| 13371 | DIO2 | 675.609013 | 2.30184448 | 0.42933404 | 5.36143017 | 8.26E-08 |
| 66990 | TMEM134 | 1285.24275 | -0.4815543 | 0.089835 | -5.360431 | 8.30E-08 |
| 241624 | EXD1 | 54.0921757 | -1.6410164 | 0.30630597 | -5.3574417 | 8.44E-08 |
| 70266 | KYAT1 | 79.6021098 | -1.0144086 | 0.18934689 | -5.357408 | 8.44E-08 |
| 21887 | TLE3 | 1956.31109 | 0.58100491 | 0.10847483 | 5.35612648 | 8.50E-08 |
| 112406 | EGLN2 | 1802.45611 | 0.42874605 | 0.08006068 | 5.35526362 | 8.54E-08 |
| 17289 | MERTK | 61.8773176 | -3.0158495 | 0.5631817 | -5.3550203 | 8.55E-08 |
| 211535 | CCDC114 | 195.730064 | 4.07688751 | 0.761589 | 5.35313339 | 8.64E-08 |
| 19291 | PURB | 10126.2159 | 0.38461719 | 0.07186545 | 5.35190695 | 8.70E-08 |
| 170822 | USP33 | 2007.13845 | -0.2793726 | 0.05220703 | -5.3512454 | 8.74E-08 |
| 327812 | Gm15663 | 7.88991815 | -6.4095306 | 1.19780438 | -5.3510662 | 8.74E-08 |
| 72170 | CHCHD4 | 1001.84123 | 0.87429366 | 0.16342253 | 5.34989665 | 8.80E-08 |
| 58250 | CHST11 | 1679.16875 | -1.2516494 | 0.23398468 | -5.3492792 | 8.83E-08 |
| 56314 | Zfp113 | 298.489291 | -0.8030264 | 0.1501531 | -5.3480509 | 8.89E-08 |
| 228715 | Gm561 | 162.359601 | 0.63889479 | 0.11952118 | 5.34545228 | 9.02E-08 |
| 12175 | BNIP2 | 4566.51608 | 0.30274751 | 0.05665223 | 5.34396441 | 9.09E-08 |
| 212647 | ALDH4A1 | 229.383672 | -0.6840031 | 0.12800974 | -5.3433677 | 9.12E-08 |
| 100038712 | Gm10516 | 42.0150051 | -1.2723764 | 0.23817175 | -5.3422643 | 9.18E-08 |
| 17533 | MRC1 | 93.7287112 | 2.6427215 | 0.49476069 | 5.34141366 | 9.22E-08 |
| 69707 | IQCG | 15.8815246 | -2.5844616 | 0.48386807 | -5.3412527 | 9.23E-08 |
| 17252 | RDH11 | 1845.58483 | 0.59722863 | 0.11184261 | 5.33990238 | 9.30E-08 |
| 244867 | ARHGAP20 | 32.5305265 | 3.49163828 | 0.6538975 | 5.33973333 | 9.31E-08 |
| 432800 | Gm5454 | 241.295629 | 1.92948125 | 0.36148042 | 5.33771989 | 9.41E-08 |
| 57895 | CCDC126 | 332.513932 | -0.8191847 | 0.15348437 | -5.3372513 | 9.44E-08 |
| 12061 | BDKRB1 | 14.888009 | 3.36219428 | 0.62997031 | 5.33706782 | 9.45E-08 |
| 116891 | DERL2 | 1536.58696 | -0.3609848 | 0.0676377 | -5.3370358 | 9.45E-08 |
| 110695 | ALDH7A1 | 1452.17229 | -1.0947216 | 0.20519012 | -5.3351577 | 9.55E-08 |
| 100169 | PHACTR4 | 2293.56931 | 0.67727017 | 0.1269776 | 5.3337766 | 9.62E-08 |
| 72519 | TMEM55A | 2348.50136 | -1.2117612 | 0.2272022 | -5.3334045 | 9.64E-08 |
| 14628 | OSTM1 | 1926.44012 | -1.1311307 | 0.21212025 | -5.3324973 | 9.69E-08 |
| 192119 | DICER1 | 3783.02969 | 0.61077721 | 0.1145767 | 5.33072774 | 9.78E-08 |
| 74018 | ALS2 | 2136.7667 | -0.3252757 | 0.06103027 | -5.3297434 | 9.84E-08 |
| 73390 | Msl3l2 | 11.7422235 | -6.2522222 | 1.17315069 | -5.329428 | 9.85E-08 |
| 18391 | SIGMAR1 | 3633.67861 | 0.62081774 | 0.1165052 | 5.32867003 | 9.89E-08 |
| 328108 | FAM179B | 2932.72262 | 0.55929599 | 0.10497038 | 5.32813152 | 9.92E-08 |
| 19716 | BEX1 | 11.8014532 | -6.9879343 | 1.31165282 | -5.3275792 | 9.95E-08 |
| 14783 | GRB10 | 271.706818 | -10.793889 | 2.02699189 | -5.3250774 | 1.01E-07 |

| | | | | | | |
|---|---|---|---|---|---|---|
| 106064 | AW549877 | 2435.6056 | 0.65381133 | 0.12279653 | 5.3243471 | 1.01E-07 |
| 100616095 | SNHG18 | 311.329367 | 0.96770929 | 0.18175185 | 5.32434365 | 1.01E-07 |
| 16646 | KPNA1 | 2598.09906 | -0.2724263 | 0.05117075 | -5.3238677 | 1.02E-07 |
| 11815 | APOD | 24.3471178 | 6.66332993 | 1.25241932 | 5.32036661 | 1.04E-07 |
| 19366 | RAD54L | 686.191824 | 1.21877313 | 0.22908813 | 5.32010609 | 1.04E-07 |
| 100504399 | C130021I20R | 342.265679 | -7.9070426 | 1.48663233 | -5.3187614 | 1.04E-07 |
| 75329 | atf7ip2 | 17.9130792 | -5.5204913 | 1.03862952 | -5.3151689 | 1.07E-07 |
| 18124 | NR4A3 | 13.776218 | 3.0870883 | 0.58109652 | 5.31252241 | 1.08E-07 |
| 78625 | 1700061G19 | 9.66139044 | -5.9576767 | 1.12146654 | -5.3123981 | 1.08E-07 |
| 12487 | CD28 | 11.7296046 | -4.4757451 | 0.8428262 | -5.310401 | 1.09E-07 |
| 107701 | SF3B4 | 920.352886 | 0.42511473 | 0.08006316 | 5.30974209 | 1.10E-07 |
| 93702 | PCDHGB5 | 212.076866 | 4.58010325 | 0.86282662 | 5.30825444 | 1.11E-07 |
| 19341 | RAB4A | 395.868122 | -0.6802845 | 0.12818796 | -5.3069295 | 1.11E-07 |
| 270685 | MTHFD1L | 4275.34788 | 0.59424797 | 0.1119802 | 5.30672366 | 1.12E-07 |
| 18996 | POU4F1 | 11.5423884 | -3.6036736 | 0.67913103 | -5.3063009 | 1.12E-07 |
| 57913 | PIDD1 | 436.894913 | -0.5402941 | 0.10182613 | -5.3060458 | 1.12E-07 |
| 74737 | PCF11 | 2053.28338 | 0.36358798 | 0.0685272 | 5.30574705 | 1.12E-07 |
| 21379 | TBRG4 | 3001.82643 | 0.24987027 | 0.04709975 | 5.30512914 | 1.13E-07 |
| 383295 | YPEL5 | 1360.3722 | 0.63142265 | 0.1191315 | 5.30021592 | 1.16E-07 |
| 54196 | PABPN1 | 4280.53339 | 0.76999746 | 0.1452895 | 5.29974614 | 1.16E-07 |
| 56708 | CLCF1 | 1262.2146 | 0.71182605 | 0.13436577 | 5.29767385 | 1.17E-07 |
| 54405 | NDUFA1 | 639.218382 | -0.428012 | 0.0808272 | -5.2953954 | 1.19E-07 |
| 72128 | 2610008E11I | 331.072309 | -0.6126044 | 0.11569944 | -5.2947914 | 1.19E-07 |
| 69048 | SLC30A5 | 2266.09961 | 0.59401776 | 0.11224313 | 5.29224153 | 1.21E-07 |
| 213783 | PLEKHG1 | 30.7032878 | -4.0859206 | 0.77205884 | -5.29224 | 1.21E-07 |
| 224794 | ENPP4 | 622.61291 | -0.9958592 | 0.18824779 | -5.2901508 | 1.22E-07 |
| 26572 | COPS3 | 2175.43797 | -0.2947687 | 0.05572485 | -5.2897177 | 1.23E-07 |
| 100129 | GPR153 | 198.231518 | 2.11174808 | 0.39922673 | 5.28959587 | 1.23E-07 |
| 11933 | ATP1B3 | 4539.27614 | 0.5284895 | 0.09992977 | 5.28860941 | 1.23E-07 |
| 66355 | GMPR | 45.9511091 | 1.82044389 | 0.3444727 | 5.28472612 | 1.26E-07 |
| 232791 | CNOT3 | 1794.46294 | 0.83635526 | 0.15827305 | 5.28425563 | 1.26E-07 |
| 106840 | UNC119B | 1776.64281 | -0.6099601 | 0.11544484 | -5.2835629 | 1.27E-07 |
| 67647 | 4930523C07I | 597.553865 | -1.555867 | 0.29449391 | -5.2831891 | 1.27E-07 |
| 24082 | Map2k3os | 134.389861 | 1.05152932 | 0.19907843 | 5.28198531 | 1.28E-07 |
| 24116 | NELFA | 924.632025 | 0.39238492 | 0.07428875 | 5.28188893 | 1.28E-07 |
| 24066 | SPRY4 | 1169.71438 | 2.85713399 | 0.54096733 | 5.28152776 | 1.28E-07 |
| 668173 | PEX10 | 138.432299 | 0.96985851 | 0.18375499 | 5.2779981 | 1.31E-07 |
| 67163 | CCDC47 | 3894.75721 | -0.2834746 | 0.05371464 | -5.2774177 | 1.31E-07 |
| 59042 | COPE | 1772.38081 | -0.3461914 | 0.06560051 | -5.2772666 | 1.31E-07 |
| 74840 | MANF | 2717.58784 | 0.53263639 | 0.10095305 | 5.27608028 | 1.32E-07 |
| 329910 | ACOT11 | 30.0759462 | -5.890754 | 1.116699 | -5.2751493 | 1.33E-07 |
| 68554 | CEBPZOS | 215.954831 | -0.6297377 | 0.11959384 | -5.2656367 | 1.40E-07 |
| 54352 | IRX5 | 1119.25066 | -0.4303314 | 0.08173515 | -5.2649489 | 1.40E-07 |
| 69660 | TMBIM1 | 3322.91976 | -0.3732412 | 0.07089611 | -5.2646213 | 1.40E-07 |
| 78263 | 4632411P08I | 7.66120956 | -6.3636344 | 1.2088027 | -5.2644112 | 1.41E-07 |

| | | | | | | |
|---|---|---|---|---|---|---|
| 242509 | BNC2 | 53.9544341 | 4.77761844 | 0.90758719 | 5.26408756 | 1.41E-07 |
| 245638 | TBC1D8B | 1641.56536 | 0.60795986 | 0.1155241 | 5.26262347 | 1.42E-07 |
| 56222 | CITED4 | 54.4880659 | -7.2105949 | 1.37091183 | -5.2597072 | 1.44E-07 |
| 103466 | NT5DC3 | 2917.42448 | 0.94723014 | 0.18021556 | 5.25609527 | 1.47E-07 |
| 231946 | FAM221A | 20.4025095 | -2.4620603 | 0.46842299 | -5.256062 | 1.47E-07 |
| 237400 | MEX3D | 2377.95119 | 0.45954721 | 0.0874358 | 5.25582422 | 1.47E-07 |
| 73845 | ANKRD42 | 163.61623 | -1.056339 | 0.20102632 | -5.2547296 | 1.48E-07 |
| 66368 | RTCA | 1376.10032 | -0.3802435 | 0.0723713 | -5.2540654 | 1.49E-07 |
| 57444 | ISG20 | 393.728431 | -1.3751922 | 0.26193665 | -5.2500944 | 1.52E-07 |
| 12400 | CBFB | 4913.40905 | 0.39921121 | 0.0760466 | 5.24956007 | 1.52E-07 |
| 319518 | PDPR | 518.056468 | 0.63715615 | 0.12138129 | 5.24921204 | 1.53E-07 |
| 239250 | SLITRK6 | 6.51330005 | -6.1350819 | 1.16883604 | -5.2488815 | 1.53E-07 |
| 380993 | ZFAT | 327.89858 | 0.82045013 | 0.15631175 | 5.24880652 | 1.53E-07 |
| 27416 | ABCC5 | 1614.13271 | -0.8276194 | 0.15769505 | -5.2482273 | 1.54E-07 |
| 620883 | Gm12696 | 680.532611 | 0.46038377 | 0.087743 | 5.24695707 | 1.55E-07 |
| 66576 | Uqcrh | 1120.78552 | 0.74216359 | 0.14145883 | 5.24649898 | 1.55E-07 |
| 71130 | SH2D6 | 13.9065164 | 3.73253107 | 0.71147402 | 5.24619445 | 1.55E-07 |
| 67582 | SLC25A26 | 210.246506 | 0.5391216 | 0.10279908 | 5.24442066 | 1.57E-07 |
| 226182 | TAF5 | 358.054471 | 0.49739095 | 0.09484719 | 5.24412933 | 1.57E-07 |
| 233545 | EMSY | 1193.35843 | 0.7499559 | 0.14305653 | 5.24237456 | 1.59E-07 |
| 75747 | SESN3 | 388.975698 | -0.7723433 | 0.14735286 | -5.2414544 | 1.59E-07 |
| 17060 | BLNK | 107.477089 | -6.3885377 | 1.21887793 | -5.241327 | 1.59E-07 |
| 76432 | 2310001H17 | 15.68631 | -2.7625499 | 0.52715815 | -5.2404575 | 1.60E-07 |
| 234023 | ARGLU1 | 1731.89934 | 0.34630531 | 0.06610505 | 5.23871183 | 1.62E-07 |
| 67996 | SRSF6 | 8318.14397 | 0.31374511 | 0.05990377 | 5.23748481 | 1.63E-07 |
| 70834 | SPAG9 | 5954.95926 | -0.5036206 | 0.0961769 | -5.2363987 | 1.64E-07 |
| 13121 | CYP51 | 8037.10061 | 0.15716795 | 0.03002061 | 5.23533476 | 1.65E-07 |
| 208650 | CBLB | 1668.10795 | -0.6597282 | 0.1260227 | -5.2349953 | 1.65E-07 |
| 17222 | ANAPC1 | 9137.90394 | 0.53324574 | 0.10190767 | 5.23263572 | 1.67E-07 |
| 210297 | LRCH2 | 309.046879 | 4.40049683 | 0.84102821 | 5.23228206 | 1.67E-07 |
| 77035 | KDM8 | 239.640961 | 0.90373148 | 0.17274301 | 5.23165305 | 1.68E-07 |
| 11736 | ANKFY1 | 4626.03845 | -0.3530856 | 0.06749175 | -5.231538 | 1.68E-07 |
| 100039419 | Gm2223 | 1945.65611 | -0.5436754 | 0.10394218 | -5.2305565 | 1.69E-07 |
| 227331 | GIGYF2 | 3158.8443 | -0.2869795 | 0.05487144 | -5.2300346 | 1.69E-07 |
| 66435 | UGGT2 | 386.73314 | -8.4014939 | 1.60724588 | -5.2272611 | 1.72E-07 |
| 71770 | AP2B1 | 7213.52123 | -0.4092664 | 0.0782958 | -5.227182 | 1.72E-07 |
| 13350 | DGAT1 | 530.029752 | -0.6868445 | 0.13140151 | -5.2270668 | 1.72E-07 |
| 14201 | FHL3 | 1331.44547 | 0.90659153 | 0.17346954 | 5.22622905 | 1.73E-07 |
| 319211 | NOL4 | 71.7287666 | 2.51858012 | 0.48191679 | 5.2261722 | 1.73E-07 |
| 75739 | MPP7 | 693.796787 | -0.428645 | 0.08202567 | -5.2257416 | 1.73E-07 |
| 213417 | KLHDC8A | 438.298687 | 1.75364617 | 0.33579479 | 5.22237454 | 1.77E-07 |
| 74485 | LRRC71 | 13.8746786 | 5.80144317 | 1.1112037 | 5.22086378 | 1.78E-07 |
| 74098 | 0610037L13I | 473.295579 | 0.95938782 | 0.18395813 | 5.2152509 | 1.84E-07 |
| 320635 | CYB5R2 | 8.0423846 | -6.4342371 | 1.23413224 | -5.2135718 | 1.85E-07 |
| 382056 | CRTC1 | 921.440902 | 0.37529962 | 0.07198882 | 5.21330445 | 1.86E-07 |

| | | | | | | |
|---|---|---|---|---|---|---|
| 104601 | mycbpap | 13.1978051 | -5.6833378 | 1.09018648 | -5.2131795 | 1.86E-07 |
| 277396 | KLHL23 | 130.215367 | 1.16248908 | 0.22309003 | 5.21085168 | 1.88E-07 |
| 68918 | 1190005I06R | 109.004514 | -3.0352789 | 0.58257609 | -5.2100987 | 1.89E-07 |
| 66448 | MRPL20 | 1286.05744 | 0.85669587 | 0.1644596 | 5.20915689 | 1.90E-07 |
| 14159 | FES | 367.89397 | 1.30109533 | 0.2498959 | 5.20654942 | 1.92E-07 |
| 76763 | MOSPD2 | 636.170938 | -0.5271485 | 0.10125735 | -5.2060269 | 1.93E-07 |
| 20301 | Ccl27a | 10.103924 | -6.0262205 | 1.15784201 | -5.2047002 | 1.94E-07 |
| 223754 | TBC1D22A | 784.978464 | -0.3320677 | 0.06380625 | -5.2043124 | 1.95E-07 |
| 17287 | MEP1A | 26.530623 | -2.8153764 | 0.54102196 | -5.2038117 | 1.95E-07 |
| 54151 | CYHR1 | 915.658278 | -0.4844439 | 0.09312983 | -5.2018122 | 1.97E-07 |
| 210766 | BRCC3 | 897.053894 | -0.3529661 | 0.06787736 | -5.2000558 | 1.99E-07 |
| 99543 | OLFML3 | 274.385913 | 1.75655961 | 0.3378397 | 5.19938786 | 2.00E-07 |
| 11555 | ADRB2 | 79.168331 | -1.3920977 | 0.26782079 | -5.1978701 | 2.02E-07 |
| 54725 | CADM1 | 2839.9046 | -4.7274902 | 0.90955005 | -5.1976141 | 2.02E-07 |
| 14082 | FADD | 599.202075 | 0.87749459 | 0.16884655 | 5.19699436 | 2.03E-07 |
| 64010 | SAV1 | 3472.52784 | 0.58797139 | 0.11314554 | 5.19659368 | 2.03E-07 |
| 20726 | Serpinb9d | 12.2540196 | -4.2153805 | 0.81153631 | -5.1943215 | 2.05E-07 |
| 69812 | Abhd11os | 7.39567883 | -6.3133992 | 1.2156041 | -5.1936311 | 2.06E-07 |
| 216443 | MARS | 3857.54632 | 0.18197186 | 0.03506154 | 5.19007106 | 2.10E-07 |
| 230126 | SHB | 961.223107 | -1.282874 | 0.2472844 | -5.1878485 | 2.13E-07 |
| 192236 | HPS1 | 551.903058 | -0.8194494 | 0.15800441 | -5.186244 | 2.15E-07 |
| 75607 | WNK2 | 65.1532515 | -7.5171836 | 1.44951003 | -5.1860169 | 2.15E-07 |
| 110809 | SRSF1 | 13683.659 | 0.44184304 | 0.08520284 | 5.18577833 | 2.15E-07 |
| 73825 | PPP1R21 | 1033.52427 | -0.3244168 | 0.0625627 | -5.1854661 | 2.15E-07 |
| 329795 | TMEM67 | 259.547164 | -1.2833118 | 0.24749052 | -5.1852969 | 2.16E-07 |
| 225288 | FHOD3 | 14.7819138 | -3.2587381 | 0.62854113 | -5.1846059 | 2.16E-07 |
| 80288 | BCL9L | 3378.76813 | 0.37143576 | 0.07164497 | 5.18439423 | 2.17E-07 |
| 69227 | SELT | 6400.44629 | -0.3101537 | 0.05982538 | -5.1843168 | 2.17E-07 |
| 20525 | SLC2A1 | 3118.05948 | 0.44921232 | 0.08665773 | 5.18375387 | 2.17E-07 |
| 27966 | RRP9 | 1276.44619 | 0.52363213 | 0.1010482 | 5.18200377 | 2.20E-07 |
| 14865 | GSTM4 | 98.0271509 | -2.3414479 | 0.45196324 | -5.1806159 | 2.21E-07 |
| 72123 | CCDC71L | 828.366131 | 0.61162732 | 0.11806641 | 5.18036669 | 2.21E-07 |
| 16453 | JAK3 | 196.8566 | -1.6206682 | 0.3129049 | -5.1794274 | 2.23E-07 |
| 238011 | ENPP7 | 8.91894715 | -6.5823693 | 1.27104713 | -5.178698 | 2.23E-07 |
| 76559 | atg2b | 3154.00812 | 0.72887165 | 0.14075124 | 5.17843846 | 2.24E-07 |
| 70012 | CEP85 | 1060.03092 | 0.8342 | 0.16110908 | 5.17785854 | 2.24E-07 |
| 22385 | BAZ1B | 8396.11975 | 0.36045896 | 0.0696654 | 5.17414601 | 2.29E-07 |
| 208869 | DOCK3 | 38.8872483 | -8.7106467 | 1.68403768 | -5.1724773 | 2.31E-07 |
| 104732 | 4930427A07 | 1341.97035 | 0.37717079 | 0.07295048 | 5.17023037 | 2.34E-07 |
| 51801 | RAMP1 | 20.2576311 | -7.7691521 | 1.50410232 | -5.1653082 | 2.40E-07 |
| 13629 | EEF2 | 77270.5397 | -0.1664661 | 0.03222798 | -5.1652659 | 2.40E-07 |
| 109689 | ARRB1 | 331.87481 | -0.6956954 | 0.13471221 | -5.1643085 | 2.41E-07 |
| 74131 | SASH3 | 10.9442365 | -4.3816134 | 0.84875686 | -5.1623894 | 2.44E-07 |
| 381232 | Mirt1 | 115.123465 | -2.762977 | 0.53528715 | -5.1616726 | 2.45E-07 |
| 67236 | cinp | 1136.22664 | 0.38553877 | 0.07471667 | 5.16001002 | 2.47E-07 |

| | | | | | | |
|---|---|---|---|---|---|---|
| 71870 | CFAP45 | 13.9715813 | -7.2367483 | 1.4027001 | -5.1591558 | 2.48E-07 |
| 207175 | cetn4 | 26.8593371 | -1.7026061 | 0.33023581 | -5.1557284 | 2.53E-07 |
| 67073 | PI4K2B | 1955.53684 | -0.4582904 | 0.08890111 | -5.1550581 | 2.54E-07 |
| 17957 | NAPB | 132.929506 | -0.9764786 | 0.18942599 | -5.1549349 | 2.54E-07 |
| 100504104 | Gm16062 | 62.4735213 | 1.29573317 | 0.25144107 | 5.15322804 | 2.56E-07 |
| 12560 | CDH3 | 10.6077129 | -4.341537 | 0.84279412 | -5.1513613 | 2.59E-07 |
| 72354 | TTC4 | 938.415756 | 0.71874112 | 0.13953911 | 5.15082183 | 2.59E-07 |
| 621430 | Vmn2r79 | 11.3477904 | -4.8512661 | 0.94225341 | -5.148579 | 2.62E-07 |
| 93684 | 41896 | 7437.39707 | -0.2822886 | 0.05484408 | -5.1471119 | 2.65E-07 |
| 26909 | EXO1 | 1270.17652 | 0.55769515 | 0.10841883 | 5.14389587 | 2.69E-07 |
| 76917 | FLYWCH2 | 10.8636835 | -6.8680385 | 1.33530784 | -5.1434121 | 2.70E-07 |
| 17389 | MMP16 | 93.8859232 | -2.1244175 | 0.41324809 | -5.14078 | 2.74E-07 |
| 224250 | CLDND1 | 2104.48543 | 0.53313422 | 0.1037571 | 5.1382916 | 2.77E-07 |
| 170676 | PEG10 | 1384.07283 | 5.05569235 | 0.98404379 | 5.13767009 | 2.78E-07 |
| 74026 | MSL1 | 2282.49053 | 0.43372605 | 0.0844333 | 5.13690751 | 2.79E-07 |
| 59045 | STARD3 | 611.554191 | -0.6024171 | 0.11727918 | -5.1366069 | 2.80E-07 |
| 117150 | PIP4K2C | 3505.71796 | -0.8048065 | 0.15668109 | -5.1365895 | 2.80E-07 |
| 98193 | dcaf8 | 2537.28954 | -0.268575 | 0.05229769 | -5.1355041 | 2.81E-07 |
| 71375 | FOXN3 | 487.938106 | 0.74748103 | 0.14556292 | 5.13510611 | 2.82E-07 |
| 238505 | MTR | 1092.12848 | 1.29114003 | 0.2515604 | 5.13252506 | 2.86E-07 |
| 100271882 | Gm14139 | 10.4777647 | 4.03635186 | 0.78653674 | 5.1318033 | 2.87E-07 |
| 19139 | PRPS1 | 1552.35122 | 0.58910113 | 0.11482263 | 5.13053148 | 2.89E-07 |
| 13872 | ERCC3 | 1954.07523 | -0.4436357 | 0.0864796 | -5.1299462 | 2.90E-07 |
| 16913 | PSMB8 | 284.776879 | -2.1893095 | 0.42680653 | -5.1295126 | 2.90E-07 |
| 83768 | DPP7 | 182.111715 | 1.94924427 | 0.38021497 | 5.12668994 | 2.95E-07 |
| 211586 | TFDP2 | 757.638024 | 0.52713911 | 0.10284071 | 5.12578228 | 2.96E-07 |
| 18972 | POLD2 | 1206.01306 | -0.3597651 | 0.07020767 | -5.1242993 | 2.99E-07 |
| 99031 | osbpl6 | 15.8830747 | 5.39358831 | 1.05255849 | 5.12426467 | 2.99E-07 |
| 239839 | CCDC14 | 416.072213 | -0.6186474 | 0.12075408 | -5.123201 | 3.00E-07 |
| 102871 | D330045A20 | 10.0360086 | -6.0199976 | 1.17508998 | -5.1230099 | 3.01E-07 |
| 638566 | TRAV13-2 | 11.852387 | -4.1567067 | 0.81142471 | -5.1227263 | 3.01E-07 |
| 100910 | chpf2 | 3446.51261 | 0.89467488 | 0.17466111 | 5.12234728 | 3.02E-07 |
| 98956 | NAT10 | 2280.68636 | 0.48605713 | 0.09490715 | 5.1213962 | 3.03E-07 |
| 67510 | TVP23B | 831.358204 | -0.5937955 | 0.11594991 | -5.1211383 | 3.04E-07 |
| 17089 | LYAR | 1621.30948 | 0.66078324 | 0.12913161 | 5.11712994 | 3.10E-07 |
| 665270 | PLB1 | 22.1989674 | -2.1863542 | 0.42737556 | -5.115768 | 3.12E-07 |
| 12291 | CACNA1G | 96.2711916 | 4.75270502 | 0.92918446 | 5.11492088 | 3.14E-07 |
| 170644 | UBN1 | 3319.88439 | 0.54397192 | 0.10635751 | 5.11456037 | 3.14E-07 |
| 12354 | Car7 | 12.5110706 | -3.1680379 | 0.61957647 | -5.1132316 | 3.17E-07 |
| 22361 | VNN1 | 15.2787283 | 2.995435 | 0.58601134 | 5.11156488 | 3.20E-07 |
| 320237 | Smim10l2a | 23.4710308 | -5.4907176 | 1.07429323 | -5.1110046 | 3.20E-07 |
| 20663 | SOS2 | 1312.76124 | -0.4027919 | 0.07881418 | -5.1106526 | 3.21E-07 |
| 72454 | CCDC71 | 920.578177 | 0.47413286 | 0.09280002 | 5.10918894 | 3.24E-07 |
| 74440 | CMIP | 2528.1733 | -0.3450008 | 0.06754108 | -5.1080141 | 3.26E-07 |
| 54484 | MKRN1 | 1025.74473 | -0.4393635 | 0.0860211 | -5.1076252 | 3.26E-07 |

| | | | | | | |
|---|---|---|---|---|---|---|
| 193385 | FAM65B | 16.3506924 | -6.0223627 | 1.1799335 | -5.1039849 | 3.33E-07 |
| 100040724 | Mirg | 8.93395075 | 6.63846458 | 1.30098393 | 5.10264918 | 3.35E-07 |
| 223739 | 5031439G07 | 3655.28278 | -0.3832524 | 0.07514293 | -5.1003118 | 3.39E-07 |
| 16627 | Klra1 | 140.708077 | -1.8836173 | 0.36937285 | -5.0995012 | 3.41E-07 |
| 18041 | NFS1 | 889.562142 | -0.4435493 | 0.08698017 | -5.0994302 | 3.41E-07 |
| 15452 | hprT | 3671.24944 | -0.4061022 | 0.07965492 | -5.0982685 | 3.43E-07 |
| 74270 | USP20 | 681.518137 | -0.6888056 | 0.13513732 | -5.0970792 | 3.45E-07 |
| 50766 | CRIM1 | 5620.33771 | -0.8495596 | 0.16674548 | -5.0949485 | 3.49E-07 |
| 66129 | AAED1 | 385.065888 | 0.86421869 | 0.16962714 | 5.09481361 | 3.49E-07 |
| 433956 | DNAAF5 | 1165.6458 | -0.5344505 | 0.10490762 | -5.094487 | 3.50E-07 |
| 235533 | GK5 | 80.211538 | 0.93485906 | 0.18351663 | 5.09413804 | 3.50E-07 |
| 21351 | TALDO1 | 6530.23879 | 0.58724003 | 0.11528498 | 5.09381228 | 3.51E-07 |
| 66566 | NTPCR | 154.78035 | -1.7160494 | 0.33694775 | -5.0929243 | 3.53E-07 |
| 74316 | ISCA2 | 364.528301 | 0.44045239 | 0.08649199 | 5.09240692 | 3.54E-07 |
| 16531 | KCNMA1 | 38.9719408 | -5.9045676 | 1.15957202 | -5.0920232 | 3.54E-07 |
| 71707 | UBIAD1 | 766.049245 | 0.62372147 | 0.1224951 | 5.09180737 | 3.55E-07 |
| 74200 | 2810403A07 | 2285.54385 | -0.4585757 | 0.0900663 | -5.091535 | 3.55E-07 |
| 69241 | POLR2D | 873.725292 | 0.33747553 | 0.06630287 | 5.08990827 | 3.58E-07 |
| 244238 | MRGPRE | 137.20828 | -2.206657 | 0.43368658 | -5.0881377 | 3.62E-07 |
| 414758 | Zfp950 | 108.497781 | -0.8450111 | 0.16608496 | -5.0878245 | 3.62E-07 |
| 668225 | FIGNL2 | 64.0480383 | -2.8486109 | 0.56004161 | -5.0864273 | 3.65E-07 |
| 19270 | PTPRG | 3951.88653 | -0.5891553 | 0.1158462 | -5.0856679 | 3.66E-07 |
| 75423 | ARL5A | 2803.24766 | 0.30595012 | 0.06016685 | 5.08502781 | 3.68E-07 |
| 12840 | COL9A2 | 13.0932375 | -5.66362 | 1.11395645 | -5.0842382 | 3.69E-07 |
| 17858 | MX2 | 37.9914718 | -4.1500493 | 0.81698538 | -5.0797106 | 3.78E-07 |
| 77809 | LRRC42 | 916.617794 | 0.54187205 | 0.10670891 | 5.07803948 | 3.81E-07 |
| 18788 | SERPINB2 | 1628.23442 | -4.3548045 | 0.85790242 | -5.0761071 | 3.85E-07 |
| 72160 | TMEM163 | 143.530416 | -10.595369 | 2.08734477 | -5.0760036 | 3.85E-07 |
| 69562 | CDK13 | 2997.19955 | 0.42647129 | 0.08402695 | 5.07541097 | 3.87E-07 |
| 217138 | PRR15L | 6.6082425 | -6.1560679 | 1.21359212 | -5.0726005 | 3.92E-07 |
| 68828 | SYNC | 125.184558 | -2.686978 | 0.52976154 | -5.0720518 | 3.94E-07 |
| 100041106 | Gm3141 | 289.634622 | 0.58834221 | 0.11601461 | 5.07127686 | 3.95E-07 |
| 246277 | CSAD | 230.193912 | -0.8797548 | 0.173493 | -5.0708375 | 3.96E-07 |
| 76633 | 1700112E06I | 24.9750532 | -3.0053334 | 0.59279514 | -5.0697672 | 3.98E-07 |
| 68428 | STEAP3 | 4124.13701 | -1.3104326 | 0.2585475 | -5.0684405 | 4.01E-07 |
| 75909 | VMP1 | 3352.49984 | -0.406318 | 0.08019614 | -5.0665534 | 4.05E-07 |
| 68499 | MRPL53 | 703.285273 | 0.69230004 | 0.13665132 | 5.06617899 | 4.06E-07 |
| 57265 | FZD2 | 1557.50448 | 0.53498134 | 0.10561971 | 5.06516561 | 4.08E-07 |
| 15284 | HLX | 118.462887 | 1.46882885 | 0.29001317 | 5.06469708 | 4.09E-07 |
| 78405 | ntf5 | 30.7264508 | -2.1632325 | 0.42732451 | -5.0622711 | 4.14E-07 |
| 17755 | MAP1B | 6243.98826 | -0.9213458 | 0.18201605 | -5.0618934 | 4.15E-07 |
| 14700 | GNG10 | 1440.79674 | 0.86872383 | 0.17165857 | 5.06076594 | 4.18E-07 |
| 51800 | BOK | 819.468066 | 1.39229951 | 0.27522366 | 5.05879292 | 4.22E-07 |
| 73419 | ARMT1 | 931.735748 | 0.40322035 | 0.07971788 | 5.05809174 | 4.23E-07 |
| 16882 | LIG3 | 2338.14142 | 0.20144604 | 0.03983685 | 5.05677686 | 4.26E-07 |

| | | | | | | |
|---|---|---|---|---|---|---|
| 68203 | DIRAS2 | 18.0659199 | -5.5455235 | 1.09668829 | -5.0566086 | 4.27E-07 |
| 19340 | RAB3D | 116.260435 | -2.0643815 | 0.40829117 | -5.0561504 | 4.28E-07 |
| 54160 | COPG2 | 1048.34911 | -0.8067821 | 0.1595688 | -5.0560143 | 4.28E-07 |
| 17127 | SMAD3 | 1439.11717 | -0.7825059 | 0.15479207 | -5.0552065 | 4.30E-07 |
| 69654 | DCTN2 | 4413.77939 | -0.4591551 | 0.09083032 | -5.0550865 | 4.30E-07 |
| 74106 | DCAF6 | 887.680436 | -0.7590847 | 0.1501671 | -5.0549335 | 4.31E-07 |
| 99696 | ANKRD50 | 1865.94637 | 0.49046997 | 0.09702895 | 5.05488275 | 4.31E-07 |
| 109674 | AMPD2 | 2714.65782 | 0.37270271 | 0.07383764 | 5.04759786 | 4.47E-07 |
| 22351 | VILL | 56.2666687 | -5.1865316 | 1.02763831 | -5.04704 | 4.49E-07 |
| 72560 | NAALAD2 | 11.7834018 | 5.56251394 | 1.10222613 | 5.04661775 | 4.50E-07 |
| 218820 | Zfp503 | 1597.68409 | 0.23428237 | 0.04642742 | 5.04620729 | 4.51E-07 |
| 93840 | VANGL2 | 243.706188 | 0.92430398 | 0.18317061 | 5.04613688 | 4.51E-07 |
| 327942 | PIGL | 210.906574 | -0.5739412 | 0.11383087 | -5.0420522 | 4.61E-07 |
| 70997 | SPEF1 | 202.605098 | 1.46373477 | 0.29033117 | 5.04160395 | 4.62E-07 |
| 26428 | ORC4 | 1611.18311 | -0.3918552 | 0.07773002 | -5.0412336 | 4.63E-07 |
| 244650 | PHLPP2 | 870.991969 | -0.3126789 | 0.06202989 | -5.0407779 | 4.64E-07 |
| 100503915 | SMPD5 | 8.21520457 | -5.726099 | 1.13597907 | -5.040673 | 4.64E-07 |
| 235169 | FOXRED1 | 453.153794 | 0.54342163 | 0.10782025 | 5.04007037 | 4.65E-07 |
| 74196 | TTC27 | 1226.41318 | 0.2928695 | 0.05812255 | 5.03882712 | 4.68E-07 |
| 243339 | TMEM130 | 9.8943506 | -3.6002039 | 0.71449999 | -5.0387738 | 4.69E-07 |
| 18563 | Pcx | 544.563468 | 0.62245076 | 0.12353746 | 5.03855887 | 4.69E-07 |
| 217721 | Mfsd7c | 13.322072 | -5.7090824 | 1.13321646 | -5.0379452 | 4.71E-07 |
| 102182 | PRMT9 | 745.660095 | -0.3197274 | 0.06349251 | -5.0356714 | 4.76E-07 |
| 100038412 | Faddos | 13.2339609 | -4.0311683 | 0.80070828 | -5.034503 | 4.79E-07 |
| 22154 | TUBB5 | 48735.5635 | 0.32089255 | 0.06374176 | 5.03425938 | 4.80E-07 |
| 11545 | PARP1 | 4474.128 | 0.40991971 | 0.08145119 | 5.032704 | 4.84E-07 |
| 76916 | TIMMDC1 | 574.386398 | -0.640512 | 0.1272742 | -5.0325358 | 4.84E-07 |
| 71723 | DHX34 | 808.726324 | 0.86317298 | 0.17158106 | 5.03070093 | 4.89E-07 |
| 77683 | EHMT1 | 1872.66362 | 0.35037216 | 0.0696659 | 5.0293205 | 4.92E-07 |
| 399510 | MAP4K5 | 3918.74475 | 0.7474484 | 0.14865942 | 5.02792492 | 4.96E-07 |
| 13361 | DHFR | 1917.95671 | 0.54258687 | 0.10791844 | 5.02774962 | 4.96E-07 |
| 69773 | 1810026J23F | 1456.35984 | 0.57448197 | 0.11426351 | 5.02769404 | 4.96E-07 |
| 621961 | Anp32b-ps1 | 595.147988 | 0.69791132 | 0.13882094 | 5.02742126 | 4.97E-07 |
| 14776 | GPX2 | 104.570714 | -10.138255 | 2.01742851 | -5.0253353 | 5.03E-07 |
| 11430 | ACOX1 | 3221.67898 | -0.7608751 | 0.15145977 | -5.0236121 | 5.07E-07 |
| 12934 | DPYSL2 | 2255.80809 | 0.90074695 | 0.17940639 | 5.02070708 | 5.15E-07 |
| 16783 | LAMP1 | 28241.2174 | -0.8838365 | 0.17623865 | -5.0149983 | 5.30E-07 |
| 73451 | Zfp763 | 144.234353 | -0.6928538 | 0.13817635 | -5.0142718 | 5.32E-07 |
| 245174 | Zfp937 | 157.995737 | -1.2618316 | 0.25168789 | -5.0134779 | 5.35E-07 |
| 107476 | ACACA | 1721.66935 | 0.63802231 | 0.12732099 | 5.01113201 | 5.41E-07 |
| 18148 | NPM1 | 9659.93068 | 0.39311274 | 0.07850022 | 5.0077913 | 5.51E-07 |
| 66208 | NENF | 636.620433 | -0.5781845 | 0.11546721 | -5.0073482 | 5.52E-07 |
| 18021 | NFATC3 | 3787.60088 | 0.3701233 | 0.07393394 | 5.00613528 | 5.55E-07 |
| 227446 | 2310035C23I | 1477.41463 | -0.7549664 | 0.15083447 | -5.0052643 | 5.58E-07 |
| 12428 | CCNA2 | 7559.56362 | 0.50821793 | 0.1015447 | 5.00486929 | 5.59E-07 |

| | | | | | | |
|---|---|---|---|---|---|---|
| 271457 | RAB5A | 1475.45769 | -0.3721634 | 0.07437926 | -5.0035907 | 5.63E-07 |
| 666038 | Gm7902 | 44.8528765 | -1.1711505 | 0.23419409 | -5.0007687 | 5.71E-07 |
| 78323 | FAM219B | 446.810256 | 0.62413912 | 0.12485043 | 4.99909453 | 5.76E-07 |
| 100042918 | Uqcrh-ps1 | 116.249694 | 0.89778006 | 0.17959124 | 4.9990193 | 5.76E-07 |
| 214253 | ETNK2 | 6.04906572 | -6.0276088 | 1.20586617 | -4.9985719 | 5.78E-07 |
| 233871 | ATXN2L | 6716.99095 | 0.61969186 | 0.12402902 | 4.9963459 | 5.84E-07 |
| 53378 | SDCBP | 6927.93657 | -1.2024962 | 0.24067986 | -4.9962477 | 5.85E-07 |
| 66894 | WWP2 | 3060.12759 | 0.5908631 | 0.1182651 | 4.9960901 | 5.85E-07 |
| 51875 | tmem141 | 113.549306 | -1.5150368 | 0.30327964 | -4.995511 | 5.87E-07 |
| 14009 | ETV1 | 616.473189 | 1.83594592 | 0.36755502 | 4.99502337 | 5.88E-07 |
| 21416 | TCF7L2 | 570.32075 | 0.57998772 | 0.11611986 | 4.99473324 | 5.89E-07 |
| 330695 | CTXN1 | 395.46527 | -1.1193352 | 0.22413321 | -4.994062 | 5.91E-07 |
| 381677 | VGF | 30.9639267 | -2.0961774 | 0.41985615 | -4.9926085 | 5.96E-07 |
| 26399 | MAP2K6 | 87.6764466 | 1.2411551 | 0.24863734 | 4.99182911 | 5.98E-07 |
| 194388 | tet3 | 1689.87118 | 0.60869335 | 0.12197827 | 4.99017861 | 6.03E-07 |
| 16773 | LAMA2 | 14.1685886 | -2.606582 | 0.52241479 | -4.9894873 | 6.05E-07 |
| 72308 | BRF1 | 1225.43597 | 0.3690444 | 0.07400296 | 4.98688696 | 6.14E-07 |
| 16398 | ITGA2 | 3039.35371 | -1.9450531 | 0.39009781 | -4.9860651 | 6.16E-07 |
| 66437 | FIS1 | 1487.97808 | -0.4657767 | 0.09342306 | -4.9856715 | 6.17E-07 |
| 68015 | TRAP1 | 3088.78071 | -0.2592169 | 0.0519993 | -4.985008 | 6.20E-07 |
| 216190 | appl2 | 977.332847 | -0.5412047 | 0.10857137 | -4.9847829 | 6.20E-07 |
| 56297 | ARL6 | 535.442906 | -0.6040177 | 0.12117283 | -4.9847615 | 6.20E-07 |
| 13864 | NR2F6 | 1382.72705 | -0.5141331 | 0.10316904 | -4.9834045 | 6.25E-07 |
| 634650 | Gbp11 | 8.86535748 | -5.8353542 | 1.17107529 | -4.9829027 | 6.26E-07 |
| 13642 | EFNB2 | 7102.11117 | -0.7350192 | 0.14755866 | -4.9812001 | 6.32E-07 |
| 66317 | WDR61 | 1371.32827 | 0.4959165 | 0.09956204 | 4.98097962 | 6.33E-07 |
| 114601 | EHBP1L1 | 3845.00499 | 2.04556265 | 0.41071001 | 4.98055221 | 6.34E-07 |
| 19267 | PTPRE | 1199.44582 | 1.10469245 | 0.22191344 | 4.97803303 | 6.42E-07 |
| 66259 | CAMK2N1 | 46.1199591 | -3.0830301 | 0.61952023 | -4.9764801 | 6.48E-07 |
| 69583 | TNFSF13 | 14.2609951 | -4.4045973 | 0.88511198 | -4.9763164 | 6.48E-07 |
| 433931 | PIGG | 449.904101 | -0.7947574 | 0.15972966 | -4.9756408 | 6.50E-07 |
| 215193 | DIEXF | 921.879974 | 0.56140963 | 0.11284513 | 4.97504535 | 6.52E-07 |
| 12215 | BSG | 13621.2951 | -0.5928542 | 0.11918456 | -4.9742536 | 6.55E-07 |
| 240888 | GPR161 | 309.632746 | -0.5799267 | 0.11660106 | -4.9735975 | 6.57E-07 |
| 100503583 | FSBP | 23.835877 | -1.8426829 | 0.37060867 | -4.9720448 | 6.63E-07 |
| 631797 | FER1L6 | 28.3173811 | -5.7874709 | 1.16418758 | -4.9712529 | 6.65E-07 |
| 224014 | FGD4 | 144.758942 | -2.751981 | 0.55366763 | -4.9704567 | 6.68E-07 |
| 72393 | FAIM2 | 85.258975 | -7.4608755 | 1.5011425 | -4.9701314 | 6.69E-07 |
| 20416 | SHC1 | 2986.48799 | 0.48646012 | 0.0978874 | 4.96958846 | 6.71E-07 |
| 101835 | AW146154 | 62.1411196 | -1.3592928 | 0.27353944 | -4.9692754 | 6.72E-07 |
| 66627 | OGFOD2 | 325.908294 | -0.8477032 | 0.17060057 | -4.9689353 | 6.73E-07 |
| 27204 | SYN3 | 21.2074738 | -6.3660686 | 1.28124005 | -4.9686775 | 6.74E-07 |
| 52187 | RRAGD | 257.925229 | -4.0082076 | 0.8067276 | -4.9684771 | 6.75E-07 |
| 27403 | ABCA7 | 1216.07524 | -1.0550714 | 0.2123654 | -4.9681886 | 6.76E-07 |
| 20787 | SREBF1 | 3722.83501 | -0.3073317 | 0.06187254 | -4.9671747 | 6.79E-07 |

| | | | | | | |
|---|---|---|---|---|---|---|
| 27407 | ABCF2 | 5258.74689 | 0.24881484 | 0.05009437 | 4.96692209 | 6.80E-07 |
| 100702 | gbp6 | 15.6427546 | -2.8854435 | 0.58094721 | -4.9667912 | 6.81E-07 |
| 215008 | VEZT | 843.463884 | -0.4882961 | 0.09833712 | -4.9655315 | 6.85E-07 |
| 12075 | BFSP1 | 22.3735086 | -2.3986743 | 0.48315919 | -4.9645631 | 6.89E-07 |
| 80902 | Zfp202 | 235.38719 | 0.60742893 | 0.12236925 | 4.96390158 | 6.91E-07 |
| 64945 | CLDN12 | 2422.08411 | -0.6614531 | 0.13326797 | -4.9633312 | 6.93E-07 |
| 56458 | FOXO1 | 983.565035 | -1.1751722 | 0.23682892 | -4.9621144 | 6.97E-07 |
| 621080 | AI429214 | 102.253907 | -1.2028799 | 0.2424627 | -4.9610927 | 7.01E-07 |
| 75475 | OPLAH | 79.5925954 | -2.0371826 | 0.41069202 | -4.9603657 | 7.04E-07 |
| 68927 | PTCD2 | 774.298834 | 0.74109839 | 0.14943667 | 4.95928058 | 7.08E-07 |
| 208092 | CHMP6 | 1572.49103 | -0.6740597 | 0.13592592 | -4.9590227 | 7.08E-07 |
| 23970 | PACSIN2 | 5472.35191 | -0.2244061 | 0.0452623 | -4.9579023 | 7.13E-07 |
| 11835 | ar | 73.6273484 | -4.6057789 | 0.92901246 | -4.9577148 | 7.13E-07 |
| 14594 | GGTA1 | 2122.61126 | -0.7077365 | 0.1427552 | -4.9576931 | 7.13E-07 |
| 58994 | SMPD3 | 7.66953875 | -5.6207393 | 1.13387922 | -4.9570882 | 7.16E-07 |
| 67120 | TTC14 | 1943.92021 | -0.4834414 | 0.09754227 | -4.9562248 | 7.19E-07 |
| 20869 | STK11 | 1905.68018 | -0.4369302 | 0.08816494 | -4.9558265 | 7.20E-07 |
| 66144 | ATP6V1F | 868.763773 | -0.835796 | 0.16868141 | -4.9548791 | 7.24E-07 |
| 230861 | EIF4G3 | 6505.8804 | 0.78823844 | 0.15913796 | 4.95317673 | 7.30E-07 |
| 15451 | HPN | 15.8292972 | -5.9430236 | 1.19988811 | -4.9529815 | 7.31E-07 |
| 70166 | LIPN | 26.6969543 | -4.8772697 | 0.98481108 | -4.9524927 | 7.33E-07 |
| 83770 | tas1r2 | 16.817971 | 4.03129583 | 0.81421196 | 4.95116261 | 7.38E-07 |
| 12167 | BMPR1B | 15.9884456 | 3.07754393 | 0.62181843 | 4.94926459 | 7.45E-07 |
| 20193 | S100A1 | 390.421687 | -0.6372465 | 0.12875726 | -4.9492087 | 7.45E-07 |
| 68709 | CILP2 | 31.3792747 | -3.5926648 | 0.72617718 | -4.9473667 | 7.52E-07 |
| 23849 | KLF6 | 7715.90323 | -0.8898046 | 0.17987518 | -4.946789 | 7.54E-07 |
| 17714 | grpel2 | 953.166103 | -0.7957468 | 0.16086967 | -4.946531 | 7.55E-07 |
| 70208 | MED23 | 1420.24047 | -0.2603794 | 0.05266947 | -4.9436485 | 7.67E-07 |
| 78309 | CUL9 | 51.3312394 | -4.7207831 | 0.95495614 | -4.9434554 | 7.67E-07 |
| 19220 | PTGFR | 41.282105 | 5.27122918 | 1.06634499 | 4.94326811 | 7.68E-07 |
| 11477 | ACVR1 | 1499.84664 | 0.44073729 | 0.08916048 | 4.94319094 | 7.69E-07 |
| 71302 | ARHGAP26 | 118.092173 | -1.8911167 | 0.38260367 | -4.9427564 | 7.70E-07 |
| 50797 | COPB2 | 7258.89487 | 0.35284086 | 0.07139793 | 4.94189209 | 7.74E-07 |
| 16418 | EIF6 | 5564.39693 | -0.6472346 | 0.1309886 | -4.9411521 | 7.77E-07 |
| 70375 | ICA1L | 10.9333165 | -3.3070869 | 0.66950202 | -4.9396221 | 7.83E-07 |
| 75729 | FAM227A | 13.2556364 | -3.0860237 | 0.62501728 | -4.9375014 | 7.91E-07 |
| 59022 | EDF1 | 1831.09191 | -0.5278051 | 0.10693121 | -4.9359316 | 7.98E-07 |
| 20866 | STIM1 | 1185.80439 | -0.6155283 | 0.12471355 | -4.9355369 | 7.99E-07 |
| 14751 | Gpi1 | 5696.65346 | 0.46293465 | 0.09381936 | 4.93431896 | 8.04E-07 |
| 52653 | NUDCD2 | 2173.15586 | 0.23661786 | 0.0479762 | 4.93198432 | 8.14E-07 |
| 56381 | SPEN | 3271.19649 | 0.70535732 | 0.14303785 | 4.93126358 | 8.17E-07 |
| 23829 | C1QL1 | 10.7423072 | 5.45088803 | 1.10537948 | 4.93123686 | 8.17E-07 |
| 633640 | TMEM267 | 474.964161 | 0.92336383 | 0.18726569 | 4.93076898 | 8.19E-07 |
| 226977 | ACTR1B | 3607.95421 | -0.2338724 | 0.04743244 | -4.930642 | 8.20E-07 |
| 331374 | DGKK | 251.318518 | -9.9839687 | 2.02503704 | -4.9302648 | 8.21E-07 |

| | | | | | | |
|---|---|---|---|---|---|---|
| 13654 | EGR2 | 61.7847993 | 2.14413483 | 0.43519033 | 4.92688992 | 8.35E-07 |
| 72552 | HSDL1 | 943.474752 | -0.6686219 | 0.13571425 | -4.9266892 | 8.36E-07 |
| 71584 | GDPD2 | 10.6938582 | -4.7645093 | 0.96719537 | -4.9261085 | 8.39E-07 |
| 17183 | MATN4 | 18.5918884 | -2.7019737 | 0.54862628 | -4.9249805 | 8.44E-07 |
| 212114 | NHLRC3 | 428.701371 | -0.9849027 | 0.19998122 | -4.9249762 | 8.44E-07 |
| 14860 | GSTA4 | 2209.12586 | -0.949422 | 0.1928343 | -4.9235122 | 8.50E-07 |
| 21975 | TOP3A | 811.798003 | 0.30902725 | 0.06277007 | 4.92316232 | 8.52E-07 |
| 69547 | NKPD1 | 17.5403787 | -2.6666245 | 0.54171147 | -4.9225919 | 8.54E-07 |
| 77700 | 9130208D14 | 6.77756859 | -6.1880207 | 1.25711413 | -4.9224017 | 8.55E-07 |
| 225215 | RSL24D1 | 713.629752 | 0.5931704 | 0.12051765 | 4.92185492 | 8.57E-07 |
| 216705 | CLINT1 | 5140.18516 | -0.2662601 | 0.05410308 | -4.9213481 | 8.60E-07 |
| 66109 | TSPAN13 | 80.2192116 | -2.5036131 | 0.50875518 | -4.9210567 | 8.61E-07 |
| 227094 | NEMP2 | 130.535443 | -0.7107017 | 0.14443403 | -4.9205973 | 8.63E-07 |
| 269966 | NUP98 | 3577.72338 | 0.57783831 | 0.11743769 | 4.92038223 | 8.64E-07 |
| 208043 | SETD1B | 1654.84079 | 0.42742087 | 0.08688037 | 4.91964867 | 8.67E-07 |
| 14705 | BSCL2 | 510.880242 | -1.2256505 | 0.24916125 | -4.9191056 | 8.69E-07 |
| 57748 | JMY | 2059.58265 | 0.60011903 | 0.12205622 | 4.91674254 | 8.80E-07 |
| 74489 | 5430421F17I | 12.793852 | 4.65022413 | 0.94608432 | 4.91523224 | 8.87E-07 |
| 399568 | BC052040 | 382.300679 | -0.6685136 | 0.1360143 | -4.9150242 | 8.88E-07 |
| 68364 | 0610030E20I | 645.524575 | 0.43074793 | 0.08766476 | 4.9135816 | 8.94E-07 |
| 319190 | HIST2H2BE | 30.5144221 | -1.4738076 | 0.30000795 | -4.912562 | 8.99E-07 |
| 12393 | RUNX2 | 1060.0773 | 0.6745067 | 0.13731213 | 4.91221478 | 9.01E-07 |
| 625409 | Gm6581 | 169.804723 | -0.6938888 | 0.14128264 | -4.9113524 | 9.05E-07 |
| 269593 | LUZP1 | 3711.93013 | 0.75741491 | 0.15423156 | 4.91089457 | 9.07E-07 |
| 104015 | SYNJ1 | 1487.15452 | -0.6811039 | 0.13872585 | -4.9097111 | 9.12E-07 |
| 67182 | PDZK1IP1 | 7.52546067 | -6.3462429 | 1.29351395 | -4.9062037 | 9.29E-07 |
| 110821 | PCCA | 250.407572 | -1.2923158 | 0.26340807 | -4.9061361 | 9.29E-07 |
| 19765 | RALBP1 | 2790.91088 | -0.3234886 | 0.06597319 | -4.9033351 | 9.42E-07 |
| 224105 | PAK2 | 6765.84095 | -0.4688951 | 0.09563247 | -4.9030954 | 9.43E-07 |
| 217866 | CDC42BPB | 7900.80323 | 0.58660055 | 0.11965767 | 4.90232299 | 9.47E-07 |
| 243853 | FKRP | 1269.7966 | 0.68366372 | 0.13955073 | 4.89903368 | 9.63E-07 |
| 66870 | SERBP1 | 25140.1765 | 0.95999194 | 0.19597882 | 4.89844729 | 9.66E-07 |
| 59029 | PSMD14 | 4359.72092 | 0.29510297 | 0.06025811 | 4.89731524 | 9.72E-07 |
| 12638 | CFTR | 12.8158067 | -7.1071452 | 1.45148878 | -4.8964521 | 9.76E-07 |
| 68598 | DNAJC8 | 1677.74501 | 0.73826538 | 0.15080268 | 4.89557218 | 9.80E-07 |
| 192173 | fam195b | 697.748663 | 0.4197 | 0.08574413 | 4.89479576 | 9.84E-07 |
| 76483 | LMF1 | 410.075268 | 0.87111852 | 0.17804388 | 4.89271824 | 9.95E-07 |
| 66066 | GNG11 | 434.285437 | 1.79215492 | 0.36629828 | 4.89261083 | 9.95E-07 |
| 27389 | DUSP13 | 12.8636838 | -6.3814883 | 1.30445194 | -4.8920839 | 9.98E-07 |
| 76132 | FAXC | 1641.98794 | -0.9673165 | 0.1977852 | -4.8907426 | 1.00E-06 |
| 231868 | E130309D02 | 1178.42667 | 0.40791493 | 0.0834127 | 4.89032173 | 1.01E-06 |
| 12829 | COL4A4 | 9.40934214 | -6.661198 | 1.36247558 | -4.8890403 | 1.01E-06 |
| 20723 | SERPINB9 | 1214.40474 | -1.3160846 | 0.26924128 | -4.8881234 | 1.02E-06 |
| 71492 | BBS7 | 456.694952 | -0.6112946 | 0.12508513 | -4.8870281 | 1.02E-06 |
| 16511 | KCNH2 | 103.458425 | -5.8379815 | 1.19512205 | -4.8848412 | 1.04E-06 |

| | | | | | | |
|---|---|---|---|---|---|---|
| 98711 | RDH10 | 1356.9251 | -1.1476521 | 0.23494193 | -4.884833 | 1.04E-06 |
| 73827 | TMEM198B | 430.611579 | 0.84978031 | 0.17398244 | 4.88428777 | 1.04E-06 |
| 14198 | FHIT | 6.97093828 | -6.2297006 | 1.27564964 | -4.8835514 | 1.04E-06 |
| 100042547 | Gm3897 | 18.6571818 | -1.9866757 | 0.40692128 | -4.8822114 | 1.05E-06 |
| 50785 | HS6ST1 | 942.836578 | -0.5680522 | 0.1163539 | -4.8821073 | 1.05E-06 |
| 66204 | ACYP1 | 108.563484 | 0.85149744 | 0.17445597 | 4.88087315 | 1.06E-06 |
| 217378 | DNAJC27 | 371.741603 | 1.03173816 | 0.21138832 | 4.88077193 | 1.06E-06 |
| 17775 | laptm4a | 11966.9894 | 0.4186114 | 0.0857709 | 4.88057626 | 1.06E-06 |
| 69185 | DTWD1 | 497.68602 | -0.7830604 | 0.16045925 | -4.8801202 | 1.06E-06 |
| 13480 | DPM1 | 662.191341 | -0.3039307 | 0.06228155 | -4.8799473 | 1.06E-06 |
| 320718 | SLC26A9 | 7.47507072 | -6.3353526 | 1.29864868 | -4.8784192 | 1.07E-06 |
| 216169 | ABHD17A | 2047.82208 | -0.9225114 | 0.18911254 | -4.8781079 | 1.07E-06 |
| 226519 | LAMC1 | 10845.4588 | 0.65684457 | 0.13468629 | 4.87684804 | 1.08E-06 |
| 102693 | PHLDB1 | 6553.00765 | 1.24254134 | 0.25480278 | 4.87648256 | 1.08E-06 |
| 268739 | ARHGEF40 | 3170.4578 | 0.99204808 | 0.20344621 | 4.87621808 | 1.08E-06 |
| 17357 | MARCKSL1 | 1819.98066 | -0.7287883 | 0.14949841 | -4.8748901 | 1.09E-06 |
| 58522 | TRIM54 | 151.474484 | -4.6906745 | 0.9623333 | -4.8742722 | 1.09E-06 |
| 14675 | GNA14 | 29.7876332 | -2.1494726 | 0.44104337 | -4.8736082 | 1.10E-06 |
| 76872 | CCDC116 | 24.3733909 | 2.58219901 | 0.52984172 | 4.873529 | 1.10E-06 |
| 18295 | OGN | 69.5056056 | 4.52216145 | 0.92807657 | 4.87261678 | 1.10E-06 |
| 66266 | EAPP | 794.336139 | 0.32363473 | 0.06642718 | 4.87202279 | 1.10E-06 |
| 69151 | LZIC | 986.11089 | 0.32956339 | 0.06764804 | 4.87173579 | 1.11E-06 |
| 69169 | FCMR | 14.0828582 | -5.1542244 | 1.05799521 | -4.8716897 | 1.11E-06 |
| 100217466 | Scarna17 | 50.522544 | -1.3682845 | 0.28096439 | -4.8699571 | 1.12E-06 |
| 52570 | CCDC69 | 24.5167277 | -2.0600934 | 0.42304086 | -4.8697267 | 1.12E-06 |
| 66585 | SNRNP40 | 1450.75896 | 1.0176561 | 0.20898651 | 4.86948232 | 1.12E-06 |
| 434233 | Gm5601 | 337.785643 | 0.49475425 | 0.10161847 | 4.86874338 | 1.12E-06 |
| 68837 | FOXK2 | 5075.12291 | 0.47322492 | 0.09721077 | 4.86802991 | 1.13E-06 |
| 103806 | MAML1 | 2303.83763 | 0.67720241 | 0.13914003 | 4.86705655 | 1.13E-06 |
| 20823 | SSB | 5884.43162 | 0.37284606 | 0.07662422 | 4.86590337 | 1.14E-06 |
| 16573 | KIF5B | 10063.843 | -0.2033189 | 0.04179176 | -4.8650477 | 1.14E-06 |
| 93760 | ARID1A | 4204.20483 | 0.85599916 | 0.17596013 | 4.86473375 | 1.15E-06 |
| 81500 | SIL1 | 750.980347 | -0.6457431 | 0.13277381 | -4.8634826 | 1.15E-06 |
| 76719 | KANSL1 | 2210.64997 | -0.3742467 | 0.0769558 | -4.8631388 | 1.16E-06 |
| 56752 | ALDH9A1 | 1422.8879 | -0.5160368 | 0.10613126 | -4.8622504 | 1.16E-06 |
| 73804 | KIF2C | 2146.44558 | 1.01959585 | 0.20976944 | 4.86055486 | 1.17E-06 |
| 66354 | SNW1 | 2720.11529 | 0.47315671 | 0.09734953 | 4.86039051 | 1.17E-06 |
| 74355 | SMCHD1 | 4400.52313 | 0.24737871 | 0.05089954 | 4.86013611 | 1.17E-06 |
| 622320 | KCTD21 | 54.5906906 | -1.4387331 | 0.29608733 | -4.8591512 | 1.18E-06 |
| 12267 | C3AR1 | 57.6594004 | -8.5558553 | 1.76106855 | -4.8583318 | 1.18E-06 |
| 224020 | PI4KA | 3786.89714 | 0.4658602 | 0.09594876 | 4.85530175 | 1.20E-06 |
| 67603 | DUSP6 | 3172.94788 | 0.93714273 | 0.19303027 | 4.8549004 | 1.20E-06 |
| 66942 | DDX18 | 761.717861 | 0.4395382 | 0.09053623 | 4.85483242 | 1.20E-06 |
| 101118 | TMEM168 | 1885.81212 | 0.36316152 | 0.07484804 | 4.85198453 | 1.22E-06 |
| 100504127 | Gm17690 | 39.975748 | -1.4230838 | 0.29334849 | -4.8511714 | 1.23E-06 |

| | | | | | | |
|---|---|---|---|---|---|---|
| 195522 | Zfp691 | 99.7597469 | 0.84219298 | 0.17361686 | 4.85087091 | 1.23E-06 |
| 245522 | ZC4H2 | 157.527477 | -1.3520864 | 0.27879359 | -4.8497756 | 1.24E-06 |
| 18220 | NUCB1 | 4269.78052 | -0.3380849 | 0.06973102 | -4.8484145 | 1.24E-06 |
| 243864 | Mill2 | 12.6933581 | -4.5846668 | 0.94561184 | -4.8483602 | 1.24E-06 |
| 70005 | Znf41-ps | 29.613045 | -3.5367445 | 0.73008429 | -4.8442962 | 1.27E-06 |
| 232431 | GPRC5A | 2855.87174 | -0.4641594 | 0.09585376 | -4.8423702 | 1.28E-06 |
| 210789 | TBC1D4 | 873.516125 | 0.51942112 | 0.10728414 | 4.8415461 | 1.29E-06 |
| 16576 | KIF7 | 294.328757 | 0.87106113 | 0.17996245 | 4.84023817 | 1.30E-06 |
| 246738 | DNAJC28 | 36.7847658 | -1.4026741 | 0.2898316 | -4.8396177 | 1.30E-06 |
| 101502 | HSD3B7 | 1064.98737 | -0.5861322 | 0.12114853 | -4.8381288 | 1.31E-06 |
| 217827 | NRDE2 | 880.627993 | 0.73025953 | 0.15097648 | 4.83690922 | 1.32E-06 |
| 383619 | AIM2 | 13.0404287 | -5.6541503 | 1.16914529 | -4.8361401 | 1.32E-06 |
| 83490 | pik3ap1 | 25.2331512 | -8.0862253 | 1.67209698 | -4.8359786 | 1.32E-06 |
| 54411 | ATP6AP1 | 7475.64891 | -0.5449453 | 0.11268905 | -4.8358313 | 1.33E-06 |
| 66441 | MAGOHB | 313.707287 | 0.91947375 | 0.19016703 | 4.8350849 | 1.33E-06 |
| 232339 | ANKRD26 | 726.352032 | 0.52701641 | 0.10904998 | 4.83279686 | 1.35E-06 |
| 71989 | RPUSD4 | 432.256973 | 0.49340357 | 0.10212516 | 4.83136158 | 1.36E-06 |
| 94045 | P2RX5 | 88.9523713 | 1.48710193 | 0.30781395 | 4.83117127 | 1.36E-06 |
| 27681 | SNF8 | 767.700786 | -0.3896979 | 0.08067591 | -4.8304131 | 1.36E-06 |
| 69956 | PTCD3 | 2511.99664 | 0.5804806 | 0.12019428 | 4.82951934 | 1.37E-06 |
| 23882 | GADD45G | 1399.81127 | -1.2095945 | 0.25047244 | -4.829252 | 1.37E-06 |
| 72578 | 2700054A10 | 42.3827174 | -2.0676359 | 0.42817261 | -4.8289775 | 1.37E-06 |
| 93739 | GABARAPL2 | 123.756691 | -0.7734445 | 0.16020312 | -4.8278989 | 1.38E-06 |
| 20624 | EFTUD2 | 7525.28753 | 0.17495203 | 0.03624559 | 4.82685048 | 1.39E-06 |
| 12568 | CDK5 | 745.075717 | -0.4212646 | 0.08727837 | -4.8266783 | 1.39E-06 |
| 384275 | Gm5297 | 686.697175 | 0.37724801 | 0.07816069 | 4.82656959 | 1.39E-06 |
| 73738 | HAUS7 | 746.334393 | 0.50630098 | 0.10490311 | 4.82636754 | 1.39E-06 |
| 71213 | CAGE1 | 51.1879834 | -1.2094614 | 0.25066829 | -4.8249478 | 1.40E-06 |
| 30960 | VAPA | 3963.44177 | 0.28186763 | 0.05843229 | 4.82383378 | 1.41E-06 |
| 55944 | EIF3D | 7454.52425 | -0.2660825 | 0.05518625 | -4.8215365 | 1.42E-06 |
| 100041655 | Gm10863 | 6.76839033 | -6.1834839 | 1.28247324 | -4.8215306 | 1.42E-06 |
| 209478 | TBC1D12 | 518.040428 | -0.4808794 | 0.09974824 | -4.8209306 | 1.43E-06 |
| 74094 | TJAP1 | 2447.24335 | 0.47560001 | 0.0986588 | 4.82065445 | 1.43E-06 |
| 15519 | HSP90AA1 | 6376.02996 | 0.47895865 | 0.09937161 | 4.81987406 | 1.44E-06 |
| 70423 | tspan15 | 327.00464 | -2.7766275 | 0.57621477 | -4.8187371 | 1.44E-06 |
| 55946 | AP3M1 | 1397.51523 | -0.5318414 | 0.11037984 | -4.8182833 | 1.45E-06 |
| 20810 | SRM | 3406.41746 | 1.18977061 | 0.24697482 | 4.81737627 | 1.45E-06 |
| 52710 | SLC52A2 | 687.066119 | 0.78886195 | 0.16379395 | 4.81618493 | 1.46E-06 |
| 228790 | ASXL1 | 3194.40348 | 0.33196337 | 0.06894606 | 4.81482718 | 1.47E-06 |
| 239546 | Zfp647 | 80.0042234 | 0.89300598 | 0.18547355 | 4.81473481 | 1.47E-06 |
| 101831 | FAAP24 | 673.870129 | 1.1254001 | 0.23376476 | 4.81424184 | 1.48E-06 |
| 109284 | R3HDM4 | 2173.30823 | -0.3806105 | 0.07906527 | -4.8138778 | 1.48E-06 |
| 57434 | XRCC2 | 588.18642 | 0.33791257 | 0.07021304 | 4.81267533 | 1.49E-06 |
| 80509 | MED8 | 976.965957 | 0.57155905 | 0.11878 | 4.81191313 | 1.49E-06 |
| 16319 | INCENP | 4692.0381 | 0.31407363 | 0.06527167 | 4.81179122 | 1.50E-06 |

| | | | | | | |
|---|---|---|---|---|---|---|
| 56469 | PIAS1 | 951.13063 | 0.35457632 | 0.07373203 | 4.80898651 | 1.52E-06 |
| 228998 | ARFGAP1 | 1713.47338 | 0.49790464 | 0.10355841 | 4.8079596 | 1.52E-06 |
| 83924 | GPR137B | 1853.63068 | -1.0317964 | 0.21461891 | -4.8075746 | 1.53E-06 |
| 236792 | MMGT1 | 2574.66906 | 0.33970583 | 0.07066059 | 4.80757162 | 1.53E-06 |
| 75030 | 4930503E24I | 123.948065 | 1.04250465 | 0.21686984 | 4.80705235 | 1.53E-06 |
| 20768 | SEPHS2 | 2284.5633 | 0.22205459 | 0.04620907 | 4.80543339 | 1.54E-06 |
| 94275 | MAGED1 | 7865.932 | 0.55424971 | 0.11537303 | 4.80397996 | 1.56E-06 |
| 373864 | COL27A1 | 170.332273 | -3.1749469 | 0.66111538 | -4.8024097 | 1.57E-06 |
| 20650 | SNTB2 | 1853.46264 | 0.38619048 | 0.08041657 | 4.80237412 | 1.57E-06 |
| 109263 | RLF | 1348.24641 | 0.48536645 | 0.10106948 | 4.80230501 | 1.57E-06 |
| 77116 | MTMR2 | 2775.26065 | 0.3048456 | 0.06348322 | 4.80198687 | 1.57E-06 |
| 105428 | Fam149b | 661.636449 | -0.3530015 | 0.07353208 | -4.8006469 | 1.58E-06 |
| 22779 | IKZF2 | 172.51341 | -1.9602669 | 0.40834736 | -4.8004888 | 1.58E-06 |
| 26397 | MAP2K3 | 7183.25392 | 0.68061591 | 0.14182077 | 4.79912718 | 1.59E-06 |
| 28075 | DESI1 | 2928.3196 | -0.4218383 | 0.08790488 | -4.7988046 | 1.60E-06 |
| 66495 | NDUFB3 | 418.015004 | -0.4369612 | 0.09106088 | -4.7985616 | 1.60E-06 |
| 212139 | cc2d1a | 1348.13983 | -0.3554018 | 0.07408015 | -4.7975309 | 1.61E-06 |
| 99382 | ABTB2 | 853.46959 | -0.8104069 | 0.16895494 | -4.7965862 | 1.61E-06 |
| 69408 | DNAJC17 | 120.370015 | 0.95815354 | 0.19975828 | 4.79656482 | 1.61E-06 |
| 12804 | CNTFR | 9.19310146 | -5.8913794 | 1.22863447 | -4.7950628 | 1.63E-06 |
| 18703 | PIGR | 8.81637959 | -6.5743413 | 1.37136627 | -4.7940083 | 1.63E-06 |
| 12055 | BCL7C | 525.230913 | 1.01623378 | 0.21203116 | 4.792851 | 1.64E-06 |
| 233107 | KCTD15 | 499.910853 | 1.81115986 | 0.3778957 | 4.79275069 | 1.65E-06 |
| 75221 | DPP3 | 2593.33191 | -0.5793598 | 0.12090405 | -4.7918973 | 1.65E-06 |
| 53319 | NXF1 | 2858.05232 | 0.20860364 | 0.04354234 | 4.79082292 | 1.66E-06 |
| 22668 | SF1 | 7886.67782 | 0.23112163 | 0.04824375 | 4.79070642 | 1.66E-06 |
| 11548 | ADRA1B | 174.546723 | -2.2251248 | 0.46463709 | -4.7889521 | 1.68E-06 |
| 16010 | IGFBP4 | 18692.2951 | 1.48643038 | 0.31044772 | 4.78802159 | 1.68E-06 |
| 26398 | MAP2K4 | 2402.71298 | -0.3068152 | 0.06408948 | -4.787294 | 1.69E-06 |
| 54006 | DEAF1 | 687.973425 | 0.54109733 | 0.11303662 | 4.78692077 | 1.69E-06 |
| 231630 | FICD | 190.181493 | -0.8185213 | 0.17101482 | -4.7862593 | 1.70E-06 |
| 16599 | KLF3 | 1750.89078 | -0.3653476 | 0.07633518 | -4.7860973 | 1.70E-06 |
| 319601 | Zfp653 | 255.945437 | 0.7211657 | 0.1507114 | 4.78507744 | 1.71E-06 |
| 17189 | MB | 13.1347737 | -7.1431793 | 1.49307522 | -4.7842059 | 1.72E-06 |
| 13006 | SMC3 | 4229.30725 | 0.30801227 | 0.06438206 | 4.78413164 | 1.72E-06 |
| 56050 | CYP39A1 | 95.4747268 | -2.9903423 | 0.62527895 | -4.7824133 | 1.73E-06 |
| 106529 | TECR | 2986.7293 | -0.5063105 | 0.10589486 | -4.7812563 | 1.74E-06 |
| 224792 | ADGRF5 | 12.4012296 | -2.8287637 | 0.59168075 | -4.7808952 | 1.75E-06 |
| 83554 | FSTL3 | 466.516524 | -1.5985721 | 0.33438216 | -4.7806739 | 1.75E-06 |
| 212153 | CCDC191 | 142.715702 | -0.9817586 | 0.20537407 | -4.7803433 | 1.75E-06 |
| 233066 | SYNE4 | 23.6529101 | -4.9691335 | 1.03967141 | -4.7795231 | 1.76E-06 |
| 233033 | SAMD4B | 5464.09503 | 0.3523682 | 0.07376131 | 4.77714108 | 1.78E-06 |
| 74782 | GLT8D2 | 69.1517676 | 1.55233487 | 0.32496108 | 4.77698704 | 1.78E-06 |
| 230866 | EMC1 | 2329.9881 | 0.46269986 | 0.09686644 | 4.77667866 | 1.78E-06 |
| 17701 | MSX1 | 55.4910279 | 1.83875586 | 0.38510254 | 4.7747176 | 1.80E-06 |

| | | | | | | |
|---|---|---|---|---|---|---|
| 59054 | MRPS30 | 1259.48753 | 0.80688603 | 0.16916901 | 4.76970367 | 1.84E-06 |
| 381270 | 41701 | 10.6016721 | 5.40604742 | 1.13372561 | 4.76839138 | 1.86E-06 |
| 20821 | TRIM21 | 164.840488 | -1.296481 | 0.27189643 | -4.7682898 | 1.86E-06 |
| 52430 | ECHDC2 | 270.393806 | -0.5229997 | 0.10968992 | -4.7679836 | 1.86E-06 |
| 19165 | PSEN2 | 165.413582 | -1.2687111 | 0.26609926 | -4.7678114 | 1.86E-06 |
| 18510 | PAX8 | 7.87756355 | -6.4024982 | 1.34288712 | -4.7677114 | 1.86E-06 |
| 77805 | ESCO1 | 738.659294 | -0.4532392 | 0.09507323 | -4.7672637 | 1.87E-06 |
| 234258 | NEIL3 | 537.340813 | -0.499386 | 0.10476565 | -4.7666964 | 1.87E-06 |
| 73368 | COL20A1 | 66.2857986 | -2.8843568 | 0.60514726 | -4.7663718 | 1.88E-06 |
| 170758 | RAC3 | 70.3890371 | -1.0076755 | 0.21144857 | -4.7655821 | 1.88E-06 |
| 110854 | PTPA | 6308.80674 | 0.39844479 | 0.08360939 | 4.765551 | 1.88E-06 |
| 227485 | CDH19 | 16.3216594 | -5.3881983 | 1.13068331 | -4.7654354 | 1.88E-06 |
| 72775 | FANCE | 683.968407 | 1.27552944 | 0.26774223 | 4.76402035 | 1.90E-06 |
| 67937 | TMEM59L | 9.6411464 | 5.27641608 | 1.10778756 | 4.7630216 | 1.91E-06 |
| 227624 | RABL6 | 2504.26294 | 0.5637246 | 0.1183875 | 4.7616903 | 1.92E-06 |
| 16560 | KIF1A | 13.3411325 | -3.8175397 | 0.80186575 | -4.7608214 | 1.93E-06 |
| 228730 | KIZ | 251.976434 | -0.8164818 | 0.17156595 | -4.7589968 | 1.95E-06 |
| 19934 | RPL22 | 3366.72142 | 0.65871579 | 0.13846387 | 4.75731169 | 1.96E-06 |
| 320873 | CDH10 | 24.4160121 | -7.3093317 | 1.53708587 | -4.7553177 | 1.98E-06 |
| 214345 | LRRC1 | 489.478478 | -0.616038 | 0.12956149 | -4.754792 | 1.99E-06 |
| 102954 | NUDT10 | 16.165043 | -7.4466473 | 1.56661404 | -4.7533388 | 2.00E-06 |
| 56360 | ACOT9 | 1659.82391 | -0.3941352 | 0.08298598 | -4.7494195 | 2.04E-06 |
| 16828 | LDHA | 18882.7784 | 0.45745515 | 0.09632591 | 4.74903511 | 2.04E-06 |
| 68955 | SRRM4 | 138.931974 | 3.37325145 | 0.71045091 | 4.74804296 | 2.05E-06 |
| 71340 | RIOK1 | 882.339707 | 0.6457256 | 0.1360228 | 4.7471866 | 2.06E-06 |
| 74157 | CMTR1 | 3096.18261 | -0.3358867 | 0.07075543 | -4.7471501 | 2.06E-06 |
| 52838 | DNLZ | 956.472501 | 0.30603195 | 0.06447061 | 4.74684424 | 2.07E-06 |
| 108154 | ADAMTS6 | 83.4858069 | -1.4603396 | 0.30765773 | -4.746637 | 2.07E-06 |
| 276829 | SMTNL2 | 29.6228654 | -3.5152332 | 0.74080204 | -4.7451722 | 2.08E-06 |
| 226970 | ARHGEF4 | 21.5349639 | -4.0060133 | 0.84437612 | -4.7443469 | 2.09E-06 |
| 215512 | FAM117A | 36.3525319 | -2.9488524 | 0.62156492 | -4.7442388 | 2.09E-06 |
| 67991 | NACC2 | 1312.6435 | -0.94929 | 0.20012282 | -4.7435372 | 2.10E-06 |
| 98732 | RAB3GAP2 | 3510.37064 | 0.21437732 | 0.04519789 | 4.74308208 | 2.10E-06 |
| 66343 | TMEM177 | 238.040837 | -0.5762963 | 0.12152193 | -4.7423238 | 2.11E-06 |
| 26396 | MAP2K2 | 4414.67456 | -0.4738404 | 0.09997553 | -4.7395639 | 2.14E-06 |
| 270162 | ELMOD1 | 23.922157 | -8.0114076 | 1.6905402 | -4.7389631 | 2.15E-06 |
| 278097 | ARMCX6 | 110.862615 | 1.0389999 | 0.21928453 | 4.73813598 | 2.16E-06 |
| 546387 | Gm5946 | 129.589242 | 0.63108843 | 0.13321149 | 4.73749251 | 2.16E-06 |
| 243529 | H1FX | 315.557801 | 2.82816272 | 0.5973389 | 4.73460333 | 2.19E-06 |
| 216724 | RUFY1 | 754.913537 | -0.6835966 | 0.14438677 | -4.7344823 | 2.20E-06 |
| 70381 | TECPR1 | 1334.83952 | -2.0303841 | 0.42892808 | -4.7336236 | 2.21E-06 |
| 73826 | POLDIP3 | 4200.81889 | 0.2593773 | 0.05479866 | 4.73327795 | 2.21E-06 |
| 432964 | K230010J24F | 12.1631488 | -2.8114827 | 0.59402451 | -4.7329406 | 2.21E-06 |
| 105351 | AW209491 | 465.121152 | -0.6561637 | 0.13867333 | -4.7317224 | 2.23E-06 |
| 70567 | FRA10AC1 | 301.192249 | -0.551254 | 0.11651201 | -4.7313058 | 2.23E-06 |

| | | | | | | |
|---|---|---|---|---|---|---|
| 170716 | Cyp4f13 | 184.023958 | -1.9787943 | 0.41830111 | -4.73055 | 2.24E-06 |
| 229949 | ak5 | 17.6012569 | 2.01486871 | 0.42592879 | 4.73052945 | 2.24E-06 |
| 17967 | NCAM1 | 1803.10729 | -4.0152445 | 0.84917856 | -4.7283866 | 2.26E-06 |
| 233876 | hirip3 | 1459.59022 | 0.99764182 | 0.21105139 | 4.72700905 | 2.28E-06 |
| 20510 | SLC1A1 | 7.48935908 | -6.3313834 | 1.33987969 | -4.7253372 | 2.30E-06 |
| 71801 | PLEKHF2 | 605.437869 | -1.0368506 | 0.21944938 | -4.7247824 | 2.30E-06 |
| 69097 | TRIM15 | 8.8608656 | -5.8398853 | 1.23602825 | -4.7247183 | 2.30E-06 |
| 19891 | RPA2 | 1148.95051 | 0.85813284 | 0.18163868 | 4.72439475 | 2.31E-06 |
| 100861774 | #N/A | 1239.20792 | 0.4227893 | 0.08952033 | 4.72283007 | 2.33E-06 |
| 270198 | PFKFB4 | 178.187706 | -1.734605 | 0.36734198 | -4.722044 | 2.33E-06 |
| 236193 | Zfp709 | 134.254758 | -1.0964894 | 0.23221349 | -4.7219022 | 2.34E-06 |
| 54519 | APBB1IP | 1897.25083 | 2.94864842 | 0.62447132 | 4.72183159 | 2.34E-06 |
| 330149 | HFM1 | 6.85787938 | -5.4591669 | 1.15645272 | -4.720614 | 2.35E-06 |
| 67590 | TCTN3 | 166.575736 | -0.8235656 | 0.17446374 | -4.7205543 | 2.35E-06 |
| 21769 | ZFAND3 | 2450.95039 | -0.4498733 | 0.09530863 | -4.7201742 | 2.36E-06 |
| 14260 | FMN1 | 2543.953 | 0.95937778 | 0.20325158 | 4.72014906 | 2.36E-06 |
| 214804 | SYDE2 | 536.972144 | 0.98433863 | 0.20856105 | 4.71966669 | 2.36E-06 |
| 79554 | CPTP | 807.099081 | 0.46258616 | 0.09805801 | 4.71747436 | 2.39E-06 |
| 225994 | NMRK1 | 78.6448476 | -1.0472815 | 0.22213877 | -4.7145371 | 2.42E-06 |
| 71086 | 4933412E12I | 109.968554 | -1.3757958 | 0.29185538 | -4.7139642 | 2.43E-06 |
| 29806 | LIMD1 | 2824.63102 | 0.24277081 | 0.05150228 | 4.71378763 | 2.43E-06 |
| 56294 | PTPN9 | 2598.94343 | 0.37674786 | 0.07992934 | 4.71351147 | 2.43E-06 |
| 71713 | CDC40 | 1440.12899 | 0.39857995 | 0.08456533 | 4.71327865 | 2.44E-06 |
| 56722 | LITAF | 5386.97941 | -1.0037394 | 0.21300281 | -4.7123292 | 2.45E-06 |
| 54003 | NELL2 | 14.5153599 | -5.229577 | 1.10985487 | -4.7119467 | 2.45E-06 |
| 12314 | CALM2 | 5876.98823 | -0.6035763 | 0.12814229 | -4.7102038 | 2.47E-06 |
| 331461 | IL1RAPL1 | 9.14804814 | 5.18520499 | 1.10120802 | 4.70865168 | 2.49E-06 |
| 233904 | SETD1A | 3298.41252 | 0.59630833 | 0.12665133 | 4.70826739 | 2.50E-06 |
| 67776 | VWA5A | 748.315832 | -1.0950724 | 0.23258552 | -4.7082571 | 2.50E-06 |
| 83922 | CEP41 | 206.071059 | -1.1928675 | 0.25343662 | -4.7067683 | 2.52E-06 |
| 74430 | 4930452B06 | 53.9907559 | -3.8806336 | 0.82475284 | -4.705208 | 2.54E-06 |
| 240518 | PELI3 | 31.9646686 | -2.537591 | 0.53936885 | -4.7047415 | 2.54E-06 |
| 328365 | ZMIZ1 | 7400.36217 | -0.4506575 | 0.09580615 | -4.703847 | 2.55E-06 |
| 54138 | ATXN10 | 4960.28106 | -0.5714061 | 0.12147659 | -4.7038368 | 2.55E-06 |
| 319263 | PCMTD1 | 966.30746 | -0.5786908 | 0.12307189 | -4.7020548 | 2.58E-06 |
| 67238 | FAM220A | 651.400531 | 0.37260541 | 0.07928419 | 4.69961783 | 2.61E-06 |
| 69681 | Cdk3-ps | 16.6220703 | -2.7238728 | 0.57972049 | -4.6985968 | 2.62E-06 |
| 22710 | Zfp52 | 1060.14944 | 0.69447181 | 0.14782078 | 4.69806614 | 2.63E-06 |
| 21414 | TCF7 | 513.85488 | 1.85919192 | 0.39575751 | 4.69780574 | 2.63E-06 |
| 320438 | ALG6 | 367.884186 | 0.62223538 | 0.13247678 | 4.69693907 | 2.64E-06 |
| 14431 | GAMT | 86.4967884 | 1.25003482 | 0.26615044 | 4.69672278 | 2.64E-06 |
| 52837 | TMX4 | 1107.51005 | -0.643788 | 0.13710319 | -4.6956456 | 2.66E-06 |
| 15926 | idh1 | 3529.29981 | 0.41027457 | 0.08737404 | 4.69561201 | 2.66E-06 |
| 66320 | TMEM208 | 656.129594 | -0.4177389 | 0.08900014 | -4.6936883 | 2.68E-06 |
| 100039239 | Gm2115 | 448.945728 | 2.92611276 | 0.62376585 | 4.69104354 | 2.72E-06 |

| | | | | | | |
|---|---|---|---|---|---|---|
| 170787 | HDAC10 | 405.284657 | 0.50329933 | 0.10735799 | 4.68804727 | 2.76E-06 |
| 69940 | EXOC1 | 1852.79537 | -0.2971418 | 0.0633862 | -4.6877991 | 2.76E-06 |
| 242362 | MANEA | 1222.4617 | -1.3410615 | 0.28619334 | -4.6858586 | 2.79E-06 |
| 67387 | UNC50 | 953.047137 | -0.3347928 | 0.07146561 | -4.6846699 | 2.80E-06 |
| 227622 | BC029214 | 115.977056 | -0.8256836 | 0.17627036 | -4.6841884 | 2.81E-06 |
| 67116 | CUEDC2 | 1127.47477 | -1.1715747 | 0.25011727 | -4.6841015 | 2.81E-06 |
| 12799 | CNP | 362.459102 | -0.4763683 | 0.10177077 | -4.6807965 | 2.86E-06 |
| 229699 | SLC16A4 | 10.8526008 | -5.4011705 | 1.15390401 | -4.6807797 | 2.86E-06 |
| 218543 | SREK1 | 1742.87893 | 0.56597994 | 0.12095497 | 4.67926161 | 2.88E-06 |
| 22758 | ZSCAN12 | 269.598588 | 0.60110283 | 0.1285063 | 4.67761385 | 2.90E-06 |
| 228880 | ZMYND8 | 1059.54387 | 1.16750295 | 0.24961209 | 4.67726917 | 2.91E-06 |
| 100502853 | Gm14002 | 16.8692127 | 1.95083041 | 0.41711407 | 4.67697095 | 2.91E-06 |
| 71785 | PDGFD | 44.9640255 | -4.4746371 | 0.95680172 | -4.6766607 | 2.92E-06 |
| 71920 | epgn | 47.8803885 | -2.0752282 | 0.4438033 | -4.676009 | 2.93E-06 |
| 171530 | ucn2 | 29.7918235 | -2.7031278 | 0.57825505 | -4.674629 | 2.94E-06 |
| 100502772 | #N/A | 103.137213 | 0.73060214 | 0.15630128 | 4.67431962 | 2.95E-06 |
| 20964 | SYN1 | 78.5166477 | -2.6576253 | 0.56858791 | -4.6740799 | 2.95E-06 |
| 231997 | FKBP14 | 319.386601 | 0.69602498 | 0.1489213 | 4.67377729 | 2.96E-06 |
| 67268 | MYL12A | 8813.71616 | -0.4651119 | 0.09951752 | -4.6736678 | 2.96E-06 |
| 72113 | ADCK1 | 424.068161 | 0.53587333 | 0.11467551 | 4.67295344 | 2.97E-06 |
| 13179 | DCN | 15.2467 | 4.37793286 | 0.93689876 | 4.67279184 | 2.97E-06 |
| 71664 | METTL7B | 41.7587613 | -1.9168801 | 0.41038862 | -4.67089 | 3.00E-06 |
| 170770 | BBC3 | 260.693546 | -0.672649 | 0.14403653 | -4.6699891 | 3.01E-06 |
| 72136 | CHST14 | 714.112233 | 0.45224406 | 0.09684851 | 4.66960267 | 3.02E-06 |
| 69276 | SEC62 | 3821.84564 | -0.3086242 | 0.06615386 | -4.6652483 | 3.08E-06 |
| 667764 | Gm15854 | 75.3222277 | -1.8017305 | 0.3862085 | -4.6651755 | 3.08E-06 |
| 66556 | DRAP1 | 2457.13081 | -0.234647 | 0.05031117 | -4.663915 | 3.10E-06 |
| 666048 | TRABD2B | 328.866508 | -5.2705349 | 1.13018905 | -4.66341 | 3.11E-06 |
| 14345 | FUT4 | 67.7344506 | 1.82030735 | 0.39035153 | 4.6632515 | 3.11E-06 |
| 12036 | BCAT2 | 422.579244 | 0.5660805 | 0.12139266 | 4.66321838 | 3.11E-06 |
| 18590 | PDGFA | 978.783631 | -0.4957005 | 0.10633846 | -4.6615355 | 3.14E-06 |
| 240328 | F830016B08 | 8.49966866 | -5.7720733 | 1.23831049 | -4.6612488 | 3.14E-06 |
| 79196 | osbpl5 | 1510.6516 | -0.4578583 | 0.09823726 | -4.6607401 | 3.15E-06 |
| 72899 | MACROD2 | 65.3279307 | 1.37282732 | 0.29457646 | 4.66034287 | 3.16E-06 |
| 13209 | DDX6 | 7820.19568 | 0.25202424 | 0.05408444 | 4.65982864 | 3.16E-06 |
| 56470 | RGS19 | 893.408224 | 1.06913163 | 0.22949542 | 4.65861867 | 3.18E-06 |
| 21982 | TMEM165 | 2439.58858 | -0.19504 | 0.04188396 | -4.6566745 | 3.21E-06 |
| 18844 | plxna1 | 9273.45065 | 0.65223641 | 0.14009912 | 4.65553538 | 3.23E-06 |
| 16589 | UHMK1 | 1008.57641 | -0.4068215 | 0.08739109 | -4.6551826 | 3.24E-06 |
| 15184 | HDAC5 | 2224.218 | 1.04176633 | 0.22380467 | 4.65480164 | 3.24E-06 |
| 74239 | IQCE | 575.044784 | -0.5342245 | 0.11478383 | -4.6541786 | 3.25E-06 |
| 207565 | CAMKK2 | 2188.89732 | 0.80108091 | 0.17212776 | 4.65399015 | 3.26E-06 |
| 56087 | DNAH10 | 9.75470018 | -3.1020752 | 0.66654256 | -4.6539792 | 3.26E-06 |
| 226539 | DARS2 | 529.219002 | -0.3543184 | 0.0761554 | -4.6525705 | 3.28E-06 |
| 20201 | S100A8 | 530.261899 | -3.5730322 | 0.76799742 | -4.6524013 | 3.28E-06 |

| | | | | | | |
|---|---|---|---|---|---|---|
| 241296 | LRRC8A | 2212.0046 | -0.5465261 | 0.11747191 | -4.6523983 | 3.28E-06 |
| 74325 | CLTB | 2401.7831 | -0.573339 | 0.12326814 | -4.6511534 | 3.30E-06 |
| 75761 | Apol7a | 72.8363788 | 1.53994794 | 0.33113757 | 4.65047782 | 3.31E-06 |
| 69256 | Zfp397 | 1221.64159 | -0.3312131 | 0.07122658 | -4.6501342 | 3.32E-06 |
| 269615 | PLCH2 | 17.1592413 | -2.9700571 | 0.63886675 | -4.6489462 | 3.34E-06 |
| 14747 | CMKLR1 | 37.0990941 | 4.19608007 | 0.90264873 | 4.64863009 | 3.34E-06 |
| 210544 | TBC1D31 | 1088.63201 | 0.62776619 | 0.13509337 | 4.64690609 | 3.37E-06 |
| 52666 | ARHGEF25 | 683.406492 | 1.00538095 | 0.21636727 | 4.64664065 | 3.37E-06 |
| 328580 | TUBGCP6 | 1850.05316 | -0.2467874 | 0.05313008 | -4.6449652 | 3.40E-06 |
| 246102 | RTTN | 543.232268 | 0.37323105 | 0.08036824 | 4.64401153 | 3.42E-06 |
| 18226 | NUP62 | 3478.78141 | 0.55537303 | 0.11963494 | 4.64223098 | 3.45E-06 |
| 29877 | hdgfrp3 | 1520.2587 | 4.89890529 | 1.05534794 | 4.64198121 | 3.45E-06 |
| 17190 | MBD1 | 1306.26731 | -0.7080725 | 0.15257393 | -4.6408487 | 3.47E-06 |
| 13107 | Cyp2f2 | 4.92720496 | -5.7266823 | 1.23397691 | -4.6408342 | 3.47E-06 |
| 16363 | IRF2 | 632.862723 | -0.3957437 | 0.08529329 | -4.6397994 | 3.49E-06 |
| 106757 | CATSPERD | 15.571582 | -3.221042 | 0.69429201 | -4.6393188 | 3.50E-06 |
| 100503793 | LOC1005037 | 30.9307889 | -1.3937513 | 0.30047592 | -4.6384793 | 3.51E-06 |
| 217946 | CDCA7L | 1510.27908 | 0.60090199 | 0.12956008 | 4.63801802 | 3.52E-06 |
| 66132 | 1110008L16 | 406.576182 | 0.39087077 | 0.08427897 | 4.6378209 | 3.52E-06 |
| 14701 | GNG12 | 5005.306 | 0.63611052 | 0.13716075 | 4.63770094 | 3.52E-06 |
| 68303 | FAM114A1 | 1258.5713 | 0.40937446 | 0.08828952 | 4.63672794 | 3.54E-06 |
| 102632 | ACAD11 | 354.189276 | -1.2171234 | 0.26250645 | -4.6365465 | 3.54E-06 |
| 74479 | SNX11 | 724.854231 | -0.3835442 | 0.08272652 | -4.6362913 | 3.55E-06 |
| 66153 | FBXO36 | 40.5479811 | -2.7137714 | 0.58534106 | -4.6362224 | 3.55E-06 |
| 59005 | TRAPPC2L | 498.254372 | -0.5003742 | 0.10793474 | -4.6358955 | 3.55E-06 |
| 240261 | CCDC112 | 149.033293 | -1.2918949 | 0.27871797 | -4.6351331 | 3.57E-06 |
| 231130 | TNIP2 | 264.383769 | -0.9829415 | 0.21219027 | -4.6323587 | 3.62E-06 |
| 52915 | ZMIZ2 | 2511.32481 | -0.323641 | 0.06987262 | -4.6318717 | 3.62E-06 |
| 69981 | TMEM30A | 4672.89972 | 0.28432745 | 0.06139444 | 4.63115951 | 3.64E-06 |
| 68127 | B230217C12 | 33.4150823 | 3.28075923 | 0.70894808 | 4.62764384 | 3.70E-06 |
| 77591 | DDX10 | 1288.85927 | 0.3825636 | 0.08274996 | 4.62312726 | 3.78E-06 |
| 244859 | ANKK1 | 19.4962564 | -3.0903849 | 0.6684989 | -4.6228721 | 3.78E-06 |
| 13871 | ERCC2 | 1002.53697 | 0.40815647 | 0.08829838 | 4.62246859 | 3.79E-06 |
| 212123 | DCAF15 | 1146.40214 | 0.37749604 | 0.08167847 | 4.62173233 | 3.81E-06 |
| 214048 | LARP1B | 383.118016 | -0.6637449 | 0.14361661 | -4.6216449 | 3.81E-06 |
| 76938 | RBM17 | 1963.96289 | 0.25599031 | 0.05543223 | 4.61807697 | 3.87E-06 |
| 239447 | COLEC10 | 9.360909 | 5.96997466 | 1.29324906 | 4.61626057 | 3.91E-06 |
| 93887 | PCDHB16 | 233.300798 | 2.3699399 | 0.51357672 | 4.61457817 | 3.94E-06 |
| 29813 | Zfp385a | 599.631738 | -1.4851326 | 0.32188555 | -4.6138531 | 3.95E-06 |
| 18813 | PA2G4 | 10239.3237 | 0.38462711 | 0.08338057 | 4.61291036 | 3.97E-06 |
| 56357 | IVD | 662.303772 | -0.6804793 | 0.14752896 | -4.6125132 | 3.98E-06 |
| 67198 | SPATS2L | 36.7415016 | -7.903059 | 1.71353542 | -4.6121364 | 3.99E-06 |
| 108911 | RCC2 | 12544.2779 | 0.96279208 | 0.20879035 | 4.6112862 | 4.00E-06 |
| 170791 | RBM39 | 5248.80655 | 0.17961331 | 0.03897973 | 4.60786422 | 4.07E-06 |
| 13367 | DIAPH1 | 3595.27835 | -0.2707947 | 0.05877579 | -4.6072479 | 4.08E-06 |

| | | | | | | |
|---|---|---|---|---|---|---|
| 268970 | ARHGAP28 | 641.468798 | 1.67801388 | 0.36422956 | 4.60702276 | 4.08E-06 |
| 12402 | CBL | 1327.13798 | 0.62344578 | 0.13535133 | 4.60612959 | 4.10E-06 |
| 56749 | DHODH | 555.072706 | 0.58571579 | 0.12716925 | 4.60579749 | 4.11E-06 |
| 211548 | nomo1 | 7138.04313 | 0.53755736 | 0.11673885 | 4.60478536 | 4.13E-06 |
| 381413 | GPR176 | 1039.00389 | 4.42456119 | 0.96091156 | 4.60454569 | 4.13E-06 |
| 226641 | ATF6 | 3279.05398 | -0.572449 | 0.12434078 | -4.6038717 | 4.15E-06 |
| 240660 | SLC35G1 | 402.664295 | -1.7059512 | 0.37057631 | -4.6035084 | 4.15E-06 |
| 15951 | Ifi204 | 593.166257 | -0.6543956 | 0.14216102 | -4.6031999 | 4.16E-06 |
| 217294 | BC006965 | 17.1902491 | -6.0525391 | 1.3151431 | -4.6021905 | 4.18E-06 |
| 66757 | ADAT2 | 399.462941 | -0.4272331 | 0.09283873 | -4.6018843 | 4.19E-06 |
| 381038 | PARL | 1085.78008 | -0.4887032 | 0.10620595 | -4.6014673 | 4.20E-06 |
| 100503104 | Gm19557 | 12.5505205 | 3.11725068 | 0.6775712 | 4.60062452 | 4.21E-06 |
| 81018 | RNF114 | 1173.57649 | -0.671953 | 0.14610414 | -4.5991374 | 4.24E-06 |
| 232078 | THNSL2 | 28.8384108 | -2.7620555 | 0.60066386 | -4.5983381 | 4.26E-06 |
| 319675 | CEP295 | 1498.71717 | 0.31451053 | 0.06841761 | 4.5969238 | 4.29E-06 |
| 69666 | PSMG4 | 299.502134 | 0.82572587 | 0.17967045 | 4.59578002 | 4.31E-06 |
| 76884 | cyfip2 | 242.330026 | -5.9544897 | 1.29607963 | -4.5942313 | 4.34E-06 |
| 17920 | MYO6 | 1518.85562 | 0.8403081 | 0.18294593 | 4.59320475 | 4.36E-06 |
| 257635 | SDSL | 21.5000279 | -3.3874199 | 0.7376678 | -4.592067 | 4.39E-06 |
| 320655 | PGAP3 | 237.027731 | -0.4509718 | 0.09823742 | -4.5906312 | 4.42E-06 |
| 241694 | RALGAPA2 | 583.372941 | -0.6246748 | 0.13608635 | -4.5902827 | 4.43E-06 |
| 207352 | SEC23IP | 3217.2547 | 0.64711389 | 0.14097561 | 4.59025438 | 4.43E-06 |
| 170741 | Pilrb1 | 7.02959653 | -6.2393315 | 1.35960339 | -4.5890821 | 4.45E-06 |
| 270192 | RAB6B | 33.0103811 | 2.26852699 | 0.49434227 | 4.58898042 | 4.45E-06 |
| 140571 | PLXNB3 | 4.63685801 | -5.64151 | 1.22974136 | -4.5875582 | 4.48E-06 |
| 330788 | Zfp866 | 428.253683 | -0.5547157 | 0.12093706 | -4.586813 | 4.50E-06 |
| 381605 | TBC1D2 | 1137.00262 | -2.117778 | 0.46172705 | -4.5866447 | 4.50E-06 |
| 66905 | PLIN3 | 2340.80958 | -1.0303051 | 0.22465114 | -4.5862447 | 4.51E-06 |
| 11938 | ATP2A2 | 17527.1704 | 0.28055191 | 0.06118111 | 4.58559704 | 4.53E-06 |
| 11352 | ABL2 | 8540.69134 | 0.55277884 | 0.1205601 | 4.58508925 | 4.54E-06 |
| 69538 | ANTXR1 | 1725.12781 | 3.77131227 | 0.82259952 | 4.58462735 | 4.55E-06 |
| 78283 | MAP7D2 | 18.6229024 | 2.83859135 | 0.61943189 | 4.58257216 | 4.59E-06 |
| 211007 | TRIM41 | 1142.02951 | -0.3527966 | 0.07700113 | -4.5817072 | 4.61E-06 |
| 14711 | GNMT | 12.3732647 | -4.2221855 | 0.92156 | -4.5815634 | 4.62E-06 |
| 52398 | 41892 | 11376.5109 | 0.70141386 | 0.153125 | 4.58066192 | 4.64E-06 |
| 12566 | CDK2 | 1605.09157 | 0.31638271 | 0.06909039 | 4.57925767 | 4.67E-06 |
| 20927 | ABCC8 | 7.24864457 | -6.2845437 | 1.37280572 | -4.5778828 | 4.70E-06 |
| 50883 | CHEK2 | 556.86196 | -0.6468871 | 0.14131521 | -4.5776186 | 4.70E-06 |
| 14050 | EYA3 | 1340.87719 | 0.60442711 | 0.13205541 | 4.577072 | 4.72E-06 |
| 109785 | PGM3 | 537.258821 | 0.41347465 | 0.09035251 | 4.5762387 | 4.73E-06 |
| 78798 | EML4 | 1205.77933 | 0.42319894 | 0.09247865 | 4.57617969 | 4.74E-06 |
| 72726 | TBCC | 506.423664 | 0.43405619 | 0.09488255 | 4.5746682 | 4.77E-06 |
| 19356 | RAD17 | 810.857474 | 0.51104494 | 0.11173458 | 4.57374007 | 4.79E-06 |
| 14181 | fgfbp1 | 123.594311 | -3.2244663 | 0.70504479 | -4.5734205 | 4.80E-06 |
| 214531 | TMPRSS13 | 6.37347216 | -6.1061206 | 1.33551759 | -4.5721005 | 4.83E-06 |

| | | | | | | |
|---|---|---|---|---|---|---|
| 103149 | UPB1 | 19.4483235 | -7.7129822 | 1.68726444 | -4.5712942 | 4.85E-06 |
| 71767 | TYSND1 | 834.605928 | 0.56550647 | 0.12371169 | 4.5711644 | 4.85E-06 |
| 74392 | SPECC1L | 2490.83406 | -0.2549528 | 0.05577537 | -4.571064 | 4.85E-06 |
| 93697 | ice2 | 419.890346 | 0.50878727 | 0.11131415 | 4.5707332 | 4.86E-06 |
| 20851 | STAT5B | 823.932064 | 0.32182435 | 0.07041159 | 4.57061595 | 4.86E-06 |
| 23859 | DLG2 | 18.5290953 | 2.33908093 | 0.51184223 | 4.56992565 | 4.88E-06 |
| 53867 | COL5A3 | 91.0582643 | 1.49729396 | 0.32770083 | 4.56908812 | 4.90E-06 |
| 100042773 | Tmed10-ps | 682.354128 | 0.48757042 | 0.10671674 | 4.56882793 | 4.90E-06 |
| 50927 | NASP | 3322.93783 | 1.10105413 | 0.2410809 | 4.56715621 | 4.94E-06 |
| 27214 | DBF4 | 3941.9425 | 0.30124993 | 0.06596428 | 4.5668644 | 4.95E-06 |
| 74998 | RAB11FIP2 | 628.109294 | -0.7054385 | 0.15446956 | -4.5668449 | 4.95E-06 |
| 244329 | mcph1 | 1590.98538 | 0.42791581 | 0.0937063 | 4.56656406 | 4.96E-06 |
| 230484 | USP1 | 2954.30488 | 0.75096197 | 0.16448838 | 4.56544095 | 4.98E-06 |
| 238323 | RPS6KL1 | 8.57740748 | -6.5343328 | 1.43143801 | -4.5648731 | 5.00E-06 |
| 19646 | RBBP4 | 6966.21565 | 0.67262556 | 0.14736421 | 4.5643753 | 5.01E-06 |
| 108960 | IRAK2 | 287.511513 | -0.616109 | 0.13499157 | -4.5640555 | 5.02E-06 |
| 78308 | GPR108 | 1250.06766 | -0.4640522 | 0.10168362 | -4.5636871 | 5.03E-06 |
| 70767 | PRPF3 | 1301.38557 | 0.24778036 | 0.05430351 | 4.56287896 | 5.05E-06 |
| 68493 | NDUFAF4 | 1111.25312 | -0.8585485 | 0.18819103 | -4.562112 | 5.06E-06 |
| 331487 | UPRT | 343.409113 | -0.5937102 | 0.13018026 | -4.5606777 | 5.10E-06 |
| 381823 | APOLD1 | 51.0264544 | 3.21642733 | 0.70542723 | 4.55954522 | 5.13E-06 |
| 231329 | POLR2B | 4434.61273 | 0.27215444 | 0.05968938 | 4.55951163 | 5.13E-06 |
| 215693 | ZMAT1 | 45.5537618 | -1.8480485 | 0.40568096 | -4.5554232 | 5.23E-06 |
| 17354 | MLLT10 | 1228.88912 | 0.25435901 | 0.0558492 | 4.55438963 | 5.25E-06 |
| 68401 | G6PC3 | 903.642446 | 0.49635134 | 0.10900318 | 4.55354916 | 5.27E-06 |
| 74041 | DDIAS | 1702.99578 | -0.6713269 | 0.14743586 | -4.553349 | 5.28E-06 |
| 100038614 | Gm10791 | 5.61249975 | -5.9137724 | 1.29884137 | -4.5531136 | 5.29E-06 |
| 20305 | Ccl6 | 6.25591551 | -5.3272034 | 1.17021998 | -4.5523093 | 5.31E-06 |
| 320541 | SLC35E2 | 677.246272 | 0.37341651 | 0.0820395 | 4.55166724 | 5.32E-06 |
| 56219 | EXTL1 | 10.1648711 | -3.4222161 | 0.75186459 | -4.5516389 | 5.32E-06 |
| 60611 | FOXJ2 | 1052.10333 | 0.46133914 | 0.1014072 | 4.5493727 | 5.38E-06 |
| 15365 | Hmga2-ps1 | 783.701882 | -1.0282099 | 0.22603125 | -4.5489721 | 5.39E-06 |
| 243385 | GPRIN3 | 38.3018493 | -3.3193799 | 0.72981025 | -4.548278 | 5.41E-06 |
| 227058 | Dnah7b | 10.448074 | -6.0719823 | 1.3350842 | -4.5480145 | 5.42E-06 |
| 100504586 | Gm20300 | 151.45612 | -0.7653206 | 0.16827602 | -4.5480078 | 5.42E-06 |
| 381224 | Gm340 | 630.046686 | 0.65877748 | 0.1449014 | 4.54638455 | 5.46E-06 |
| 17776 | mast2 | 4568.70566 | 0.67139522 | 0.1476815 | 4.54623777 | 5.46E-06 |
| 13711 | ELF5 | 29.1338775 | -8.2939738 | 1.82480721 | -4.5451233 | 5.49E-06 |
| 230753 | THRAP3 | 3167.62666 | 0.57539866 | 0.12661037 | 4.54464073 | 5.50E-06 |
| 16653 | kras | 2353.80754 | -0.4041674 | 0.0889334 | -4.544607 | 5.50E-06 |
| 329078 | EMX2OS | 4.53927014 | 5.66217511 | 1.2460204 | 4.54420737 | 5.51E-06 |
| 19942 | RPL27 | 252.34258 | 0.55104116 | 0.12126455 | 4.54412418 | 5.52E-06 |
| 236915 | ARHGEF9 | 82.5135201 | -1.1002918 | 0.24215805 | -4.5436927 | 5.53E-06 |
| 102124 | ENKD1 | 414.841474 | -0.4318213 | 0.09504003 | -4.5435732 | 5.53E-06 |
| 13859 | EPS15L1 | 1444.81786 | 0.26114429 | 0.05749059 | 4.54238289 | 5.56E-06 |

| | | | | | | |
|---|---|---|---|---|---|---|
| 64602 | IREB2 | 3349.11003 | 0.24936068 | 0.05490493 | 4.54168119 | 5.58E-06 |
| 66052 | SDHC | 2453.19506 | -0.3953281 | 0.08707023 | -4.5403352 | 5.62E-06 |
| 22418 | WNT5A | 278.426021 | 5.54039908 | 1.22046432 | 4.539583 | 5.64E-06 |
| 229534 | PBXIP1 | 1361.69484 | -1.2198779 | 0.26872305 | -4.5395355 | 5.64E-06 |
| 114615 | ELAC1 | 283.085612 | 0.43763217 | 0.09642923 | 4.53837668 | 5.67E-06 |
| 29862 | NINJ2 | 8.64027232 | -5.0626314 | 1.11572233 | -4.5375371 | 5.69E-06 |
| 54170 | RRAGC | 4954.0717 | 0.46102418 | 0.10160705 | 4.53732489 | 5.70E-06 |
| 16195 | il6st | 6617.46835 | 0.79786628 | 0.17586822 | 4.53672795 | 5.71E-06 |
| 216136 | ILVBL | 839.278991 | -0.6010869 | 0.13251757 | -4.5359039 | 5.74E-06 |
| 66932 | REXO1 | 1780.11544 | 0.19746539 | 0.04353775 | 4.53549796 | 5.75E-06 |
| 14660 | GLS | 9011.41786 | 0.41463495 | 0.09143337 | 4.53483158 | 5.76E-06 |
| 68877 | MAF1 | 1209.40532 | -0.3376577 | 0.07446652 | -4.5343552 | 5.78E-06 |
| 17991 | NDUFA2 | 1046.04879 | -0.3871991 | 0.08541464 | -4.5331698 | 5.81E-06 |
| 330050 | FAM185A | 351.285785 | -0.5465252 | 0.12061556 | -4.531134 | 5.87E-06 |
| 71782 | ANKLE2 | 2116.37638 | -0.2447713 | 0.05403287 | -4.5300442 | 5.90E-06 |
| 93691 | KLF7 | 299.760628 | -0.9999106 | 0.22080334 | -4.5285123 | 5.94E-06 |
| 15465 | HRH1 | 9.98137964 | -3.1742868 | 0.70098863 | -4.5283 | 5.95E-06 |
| 240058 | CPNE5 | 9.89869861 | -2.6290746 | 0.58082663 | -4.5264359 | 6.00E-06 |
| 11622 | AHR | 946.873874 | 0.74275299 | 0.16410571 | 4.52606442 | 6.01E-06 |
| 229731 | SLC25A24 | 6407.53896 | -0.4882848 | 0.10789042 | -4.525748 | 6.02E-06 |
| 214944 | MOB3B | 235.053328 | 1.24425687 | 0.27497147 | 4.52503982 | 6.04E-06 |
| 11771 | AP2A1 | 4992.8724 | 0.37413085 | 0.0826993 | 4.52399077 | 6.07E-06 |
| 109270 | PRR5 | 28.3819279 | -7.5298635 | 1.66448957 | -4.5238274 | 6.07E-06 |
| 24135 | Zfp68 | 1217.71725 | -0.2946835 | 0.06515041 | -4.5231262 | 6.09E-06 |
| 245555 | C77370 | 7.28618368 | -6.2915779 | 1.39121175 | -4.5223726 | 6.12E-06 |
| 26433 | PLOD3 | 3876.19985 | 0.51316415 | 0.11348875 | 4.5217181 | 6.13E-06 |
| 268480 | RAPGEFL1 | 47.2047663 | 1.16924716 | 0.25874618 | 4.51889626 | 6.22E-06 |
| 71711 | MUS81 | 404.460741 | -0.7024327 | 0.15549083 | -4.5175187 | 6.26E-06 |
| 240665 | CCNJ | 232.730132 | -0.5519231 | 0.12217604 | -4.5174416 | 6.26E-06 |
| 70598 | FILIP1 | 15.0943548 | -5.8966189 | 1.30539444 | -4.5171166 | 6.27E-06 |
| 246788 | TRPV3 | 11.721717 | -5.5141867 | 1.22113461 | -4.5156256 | 6.31E-06 |
| 110198 | Akr7a5 | 554.335555 | 0.36947922 | 0.08186944 | 4.51302982 | 6.39E-06 |
| 227154 | STRADB | 265.511405 | -0.5720055 | 0.12674789 | -4.5129391 | 6.39E-06 |
| 16578 | KIF9 | 22.1678178 | -1.9945435 | 0.44198811 | -4.5126634 | 6.40E-06 |
| 20257 | STMN2 | 6.64724883 | 6.20917932 | 1.37637522 | 4.51125481 | 6.44E-06 |
| 13854 | EPN1 | 3369.6596 | 0.2743486 | 0.06081765 | 4.51100265 | 6.45E-06 |
| 50911 | EXOSC9 | 1431.56148 | -0.2866464 | 0.06354443 | -4.5109608 | 6.45E-06 |
| 19062 | INPP5K | 555.40876 | -0.5515569 | 0.12227798 | -4.51068 | 6.46E-06 |
| 666311 | ZSCAN25 | 451.447317 | -0.3679342 | 0.08159532 | -4.5092568 | 6.51E-06 |
| 233103 | 4931406P16I | 1679.35747 | 0.72288995 | 0.16034677 | 4.50829122 | 6.54E-06 |
| 226075 | GLIS3 | 557.087615 | 0.99747877 | 0.22128285 | 4.50770931 | 6.55E-06 |
| 381651 | Gm1045 | 223.997197 | -0.944327 | 0.20949892 | -4.5075505 | 6.56E-06 |
| 67946 | SPATA6 | 374.984461 | -1.1711328 | 0.2598321 | -4.5072676 | 6.57E-06 |
| 22030 | TRAF2 | 928.155845 | -0.3299455 | 0.07321425 | -4.5065747 | 6.59E-06 |
| 72361 | Ces2g | 8.18124108 | -3.9426424 | 0.87500157 | -4.5058689 | 6.61E-06 |

| | | | | | | |
|---|---|---|---|---|---|---|
| 231382 | TMPRSS11D | 9.29420277 | -3.7738242 | 0.83755199 | -4.505779 | 6.61E-06 |
| 66684 | TCEAL8 | 2726.13619 | 0.50246219 | 0.11152383 | 4.50542429 | 6.62E-06 |
| 66341 | EID3 | 110.794421 | 1.34603536 | 0.29887296 | 4.50370402 | 6.68E-06 |
| 52276 | CDCA8 | 2267.14758 | 0.82302181 | 0.1828744 | 4.50047584 | 6.78E-06 |
| 223672 | Apol9a | 171.93029 | -0.837307 | 0.18606795 | -4.5000068 | 6.80E-06 |
| 30878 | apln | 793.788419 | 1.28382558 | 0.28533151 | 4.49941746 | 6.81E-06 |
| 23892 | GREM1 | 50.4780829 | 2.7845719 | 0.61893303 | 4.49898739 | 6.83E-06 |
| 17329 | CXCL9 | 7.55360406 | -6.343624 | 1.41008186 | -4.4987629 | 6.84E-06 |
| 321000 | LRIF1 | 496.577716 | -0.6485937 | 0.14418636 | -4.4983013 | 6.85E-06 |
| 76563 | QRSL1 | 396.268623 | -0.3684222 | 0.08191942 | -4.4973725 | 6.88E-06 |
| 668504 | Gm9209 | 53.0443409 | -1.1166326 | 0.24828706 | -4.4973452 | 6.88E-06 |
| 68991 | SSU72 | 1193.67939 | 0.32741898 | 0.07280834 | 4.49699831 | 6.89E-06 |
| 76614 | IMMT | 6497.0244 | 0.43991817 | 0.09790146 | 4.49347922 | 7.01E-06 |
| 12121 | BICD1 | 279.195987 | -1.0549787 | 0.23479145 | -4.4932587 | 7.01E-06 |
| 75788 | SMURF1 | 3927.84783 | -0.6751109 | 0.15026297 | -4.4928629 | 7.03E-06 |
| 231861 | TNRC18 | 7338.12451 | 0.39311388 | 0.08750157 | 4.49264921 | 7.03E-06 |
| 18035 | NFKBIA | 2556.88514 | 0.52477517 | 0.11682728 | 4.49188877 | 7.06E-06 |
| 20520 | slc22a5 | 193.542355 | -0.7886362 | 0.17569104 | -4.4887676 | 7.16E-06 |
| 69428 | 1700016C15I | 15.5714811 | -6.6573069 | 1.48391703 | -4.4863067 | 7.25E-06 |
| 269610 | CHD5 | 59.7962654 | 1.37213927 | 0.30600517 | 4.48403954 | 7.32E-06 |
| 66079 | TMEM42 | 80.2087198 | 0.98886714 | 0.22060727 | 4.48247754 | 7.38E-06 |
| 109093 | RARS2 | 524.162607 | -1.2999198 | 0.29005175 | -4.4816824 | 7.41E-06 |
| 14407 | GABRG3 | 4.93779266 | -5.7406563 | 1.28102836 | -4.4812875 | 7.42E-06 |
| 108841 | RDH13 | 976.749505 | 0.66279071 | 0.14794854 | 4.4798733 | 7.47E-06 |
| 230917 | TMEM201 | 1254.5117 | 0.55631345 | 0.12419867 | 4.47922234 | 7.49E-06 |
| 209540 | RGAG1 | 6.6601136 | 5.46094847 | 1.21920326 | 4.47911242 | 7.50E-06 |
| 170748 | SMCO4 | 211.191261 | 0.67609902 | 0.15102349 | 4.47678062 | 7.58E-06 |
| 17187 | MAX | 3230.24776 | 0.56841949 | 0.12700431 | 4.47559212 | 7.62E-06 |
| 18617 | Rhox5 | 39.2322689 | 5.70919024 | 1.27567912 | 4.47541248 | 7.63E-06 |
| 56543 | KCND3 | 4.55142886 | -5.6197346 | 1.25643103 | -4.472776 | 7.72E-06 |
| 22687 | ZPR1 | 1630.43989 | 0.40194448 | 0.08988335 | 4.47184561 | 7.75E-06 |
| 14712 | GNPAT | 2200.17357 | 0.41300757 | 0.09235956 | 4.47173595 | 7.76E-06 |
| 442834 | D830031N03 | 1707.42598 | 0.74654087 | 0.16703644 | 4.46932938 | 7.85E-06 |
| 320234 | CCDC66 | 507.346329 | -0.3694318 | 0.08266687 | -4.4689214 | 7.86E-06 |
| 18221 | NUDC | 1995.39896 | 0.63815089 | 0.14281356 | 4.46841951 | 7.88E-06 |
| 238074 | Gm4928 | 240.577978 | 0.76509032 | 0.17125417 | 4.46757196 | 7.91E-06 |
| 57741 | NOC2L | 3570.71668 | 0.93991426 | 0.21050933 | 4.46495298 | 8.01E-06 |
| 19258 | PTPN4 | 712.403102 | -0.5815004 | 0.13027216 | -4.4637347 | 8.05E-06 |
| 13007 | CSRP1 | 3515.75602 | -1.9550621 | 0.4380904 | -4.4626911 | 8.09E-06 |
| 69156 | COMTD1 | 505.228478 | -0.5650289 | 0.1266134 | -4.4626309 | 8.10E-06 |
| 13805 | ENG | 406.494316 | -3.8259666 | 0.85750725 | -4.4617309 | 8.13E-06 |
| 231147 | SH3TC1 | 18.982108 | -4.1594271 | 0.93230332 | -4.4614526 | 8.14E-06 |
| 72522 | ATXN7L2 | 216.965242 | 0.4567348 | 0.1023878 | 4.46083211 | 8.16E-06 |
| 242819 | RUNDC3B | 54.4705991 | -1.3763493 | 0.30854776 | -4.4607333 | 8.17E-06 |
| 100503197 | Gm19589 | 21.587624 | -3.4646816 | 0.77676082 | -4.4604227 | 8.18E-06 |

| | | | | | | |
|---|---|---|---|---|---|---|
| 432442 | AKAP7 | 264.293811 | 0.49661519 | 0.1113722 | 4.45905864 | 8.23E-06 |
| 66979 | POLE4 | 842.239132 | 0.91283457 | 0.20471849 | 4.45897475 | 8.24E-06 |
| 67769 | GPATCH2 | 705.176175 | -0.3908525 | 0.08774418 | -4.4544549 | 8.41E-06 |
| 59095 | FXYD6 | 8.66893273 | -5.8085347 | 1.30400874 | -4.454368 | 8.41E-06 |
| 57276 | VSIG2 | 13.3231292 | 5.73549392 | 1.28781986 | 4.4536461 | 8.44E-06 |
| 75302 | ASXL2 | 3099.09316 | 0.77399203 | 0.17379913 | 4.45337104 | 8.45E-06 |
| 20589 | IGHMBP2 | 535.600539 | 0.46758739 | 0.10501383 | 4.45262696 | 8.48E-06 |
| 74386 | RMI1 | 204.235981 | 0.53227735 | 0.1195633 | 4.45184568 | 8.51E-06 |
| 278795 | LRRC10B | 6.46275485 | 5.42101879 | 1.21778094 | 4.45155495 | 8.53E-06 |
| 18378 | OMP | 29.5431314 | -2.548179 | 0.57244267 | -4.4514135 | 8.53E-06 |
| 319236 | Trim12c | 220.066249 | -1.0567534 | 0.23742383 | -4.4509156 | 8.55E-06 |
| 18619 | PENK | 17.8720016 | 3.32645466 | 0.74737178 | 4.45087005 | 8.55E-06 |
| 20408 | SH3GL3 | 84.6650252 | 1.23133749 | 0.27668467 | 4.45032788 | 8.57E-06 |
| 319953 | TTLL1 | 378.993427 | -0.655432 | 0.14730698 | -4.4494292 | 8.61E-06 |
| 69684 | AARSD1 | 1024.61857 | 0.30881408 | 0.06941159 | 4.44902758 | 8.63E-06 |
| 73847 | fam110a | 661.620325 | -0.5576977 | 0.12544004 | -4.4459301 | 8.75E-06 |
| 19192 | PSME3 | 5225.27432 | 0.23299588 | 0.05241911 | 4.44486503 | 8.79E-06 |
| 100862060 | #N/A | 45.4446902 | 1.40937862 | 0.31714201 | 4.44399847 | 8.83E-06 |
| 73724 | MCEE | 109.651215 | -0.742985 | 0.16723912 | -4.442651 | 8.89E-06 |
| 69116 | UBR4 | 12242.3714 | 0.73194214 | 0.16484734 | 4.4401211 | 8.99E-06 |
| 100126243 | A030001D20 | 16.522366 | 2.21543901 | 0.49903293 | 4.43946456 | 9.02E-06 |
| 66422 | DCTPP1 | 654.049188 | 0.84721665 | 0.19084231 | 4.43935443 | 9.02E-06 |
| 100503397 | Gm11714 | 6.32171867 | -5.3475858 | 1.20495439 | -4.4379985 | 9.08E-06 |
| 545622 | PTPN3 | 743.066115 | -1.0567881 | 0.23812706 | -4.4379168 | 9.08E-06 |
| 73692 | 2410089E03I | 2475.67008 | 0.71389591 | 0.1608887 | 4.43720358 | 9.11E-06 |
| 20641 | SNRPD1 | 1531.70559 | 0.36829402 | 0.08304393 | 4.43493004 | 9.21E-06 |
| 98985 | CLP1 | 423.15696 | 0.35443501 | 0.07994664 | 4.43339478 | 9.28E-06 |
| 382118 | ZKSCAN7 | 67.2764378 | -1.6136562 | 0.36403543 | -4.4326899 | 9.31E-06 |
| 12469 | CCT8 | 13712.4608 | 0.26944784 | 0.06079614 | 4.43198914 | 9.34E-06 |
| 547176 | zc3h12b | 75.6338805 | -1.1512122 | 0.25979623 | -4.4312123 | 9.37E-06 |
| 27413 | ABCB11 | 40.6138212 | -3.2570423 | 0.73511677 | -4.4306462 | 9.40E-06 |
| 16181 | IL1RN | 178.68858 | -3.0514413 | 0.68872458 | -4.4305683 | 9.40E-06 |
| 328977 | Zfp532 | 862.535474 | -0.3510263 | 0.079229 | -4.4305282 | 9.40E-06 |
| 18759 | PRKCI | 3560.51847 | -0.5168091 | 0.11667053 | -4.4296453 | 9.44E-06 |
| 57314 | NELFCD | 1495.06307 | 0.40006151 | 0.09031822 | 4.42946642 | 9.45E-06 |
| 27357 | Gyg | 1144.16258 | -0.5309173 | 0.11987247 | -4.429018 | 9.47E-06 |
| 140919 | SLC17A6 | 6.64303152 | 5.46767905 | 1.23469008 | 4.42838178 | 9.49E-06 |
| 69372 | MOCS3 | 344.331955 | 0.4816592 | 0.10877475 | 4.42804236 | 9.51E-06 |
| 104836 | CBLL1 | 995.546859 | 0.40236299 | 0.09087377 | 4.427713 | 9.52E-06 |
| 17910 | Myo15 | 9.12203916 | -4.0947106 | 0.92481696 | -4.4275903 | 9.53E-06 |
| 107895 | MGAT5 | 1543.59823 | -0.9294651 | 0.209951 | -4.4270574 | 9.55E-06 |
| 268902 | ROBO2 | 155.627583 | 4.40408654 | 0.99482898 | 4.42697856 | 9.56E-06 |
| 56079 | ASTN2 | 5.25730486 | -5.8311601 | 1.31737915 | -4.426334 | 9.58E-06 |
| 224671 | BTBD9 | 828.230314 | -0.5134714 | 0.11601559 | -4.4258829 | 9.60E-06 |
| 93891 | Pcdhb20 | 37.6032647 | 1.51051031 | 0.34132317 | 4.42545499 | 9.62E-06 |

| | | | | | | |
|---|---|---|---|---|---|---|
| 28254 | Slco1a6 | 37.4067441 | -2.7763481 | 0.6274545 | -4.4247799 | 9.65E-06 |
| 100206 | ADPRHL2 | 520.601781 | 0.43497776 | 0.098307 | 4.42468777 | 9.66E-06 |
| 106025 | SHARPIN | 697.970768 | -0.4885858 | 0.1104321 | -4.4243098 | 9.68E-06 |
| 15936 | IER2 | 1181.75198 | 0.53117958 | 0.12008882 | 4.42322247 | 9.72E-06 |
| 235626 | SETD2 | 3270.27323 | 0.33643606 | 0.07607093 | 4.42266295 | 9.75E-06 |
| 17427 | MNS1 | 183.391503 | 0.56125255 | 0.1269306 | 4.42172766 | 9.79E-06 |
| 257633 | ACSF3 | 413.989869 | -0.6418821 | 0.14521748 | -4.4201432 | 9.86E-06 |
| 218441 | ZFYVE16 | 1128.14052 | 0.35891286 | 0.08121227 | 4.41944117 | 9.90E-06 |
| 18415 | HSPA4L | 3457.45115 | 0.65670819 | 0.14859814 | 4.41935679 | 9.90E-06 |
| 12834 | COL6A2 | 380.916913 | 2.1070943 | 0.47687786 | 4.41851991 | 9.94E-06 |
| 18951 | 41886 | 765.634364 | 0.71800241 | 0.16250978 | 4.41821041 | 9.95E-06 |
| 105000 | DNAL1 | 539.37113 | -0.5669638 | 0.12836304 | -4.4168774 | 1.00E-05 |
| 319478 | cxxc4 | 10.9153273 | 2.82657982 | 0.64005431 | 4.41615624 | 1.00E-05 |
| 22200 | UBA3 | 2984.74548 | 0.48103059 | 0.10898283 | 4.41381975 | 1.02E-05 |
| 244713 | Zfp317 | 1039.59566 | 0.29471545 | 0.06679034 | 4.41254616 | 1.02E-05 |
| 213391 | RASSF4 | 1723.24987 | -4.7612779 | 1.07929056 | -4.4114885 | 1.03E-05 |
| 70350 | BASP1 | 2925.04112 | -3.3861457 | 0.76784114 | -4.409956 | 1.03E-05 |
| 74251 | ANKRD9 | 278.190488 | -0.591916 | 0.13424717 | -4.4091507 | 1.04E-05 |
| 75625 | MAGEH1 | 11.3724459 | -6.9353039 | 1.57296255 | -4.4090712 | 1.04E-05 |
| 20278 | SCNN1G | 4.52214085 | -5.6118481 | 1.27291778 | -4.4086493 | 1.04E-05 |
| 71279 | SLC29A3 | 760.929549 | -0.7175477 | 0.16278234 | -4.4080191 | 1.04E-05 |
| 69085 | ZCCHC9 | 693.418335 | 0.4326534 | 0.09817719 | 4.40686287 | 1.05E-05 |
| 68145 | ETAA1 | 528.789708 | 0.44480449 | 0.10094315 | 4.40648504 | 1.05E-05 |
| 106522 | PKDCC | 78.4914504 | 1.23985148 | 0.28137118 | 4.4064622 | 1.05E-05 |
| 319901 | DSEL | 1096.43659 | -0.4032362 | 0.09153011 | -4.4055035 | 1.06E-05 |
| 219151 | SCARA3 | 756.665449 | 4.81873222 | 1.0940929 | 4.40431723 | 1.06E-05 |
| 68948 | FAM216A | 723.843632 | -0.4899721 | 0.1112563 | -4.4039942 | 1.06E-05 |
| 170677 | CDHR1 | 179.098451 | -4.2110628 | 0.95635483 | -4.4032431 | 1.07E-05 |
| 14165 | FGF10 | 867.898914 | 6.625623 | 1.50487033 | 4.40278664 | 1.07E-05 |
| 268297 | SCML4 | 57.8913301 | -3.6227535 | 0.8230837 | -4.40144 | 1.08E-05 |
| 56321 | AATF | 1448.64075 | 0.2918588 | 0.06632446 | 4.40046997 | 1.08E-05 |
| 432450 | NKAIN2 | 11.1765591 | -6.9104862 | 1.57101428 | -4.3987418 | 1.09E-05 |
| 75033 | MEI4 | 14.1139846 | -7.2474542 | 1.64802087 | -4.3976714 | 1.09E-05 |
| 320683 | Zfp629 | 547.908459 | 0.54920087 | 0.12490054 | 4.39710577 | 1.10E-05 |
| 16581 | KIFC2 | 24.4796367 | -1.6531757 | 0.3759875 | -4.39689 | 1.10E-05 |
| 320302 | Glt28d2 | 107.325785 | -1.0619957 | 0.24156781 | -4.3962632 | 1.10E-05 |
| 17886 | MYH9 | 56545.3521 | -0.3613032 | 0.08218555 | -4.3961882 | 1.10E-05 |
| 17711 | CYTB | 67448.2556 | -0.5814074 | 0.13226571 | -4.395753 | 1.10E-05 |
| 17314 | MGMT | 8.29747488 | -6.4790269 | 1.4742738 | -4.3947243 | 1.11E-05 |
| 75593 | MALSU1 | 353.351528 | -0.4687767 | 0.10667796 | -4.3943162 | 1.11E-05 |
| 216033 | CTNNA3 | 7.66780285 | -3.8312005 | 0.87204832 | -4.3933352 | 1.12E-05 |
| 12043 | BCL2 | 482.607941 | -0.6423687 | 0.14628652 | -4.3911681 | 1.13E-05 |
| 54709 | EIF3I | 3209.6167 | 0.74367147 | 0.16941791 | 4.38956826 | 1.14E-05 |
| 69126 | 1810022K09I | 177.413729 | -0.5224003 | 0.1190355 | -4.3886092 | 1.14E-05 |
| 20751 | SPR | 1248.64808 | 0.5496906 | 0.12526398 | 4.38825735 | 1.14E-05 |

| | | | | | | |
|---|---|---|---|---|---|---|
| 434232 | IQCK | 31.0099478 | -2.2715331 | 0.51798117 | -4.3853584 | 1.16E-05 |
| 67446 | dusp28 | 66.5233191 | -1.5129767 | 0.34501626 | -4.3852329 | 1.16E-05 |
| 99683 | SEC24B | 3161.32271 | 0.24171356 | 0.05516638 | 4.38153722 | 1.18E-05 |
| 100126768 | Esp31 | 4.59898595 | -5.637538 | 1.2869085 | -4.3806828 | 1.18E-05 |
| 52440 | TAX1BP1 | 4855.18161 | -0.3168424 | 0.07232946 | -4.3805436 | 1.18E-05 |
| 12537 | CDK11B | 1772.36484 | 0.41098377 | 0.09382286 | 4.38042241 | 1.18E-05 |
| 22630 | YWHAQ | 1030.69549 | 0.52534813 | 0.11995788 | 4.37943819 | 1.19E-05 |
| 75659 | WDR54 | 154.60297 | 0.67346677 | 0.15378394 | 4.37930496 | 1.19E-05 |
| 213541 | YTHDF2 | 1911.39835 | 0.63350624 | 0.14468239 | 4.37859938 | 1.19E-05 |
| 14794 | SPSB2 | 557.668913 | 1.20262778 | 0.27466537 | 4.37851989 | 1.19E-05 |
| 68267 | SLC25A22 | 907.053033 | 0.60400561 | 0.13796366 | 4.37800515 | 1.20E-05 |
| 18700 | PIGA | 669.031388 | 0.53578421 | 0.12238355 | 4.37791043 | 1.20E-05 |
| 100715 | PAPD4 | 424.39353 | 0.60668507 | 0.13858485 | 4.37771566 | 1.20E-05 |
| 68666 | SVOP | 11.9221236 | -4.5305317 | 1.0349865 | -4.3773824 | 1.20E-05 |
| 106073 | MFSD5 | 1768.9999 | -0.7131242 | 0.16296376 | -4.375968 | 1.21E-05 |
| 58186 | RAD18 | 1524.12686 | 0.65670118 | 0.15007364 | 4.37585962 | 1.21E-05 |
| 118445 | KLF16 | 1338.22737 | 0.3748881 | 0.08568582 | 4.37514728 | 1.21E-05 |
| 67628 | ANP32B | 5320.37495 | 0.61450176 | 0.14045898 | 4.3749554 | 1.21E-05 |
| 30955 | PIK3CG | 13.9511936 | -4.7412948 | 1.0841289 | -4.3733682 | 1.22E-05 |
| 66464 | TAF12 | 658.589518 | 0.79612078 | 0.18210573 | 4.37175022 | 1.23E-05 |
| 269713 | CLIP2 | 2017.36937 | 0.65325719 | 0.14947198 | 4.37043237 | 1.24E-05 |
| 11702 | AMD1 | 775.422671 | -0.3438381 | 0.07867521 | -4.3703488 | 1.24E-05 |
| 228869 | NCOA5 | 1716.4271 | 0.36977537 | 0.0846137 | 4.37015947 | 1.24E-05 |
| 20930 | SURF1 | 428.261834 | -0.5145627 | 0.11775464 | -4.3697875 | 1.24E-05 |
| 13427 | DYNC1I2 | 4066.16844 | -0.3013611 | 0.06896696 | -4.3696444 | 1.24E-05 |
| 12836 | COL7A1 | 1690.64015 | -2.2990394 | 0.52636213 | -4.3677904 | 1.26E-05 |
| 26939 | POLR3E | 1285.11871 | 0.66654273 | 0.15266961 | 4.36591628 | 1.27E-05 |
| 68180 | HYI | 90.1719178 | 1.3763632 | 0.31528861 | 4.36540725 | 1.27E-05 |
| 14731 | GPAA1 | 1857.13346 | -0.3044101 | 0.06975945 | -4.3637105 | 1.28E-05 |
| 78232 | TRAPPC6B | 841.024586 | 0.377268 | 0.08646922 | 4.36303216 | 1.28E-05 |
| 211896 | DEPDC7 | 939.168801 | -2.1770553 | 0.49899023 | -4.3629216 | 1.28E-05 |
| 93876 | PCDHB5 | 37.929585 | 2.59747565 | 0.59537371 | 4.36276508 | 1.28E-05 |
| 76453 | PRSS23 | 3272.13644 | -1.8456002 | 0.42306507 | -4.3624499 | 1.29E-05 |
| 320977 | A330023F24 | 20.7219445 | -2.5837159 | 0.59229052 | -4.3622442 | 1.29E-05 |
| 74626 | TMEM81 | 94.3777367 | -1.0388418 | 0.2382009 | -4.3612 | 1.29E-05 |
| 170756 | SLC8B1 | 854.440164 | -0.9662989 | 0.22156987 | -4.3611478 | 1.29E-05 |
| 11306 | ABCB7 | 1921.0654 | -0.4725465 | 0.10837695 | -4.3602124 | 1.30E-05 |
| 72415 | sgol1 | 1321.83499 | -0.2205848 | 0.05059662 | -4.3596739 | 1.30E-05 |
| 11350 | abl1 | 4477.93203 | 0.5440879 | 0.1248579 | 4.35765681 | 1.31E-05 |
| 19281 | PTPRT | 5.62641982 | -5.9177737 | 1.35811712 | -4.3573368 | 1.32E-05 |
| 52389 | ADGRA1 | 22.8104023 | -4.2499331 | 0.97555314 | -4.3564343 | 1.32E-05 |
| 245886 | ANKRD27 | 1502.03063 | 0.66858053 | 0.15348159 | 4.35609583 | 1.32E-05 |
| 14828 | HSPA5 | 20776.5954 | 0.64884663 | 0.14896372 | 4.35573603 | 1.33E-05 |
| 74108 | PARN | 1234.48416 | -0.2951199 | 0.06776976 | -4.3547434 | 1.33E-05 |
| 213350 | PDDC1 | 300.287826 | 0.44904179 | 0.10312324 | 4.35441912 | 1.33E-05 |

| | | | | | | |
|---|---|---|---|---|---|---|
| 192185 | NADK | 6087.3723 | 0.50100417 | 0.11508054 | 4.35350898 | 1.34E-05 |
| 140858 | WDR5 | 2765.01844 | 0.31477826 | 0.07230962 | 4.35320026 | 1.34E-05 |
| 12269 | c4bp | 36.2876832 | -4.4509089 | 1.02271852 | -4.3520371 | 1.35E-05 |
| 54383 | PHC2 | 6886.53464 | 0.82432774 | 0.18949352 | 4.35016335 | 1.36E-05 |
| 98386 | LBR | 3948.32075 | 0.49561281 | 0.11395987 | 4.34901162 | 1.37E-05 |
| 329504 | lcmt2 | 431.584442 | 0.52099123 | 0.11982673 | 4.34787147 | 1.37E-05 |
| 100568459 | Bc1 | 1375.0752 | 1.42817172 | 0.32851123 | 4.34740608 | 1.38E-05 |
| 18634 | PEX7 | 406.162613 | -0.530529 | 0.12204045 | -4.3471566 | 1.38E-05 |
| 109880 | BRAF | 1487.90351 | 0.39577229 | 0.09106983 | 4.34581105 | 1.39E-05 |
| 110616 | ATXN3 | 1527.62438 | 0.53875446 | 0.1240024 | 4.34470976 | 1.39E-05 |
| 68813 | DOCK5 | 5136.87149 | 0.70639515 | 0.16260764 | 4.34416956 | 1.40E-05 |
| 20716 | Serpina3n | 374.994256 | -3.085076 | 0.71030878 | -4.3432886 | 1.40E-05 |
| 93837 | DACH2 | 16.8164698 | -6.7677824 | 1.55883544 | -4.3415631 | 1.41E-05 |
| 11767 | AP1M1 | 1738.67365 | 0.43956377 | 0.1012491 | 4.34140904 | 1.42E-05 |
| 16790 | ANPEP | 174.136414 | -5.364649 | 1.23595813 | -4.340478 | 1.42E-05 |
| 242687 | WASF2 | 3078.46469 | 0.65630987 | 0.15120944 | 4.34040276 | 1.42E-05 |
| 11857 | ARHGDIB | 7974.33343 | 4.80247693 | 1.10663569 | 4.33970907 | 1.43E-05 |
| 17685 | MSH2 | 1602.94809 | -0.2909311 | 0.06704244 | -4.3395058 | 1.43E-05 |
| 18570 | PDCD6 | 2127.12536 | -0.2940752 | 0.06777701 | -4.3388627 | 1.43E-05 |
| 210126 | LPP | 5491.28588 | -0.5959991 | 0.13736432 | -4.3388203 | 1.43E-05 |
| 70510 | RNF167 | 427.016849 | -0.7777254 | 0.17926344 | -4.3384495 | 1.43E-05 |
| 67819 | DERL1 | 3010.26001 | 0.42531143 | 0.09805457 | 4.33749708 | 1.44E-05 |
| 22393 | WFS1 | 735.922951 | -0.8569658 | 0.19757788 | -4.3373568 | 1.44E-05 |
| 72201 | OTUD6B | 1883.63055 | -0.7659803 | 0.17662252 | -4.3368212 | 1.45E-05 |
| 16553 | KIF13A | 1313.11429 | 0.6413296 | 0.14795963 | 4.3344904 | 1.46E-05 |
| 78697 | PUS7 | 1460.19695 | 1.15587444 | 0.26677431 | 4.33278019 | 1.47E-05 |
| 208146 | YEATS2 | 1591.84748 | -0.3462327 | 0.07993052 | -4.3316713 | 1.48E-05 |
| 66965 | CTU2 | 727.456497 | -0.4236052 | 0.097824 | -4.3302793 | 1.49E-05 |
| 225594 | Gm4841 | 6.09435451 | -5.2885425 | 1.22159349 | -4.3292163 | 1.50E-05 |
| 14942 | Gzme | 5.27006099 | -5.8222712 | 1.34496597 | -4.3289357 | 1.50E-05 |
| 242425 | GABBR2 | 7.76506542 | -6.3907063 | 1.47698229 | -4.3268673 | 1.51E-05 |
| 399558 | FLRT2 | 4855.57287 | 4.57710301 | 1.05798069 | 4.32626328 | 1.52E-05 |
| 107939 | pom121 | 4327.25043 | 0.31919378 | 0.07382006 | 4.32394366 | 1.53E-05 |
| 207375 | FAM120C | 568.845647 | 0.7663893 | 0.17724458 | 4.3239082 | 1.53E-05 |
| 23928 | LAMC3 | 30.7327886 | -3.9292282 | 0.90882378 | -4.3234215 | 1.54E-05 |
| 73218 | SPPL2B | 1177.43808 | 0.38791464 | 0.08975973 | 4.32170006 | 1.55E-05 |
| 98999 | ZNFX1 | 1316.43087 | -0.5271869 | 0.12198983 | -4.3215644 | 1.55E-05 |
| 19167 | PSMA3 | 290.037264 | 0.56328091 | 0.13034216 | 4.32155585 | 1.55E-05 |
| 17691 | SIK1 | 756.215372 | -1.7234413 | 0.39887184 | -4.3207896 | 1.55E-05 |
| 26356 | ING1 | 1301.95735 | 0.30212216 | 0.06994406 | 4.31948287 | 1.56E-05 |
| 237412 | Gm4924 | 148.451186 | -0.5644786 | 0.1307007 | -4.3188645 | 1.57E-05 |
| 76779 | CLUAP1 | 269.316276 | -0.6454694 | 0.14947432 | -4.3182627 | 1.57E-05 |
| 244562 | ABCC12 | 4.29168986 | -5.5290698 | 1.28057511 | -4.3176458 | 1.58E-05 |
| 67115 | RPL14 | 2279.60602 | 0.25610453 | 0.05931879 | 4.31742681 | 1.58E-05 |
| 101612 | GRWD1 | 2057.02346 | 0.65373463 | 0.15147145 | 4.31589338 | 1.59E-05 |

| | | | | | | |
|---|---|---|---|---|---|---|
| 13211 | DHX9 | 14026.017 | 0.31030649 | 0.0719036 | 4.31559043 | 1.59E-05 |
| 50768 | DLC1 | 1521.7092 | 1.97114346 | 0.45676999 | 4.31539616 | 1.59E-05 |
| 66500 | SLC30A7 | 1869.56063 | -0.4266271 | 0.09889658 | -4.3138716 | 1.60E-05 |
| 100042625 | Gstp-ps | 1227.15021 | 0.42045842 | 0.09746985 | 4.31372779 | 1.61E-05 |
| 108073 | GRM7 | 5.451428 | 5.16752131 | 1.19815937 | 4.3128831 | 1.61E-05 |
| 71927 | ITFG1 | 2361.32453 | -0.49845 | 0.11557259 | -4.3128736 | 1.61E-05 |
| 26400 | MAP2K7 | 1809.07018 | 0.25940388 | 0.06014858 | 4.3127184 | 1.61E-05 |
| 13138 | DAG1 | 6888.92847 | -0.5608121 | 0.13010534 | -4.3104468 | 1.63E-05 |
| 226442 | Zfp281 | 1954.3308 | -0.4439453 | 0.10301384 | -4.3095694 | 1.64E-05 |
| 73158 | LARP1 | 11520.8215 | 0.30668996 | 0.07117366 | 4.30903709 | 1.64E-05 |
| 77065 | INTS7 | 1994.93086 | 0.34350639 | 0.07972049 | 4.30888442 | 1.64E-05 |
| 241226 | ITGA8 | 682.716556 | -2.779636 | 0.64510392 | -4.3088189 | 1.64E-05 |
| 58520 | 0610007P14I | 1568.92872 | 0.43557584 | 0.1010962 | 4.30852832 | 1.64E-05 |
| 434402 | Gm5617 | 33.858223 | 1.18903731 | 0.2759776 | 4.30845584 | 1.64E-05 |
| 235048 | Zfp599 | 112.277865 | -0.9055687 | 0.2101934 | -4.3082643 | 1.65E-05 |
| 14534 | KAT2A | 2949.87036 | 0.37227167 | 0.0864517 | 4.306123 | 1.66E-05 |
| 12606 | CEBPA | 735.553333 | 2.94333397 | 0.68353859 | 4.30602459 | 1.66E-05 |
| 207592 | TBC1D16 | 640.443478 | -2.0201765 | 0.46925306 | -4.3050897 | 1.67E-05 |
| 102442 | DENND4A | 1001.16888 | 0.42371977 | 0.09843319 | 4.30464324 | 1.67E-05 |
| 68355 | 2010204K13I | 99.7836547 | -0.6889025 | 0.16007097 | -4.3037315 | 1.68E-05 |
| 63913 | FAM129A | 968.554956 | -1.5551484 | 0.36137887 | -4.3033741 | 1.68E-05 |
| 11637 | AK2 | 2506.85099 | 0.68377949 | 0.15891823 | 4.30271275 | 1.69E-05 |
| 17535 | mre11a | 1645.5422 | 0.37512373 | 0.08718364 | 4.30268475 | 1.69E-05 |
| 171504 | APOBR | 564.708036 | 2.38504121 | 0.55448012 | 4.30140079 | 1.70E-05 |
| 30940 | USP25 | 2278.20207 | -0.4101117 | 0.09535801 | -4.3007576 | 1.70E-05 |
| 77827 | KRBA1 | 221.588449 | -2.4318781 | 0.56547403 | -4.3006009 | 1.70E-05 |
| 70967 | EVA1C | 215.715644 | -2.3090112 | 0.53698999 | -4.2999148 | 1.71E-05 |
| 226646 | NDUFS2 | 3568.44993 | -0.3567076 | 0.08297228 | -4.299118 | 1.71E-05 |
| 100504564 | #N/A | 9.26710922 | -3.7740021 | 0.87803493 | -4.2982368 | 1.72E-05 |
| 69597 | AFG3L2 | 3040.67082 | -0.1834306 | 0.04267765 | -4.298049 | 1.72E-05 |
| 66290 | ATP6V1G1 | 1411.15894 | -0.9192298 | 0.21390315 | -4.2974114 | 1.73E-05 |
| 110460 | ACAT2 | 1454.97939 | 0.84541853 | 0.19673735 | 4.29719378 | 1.73E-05 |
| 252875 | MIOS | 678.186795 | 0.40975048 | 0.09536736 | 4.29654862 | 1.73E-05 |
| 56464 | CTSF | 11.0757685 | -4.8632191 | 1.13210782 | -4.2957208 | 1.74E-05 |
| 229672 | BCL2L15 | 9.88269888 | -3.3392091 | 0.7775408 | -4.2945773 | 1.75E-05 |
| 68028 | RPL22L1 | 612.182914 | -0.5595385 | 0.13032283 | -4.2934804 | 1.76E-05 |
| 109006 | CIAPIN1 | 2157.26575 | 0.41285625 | 0.09616567 | 4.2931773 | 1.76E-05 |
| 68183 | BCAS2 | 1860.65098 | -0.2903159 | 0.06762306 | -4.2931484 | 1.76E-05 |
| 80876 | IFITM2 | 1751.5067 | 0.94792178 | 0.22080595 | 4.29300832 | 1.76E-05 |
| 19188 | PSME2 | 187.520615 | -0.8808994 | 0.205241 | -4.2920244 | 1.77E-05 |
| 269582 | CLSPN | 1959.83621 | 0.93409351 | 0.21768854 | 4.29096314 | 1.78E-05 |
| 19072 | PREP | 5057.85441 | 0.4064876 | 0.09475688 | 4.28979524 | 1.79E-05 |
| 626870 | Gm11992 | 13.348476 | -3.0080925 | 0.70125543 | -4.2895818 | 1.79E-05 |
| 19415 | RASAL1 | 10.6043171 | -6.0936945 | 1.42063599 | -4.2894131 | 1.79E-05 |
| 18286 | ODF2 | 2466.462 | 0.5950064 | 0.13871924 | 4.28928538 | 1.79E-05 |

| | | | | | | |
|---|---|---|---|---|---|---|
| 72318 | CYTH4 | 20.2763399 | -3.8045772 | 0.88703212 | -4.2891088 | 1.79E-05 |
| 13836 | EPHA2 | 5870.12391 | -0.2844983 | 0.06635008 | -4.287837 | 1.80E-05 |
| 100042165 | BC005561 | 323.413532 | -0.4134913 | 0.09643568 | -4.2877415 | 1.80E-05 |
| 279653 | PCDH19 | 4239.03167 | 3.66791825 | 0.85553455 | 4.28728241 | 1.81E-05 |
| 230673 | IPO13 | 1866.20608 | 0.67021335 | 0.15632951 | 4.28718371 | 1.81E-05 |
| 68366 | TMEM129 | 881.867726 | -0.4147786 | 0.09675041 | -4.2870992 | 1.81E-05 |
| 223722 | MCAT | 612.938441 | 0.31751116 | 0.07406703 | 4.28680855 | 1.81E-05 |
| 215951 | LACE1 | 205.857472 | -0.7833444 | 0.18276634 | -4.2860432 | 1.82E-05 |
| 72309 | TMEM158 | 217.687859 | 1.24053218 | 0.28944 | 4.28597356 | 1.82E-05 |
| 76784 | MTIF2 | 701.16619 | -0.3581712 | 0.08357067 | -4.2858488 | 1.82E-05 |
| 22608 | YBX1 | 8983.54906 | 0.92989594 | 0.21699045 | 4.28542339 | 1.82E-05 |
| 69601 | DAB2IP | 2037.02749 | -0.6097349 | 0.14229614 | -4.2849712 | 1.83E-05 |
| 231855 | AP5Z1 | 473.851065 | -0.896233 | 0.20919549 | -4.2841887 | 1.83E-05 |
| 27404 | Abca8b | 207.641859 | -1.4763221 | 0.34466415 | -4.2833641 | 1.84E-05 |
| 210710 | GAB3 | 16.9441407 | -1.9996089 | 0.46693126 | -4.2824481 | 1.85E-05 |
| 18947 | PNLIPRP2 | 7.50901103 | -6.3363253 | 1.47968529 | -4.2822114 | 1.85E-05 |
| 72301 | 1810041L15R | 24.9401478 | -8.0711611 | 1.88504589 | -4.2816789 | 1.85E-05 |
| 103742 | MIEN1 | 561.259851 | -0.4976152 | 0.11625446 | -4.2803965 | 1.87E-05 |
| 57317 | SRSF4 | 1477.20134 | 0.86664644 | 0.20253611 | 4.27897255 | 1.88E-05 |
| 20190 | RYR1 | 33.5294576 | -1.3803702 | 0.32260581 | -4.2788136 | 1.88E-05 |
| 381822 | Lockd | 168.126792 | 0.87295158 | 0.20404416 | 4.27824823 | 1.88E-05 |
| 234673 | Ces2e | 3.97910882 | -5.4238026 | 1.26779625 | -4.2781343 | 1.88E-05 |
| 22097 | Tsix | 3.97085772 | -5.4217519 | 1.26749812 | -4.2775226 | 1.89E-05 |
| 223920 | SOAT2 | 12.302817 | -2.1801044 | 0.50973101 | -4.2769702 | 1.89E-05 |
| 76137 | MCUR1 | 1137.92226 | 0.56435581 | 0.13196553 | 4.2765396 | 1.90E-05 |
| 11927 | ATOX1 | 1551.23478 | -0.308468 | 0.07213118 | -4.2764864 | 1.90E-05 |
| 66308 | mplkip | 244.887956 | 0.67407674 | 0.15763409 | 4.27621168 | 1.90E-05 |
| 11610 | AGTRAP | 709.754223 | -0.8096392 | 0.18934811 | -4.2759296 | 1.90E-05 |
| 66607 | Ms4a4d | 8.87816877 | -6.5769035 | 1.53814781 | -4.2758592 | 1.90E-05 |
| 216873 | SPAG7 | 1264.26552 | 0.20624601 | 0.04823931 | 4.27547616 | 1.91E-05 |
| 53893 | NUDT5 | 969.27703 | 0.43362222 | 0.10143116 | 4.27503966 | 1.91E-05 |
| 226744 | CNST | 680.746508 | 0.41066448 | 0.09607724 | 4.27431598 | 1.92E-05 |
| 26926 | AIFM1 | 1957.39103 | -0.1940583 | 0.04540141 | -4.2742802 | 1.92E-05 |
| 20197 | S100A3 | 112.338828 | 0.99928615 | 0.2338304 | 4.27355098 | 1.92E-05 |
| 236794 | SLC9A6 | 873.509153 | -0.3685997 | 0.0862632 | -4.2729659 | 1.93E-05 |
| 15481 | HSPA8 | 1056.9012 | 0.35123479 | 0.08220821 | 4.27250231 | 1.93E-05 |
| 240034 | Zfp760 | 184.848115 | -1.1377156 | 0.26631543 | -4.2720604 | 1.94E-05 |
| 102580 | ALG9 | 972.051265 | 0.27318141 | 0.06394985 | 4.27180714 | 1.94E-05 |
| 228410 | CSTF3 | 1320.66323 | 0.33402518 | 0.07821111 | 4.27081518 | 1.95E-05 |
| 94232 | UBQLN4 | 3672.02471 | 0.35857567 | 0.08396898 | 4.27033476 | 1.95E-05 |
| 228775 | TRIB3 | 812.069076 | -0.6378146 | 0.14935944 | -4.2703334 | 1.95E-05 |
| 140859 | NEK8 | 299.737294 | -0.4127034 | 0.09665448 | -4.2698841 | 1.96E-05 |
| 225027 | SRSF7 | 4103.08944 | 0.41718628 | 0.0977196 | 4.26921793 | 1.96E-05 |
| 328186 | Gm10336 | 403.036979 | 0.92945755 | 0.21772977 | 4.26885833 | 1.96E-05 |
| 71240 | OSBPL7 | 470.382476 | -0.7598733 | 0.17802866 | -4.2682641 | 1.97E-05 |

| | | | | | | |
|---|---|---|---|---|---|---|
| 13175 | DCLK1 | 1730.35373 | -1.9522 | 0.45742105 | -4.2678402 | 1.97E-05 |
| 268880 | XXYLT1 | 1353.73577 | 0.80754903 | 0.18932011 | 4.26552156 | 1.99E-05 |
| 11489 | ADAM12 | 2995.05191 | -4.7328226 | 1.10976946 | -4.2646899 | 2.00E-05 |
| 73711 | MVB12A | 1032.77349 | -0.8104796 | 0.19005217 | -4.2645113 | 2.00E-05 |
| 70620 | UBE2V2 | 326.918106 | -0.5690012 | 0.13344665 | -4.2638849 | 2.01E-05 |
| 269233 | FAM171A1 | 1473.69531 | 1.99915699 | 0.46897979 | 4.26277858 | 2.02E-05 |
| 224897 | DPP9 | 2911.67734 | 0.32778472 | 0.07691021 | 4.2619143 | 2.03E-05 |
| 207596 | THSD4 | 312.99446 | -2.5456774 | 0.5973423 | -4.2616727 | 2.03E-05 |
| 381511 | PDP1 | 368.044371 | -0.9902195 | 0.23238649 | -4.2610887 | 2.03E-05 |
| 233060 | Zfp382 | 190.830824 | 0.69082369 | 0.16212445 | 4.26107029 | 2.03E-05 |
| 83396 | GLIS2 | 1449.57345 | -0.6767312 | 0.15887996 | -4.2593869 | 2.05E-05 |
| 28088 | RTCB | 4657.9867 | -0.2168965 | 0.05093338 | -4.2584353 | 2.06E-05 |
| 230806 | AIM1L | 7.39223481 | -4.2041706 | 0.98729387 | -4.2582768 | 2.06E-05 |
| 53627 | porcn | 80.0203799 | -1.870786 | 0.43935907 | -4.2579888 | 2.06E-05 |
| 69202 | PTMS | 9275.69935 | 0.83557725 | 0.19634421 | 4.25567553 | 2.08E-05 |
| 16419 | ITGB5 | 3627.80156 | -0.6161057 | 0.14481253 | -4.2545052 | 2.10E-05 |
| 320799 | ZHX3 | 373.023096 | 0.56186191 | 0.13208085 | 4.2539241 | 2.10E-05 |
| 24057 | SH3YL1 | 206.932298 | -0.6585671 | 0.15482412 | -4.2536468 | 2.10E-05 |
| 327762 | DNA2 | 411.56476 | 0.42892248 | 0.10084478 | 4.25329387 | 2.11E-05 |
| 18812 | Prl2c3 | 1050.8267 | -1.9750153 | 0.4643689 | -4.253117 | 2.11E-05 |
| 212728 | TARBP1 | 669.474405 | 0.56488872 | 0.13283982 | 4.25240505 | 2.11E-05 |
| 66706 | NDUFAF3 | 160.498118 | -0.6312365 | 0.14844646 | -4.2522842 | 2.12E-05 |
| 53897 | GAL3ST1 | 19.3163941 | -3.8785905 | 0.9125103 | -4.2504622 | 2.13E-05 |
| 381126 | GAREM1 | 228.171143 | 0.56181135 | 0.1321829 | 4.25025748 | 2.14E-05 |
| 97487 | CMTM4 | 5163.80695 | -0.1923531 | 0.0452731 | -4.2487284 | 2.15E-05 |
| 74648 | S100PBP | 305.576945 | 0.46082928 | 0.10847435 | 4.2482788 | 2.15E-05 |
| 16372 | IRX2 | 2047.18273 | 0.41101994 | 0.09675074 | 4.24823559 | 2.15E-05 |
| 98365 | SLAMF9 | 6.18928282 | -6.0660815 | 1.42791859 | -4.2481984 | 2.15E-05 |
| 100038607 | Gm10636 | 9.90119907 | -2.9421096 | 0.69261723 | -4.2478146 | 2.16E-05 |
| 12972 | CRYZ | 842.654896 | 0.69573938 | 0.16384394 | 4.24635396 | 2.17E-05 |
| 13972 | GNB1L | 264.780606 | -0.7383687 | 0.17392188 | -4.2454042 | 2.18E-05 |
| 77220 | TMEM200A | 7.38279511 | 4.29215663 | 1.01121518 | 4.24455319 | 2.19E-05 |
| 69742 | TM2D2 | 877.15554 | -0.3784026 | 0.08918851 | -4.242728 | 2.21E-05 |
| 208440 | DIP2C | 1510.03627 | 0.59578888 | 0.1405012 | 4.24045407 | 2.23E-05 |
| 77446 | HEG1 | 639.346151 | -2.5914645 | 0.61146639 | -4.2381144 | 2.25E-05 |
| 381812 | CRACR2A | 8.65425234 | 3.49530995 | 0.82477922 | 4.23787343 | 2.26E-05 |
| 70645 | OIP5 | 512.915058 | -0.3282891 | 0.07746888 | -4.2376906 | 2.26E-05 |
| 22282 | USF2 | 1178.96686 | 0.54955461 | 0.12971589 | 4.23660207 | 2.27E-05 |
| 106794 | DHX57 | 777.736617 | -0.4320169 | 0.10197393 | -4.2365425 | 2.27E-05 |
| 27058 | SRP9 | 265.739983 | -0.5981919 | 0.14120022 | -4.2364798 | 2.27E-05 |
| 270120 | FAT3 | 5.14736275 | -5.7884441 | 1.36709076 | -4.234133 | 2.29E-05 |
| 21815 | TGIF1 | 2506.04191 | -0.4145281 | 0.09793637 | -4.232627 | 2.31E-05 |
| 235582 | GLYCTK | 79.5151641 | 1.00934051 | 0.2384843 | 4.23231435 | 2.31E-05 |
| 232879 | ZBTB45 | 525.779315 | 0.64873819 | 0.15338233 | 4.22954969 | 2.34E-05 |
| 14933 | gk | 1479.07739 | -0.6933527 | 0.16393696 | -4.2293858 | 2.34E-05 |

| | | | | | | |
|---|---|---|---|---|---|---|
| 22278 | USF1 | 634.709779 | -0.3111967 | 0.07362303 | -4.2268939 | 2.37E-05 |
| 67689 | ALDH3B1 | 102.759244 | -1.8005527 | 0.42603925 | -4.2262602 | 2.38E-05 |
| 252903 | AP1S3 | 155.823167 | -1.7537011 | 0.41506224 | -4.2251522 | 2.39E-05 |
| 231633 | TMEM119 | 194.809779 | 2.20987975 | 0.52305392 | 4.22495594 | 2.39E-05 |
| 209630 | FRMD4A | 2598.45953 | 0.84613848 | 0.20029917 | 4.22437347 | 2.40E-05 |
| 433702 | NCBP1 | 3769.83271 | 0.5261383 | 0.1245968 | 4.22272712 | 2.41E-05 |
| 236604 | Pisd-ps1 | 53.4602899 | -0.9210468 | 0.21813586 | -4.222354 | 2.42E-05 |
| 71735 | LRWD1 | 1071.5178 | 0.31055857 | 0.07359828 | 4.21964457 | 2.45E-05 |
| 216616 | EFEMP1 | 1369.17452 | 2.11991222 | 0.50245351 | 4.21912116 | 2.45E-05 |
| 229595 | ADAMTSL4 | 822.326937 | -1.066219 | 0.25283428 | -4.2170666 | 2.48E-05 |
| 13821 | EPB41L1 | 5102.77844 | 0.78128815 | 0.18529218 | 4.21651989 | 2.48E-05 |
| 74167 | NUDT9 | 1681.90862 | 0.71074134 | 0.16856177 | 4.21650368 | 2.48E-05 |
| 631323 | Gm12250 | 10.5266302 | -2.7349302 | 0.64893342 | -4.2145005 | 2.50E-05 |
| 23856 | DIDO1 | 3951.02016 | 0.36930878 | 0.08764612 | 4.21363507 | 2.51E-05 |
| 74918 | Iqca | 5.07325222 | -5.773875 | 1.37045138 | -4.2131192 | 2.52E-05 |
| 67035 | DNAJB4 | 1696.13563 | 0.50765234 | 0.12054524 | 4.21130127 | 2.54E-05 |
| 76969 | CHST1 | 52.6247878 | -3.1704378 | 0.75296631 | -4.2105972 | 2.55E-05 |
| 78266 | Zfp687 | 1570.62585 | -0.2941414 | 0.06986472 | -4.2101564 | 2.55E-05 |
| 319371 | D030028A08 | 118.401712 | -1.3461291 | 0.31986393 | -4.2084429 | 2.57E-05 |
| 27362 | DNAJB9 | 849.45063 | 0.67652131 | 0.16077248 | 4.20794222 | 2.58E-05 |
| 70248 | DAZAP1 | 4689.23912 | 0.38245878 | 0.09089101 | 4.2078834 | 2.58E-05 |
| 214642 | CPED1 | 249.432344 | 1.33398519 | 0.31703474 | 4.20769402 | 2.58E-05 |
| 17713 | GRPEL1 | 2131.35721 | -0.2588996 | 0.06154165 | -4.2069006 | 2.59E-05 |
| 66468 | SKA1 | 656.411153 | 0.51998921 | 0.12364839 | 4.20538602 | 2.61E-05 |
| 69310 | PACRG | 3.93982732 | -5.4121752 | 1.28705225 | -4.2050936 | 2.61E-05 |
| 218454 | LHFPL2 | 2100.32587 | -0.8804873 | 0.20940034 | -4.2048037 | 2.61E-05 |
| 100383 | BSDC1 | 533.754702 | 0.28414642 | 0.06758613 | 4.20421191 | 2.62E-05 |
| 16561 | KIF1B | 4992.12405 | 0.66093254 | 0.157224 | 4.20376363 | 2.63E-05 |
| 230935 | DNAJC11 | 2140.88241 | 0.51360876 | 0.12218527 | 4.20352429 | 2.63E-05 |
| 237362 | NPFFR1 | 9.07338759 | -6.6089055 | 1.57224389 | -4.2034862 | 2.63E-05 |
| 12617 | Cenpc1 | 953.254972 | -0.4387274 | 0.10438434 | -4.2030007 | 2.63E-05 |
| 56495 | ASNA1 | 2486.97644 | -0.364706 | 0.08684515 | -4.1994972 | 2.68E-05 |
| 14348 | FUT9 | 81.5620084 | -9.7797341 | 2.32885412 | -4.199376 | 2.68E-05 |
| 26940 | ECSIT | 475.908044 | 0.38841091 | 0.09250878 | 4.19863827 | 2.69E-05 |
| 70974 | PGM2L1 | 466.45789 | 0.90368268 | 0.21525891 | 4.19811964 | 2.69E-05 |
| 230908 | TARDBP | 7398.04195 | 0.47064119 | 0.11210819 | 4.19809839 | 2.69E-05 |
| 245865 | SPAG4 | 41.5203555 | 1.05375912 | 0.2510198 | 4.19791228 | 2.69E-05 |
| 20595 | SMN1 | 1003.98963 | 0.56138078 | 0.13383833 | 4.19446935 | 2.74E-05 |
| 217011 | NLE1 | 752.961794 | 0.32897336 | 0.0784476 | 4.19354294 | 2.75E-05 |
| 100900 | HSCB | 130.981873 | -0.635101 | 0.15145339 | -4.1933756 | 2.75E-05 |
| 18604 | PDK2 | 58.6130782 | -1.0673534 | 0.25457749 | -4.1926462 | 2.76E-05 |
| 15439 | HP | 7.82116723 | 3.27454621 | 0.78115765 | 4.19191467 | 2.77E-05 |
| 66089 | RMND5B | 779.503198 | -0.5171251 | 0.12346272 | -4.188512 | 2.81E-05 |
| 232023 | VOPP1 | 1855.59768 | -0.6102921 | 0.14571343 | -4.1883038 | 2.81E-05 |
| 66863 | LZTR1 | 2379.00672 | -0.2702473 | 0.06452524 | -4.1882423 | 2.81E-05 |

| | | | | | | |
|---|---|---|---|---|---|---|
| 116872 | SERPINB7 | 4.17508304 | -5.4958913 | 1.31223963 | -4.1881766 | 2.81E-05 |
| 16452 | JAK2 | 1078.91012 | -0.6667891 | 0.1592698 | -4.1865382 | 2.83E-05 |
| 14810 | GRIN1 | 7.91898393 | -4.9178758 | 1.17476815 | -4.1862522 | 2.84E-05 |
| 53379 | HNRNPA2B1 | 26074.582 | 0.38057718 | 0.09091654 | 4.18600589 | 2.84E-05 |
| 15353 | HMG20B | 1212.52224 | -0.4823651 | 0.11524105 | -4.1857058 | 2.84E-05 |
| 67973 | MPHOSPH10 | 1516.98239 | 0.57270121 | 0.13683902 | 4.18521852 | 2.85E-05 |
| 67755 | DDX47 | 1875.06662 | 0.3730693 | 0.08916619 | 4.1839774 | 2.86E-05 |
| 223697 | SUN2 | 4085.5352 | -0.3730429 | 0.08916533 | -4.183722 | 2.87E-05 |
| 67896 | CCDC80 | 174.801467 | 3.17975014 | 0.76045743 | 4.18136506 | 2.90E-05 |
| 103406 | ZFR2 | 14.7656906 | -2.6384671 | 0.631075 | -4.180909 | 2.90E-05 |
| 20503 | SLC16A7 | 17.8206893 | -7.5844871 | 1.81417985 | -4.1806699 | 2.91E-05 |
| 78552 | E130201H02 | 71.0544517 | 1.15988167 | 0.2774586 | 4.18037737 | 2.91E-05 |
| 381560 | XKR8 | 89.8918645 | -0.8629917 | 0.20645276 | -4.1800928 | 2.91E-05 |
| 14870 | GSTP1 | 1342.83831 | 0.36210643 | 0.08663136 | 4.17985364 | 2.92E-05 |
| 107338 | GBF1 | 3059.33673 | 0.25663084 | 0.06139904 | 4.17972055 | 2.92E-05 |
| 66592 | STOML2 | 1378.00509 | 0.4154611 | 0.09941953 | 4.17886812 | 2.93E-05 |
| 193813 | MCFD2 | 2334.36595 | -0.4820861 | 0.11541311 | -4.1770483 | 2.95E-05 |
| 71432 | 5430437J10F | 11.1317508 | 2.88543779 | 0.69082679 | 4.17678905 | 2.96E-05 |
| 74351 | DDX23 | 4709.19224 | 0.17511443 | 0.04192685 | 4.1766653 | 2.96E-05 |
| 20429 | SHOX2 | 223.047887 | 2.53157851 | 0.60625105 | 4.17579236 | 2.97E-05 |
| 19245 | PTP4A3 | 517.532514 | -1.0564447 | 0.2530201 | -4.1753389 | 2.98E-05 |
| 71807 | TARS2 | 1305.47426 | -0.5612773 | 0.13444043 | -4.1749147 | 2.98E-05 |
| 18628 | PER3 | 139.181312 | 1.02030495 | 0.24454206 | 4.1723086 | 3.02E-05 |
| 329369 | Snhg7os | 5.74112234 | -5.1902899 | 1.24408092 | -4.1719874 | 3.02E-05 |
| 18166 | NPY1R | 12.3391249 | -6.3182909 | 1.51449412 | -4.1718821 | 3.02E-05 |
| 22635 | ZAN | 66.1820028 | -2.0468719 | 0.49064925 | -4.1717621 | 3.02E-05 |
| 216739 | ACSL6 | 5.55609397 | -5.1598264 | 1.23693125 | -4.1714739 | 3.03E-05 |
| 67939 | Prorsd1 | 140.534665 | -0.5941249 | 0.14243554 | -4.1711845 | 3.03E-05 |
| 217125 | SAMD14 | 263.914808 | -1.6664838 | 0.39958603 | -4.1705257 | 3.04E-05 |
| 28071 | TWISTNB | 662.155722 | 0.68430486 | 0.16412178 | 4.1694946 | 3.05E-05 |
| 328232 | GFOD1 | 1200.24941 | 1.08210337 | 0.25963736 | 4.16774905 | 3.08E-05 |
| 217944 | RAPGEF5 | 27.6253091 | -5.1656111 | 1.23943208 | -4.1677243 | 3.08E-05 |
| 21766 | TEX261 | 2007.77043 | 0.20925457 | 0.05021851 | 4.16688123 | 3.09E-05 |
| 77627 | EFCAB6 | 4.33616013 | -5.5474749 | 1.33136345 | -4.1667622 | 3.09E-05 |
| 327958 | PITPNM3 | 8.83778219 | -5.8273936 | 1.39894047 | -4.1655765 | 3.11E-05 |
| 80289 | LYSMD3 | 862.361497 | 0.72825861 | 0.17483574 | 4.1653876 | 3.11E-05 |
| 67181 | CTDNEP1 | 2370.4048 | 0.28113917 | 0.06749682 | 4.16522125 | 3.11E-05 |
| 384185 | ARL9 | 10.8501549 | -2.2436368 | 0.5387944 | -4.1641799 | 3.12E-05 |
| 13609 | S1PR1 | 68.5916736 | 3.62595217 | 0.87094288 | 4.1632491 | 3.14E-05 |
| 14051 | EYA4 | 63.2970187 | -4.4656732 | 1.0731178 | -4.1614007 | 3.16E-05 |
| 71839 | OSGIN1 | 444.390627 | 0.75373414 | 0.18122192 | 4.15917751 | 3.19E-05 |
| 67763 | PRPSAP1 | 1946.45817 | 0.42947316 | 0.10329099 | 4.15789551 | 3.21E-05 |
| 83395 | SP6 | 8.59410875 | -2.9469425 | 0.70893135 | -4.1568799 | 3.23E-05 |
| 68087 | DCAKD | 1190.93226 | 0.31370585 | 0.07549647 | 4.15523859 | 3.25E-05 |
| 380773 | SLIRP | 946.723688 | 0.3932309 | 0.09466977 | 4.15371126 | 3.27E-05 |

| | | | | | | |
|---|---|---|---|---|---|---|
| 213409 | lemd1 | 5.41031135 | -5.8609344 | 1.41113304 | -4.1533535 | 3.28E-05 |
| 81601 | KAT5 | 1107.80238 | 0.29871703 | 0.07194651 | 4.1519322 | 3.30E-05 |
| 66676 | TMED7 | 6882.33523 | -0.2223392 | 0.05356049 | -4.1511796 | 3.31E-05 |
| 14417 | GAD2 | 7.16480662 | -4.7850367 | 1.15294688 | -4.1502664 | 3.32E-05 |
| 51793 | DDAH2 | 137.538545 | -1.4233465 | 0.34305712 | -4.1490074 | 3.34E-05 |
| 217310 | HID1 | 260.992082 | -2.5353267 | 0.61108392 | -4.1489009 | 3.34E-05 |
| 20536 | SLC4A3 | 189.523467 | -0.7562837 | 0.18237667 | -4.1468227 | 3.37E-05 |
| 109169 | IGIP | 545.463184 | -0.4483383 | 0.10814596 | -4.1456775 | 3.39E-05 |
| 54634 | MAGIX | 6.80632922 | -5.443009 | 1.3129798 | -4.145539 | 3.39E-05 |
| 218442 | SERINC5 | 778.944417 | -1.0269506 | 0.24774794 | -4.145143 | 3.40E-05 |
| 217664 | mgat2 | 4249.17073 | 0.5373301 | 0.12966445 | 4.14400465 | 3.41E-05 |
| 244653 | HYDIN | 13.9748274 | -4.7450861 | 1.14524303 | -4.1433006 | 3.42E-05 |
| 104175 | SBK1 | 226.014388 | -0.4427294 | 0.10685652 | -4.1432134 | 3.42E-05 |
| 242484 | D630039A03 | 4.22288567 | -5.5027474 | 1.32814167 | -4.1431931 | 3.43E-05 |
| 66931 | 1700010I14R | 7.15420892 | -6.2674906 | 1.51293651 | -4.1425999 | 3.43E-05 |
| 67683 | PBDC1 | 1226.20051 | -0.5063954 | 0.12224287 | -4.1425348 | 3.43E-05 |
| 12235 | BUB1 | 2913.11613 | 0.29286955 | 0.07070321 | 4.14223807 | 3.44E-05 |
| 97820 | 4833439L19F | 1673.18237 | 0.50966675 | 0.12304339 | 4.14217078 | 3.44E-05 |
| 68117 | APOOL | 297.768387 | -0.7302424 | 0.17635228 | -4.1408162 | 3.46E-05 |
| 228602 | 4930402H24 | 998.762918 | -0.4305553 | 0.10397946 | -4.1407725 | 3.46E-05 |
| 93883 | PCDHB12 | 12.0798939 | 2.79586491 | 0.67528874 | 4.14025105 | 3.47E-05 |
| 16409 | ITGAM | 11.2043288 | -4.8433179 | 1.16983799 | -4.1401613 | 3.47E-05 |
| 107993 | BFSP2 | 6.8548694 | -6.2035637 | 1.49842628 | -4.1400527 | 3.47E-05 |
| 67080 | 1700019D03 | 3.94532453 | -5.406254 | 1.30613839 | -4.1391127 | 3.49E-05 |
| 78455 | HELZ | 1537.83741 | -0.2094964 | 0.05063435 | -4.1374361 | 3.51E-05 |
| 665181 | Gm11964 | 245.531914 | -0.5976268 | 0.14445105 | -4.1372273 | 3.52E-05 |
| 74245 | CTBS | 413.454196 | -1.795443 | 0.43397548 | -4.1371992 | 3.52E-05 |
| 67242 | gemin6 | 395.753592 | 0.74325531 | 0.17967343 | 4.13670142 | 3.52E-05 |
| 192950 | NACAD | 5.30983256 | -5.0770647 | 1.227393 | -4.1364621 | 3.53E-05 |
| 100504720 | 2310002F09I | 7.96480892 | -4.9280288 | 1.19208812 | -4.1339467 | 3.57E-05 |
| 12476 | CD151 | 5780.65854 | -0.5437143 | 0.13157963 | -4.1322072 | 3.59E-05 |
| 18792 | PLAU | 732.628987 | -1.1753036 | 0.28447599 | -4.1314686 | 3.60E-05 |
| 240120 | Zfp119b | 96.1623725 | -0.7777413 | 0.18827279 | -4.1309279 | 3.61E-05 |
| 70808 | 4632415L05F | 350.018724 | -0.5759559 | 0.1394314 | -4.1307471 | 3.62E-05 |
| 319765 | IGF2BP2 | 3410.82653 | 0.56246188 | 0.13619756 | 4.12974999 | 3.63E-05 |
| 76707 | CLASP1 | 3592.4605 | -0.3394107 | 0.08220981 | -4.1285909 | 3.65E-05 |
| 217995 | HEATR1 | 4428.85289 | 0.71580278 | 0.17337818 | 4.12856335 | 3.65E-05 |
| 70099 | SMC4 | 8560.49936 | 0.41638845 | 0.10086468 | 4.12818875 | 3.66E-05 |
| 67664 | RNF125 | 87.1090878 | 1.82685728 | 0.44261348 | 4.12743253 | 3.67E-05 |
| 232853 | Zfp954 | 257.940295 | 0.61335681 | 0.14862897 | 4.12676487 | 3.68E-05 |
| 19221 | PTGFRN | 6440.31496 | -0.9782259 | 0.23705691 | -4.1265444 | 3.68E-05 |
| 99151 | CERCAM | 32.7817438 | -1.53593 | 0.37223505 | -4.1262369 | 3.69E-05 |
| 241568 | LRRC4C | 9.53376265 | -3.5450516 | 0.85932389 | -4.1253964 | 3.70E-05 |
| 667952 | Gm8894 | 213.752844 | -0.8870265 | 0.21510967 | -4.1236012 | 3.73E-05 |
| 100316685 | Mir1907 | 10.1110813 | 2.55667317 | 0.62006996 | 4.12320115 | 3.74E-05 |

| | | | | | | |
|---|---|---|---|---|---|---|
| 19094 | MAPK11 | 325.933384 | -0.7463649 | 0.18101679 | -4.1231805 | 3.74E-05 |
| 208449 | SGMS1 | 1270.11547 | -0.6947875 | 0.16851533 | -4.122993 | 3.74E-05 |
| 277463 | GPR107 | 1776.28665 | -0.5888334 | 0.14285534 | -4.1218857 | 3.76E-05 |
| 212772 | ARL14EP | 2327.4122 | -0.3424245 | 0.08307483 | -4.1218798 | 3.76E-05 |
| 68481 | MPZL1 | 2072.94735 | 0.69783871 | 0.16933127 | 4.12114489 | 3.77E-05 |
| 209773 | DENND2A | 306.444709 | 3.41455583 | 0.82857005 | 4.12102254 | 3.77E-05 |
| 69957 | CDC16 | 2287.40937 | 0.59646786 | 0.14473978 | 4.12096705 | 3.77E-05 |
| 432999 | A930007A09 | 66.7368414 | -1.2187141 | 0.29576246 | -4.1205841 | 3.78E-05 |
| 57440 | EHD3 | 132.040715 | -1.8305623 | 0.44446239 | -4.118599 | 3.81E-05 |
| 17826 | FAM89B | 1445.69116 | 0.83609365 | 0.20300941 | 4.11849701 | 3.81E-05 |
| 14070 | F8a | 221.673818 | -0.6076503 | 0.14761879 | -4.116348 | 3.85E-05 |
| 101565 | CCP110 | 1200.47437 | 0.61292099 | 0.14891482 | 4.11591654 | 3.86E-05 |
| 14077 | FABP3 | 9.93175182 | 3.92035278 | 0.95279149 | 4.11459675 | 3.88E-05 |
| 23964 | TENM2 | 4.0778935 | -5.4605311 | 1.32727388 | -4.1140952 | 3.89E-05 |
| 100503381 | Gm14022 | 8.87872302 | -2.7817378 | 0.67615592 | -4.1140478 | 3.89E-05 |
| 268663 | CDHR2 | 3.82829525 | -5.3621709 | 1.30343259 | -4.1138843 | 3.89E-05 |
| 67883 | UXS1 | 916.050629 | -0.2894976 | 0.0704178 | -4.1111426 | 3.94E-05 |
| 74665 | LRRC48 | 19.362577 | -2.2878147 | 0.55652603 | -4.1108854 | 3.94E-05 |
| 11305 | abca2 | 2036.32256 | -1.270975 | 0.30921573 | -4.1103179 | 3.95E-05 |
| 214469 | FAM168B | 7856.35952 | 0.15443611 | 0.03758867 | 4.10858104 | 3.98E-05 |
| 55927 | HES6 | 389.115854 | -0.9251517 | 0.22517726 | -4.1085485 | 3.98E-05 |
| 18029 | nfic | 10863.7577 | 0.39369823 | 0.09582702 | 4.10842624 | 3.98E-05 |
| 67884 | 1810043G02 | 339.20396 | -0.5015974 | 0.1221014 | -4.1080396 | 3.99E-05 |
| 27999 | FAM3C | 3600.4781 | -0.5237015 | 0.12749223 | -4.1077132 | 4.00E-05 |
| 72729 | CDC42SE2 | 2023.74264 | -0.4291176 | 0.10448151 | -4.1071152 | 4.01E-05 |
| 66143 | EEF1E1 | 1323.94432 | 0.65788658 | 0.16020237 | 4.10659707 | 4.02E-05 |
| 21853 | TIMELESS | 1282.21545 | 0.24107262 | 0.05870885 | 4.10624015 | 4.02E-05 |
| 382018 | UNC13A | 61.2116455 | -3.9586041 | 0.96431977 | -4.1050741 | 4.04E-05 |
| 66229 | RPL7L1 | 3373.56369 | 0.41019062 | 0.09993114 | 4.10473296 | 4.05E-05 |
| 270210 | Zfp651 | 509.267053 | 0.85617159 | 0.20859189 | 4.1045296 | 4.05E-05 |
| 59021 | RAB2A | 3944.99885 | -1.1391788 | 0.27763236 | -4.1031918 | 4.07E-05 |
| 73242 | ATAT1 | 100.701861 | -0.8912926 | 0.21723673 | -4.1028633 | 4.08E-05 |
| 230661 | TESK2 | 181.164521 | 0.64689544 | 0.15768212 | 4.10252887 | 4.09E-05 |
| 13644 | EFS | 9.31137431 | -6.651627 | 1.62154196 | -4.1020382 | 4.10E-05 |
| 12373 | CASQ2 | 17.5041013 | -3.0343169 | 0.73991184 | -4.1009168 | 4.12E-05 |
| 214897 | CSNK1G1 | 1816.1294 | -0.2419475 | 0.05900176 | -4.1006828 | 4.12E-05 |
| 59043 | WSB2 | 3131.01499 | 0.14544586 | 0.03546942 | 4.10059924 | 4.12E-05 |
| 72607 | USP13 | 5.2683186 | -5.8228984 | 1.42043956 | -4.0993637 | 4.14E-05 |
| 213556 | PLEKHH2 | 1771.74443 | 0.93282577 | 0.2275738 | 4.09900342 | 4.15E-05 |
| 16439 | ITPR2 | 442.473088 | -0.707282 | 0.17256985 | -4.098526 | 4.16E-05 |
| 100041948 | Gm9800 | 416.509825 | 0.5977998 | 0.14586092 | 4.09842345 | 4.16E-05 |
| 72948 | TPPP | 11.9938867 | -2.1028499 | 0.51325506 | -4.0970855 | 4.18E-05 |
| 235527 | PLSCR4 | 15.3254412 | 3.01983024 | 0.73708086 | 4.09701351 | 4.19E-05 |
| 623474 | RAD54B | 334.301139 | -0.9974095 | 0.24345133 | -4.0969565 | 4.19E-05 |
| 83997 | SLMAP | 4427.38918 | -0.2498277 | 0.06097992 | -4.0968846 | 4.19E-05 |

| | | | | | | |
|---|---|---|---|---|---|---|
| 12797 | CNN1 | 8.64803701 | -3.1370229 | 0.7657358 | -4.0967431 | 4.19E-05 |
| 17130 | SMAD6 | 767.633238 | 1.37174824 | 0.33486168 | 4.09646224 | 4.20E-05 |
| 217258 | Abca8a | 12.5354476 | 4.0463029 | 0.98830619 | 4.09417948 | 4.24E-05 |
| 76478 | HAUS8 | 594.673097 | 0.51863384 | 0.12668512 | 4.09388137 | 4.24E-05 |
| 268417 | Zkscan17 | 1900.93127 | 0.23840717 | 0.05824015 | 4.09351906 | 4.25E-05 |
| 66717 | CCDC96 | 47.6229253 | -0.9105446 | 0.22246451 | -4.0929881 | 4.26E-05 |
| 68188 | SYMPK | 4112.59073 | 0.484584 | 0.11840835 | 4.09248149 | 4.27E-05 |
| 103213 | TRAF3IP2 | 822.546354 | -0.4202506 | 0.10269744 | -4.0921231 | 4.27E-05 |
| 18439 | P2RX7 | 1183.87375 | -1.346019 | 0.32901108 | -4.0911055 | 4.29E-05 |
| 13984 | ESX1 | 9.56848828 | 2.7819106 | 0.68011373 | 4.09036089 | 4.31E-05 |
| 15531 | NDST1 | 5842.31188 | -0.3784088 | 0.09251531 | -4.0902296 | 4.31E-05 |
| 140577 | ANKRD6 | 53.4754506 | -2.4030854 | 0.58753507 | -4.090114 | 4.31E-05 |
| 26893 | COPS6 | 2513.93241 | -0.2604435 | 0.06368004 | -4.089876 | 4.32E-05 |
| 12877 | CPEB1 | 339.738659 | 0.63802264 | 0.15603169 | 4.08905801 | 4.33E-05 |
| 211151 | CHURC1 | 541.303659 | -0.4026962 | 0.09851799 | -4.0875401 | 4.36E-05 |
| 445007 | NUP85 | 3072.04734 | 0.30093946 | 0.07362809 | 4.08729108 | 4.36E-05 |
| 20463 | COX7A2L | 902.317723 | -0.4202737 | 0.10284069 | -4.0866483 | 4.38E-05 |
| 16168 | IL15 | 28.1883282 | -1.9135397 | 0.46825276 | -4.0865529 | 4.38E-05 |
| 244879 | NPAT | 1781.94002 | 0.23040927 | 0.05639914 | 4.08533322 | 4.40E-05 |
| 74174 | GTSF1 | 10.3849466 | -5.3466495 | 1.30875766 | -4.0852861 | 4.40E-05 |
| 231842 | AMZ1 | 36.1664223 | -1.980413 | 0.4847892 | -4.0851013 | 4.41E-05 |
| 23834 | CDC6 | 1070.54732 | 0.4980527 | 0.12194323 | 4.08429979 | 4.42E-05 |
| 56449 | YBX3 | 17712.314 | 0.64911897 | 0.15896812 | 4.08332789 | 4.44E-05 |
| 67667 | ALKBH8 | 492.601607 | 0.63241391 | 0.15489312 | 4.08290508 | 4.45E-05 |
| 192188 | STAB2 | 7.38901769 | -6.3121268 | 1.54599867 | -4.0828799 | 4.45E-05 |
| 208366 | RPP40 | 252.116275 | 0.77118064 | 0.18892948 | 4.08184395 | 4.47E-05 |
| 11565 | ADSSL1 | 176.982628 | -2.1188499 | 0.51918579 | -4.0811015 | 4.48E-05 |
| 269639 | Zfp512 | 921.816903 | -0.4607608 | 0.11290392 | -4.0809992 | 4.48E-05 |
| 21928 | TNFAIP2 | 7178.18236 | 1.98250499 | 0.4858095 | 4.08082797 | 4.49E-05 |
| 23963 | TENM1 | 6.32914397 | -6.090041 | 1.49254708 | -4.0803008 | 4.50E-05 |
| 14702 | GNG2 | 423.956734 | 2.66527603 | 0.65325999 | 4.07996215 | 4.50E-05 |
| 66367 | 2310022A10 | 852.429821 | 0.42756127 | 0.10479758 | 4.07987718 | 4.51E-05 |
| 15402 | HOXA5 | 318.379741 | -0.4684698 | 0.11484142 | -4.0792758 | 4.52E-05 |
| 14017 | EVI2A | 247.085182 | 2.662715 | 0.65301246 | 4.07758681 | 4.55E-05 |
| 381629 | ATRAID | 832.442946 | -0.3072672 | 0.07535828 | -4.0774181 | 4.55E-05 |
| 381650 | THAP6 | 208.845482 | -0.6821776 | 0.1673425 | -4.0765351 | 4.57E-05 |
| 104110 | ADCY4 | 4.9392693 | 5.03640614 | 1.23552622 | 4.07632478 | 4.58E-05 |
| 19183 | PSMC3IP | 356.926048 | -0.380625 | 0.0933785 | -4.0761521 | 4.58E-05 |
| 626067 | Gm6650 | 7.05646524 | -3.7166589 | 0.91193642 | -4.0755681 | 4.59E-05 |
| 236733 | USP11 | 175.409122 | 1.45565358 | 0.35716791 | 4.07554417 | 4.59E-05 |
| 21822 | Tgtp1 | 3.58346849 | -5.2754244 | 1.2944758 | -4.0753365 | 4.59E-05 |
| 328162 | TRMT61A | 882.042486 | 0.52579066 | 0.12904589 | 4.07444717 | 4.61E-05 |
| 17261 | MEF2D | 1859.71616 | 0.4446588 | 0.10913754 | 4.07429726 | 4.62E-05 |
| 53817 | DDX39B | 5277.9091 | 0.2600563 | 0.06383071 | 4.07415665 | 4.62E-05 |
| 14422 | B4GALNT2 | 7.0262881 | -5.4862846 | 1.3466764 | -4.0739442 | 4.62E-05 |

| | | | | | | |
|---|---|---|---|---|---|---|
| 217337 | SRP68 | 2833.4343 | 0.27264702 | 0.06692776 | 4.07375105 | 4.63E-05 |
| 210094 | IGLON5 | 11.255026 | -3.1624106 | 0.77630973 | -4.0736455 | 4.63E-05 |
| 56310 | GPS2 | 486.554111 | -0.3156945 | 0.07751865 | -4.0724979 | 4.65E-05 |
| 70568 | CPNE3 | 2661.04593 | -1.0457401 | 0.25678167 | -4.0724875 | 4.65E-05 |
| 269800 | Zfp384 | 1590.04831 | 0.5555347 | 0.13642557 | 4.07207161 | 4.66E-05 |
| 235315 | RNF214 | 783.590142 | 0.34351912 | 0.08439829 | 4.07021429 | 4.70E-05 |
| 11876 | ARTN | 84.421762 | -4.9899521 | 1.22670044 | -4.0677838 | 4.75E-05 |
| 69792 | MED6 | 781.592514 | 0.45691439 | 0.11233354 | 4.06747961 | 4.75E-05 |
| 72750 | FAM117B | 1905.96951 | -0.5212965 | 0.12821534 | -4.0657886 | 4.79E-05 |
| 17751 | MT3 | 20.2619708 | -7.0412168 | 1.7319078 | -4.0655841 | 4.79E-05 |
| 170707 | USP48 | 1913.46521 | 0.54010618 | 0.13284948 | 4.06554993 | 4.79E-05 |
| 14411 | SLC6A12 | 77.6332106 | 1.33404911 | 0.32821098 | 4.06460848 | 4.81E-05 |
| 14168 | FGF13 | 12.5508052 | -7.0785786 | 1.74191584 | -4.0636743 | 4.83E-05 |
| 114663 | IMPA2 | 488.758785 | 0.44378101 | 0.10921378 | 4.06341593 | 4.84E-05 |
| 52585 | DHRS1 | 1296.78696 | -0.9638798 | 0.23721199 | -4.0633691 | 4.84E-05 |
| 330474 | ZC3H4 | 3478.11695 | 0.55146285 | 0.13571859 | 4.06328167 | 4.84E-05 |
| 269959 | ADAMTSL3 | 185.580577 | -2.440698 | 0.60087674 | -4.0618947 | 4.87E-05 |
| 381399 | BPIFB4 | 174.925724 | -3.0108811 | 0.74133449 | -4.061434 | 4.88E-05 |
| 51796 | SRRM1 | 5247.07252 | 0.54832606 | 0.13502747 | 4.06084821 | 4.89E-05 |
| 100042807 | Eif3j2 | 725.867486 | 0.3638886 | 0.08961585 | 4.06053838 | 4.90E-05 |
| 69702 | NDUFAF1 | 258.768611 | -0.4685644 | 0.11541393 | -4.0598601 | 4.91E-05 |
| 24014 | RNASEL | 32.4622789 | -1.2844469 | 0.31646978 | -4.0586717 | 4.94E-05 |
| 209584 | TYW3 | 150.494532 | 0.60263524 | 0.1485276 | 4.05739554 | 4.96E-05 |
| 73683 | atg16l2 | 255.168721 | 0.55826896 | 0.13762109 | 4.05656559 | 4.98E-05 |
| 667214 | 9930111J21F | 20.9491633 | 2.73085806 | 0.67320576 | 4.05649836 | 4.98E-05 |
| 27384 | Akr1c13 | 13.5963509 | -3.4312075 | 0.84595487 | -4.0560172 | 4.99E-05 |
| 224826 | UBR2 | 4231.70965 | 0.35144315 | 0.08664802 | 4.05598589 | 4.99E-05 |
| 212531 | SH3BGRL2 | 1224.11973 | -0.5523952 | 0.13621619 | -4.0552829 | 5.01E-05 |
| 232664 | CCDC136 | 159.006392 | -1.1668904 | 0.28779958 | -4.0545245 | 5.02E-05 |
| 17309 | MGAT3 | 52.7231114 | -4.1461964 | 1.02315328 | -4.0523707 | 5.07E-05 |
| 56791 | UBE2L6 | 200.667245 | -1.1960985 | 0.29523963 | -4.0512802 | 5.09E-05 |
| 110593 | PRDM2 | 1948.024 | 0.58912182 | 0.14544848 | 4.05038145 | 5.11E-05 |
| 26992 | BRD7 | 1604.35978 | 0.23643949 | 0.0583807 | 4.04995955 | 5.12E-05 |
| 83984 | TSSK6 | 15.4308756 | 3.10113874 | 0.76578008 | 4.04964664 | 5.13E-05 |
| 71406 | 5430416O09 | 7.36449386 | -3.7791994 | 0.93328385 | -4.0493568 | 5.14E-05 |
| 213539 | BAG2 | 1189.37254 | 0.54033099 | 0.13345222 | 4.04887224 | 5.15E-05 |
| 21843 | TIAL1 | 2494.55209 | 0.73523808 | 0.18179772 | 4.04426466 | 5.25E-05 |
| 69737 | TTL | 1039.84048 | -0.4170767 | 0.10314055 | -4.0437702 | 5.26E-05 |
| 57757 | PGLYRP2 | 5.08324053 | -5.7695706 | 1.42686713 | -4.0435234 | 5.27E-05 |
| 216440 | OS9 | 3036.6715 | -0.561972 | 0.13898621 | -4.0433649 | 5.27E-05 |
| 229782 | SLC35A3 | 1938.93079 | -0.3138836 | 0.0776364 | -4.0429947 | 5.28E-05 |
| 329178 | UNC80 | 7.22715146 | -4.7942522 | 1.18595924 | -4.0425101 | 5.29E-05 |
| 239364 | TSPYL5 | 16.4693594 | -2.2339336 | 0.55262775 | -4.042384 | 5.29E-05 |
| 224170 | DZIP3 | 895.827822 | -0.5227134 | 0.12934995 | -4.0410797 | 5.32E-05 |
| 215653 | RASSF2 | 82.7749216 | 1.42564494 | 0.35288435 | 4.03997785 | 5.35E-05 |

| | | | | | | |
|---|---|---|---|---|---|---|
| 94066 | mrpl36 | 548.479717 | 0.7386386 | 0.18283368 | 4.03994824 | 5.35E-05 |
| 74007 | BTBD11 | 498.041922 | -0.9575033 | 0.23706656 | -4.0389641 | 5.37E-05 |
| 226778 | mark1 | 321.767704 | 2.71175471 | 0.6715379 | 4.03812606 | 5.39E-05 |
| 100418235 | C130030K03I | 3.49194792 | -5.2357545 | 1.29682982 | -4.0373489 | 5.41E-05 |
| 15183 | HDAC3 | 2576.69249 | -0.2516784 | 0.0623442 | -4.0369184 | 5.42E-05 |
| 236643 | SYTL5 | 3.5989837 | -5.280405 | 1.30807211 | -4.0367844 | 5.42E-05 |
| 382073 | CCDC84 | 140.703208 | -0.7504683 | 0.18593332 | -4.0362227 | 5.43E-05 |
| 546049 | C330021F23I | 5.75244684 | -5.9598126 | 1.47674177 | -4.0357852 | 5.44E-05 |
| 11539 | ADORA1 | 18.4322971 | -2.0469053 | 0.50725652 | -4.035247 | 5.45E-05 |
| 70821 | 4921507P07I | 11.8182153 | -4.4785821 | 1.11047731 | -4.0330244 | 5.51E-05 |
| 19016 | PPARG | 71.5672912 | -3.0074918 | 0.7458248 | -4.0324374 | 5.52E-05 |
| 108077 | SKIV2L | 2330.6631 | -0.2460732 | 0.06103583 | -4.0316185 | 5.54E-05 |
| 27096 | TRAPPC3 | 1447.39751 | 0.38155488 | 0.09466186 | 4.03071412 | 5.56E-05 |
| 433224 | Gm5512 | 85.8651924 | -0.8272368 | 0.20523676 | -4.0306462 | 5.56E-05 |
| 268281 | SHPRH | 1193.4819 | -0.321334 | 0.07972679 | -4.0304388 | 5.57E-05 |
| 13012 | CST8 | 3.77578617 | -5.346008 | 1.3264294 | -4.030375 | 5.57E-05 |
| 68073 | FAM173B | 389.223658 | 0.59332205 | 0.14721918 | 4.03019541 | 5.57E-05 |
| 68938 | ASPSCR1 | 1478.01501 | -0.3571048 | 0.08861443 | -4.0298715 | 5.58E-05 |
| 210035 | NEMP1 | 707.334707 | -0.5595053 | 0.13884577 | -4.0296891 | 5.59E-05 |
| 56613 | RPS6KA4 | 4953.74624 | 0.59184143 | 0.14688164 | 4.02937667 | 5.59E-05 |
| 320472 | PPM1E | 74.4632526 | -3.4688082 | 0.86136729 | -4.0270953 | 5.65E-05 |
| 16413 | ITGB1BP1 | 1006.55472 | 0.7214825 | 0.17917423 | 4.02670897 | 5.66E-05 |
| 66043 | ATP5D | 4254.87671 | -0.2971405 | 0.07379987 | -4.026301 | 5.67E-05 |
| 16515 | KCNJ12 | 46.8554276 | -5.1954256 | 1.29085748 | -4.0247863 | 5.70E-05 |
| 59010 | SQRDL | 1295.57273 | -1.163472 | 0.28913695 | -4.0239477 | 5.72E-05 |
| 234664 | NAE1 | 1422.36765 | -0.3472156 | 0.08628835 | -4.0238987 | 5.72E-05 |
| 14765 | GPR50 | 50.2934919 | -8.3569331 | 2.07754711 | -4.0224999 | 5.76E-05 |
| 71304 | Wbscr25 | 3.71630749 | -5.3262555 | 1.3241177 | -4.022494 | 5.76E-05 |
| 50762 | FBXO6 | 590.579929 | 0.4594321 | 0.11422691 | 4.02210033 | 5.77E-05 |
| 212516 | EFCAB12 | 4.9869449 | -5.7541835 | 1.43064292 | -4.0220963 | 5.77E-05 |
| 74600 | MRPL47 | 457.174436 | 0.41443722 | 0.1030451 | 4.02190145 | 5.77E-05 |
| 13590 | lefty1 | 11.9908201 | -5.5304894 | 1.37537194 | -4.0210864 | 5.79E-05 |
| 100040018 | Gm9835 | 6.86829694 | -5.4654426 | 1.35923498 | -4.0209697 | 5.80E-05 |
| 66286 | SEC11C | 1466.56916 | -0.6586742 | 0.16383348 | -4.0203883 | 5.81E-05 |
| 14886 | GTF2I | 5365.38751 | 0.39460212 | 0.09816057 | 4.01996572 | 5.82E-05 |
| 238831 | ppwd1 | 687.885998 | 0.65452803 | 0.16285384 | 4.01911333 | 5.84E-05 |
| 13401 | DMWD | 1300.03388 | 0.47647504 | 0.11859689 | 4.01760156 | 5.88E-05 |
| 12140 | FABP7 | 6.94582462 | 4.165772 | 1.0372834 | 4.01604038 | 5.92E-05 |
| 12503 | CD247 | 4.64481927 | -4.8804973 | 1.21546497 | -4.0153336 | 5.94E-05 |
| 74591 | ABCA12 | 4.54804233 | -5.6085711 | 1.39707459 | -4.0145109 | 5.96E-05 |
| 67727 | STX17 | 931.114285 | -0.6087975 | 0.15165727 | -4.0142984 | 5.96E-05 |
| 78783 | BRPF1 | 1690.44135 | 0.42677671 | 0.10633532 | 4.01349922 | 5.98E-05 |
| 13494 | DRG1 | 2728.6338 | -0.2675721 | 0.06668808 | -4.0122926 | 6.01E-05 |
| 59049 | SLC22A17 | 13.8198258 | -5.7690474 | 1.43823153 | -4.0112091 | 6.04E-05 |
| 20901 | STRAP | 7843.06466 | 0.32040047 | 0.07987964 | 4.0110405 | 6.05E-05 |

| | | | | | | |
|---|---|---|---|---|---|---|
| 21951 | TNKS | 3816.08445 | 0.46074653 | 0.11487891 | 4.01071484 | 6.05E-05 |
| 100504425 | Gm20219 | 48.0261592 | -1.4099753 | 0.35155629 | -4.0106671 | 6.05E-05 |
| 69354 | SLC38A4 | 7.95347895 | 4.37452718 | 1.09072629 | 4.01065529 | 6.06E-05 |
| 54683 | PRDX5 | 1851.50223 | -0.6803885 | 0.16968077 | -4.0098151 | 6.08E-05 |
| 232533 | STK38L | 723.134249 | 0.64886189 | 0.16183372 | 4.0094356 | 6.09E-05 |
| 16180 | IL1RAP | 2963.40592 | -0.3915387 | 0.09765885 | -4.0092498 | 6.09E-05 |
| 20637 | SNRNP70 | 7229.83875 | 0.43851584 | 0.10937796 | 4.00917935 | 6.09E-05 |
| 50915 | GRB14 | 1075.72861 | 0.72842447 | 0.18171786 | 4.00854636 | 6.11E-05 |
| 27762 | VWA7 | 12.8895911 | -2.8037322 | 0.69951761 | -4.0080939 | 6.12E-05 |
| 11777 | AP3S1 | 511.42801 | -0.4478012 | 0.11173565 | -4.0076842 | 6.13E-05 |
| 18798 | PLCB4 | 188.721136 | -1.7791193 | 0.44400218 | -4.0070057 | 6.15E-05 |
| 17388 | MMP15 | 62.8376389 | -2.1715843 | 0.54219417 | -4.0051783 | 6.20E-05 |
| 56040 | RPLP1 | 8284.31809 | 0.28878602 | 0.07211945 | 4.00427378 | 6.22E-05 |
| 17254 | SLC3A2 | 7841.24242 | 0.18688154 | 0.04667141 | 4.00419713 | 6.22E-05 |
| 15163 | HCLS1 | 23.0428161 | -7.9569891 | 1.98776642 | -4.0029799 | 6.25E-05 |
| 70699 | NUP205 | 7104.54804 | 0.38233943 | 0.09552198 | 4.00263318 | 6.26E-05 |
| 72306 | Zfp777 | 983.798218 | 0.25965297 | 0.06487134 | 4.00258367 | 6.27E-05 |
| 234549 | HEATR3 | 2981.30555 | 0.3683857 | 0.09204169 | 4.00237882 | 6.27E-05 |
| 245474 | DKC1 | 3341.29838 | 0.51716829 | 0.12923841 | 4.001661 | 6.29E-05 |
| 223732 | LDOC1L | 18.2229362 | -7.6188251 | 1.90470846 | -3.9999954 | 6.33E-05 |
| 266690 | CYB5R4 | 307.331124 | 0.3685608 | 0.09214738 | 3.99968832 | 6.34E-05 |
| 18518 | IGBP1 | 1055.46789 | -0.2193776 | 0.05487628 | -3.9976765 | 6.40E-05 |
| 215085 | SLC35F1 | 80.4388255 | -2.1140023 | 0.52888467 | -3.997095 | 6.41E-05 |
| 27219 | SGK2 | 14.359838 | -3.0856869 | 0.77203713 | -3.9968115 | 6.42E-05 |
| 15900 | IRF8 | 53.8922753 | -3.28128 | 0.82123046 | -3.9955654 | 6.45E-05 |
| 74455 | NSUN6 | 161.149365 | -0.4906176 | 0.12279294 | -3.9954873 | 6.46E-05 |
| 104910 | SLC25A47 | 81.2443891 | 1.72678668 | 0.43235169 | 3.99393989 | 6.50E-05 |
| 66962 | SWSAP1 | 111.252099 | 0.58122406 | 0.14552946 | 3.99385848 | 6.50E-05 |
| 21936 | TNFRSF18 | 5.50649478 | -5.134037 | 1.2856626 | -3.9933004 | 6.52E-05 |
| 102502 | PLS1 | 62.8190816 | -1.5787374 | 0.39537421 | -3.9930206 | 6.52E-05 |
| 93878 | PCDHB7 | 24.3468326 | 2.50070491 | 0.62636647 | 3.99239908 | 6.54E-05 |
| 68497 | AREL1 | 2408.96985 | -0.2856992 | 0.07156547 | -3.992137 | 6.55E-05 |
| 14533 | BLOC1S1 | 374.596386 | -0.6607296 | 0.16564804 | -3.9887557 | 6.64E-05 |
| 66531 | CMC2 | 429.899034 | 0.55127751 | 0.13823711 | 3.98791249 | 6.67E-05 |
| 12447 | CCNE1 | 1150.09538 | 0.73043593 | 0.18322942 | 3.98645551 | 6.71E-05 |
| 67552 | H2afy3 | 28.0402509 | 1.37102716 | 0.34402469 | 3.98525804 | 6.74E-05 |
| 320453 | Dlx4os | 6.08384218 | -6.0400656 | 1.51569923 | -3.9850028 | 6.75E-05 |
| 619331 | Zfp551 | 47.933127 | -0.8734096 | 0.21918749 | -3.9847602 | 6.75E-05 |
| 216151 | POLRMT | 1226.43042 | 0.32283544 | 0.08103187 | 3.98405498 | 6.77E-05 |
| 100503578 | #N/A | 46.6288279 | -1.0750137 | 0.26984741 | -3.9837836 | 6.78E-05 |
| 227699 | NUP188 | 4279.22866 | 0.2838808 | 0.07127378 | 3.9829627 | 6.81E-05 |
| 66861 | DNAJC10 | 6623.6568 | 0.21212116 | 0.05326425 | 3.98243009 | 6.82E-05 |
| 69504 | Zfp932 | 18.4039808 | -1.519 | 0.38143251 | -3.9823558 | 6.82E-05 |
| 75964 | TRAPPC8 | 2951.34268 | -0.3123742 | 0.07844194 | -3.9822347 | 6.83E-05 |
| 74257 | TSPAN17 | 370.854898 | 2.53113173 | 0.63566446 | 3.98186765 | 6.84E-05 |

| | | | | | | |
|---|---|---|---|---|---|---|
| 319314 | A930001C03 | 43.275469 | -1.8640369 | 0.46815724 | -3.981647 | 6.84E-05 |
| 76965 | SLITRK1 | 5.92902252 | -5.2396413 | 1.31601966 | -3.9814309 | 6.85E-05 |
| 268527 | GREB1 | 71.5750704 | -2.26395 | 0.56872257 | -3.9807633 | 6.87E-05 |
| 209334 | GEN1 | 1129.57727 | 0.50996481 | 0.12812378 | 3.98025099 | 6.88E-05 |
| 67602 | NECAP1 | 1374.39528 | 0.47347445 | 0.11896087 | 3.98008574 | 6.89E-05 |
| 207304 | hectd1 | 11933.783 | 0.51514971 | 0.1294574 | 3.97929894 | 6.91E-05 |
| 68942 | CHMP2B | 2686.65293 | -0.3277918 | 0.08239258 | -3.9784136 | 6.94E-05 |
| 269585 | ZSCAN20 | 159.809854 | 0.89890432 | 0.2259604 | 3.97814988 | 6.95E-05 |
| 20692 | SPARC | 12720.3828 | 0.67906521 | 0.1707195 | 3.97766632 | 6.96E-05 |
| 238673 | Zfp367 | 1652.38374 | 0.53212784 | 0.13378558 | 3.97746794 | 6.97E-05 |
| 635702 | NAALADL2 | 30.5967661 | -2.4248025 | 0.609668 | -3.9772507 | 6.97E-05 |
| 230234 | FAM206A | 1160.99068 | 0.59146276 | 0.1487352 | 3.97661579 | 6.99E-05 |
| 207958 | ALG11 | 850.999135 | -0.4083078 | 0.10268943 | -3.9761427 | 7.00E-05 |
| 53310 | DLG3 | 788.090038 | -0.779288 | 0.19604618 | -3.9750227 | 7.04E-05 |
| 237928 | phospho1 | 291.057245 | -1.7934727 | 0.45121592 | -3.9747548 | 7.05E-05 |
| 100504375 | Gm15880 | 5.26840082 | 5.12332634 | 1.28939645 | 3.97342984 | 7.08E-05 |
| 211922 | DENND6A | 936.720594 | -0.2924843 | 0.07361196 | -3.9733254 | 7.09E-05 |
| 73261 | 1700037C18I | 40.3821128 | -0.9415399 | 0.23699284 | -3.9728622 | 7.10E-05 |
| 26439 | Psg19 | 4.95431348 | -4.9734671 | 1.25283194 | -3.9697799 | 7.19E-05 |
| 14219 | CTGF | 21499.1019 | -1.967505 | 0.49567069 | -3.9693795 | 7.21E-05 |
| 233781 | XYLT1 | 221.219777 | 3.03002472 | 0.76344566 | 3.96888067 | 7.22E-05 |
| 18032 | NFIX | 5548.8846 | 0.61867902 | 0.15590499 | 3.96830803 | 7.24E-05 |
| 66073 | TXNDC12 | 1280.11196 | 0.66534565 | 0.16767189 | 3.96814079 | 7.24E-05 |
| 218461 | PDE8B | 8.42010282 | 2.74290287 | 0.69141634 | 3.96707845 | 7.28E-05 |
| 20855 | STC1 | 18.5968956 | 4.38328206 | 1.10503765 | 3.96663594 | 7.29E-05 |
| 100494 | ZFAND2A | 4466.6297 | -1.0417537 | 0.26264382 | -3.9664125 | 7.30E-05 |
| 71881 | APMAP | 971.396143 | -0.2556366 | 0.06446124 | -3.9657406 | 7.32E-05 |
| 12858 | Cox5a | 2930.43393 | 0.21681248 | 0.05468833 | 3.96451096 | 7.35E-05 |
| 56542 | ICK | 621.390901 | 0.34687065 | 0.08750412 | 3.96404957 | 7.37E-05 |
| 70153 | 2210016F16I | 757.224233 | 1.81526288 | 0.45808273 | 3.96274026 | 7.41E-05 |
| 632687 | 41707 | 11.072292 | 2.57459264 | 0.64970009 | 3.96274014 | 7.41E-05 |
| 13631 | EEF2K | 918.98477 | 0.45353593 | 0.1144517 | 3.9626842 | 7.41E-05 |
| 17423 | NDST2 | 792.680047 | 0.48463317 | 0.12231176 | 3.96227784 | 7.42E-05 |
| 227327 | B3GNT7 | 18.7463301 | -2.2370264 | 0.5646092 | -3.9620793 | 7.43E-05 |
| 230824 | GRHL3 | 5.27367393 | -5.0688714 | 1.27936542 | -3.96202 | 7.43E-05 |
| 14275 | folr1 | 13.607395 | -6.4585137 | 1.63013875 | -3.9619411 | 7.43E-05 |
| 54667 | ATP8B2 | 3015.1202 | 1.79992838 | 0.4547092 | 3.95841643 | 7.54E-05 |
| 18190 | NRXN2 | 9.18107378 | 3.55475703 | 0.89803368 | 3.9583783 | 7.55E-05 |
| 77428 | 9430083A17 | 6.40194463 | -4.0158382 | 1.01454005 | -3.9582846 | 7.55E-05 |
| 75212 | RNF121 | 657.598551 | 0.49173147 | 0.12424337 | 3.9578086 | 7.56E-05 |
| 16329 | INPP1 | 312.47899 | -0.3789418 | 0.09575004 | -3.9576146 | 7.57E-05 |
| 24060 | slc35a1 | 1016.78737 | -0.8384651 | 0.21186285 | -3.9575844 | 7.57E-05 |
| 68617 | MTCL1 | 256.901402 | 0.94829281 | 0.23961559 | 3.95755897 | 7.57E-05 |
| 243277 | ADGRD1 | 3.95666246 | -5.4105473 | 1.36777494 | -3.9557292 | 7.63E-05 |
| 11946 | ATP5A1 | 20880.9003 | 0.19380622 | 0.04900118 | 3.95513362 | 7.65E-05 |

| | | | | | | |
|---|---|---|---|---|---|---|
| 104252 | CDC42EP2 | 174.056195 | 1.19001146 | 0.30088323 | 3.95506081 | 7.65E-05 |
| 56150 | MAD2L1 | 2754.36799 | 0.67503083 | 0.17071165 | 3.95421662 | 7.68E-05 |
| 68852 | LRRN4CL | 54.7156972 | 0.92827017 | 0.23475626 | 3.95418711 | 7.68E-05 |
| 22654 | Zfp13 | 85.8911371 | -0.986852 | 0.24965408 | -3.9528776 | 7.72E-05 |
| 53382 | TXNL1 | 5080.74462 | -0.1341098 | 0.03393548 | -3.9519056 | 7.75E-05 |
| 70999 | NAA40 | 1658.60041 | -0.3748303 | 0.09485257 | -3.9517149 | 7.76E-05 |
| 56350 | ARL3 | 436.853873 | -0.4928784 | 0.12472716 | -3.9516525 | 7.76E-05 |
| 19367 | RAD9A | 1327.70948 | -0.4864025 | 0.12313535 | -3.9501448 | 7.81E-05 |
| 109552 | SRI | 1999.53279 | 0.46041911 | 0.11657566 | 3.94953046 | 7.83E-05 |
| 74513 | NETO2 | 459.157853 | 1.59628355 | 0.4041776 | 3.94946074 | 7.83E-05 |
| 14081 | ACSL1 | 760.529073 | 0.72746991 | 0.18426355 | 3.94798595 | 7.88E-05 |
| 16841 | LECT2 | 12.0681264 | -2.7642968 | 0.70067653 | -3.9451825 | 7.97E-05 |
| 14115 | FBLN2 | 44812.5821 | -0.8779913 | 0.22265522 | -3.9432775 | 8.04E-05 |
| 74134 | CYP2S1 | 11.7059327 | -3.0514202 | 0.77390515 | -3.9428865 | 8.05E-05 |
| 207182 | GGT7 | 28.6129108 | -3.2280985 | 0.81883914 | -3.9422865 | 8.07E-05 |
| 73086 | RPS6KA5 | 325.824088 | 0.79507721 | 0.20169896 | 3.94190031 | 8.08E-05 |
| 94227 | PI15 | 6.19255337 | -4.5534344 | 1.15543905 | -3.9408694 | 8.12E-05 |
| 14732 | GPAM | 650.437681 | 0.46279346 | 0.11748472 | 3.93918 | 8.18E-05 |
| 17342 | MITF | 412.684344 | 0.8929131 | 0.22668598 | 3.93898684 | 8.18E-05 |
| 219114 | SKA3 | 1053.68413 | 0.86255694 | 0.21906507 | 3.93744625 | 8.24E-05 |
| 21833 | THRA | 413.596645 | 1.0981601 | 0.27890569 | 3.93738871 | 8.24E-05 |
| 14773 | GRK5 | 532.518747 | 0.75752365 | 0.19245345 | 3.93613957 | 8.28E-05 |
| 100043272 | Inafm2 | 1331.42612 | -0.645044 | 0.16388678 | -3.9359125 | 8.29E-05 |
| 72661 | serp2 | 93.769045 | 1.6502405 | 0.41939488 | 3.93481316 | 8.33E-05 |
| 99899 | IFI44 | 92.2475928 | -3.6348133 | 0.92378856 | -3.934681 | 8.33E-05 |
| 320615 | DOPEY1 | 1093.26981 | 0.28065199 | 0.07140326 | 3.93052072 | 8.48E-05 |
| 100039304 | Gm2148 | 56.0143078 | -0.931013 | 0.23687835 | -3.9303423 | 8.48E-05 |
| 73658 | SPNS1 | 1377.41629 | -0.3494469 | 0.08891495 | -3.9301248 | 8.49E-05 |
| 231148 | ABLIM2 | 3.54040768 | -5.2605774 | 1.33869436 | -3.9296329 | 8.51E-05 |
| 320394 | CENPT | 832.11983 | -0.5171403 | 0.13160527 | -3.9294802 | 8.51E-05 |
| 100216533 | Snord55 | 17.7356324 | 1.55933543 | 0.39683631 | 3.92941718 | 8.52E-05 |
| 68479 | PHF5A | 1188.22891 | 0.48034768 | 0.12225443 | 3.92908198 | 8.53E-05 |
| 23988 | PIN1 | 1509.61871 | 0.34418871 | 0.0876121 | 3.92855207 | 8.55E-05 |
| 19142 | PRSS12 | 350.841439 | 1.5282434 | 0.38901948 | 3.92844956 | 8.55E-05 |
| 68567 | CGREF1 | 509.602295 | 0.69841888 | 0.177841 | 3.9272095 | 8.59E-05 |
| 208158 | MAP6D1 | 10.7470588 | 2.45499414 | 0.62512469 | 3.92720711 | 8.59E-05 |
| 50849 | RNF10 | 9140.43777 | 0.17520508 | 0.04462181 | 3.92644482 | 8.62E-05 |
| 208982 | HMGCLL1 | 16.5866995 | 2.22976201 | 0.56794638 | 3.92600797 | 8.64E-05 |
| 17182 | MATN3 | 33.3677561 | -2.4717938 | 0.62960265 | -3.9259583 | 8.64E-05 |
| 64659 | MRPS14 | 834.145772 | -0.3802675 | 0.09686033 | -3.9259361 | 8.64E-05 |
| 21781 | TFDP1 | 2621.75799 | 0.54786392 | 0.13955406 | 3.92581851 | 8.64E-05 |
| 68876 | xrcc6bp1 | 171.47442 | 0.62421033 | 0.15904057 | 3.92484961 | 8.68E-05 |
| 17965 | nbl1 | 834.321107 | 0.62845669 | 0.16018811 | 3.9232418 | 8.74E-05 |
| 11677 | Akr1b3 | 119.810719 | 0.70079687 | 0.17867841 | 3.92211262 | 8.78E-05 |
| 230810 | SLC30A2 | 7.32510108 | -4.2030103 | 1.07175925 | -3.9215992 | 8.80E-05 |

| | | | | | | |
|---|---|---|---|---|---|---|
| 80907 | LACTB | 331.387537 | 0.56009303 | 0.14283468 | 3.92126771 | 8.81E-05 |
| 21917 | TMPO | 11750.0474 | -0.2036158 | 0.05192656 | -3.921227 | 8.81E-05 |
| 218203 | MYLIP | 375.795883 | 1.2931616 | 0.32979324 | 3.92112832 | 8.81E-05 |
| 233744 | SPON1 | 5.91422697 | -5.2360743 | 1.33543598 | -3.9208726 | 8.82E-05 |
| 232089 | ELMOD3 | 266.288383 | 0.52048136 | 0.13275552 | 3.92060059 | 8.83E-05 |
| 19247 | PTPN11 | 9273.81804 | -0.2852272 | 0.07278182 | -3.9189344 | 8.89E-05 |
| 74253 | KLRG2 | 204.463698 | -0.9407675 | 0.2401166 | -3.9179609 | 8.93E-05 |
| 215210 | tmem120a | 1312.12018 | -0.7718331 | 0.19700153 | -3.9179043 | 8.93E-05 |
| 668829 | Gm9385 | 930.923161 | -0.3034602 | 0.0774798 | -3.9166367 | 8.98E-05 |
| 100044193 | Gm20939 | 122.735776 | -0.789866 | 0.20170203 | -3.9160045 | 9.00E-05 |
| 330010 | TTLL10 | 3.74313612 | -5.3278892 | 1.36062041 | -3.9157793 | 9.01E-05 |
| 18797 | plcb3 | 2290.15341 | -0.4924935 | 0.12579806 | -3.9149532 | 9.04E-05 |
| 76479 | SMNDC1 | 1232.49638 | 0.19919318 | 0.05088063 | 3.91491181 | 9.04E-05 |
| 77352 | AXDND1 | 15.1743281 | -2.258714 | 0.57731159 | -3.9124694 | 9.14E-05 |
| 108961 | E2F8 | 2496.71626 | 0.46524823 | 0.11893381 | 3.91182492 | 9.16E-05 |
| 13667 | EIF2B4 | 692.750851 | 0.26470445 | 0.06769559 | 3.91021709 | 9.22E-05 |
| 668501 | Zfp507 | 1638.33058 | 0.53616292 | 0.13716369 | 3.90892744 | 9.27E-05 |
| 12768 | CCR1 | 14.2800317 | -2.3613244 | 0.60415111 | -3.9084997 | 9.29E-05 |
| 50850 | SPAST | 1350.68384 | -0.2256053 | 0.05772913 | -3.907997 | 9.31E-05 |
| 70873 | CNBD2 | 61.1598496 | -0.9874213 | 0.25269286 | -3.9075948 | 9.32E-05 |
| 56745 | C1QTNF1 | 123.94268 | -1.8941856 | 0.48478052 | -3.9073056 | 9.33E-05 |
| 17347 | MKNK2 | 2847.96737 | -0.3220974 | 0.08245839 | -3.9061809 | 9.38E-05 |
| 14156 | FEN1 | 2911.67308 | 0.30154565 | 0.07720308 | 3.90587597 | 9.39E-05 |
| 26904 | Sh2d1b1 | 21.9710519 | -2.6526231 | 0.67919587 | -3.9055348 | 9.40E-05 |
| 272359 | IRF2BP1 | 1323.12622 | 0.77689201 | 0.19902247 | 3.90353922 | 9.48E-05 |
| 24070 | MPDU1 | 1204.54615 | -0.4926353 | 0.12621618 | -3.9031074 | 9.50E-05 |
| 12585 | CDR2 | 1122.80341 | 1.03944417 | 0.26634105 | 3.90268102 | 9.51E-05 |
| 14990 | H2-M2 | 5.574328 | -5.9045068 | 1.51308265 | -3.9023029 | 9.53E-05 |
| 13436 | DNMT3B | 56.0177637 | 0.9229901 | 0.23674638 | 3.89864498 | 9.67E-05 |
| 28040 | D6Wsu163e | 679.036989 | 0.36807156 | 0.09441681 | 3.89836892 | 9.68E-05 |
| 225743 | RNF165 | 44.0358603 | 3.10051865 | 0.7955466 | 3.89734384 | 9.73E-05 |
| 68729 | TRIM37 | 2798.11957 | -0.2216469 | 0.05688156 | -3.8966382 | 9.75E-05 |
| 77053 | SUN1 | 2646.23174 | -0.5070539 | 0.13014573 | -3.896047 | 9.78E-05 |
| 66765 | 4933411K16I | 11.3823605 | -2.3383875 | 0.6003146 | -3.8952701 | 9.81E-05 |
| 18100 | MRPL40 | 803.060349 | -0.2843872 | 0.07301963 | -3.8946672 | 9.83E-05 |
| 270152 | AMICA1 | 4.41073693 | -4.8088841 | 1.23523713 | -3.8930858 | 9.90E-05 |
| 66892 | EIF4E3 | 89.4540778 | 1.20964653 | 0.3107307 | 3.89290966 | 9.90E-05 |
| 241944 | D3Ertd254e | 792.623565 | -0.4401213 | 0.11305817 | -3.8928752 | 9.91E-05 |
| 11480 | ACVR2A | 856.044124 | -0.3267437 | 0.08394054 | -3.8925613 | 9.92E-05 |
| 29807 | TPK1 | 63.4876803 | -0.8400378 | 0.21580702 | -3.8925415 | 9.92E-05 |
| 626578 | Gbp10 | 8.9922704 | -3.2545675 | 0.83664724 | -3.8900116 | 0.00010024 |
| 236920 | STARD8 | 7.04365297 | -4.1070014 | 1.05589002 | -3.889611 | 0.00010041 |
| 106931 | KCTD1 | 233.782699 | 0.62673953 | 0.16117527 | 3.88855878 | 0.00010084 |
| 242557 | ATG4C | 302.499773 | 0.63792622 | 0.16405686 | 3.88844592 | 0.00010089 |
| 18113 | NNMT | 48.9602716 | 3.48739794 | 0.89686976 | 3.88841065 | 0.0001009 |

| | | | | | | |
|---|---|---|---|---|---|---|
| 72486 | rnf219 | 795.562946 | 0.9477904 | 0.24376341 | 3.88815703 | 0.00010101 |
| 52846 | CNOT11 | 1490.88986 | 0.19906904 | 0.05120473 | 3.8877076 | 0.0001012 |
| 329738 | aknad1 | 6.0401639 | 3.98978507 | 1.02642561 | 3.88706694 | 0.00010146 |
| 18799 | PLCD1 | 419.629407 | 0.98635415 | 0.25390817 | 3.88468858 | 0.00010246 |
| 11819 | NR2F2 | 2561.81375 | 0.50411566 | 0.12977472 | 3.88454431 | 0.00010252 |
| 672511 | RNF213 | 6072.7421 | -0.5699063 | 0.1467693 | -3.8830076 | 0.00010317 |
| 20918 | EIF1 | 4232.581 | 0.25192786 | 0.06491843 | 3.88068291 | 0.00010416 |
| 100504139 | Gm16685 | 6.54724574 | -5.390373 | 1.38966647 | -3.8788969 | 0.00010493 |
| 56036 | CCNL2 | 1295.22273 | 0.37496544 | 0.09668202 | 3.87833668 | 0.00010517 |
| 15412 | HOXB4 | 243.379912 | 0.75933098 | 0.19578942 | 3.87830442 | 0.00010519 |
| 20020 | POLR2A | 10703.8962 | 0.378728 | 0.09777497 | 3.87346572 | 0.0001073 |
| 17748 | Mt1 | 3805.51097 | 0.31100451 | 0.08031851 | 3.87213989 | 0.00010788 |
| 21681 | ALYREF | 3294.45435 | 0.25774124 | 0.06656338 | 3.87211783 | 0.00010789 |
| 24083 | natd1 | 679.501383 | -0.8506045 | 0.21972526 | -3.8712185 | 0.00010829 |
| 73824 | SNHG6 | 333.991011 | -0.5080814 | 0.13128073 | -3.8701904 | 0.00010875 |
| 66233 | DMAP1 | 446.54598 | 0.67802327 | 0.17520016 | 3.86999223 | 0.00010884 |
| 114249 | NPNT | 97.687112 | -1.8072589 | 0.46701452 | -3.8698131 | 0.00010892 |
| 57874 | HACD3 | 2061.9912 | -0.317496 | 0.08205526 | -3.8692951 | 0.00010915 |
| 100502601 | Gm19276 | 3.51990381 | 5.29655176 | 1.36895702 | 3.86904167 | 0.00010926 |
| 77034 | 2510039O18 | 2254.27384 | 0.47416153 | 0.12255566 | 3.86894854 | 0.00010931 |
| 71910 | PLPP5 | 492.227736 | 0.34174114 | 0.08836872 | 3.86721836 | 0.00011008 |
| 27277 | GOLGA5 | 1352.32354 | 0.39770113 | 0.10286278 | 3.86632683 | 0.00011049 |
| 109700 | ITGA1 | 115.974934 | 2.71596443 | 0.70269681 | 3.86505872 | 0.00011106 |
| 94180 | acsbg1 | 639.429071 | -1.325859 | 0.34304489 | -3.8649724 | 0.0001111 |
| 237806 | DNAH9 | 4.22696467 | -5.5146729 | 1.4268364 | -3.8649651 | 0.00011111 |
| 22596 | XRCC5 | 1432.43785 | -0.4739515 | 0.12263607 | -3.8646992 | 0.00011123 |
| 170441 | SLC2A10 | 3.55467203 | 5.31612015 | 1.37570937 | 3.86427559 | 0.00011142 |
| 23960 | Oas1g | 31.6623807 | -3.5787849 | 0.92663946 | -3.8621115 | 0.00011241 |
| 77980 | SBF1 | 5642.60335 | -0.2872046 | 0.07438117 | -3.8612536 | 0.00011281 |
| 50917 | GALNS | 615.603376 | -1.1105545 | 0.2876443 | -3.8608605 | 0.00011299 |
| 72344 | USP36 | 1854.24702 | 0.30646135 | 0.07938274 | 3.86055396 | 0.00011313 |
| 67042 | IFT27 | 268.729534 | -0.5556038 | 0.14398948 | -3.8586415 | 0.00011402 |
| 50932 | MINK1 | 2076.861 | -0.444435 | 0.11518874 | -3.8583199 | 0.00011417 |
| 435965 | LRP3 | 14.5211273 | -7.2913351 | 1.8899298 | -3.8579926 | 0.00011432 |
| 13731 | EMP2 | 3555.13959 | -0.9903947 | 0.25676015 | -3.8572759 | 0.00011466 |
| 68968 | CDAN1 | 805.036759 | 0.42562166 | 0.1103481 | 3.85708188 | 0.00011475 |
| 235330 | TTC12 | 113.96644 | -1.216864 | 0.31550862 | -3.8568328 | 0.00011487 |
| 26432 | PLOD2 | 2717.42008 | 1.22198043 | 0.31703888 | 3.85435506 | 0.00011604 |
| 13406 | DMP1 | 3.22023569 | 5.1715616 | 1.34197196 | 3.85370318 | 0.00011634 |
| 215690 | NAV1 | 3084.42378 | 0.91848423 | 0.23836352 | 3.85329199 | 0.00011654 |
| 70408 | POLR3F | 1046.21762 | 0.3245472 | 0.0842563 | 3.85190408 | 0.0001172 |
| 235040 | ATG4D | 451.137995 | 0.59394141 | 0.1542 | 3.8517602 | 0.00011727 |
| 66170 | chchd5 | 277.87526 | -0.6372819 | 0.16547369 | -3.8512584 | 0.00011751 |
| 14682 | GNAQ | 2768.50283 | -0.394979 | 0.10258447 | -3.8502807 | 0.00011798 |
| 100042802 | AI463229 | 5.08962647 | -5.7748341 | 1.50047996 | -3.8486579 | 0.00011877 |

| | | | | | | |
|---|---|---|---|---|---|---|
| 108689 | OBFC1 | 812.892098 | 0.59565673 | 0.15480883 | 3.84769231 | 0.00011924 |
| 56550 | Ube2d2a | 1153.88049 | -0.3252453 | 0.08453398 | -3.847509 | 0.00011932 |
| 434175 | Gm5593 | 231.728656 | 0.69636381 | 0.18099202 | 3.8474834 | 0.00011934 |
| 72333 | PALLD | 1534.90046 | -1.2323818 | 0.32034166 | -3.8470855 | 0.00011953 |
| 432777 | D630045M09 | 6.51605057 | -4.6449948 | 1.20781132 | -3.845795 | 0.00012016 |
| 66047 | MRPL54 | 982.264098 | -0.3314302 | 0.08619319 | -3.845201 | 0.00012045 |
| 100504464 | E230016K23I | 10.0227915 | -5.2892987 | 1.37558983 | -3.8451133 | 0.0001205 |
| 16516 | KCNJ15 | 783.468338 | -8.2615694 | 2.14922191 | -3.8439816 | 0.00012105 |
| 77087 | ANKRD11 | 6272.10122 | 0.30159993 | 0.07846712 | 3.84364742 | 0.00012122 |
| 100040736 | Foxd2os | 150.46416 | 0.947504 | 0.2465194 | 3.84352713 | 0.00012128 |
| 195018 | ZZEF1 | 2502.89917 | 0.3572856 | 0.09296026 | 3.84342307 | 0.00012133 |
| 26950 | VSNL1 | 5.87844776 | -5.9810136 | 1.55617082 | -3.8434171 | 0.00012133 |
| 69714 | TFPT | 351.340689 | 0.4382102 | 0.11401958 | 3.84328894 | 0.0001214 |
| 59083 | FETUB | 103.121706 | 0.89259115 | 0.23226494 | 3.84298697 | 0.00012155 |
| 100862396 | #N/A | 7.41396633 | 4.86990356 | 1.2675011 | 3.84212965 | 0.00012197 |
| 109685 | hyal3 | 16.86506 | -1.567213 | 0.40791909 | -3.8419703 | 0.00012205 |
| 106878 | SMIM3 | 453.607405 | -0.6304453 | 0.16412194 | -3.8413227 | 0.00012237 |
| 76357 | TRMT5 | 595.481996 | 0.4954934 | 0.129008 | 3.84079579 | 0.00012264 |
| 104370 | Snora68 | 37.8109352 | 1.14407933 | 0.29791804 | 3.84024857 | 0.00012291 |
| 20115 | RPS7 | 185.090836 | 0.62981872 | 0.16403508 | 3.83953668 | 0.00012327 |
| 109212 | FAM64A | 1672.47421 | 0.49317703 | 0.12844721 | 3.83953076 | 0.00012327 |
| 50771 | ATP9B | 2272.50948 | -0.3722278 | 0.09696203 | -3.8389025 | 0.00012359 |
| 218214 | KDM1B | 618.458767 | -0.3225001 | 0.08401142 | -3.8387654 | 0.00012365 |
| 52708 | Zfp410 | 1012.85697 | 0.23968992 | 0.06245775 | 3.83763308 | 0.00012423 |
| 217109 | UTP18 | 2055.53714 | 0.19851991 | 0.05173895 | 3.83695269 | 0.00012457 |
| 71742 | ULK3 | 356.751139 | -0.6195397 | 0.16150928 | -3.835939 | 0.00012509 |
| 77862 | THYN1 | 822.879133 | 0.26355999 | 0.06872127 | 3.83520244 | 0.00012546 |
| 12163 | BMP8A | 10.8781101 | -2.7663296 | 0.72133002 | -3.8350402 | 0.00012554 |
| 69736 | NUP37 | 947.711647 | 0.41044486 | 0.10702762 | 3.83494342 | 0.00012559 |
| 80860 | GHDC | 377.78082 | -0.7596517 | 0.19809948 | -3.834698 | 0.00012572 |
| 107817 | JMJD6 | 1302.29016 | 0.23701688 | 0.06181652 | 3.83419938 | 0.00012597 |
| 72018 | FUNDC1 | 739.918233 | -0.386506 | 0.10083581 | -3.8330233 | 0.00012658 |
| 21400 | TCEA2 | 56.6597455 | 1.85313649 | 0.4834827 | 3.832891 | 0.00012665 |
| 58246 | SLC35B4 | 2196.73396 | 0.28531206 | 0.07445015 | 3.83225636 | 0.00012697 |
| 20017 | POLR1B | 2164.87045 | 0.45359845 | 0.11836616 | 3.83216312 | 0.00012702 |
| 72193 | SCAF11 | 7317.47513 | 0.26557612 | 0.06930336 | 3.83208133 | 0.00012706 |
| 56748 | NFU1 | 572.09021 | -0.3472129 | 0.09060689 | -3.8320806 | 0.00012706 |
| 11569 | AEBP2 | 3871.28868 | 0.38052317 | 0.09933741 | 3.83061279 | 0.00012782 |
| 19014 | MED1 | 3956.79214 | 0.18790995 | 0.04905573 | 3.83054032 | 0.00012786 |
| 66506 | PSMG3 | 490.475212 | 0.39127893 | 0.10219213 | 3.82885601 | 0.00012874 |
| 75734 | MFF | 1823.42906 | -0.2652254 | 0.06927105 | -3.8288053 | 0.00012877 |
| 70223 | NARS | 16027.9629 | 0.25805424 | 0.0674098 | 3.82814147 | 0.00012911 |
| 22437 | XIRP1 | 15.5654458 | -5.9003486 | 1.5413785 | -3.8279687 | 0.00012921 |
| 338467 | MORC3 | 1933.92379 | 0.35978868 | 0.09399496 | 3.82774439 | 0.00012932 |
| 228543 | rhov | 15.5182622 | -5.8966705 | 1.54052314 | -3.8277066 | 0.00012934 |

| | | | | | | |
|---|---|---|---|---|---|---|
| 16667 | krt17 | 25.879979 | -6.1220647 | 1.5996073 | -3.8272298 | 0.00012959 |
| 21346 | TAGLN2 | 12898.526 | -1.2253512 | 0.32019035 | -3.8269461 | 0.00012974 |
| 26920 | CNTRL | 1139.34142 | 0.5150769 | 0.13465175 | 3.82525208 | 0.00013064 |
| 12815 | COL11A2 | 101.978288 | 0.65393046 | 0.17096532 | 3.82493037 | 0.00013081 |
| 13423 | Dnase2a | 306.796132 | 1.02226451 | 0.2672892 | 3.8245634 | 0.000131 |
| 243910 | NFKBID | 123.476209 | 2.07829616 | 0.54355618 | 3.82351673 | 0.00013156 |
| 27393 | MRPL39 | 1204.3194 | 0.3925077 | 0.1026703 | 3.82299178 | 0.00013184 |
| 666634 | Gm8203 | 2387.01367 | 0.33056944 | 0.08647607 | 3.82266931 | 0.00013201 |
| 193034 | TRPV1 | 4.43206297 | -5.5721127 | 1.45767586 | -3.8226007 | 0.00013205 |
| 15893 | ICA1 | 377.228101 | -0.638101 | 0.1669408 | -3.8223188 | 0.0001322 |
| 71233 | ENKUR | 12.532919 | -2.4170168 | 0.63259563 | -3.8207927 | 0.00013302 |
| 73230 | BMPER | 1169.54492 | 1.02667629 | 0.26885114 | 3.81875366 | 0.00013413 |
| 66612 | ORMDL3 | 727.708606 | -0.4453375 | 0.11667381 | -3.8169447 | 0.00013511 |
| 100504089 | Gm20056 | 260.45985 | 0.47364061 | 0.12410455 | 3.81646441 | 0.00013538 |
| 217154 | STAC2 | 1928.21888 | -0.9902351 | 0.25949116 | -3.8160649 | 0.0001356 |
| 268567 | TMEM229B | 9.43621192 | -2.4224199 | 0.63493492 | -3.8152254 | 0.00013606 |
| 66645 | PSPC1 | 890.073626 | -1.2342305 | 0.3235595 | -3.8145396 | 0.00013644 |
| 13132 | DAB2 | 8138.80629 | 2.73532885 | 0.71714949 | 3.81416832 | 0.00013664 |
| 69885 | AUNIP | 379.804726 | 0.72897882 | 0.19113717 | 3.81390393 | 0.00013679 |
| 14670 | GNL1 | 2321.51088 | 0.33050941 | 0.08667039 | 3.81340605 | 0.00013706 |
| 101471 | PHRF1 | 3458.03547 | 0.32742402 | 0.08586533 | 3.81322731 | 0.00013716 |
| 18763 | PKD1 | 3828.55726 | 0.77080322 | 0.20215333 | 3.81296318 | 0.00013731 |
| 20531 | SLC34A2 | 7.73256621 | -6.3779465 | 1.67303502 | -3.8122014 | 0.00013773 |
| 27355 | PALD1 | 575.199901 | -0.5993644 | 0.15722326 | -3.8121869 | 0.00013774 |
| 16779 | LAMB2 | 2788.62504 | 0.33566258 | 0.08805188 | 3.81209998 | 0.00013779 |
| 64934 | PES1 | 2513.65547 | 0.17242472 | 0.04523754 | 3.81154085 | 0.0001381 |
| 20382 | SRSF2 | 10946.5058 | 0.31719517 | 0.08322988 | 3.81107317 | 0.00013836 |
| 56795 | ARL10 | 209.64252 | 0.52210947 | 0.13699999 | 3.81101823 | 0.0001384 |
| 69537 | DNASE1L1 | 400.376808 | -0.4719397 | 0.12384222 | -3.8108142 | 0.00013851 |
| 77634 | SNAPC3 | 285.886722 | 0.44287836 | 0.11621706 | 3.81078593 | 0.00013853 |
| 74159 | ACBD5 | 1442.84079 | -0.326689 | 0.08572827 | -3.8107497 | 0.00013855 |
| 100043040 | 1110002L01F | 456.473194 | 0.76287753 | 0.2001958 | 3.81065696 | 0.0001386 |
| 319991 | KIF6 | 3.97331709 | -5.4266911 | 1.4242 | -3.8103434 | 0.00013877 |
| 23888 | GPC6 | 835.616921 | -0.7142359 | 0.18745116 | -3.8102504 | 0.00013883 |
| 54402 | STK19 | 291.481741 | -0.4532004 | 0.11899444 | -3.8085844 | 0.00013976 |
| 69215 | SAT2 | 25.9668732 | 1.48263894 | 0.38946985 | 3.80681317 | 0.00014077 |
| 26409 | MAP3K7 | 3479.15724 | -0.907211 | 0.23834434 | -3.8063039 | 0.00014106 |
| 68147 | GAR1 | 1232.34579 | 0.4910419 | 0.12901712 | 3.80602134 | 0.00014122 |
| 233905 | Zfp646 | 1578.09995 | 0.5757754 | 0.15128165 | 3.80598297 | 0.00014124 |
| 69009 | thap7 | 638.325451 | 0.29277149 | 0.07692533 | 3.80591776 | 0.00014128 |
| 217473 | ANKMY2 | 539.765269 | 0.42148407 | 0.11077825 | 3.80475484 | 0.00014194 |
| 66914 | VPS28 | 856.726525 | -0.3001958 | 0.07890836 | -3.8043597 | 0.00014217 |
| 57279 | SLC25A20 | 447.240625 | -0.4445209 | 0.11685536 | -3.8040263 | 0.00014236 |
| 14687 | GNAZ | 34.5669268 | -1.5936444 | 0.41894817 | -3.8039179 | 0.00014243 |
| 320376 | BCORL1 | 600.775629 | -0.5905235 | 0.15525317 | -3.8036165 | 0.0001426 |

| | | | | | | |
|---|---|---|---|---|---|---|
| 320197 | D830046C22 | 6.93342884 | -6.2266611 | 1.63708453 | -3.8035061 | 0.00014266 |
| 53332 | MTMR1 | 884.577366 | -0.2010438 | 0.05285761 | -3.8034974 | 0.00014267 |
| 72171 | SHQ1 | 543.524758 | 0.46351433 | 0.12186728 | 3.80343544 | 0.0001427 |
| 15463 | AGFG1 | 3422.55816 | 0.51959262 | 0.13662007 | 3.80319401 | 0.00014284 |
| 57738 | SLC15A2 | 73.8324698 | -1.6982344 | 0.44654442 | -3.8030582 | 0.00014292 |
| 13615 | EDN2 | 6.38599543 | -5.3488542 | 1.40646946 | -3.8030361 | 0.00014293 |
| 12764 | CMAS | 1139.22043 | -0.6995822 | 0.18396312 | -3.8028395 | 0.00014305 |
| 216877 | DHX33 | 2671.83003 | 0.38942971 | 0.10241007 | 3.80265063 | 0.00014316 |
| 16905 | LMNA | 30929.8929 | 0.45046695 | 0.11851583 | 3.8009013 | 0.00014417 |
| 13650 | RHBDF1 | 2505.32948 | -0.7738722 | 0.20372234 | -3.7986612 | 0.00014548 |
| 211323 | NRG1 | 501.549055 | 1.06975929 | 0.28162195 | 3.79856501 | 0.00014554 |
| 320806 | GFM2 | 568.407661 | 0.28543104 | 0.07514902 | 3.79820018 | 0.00014575 |
| 14595 | B4GALT1 | 5419.18052 | -0.8099402 | 0.21326135 | -3.7978763 | 0.00014594 |
| 320022 | TERB1 | 4.12620082 | -5.4701346 | 1.44077528 | -3.7966605 | 0.00014666 |
| 56805 | ZBTB33 | 1512.66035 | -0.1814794 | 0.04780082 | -3.7965757 | 0.00014671 |
| 20276 | SCNN1A | 43.9114502 | -1.4614741 | 0.38496942 | -3.7963381 | 0.00014685 |
| 11640 | AKAP1 | 2605.08024 | -0.1481799 | 0.03903741 | -3.7958445 | 0.00014714 |
| 54216 | PCDH7 | 4267.90826 | -0.9712039 | 0.2558602 | -3.795838 | 0.00014715 |
| 72147 | ZBTB46 | 37.9031517 | 2.17568522 | 0.57324663 | 3.79537377 | 0.00014742 |
| 233410 | Zfp592 | 1403.01447 | 0.51145859 | 0.13477138 | 3.79500885 | 0.00014764 |
| 100504308 | Gm15473 | 12.5832299 | 2.40849038 | 0.63479939 | 3.79409686 | 0.00014818 |
| 230249 | AI314180 | 6268.79692 | 0.37724902 | 0.09945505 | 3.79316106 | 0.00014874 |
| 94192 | C1GALT1 | 1240.61655 | 0.5541534 | 0.14612326 | 3.79236944 | 0.00014922 |
| 52633 | NIT2 | 552.408803 | 0.32837469 | 0.08659372 | 3.79213059 | 0.00014936 |
| 67657 | RABL3 | 389.096173 | -0.3170911 | 0.08361985 | -3.7920557 | 0.00014941 |
| 17380 | MME | 1316.21116 | -1.4584056 | 0.384734 | -3.7906855 | 0.00015023 |
| 12765 | CXCR2 | 4.39858376 | -4.8139949 | 1.2701609 | -3.790067 | 0.00015061 |
| 17874 | MYD88 | 842.222619 | 0.72160132 | 0.19044242 | 3.78907869 | 0.00015121 |
| 100503868 | Gm19935 | 10.9872792 | -6.8860823 | 1.81786252 | -3.7880105 | 0.00015186 |
| 24109 | UBL3 | 2245.11789 | -0.4341893 | 0.1146328 | -3.7876533 | 0.00015208 |
| 72061 | 2010111I01R | 517.730327 | 0.57022254 | 0.15055945 | 3.78735814 | 0.00015226 |
| 216227 | SLC17A8 | 5.90910257 | -3.436453 | 0.90737185 | -3.7872599 | 0.00015232 |
| 100040260 | Gm1862 | 247.08919 | 0.50106574 | 0.13230511 | 3.78719877 | 0.00015236 |
| 228993 | SLC17A9 | 8.54405139 | -2.5793264 | 0.68111307 | -3.7869283 | 0.00015252 |
| 235472 | PRTG | 39.8709623 | 2.19945484 | 0.58083526 | 3.78671026 | 0.00015265 |
| 70693 | adgra3 | 1789.67229 | 0.62758004 | 0.1658394 | 3.78426387 | 0.00015416 |
| 76477 | PCOLCE2 | 2355.80995 | 0.86176179 | 0.22776958 | 3.78348066 | 0.00015465 |
| 55983 | PDZRN3 | 2829.86798 | -0.9149747 | 0.24185076 | -3.7832202 | 0.00015481 |
| 14748 | GPR3 | 13.5784274 | 1.84949851 | 0.48889009 | 3.78305588 | 0.00015491 |
| 19660 | RBP2 | 11.0563287 | 6.20762842 | 1.6414607 | 3.78177097 | 0.00015572 |
| 434204 | WHAMM | 327.195961 | 0.32573755 | 0.08615216 | 3.78095629 | 0.00015623 |
| 237781 | MIEF2 | 291.211436 | -0.4381749 | 0.11589821 | -3.7806879 | 0.0001564 |
| 81904 | CACNG7 | 4.98967429 | -5.7535086 | 1.52187025 | -3.7805513 | 0.00015648 |
| 140488 | IGF2BP3 | 7.34672559 | -2.911868 | 0.77028973 | -3.7802244 | 0.00015669 |
| 107995 | CDC20 | 3322.12539 | 1.04443541 | 0.27632116 | 3.77978807 | 0.00015696 |

| | | | | | | |
|---|---|---|---|---|---|---|
| 66237 | ATP6V1G2 | 18.0494373 | 1.54103487 | 0.40776429 | 3.7792296 | 0.00015731 |
| 18139 | Zfp638 | 3135.48992 | 0.19611212 | 0.05190149 | 3.77854497 | 0.00015775 |
| 12282 | HYOU1 | 5743.82602 | 0.51228344 | 0.1355848 | 3.77832485 | 0.00015789 |
| 214580 | PSTK | 291.701725 | 0.63352065 | 0.16769228 | 3.77787608 | 0.00015817 |
| 67941 | RPS27L | 1351.54213 | -0.2889229 | 0.07649429 | -3.7770515 | 0.0001587 |
| 72554 | UTP14A | 1573.95119 | 0.23365492 | 0.06188328 | 3.77573617 | 0.00015954 |
| 71319 | 4933439K11I | 6.58273898 | -6.1520131 | 1.62960464 | -3.7751569 | 0.00015991 |
| 68190 | DUBR | 168.256821 | -0.4732374 | 0.12537369 | -3.774615 | 0.00016025 |
| 214424 | PARP16 | 294.933027 | -0.5053371 | 0.13390891 | -3.7737379 | 0.00016082 |
| 68053 | UBXN2B | 335.26394 | -1.0572199 | 0.28025914 | -3.772294 | 0.00016175 |
| 17129 | SMAD5 | 2876.78928 | 0.58416725 | 0.15487012 | 3.77198155 | 0.00016196 |
| 626802 | Gm14322 | 23.6457676 | 2.01808115 | 0.53512392 | 3.7712408 | 0.00016244 |
| 270906 | PRR11 | 2611.95573 | 0.30155562 | 0.07998762 | 3.77002883 | 0.00016323 |
| 74476 | 4933439C10I | 115.989275 | -0.7385486 | 0.195909 | -3.7698553 | 0.00016334 |
| 80986 | CKAP2 | 2831.62878 | -0.7134826 | 0.18929317 | -3.7691934 | 0.00016378 |
| 12306 | ANXA2 | 29216.682 | 0.41661979 | 0.11056705 | 3.7680283 | 0.00016454 |
| 100043823 | Gm4673 | 38.0962355 | 1.24281437 | 0.32986434 | 3.76765295 | 0.00016479 |
| 71998 | SLC25A35 | 18.9981506 | -1.7504362 | 0.4646077 | -3.7675575 | 0.00016485 |
| 432769 | Zfp708 | 27.9128652 | -1.660066 | 0.44066466 | -3.7671865 | 0.0001651 |
| 104082 | WDR7 | 1779.22837 | -0.4249371 | 0.1128327 | -3.7660808 | 0.00016583 |
| 107260 | OTUB1 | 3440.25039 | -0.3886497 | 0.1032153 | -3.7654275 | 0.00016626 |
| 72881 | ZDHHC4 | 550.298142 | -0.5626169 | 0.14942344 | -3.7652522 | 0.00016638 |
| 66461 | PTPMT1 | 972.59192 | -0.3425163 | 0.09097343 | -3.7650141 | 0.00016654 |
| 230598 | NRD1 | 4786.21692 | 0.73752736 | 0.19603473 | 3.76222806 | 0.00016841 |
| 100756 | USP30 | 782.994594 | -0.3587513 | 0.09537047 | -3.7616604 | 0.00016879 |
| 109934 | ABR | 5513.46417 | 1.06041135 | 0.28198426 | 3.76053389 | 0.00016955 |
| 100503250 | Gm19610 | 6.39769298 | 4.65229357 | 1.23718744 | 3.76037891 | 0.00016966 |
| 210853 | Zfp947 | 38.4721947 | -1.2181276 | 0.32394492 | -3.7602922 | 0.00016972 |
| 20585 | HLTF | 1081.92354 | -0.3934689 | 0.10466024 | -3.7594876 | 0.00017026 |
| 239408 | TMEM74 | 115.956631 | 4.20141436 | 1.11765611 | 3.75912979 | 0.00017051 |
| 319757 | SMO | 3657.39835 | 0.5280417 | 0.1404738 | 3.75900499 | 0.00017059 |
| 77929 | YIPF6 | 1134.21298 | -0.5559756 | 0.14792386 | -3.7585256 | 0.00017092 |
| 74116 | PI16 | 3.36561835 | 5.23374354 | 1.39299407 | 3.75719011 | 0.00017183 |
| 26364 | ADGRE5 | 187.57018 | -3.4929895 | 0.92971946 | -3.757036 | 0.00017194 |
| 13002 | DNAJC5 | 4357.69067 | -0.8192682 | 0.21807267 | -3.7568586 | 0.00017206 |
| 20523 | SLC25A14 | 203.465239 | -0.4488582 | 0.11950151 | -3.756088 | 0.00017259 |
| 66917 | CHORDC1 | 2395.30611 | 0.46984207 | 0.12509142 | 3.75598969 | 0.00017266 |
| 11937 | ATP2A1 | 3.9181962 | -5.3932872 | 1.43592875 | -3.7559574 | 0.00017268 |
| 100101806 | Srp54c | 304.028776 | 0.43339149 | 0.11541968 | 3.75491849 | 0.0001734 |
| 329384 | PTRH1 | 963.879398 | 0.37682263 | 0.10038939 | 3.75361016 | 0.00017431 |
| 328099 | Prps1l3 | 3226.25135 | 0.37664911 | 0.10034907 | 3.75338922 | 0.00017446 |
| 17096 | LYN | 766.50806 | -1.1637197 | 0.31005704 | -3.753244 | 0.00017456 |
| 277854 | DEPDC5 | 960.521942 | 0.28619468 | 0.07626332 | 3.75271707 | 0.00017493 |
| 320769 | Prdx6b | 62.5972749 | 0.82193436 | 0.21906406 | 3.75202744 | 0.00017541 |
| 53424 | TSNAX | 1914.71371 | -0.3289642 | 0.087694 | -3.7512738 | 0.00017594 |

| | | | | | | |
|---|---|---|---|---|---|---|
| 70536 | QPCT | 98.0766768 | -2.0487937 | 0.54622497 | -3.7508239 | 0.00017625 |
| 74356 | 4931428F04I | 77.0210757 | 0.79020336 | 0.21069836 | 3.750401 | 0.00017655 |
| 21676 | TEAD1 | 10064.4037 | 0.65219082 | 0.17391132 | 3.75013429 | 0.00017674 |
| 665700 | HMCN2 | 44.3855158 | -1.0002209 | 0.26676354 | -3.7494664 | 0.00017721 |
| 67027 | MKRN2 | 565.211746 | 0.5237445 | 0.13968754 | 3.74940018 | 0.00017726 |
| 21897 | TLR1 | 4.91364701 | -4.9645538 | 1.32413856 | -3.7492706 | 0.00017735 |
| 432649 | Gm5434 | 62.7655153 | -0.7945988 | 0.21205017 | -3.7472209 | 0.0001788 |
| 20362 | 41889 | 3805.64274 | -0.3628112 | 0.0968231 | -3.7471556 | 0.00017885 |
| 81701 | EGFL8 | 14.3551093 | -1.7829916 | 0.4758892 | -3.7466528 | 0.00017921 |
| 26558 | HOMER3 | 1877.75575 | -0.7522479 | 0.20078385 | -3.7465558 | 0.00017928 |
| 69183 | C1QTNF2 | 9.7710326 | -3.3696273 | 0.89951091 | -3.7460661 | 0.00017963 |
| 59002 | WRAP73 | 350.13025 | 0.59137322 | 0.15791978 | 3.74476978 | 0.00018056 |
| 67867 | LRRC28 | 438.162789 | -0.3908884 | 0.104384 | -3.7447158 | 0.0001806 |
| 72668 | skida1 | 105.932332 | 0.83566764 | 0.22327981 | 3.74269234 | 0.00018206 |
| 17295 | MET | 6201.0232 | -0.6141396 | 0.16409879 | -3.742499 | 0.0001822 |
| 333050 | KSR2 | 4.64328776 | -4.893206 | 1.30778962 | -3.741585 | 0.00018286 |
| 30049 | Scd3 | 8.01819061 | -3.5118512 | 0.93882821 | -3.7406749 | 0.00018353 |
| 108089 | RNF144A | 23.1496286 | 4.75937444 | 1.27238849 | 3.74050417 | 0.00018365 |
| 30946 | ABT1 | 586.02681 | 0.47554152 | 0.12714151 | 3.74025395 | 0.00018383 |
| 97165 | HMGB2 | 521.067338 | 0.41775572 | 0.11170652 | 3.73976141 | 0.0001842 |
| 12452 | CCNG2 | 537.24943 | -0.9593593 | 0.25654742 | -3.739501 | 0.00018439 |
| 76167 | SNRNP35 | 275.321515 | 0.55582219 | 0.14870474 | 3.737757 | 0.00018567 |
| 627049 | Zfp800 | 850.797236 | -0.4203043 | 0.11246513 | -3.737197 | 0.00018608 |
| 234384 | MPV17L2 | 703.33192 | -0.2987033 | 0.07995428 | -3.7359264 | 0.00018703 |
| 18991 | POU3F1 | 6.18109365 | -3.9434462 | 1.05566625 | -3.7355047 | 0.00018734 |
| 319554 | IDI1 | 211.60763 | 0.67906576 | 0.18179939 | 3.73524773 | 0.00018753 |
| 13411 | DNAH11 | 5.68451875 | -3.794406 | 1.0160662 | -3.7344082 | 0.00018816 |
| 56462 | MTCH1 | 3421.78321 | -0.5038603 | 0.13500337 | -3.7322054 | 0.00018981 |
| 67247 | 41699 | 841.155297 | -0.3337604 | 0.08943958 | -3.7316861 | 0.0001902 |
| 76123 | GPSM2 | 1810.83757 | 0.45255019 | 0.12137399 | 3.72855983 | 0.00019258 |
| 67041 | OXCT1 | 7018.24105 | 0.73351638 | 0.19675003 | 3.72816393 | 0.00019288 |
| 233057 | Zfp940 | 85.6377988 | 1.10519658 | 0.29648533 | 3.72766023 | 0.00019327 |
| 67164 | LIPT2 | 122.795263 | 0.83822171 | 0.2248904 | 3.72724543 | 0.00019358 |
| 72098 | TMEM68 | 905.930036 | -1.0153575 | 0.27245213 | -3.7267373 | 0.00019397 |
| 56700 | GLMP | 3704.13845 | -0.8131235 | 0.2181868 | -3.726731 | 0.00019398 |
| 21899 | TLR6 | 122.511449 | -0.8511587 | 0.22841128 | -3.7264304 | 0.00019421 |
| 15490 | HSD17B7 | 3175.04264 | -0.5592598 | 0.1500862 | -3.7262573 | 0.00019434 |
| 668137 | Gm8994 | 5.505734 | -5.8863894 | 1.57977458 | -3.7260945 | 0.00019447 |
| 17865 | MYBL2 | 1203.35034 | 0.3325814 | 0.08926594 | 3.72573685 | 0.00019475 |
| 218734 | 3830406C13I | 269.280051 | -0.7129804 | 0.19137342 | -3.7255978 | 0.00019485 |
| 665928 | Gm7850 | 8.92508272 | -5.0916043 | 1.36675413 | -3.7253257 | 0.00019506 |
| 18105 | NQO2 | 993.73367 | 0.52198303 | 0.14012825 | 3.72503796 | 0.00019529 |
| 26875 | PCLO | 3.61260925 | -5.2844324 | 1.41926431 | -3.7233603 | 0.00019659 |
| 76501 | COMMD9 | 517.295731 | -0.4877366 | 0.13101457 | -3.7227661 | 0.00019705 |
| 16000 | IGF1 | 6.35240932 | -4.577375 | 1.23007393 | -3.7212194 | 0.00019826 |

| | | | | | | |
|---|---|---|---|---|---|---|
| 15356 | HMGCL | 372.64881 | 0.5592675 | 0.15032344 | 3.72042778 | 0.00019889 |
| 192656 | RIPK2 | 1555.9989 | -1.3176576 | 0.35430898 | -3.7189506 | 0.00020005 |
| 12190 | brca2 | 1659.40628 | -0.4562064 | 0.12267606 | -3.7187893 | 0.00020018 |
| 71878 | FAM83D | 1568.58506 | 0.45979041 | 0.12368444 | 3.71744756 | 0.00020125 |
| 18786 | PLAA | 4347.42094 | 0.7704328 | 0.20731894 | 3.71617174 | 0.00020226 |
| 69478 | 2300009A05 | 121.441479 | 0.62741433 | 0.16886313 | 3.71552002 | 0.00020279 |
| 13641 | EFNB1 | 1333.07727 | -1.4491568 | 0.3900446 | -3.7153617 | 0.00020291 |
| 52466 | SLC46A1 | 159.229615 | -0.6991395 | 0.18821292 | -3.71462 | 0.00020351 |
| 15458 | HPX | 9.89647131 | -2.2071085 | 0.59439922 | -3.7131754 | 0.00020467 |
| 83409 | LAMTOR2 | 643.851917 | -0.4224405 | 0.113815 | -3.7116418 | 0.00020592 |
| 213452 | DSTYK | 918.982386 | -0.3924276 | 0.10573963 | -3.7112631 | 0.00020623 |
| 381626 | RBM33 | 2722.70712 | 0.36344875 | 0.09794454 | 3.71076057 | 0.00020664 |
| 83704 | SLC12A9 | 592.287417 | -0.8004423 | 0.21581834 | -3.7088706 | 0.00020819 |
| 76804 | KDM4C | 801.922237 | -0.559615 | 0.15088901 | -3.7087854 | 0.00020826 |
| 20850 | STAT5A | 362.407691 | -0.3782346 | 0.10200183 | -3.7081161 | 0.00020881 |
| 110265 | MSRA | 200.204735 | 0.56750336 | 0.15304702 | 3.70803277 | 0.00020888 |
| 74761 | MXRA8 | 865.880192 | 2.23338595 | 0.6023487 | 3.70779577 | 0.00020907 |
| 74694 | TBC1D30 | 66.5913205 | -1.819897 | 0.49091374 | -3.7071625 | 0.00020959 |
| 232969 | Zfp428 | 387.143711 | 0.93159028 | 0.25134951 | 3.70635408 | 0.00021026 |
| 72535 | ALDH1B1 | 21.1169204 | -2.872003 | 0.7749496 | -3.7060513 | 0.00021052 |
| 27368 | TBL2 | 1354.73643 | 0.33714285 | 0.09101561 | 3.70423117 | 0.00021203 |
| 94254 | WBSCR16 | 885.51051 | 0.25294915 | 0.06828907 | 3.70409429 | 0.00021215 |
| 241727 | SNPH | 8.5634928 | -5.791858 | 1.56369548 | -3.7039552 | 0.00021226 |
| 12144 | BLM | 1448.71426 | 0.71815466 | 0.19394916 | 3.70279848 | 0.00021323 |
| 77580 | 5730596B20 | 4.27185012 | -5.5242209 | 1.49220531 | -3.7020515 | 0.00021386 |
| 51813 | CCNC | 1120.12038 | -0.9819707 | 0.26532833 | -3.7009643 | 0.00021478 |
| 11865 | ARNTL | 783.635318 | -0.4119131 | 0.11130542 | -3.700746 | 0.00021497 |
| 58869 | PEX5L | 5.57090613 | -5.9046146 | 1.59584262 | -3.6999981 | 0.0002156 |
| 71446 | WRB | 824.232467 | -0.2367146 | 0.06398697 | -3.6994183 | 0.00021609 |
| 226169 | pprc1 | 3655.64123 | 0.25972254 | 0.07030866 | 3.69403346 | 0.00022072 |
| 78016 | CCDC150 | 5.12265718 | -4.2814223 | 1.15906467 | -3.6938597 | 0.00022088 |
| 226999 | SLC9A2 | 134.196838 | 3.38468844 | 0.9164362 | 3.69331596 | 0.00022135 |
| 20249 | Scd1 | 8676.63607 | 0.48105326 | 0.13025809 | 3.69307784 | 0.00022156 |
| 217039 | GGNBP2 | 2733.94178 | -0.1875922 | 0.05080252 | -3.6925772 | 0.00022199 |
| 104401 | PCNX3 | 3890.47304 | 0.49943425 | 0.13525976 | 3.69240813 | 0.00022214 |
| 320214 | MAATS1 | 36.0954118 | 2.14668378 | 0.58148922 | 3.69170003 | 0.00022276 |
| 171180 | SYT12 | 138.822053 | -2.5162733 | 0.68182567 | -3.6904937 | 0.00022382 |
| 15416 | HOXB8 | 27.0281662 | -7.457761 | 2.02101522 | -3.6901063 | 0.00022416 |
| 108159 | UBXN8 | 747.81101 | 0.33381389 | 0.09047872 | 3.68941889 | 0.00022477 |
| 81896 | IFT122 | 426.957517 | -0.9094388 | 0.24652739 | -3.688997 | 0.00022514 |
| 66082 | ABHD6 | 367.992194 | -0.9395633 | 0.25492164 | -3.6856947 | 0.00022808 |
| 66681 | PGM1 | 1222.84581 | 0.25820851 | 0.07005804 | 3.68563704 | 0.00022813 |
| 17122 | MXD4 | 249.361956 | -0.7517909 | 0.20400641 | -3.6851334 | 0.00022858 |
| 109346 | ankrd39 | 177.362576 | -0.5787546 | 0.15710378 | -3.6839002 | 0.00022969 |
| 22365 | VPS45 | 417.58534 | -0.3894387 | 0.105729 | -3.6833664 | 0.00023017 |

| | | | | | | |
|---|---|---|---|---|---|---|
| 70592 | 5730480H06 | 27.7577564 | -1.1900225 | 0.32309504 | -3.6831964 | 0.00023033 |
| 26424 | NR5A2 | 32.9198598 | -7.7451889 | 2.10300583 | -3.6829137 | 0.00023058 |
| 20558 | Slfn4 | 3.66335144 | -5.2958689 | 1.43805274 | -3.6826667 | 0.00023081 |
| 216860 | neurl4 | 1219.70534 | 0.33294097 | 0.09044657 | 3.68107886 | 0.00023225 |
| 70503 | DDO | 9.39172871 | -6.6638206 | 1.81045289 | -3.6807479 | 0.00023255 |
| 72640 | MEX3A | 803.385822 | 0.65164288 | 0.17705296 | 3.68049692 | 0.00023278 |
| 104215 | RHOQ | 4340.53744 | 0.35666388 | 0.09690853 | 3.68041776 | 0.00023285 |
| 240667 | SEC31B | 19.0188537 | -1.6001697 | 0.43481813 | -3.6800896 | 0.00023315 |
| 72175 | MFSD8 | 188.118343 | -0.5338857 | 0.1451328 | -3.6786011 | 0.00023452 |
| 70461 | CRTC3 | 835.343632 | 0.51913608 | 0.14114274 | 3.6780927 | 0.00023498 |
| 100036521 | Umad1 | 249.109716 | 0.53996504 | 0.14681647 | 3.67782332 | 0.00023523 |
| 56484 | FOXO3 | 1181.13839 | -0.4189435 | 0.11391137 | -3.6778021 | 0.00023525 |
| 78908 | igsf3 | 1120.6454 | 0.76506414 | 0.20804076 | 3.67747235 | 0.00023556 |
| 209378 | ITIH5 | 4.90614823 | -4.9572636 | 1.34816702 | -3.6770397 | 0.00023596 |
| 66809 | KRT20 | 4.85448119 | -4.9550139 | 1.34789811 | -3.6761042 | 0.00023682 |
| 12359 | CAT | 4613.80613 | 0.62183194 | 0.16917429 | 3.67568822 | 0.00023721 |
| 338369 | TMEM220 | 21.4523268 | -1.4044805 | 0.3821589 | -3.6751217 | 0.00023774 |
| 77583 | NOTUM | 255.319872 | -1.7205537 | 0.46824063 | -3.6745075 | 0.00023831 |
| 20280 | SCP2 | 513.410785 | 0.39059494 | 0.10630603 | 3.67425012 | 0.00023855 |
| 21809 | TGFB3 | 720.548453 | -0.7073901 | 0.19253572 | -3.6740722 | 0.00023872 |
| 94184 | PDXDC1 | 4133.55072 | -0.2411246 | 0.06563655 | -3.6736335 | 0.00023913 |
| 114872 | Psg29 | 4.98361612 | -5.7421918 | 1.56402221 | -3.671426 | 0.0002412 |
| 20224 | SAR1A | 5923.98018 | 0.13805921 | 0.03761186 | 3.67062919 | 0.00024195 |
| 74080 | NMNAT3 | 56.679614 | -0.7922999 | 0.21585717 | -3.6704823 | 0.00024209 |
| 56551 | TXN2 | 2851.75205 | -0.2012775 | 0.05484908 | -3.6696604 | 0.00024287 |
| 69305 | DCPS | 790.525655 | 0.27266429 | 0.07430818 | 3.66937109 | 0.00024315 |
| 26357 | ABCG2 | 886.022768 | -0.477279 | 0.13009805 | -3.6686101 | 0.00024387 |
| 66136 | ZNRD1 | 384.166583 | 0.60047722 | 0.16368638 | 3.66846172 | 0.00024401 |
| 72057 | PHF10 | 1218.88869 | 0.90009975 | 0.24537423 | 3.6682733 | 0.00024419 |
| 19664 | RBPJ | 1170.87243 | 0.80574548 | 0.21966779 | 3.66801838 | 0.00024444 |
| 100662 | D930016D06 | 225.242258 | -0.5020076 | 0.13686808 | -3.6678209 | 0.00024463 |
| 14787 | RHPN1 | 24.8313473 | -2.6200671 | 0.71439017 | -3.6675576 | 0.00024488 |
| 110083 | DNAH12 | 10.1149437 | -4.2414584 | 1.15663586 | -3.6670646 | 0.00024535 |
| 15159 | HCCS | 1177.6964 | 0.27885501 | 0.0760524 | 3.66661666 | 0.00024578 |
| 12192 | ZFP36L1 | 2060.44248 | 0.3333081 | 0.09091125 | 3.66630206 | 0.00024608 |
| 68441 | RRAGA | 2229.97612 | 0.39282558 | 0.10719387 | 3.66462714 | 0.0002477 |
| 225182 | RBBP8 | 1395.91406 | 0.42192328 | 0.11515055 | 3.66410121 | 0.00024821 |
| 66276 | 1810009A15 | 1016.31742 | 0.47047778 | 0.12843394 | 3.66318895 | 0.00024909 |
| 13194 | DDB1 | 36943.2478 | -0.2848842 | 0.07778618 | -3.6624011 | 0.00024986 |
| 74453 | CFAP53 | 6.64499107 | -4.7040427 | 1.28472838 | -3.6615076 | 0.00025074 |
| 80888 | HSPB8 | 3057.62459 | 2.81156513 | 0.7678767 | 3.66147995 | 0.00025076 |
| 208263 | TOR1AIP1 | 2860.68344 | -0.2402095 | 0.06561116 | -3.6611078 | 0.00025113 |
| 94315 | PRCC | 1591.49841 | 0.27821723 | 0.0760026 | 3.66062765 | 0.0002516 |
| 78903 | WRNIP1 | 1762.96529 | 0.43929377 | 0.12001778 | 3.66023898 | 0.00025198 |
| 22184 | zrsr2 | 1176.51233 | 0.29601488 | 0.08088336 | 3.65977497 | 0.00025244 |

| | | | | | | |
|---|---|---|---|---|---|---|
| 16675 | KRT27 | 3.4676119 | -5.2200917 | 1.42649701 | -3.6593779 | 0.00025283 |
| 73420 | CCSAP | 453.505464 | 0.68090147 | 0.18609308 | 3.65892954 | 0.00025327 |
| 170472 | RECQL5 | 572.254752 | -0.4495112 | 0.12285789 | -3.65879 | 0.00025341 |
| 15526 | HSPA9 | 22336.298 | -0.1577193 | 0.04311013 | -3.6585216 | 0.00025367 |
| 106894 | HMGXB3 | 2129.26931 | 0.30267853 | 0.08275018 | 3.65773855 | 0.00025445 |
| 242083 | PPM1L | 667.510455 | 1.07776091 | 0.29466602 | 3.65756766 | 0.00025462 |
| 19347 | DENND5A | 3888.905 | 0.39211038 | 0.10722658 | 3.65683952 | 0.00025534 |
| 72482 | ACBD6 | 1329.75055 | -0.1646176 | 0.04502616 | -3.656044 | 0.00025614 |
| 71856 | WFDC3 | 13.6448525 | -4.3651657 | 1.19403365 | -3.6558146 | 0.00025637 |
| 100504344 | Gm20186 | 29.0680687 | -1.1880217 | 0.32497987 | -3.6556778 | 0.0002565 |
| 55982 | PAXIP1 | 1980.44869 | 0.30053924 | 0.08223015 | 3.65485448 | 0.00025733 |
| 211660 | CSPP1 | 578.590837 | -0.3895243 | 0.10658706 | -3.6545175 | 0.00025767 |
| 74359 | 4931414P19I | 102.340876 | 0.70259003 | 0.19226817 | 3.65421913 | 0.00025797 |
| 21814 | TGFBR3 | 2164.42524 | 1.4696624 | 0.40222097 | 3.65386815 | 0.00025832 |
| 72307 | 2510002D24 | 169.432158 | -0.8882519 | 0.24314411 | -3.6531911 | 0.000259 |
| 64540 | TSPAN4 | 3755.76463 | 0.60088276 | 0.16453492 | 3.65200743 | 0.0002602 |
| 232807 | PPP1R12C | 1634.77807 | 0.5945057 | 0.16281084 | 3.65151175 | 0.0002607 |
| 619289 | RFX8 | 4.03019383 | -4.6706405 | 1.27938154 | -3.6507018 | 0.00026152 |
| 232811 | KMT5C | 331.05443 | 0.4945851 | 0.13548388 | 3.65050895 | 0.00026172 |
| 218343 | TTC37 | 1813.85276 | 0.49450369 | 0.13546644 | 3.6503778 | 0.00026185 |
| 107734 | MRPL30 | 1110.54996 | -0.2989811 | 0.08193684 | -3.6489215 | 0.00026334 |
| 83946 | PHIP | 3502.02793 | 0.55064446 | 0.15094676 | 3.64793828 | 0.00026435 |
| 14013 | MECOM | 45.1035928 | 1.04047068 | 0.28526139 | 3.64742904 | 0.00026488 |
| 19734 | RGS16 | 3095.21231 | 0.76656379 | 0.21018273 | 3.6471303 | 0.00026519 |
| 23830 | CAPN10 | 524.029318 | -0.2640786 | 0.07242155 | -3.6464094 | 0.00026593 |
| 20170 | HPS6 | 561.435879 | 0.47804174 | 0.13113723 | 3.64535477 | 0.00026702 |
| 22417 | WNT4 | 10.2977851 | -2.8435525 | 0.78030588 | -3.644151 | 0.00026828 |
| 66120 | FKBP11 | 402.551752 | 0.97465508 | 0.26745758 | 3.64414825 | 0.00026828 |
| 434325 | TMEM221 | 6.98061708 | -5.4868561 | 1.50627281 | -3.6426709 | 0.00026982 |
| 22295 | CDH23 | 11.77918 | -2.6819294 | 0.73638471 | -3.642022 | 0.0002705 |
| 14897 | TRIP12 | 12372.2276 | -0.3232068 | 0.08874963 | -3.6417815 | 0.00027076 |
| 72735 | ELDR | 12.1377126 | -2.5150549 | 0.69061503 | -3.6417611 | 0.00027078 |
| 13803 | ENC1 | 4571.85109 | 0.5449232 | 0.14966472 | 3.64095951 | 0.00027162 |
| 20682 | SOX9 | 5293.42506 | 0.43285248 | 0.11889717 | 3.64056157 | 0.00027204 |
| 225055 | FBXO11 | 2587.12107 | 0.26269801 | 0.0722319 | 3.63686956 | 0.00027597 |
| 239463 | FAM83A | 151.622456 | -9.9517014 | 2.73667067 | -3.6364264 | 0.00027645 |
| 237073 | RBM41 | 672.098498 | -0.3082455 | 0.08477446 | -3.6360661 | 0.00027683 |
| 104009 | qsox1 | 3113.68138 | 0.35745566 | 0.09833308 | 3.63515152 | 0.00027782 |
| 20199 | S100A5 | 4.06219602 | 4.73439395 | 1.30244721 | 3.63499874 | 0.00027798 |
| 12544 | CDC45 | 1345.69404 | 0.2500728 | 0.06879588 | 3.63499683 | 0.00027798 |
| 17764 | MTF1 | 817.789748 | 0.4666205 | 0.12838212 | 3.6346222 | 0.00027839 |
| 15413 | HOXB5 | 27.2480116 | -8.1976819 | 2.25548714 | -3.6345505 | 0.00027847 |
| 22143 | TUBA1B | 1795.91139 | 0.40689277 | 0.11195239 | 3.63451621 | 0.0002785 |
| 13134 | DACH1 | 3.89079784 | -5.3849223 | 1.48175982 | -3.6341398 | 0.00027891 |
| 209354 | EIF2B1 | 1553.03964 | 0.41500524 | 0.11420036 | 3.63400997 | 0.00027905 |

| | | | | | | |
|---|---|---|---|---|---|---|
| 93759 | SIRT1 | 1262.58755 | 0.31995561 | 0.08806836 | 3.63303687 | 0.0002801 |
| 12631 | CFL1 | 15760.5221 | -0.2287289 | 0.06296517 | -3.6326252 | 0.00028055 |
| 66241 | TMEM9 | 1053.96263 | -0.5525201 | 0.15212323 | -3.632056 | 0.00028117 |
| 81702 | ANKRD17 | 9223.13479 | 0.27982655 | 0.0770441 | 3.63203079 | 0.0002812 |
| 93717 | PCDHGA9 | 8.2141412 | -3.6282713 | 0.99900485 | -3.6318856 | 0.00028136 |
| 67074 | MON2 | 3362.69659 | -0.1551346 | 0.04271991 | -3.6314354 | 0.00028185 |
| 14129 | Fcgr1 | 8.03011434 | -6.4327295 | 1.77148397 | -3.631266 | 0.00028203 |
| 239706 | METTL22 | 132.213272 | -0.586904 | 0.16165359 | -3.6306273 | 0.00028273 |
| 22035 | TNFSF10 | 5.72484039 | -3.8297803 | 1.05501349 | -3.630077 | 0.00028334 |
| 109275 | ACTR5 | 327.857862 | 0.39915546 | 0.11001071 | 3.62833286 | 0.00028526 |
| 12667 | CHRD | 51.1397726 | -2.8585968 | 0.787989 | -3.6277115 | 0.00028594 |
| 70788 | KLHL30 | 68.0993815 | -2.4740273 | 0.68211815 | -3.6269776 | 0.00028676 |
| 51810 | HNRNPU | 28100.4743 | 0.30416844 | 0.08386667 | 3.6268094 | 0.00028694 |
| 75116 | 4930520O04 | 19.8483318 | 1.35499108 | 0.37361905 | 3.62666489 | 0.00028711 |
| 70432 | RUFY2 | 737.93623 | -0.4980799 | 0.13734547 | -3.626475 | 0.00028732 |
| 14999 | H2-DMb1 | 9.53768247 | -3.1014785 | 0.85537654 | -3.6258634 | 0.000288 |
| 68041 | MID1IP1 | 2030.81387 | 0.36205581 | 0.09985906 | 3.625668 | 0.00028822 |
| 78825 | DESI2 | 1876.74766 | 0.29451141 | 0.08123973 | 3.62521409 | 0.00028872 |
| 212547 | NEPRO | 1130.21568 | 0.34652626 | 0.09559253 | 3.62503509 | 0.00028892 |
| 27973 | vkorc1 | 347.012297 | 0.78169552 | 0.21564451 | 3.62492663 | 0.00028904 |
| 17448 | MDH2 | 9183.6119 | -0.2081262 | 0.05742065 | -3.624588 | 0.00028942 |
| 17534 | mrc2 | 4032.77383 | 3.55969802 | 0.98220562 | 3.6241882 | 0.00028987 |
| 232371 | C1RL | 69.5968314 | -0.8866254 | 0.24464312 | -3.6241584 | 0.0002899 |
| 227929 | CYTIP | 207.183764 | 2.95440276 | 0.81530655 | 3.62367106 | 0.00029045 |
| 112403 | DXO | 391.78525 | -0.344405 | 0.09505742 | -3.6231256 | 0.00029106 |
| 414068 | Bach2os | 3.65902691 | -5.2948491 | 1.4615032 | -3.6228789 | 0.00029134 |
| 232337 | Zfp637 | 647.940431 | 0.84069597 | 0.23207009 | 3.62259511 | 0.00029166 |
| 20750 | SPP1 | 27671.5319 | -0.8009383 | 0.22110057 | -3.6225066 | 0.00029176 |
| 12389 | CAV1 | 8640.60754 | -0.9557305 | 0.26384989 | -3.6222508 | 0.00029205 |
| 67723 | Cep83os | 134.075611 | -0.7593817 | 0.209675 | -3.6217083 | 0.00029266 |
| 67230 | Zfp329 | 392.984269 | -0.4466292 | 0.12332278 | -3.6216279 | 0.00029276 |
| 101739 | PSIP1 | 2009.42023 | 0.44700361 | 0.12344527 | 3.62106706 | 0.00029339 |
| 74365 | LONRF3 | 1112.65901 | -0.8289708 | 0.22894045 | -3.6209014 | 0.00029358 |
| 227715 | EXOSC2 | 1487.09753 | 0.32838233 | 0.09069846 | 3.62059421 | 0.00029393 |
| 73068 | FUT11 | 473.66204 | 0.70983495 | 0.19609827 | 3.61979202 | 0.00029484 |
| 68099 | FAM92A | 939.249334 | -0.7980011 | 0.22047238 | -3.6195059 | 0.00029517 |
| 12091 | GLB1 | 1317.23602 | 0.35648997 | 0.09852029 | 3.61844236 | 0.00029638 |
| 73942 | FAM151B | 49.5879889 | -1.6551515 | 0.4574512 | -3.6182035 | 0.00029666 |
| 69072 | EBNA1BP2 | 2172.22648 | 0.74456949 | 0.20586877 | 3.61671895 | 0.00029836 |
| 675812 | Zfp605 | 528.106227 | -0.4964862 | 0.13738517 | -3.6138264 | 0.00030171 |
| 20182 | RXRB | 1291.1512 | 0.45650045 | 0.12632959 | 3.61356719 | 0.00030201 |
| 624662 | Gm6517 | 693.350256 | -0.41149 | 0.11389381 | -3.612927 | 0.00030276 |
| 70638 | FAM189A1 | 9.49902686 | -5.2099739 | 1.44213346 | -3.612685 | 0.00030304 |
| 67767 | JAGN1 | 1161.81085 | 0.28301565 | 0.0783436 | 3.61249253 | 0.00030327 |
| 70804 | PGRMC2 | 2851.33885 | 0.16015033 | 0.0443425 | 3.611667 | 0.00030424 |

| | | | | | | |
|---|---|---|---|---|---|---|
| 58249 | FIBP | 861.465823 | -0.5210588 | 0.14429828 | -3.6109839 | 0.00030504 |
| 58859 | EFEMP2 | 3249.61979 | 1.62508593 | 0.45013599 | 3.6102111 | 0.00030595 |
| 16149 | CD74 | 5.50596867 | -5.888866 | 1.63190302 | -3.6085882 | 0.00030787 |
| 66915 | Myeov2 | 678.500127 | -0.4879328 | 0.13523206 | -3.6081145 | 0.00030843 |
| 19698 | RELB | 734.095965 | -0.3993749 | 0.11071651 | -3.6071847 | 0.00030954 |
| 18117 | EMC8 | 1697.71749 | -0.337499 | 0.09356931 | -3.6069411 | 0.00030983 |
| 60455 | Tmem8 | 196.824415 | -0.4177709 | 0.1159004 | -3.6045683 | 0.00031267 |
| 77697 | MMAB | 480.285514 | 0.38263002 | 0.10616985 | 3.60394248 | 0.00031343 |
| 66805 | TSPAN1 | 3.51430477 | -5.2374475 | 1.45327378 | -3.603896 | 0.00031348 |
| 20747 | spop | 3900.67277 | 0.37030086 | 0.10275377 | 3.60376923 | 0.00031364 |
| 76709 | ARPC2 | 8302.87613 | -0.3612876 | 0.10026307 | -3.6033969 | 0.00031409 |
| 11991 | HNRNPD | 7888.88785 | 0.35013701 | 0.0971806 | 3.60295161 | 0.00031462 |
| 68010 | BAMBI | 5.9862742 | -4.5445442 | 1.26134147 | -3.6029452 | 0.00031463 |
| 668212 | EFR3B | 299.268607 | -2.8117979 | 0.78042878 | -3.6028885 | 0.0003147 |
| 66152 | Uqcr10 | 1213.34115 | -0.2710636 | 0.07525233 | -3.6020628 | 0.0003157 |
| 20971 | SDC4 | 10482.5689 | -0.4417072 | 0.12263856 | -3.6016992 | 0.00031614 |
| 17719 | ND4 | 21912.3551 | -0.6921585 | 0.19221124 | -3.6010301 | 0.00031696 |
| 57436 | GABARAPL1 | 2402.34937 | 0.58042221 | 0.16118889 | 3.60088219 | 0.00031714 |
| 70385 | SPDL1 | 1247.01013 | 0.42790291 | 0.11884284 | 3.60057807 | 0.00031751 |
| 56486 | GABARAP | 5042.77799 | -0.4529747 | 0.12581403 | -3.600351 | 0.00031779 |
| 66632 | DPH6 | 1242.56713 | 0.30582862 | 0.08494492 | 3.60031658 | 0.00031783 |
| 209387 | Trim30d | 8.16679556 | -6.4572803 | 1.79400413 | -3.5993676 | 0.00031899 |
| 22185 | U2AF2 | 4101.51915 | 0.40762921 | 0.11328178 | 3.59836508 | 0.00032022 |
| 218476 | GCNT4 | 831.903249 | 2.34056641 | 0.65050415 | 3.59808067 | 0.00032057 |
| 67749 | mgarp | 7.79398445 | -3.000077 | 0.83380386 | -3.5980608 | 0.0003206 |
| 53620 | VAMP5 | 241.091192 | -0.7101182 | 0.19738801 | -3.5975749 | 0.0003212 |
| 235345 | 4833427G06 | 21.6300528 | -1.6223431 | 0.45097455 | -3.5974162 | 0.00032139 |
| 76000 | 5033430I15R | 42.8970793 | 1.22430096 | 0.34047873 | 3.59582218 | 0.00032337 |
| 74574 | LVRN | 37.121269 | -1.3613407 | 0.37873386 | -3.5944521 | 0.00032507 |
| 226041 | PGM5 | 5.26684536 | -4.3219393 | 1.20253517 | -3.5940232 | 0.00032561 |
| 71955 | IST1 | 2251.2497 | -0.2834036 | 0.07885848 | -3.5938249 | 0.00032586 |
| 23966 | TENM4 | 390.090459 | 2.76315306 | 0.76897039 | 3.59331531 | 0.0003265 |
| 18830 | PLTP | 335.092977 | -2.4864835 | 0.69201622 | -3.5931001 | 0.00032677 |
| 17749 | POLR2K | 26.17207 | 1.13562932 | 0.31605887 | 3.59309427 | 0.00032677 |
| 68550 | TEFM | 219.704634 | -0.3837049 | 0.10680474 | -3.5925835 | 0.00032742 |
| 69694 | TATDN1 | 374.029772 | 0.44815226 | 0.12476149 | 3.59207201 | 0.00032806 |
| 214523 | TMPRSS4 | 10.5615112 | -6.8293268 | 1.90144531 | -3.5916504 | 0.00032859 |
| 74377 | HSF2BP | 17.8340432 | -2.258322 | 0.62877918 | -3.591598 | 0.00032866 |
| 68080 | gpn3 | 807.384857 | 0.29359589 | 0.0817524 | 3.59128146 | 0.00032906 |
| 20222 | SF3A2 | 2566.80962 | 0.35532644 | 0.09895088 | 3.59093778 | 0.00032949 |
| 99311 | COMMD7 | 867.581502 | 0.37254637 | 0.10376015 | 3.59045725 | 0.0003301 |
| 320571 | Atp8b5 | 8.58554653 | -3.1153242 | 0.86784896 | -3.5897078 | 0.00033105 |
| 378460 | PRAM1 | 9.28043038 | -2.3312484 | 0.64955537 | -3.5889911 | 0.00033196 |
| 59044 | RNF130 | 1057.11477 | -0.5290614 | 0.14748007 | -3.5873417 | 0.00033407 |
| 229096 | YTHDF3 | 4305.89936 | -0.2125746 | 0.05926448 | -3.5868805 | 0.00033466 |

| | | | | | | |
|---|---|---|---|---|---|---|
| 225998 | RORB | 9.62388315 | -4.1651325 | 1.16121954 | -3.5868605 | 0.00033468 |
| 68339 | CCDC88C | 231.329481 | -1.1364607 | 0.31689971 | -3.586184 | 0.00033555 |
| 17144 | MAGEA8 | 3.35736664 | 5.22962376 | 1.45827307 | 3.5861759 | 0.00033556 |
| 240427 | SETBP1 | 413.00874 | 2.5411207 | 0.70875477 | 3.58533135 | 0.00033665 |
| 13885 | ESD | 11239.7618 | 0.95940869 | 0.26759992 | 3.58523532 | 0.00033677 |
| 56395 | TMEM115 | 1342.80875 | 0.28600245 | 0.07977804 | 3.58497727 | 0.00033711 |
| 319996 | CASC4 | 561.1003 | -0.8201684 | 0.22879005 | -3.584808 | 0.00033733 |
| 74919 | SLC35F6 | 1590.94892 | -0.4162847 | 0.1161264 | -3.5847548 | 0.0003374 |
| 381510 | DPY19L4 | 1033.56024 | -0.9262334 | 0.25838736 | -3.58467 | 0.0003375 |
| 14693 | GNB2 | 9627.02418 | -0.2299493 | 0.0641679 | -3.583557 | 0.00033895 |
| 30927 | SNAI3 | 7.41979098 | -3.7867914 | 1.057316 | -3.5815134 | 0.00034161 |
| 69035 | ZDHHC3 | 3126.89588 | 0.42883209 | 0.11974021 | 3.58135409 | 0.00034182 |
| 20316 | SDF2 | 966.024253 | -0.3086663 | 0.08618926 | -3.5812614 | 0.00034194 |
| 101867 | RRP8 | 1623.93179 | 0.79783104 | 0.22278626 | 3.58115005 | 0.00034209 |
| 12462 | CCT3 | 10274.6134 | 0.23073266 | 0.06444207 | 3.58046603 | 0.00034298 |
| 338366 | MIA3 | 3930.17391 | 0.230373 | 0.06434289 | 3.58039536 | 0.00034307 |
| 216987 | UTP6 | 3115.30945 | 0.25972011 | 0.07254643 | 3.58005361 | 0.00034352 |
| 263803 | PKN3 | 377.886959 | 0.99188588 | 0.27710746 | 3.57942682 | 0.00034435 |
| 17149 | magoh | 1115.80633 | 0.68710029 | 0.19210184 | 3.57675008 | 0.00034789 |
| 67876 | COQ10B | 982.957372 | -0.3616924 | 0.10118753 | -3.5744757 | 0.00035093 |
| 20678 | SOX5 | 199.664459 | -1.104313 | 0.30896559 | -3.5742263 | 0.00035126 |
| 214987 | CHTF8 | 3962.41491 | 0.28597037 | 0.08001426 | 3.57399251 | 0.00035158 |
| 11990 | ATRN | 2790.90094 | -0.3911774 | 0.1094672 | -3.5734667 | 0.00035229 |
| 100042049 | Gm15421 | 323.615428 | -0.6294785 | 0.17619763 | -3.5725708 | 0.00035349 |
| 20128 | Trim30a | 331.784338 | -2.8057769 | 0.78551298 | -3.571904 | 0.0003544 |
| 17258 | MEF2A | 1724.74001 | 0.41018612 | 0.11485518 | 3.57133316 | 0.00035517 |
| 191578 | HELQ | 776.869804 | -0.287599 | 0.08053148 | -3.5712621 | 0.00035527 |
| 625530 | Usp17le | 10.5011201 | -4.3390724 | 1.2153107 | -3.57034 | 0.00035652 |
| 22217 | USP12 | 1576.05503 | 0.46759058 | 0.13098115 | 3.56990751 | 0.00035711 |
| 237911 | BRIP1 | 1008.15338 | -0.3136207 | 0.08785328 | -3.5698238 | 0.00035722 |
| 100855 | TBC1D14 | 1314.78366 | -0.3161748 | 0.08858615 | -3.5691221 | 0.00035818 |
| 72692 | HNRNPLL | 3360.71072 | 0.2918355 | 0.08177399 | 3.56880588 | 0.00035861 |
| 21683 | TECTA | 6.70787138 | 4.1410155 | 1.16070133 | 3.56768394 | 0.00036015 |
| 216019 | HKDC1 | 8.50650043 | 2.92361422 | 0.81950294 | 3.56754574 | 0.00036034 |
| 16795 | LARGE1 | 1673.1386 | -0.2560541 | 0.0717788 | -3.5672659 | 0.00036073 |
| 394432 | Ugt1a7c | 31.3164028 | 3.2134008 | 0.90089375 | 3.56690321 | 0.00036122 |
| 230700 | FOXJ3 | 1478.75811 | 0.55485571 | 0.15556013 | 3.56682472 | 0.00036133 |
| 105203 | fam208b | 2960.89371 | 0.36973564 | 0.10367122 | 3.56642513 | 0.00036188 |
| 70791 | HARS2 | 693.955201 | -0.2337697 | 0.06554744 | -3.5664196 | 0.00036189 |
| 241846 | LSM14B | 2377.91119 | 0.37727545 | 0.10578687 | 3.56637309 | 0.00036196 |
| 52700 | TXNDC17 | 1919.38027 | -0.3127677 | 0.08771173 | -3.5658597 | 0.00036267 |
| 225392 | RELL2 | 33.0871529 | -1.3828155 | 0.38780416 | -3.5657572 | 0.00036281 |
| 100861649 | #N/A | 12.6911673 | 1.75775666 | 0.49299455 | 3.56546877 | 0.00036321 |
| 68718 | RNF166 | 529.067587 | -0.3406719 | 0.09559708 | -3.5636228 | 0.00036577 |
| 17123 | MADCAM1 | 6.33386128 | -6.0894788 | 1.70915545 | -3.5628583 | 0.00036684 |

| | | | | | | |
|---|---|---|---|---|---|---|
| 56857 | SLC37A2 | 62.6036606 | -1.3837491 | 0.38838518 | -3.5628268 | 0.00036688 |
| 433619 | KPRP | 6.53152104 | 3.32323554 | 0.93289233 | 3.56229272 | 0.00036763 |
| 240960 | dnah14 | 3.59982343 | -5.279232 | 1.48224439 | -3.5616475 | 0.00036854 |
| 56398 | CHP1 | 2033.24472 | 0.40284266 | 0.11312709 | 3.56097432 | 0.00036948 |
| 72947 | PHYKPL | 269.450054 | -0.425007 | 0.11936053 | -3.5606998 | 0.00036987 |
| 208908 | CCDC62 | 22.9356174 | -1.4876627 | 0.4178033 | -3.5606773 | 0.0003699 |
| 76816 | SDCCAG8 | 313.325594 | -0.5416197 | 0.1521533 | -3.5596974 | 0.00037128 |
| 99982 | KDM1A | 3082.39405 | 0.7201777 | 0.20232647 | 3.55948328 | 0.00037159 |
| 70470 | RPRD1B | 1459.5496 | 0.41007396 | 0.1152109 | 3.55933302 | 0.0003718 |
| 105243 | SLC9A3 | 3.99987571 | -5.4342653 | 1.52683437 | -3.5591714 | 0.00037203 |
| 100502819 | BC065403 | 7.79487512 | 2.99348249 | 0.84117582 | 3.55868821 | 0.00037271 |
| 237979 | SDK2 | 3.86745938 | -4.626501 | 1.30019496 | -3.5583133 | 0.00037324 |
| 238037 | BC068281 | 185.816548 | 0.68411837 | 0.19229272 | 3.55769252 | 0.00037413 |
| 56353 | RYBP | 367.730498 | 0.5965239 | 0.16774401 | 3.5561562 | 0.00037632 |
| 27053 | ASNS | 4364.917 | 0.4755383 | 0.13375338 | 3.55533681 | 0.0003775 |
| 12583 | CDO1 | 20.6202149 | -2.662089 | 0.7489774 | -3.5542982 | 0.00037899 |
| 53625 | B3GNT2 | 2185.75166 | -0.6666041 | 0.1875567 | -3.5541471 | 0.00037921 |
| 241035 | PKHD1 | 3.40985112 | -5.1923777 | 1.46137924 | -3.5530666 | 0.00038077 |
| 108671 | DNAJC9 | 1657.66098 | -0.2623848 | 0.07386333 | -3.5523009 | 0.00038188 |
| 94230 | CPSF1 | 3234.9178 | -0.2605713 | 0.07335362 | -3.5522615 | 0.00038194 |
| 246196 | Zfp277 | 699.958238 | 0.27445595 | 0.07726475 | 3.5521495 | 0.0003821 |
| 330286 | D630045J12F | 1256.75059 | 0.55809921 | 0.15711842 | 3.55209275 | 0.00038218 |
| 21406 | TCF12 | 3469.63043 | 0.23338491 | 0.06571493 | 3.55147473 | 0.00038308 |
| 16918 | MYCL | 37.9732659 | -3.5264542 | 0.99321473 | -3.5505456 | 0.00038443 |
| 13163 | DAXX | 781.584715 | -0.2416395 | 0.06807047 | -3.5498441 | 0.00038546 |
| 666173 | VPS13B | 1912.74846 | 0.47129974 | 0.13277097 | 3.5497198 | 0.00038564 |
| 546143 | Ccpg1os | 27.6361716 | -1.5020258 | 0.42321243 | -3.5491061 | 0.00038654 |
| 13929 | AMZ2 | 825.289523 | -0.2654531 | 0.07479564 | -3.5490447 | 0.00038663 |
| 68095 | OCIAD1 | 4061.73801 | -0.2078736 | 0.05858351 | -3.5483302 | 0.00038768 |
| 70088 | MEAF6 | 317.446527 | 0.8686454 | 0.24481163 | 3.54821948 | 0.00038784 |
| 380711 | RAP1GAP2 | 787.823103 | 1.89421793 | 0.53385864 | 3.54816386 | 0.00038793 |
| 11409 | ACADS | 925.121994 | -0.3218384 | 0.09071276 | -3.5478844 | 0.00038834 |
| 17857 | MX1 | 4.18117914 | -4.7365228 | 1.33509732 | -3.5476986 | 0.00038861 |
| 106205 | ZC3H7A | 2136.07609 | -0.3537571 | 0.0997151 | -3.5476783 | 0.00038864 |
| 14852 | GSPT1 | 10408.9213 | 0.16574058 | 0.0467304 | 3.54674022 | 0.00039003 |
| 268949 | DPCR1 | 7.67718595 | 4.91170541 | 1.3848628 | 3.54670905 | 0.00039008 |
| 53972 | NGEF | 634.468215 | 2.43497607 | 0.68670971 | 3.54585938 | 0.00039133 |
| 68021 | BPHL | 185.280768 | -0.6930586 | 0.19548184 | -3.545386 | 0.00039204 |
| 116701 | FGFRL1 | 919.592342 | 0.2334308 | 0.06584645 | 3.54507783 | 0.0003925 |
| 64176 | SV2B | 8.65707869 | -5.0405967 | 1.42227942 | -3.544027 | 0.00039406 |
| 72935 | DDX41 | 1747.06589 | 0.32801581 | 0.09256508 | 3.54362374 | 0.00039467 |
| 100678 | PSPH | 1128.01837 | 0.21942536 | 0.06194283 | 3.54238524 | 0.00039653 |
| 76972 | SNHG20 | 125.818406 | -0.6812632 | 0.1924068 | -3.5407437 | 0.000399 |
| 192885 | BC005512 | 3.78523982 | -4.5923015 | 1.29706939 | -3.5405211 | 0.00039934 |
| 381356 | CACFD1 | 602.925106 | -0.4829392 | 0.13641659 | -3.5401796 | 0.00039985 |

| | | | | | | |
|---|---|---|---|---|---|---|
| 544752 | Tug1 | 3659.92324 | -0.2134828 | 0.06032756 | -3.5387276 | 0.00040206 |
| 26447 | POLI | 158.414694 | -1.0795023 | 0.30511146 | -3.538059 | 0.00040308 |
| 52335 | ATXN1L | 2965.5395 | -0.170177 | 0.04810032 | -3.53796 | 0.00040323 |
| 24045 | SCAMP3 | 2006.68987 | -0.4264663 | 0.12054459 | -3.5378304 | 0.00040343 |
| 15937 | IER3 | 5026.88775 | 0.75200914 | 0.21263067 | 3.53669178 | 0.00040517 |
| 98267 | STK17B | 1206.68321 | -0.2683556 | 0.0758782 | -3.5366627 | 0.00040522 |
| 11863 | ARNT | 1575.09165 | 0.24515652 | 0.06932523 | 3.53632486 | 0.00040574 |
| 81910 | RRBP1 | 7536.26028 | 0.39866145 | 0.1127408 | 3.53608865 | 0.0004061 |
| 171429 | slc26a6 | 143.856095 | 1.06271003 | 0.30053313 | 3.53608274 | 0.00040611 |
| 14824 | GRN | 12777.1137 | -1.0330794 | 0.29220537 | -3.5354565 | 0.00040707 |
| 68607 | serhl | 247.417948 | -0.4138883 | 0.11707206 | -3.5353297 | 0.00040727 |
| 80284 | SMIM12 | 491.055191 | 0.63226943 | 0.17884688 | 3.53525556 | 0.00040738 |
| 231830 | MICALL2 | 3155.23926 | 0.47931158 | 0.13559575 | 3.53485697 | 0.000408 |
| 216188 | ALDH1L2 | 2573.66582 | 0.89694098 | 0.25378498 | 3.53425556 | 0.00040893 |
| 57814 | KCNE4 | 4.0813783 | -5.4607316 | 1.5454933 | -3.533326 | 0.00041037 |
| 105348 | GOLM1 | 889.669982 | 0.37662119 | 0.1065938 | 3.53323743 | 0.0004105 |
| 51869 | RIF1 | 3585.03994 | -0.3031509 | 0.08580387 | -3.5330681 | 0.00041077 |
| 68040 | Zfp593 | 619.435126 | 0.66180766 | 0.18732433 | 3.53295096 | 0.00041095 |
| 140742 | SESN1 | 296.146363 | 0.64576908 | 0.18281445 | 3.53237435 | 0.00041185 |
| 208117 | APH1B | 58.6780513 | -0.9902744 | 0.28036419 | -3.5321002 | 0.00041227 |
| 545481 | ARHGAP40 | 3.80570701 | -4.5904421 | 1.30012125 | -3.53078 | 0.00041434 |
| 320844 | amigo3 | 41.7196591 | 0.89123134 | 0.25243947 | 3.53047538 | 0.00041481 |
| 71583 | 9130008F23I | 155.870722 | -1.4488428 | 0.41038759 | -3.5304255 | 0.00041489 |
| 66394 | NOSIP | 1155.24423 | 0.43456527 | 0.12310902 | 3.52992238 | 0.00041568 |
| 11810 | APOBEC1 | 118.563288 | -1.7421366 | 0.49359023 | -3.5295199 | 0.00041631 |
| 271005 | KLHDC1 | 132.597265 | -0.8692868 | 0.24636206 | -3.5284928 | 0.00041793 |
| 70523 | 5730420D15 | 4.75860892 | -4.9245126 | 1.39574914 | -3.5282218 | 0.00041836 |
| 102334 | ANKRD10 | 1396.46988 | 0.24860874 | 0.07048225 | 3.52725299 | 0.0004199 |
| 66790 | GRTP1 | 39.1479791 | -2.5934005 | 0.73526234 | -3.5271771 | 0.00042002 |
| 52563 | CDC23 | 1793.56937 | 0.20030706 | 0.05679388 | 3.52691292 | 0.00042044 |
| 116940 | TGS1 | 911.874875 | -0.7560759 | 0.2143895 | -3.5266462 | 0.00042086 |
| 22419 | WNT5B | 47.7798452 | -3.2201569 | 0.91327861 | -3.5259305 | 0.000422 |
| 67733 | ITGB3BP | 53.2992876 | -0.8678204 | 0.24615612 | -3.5254877 | 0.0004227 |
| 76551 | CCDC6 | 4206.18421 | 0.21768261 | 0.06176729 | 3.52423739 | 0.0004247 |
| 215445 | RAB11FIP3 | 1278.16962 | 0.56654143 | 0.16077925 | 3.5237222 | 0.00042553 |
| 234309 | CBR4 | 175.624264 | -0.744021 | 0.21121988 | -3.5224954 | 0.0004275 |
| 100163 | PAFAH2 | 965.691276 | 0.37937806 | 0.10770984 | 3.52222299 | 0.00042794 |
| 791329 | Gm9968 | 14.2817857 | 1.76113378 | 0.50004519 | 3.52194927 | 0.00042839 |
| 209815 | TBC1D25 | 446.384344 | -0.3006229 | 0.08537192 | -3.5213328 | 0.00042938 |
| 240067 | Zfp952 | 158.145265 | -0.4194619 | 0.11915363 | -3.5203454 | 0.00043099 |
| 237107 | GNL3L | 4624.38719 | 0.27029727 | 0.07679275 | 3.51982779 | 0.00043183 |
| 22092 | rsph1 | 6.08820983 | -5.2869264 | 1.50227645 | -3.5192766 | 0.00043273 |
| 233210 | PRR12 | 2224.06128 | 0.29015062 | 0.08244981 | 3.51911805 | 0.00043298 |
| 692132 | TRAPPC3L | 4.77586134 | -4.920799 | 1.39836081 | -3.5189766 | 0.00043321 |
| 19766 | RIPK1 | 1793.16091 | 0.25733354 | 0.07313282 | 3.51871476 | 0.00043364 |

| | | | | | | |
|---|---|---|---|---|---|---|
| 215814 | CCDC28A | 74.0433886 | -0.8351443 | 0.2374975 | -3.516434 | 0.00043739 |
| 26427 | CREB3L1 | 90.7414043 | -1.7974463 | 0.5112044 | -3.5161009 | 0.00043793 |
| 80517 | HERPUD2 | 1040.49591 | 0.25844649 | 0.07351789 | 3.51542321 | 0.00043905 |
| 12994 | CSN3 | 3.96314667 | -5.4098583 | 1.53902431 | -3.5151221 | 0.00043955 |
| 227743 | MAPKAP1 | 1417.46288 | -0.2500174 | 0.07113179 | -3.5148478 | 0.00044001 |
| 217707 | COQ6 | 418.421559 | 0.32996726 | 0.09388482 | 3.51459631 | 0.00044042 |
| 74302 | MTMR3 | 2097.502 | -0.26459 | 0.0752898 | -3.5142877 | 0.00044093 |
| 59032 | PPP2R3C | 449.307541 | 0.39294189 | 0.11188075 | 3.51214923 | 0.0004445 |
| 633395 | Gm10548 | 10.0049353 | -1.8731772 | 0.53335313 | -3.5120768 | 0.00044462 |
| 14950 | H13 | 4561.26598 | -0.422531 | 0.12036346 | -3.5104596 | 0.00044733 |
| 93893 | Pcdhb22 | 79.9721982 | 1.64159831 | 0.46768015 | 3.51008764 | 0.00044796 |
| 51798 | ECH1 | 643.293796 | -0.6471827 | 0.18444156 | -3.5088769 | 0.00045 |
| 245450 | SLITRK2 | 7.88469971 | -6.406686 | 1.82617554 | -3.5082531 | 0.00045106 |
| 67972 | ATP2B1 | 8234.56468 | -0.2431881 | 0.06934426 | -3.5069676 | 0.00045324 |
| 20116 | RPS8 | 1364.736 | 0.56343137 | 0.1606826 | 3.50648656 | 0.00045406 |
| 20541 | SLC8A1 | 13.5384638 | 3.12268192 | 0.89062495 | 3.50616938 | 0.00045461 |
| 71846 | SYCE2 | 46.8079195 | -1.93154 | 0.5511288 | -3.504698 | 0.00045713 |
| 269003 | SAP130 | 2104.0684 | 0.17882223 | 0.05102363 | 3.50469423 | 0.00045713 |
| 26362 | AXL | 34644.5555 | 0.2204117 | 0.06292555 | 3.50273763 | 0.0004605 |
| 20738 | Spn-ps | 8.62166496 | 2.1540582 | 0.61509959 | 3.50196656 | 0.00046184 |
| 67655 | CTDP1 | 1707.22319 | 0.31875639 | 0.09102574 | 3.50182714 | 0.00046208 |
| 71176 | FBXO24 | 10.6064791 | -2.2113062 | 0.6314829 | -3.5017674 | 0.00046218 |
| 12338 | CAPN6 | 8.19380036 | 2.47977737 | 0.70819396 | 3.50155115 | 0.00046256 |
| 104111 | ADCY3 | 317.233367 | 1.54198026 | 0.44047298 | 3.50073749 | 0.00046397 |
| 71242 | SPATA24 | 122.226419 | -0.8586116 | 0.24528915 | -3.500406 | 0.00046455 |
| 66950 | TMEM206 | 476.782515 | -0.5401853 | 0.1543472 | -3.4998062 | 0.0004656 |
| 75420 | SECISBP2 | 506.780039 | 0.38431188 | 0.10983819 | 3.49889109 | 0.0004672 |
| 24136 | ZEB2 | 2164.13148 | 0.54494668 | 0.15575671 | 3.49870442 | 0.00046752 |
| 67839 | GPSM1 | 501.513166 | 0.81136641 | 0.23191159 | 3.49860222 | 0.0004677 |
| 57908 | Zfp318 | 974.952596 | 0.50553255 | 0.14451651 | 3.49809552 | 0.00046859 |
| 328133 | SLC39A9 | 2645.23401 | 0.28189611 | 0.08059511 | 3.49768243 | 0.00046932 |
| 68564 | NUFIP2 | 4715.33553 | -0.172581 | 0.04935218 | -3.4969269 | 0.00047065 |
| 65254 | DPYSL5 | 6.33393712 | -3.5413669 | 1.0127185 | -3.4968917 | 0.00047071 |
| 230027 | COQ3 | 527.44407 | -0.9972034 | 0.28519659 | -3.4965475 | 0.00047132 |
| 22275 | UROD | 743.802891 | 0.48866205 | 0.13984865 | 3.49422069 | 0.00047545 |
| 67379 | DEDD2 | 217.554705 | -0.7132524 | 0.20414552 | -3.4938433 | 0.00047612 |
| 100043882 | Gm14277 | 226.19517 | 0.35297603 | 0.10105013 | 3.49307839 | 0.00047749 |
| 22626 | SLC23A3 | 15.2554603 | -1.430669 | 0.40966889 | -3.492257 | 0.00047896 |
| 66158 | Cxx1a | 38.8553199 | -1.1037758 | 0.31606821 | -3.4922075 | 0.00047905 |
| 11426 | MACF1 | 8448.8175 | 0.61133488 | 0.17506231 | 3.49209872 | 0.00047924 |
| 100504265 | Gm20139 | 3.93112926 | -5.399518 | 1.54637157 | -3.4917339 | 0.0004799 |
| 21854 | TIMM17A | 1579.09073 | -0.3188706 | 0.09132294 | -3.4916811 | 0.00047999 |
| 67283 | SLC25A19 | 376.001943 | -0.581891 | 0.1666527 | -3.4916387 | 0.00048007 |
| 68981 | SNRPA1 | 1668.11148 | 0.61326024 | 0.17564055 | 3.49156406 | 0.0004802 |
| 12828 | COL4A3 | 7.15624992 | -3.6917306 | 1.05741999 | -3.4912624 | 0.00048074 |

| | | | | | | |
|---|---|---|---|---|---|---|
| 208228 | MOB3A | 1194.46051 | -0.2916004 | 0.08352982 | -3.4909738 | 0.00048126 |
| 213484 | NUDT18 | 221.334795 | -1.1558871 | 0.33113625 | -3.4906692 | 0.00048181 |
| 381924 | ITGAD | 3.14879219 | -5.0772609 | 1.45465409 | -3.4903562 | 0.00048238 |
| 67101 | 2310039H08 | 147.278394 | -0.6598259 | 0.18904356 | -3.4903382 | 0.00048241 |
| 67264 | NDUFB8 | 2035.32101 | -0.2516955 | 0.07212125 | -3.4898933 | 0.00048321 |
| 233899 | CCDC189 | 24.6533132 | -1.3176802 | 0.37765495 | -3.4891113 | 0.00048463 |
| 117160 | TTYH2 | 882.627813 | 0.57793808 | 0.16564886 | 3.48893477 | 0.00048495 |
| 15168 | HCN3 | 8.47027107 | -3.3497216 | 0.96041302 | -3.4877928 | 0.00048703 |
| 72373 | PSCA | 3.57956642 | -5.2773018 | 1.51337618 | -3.4871051 | 0.00048828 |
| 67310 | Prl8a9 | 4.79406932 | -3.5591192 | 1.02078408 | -3.4866523 | 0.00048911 |
| 235339 | DLAT | 2577.00754 | 0.79905118 | 0.22921065 | 3.48609972 | 0.00049012 |
| 80720 | PBX4 | 18.3017038 | -2.8030584 | 0.80435631 | -3.4848466 | 0.00049242 |
| 433759 | HDAC1 | 256.233122 | 0.75474376 | 0.21660781 | 3.48437927 | 0.00049328 |
| 52118 | PVR | 1927.65051 | 0.18232284 | 0.05232673 | 3.48431552 | 0.0004934 |
| 109135 | PLEKHA5 | 449.745739 | 0.84448098 | 0.24238329 | 3.48407255 | 0.00049385 |
| 52715 | CCDC43 | 1272.07494 | -0.2378805 | 0.06828199 | -3.4837962 | 0.00049436 |
| 110351 | RAP1GAP | 457.666912 | -1.4980507 | 0.43002031 | -3.4836744 | 0.00049458 |
| 75778 | THEM4 | 109.461201 | -0.7080188 | 0.2032405 | -3.4836502 | 0.00049463 |
| 13804 | endog | 138.600725 | 0.65822201 | 0.18895441 | 3.48349638 | 0.00049491 |
| 52004 | CDK2AP2 | 584.543777 | -0.3470504 | 0.09964046 | -3.4830274 | 0.00049578 |
| 620986 | Gm6195 | 50.1988607 | 0.86724939 | 0.24902662 | 3.48255696 | 0.00049665 |
| 75577 | DYNAP | 117.189227 | -1.7043992 | 0.48949213 | -3.4819747 | 0.00049773 |
| 80291 | rilpl2 | 292.995288 | -0.765775 | 0.21992633 | -3.4819613 | 0.00049776 |
| 29870 | GTSE1 | 2803.88117 | -0.3854121 | 0.11070783 | -3.4813446 | 0.0004989 |
| 26373 | CLCN7 | 1869.42508 | -0.3748337 | 0.10767262 | -3.4812349 | 0.00049911 |
| 56429 | DPT | 17.6157119 | 6.88854375 | 1.97900171 | 3.48081748 | 0.00049989 |
| 54644 | OTUD5 | 1972.33109 | -0.2681756 | 0.07712409 | -3.4771963 | 0.00050669 |
| 12807 | HPS3 | 672.136337 | -0.5370266 | 0.15444565 | -3.4771235 | 0.00050682 |
| 171531 | MLPH | 984.224951 | -1.653975 | 0.47584817 | -3.4758461 | 0.00050924 |
| 231874 | CCZ1 | 2013.46504 | -0.2438053 | 0.07016598 | -3.4746933 | 0.00051144 |
| 74337 | PALM3 | 16.6036547 | 1.81788214 | 0.52318876 | 3.47462002 | 0.00051158 |
| 14376 | GANAB | 8568.52299 | -0.2796725 | 0.08050371 | -3.4740328 | 0.0005127 |
| 242109 | Zfp697 | 834.633846 | -0.6936229 | 0.19971063 | -3.4731396 | 0.00051441 |
| 66493 | MRPL51 | 667.254377 | 0.3673968 | 0.10578728 | 3.47297701 | 0.00051472 |
| 110213 | TMBIM6 | 13098.373 | -0.2723953 | 0.07843976 | -3.4726692 | 0.00051531 |
| 13424 | DYNC1H1 | 25702.5741 | 0.57502232 | 0.16559298 | 3.47250415 | 0.00051563 |
| 50782 | RGS11 | 18.0090849 | -1.6043032 | 0.46214551 | -3.4714244 | 0.00051771 |
| 227526 | CDNF | 41.2479216 | -1.3770285 | 0.39676604 | -3.4706311 | 0.00051924 |
| 229571 | Gm4858 | 3.70208773 | -5.3111488 | 1.53112843 | -3.4687807 | 0.00052283 |
| 72479 | HSDL2 | 468.252836 | -0.63659 | 0.18354446 | -3.4683148 | 0.00052373 |
| 53895 | CLPP | 926.862119 | -0.3131001 | 0.09028537 | -3.4678936 | 0.00052455 |
| 223970 | RMI2 | 145.540175 | 0.75168508 | 0.21679814 | 3.46721185 | 0.00052589 |
| 74341 | G630025P09 | 20.3854785 | -1.9093784 | 0.55071134 | -3.4671129 | 0.00052608 |
| 110172 | SLC35B1 | 1590.23443 | -0.2256136 | 0.06507255 | -3.4671084 | 0.00052609 |
| 100503117 | 2010010A06 | 3.11301997 | -5.0710409 | 1.46312435 | -3.4658988 | 0.00052846 |

| | | | | | | |
|---|---|---|---|---|---|---|
| 212679 | MARS2 | 612.969902 | -0.3899423 | 0.11254026 | -3.4649134 | 0.0005304 |
| 226025 | TRPM3 | 12.7358973 | -1.528969 | 0.4412914 | -3.4647604 | 0.0005307 |
| 240047 | MMP25 | 3.71370624 | -4.5599053 | 1.31618978 | -3.464474 | 0.00053127 |
| 16848 | LFNG | 413.728592 | -0.8943476 | 0.25819623 | -3.4638292 | 0.00053254 |
| 280408 | rilp | 37.5394665 | -0.8744631 | 0.25247927 | -3.4635044 | 0.00053319 |
| 71919 | RPAP3 | 796.117027 | 0.22713099 | 0.06559419 | 3.46266932 | 0.00053485 |
| 66205 | CD302 | 187.572309 | 2.64060052 | 0.76294664 | 3.4610553 | 0.00053806 |
| 69125 | CNOT8 | 991.966081 | 0.18054523 | 0.05217302 | 3.46050969 | 0.00053915 |
| 72181 | NSUN4 | 357.192269 | 0.61102566 | 0.17659705 | 3.45999925 | 0.00054018 |
| 22670 | TRIM26 | 1576.51485 | -0.3724966 | 0.10765953 | -3.45995 | 0.00054028 |
| 71949 | CERS5 | 2494.76191 | 0.20286257 | 0.05863862 | 3.45953861 | 0.0005411 |
| 56258 | HNRNPH2 | 1861.88256 | 0.28172431 | 0.08144677 | 3.45899933 | 0.00054219 |
| 382137 | FDXACB1 | 224.534726 | 0.46931991 | 0.13573153 | 3.45770744 | 0.00054479 |
| 234695 | CARMIL2 | 38.8998911 | 1.43945767 | 0.41644523 | 3.45653538 | 0.00054717 |
| 107035 | FBXO38 | 2116.96291 | -0.2762489 | 0.07992603 | -3.4563067 | 0.00054763 |
| 74326 | HNRNPR | 2172.66912 | 0.69398468 | 0.20091992 | 3.45403623 | 0.00055226 |
| 78890 | TRMT44 | 406.301373 | -0.3344367 | 0.09684662 | -3.4532615 | 0.00055385 |
| 100503178 | 2810013P06I | 85.2602116 | -0.6077767 | 0.17601563 | -3.4529699 | 0.00055445 |
| 19231 | PTMA | 2326.88572 | 0.3841989 | 0.11126737 | 3.45293405 | 0.00055452 |
| 19038 | PPIC | 2009.92169 | 1.21177739 | 0.35099141 | 3.45244178 | 0.00055554 |
| 14283 | fosl1 | 6985.74836 | 1.04115723 | 0.30159181 | 3.45220659 | 0.00055602 |
| 224805 | AARS2 | 512.652516 | 0.31261976 | 0.09056627 | 3.45183453 | 0.00055679 |
| 13637 | EFNA2 | 28.3554619 | 1.68849936 | 0.48925084 | 3.45119358 | 0.00055811 |
| 67976 | TRABD | 1632.89309 | -0.2121213 | 0.06146696 | -3.4509808 | 0.00055855 |
| 320802 | IFITM10 | 18.9229414 | -2.5305835 | 0.73335059 | -3.4507145 | 0.0005591 |
| 19035 | PPIB | 3841.44163 | 0.44703505 | 0.12964869 | 3.44804911 | 0.00056465 |
| 103220 | ttc41 | 45.2074982 | -0.7861662 | 0.22800672 | -3.4479959 | 0.00056476 |
| 18987 | POU2F2 | 5.70606834 | -3.4103982 | 0.98913739 | -3.4478509 | 0.00056507 |
| 68119 | CMTM3 | 2160.28979 | 0.93844382 | 0.27222758 | 3.44727685 | 0.00056627 |
| 19989 | RPL7 | 3650.61696 | -0.2408583 | 0.06988857 | -3.4463184 | 0.00056828 |
| 665119 | SEC14L5 | 7.6682671 | -4.8601351 | 1.41081884 | -3.4449037 | 0.00057126 |
| 65970 | LIMA1 | 4877.44512 | 0.55794251 | 0.16196242 | 3.4448887 | 0.00057129 |
| 101700 | TRIM68 | 196.761526 | -0.8958315 | 0.26004673 | -3.4448866 | 0.0005713 |
| 104002 | Ctsq | 3.08337452 | -5.0582758 | 1.4689986 | -3.4433496 | 0.00057456 |
| 66679 | RAE1 | 1402.45216 | 0.30719012 | 0.08921743 | 3.44316247 | 0.00057495 |
| 330401 | TMCC1 | 713.928284 | -0.3797389 | 0.11031734 | -3.442241 | 0.00057692 |
| 170767 | rfxap | 442.587226 | -0.390956 | 0.1136118 | -3.4411565 | 0.00057923 |
| 52530 | nhp2 | 3410.66308 | 0.15246983 | 0.04430808 | 3.44112896 | 0.00057929 |
| 100041797 | #N/A | 205.356812 | -0.4447827 | 0.12927464 | -3.4406029 | 0.00058042 |
| 110895 | SLC9A4 | 13.6478246 | 2.53960876 | 0.73822132 | 3.44017259 | 0.00058134 |
| 66460 | SYS1 | 927.137106 | -0.3585947 | 0.10424756 | -3.4398378 | 0.00058206 |
| 69524 | ESAM | 19.8632286 | 1.36078696 | 0.39576462 | 3.43837444 | 0.00058522 |
| 230971 | MEGF6 | 141.31208 | -1.9702534 | 0.57314491 | -3.4376183 | 0.00058685 |
| 381535 | Gm12800 | 5.14763278 | -5.7895366 | 1.68451365 | -3.4369188 | 0.00058837 |
| 94280 | SFXN3 | 2127.52556 | -0.5148858 | 0.14981445 | -3.4368232 | 0.00058858 |

| | | | | | | |
|---|---|---|---|---|---|---|
| 74901 | KBTBD11 | 12.9144503 | 3.49631242 | 1.0173734 | 3.43660687 | 0.00058905 |
| 67378 | BBS2 | 153.340669 | -0.4771667 | 0.13892043 | -3.4348205 | 0.00059295 |
| 74195 | ELP3 | 1744.90888 | 0.68811011 | 0.20037944 | 3.43403542 | 0.00059467 |
| 268445 | ANKRD13B | 2073.1938 | 0.53745207 | 0.15652193 | 3.43371746 | 0.00059536 |
| 84035 | KREMEN1 | 944.745158 | 0.41018281 | 0.11947946 | 3.43308233 | 0.00059676 |
| 54645 | GRIPAP1 | 902.900785 | 0.23278764 | 0.06782428 | 3.43221717 | 0.00059867 |
| 16912 | PSMB9 | 35.384804 | -2.3908137 | 0.69663101 | -3.4319657 | 0.00059922 |
| 11514 | ADCY8 | 36.7950309 | -1.7423412 | 0.507732 | -3.4316159 | 0.0006 |
| 226499 | BC003331 | 996.919815 | -0.3662061 | 0.10672281 | -3.4313757 | 0.00060053 |
| 12751 | TPP1 | 4176.13257 | -0.6895516 | 0.20100692 | -3.4304868 | 0.0006025 |
| 73516 | 1700086O06 | 193.751148 | -0.5740927 | 0.16736577 | -3.4301677 | 0.00060321 |
| 94178 | MCOLN1 | 1255.59483 | -0.67037 | 0.19545622 | -3.4297707 | 0.00060409 |
| 73822 | MFSD12 | 178.702326 | -0.8448471 | 0.24637741 | -3.4290771 | 0.00060564 |
| 66096 | LAMTOR4 | 467.904676 | -0.3247298 | 0.09470406 | -3.4288901 | 0.00060605 |
| 11881 | ARSB | 2233.71582 | -0.2146773 | 0.06261118 | -3.4287378 | 0.00060639 |
| 50754 | FBXW7 | 616.444718 | -0.3399159 | 0.099143 | -3.428542 | 0.00060683 |
| 208768 | SDE2 | 1737.71975 | 0.32169806 | 0.09383852 | 3.42820919 | 0.00060758 |
| 71941 | CARS2 | 409.408976 | -0.3178646 | 0.0927248 | -3.4280427 | 0.00060795 |
| 56376 | PDLIM5 | 11654.0694 | -0.6467951 | 0.18869319 | -3.4277605 | 0.00060858 |
| 67430 | 4921536K21I | 50.5880869 | -0.7470102 | 0.21800515 | -3.4265715 | 0.00061125 |
| 22719 | Zfp61 | 151.830513 | 1.06812866 | 0.31173061 | 3.42644777 | 0.00061153 |
| 320685 | DCTD | 576.640216 | 0.70608034 | 0.20609335 | 3.42602194 | 0.00061249 |
| 56444 | ACTR10 | 2613.01772 | 0.24342174 | 0.07106849 | 3.42517089 | 0.00061441 |
| 15568 | ELAVL1 | 4591.89367 | 0.36820875 | 0.10750709 | 3.42497191 | 0.00061486 |
| 230576 | TTC22 | 3.70673645 | -5.3262008 | 1.55527512 | -3.4246036 | 0.0006157 |
| 73212 | 3110082I17R | 694.159434 | -0.4389819 | 0.12821622 | -3.4237623 | 0.00061761 |
| 67726 | FAM114A2 | 1308.48584 | 0.20516284 | 0.05992757 | 3.42351332 | 0.00061817 |
| 13001 | CSNK2B | 2770.14713 | -0.2320286 | 0.06777636 | -3.4234447 | 0.00061833 |
| 11848 | RHOA | 3533.99892 | 0.12913373 | 0.03772338 | 3.423175 | 0.00061894 |
| 16656 | HIVEP3 | 71.1708988 | -1.782128 | 0.52068518 | -3.4226593 | 0.00062012 |
| 235416 | LMAN1L | 21.5828473 | 1.72791828 | 0.50485715 | 3.42258849 | 0.00062028 |
| 235406 | snx33 | 1140.22818 | 0.51011448 | 0.14910354 | 3.4212097 | 0.00062343 |
| 56418 | YKT6 | 5162.84656 | -0.2724197 | 0.0796357 | -3.4208231 | 0.00062432 |
| 16978 | LRRFIP1 | 6286.38541 | -0.1107537 | 0.03238009 | -3.420427 | 0.00062523 |
| 22247 | UMPS | 3281.51888 | 0.46228134 | 0.13515944 | 3.4202667 | 0.0006256 |
| 668940 | MYH7B | 6.97812881 | -4.783974 | 1.39921537 | -3.4190405 | 0.00062842 |
| 20192 | RYR3 | 5.12117942 | -5.0227226 | 1.47003575 | -3.416735 | 0.00063377 |
| 54633 | PQBP1 | 977.762151 | 0.24867688 | 0.07280513 | 3.41565072 | 0.0006363 |
| 319266 | A130010J15F | 360.029204 | 0.4693155 | 0.13742464 | 3.41507534 | 0.00063764 |
| 26936 | MPRIP | 13742.1738 | 0.17339848 | 0.05078901 | 3.4140947 | 0.00063994 |
| 230612 | SLC5A9 | 7.89211107 | -6.4082121 | 1.87710972 | -3.4138719 | 0.00064047 |
| 56503 | ANKRD49 | 570.480896 | 0.31161299 | 0.09130876 | 3.41273925 | 0.00064313 |
| 230863 | SH2D5 | 1214.15308 | -1.4742106 | 0.43199839 | -3.4125374 | 0.00064361 |
| 53602 | HPCAL1 | 749.535542 | -0.6456798 | 0.18922321 | -3.4122654 | 0.00064425 |
| 21355 | tap2 | 718.613267 | -0.757474 | 0.22199786 | -3.412078 | 0.0006447 |

| | | | | | | |
|---|---|---|---|---|---|---|
| 66948 | ACAD8 | 467.189468 | 0.27576321 | 0.0808367 | 3.41136169 | 0.00064639 |
| 69617 | PITRM1 | 4290.72418 | 0.42359039 | 0.12419624 | 3.41065397 | 0.00064807 |
| 76781 | METTL4 | 392.511379 | 0.42447945 | 0.12457095 | 3.40753152 | 0.00065553 |
| 110033 | KIF22 | 2100.36067 | 0.70639143 | 0.20731361 | 3.4073568 | 0.00065595 |
| 15574 | HUS1 | 1087.47014 | 0.26252364 | 0.07708153 | 3.40579161 | 0.00065973 |
| 20709 | Serpinb9f | 3.44324263 | -5.2220404 | 1.53331514 | -3.4057189 | 0.0006599 |
| 67586 | UBXN11 | 11.6911572 | -2.0604083 | 0.60504553 | -3.4053773 | 0.00066073 |
| 69104 | 41702 | 2035.23481 | -0.2253187 | 0.06617382 | -3.4049525 | 0.00066176 |
| 68653 | SAMM50 | 3270.30302 | -0.1703187 | 0.05002469 | -3.4046934 | 0.00066238 |
| 67942 | ATP5G2 | 12.2537788 | -1.7911037 | 0.52608318 | -3.4046017 | 0.00066261 |
| 328967 | ARHGEF37 | 8.12002948 | -2.6365262 | 0.7744705 | -3.4042952 | 0.00066335 |
| 76824 | MTFR1L | 1235.46204 | 0.46079758 | 0.1353791 | 3.40375717 | 0.00066466 |
| 235574 | ATP2C1 | 2354.08357 | 0.22833419 | 0.06709371 | 3.4032129 | 0.00066598 |
| 14466 | GBA | 2417.12701 | -0.9072214 | 0.26668373 | -3.4018627 | 0.00066928 |
| 74178 | STK40 | 600.599136 | 1.03784301 | 0.30514886 | 3.40110404 | 0.00067114 |
| 18846 | PLXNA3 | 436.090539 | -0.6454431 | 0.18978038 | -3.401 | 0.0006714 |
| 100503467 | AA667203 | 6.40695692 | 2.69813627 | 0.79336505 | 3.40087614 | 0.0006717 |
| 13684 | EIF4E | 2459.1553 | 0.7312606 | 0.21503713 | 3.40062486 | 0.00067232 |
| 378431 | TXLNB | 8.9478091 | -2.3145144 | 0.68069317 | -3.4002316 | 0.00067329 |
| 58222 | RAB37 | 6.32882199 | -3.9654776 | 1.16631441 | -3.4000074 | 0.00067384 |
| 26903 | DYSF | 258.506053 | 1.34623225 | 0.39598205 | 3.39973053 | 0.00067452 |
| 235567 | DNAJC13 | 5783.22792 | 0.34059714 | 0.10018455 | 3.39969729 | 0.0006746 |
| 20446 | st6galnac2 | 14.957249 | -3.4652516 | 1.01934013 | -3.3995047 | 0.00067508 |
| 622408 | MCIDAS | 7.00793943 | 2.67822415 | 0.78783543 | 3.39947159 | 0.00067516 |
| 68239 | KRT42 | 2.99407654 | -5.0069076 | 1.47293502 | -3.3992726 | 0.00067565 |
| 16136 | IGLL1 | 2.99407654 | -5.0069076 | 1.47293502 | -3.3992726 | 0.00067565 |
| 23894 | GTF2H2 | 472.322886 | 0.50770218 | 0.1493751 | 3.39884072 | 0.00067672 |
| 245867 | PCMTD2 | 533.586433 | -0.5617585 | 0.16530313 | -3.3983536 | 0.00067793 |
| 68480 | CARD19 | 644.496321 | -0.7172564 | 0.21112619 | -3.3972877 | 0.00068057 |
| 791294 | Gm9958 | 17.0484008 | -1.3154839 | 0.3872439 | -3.3970424 | 0.00068118 |
| 27376 | SLC25A10 | 1932.10853 | 0.44406379 | 0.1307316 | 3.39675948 | 0.00068189 |
| 68802 | MYPN | 4.88677469 | -4.9486842 | 1.45705401 | -3.3963629 | 0.00068288 |
| 68721 | 1110032A03 | 78.2031827 | -0.7571769 | 0.22294536 | -3.3962443 | 0.00068317 |
| 69878 | SNRPF | 418.591599 | 0.56375516 | 0.16600363 | 3.39604115 | 0.00068368 |
| 68512 | TOMM5 | 417.508617 | 0.59891977 | 0.17642959 | 3.39466737 | 0.00068712 |
| 14998 | H2-DMa | 28.5984507 | -1.0795958 | 0.31808057 | -3.3940955 | 0.00068856 |
| 11307 | ABCG1 | 123.701512 | -2.8149022 | 0.82944328 | -3.3937248 | 0.00068949 |
| 20324 | SDPR | 5759.09699 | -1.0036803 | 0.29575016 | -3.3936764 | 0.00068961 |
| 12633 | CFLAR | 3673.58007 | -0.3743041 | 0.11029825 | -3.3935638 | 0.00068989 |
| 242409 | TMEM8B | 145.462456 | -0.7403845 | 0.21818223 | -3.3934224 | 0.00069025 |
| 104103 | AIRN | 40.2921264 | -1.0997824 | 0.3241008 | -3.3933345 | 0.00069047 |
| 17869 | MYC | 3878.59633 | 0.65291838 | 0.19241365 | 3.39330598 | 0.00069054 |
| 235493 | FAM214A | 237.331847 | 0.4228022 | 0.12463536 | 3.39231333 | 0.00069305 |
| 27356 | INSL6 | 33.7613589 | -1.0887387 | 0.32096711 | -3.392057 | 0.0006937 |
| 16492 | KCNA4 | 17.921798 | -2.5894984 | 0.76346815 | -3.391757 | 0.00069446 |

| | | | | | | |
|---|---|---|---|---|---|---|
| 76589 | UNC5CL | 5.31257659 | -5.8366534 | 1.72106912 | -3.3912952 | 0.00069563 |
| 20174 | RUVBL2 | 2581.39452 | 0.6147676 | 0.18129005 | 3.39107188 | 0.0006962 |
| 12984 | Csf2rb2 | 7.13039544 | -5.5244794 | 1.62940522 | -3.3904883 | 0.00069768 |
| 231713 | NAA25 | 2410.12107 | 0.29755186 | 0.08779727 | 3.38907883 | 0.00070128 |
| 75291 | zbtb3 | 80.1457836 | 0.71430451 | 0.21076903 | 3.38903918 | 0.00070138 |
| 240068 | Zfp563 | 230.613671 | -0.3837221 | 0.11323048 | -3.3888586 | 0.00070184 |
| 69748 | ALDH16A1 | 1117.50804 | 0.56500157 | 0.16674095 | 3.38849921 | 0.00070276 |
| 630836 | 2010315B03 | 157.915719 | -0.5057441 | 0.14926674 | -3.3881905 | 0.00070355 |
| 72843 | prdm4 | 2086.56297 | -0.2282363 | 0.06736664 | -3.3879723 | 0.00070411 |
| 228071 | SESTD1 | 2785.31003 | 0.44724919 | 0.13203897 | 3.38725129 | 0.00070597 |
| 93880 | PCDHB9 | 19.3829532 | 2.78748041 | 0.82306142 | 3.38672223 | 0.00070733 |
| 246728 | OAS2 | 19.2160576 | -3.3430469 | 0.98713206 | -3.3866258 | 0.00070758 |
| 94092 | TRIM16 | 1833.78983 | -0.4919169 | 0.14529549 | -3.3856307 | 0.00071015 |
| 241311 | zbtb34 | 261.471698 | -0.51775 | 0.1529482 | -3.3851332 | 0.00071144 |
| 68576 | LAMTOR5 | 480.528827 | -0.2611126 | 0.07713514 | -3.385131 | 0.00071144 |
| 16475 | AJUBA | 3034.15665 | -0.9598864 | 0.28358099 | -3.3848756 | 0.00071211 |
| 212986 | SCFD2 | 366.321729 | 0.30586247 | 0.09038954 | 3.383826 | 0.00071483 |
| 244962 | SNX14 | 907.156117 | 0.2907397 | 0.08592954 | 3.38346623 | 0.00071577 |
| 19724 | RFX1 | 795.416783 | 0.31946973 | 0.09442841 | 3.38319521 | 0.00071648 |
| 12569 | CDK5R1 | 62.7625801 | 0.89125408 | 0.2634429 | 3.38310148 | 0.00071672 |
| 238871 | PDE4D | 297.83463 | 0.59970056 | 0.17737294 | 3.38101493 | 0.00072219 |
| 81006 | GPR63 | 85.9582917 | -0.8951087 | 0.26475175 | -3.3809357 | 0.00072239 |
| 83996 | Mmp1b | 27.2354711 | 1.72710996 | 0.51098035 | 3.37999289 | 0.00072488 |
| 80905 | POLH | 989.373836 | 0.43055208 | 0.12740271 | 3.37945782 | 0.00072629 |
| 53356 | EIF3G | 2567.30389 | 0.26105495 | 0.07725956 | 3.37893394 | 0.00072767 |
| 83962 | BTBD1 | 1123.53818 | 0.85965898 | 0.25449609 | 3.37788671 | 0.00073045 |
| 266692 | CPNE1 | 3865.31691 | 0.58540706 | 0.17331729 | 3.37766116 | 0.00073105 |
| 210741 | KCNK12 | 19.897956 | -6.3434802 | 1.87879135 | -3.3763623 | 0.00073451 |
| 243083 | TMPRSS11F | 19.7517952 | -1.2670352 | 0.37531096 | -3.3759611 | 0.00073558 |
| 28018 | UBFD1 | 1682.81365 | 0.5693188 | 0.16874715 | 3.37379794 | 0.00074139 |
| 105833 | CCDC65 | 20.8447882 | -2.0270738 | 0.60093986 | -3.3731725 | 0.00074307 |
| 104394 | E2F4 | 2268.35284 | 0.25721466 | 0.07626919 | 3.37245822 | 0.000745 |
| 72053 | TMUB2 | 708.985882 | -0.2211039 | 0.06557581 | -3.37173 | 0.00074698 |
| 70599 | SSFA2 | 2484.55618 | 0.44682086 | 0.13255496 | 3.37083475 | 0.00074941 |
| 216345 | ZFC3H1 | 2528.5785 | -0.5572359 | 0.16532045 | -3.3706412 | 0.00074993 |
| 66407 | MRPS15 | 817.825719 | 0.53686529 | 0.15929186 | 3.37032464 | 0.0007508 |
| 17356 | AFDN | 7059.50107 | -0.6481076 | 0.19231342 | -3.3700589 | 0.00075152 |
| 328035 | FADS6 | 18.9147586 | -5.5307871 | 1.64120873 | -3.3699474 | 0.00075183 |
| 208795 | TMEM63A | 2043.63271 | -0.781383 | 0.23188094 | -3.3697598 | 0.00075234 |
| 67008 | YAE1D1 | 448.429054 | 0.51875598 | 0.15395766 | 3.36947171 | 0.00075312 |
| 230837 | ASAP3 | 116.325532 | -1.7256936 | 0.51222254 | -3.369031 | 0.00075433 |
| 17863 | MYB | 28.4614449 | -2.9380918 | 0.87212393 | -3.3688925 | 0.00075471 |
| 216516 | CCDC157 | 393.663739 | -0.4616975 | 0.13706516 | -3.3684525 | 0.00075591 |
| 14884 | GTF2H1 | 3178.81053 | 0.32060043 | 0.09521493 | 3.36712343 | 0.00075957 |
| 100503800 | #N/A | 4.75057458 | -4.1575155 | 1.23485918 | -3.3667932 | 0.00076048 |

| | | | | | | |
|---|---|---|---|---|---|---|
| 13822 | EPB41L2 | 6159.88908 | -0.4988685 | 0.14818394 | -3.3665489 | 0.00076115 |
| 71544 | ARHGAP42 | 557.57236 | 0.59713985 | 0.17737622 | 3.36651581 | 0.00076124 |
| 27390 | MMEL1 | 6.53027135 | -2.6914614 | 0.79968879 | -3.365636 | 0.00076367 |
| 16777 | LAMB1 | 16849.1829 | 0.44159305 | 0.13127978 | 3.36375526 | 0.0007689 |
| 239759 | LIPH | 6.72486955 | -2.9526478 | 0.87790085 | -3.3633044 | 0.00077015 |
| 20018 | POLR1D | 1618.73509 | 0.30284048 | 0.09007167 | 3.36221673 | 0.00077319 |
| 106821 | OARD1 | 533.917425 | 0.6076034 | 0.18073032 | 3.36193395 | 0.00077399 |
| 245684 | CNKSR2 | 9.18105083 | -6.6270586 | 1.97146811 | -3.361484 | 0.00077525 |
| 68205 | URM1 | 1091.02699 | 0.55775381 | 0.16593979 | 3.36118193 | 0.0007761 |
| 108811 | CCDC122 | 173.164622 | -0.8573141 | 0.2552081 | -3.3592748 | 0.00078147 |
| 59041 | STK25 | 1637.8306 | -0.2543213 | 0.07570936 | -3.3591784 | 0.00078175 |
| 101437 | DHX32 | 1950.41429 | -0.4322665 | 0.12868806 | -3.3590258 | 0.00078218 |
| 78878 | FTX | 278.816925 | -0.5903587 | 0.17576215 | -3.3588502 | 0.00078267 |
| 72832 | CRTAC1 | 20.1806003 | -7.7653908 | 2.31295889 | -3.3573406 | 0.00078696 |
| 219134 | SHISA2 | 9.67413893 | -4.2093617 | 1.25400051 | -3.3567464 | 0.00078865 |
| 67698 | FAM174A | 369.903271 | -0.6740514 | 0.20082935 | -3.3563393 | 0.00078982 |
| 225861 | SNX32 | 33.7358487 | -1.1468724 | 0.34172375 | -3.3561391 | 0.00079039 |
| 73739 | CBY1 | 335.724269 | -0.4710874 | 0.14038321 | -3.3557244 | 0.00079157 |
| 20972 | SYNGR1 | 259.941196 | -2.0656219 | 0.61569176 | -3.354961 | 0.00079376 |
| 50784 | PLPP2 | 4012.64479 | -0.8013301 | 0.23893598 | -3.3537441 | 0.00079726 |
| 20683 | SP1 | 4105.00183 | -0.2416361 | 0.07205737 | -3.3533847 | 0.0007983 |
| 100038730 | Gm10387 | 26.9250754 | 1.92764686 | 0.57487678 | 3.35314788 | 0.00079898 |
| 72351 | PTAR1 | 328.885741 | 0.52820578 | 0.1575305 | 3.35303827 | 0.0007993 |
| 66615 | ATG4B | 1247.95333 | -0.2941295 | 0.08772348 | -3.3529167 | 0.00079965 |
| 329421 | MYO3B | 3.19074862 | -4.3393114 | 1.29438322 | -3.3524163 | 0.00080109 |
| 68152 | FAM133B | 642.480227 | 0.25392898 | 0.07575941 | 3.35178122 | 0.00080293 |
| 66154 | TMEM14C | 1486.57252 | 0.29075267 | 0.08674974 | 3.35162587 | 0.00080339 |
| 108899 | 2700081O15 | 1413.3079 | 0.91018323 | 0.27156618 | 3.3516075 | 0.00080344 |
| 21357 | TARBP2 | 224.558316 | -0.5663878 | 0.16900631 | -3.3512818 | 0.00080438 |
| 108105 | B3GNT5 | 15.8368187 | -7.4146073 | 2.21329334 | -3.3500337 | 0.00080802 |
| 66815 | ccdc109b | 312.198066 | 1.65008888 | 0.49257296 | 3.34993801 | 0.0008083 |
| 67529 | FGFR1OP2 | 1855.21371 | 0.29506753 | 0.08809632 | 3.34937409 | 0.00080994 |
| 56693 | CRTAP | 2095.14668 | 0.55506359 | 0.16572907 | 3.34922291 | 0.00081039 |
| 319475 | Zfp672 | 362.129745 | -0.4242526 | 0.12667329 | -3.3491876 | 0.00081049 |
| 69228 | Zfp746 | 1127.46907 | 0.27212756 | 0.08128068 | 3.34799824 | 0.00081398 |
| 81877 | TNXB | 251.281855 | 2.28368424 | 0.68231058 | 3.34698643 | 0.00081695 |
| 67219 | MED18 | 407.679087 | 0.60553933 | 0.18098191 | 3.34585552 | 0.00082029 |
| 78108 | PARTICL | 46.3797495 | 0.93817799 | 0.28045779 | 3.3451664 | 0.00082233 |
| 68055 | ATP5S | 87.1604568 | -0.5892096 | 0.1761668 | -3.3446119 | 0.00082398 |
| 237898 | USP32 | 2726.37646 | -0.4263242 | 0.12746701 | -3.3445849 | 0.00082406 |
| 69837 | PCGF1 | 86.2028711 | 0.65057523 | 0.19453461 | 3.34426475 | 0.00082501 |
| 67278 | Pagr1a | 730.926913 | 0.68550846 | 0.20502339 | 3.34356221 | 0.0008271 |
| 13607 | EDA | 6.09209686 | -6.0336384 | 1.80458012 | -3.3435137 | 0.00082725 |
| 101809 | SPRED3 | 410.689263 | -0.8942278 | 0.26752243 | -3.3426272 | 0.00082989 |
| 17313 | MGP | 1849.84091 | 2.73257016 | 0.81754115 | 3.34242523 | 0.0008305 |

| | | | | | | |
|---|---|---|---|---|---|---|
| 381379 | MED19 | 1172.64394 | -0.3905182 | 0.11685749 | -3.3418332 | 0.00083227 |
| 66396 | CCDC82 | 443.927001 | -0.3543556 | 0.10604636 | -3.3415155 | 0.00083322 |
| 22209 | UBE2A | 1082.93563 | -0.2340529 | 0.07004569 | -3.3414325 | 0.00083347 |
| 116838 | RIMS2 | 387.651189 | 1.73871951 | 0.52050963 | 3.34041759 | 0.00083653 |
| 384281 | GATC | 706.048287 | 0.23382015 | 0.07002654 | 3.33902207 | 0.00084074 |
| 319909 | ISM1 | 94.7557707 | 1.95065569 | 0.58424197 | 3.3387805 | 0.00084147 |
| 67136 | KBTBD4 | 722.103117 | 0.4152738 | 0.12439699 | 3.3382947 | 0.00084294 |
| 269774 | AAK1 | 6259.72383 | 0.32019877 | 0.09593753 | 3.33757559 | 0.00084513 |
| 16173 | IL18 | 39.6393721 | -0.9040638 | 0.2708968 | -3.3372995 | 0.00084597 |
| 69723 | RPAIN | 227.997974 | -0.514379 | 0.15415152 | -3.3368401 | 0.00084737 |
| 69207 | SRSF11 | 3401.39307 | 0.2347865 | 0.0703835 | 3.33581757 | 0.00085049 |
| 93890 | Pcdhb19 | 51.3519539 | 1.45763141 | 0.43705298 | 3.33513668 | 0.00085257 |
| 13829 | DMTN | 56.5921727 | 0.78165045 | 0.23439693 | 3.33472993 | 0.00085382 |
| 12295 | CACNB1 | 43.4963676 | 1.00485033 | 0.301341 | 3.33459548 | 0.00085424 |
| 20562 | SLIT1 | 6.38890307 | -3.5546045 | 1.06633503 | -3.3334782 | 0.00085767 |
| 110380 | SHROOM2 | 656.76308 | -4.4101447 | 1.32376616 | -3.3315134 | 0.00086375 |
| 71354 | WDR31 | 34.7351711 | -2.587938 | 0.77683421 | -3.3313903 | 0.00086413 |
| 67475 | ERO1LB | 389.457048 | -0.4472216 | 0.13428587 | -3.3303694 | 0.00086731 |
| 544678 | CFAP74 | 22.6379379 | 1.1520541 | 0.34595258 | 3.33009256 | 0.00086817 |
| 625342 | Gm13315 | 164.516976 | 0.57439488 | 0.17248823 | 3.3300525 | 0.0008683 |
| 73991 | ATL1 | 76.4434756 | -0.6712607 | 0.20158319 | -3.3299439 | 0.00086863 |
| 19733 | RGN | 5.25586573 | -5.8204583 | 1.74793215 | -3.3299109 | 0.00086874 |
| 51960 | KCTD18 | 327.581756 | 0.37822035 | 0.11359383 | 3.32958529 | 0.00086975 |
| 408057 | BC039771 | 15.0255933 | 1.41851178 | 0.42610118 | 3.32904914 | 0.00087143 |
| 216810 | TOM1L2 | 2372.09826 | -0.4449047 | 0.13365869 | -3.3286628 | 0.00087264 |
| 68059 | TM9SF2 | 4309.17784 | -0.8928275 | 0.26824292 | -3.328429 | 0.00087337 |
| 80859 | nfkbiz | 3052.94429 | 0.6290188 | 0.18898584 | 3.32839122 | 0.00087349 |
| 67463 | POC5 | 624.637919 | 0.53745371 | 0.1614818 | 3.32826191 | 0.0008739 |
| 12824 | COL2A1 | 120.019692 | -2.8862812 | 0.8676061 | -3.3267184 | 0.00087875 |
| 15441 | HP1BP3 | 4576.79899 | 0.37258603 | 0.11199873 | 3.32669884 | 0.00087881 |
| 75665 | CCDC64 | 57.8619039 | -2.5160894 | 0.75634769 | -3.3266307 | 0.00087903 |
| 100233175 | AK010878 | 152.791899 | 0.65285009 | 0.19626562 | 3.32635984 | 0.00087988 |
| 17248 | MDM4 | 919.446142 | -0.5262605 | 0.15825682 | -3.3253575 | 0.00088305 |
| 51885 | TUBGCP4 | 1335.32144 | 0.37608297 | 0.11309927 | 3.32524676 | 0.0008834 |
| 230101 | GBA2 | 302.451315 | -0.9411178 | 0.28307452 | -3.3246293 | 0.00088536 |
| 29817 | IGFBP7 | 2326.07982 | -1.7840845 | 0.53686181 | -3.3231727 | 0.00089 |
| 105246 | BRD9 | 996.632208 | 0.64673162 | 0.19461835 | 3.32307629 | 0.00089031 |
| 381066 | Zfp948 | 409.636113 | -1.4416891 | 0.43397323 | -3.3220691 | 0.00089353 |
| 320709 | TMEM117 | 226.518228 | 0.92443058 | 0.27828265 | 3.32191234 | 0.00089403 |
| 68137 | KDELR1 | 3452.72417 | 0.57944805 | 0.17449273 | 3.3207576 | 0.00089773 |
| 320435 | RINL | 8.29326506 | -2.3383993 | 0.70426928 | -3.3203199 | 0.00089914 |
| 67525 | TRIM62 | 169.675362 | 0.57815509 | 0.17417809 | 3.31933304 | 0.00090233 |
| 218138 | GMDS | 531.211922 | -0.6353261 | 0.19142737 | -3.3188882 | 0.00090377 |
| 68118 | ATG101 | 689.163336 | -0.3894911 | 0.11735947 | -3.3187869 | 0.00090409 |
| 66664 | TMEM41A | 274.593667 | -0.5323162 | 0.16044141 | -3.3178233 | 0.00090722 |

| | | | | | | |
|---|---|---|---|---|---|---|
| 66409 | RSL1D1 | 6599.48355 | 0.25638726 | 0.07728609 | 3.31737905 | 0.00090866 |
| 13612 | EDIL3 | 345.331576 | 2.41241518 | 0.72728939 | 3.31699489 | 0.00090991 |
| 67841 | ATG3 | 1263.99742 | -0.236468 | 0.07129182 | -3.316902 | 0.00091022 |
| 214505 | GNPTG | 424.670295 | -0.7951819 | 0.23984664 | -3.3153766 | 0.0009152 |
| 11982 | ATP10A | 1950.08458 | 0.5349742 | 0.16139599 | 3.31466846 | 0.00091752 |
| 664862 | Gpr137b-ps | 1118.02691 | -0.8044122 | 0.24274807 | -3.3137741 | 0.00092046 |
| 67512 | AGPAT2 | 495.087362 | -0.5712321 | 0.17238511 | -3.3136975 | 0.00092071 |
| 268354 | FAM19A2 | 5.1800908 | -2.8640538 | 0.86447424 | -3.3130586 | 0.00092282 |
| 107589 | MYLK | 41.6496196 | -1.4507487 | 0.43804532 | -3.311869 | 0.00092675 |
| 56438 | RBX1 | 1251.53772 | -0.2208241 | 0.06670917 | -3.3102517 | 0.00093212 |
| 622675 | Zfp827 | 877.170409 | 0.44213623 | 0.13358965 | 3.30965935 | 0.0009341 |
| 276905 | ARMC7 | 142.475586 | -0.4276947 | 0.12922914 | -3.3095839 | 0.00093435 |
| 20725 | SERPINB8 | 884.434546 | -1.0462647 | 0.31613235 | -3.3095782 | 0.00093437 |
| 94229 | SLC4A10 | 13.9187181 | -4.3143636 | 1.30368247 | -3.3093669 | 0.00093507 |
| 16885 | LIMK1 | 888.72088 | 1.16611336 | 0.35249963 | 3.30812648 | 0.00093922 |
| 109299 | C330006A16 | 1081.95299 | 0.27404964 | 0.08287737 | 3.30668826 | 0.00094406 |
| 21679 | TEAD4 | 1826.35503 | 0.57294959 | 0.17328478 | 3.30640466 | 0.00094502 |
| 22059 | Trp53 | 4600.03295 | 0.20264764 | 0.06130677 | 3.30546934 | 0.00094817 |
| 19326 | RAB11B | 830.036793 | -0.3694077 | 0.11176039 | -3.3053545 | 0.00094856 |
| 229593 | GOLPH3L | 707.885171 | -0.292442 | 0.08848056 | -3.3051558 | 0.00094924 |
| 68146 | ARL13B | 790.014291 | -0.3660438 | 0.11074972 | -3.3051444 | 0.00094927 |
| 170755 | SGK3 | 302.119602 | -1.9103694 | 0.57811928 | -3.3044555 | 0.00095161 |
| 12322 | CAMK2A | 13.3447231 | -4.6211992 | 1.39883276 | -3.3036109 | 0.00095448 |
| 66958 | TMX2 | 411.879993 | -0.4247146 | 0.12856086 | -3.3036072 | 0.00095449 |
| 71592 | POGK | 756.048223 | 2.30121496 | 0.69674023 | 3.30283063 | 0.00095714 |
| 243574 | KBTBD8 | 248.456907 | 0.741572 | 0.2245499 | 3.30248204 | 0.00095833 |
| 327747 | mettl24 | 26.1046341 | -1.9032108 | 0.57631147 | -3.3023997 | 0.00095861 |
| 66126 | ELOF1 | 1219.92947 | 0.25271461 | 0.07655133 | 3.30124389 | 0.00096257 |
| 100465 | MOB3C | 133.289866 | 0.49018034 | 0.14852651 | 3.30028861 | 0.00096585 |
| 264134 | TTC26 | 311.26724 | -0.5764717 | 0.17472196 | -3.2993659 | 0.00096904 |
| 230376 | HAUS6 | 1858.8673 | 0.6720032 | 0.20369889 | 3.29900283 | 0.00097029 |
| 242627 | Skint5 | 19.3409991 | -1.5640434 | 0.47416419 | -3.2985269 | 0.00097194 |
| 320534 | TMEM104 | 613.823966 | -0.5583951 | 0.16938476 | -3.2966074 | 0.0009786 |
| 236546 | AF067061 | 3.51445202 | -5.238939 | 1.58925525 | -3.2964742 | 0.00097907 |
| 26905 | Eif2s3x | 475.39775 | -0.5113911 | 0.15521438 | -3.2947408 | 0.00098513 |
| 101966 | D8Ertd738e | 1354.86961 | -0.3104145 | 0.09422742 | -3.294312 | 0.00098663 |
| 244723 | OLFM2 | 25.5332096 | 3.37990817 | 1.0259914 | 3.29428509 | 0.00098672 |
| 69253 | HSPB2 | 119.058805 | 1.89082175 | 0.57411727 | 3.29344167 | 0.00098969 |
| 381678 | ZCWPW1 | 67.8113746 | -1.4706866 | 0.44658124 | -3.2932118 | 0.0009905 |
| 69663 | DDX51 | 1165.44245 | -0.2014466 | 0.06117454 | -3.2929812 | 0.00099131 |
| 22228 | UCP2 | 2265.95171 | -1.332048 | 0.40453733 | -3.2927691 | 0.00099206 |
| 68170 | B230118H07 | 317.144334 | -0.4858357 | 0.14754689 | -3.2927547 | 0.00099211 |
| 110379 | SEC13 | 3811.5187 | 0.45462981 | 0.13809636 | 3.29212011 | 0.00099435 |
| 63986 | GMFG | 4.62942914 | -4.8832309 | 1.48369731 | -3.2912582 | 0.0009974 |
| 54604 | pcnx | 2536.11785 | 0.66957239 | 0.20344594 | 3.29115637 | 0.00099776 |

| | | | | | | |
|---|---|---|---|---|---|---|
| 76947 | NDUFAF6 | 170.508956 | -0.7950004 | 0.24157394 | -3.2909195 | 0.0009986 |
| 66964 | GOLT1B | 2956.44867 | 0.48555861 | 0.14754587 | 3.29089939 | 0.00099868 |
| 15162 | HCK | 2.82831749 | -4.9276356 | 1.4974518 | -3.2906806 | 0.00099945 |
| 71146 | GOLGA7B | 11.5989146 | -1.9532925 | 0.59362017 | -3.2904753 | 0.00100018 |
| 12348 | Car11 | 3.9706317 | 4.70511855 | 1.4303336 | 3.2895253 | 0.00100357 |
| 11304 | ABCA4 | 39.0859816 | -3.6243237 | 1.10182787 | -3.2893738 | 0.00100411 |
| 238257 | TMEM30B | 3.42833039 | -5.2064419 | 1.58311185 | -3.2887392 | 0.00100637 |
| 69863 | TTC39B | 4245.56031 | 0.54901533 | 0.16693803 | 3.28873725 | 0.00100638 |
| 12866 | COX7A2 | 1572.06819 | 0.22996856 | 0.06993143 | 3.2884862 | 0.00100728 |
| 19330 | RAB18 | 4908.56462 | -0.1738033 | 0.0528532 | -3.2884168 | 0.00100753 |
| 55989 | NOP58 | 5130.61857 | -0.506417 | 0.15403459 | -3.2876835 | 0.00101015 |
| 665574 | Gm7694 | 6.22420904 | -2.9103906 | 0.88527998 | -3.2875369 | 0.00101068 |
| 12315 | CALM3 | 5666.76419 | 0.51825522 | 0.15766123 | 3.28714432 | 0.00101209 |
| 56808 | CACNA2D2 | 5.65918511 | -4.4022393 | 1.33928023 | -3.2870188 | 0.00101254 |
| 79221 | HDAC9 | 9.61986951 | 2.08822651 | 0.63530646 | 3.28695936 | 0.00101275 |
| 619605 | ZCCHC17 | 526.89613 | 0.63003855 | 0.19171171 | 3.2863853 | 0.00101482 |
| 19134 | PRPF4B | 3138.93172 | 0.35538007 | 0.10815476 | 3.28584787 | 0.00101676 |
| 212377 | MMS22L | 1500.94649 | -0.9141395 | 0.27824219 | -3.2854094 | 0.00101834 |
| 54125 | POLM | 1734.39925 | 0.22985044 | 0.07000867 | 3.28317115 | 0.00102646 |
| 109232 | SCCPDH | 964.035132 | -0.2217659 | 0.0675493 | -3.2830235 | 0.001027 |
| 433466 | JMJD7 | 181.042664 | 0.39928993 | 0.12173314 | 3.28004303 | 0.00103791 |
| 67304 | 3110070M22 | 12.9147811 | 1.67880924 | 0.51183556 | 3.27997773 | 0.00103815 |
| 235459 | GTF2A2 | 879.405534 | 0.18792144 | 0.05729621 | 3.27982328 | 0.00103872 |
| 215708 | MIGA1 | 925.637968 | -0.6558215 | 0.19997587 | -3.279503 | 0.0010399 |
| 13638 | EFNA3 | 23.5353599 | 1.3509315 | 0.41199944 | 3.27896439 | 0.00104189 |
| 226409 | ZRANB3 | 283.026048 | -0.4623777 | 0.14105746 | -3.2779386 | 0.00104568 |
| 239096 | cdh24 | 195.260753 | -0.9791601 | 0.29878155 | -3.2771772 | 0.00104851 |
| 17188 | MAZ | 4392.4714 | 0.72584758 | 0.22151072 | 3.27680559 | 0.00104989 |
| 66889 | RNF128 | 532.133816 | -1.4355444 | 0.43811056 | -3.2766715 | 0.00105039 |
| 74778 | RRP7A | 1423.41217 | 0.20829836 | 0.06357912 | 3.27620699 | 0.00105211 |
| 207965 | VCPKMT | 539.095932 | 0.39640784 | 0.12102162 | 3.27551266 | 0.0010547 |
| 72477 | TMEM87B | 875.991504 | -0.4284022 | 0.13079017 | -3.2754924 | 0.00105478 |
| 233529 | KCTD14 | 6.89732241 | -4.702619 | 1.43572833 | -3.2754239 | 0.00105504 |
| 121022 | mrps6 | 1613.30643 | -0.4201768 | 0.12828287 | -3.2753928 | 0.00105515 |
| 67917 | ZCCHC3 | 278.902392 | 0.55172678 | 0.1684563 | 3.27519232 | 0.0010559 |
| 100986 | AKAP9 | 2032.87115 | -0.5630097 | 0.17190725 | -3.275078 | 0.00105633 |
| 70823 | HMGXB4 | 721.254239 | 0.36334428 | 0.11094419 | 3.27501855 | 0.00105655 |
| 12562 | CDH5 | 28.4663575 | 3.00967786 | 0.91900021 | 3.27494794 | 0.00105681 |
| 20474 | SIX4 | 575.728325 | 0.61059114 | 0.1864877 | 3.27416311 | 0.00105975 |
| 13360 | DHCR7 | 2868.01567 | -0.554611 | 0.16943516 | -3.2732935 | 0.00106302 |
| 56089 | RAMP3 | 56.5442938 | -0.7678584 | 0.23466295 | -3.2721758 | 0.00106723 |
| 21933 | TNFRSF10B | 1707.97697 | -1.189365 | 0.36349518 | -3.2720242 | 0.0010678 |
| 53607 | SNRPA | 1203.99468 | 0.36035671 | 0.11014686 | 3.27160224 | 0.0010694 |
| 225849 | PPP2R5B | 702.266869 | -0.4898859 | 0.14974394 | -3.271491 | 0.00106982 |
| 56456 | ACTL6A | 2075.03735 | 0.40780435 | 0.12465402 | 3.27148981 | 0.00106982 |

| | | | | | | |
|---|---|---|---|---|---|---|
| 381933 | 6430531B16 | 4.07108978 | -4.6788308 | 1.4304644 | -3.2708474 | 0.00107226 |
| 70335 | REEP6 | 85.6595563 | 1.21900803 | 0.37276481 | 3.27018004 | 0.00107479 |
| 77521 | MTUS2 | 4.23832653 | -4.7356081 | 1.44861643 | -3.2690559 | 0.00107907 |
| 68273 | POMGNT1 | 1863.85683 | 0.79814381 | 0.24417114 | 3.26878847 | 0.00108009 |
| 170719 | OXR1 | 5821.8519 | -0.9181551 | 0.28089143 | -3.2687188 | 0.00108036 |
| 98363 | EFHD1 | 16.0884196 | -2.1716618 | 0.66440497 | -3.2685814 | 0.00108088 |
| 101985 | USB1 | 574.358947 | 0.30923953 | 0.09462479 | 3.26806039 | 0.00108287 |
| 69772 | BDH2 | 30.6283338 | -1.7679136 | 0.54098053 | -3.26798 | 0.00108318 |
| 50723 | Icosl | 45.8605942 | 0.92761451 | 0.28386226 | 3.26783313 | 0.00108374 |
| 66451 | 2610528J11F | 3.6373027 | -5.2971205 | 1.62098878 | -3.2678329 | 0.00108374 |
| 100503670 | RPL5 | 516.037854 | -0.3251088 | 0.09948918 | -3.2677808 | 0.00108394 |
| 58909 | FAM13A | 128.521315 | -1.3472761 | 0.41235679 | -3.2672581 | 0.00108595 |
| 223431 | Tiaf2 | 4.61352195 | 4.92918708 | 1.50867064 | 3.26723869 | 0.00108602 |
| 667253 | Gm8539 | 105.659573 | 0.61927283 | 0.1896128 | 3.2659864 | 0.00109083 |
| 229841 | CENPE | 5781.59825 | 0.36375703 | 0.11138198 | 3.2658517 | 0.00109135 |
| 13386 | DLK1 | 3.43460534 | 5.25888086 | 1.61054227 | 3.26528584 | 0.00109354 |
| 21372 | TBL1X | 2400.29646 | -0.3057795 | 0.09365561 | -3.2649352 | 0.00109489 |
| 244421 | LONRF1 | 165.34546 | -1.1378989 | 0.34855535 | -3.2646145 | 0.00109613 |
| 242702 | MYOM3 | 4.3652259 | -4.01722 | 1.23067993 | -3.2642281 | 0.00109763 |
| 71898 | Apol9b | 121.698023 | -0.7734439 | 0.23695001 | -3.2641647 | 0.00109787 |
| 93742 | PARD3 | 2812.50785 | -0.28192 | 0.08637167 | -3.264033 | 0.00109838 |
| 18451 | P4HA1 | 2622.44322 | 1.02064921 | 0.31279102 | 3.26303868 | 0.00110224 |
| 546546 | Serpina3h | 4274.98762 | -1.1238988 | 0.34444711 | -3.2629066 | 0.00110276 |
| 108954 | PPP1R15B | 5747.93561 | 0.24794833 | 0.07599076 | 3.26287475 | 0.00110288 |
| 71877 | EFHC1 | 16.2710518 | -1.7933049 | 0.54972596 | -3.2621798 | 0.00110559 |
| 66053 | PPIL2 | 1400.95154 | -0.1867388 | 0.05724716 | -3.2619747 | 0.00110639 |
| 66171 | PGLS | 1425.41008 | 0.29078039 | 0.08917259 | 3.26087195 | 0.0011107 |
| 208677 | CREB3L3 | 5.25808894 | -3.7037205 | 1.13587271 | -3.2606827 | 0.00111144 |
| 56445 | DNAJA2 | 6792.88364 | 0.25021064 | 0.07674479 | 3.2602949 | 0.00111296 |
| 544971 | BDP1 | 2277.57337 | 0.59829326 | 0.18352067 | 3.26008648 | 0.00111378 |
| 77605 | H2AFV | 582.651326 | -0.4370387 | 0.13408073 | -3.2595195 | 0.00111601 |
| 235497 | LEO1 | 1197.14673 | 0.38582827 | 0.11837045 | 3.25949804 | 0.0011161 |
| 72701 | Zfp618 | 23.7675687 | 1.47651186 | 0.45300476 | 3.25937381 | 0.00111658 |
| 269643 | PPP2R2C | 3.81857178 | -5.3579062 | 1.6439142 | -3.2592372 | 0.00111712 |
| 627214 | FAM196A | 3.21033227 | -5.1102375 | 1.5687223 | -3.2575794 | 0.00112367 |
| 66202 | 1110059G10 | 239.494298 | 0.40955072 | 0.12575769 | 3.25666549 | 0.00112729 |
| 21664 | PHLDA1 | 2594.82399 | 0.56488171 | 0.17348757 | 3.25603561 | 0.0011298 |
| 234135 | whsc1l1 | 2655.55375 | 0.21797655 | 0.06696008 | 3.25532101 | 0.00113264 |
| 109181 | TRIP11 | 1778.36182 | 0.26844312 | 0.0824685 | 3.25509869 | 0.00113353 |
| 30045 | DNAJC12 | 74.8446449 | -0.7956938 | 0.24450691 | -3.2542795 | 0.0011368 |
| 625631 | Bloc1s2-ps | 234.459316 | -0.4087966 | 0.12564445 | -3.2535987 | 0.00113953 |
| 380969 | NCKAP5L | 250.268824 | 1.40279227 | 0.43122295 | 3.25305572 | 0.00114171 |
| 225432 | RBM27 | 2714.62278 | 0.21479416 | 0.06603738 | 3.25261462 | 0.00114348 |
| 56200 | DDX21 | 12987.4547 | 0.30319286 | 0.09323906 | 3.25177949 | 0.00114685 |
| 74385 | AP5M1 | 456.177219 | 0.7493873 | 0.23051893 | 3.25087094 | 0.00115052 |

| | | | | | | |
|---|---|---|---|---|---|---|
| 216049 | Zfp365 | 1854.96538 | -0.6266739 | 0.19280212 | -3.2503479 | 0.00115264 |
| 269113 | NUP54 | 991.538275 | 0.27207929 | 0.08372219 | 3.24978698 | 0.00115491 |
| 66487 | smim4 | 116.055779 | 0.94278681 | 0.29013212 | 3.24950856 | 0.00115605 |
| 100504307 | Gm20163 | 11.4641238 | 2.5680839 | 0.79035479 | 3.24927986 | 0.00115698 |
| 109857 | cbr3 | 491.792101 | 1.80733345 | 0.55636477 | 3.24846855 | 0.00116028 |
| 320770 | A630072M18 | 102.973959 | 0.59197337 | 0.18226999 | 3.24778297 | 0.00116308 |
| 18201 | NSMAF | 1274.45447 | -0.7905569 | 0.24345503 | -3.24724 | 0.0011653 |
| 75234 | RNF19B | 1782.03468 | -0.7810448 | 0.24054793 | -3.2469406 | 0.00116653 |
| 104271 | TEX15 | 81.774216 | -3.3320168 | 1.02638102 | -3.2463741 | 0.00116885 |
| 353282 | SFMBT2 | 661.403922 | 0.66886412 | 0.20605059 | 3.24611607 | 0.00116991 |
| 53418 | B4GALT2 | 755.132783 | 3.3496691 | 1.03197204 | 3.24589133 | 0.00117084 |
| 70769 | NOLC1 | 5978.01843 | 0.4079205 | 0.12567655 | 3.24579642 | 0.00117123 |
| 18971 | POLD1 | 1670.17766 | 0.67825344 | 0.20898731 | 3.24542878 | 0.00117274 |
| 100322896 | DTHD1 | 4.70998155 | -5.6623884 | 1.74490968 | -3.2450897 | 0.00117414 |
| 238023 | HEXDC | 209.28427 | -0.6479595 | 0.19976848 | -3.2435522 | 0.00118049 |
| 110094 | PHKA2 | 713.140847 | -0.2339904 | 0.07216853 | -3.2422768 | 0.00118579 |
| 56426 | PDCD10 | 446.649054 | -0.4395948 | 0.13561301 | -3.2415387 | 0.00118886 |
| 332397 | NANOS1 | 216.759333 | 0.48156831 | 0.14856182 | 3.24153486 | 0.00118888 |
| 229228 | NUDT6 | 97.5964789 | 0.5531042 | 0.17063309 | 3.24148262 | 0.0011891 |
| 14149 | FDXR | 317.22562 | 0.52935511 | 0.16331575 | 3.24129854 | 0.00118987 |
| 18126 | NOS2 | 114.944119 | -2.1782608 | 0.6721046 | -3.2409551 | 0.0011913 |
| 217143 | GPR179 | 11.3873164 | -2.2158121 | 0.68372524 | -3.2407932 | 0.00119198 |
| 64929 | scel | 16.6732476 | -2.648538 | 0.81732496 | -3.2404957 | 0.00119322 |
| 12788 | CNGA1 | 12.5295317 | -7.0785761 | 2.18477466 | -3.2399571 | 0.00119548 |
| 240613 | 9930021J03R | 1066.23441 | -0.2250359 | 0.06945729 | -3.2399174 | 0.00119564 |
| 320742 | A230072C01 | 28.799828 | -1.3030956 | 0.40227692 | -3.2392999 | 0.00119824 |
| 634834 | Gm11821 | 69.8623657 | -0.6665109 | 0.20577871 | -3.2389693 | 0.00119962 |
| 74238 | MTERF2 | 80.8395033 | -0.6953922 | 0.21470816 | -3.2387786 | 0.00120043 |
| 218341 | RFESD | 99.7902857 | -0.5515741 | 0.17033669 | -3.2381406 | 0.00120312 |
| 66594 | UQCR11 | 755.713816 | -0.3700341 | 0.11430635 | -3.237214 | 0.00120703 |
| 66054 | CNDP2 | 2191.10724 | -0.7652507 | 0.23644056 | -3.2365458 | 0.00120986 |
| 15370 | NR4A1 | 528.653592 | 1.28034457 | 0.39562363 | 3.23626922 | 0.00121103 |
| 102141 | SNX25 | 1024.67664 | 0.2248883 | 0.06949455 | 3.23605674 | 0.00121193 |
| 142688 | ASB13 | 442.462242 | 0.40729624 | 0.12587162 | 3.23580678 | 0.00121299 |
| 15368 | HMOX1 | 7966.69618 | 1.00067995 | 0.30925543 | 3.23577167 | 0.00121314 |
| 56743 | LAT2 | 89.5459108 | -2.0476206 | 0.63283658 | -3.235623 | 0.00121378 |
| 225609 | Gm9949 | 14.7208837 | -2.2223275 | 0.68687696 | -3.2354085 | 0.00121469 |
| 66480 | RPL15 | 680.451444 | -0.6437157 | 0.19899738 | -3.2347949 | 0.0012173 |
| 100689703 | 9330133O14 | 82.6207736 | 1.02823784 | 0.31797987 | 3.23365705 | 0.00122216 |
| 100502670 | Gm16010 | 3.00973901 | -5.0143849 | 1.55077692 | -3.2334663 | 0.00122298 |
| 227933 | CCDC148 | 7.80570467 | -3.0003669 | 0.92799058 | -3.2331868 | 0.00122417 |
| 94218 | CNNM3 | 697.964575 | -0.2312937 | 0.07154743 | -3.232732 | 0.00122613 |
| 230737 | GNL2 | 2019.84859 | 0.47014824 | 0.14543851 | 3.23262556 | 0.00122658 |
| 380698 | OBSCN | 4.80846266 | -3.582578 | 1.10838476 | -3.2322512 | 0.00122819 |
| 20742 | SPTBN1 | 18462.7727 | 0.32812221 | 0.10151732 | 3.23217974 | 0.0012285 |

| | | | | | | |
|---|---|---|---|---|---|---|
| 226178 | WBP1L | 3319.39207 | -0.3480608 | 0.10770897 | -3.2314929 | 0.00123145 |
| 18845 | PLXNA2 | 1588.12191 | -1.0545379 | 0.32643533 | -3.2304649 | 0.00123589 |
| 67064 | CHMP1B | 1365.57558 | -0.2898825 | 0.08975831 | -3.22959 | 0.00123968 |
| 320466 | A230103J11F | 10.2745961 | -2.4245173 | 0.75076765 | -3.2293843 | 0.00124057 |
| 12039 | BCKDHA | 503.900427 | -0.2371043 | 0.07342529 | -3.2291914 | 0.00124141 |
| 433408 | Gm13375 | 123.619239 | 0.8390498 | 0.25989967 | 3.22836041 | 0.00124502 |
| 19263 | PTPRB | 14.17826 | 3.07300451 | 0.95188129 | 3.22834848 | 0.00124507 |
| 74772 | ATP13A2 | 665.160714 | -0.7163918 | 0.22193812 | -3.2278898 | 0.00124707 |
| 216874 | CAMTA2 | 792.949332 | 0.49340974 | 0.15287009 | 3.2276409 | 0.00124816 |
| 12567 | CDK4 | 9274.84591 | 0.18389326 | 0.05698731 | 3.22691565 | 0.00125132 |
| 269587 | EPB41 | 1059.44879 | -0.8799416 | 0.27273639 | -3.2263447 | 0.00125382 |
| 238799 | TNPO1 | 5975.95691 | 0.46176253 | 0.14314071 | 3.22593431 | 0.00125562 |
| 227937 | PKP4 | 2812.52334 | 0.19444014 | 0.06027572 | 3.225845 | 0.00125601 |
| 235461 | FAM63B | 1439.22807 | -0.7795171 | 0.24167774 | -3.2254404 | 0.00125779 |
| 17001 | LTC4S | 2.73490726 | -4.8770368 | 1.51209403 | -3.2253528 | 0.00125818 |
| 66116 | Nat8f1 | 12.3002269 | 1.7346268 | 0.53790428 | 3.22478717 | 0.00126066 |
| 170829 | TRAM2 | 317.884616 | 0.43407478 | 0.13460584 | 3.22478427 | 0.00126068 |
| 545812 | Pilrb2 | 2.93079406 | -4.9726892 | 1.54265515 | -3.2234613 | 0.00126651 |
| 100113398 | ADAT3 | 162.92598 | 0.39117039 | 0.12135534 | 3.2233472 | 0.00126702 |
| 69155 | 1810030O07 | 934.749585 | -0.3283075 | 0.10192945 | -3.2209289 | 0.00127776 |
| 17299 | METTL1 | 484.872258 | 0.38169074 | 0.11851971 | 3.22048317 | 0.00127975 |
| 330577 | SAXO2 | 7.715779 | -2.6994305 | 0.83825514 | -3.220297 | 0.00128058 |
| 319945 | FLAD1 | 434.126906 | -0.5616468 | 0.17442441 | -3.2200014 | 0.0012819 |
| 80515 | Chd3os | 118.311783 | 1.29209671 | 0.40133045 | 3.21953319 | 0.00128399 |
| 12632 | CFL2 | 3518.53973 | 0.29803597 | 0.09257282 | 3.21947607 | 0.00128425 |
| 68708 | rabl2 | 120.421383 | -0.5527684 | 0.17170219 | -3.219344 | 0.00128484 |
| 19206 | PTCH1 | 453.887856 | 0.63388085 | 0.19702065 | 3.21733209 | 0.00129389 |
| 17475 | MPDZ | 2894.79677 | 1.79253692 | 0.55743756 | 3.215673 | 0.00130139 |
| 51938 | CCDC39 | 18.0704298 | -2.0765581 | 0.64581469 | -3.2154087 | 0.00130259 |
| 75462 | 1700001C19I | 3.75029264 | -4.5684538 | 1.42125593 | -3.214378 | 0.00130727 |
| 69237 | GTPBP4 | 3039.19372 | 0.56640773 | 0.17622146 | 3.21418142 | 0.00130817 |
| 259300 | EHD2 | 5570.91734 | -0.542135 | 0.16866999 | -3.2141756 | 0.0013082 |
| 66165 | BCCIP | 2056.23377 | 0.56778256 | 0.17666938 | 3.21381424 | 0.00130984 |
| 52323 | KLHL7 | 941.169941 | 0.78935881 | 0.24564804 | 3.21337319 | 0.00131186 |
| 15414 | HOXB6 | 42.4940781 | -4.9071329 | 1.52738376 | -3.2127701 | 0.00131461 |
| 230674 | KDM4A | 2273.33274 | 0.50173019 | 0.15619683 | 3.21216632 | 0.00131738 |
| 72544 | EXOSC6 | 697.680302 | 0.40508496 | 0.12612284 | 3.2118287 | 0.00131893 |
| 22258 | USP4 | 3276.49361 | 0.27442611 | 0.08544455 | 3.21174496 | 0.00131931 |
| 104859 | TECPR2 | 787.704368 | -0.3478822 | 0.10832538 | -3.2114565 | 0.00132064 |
| 387524 | ZNRF2 | 1031.10149 | 0.5218895 | 0.16259386 | 3.20977368 | 0.0013284 |
| 140810 | TTBK2 | 853.901243 | 0.30128092 | 0.09388282 | 3.20911681 | 0.00133143 |
| 76799 | TMEM234 | 943.003986 | 0.4615365 | 0.14382177 | 3.20908652 | 0.00133157 |
| 109042 | PRKCDBP | 1287.97348 | 2.92252395 | 0.91152815 | 3.20618069 | 0.00134509 |
| 20399 | SH2B1 | 1316.92823 | -0.158747 | 0.04951358 | -3.2061309 | 0.00134533 |
| 68420 | ANKRD13A | 3466.87605 | 0.17504696 | 0.05459907 | 3.20604288 | 0.00134574 |

| | | | | | | |
|---|---|---|---|---|---|---|
| 72046 | URGCP | 1961.36814 | 0.2380293 | 0.07425692 | 3.20548307 | 0.00134836 |
| 26927 | FOXL2 | 129.627668 | 3.17202294 | 0.98996423 | 3.20417936 | 0.00135448 |
| 319480 | ITGA11 | 6.45222411 | -3.5968077 | 1.12280354 | -3.2034168 | 0.00135807 |
| 272589 | TBCEL | 922.487377 | 0.97643675 | 0.30481612 | 3.20336323 | 0.00135833 |
| 320538 | UBN2 | 1592.56278 | 0.38979096 | 0.12168761 | 3.20320994 | 0.00135905 |
| 72096 | METTL10 | 331.35638 | 0.4949842 | 0.15459442 | 3.20182451 | 0.0013656 |
| 56048 | LGALS8 | 2946.23798 | 0.62448531 | 0.19505435 | 3.20159643 | 0.00136668 |
| 14593 | GGPS1 | 559.293606 | 0.41517397 | 0.12968636 | 3.20136963 | 0.00136776 |
| 77951 | CYP20A1 | 1322.79185 | -0.2558369 | 0.07992427 | -3.2009916 | 0.00136956 |
| 100503099 | Gm19554 | 5.01892121 | -4.9899639 | 1.55900726 | -3.2007317 | 0.00137079 |
| 107932 | CHD4 | 17253.8697 | 0.27042522 | 0.08451288 | 3.19981053 | 0.00137518 |
| 277328 | TRPA1 | 16.7475229 | -7.4953873 | 2.34333051 | -3.1986044 | 0.00138095 |
| 22256 | UNG | 448.053142 | 0.27730807 | 0.08670921 | 3.19813861 | 0.00138318 |
| 117149 | TIRAP | 942.945667 | 0.31504257 | 0.09855037 | 3.19676685 | 0.00138977 |
| 237246 | BC022960 | 25.5073221 | -1.2664659 | 0.39622137 | -3.1963593 | 0.00139174 |
| 233913 | BC017158 | 604.29225 | 0.51392646 | 0.16079587 | 3.19614218 | 0.00139278 |
| 208643 | EIF4G1 | 38326.795 | 0.10767781 | 0.03369906 | 3.19527618 | 0.00139697 |
| 330599 | Gm16938 | 5.58875539 | -2.7196566 | 0.85126953 | -3.1948243 | 0.00139916 |
| 83964 | JAM3 | 5.73868299 | -5.1974232 | 1.62693938 | -3.1946016 | 0.00140024 |
| 22193 | UBE2E3 | 1309.96214 | 0.3915708 | 0.12261982 | 3.19337277 | 0.00140621 |
| 26554 | CUL3 | 5612.02848 | -0.2409102 | 0.07545704 | -3.1926799 | 0.00140959 |
| 214663 | SLC25A29 | 238.072495 | 0.76633895 | 0.24003323 | 3.19263689 | 0.0014098 |
| 107197 | UQCC3 | 268.14396 | 0.33511398 | 0.10496823 | 3.19252756 | 0.00141033 |
| 15185 | HDAC6 | 1116.46494 | -0.4059962 | 0.12717149 | -3.1925097 | 0.00141042 |
| 67037 | PMF1 | 1226.9604 | 0.20343227 | 0.06373036 | 3.19207783 | 0.00141253 |
| 52477 | ANGEL2 | 1469.88498 | 0.15752946 | 0.04935723 | 3.19161884 | 0.00141478 |
| 68597 | CCDC167 | 240.726981 | -0.519985 | 0.16293215 | -3.1914206 | 0.00141575 |
| 241576 | LDLRAD3 | 473.349538 | 2.15998164 | 0.67686996 | 3.19113238 | 0.00141716 |
| 72275 | 2200002D01 | 854.53894 | -0.7174363 | 0.22485092 | -3.19072 | 0.00141919 |
| 18010 | NEU1 | 1341.05241 | -0.9850266 | 0.30884076 | -3.189432 | 0.00142553 |
| 319565 | SYNE2 | 2062.02089 | 0.96455071 | 0.30244283 | 3.18920015 | 0.00142667 |
| 66999 | MED28 | 1343.12903 | -0.2298883 | 0.07209467 | -3.1887004 | 0.00142914 |
| 268822 | adck5 | 498.691484 | -0.2830453 | 0.0887796 | -3.1881799 | 0.00143171 |
| 12237 | BUB3 | 3890.0475 | 0.54662902 | 0.17154627 | 3.18648147 | 0.00144015 |
| 13417 | DNAH8 | 8.07593916 | 2.19221715 | 0.68828706 | 3.18503323 | 0.00144737 |
| 94179 | KRT23 | 15.2900756 | -7.3653415 | 2.31360758 | -3.1834878 | 0.00145512 |
| 269254 | SETX | 4492.51702 | -0.3250127 | 0.10211137 | -3.1829238 | 0.00145796 |
| 171543 | BMF | 46.2481921 | -1.663989 | 0.52281529 | -3.1827474 | 0.00145885 |
| 18164 | NPTX1 | 2972.18105 | -5.1681394 | 1.62399637 | -3.1823589 | 0.00146081 |
| 668773 | Gm9349 | 46.139187 | -0.8446812 | 0.26543683 | -3.1822304 | 0.00146146 |
| 105148 | IARS | 6029.75516 | 0.54505425 | 0.17129838 | 3.18189959 | 0.00146313 |
| 207618 | Zfp804b | 4.04374659 | -5.4390578 | 1.70975991 | -3.1811822 | 0.00146675 |
| 208439 | KLHL29 | 3.61462634 | -4.5032711 | 1.41559842 | -3.1811784 | 0.00146677 |
| 71208 | 4933440M02 | 6.45762687 | 2.27226681 | 0.71442201 | 3.18056666 | 0.00146987 |
| 12369 | CASP7 | 295.249285 | -0.5524603 | 0.1737205 | -3.1801679 | 0.0014719 |

| | | | | | | |
|---|---|---|---|---|---|---|
| 12531 | cdc25b | 2371.38863 | 0.4295417 | 0.1350807 | 3.17988957 | 0.00147331 |
| 68952 | FAM57B | 8.50047328 | -2.9201556 | 0.91833596 | -3.1798337 | 0.0014736 |
| 108014 | SRSF9 | 2231.18074 | 0.18399784 | 0.05787678 | 3.17913026 | 0.00147718 |
| 231128 | FAM193A | 1402.68021 | -0.1715725 | 0.0539985 | -3.1773557 | 0.00148625 |
| 234736 | RFWD3 | 1616.44009 | 0.3482198 | 0.10960633 | 3.17700449 | 0.00148805 |
| 68463 | MRPL14 | 639.545412 | -0.2527994 | 0.0795758 | -3.1768381 | 0.0014889 |
| 76497 | PPP1R11 | 1716.81793 | 0.39198206 | 0.12340049 | 3.17650319 | 0.00149062 |
| 19047 | PPP1CC | 291.561638 | 0.42892666 | 0.13506256 | 3.17576292 | 0.00149443 |
| 11308 | ABI1 | 4282.1924 | 0.22070539 | 0.069497 | 3.17575433 | 0.00149447 |
| 68077 | GLTSCR2 | 1771.46499 | 0.48012568 | 0.15121137 | 3.17519569 | 0.00149735 |
| 114654 | LY6G6D | 4.18186694 | -5.4979118 | 1.7315519 | -3.1751354 | 0.00149767 |
| 11504 | adamts1 | 2144.13801 | 0.97555473 | 0.30726526 | 3.17495946 | 0.00149857 |
| 73284 | DDIT4L | 803.839178 | -0.5196795 | 0.16368598 | -3.1748567 | 0.00149911 |
| 100039707 | Mthfsl | 37.9029677 | 0.85830231 | 0.27036067 | 3.17465671 | 0.00150014 |
| 14272 | FNTA | 1760.01903 | 0.31672045 | 0.0998004 | 3.17353892 | 0.00150593 |
| 67442 | RETSAT | 674.212902 | 0.48013873 | 0.15130871 | 3.17323923 | 0.00150748 |
| 16332 | INPPL1 | 5579.73658 | 0.68423493 | 0.21567542 | 3.17252166 | 0.00151121 |
| 68051 | NUTF2 | 209.197013 | -0.4651591 | 0.14663065 | -3.1723186 | 0.00151227 |
| 22341 | VEGFC | 617.404902 | -1.7676093 | 0.55726239 | -3.1719516 | 0.00151418 |
| 230789 | FAM76A | 765.943077 | 0.47930419 | 0.1511085 | 3.17192074 | 0.00151434 |
| 66625 | PNISR | 1426.77889 | -0.716929 | 0.22607883 | -3.1711461 | 0.00151839 |
| 50928 | KLRG1 | 4.08704208 | -5.4566989 | 1.72178171 | -3.1692164 | 0.00152851 |
| 232016 | CCDC129 | 19.1221106 | -5.6435905 | 1.78127156 | -3.1682931 | 0.00153337 |
| 21939 | CD40 | 4.13457468 | -5.4723657 | 1.7274651 | -3.1678589 | 0.00153566 |
| 18141 | NUP50 | 3333.72675 | 0.16646763 | 0.05255247 | 3.16764606 | 0.00153678 |
| 52468 | CTDSP2 | 3051.97459 | 0.17537859 | 0.05538408 | 3.16658834 | 0.00154238 |
| 69076 | TRIAP1 | 531.852494 | -0.35375 | 0.11175708 | -3.1653474 | 0.00154898 |
| 231570 | A830010M2( | 134.08789 | -0.6105635 | 0.19297692 | -3.1639199 | 0.0015566 |
| 69082 | ZC3H15 | 1689.56782 | 0.22366854 | 0.07071702 | 3.16286695 | 0.00156224 |
| 14470 | RABAC1 | 1255.28443 | -0.5610204 | 0.17740986 | -3.1622843 | 0.00156537 |
| 93705 | Pcdhgb8 | 69.1408562 | -7.1295827 | 2.25531554 | -3.1612351 | 0.00157102 |
| 103967 | DNM3 | 33.3208427 | -1.6330164 | 0.51660984 | -3.1610245 | 0.00157215 |
| 56715 | RABGEF1 | 1169.03592 | -0.3730038 | 0.11801356 | -3.1606859 | 0.00157398 |
| 111173 | ERC1 | 3301.5198 | 0.39738865 | 0.12574627 | 3.16024209 | 0.00157638 |
| 77110 | GPBP1L1 | 743.336117 | 0.45963155 | 0.14546692 | 3.15969802 | 0.00157933 |
| 64707 | SUV39H2 | 602.894928 | 0.4983396 | 0.15772687 | 3.15950989 | 0.00158035 |
| 57423 | ATP5J2 | 1255.18547 | -0.3288934 | 0.10409673 | -3.1594975 | 0.00158041 |
| 73102 | SLC22A23 | 1976.76861 | -1.0589036 | 0.33521144 | -3.1589124 | 0.00158359 |
| 56176 | PIGP | 132.937406 | -0.4842183 | 0.15330245 | -3.1585818 | 0.00158539 |
| 70550 | 5730416F02I | 41.0613853 | 0.75588103 | 0.23957372 | 3.15510835 | 0.00160439 |
| 17318 | MID1 | 137.771858 | -0.5758995 | 0.18261481 | -3.1536296 | 0.00161254 |
| 64074 | SMOC2 | 20.4514532 | -3.3691598 | 1.06847375 | -3.1532453 | 0.00161466 |
| 22612 | YES1 | 1725.14379 | -0.2153474 | 0.06830598 | -3.1526868 | 0.00161775 |
| 19224 | PTGS1 | 5875.02537 | 2.25869274 | 0.7164369 | 3.15267505 | 0.00161782 |
| 74107 | CEP55 | 2349.54715 | 0.28303434 | 0.08980631 | 3.15160865 | 0.00162374 |

| | | | | | | |
|---|---|---|---|---|---|---|
| 230676 | SZT2 | 934.756796 | 0.33208393 | 0.10538075 | 3.15127684 | 0.00162558 |
| 15469 | PRMT1 | 5078.902 | 0.54889999 | 0.17419773 | 3.15101682 | 0.00162703 |
| 230654 | LRRC41 | 1744.63703 | 0.4083865 | 0.12963664 | 3.15023981 | 0.00163136 |
| 66315 | SENP7 | 403.68605 | -0.5614451 | 0.17822681 | -3.150172 | 0.00163174 |
| 18479 | PAK1 | 1415.69577 | -0.9254125 | 0.29378013 | -3.1500174 | 0.00163261 |
| 12649 | CHEK1 | 847.613603 | 0.64728548 | 0.20562211 | 3.14793717 | 0.00164427 |
| 67285 | CWC27 | 649.159387 | 0.53405967 | 0.16968991 | 3.14726834 | 0.00164804 |
| 15525 | HSPA4 | 16207.2539 | 0.17287329 | 0.05493944 | 3.14661539 | 0.00165172 |
| 244310 | DLGAP2 | 4.86364501 | -3.5980886 | 1.14373372 | -3.1459146 | 0.00165568 |
| 546100 | Gm5914 | 14.6865451 | 1.54910593 | 0.49252757 | 3.14521668 | 0.00165964 |
| 209586 | NUDCD3 | 3913.93821 | -0.248994 | 0.07917562 | -3.144832 | 0.00166182 |
| 223665 | C030006K11I | 271.214492 | 0.43713589 | 0.13900468 | 3.14475656 | 0.00166225 |
| 94284 | Ugt1a6a | 114.770913 | 1.15445174 | 0.36712015 | 3.14461557 | 0.00166305 |
| 217370 | BC017643 | 983.260156 | -0.4071317 | 0.12948189 | -3.1443136 | 0.00166477 |
| 246707 | EMILIN2 | 2655.96287 | 3.34231616 | 1.06411404 | 3.14093793 | 0.00168408 |
| 73736 | FCF1 | 406.055246 | 0.28321381 | 0.09017528 | 3.14070322 | 0.00168543 |
| 232875 | ZSCAN18 | 5.2042872 | -3.6823228 | 1.17245799 | -3.1406864 | 0.00168552 |
| 229279 | HNRNPA3 | 675.818461 | -0.4060169 | 0.12929142 | -3.1403233 | 0.00168761 |
| 75547 | AKAP13 | 2177.71418 | 0.75116228 | 0.23923263 | 3.13988229 | 0.00169016 |
| 20462 | TRA2B | 1343.03595 | 0.33760592 | 0.10752593 | 3.1397628 | 0.00169085 |
| 76411 | IFT43 | 386.333692 | -0.4665134 | 0.14858524 | -3.1397019 | 0.0016912 |
| 224697 | ADAMTS10 | 199.939349 | 0.87359343 | 0.27831843 | 3.13882713 | 0.00169625 |
| 628900 | Serpina3i | 1024.83498 | -1.2126624 | 0.38658038 | -3.1368959 | 0.00170747 |
| 269224 | PASK | 746.044588 | -0.3798241 | 0.12108512 | -3.1368355 | 0.00170782 |
| 19647 | RBBP6 | 3571.52382 | 0.36772058 | 0.11723198 | 3.13669157 | 0.00170866 |
| 67382 | BRD3 | 2221.05465 | 0.19857507 | 0.06331203 | 3.13645102 | 0.00171006 |
| 110157 | RAF1 | 3237.4345 | 0.53316242 | 0.17000248 | 3.13620367 | 0.0017115 |
| 268512 | SLC26A11 | 168.584806 | -0.8415168 | 0.26833362 | -3.136084 | 0.0017122 |
| 12313 | CALM1 | 17139.6949 | 0.3067305 | 0.09780738 | 3.13606693 | 0.0017123 |
| 30930 | VPS26A | 5206.64813 | -0.5183264 | 0.16532083 | -3.135276 | 0.00171692 |
| 30939 | PTTG1 | 1014.36207 | -0.4018928 | 0.12818495 | -3.1352574 | 0.00171703 |
| 20238 | ATXN1 | 984.827937 | 0.82062673 | 0.26175551 | 3.13508861 | 0.00171802 |
| 229615 | PIAS3 | 745.644571 | -0.2395569 | 0.07642535 | -3.1345219 | 0.00172134 |
| 18212 | NTRK2 | 15.0505904 | -7.3410164 | 2.34238747 | -3.1339889 | 0.00172447 |
| 22644 | RNF103 | 1178.79345 | 0.21862452 | 0.06977994 | 3.133057 | 0.00172996 |
| 66164 | NIP7 | 1566.28514 | 0.31333641 | 0.10001627 | 3.1328544 | 0.00173115 |
| 16664 | KRT14 | 3291.99306 | -4.185132 | 1.33593546 | -3.1327352 | 0.00173186 |
| 225888 | KMT5B | 1345.3803 | -0.3615974 | 0.11542994 | -3.1326135 | 0.00173257 |
| 69656 | PIR | 72.9405487 | -7.510492 | 2.3977933 | -3.1322517 | 0.00173471 |
| 53621 | CNOT4 | 1182.92782 | 0.25126368 | 0.08021951 | 3.13220152 | 0.00173501 |
| 69528 | 1700030J22F | 32.8876765 | -1.1697367 | 0.37354006 | -3.1314892 | 0.00173922 |
| 233064 | WDR62 | 1504.28242 | 0.59846535 | 0.19115695 | 3.13075386 | 0.00174358 |
| 209456 | Trp53bp2 | 1109.50481 | -0.306376 | 0.09786299 | -3.1306631 | 0.00174412 |
| 14911 | THUMPD3 | 923.55536 | -0.3016444 | 0.09635821 | -3.1304488 | 0.00174539 |
| 78303 | HIST3H2BA | 9.26659062 | -5.1796173 | 1.65459694 | -3.1304405 | 0.00174544 |

| | | | | | | |
|---|---|---|---|---|---|---|
| 18045 | NFYB | 975.094918 | -0.1992284 | 0.06364632 | -3.130243 | 0.00174662 |
| 23991 | CIB1 | 678.284991 | -0.3932399 | 0.12566114 | -3.1293672 | 0.00175183 |
| 320159 | FAM179A | 7.50730933 | -1.8435756 | 0.58918109 | -3.1290475 | 0.00175374 |
| 381308 | Ifi211 | 79.9663223 | -0.8586768 | 0.27444394 | -3.1287874 | 0.00175529 |
| 93885 | PCDHB14 | 46.1996827 | 1.09265149 | 0.34935946 | 3.12758521 | 0.00176249 |
| 14961 | H2-Ab1 | 5.8925987 | -3.850355 | 1.23134572 | -3.1269488 | 0.00176631 |
| 11792 | APEX1 | 2866.76102 | 0.82807277 | 0.26484492 | 3.12663262 | 0.00176821 |
| 13047 | CUX1 | 2659.29833 | -0.2150778 | 0.06878974 | -3.1265976 | 0.00176842 |
| 320940 | ATP11C | 1272.28335 | 0.47991777 | 0.15355061 | 3.12546961 | 0.00177521 |
| 19205 | PTBP1 | 13365.7448 | 0.36254706 | 0.11604666 | 3.12414908 | 0.0017832 |
| 70495 | ATP6AP2 | 4988.15748 | -0.682497 | 0.21847042 | -3.1239792 | 0.00178423 |
| 12301 | CACYBP | 3852.56559 | 0.39334244 | 0.12592067 | 3.12373211 | 0.00178573 |
| 231821 | ADAP1 | 18.7659662 | -2.2019871 | 0.70509153 | -3.1229805 | 0.0017903 |
| 71751 | MAP3K13 | 38.3463165 | -1.1929365 | 0.38210888 | -3.1219805 | 0.00179639 |
| 56874 | RNF32 | 9.18183647 | -2.16451 | 0.69385905 | -3.1195241 | 0.00181143 |
| 106763 | TTBK1 | 2.92968969 | -4.213304 | 1.35095544 | -3.1187586 | 0.00181615 |
| 20495 | SLC12A1 | 9.15775161 | 2.34325558 | 0.75139141 | 3.1185552 | 0.0018174 |
| 66294 | FAM3A | 409.746529 | -0.4510706 | 0.14476602 | -3.1158596 | 0.0018341 |
| 218973 | WDHD1 | 1789.57753 | 0.76393924 | 0.24520275 | 3.11554104 | 0.00183608 |
| 14619 | GJB2 | 2.66973512 | -4.8379027 | 1.55352889 | -3.1141376 | 0.00184483 |
| 114741 | Supt16 | 6505.91621 | 0.76524977 | 0.24575511 | 3.11387126 | 0.0018465 |
| 436022 | DNAAF3 | 42.1962048 | -0.9975475 | 0.32047066 | -3.1127576 | 0.00185348 |
| 57258 | XPO4 | 1287.79656 | 0.76330726 | 0.24524038 | 3.11248603 | 0.00185519 |
| 232599 | Gm4876 | 3.38270529 | -4.4057227 | 1.41557639 | -3.1123172 | 0.00185625 |
| 24056 | SH3BP5 | 403.492952 | -1.0379132 | 0.33363824 | -3.1108938 | 0.00186522 |
| 226432 | IPO9 | 8805.28278 | 0.20223673 | 0.06502277 | 3.11024485 | 0.00186932 |
| 331623 | BEND3 | 1428.52106 | 0.48334205 | 0.15541398 | 3.11002934 | 0.00187069 |
| 73132 | SLC25A16 | 783.926859 | -0.2685602 | 0.08635603 | -3.1099182 | 0.00187139 |
| 53608 | MAP3K6 | 370.304176 | 0.30961061 | 0.09959355 | 3.10874165 | 0.00187886 |
| 235134 | NFRKB | 900.220062 | 0.24883471 | 0.08006371 | 3.10795897 | 0.00188384 |
| 17217 | MCM4 | 6605.65977 | 0.18151268 | 0.05841777 | 3.10714845 | 0.00188902 |
| 18575 | PDE1C | 10.1279553 | 1.66673458 | 0.53642215 | 3.10713227 | 0.00188912 |
| 50905 | IL17RB | 7.05119164 | -3.3518448 | 1.07877264 | -3.1070911 | 0.00188938 |
| 13603 | OPN3 | 111.847434 | 0.71100195 | 0.22885486 | 3.10678108 | 0.00189136 |
| 77254 | YIF1B | 850.697134 | 0.41705105 | 0.13428553 | 3.1057036 | 0.00189827 |
| 20719 | Serpinb6a | 9235.86955 | -0.3130132 | 0.10080195 | -3.1052292 | 0.00190132 |
| 14408 | GABRR1 | 4.89587485 | -5.7180293 | 1.84186431 | -3.1044791 | 0.00190614 |
| 215900 | FAM26F | 4.75863732 | 3.56747723 | 1.14934471 | 3.10392277 | 0.00190973 |
| 77832 | TCHP | 348.591852 | -0.3193581 | 0.10290261 | -3.1034985 | 0.00191247 |
| 13200 | DDOST | 5680.24017 | 0.47041321 | 0.15158951 | 3.10320434 | 0.00191437 |
| 16594 | KLC2 | 1126.38368 | 0.40495799 | 0.13050437 | 3.10302239 | 0.00191555 |
| 23993 | KLK7 | 4.05159886 | -3.922601 | 1.26430814 | -3.1025672 | 0.0019185 |
| 66526 | TCEANC2 | 183.602636 | 0.55649734 | 0.17945861 | 3.10097881 | 0.00192882 |
| 103583 | FBXW11 | 4088.20981 | 0.10492425 | 0.03383602 | 3.1009634 | 0.00192892 |
| 14349 | Fv1 | 122.954356 | 0.56080711 | 0.18085712 | 3.10082951 | 0.00192979 |

| | | | | | | |
|---|---|---|---|---|---|---|
| 80744 | CWC22 | 632.457161 | 0.36580955 | 0.1179878 | 3.1004015 | 0.00193258 |
| 13531 | Usp17la | 37.4653293 | -2.6037008 | 0.83993382 | -3.099888 | 0.00193594 |
| 21673 | DNTT | 10.856458 | -6.1330352 | 1.97855936 | -3.0997479 | 0.00193685 |
| 320273 | B230208H11 | 41.762759 | -1.0226769 | 0.32996046 | -3.0993922 | 0.00193918 |
| 72179 | FBXL2 | 213.878075 | 0.80210452 | 0.25891587 | 3.09793495 | 0.00194874 |
| 66515 | CUL7 | 2005.12275 | 1.39416403 | 0.45014255 | 3.09716115 | 0.00195384 |
| 108655 | FOXP1 | 4143.70309 | 0.60458158 | 0.19526993 | 3.09613259 | 0.00196063 |
| 52551 | SGTA | 3317.56874 | -0.194489 | 0.06281746 | -3.0960976 | 0.00196086 |
| 24084 | TEKT2 | 10.0352664 | -1.87545 | 0.60576023 | -3.0960269 | 0.00196133 |
| 209707 | LCORL | 538.991541 | -0.6023275 | 0.19457084 | -3.0956721 | 0.00196367 |
| 15931 | IDS | 1738.83165 | -0.513856 | 0.16601052 | -3.0953221 | 0.00196599 |
| 67773 | kat8 | 362.74197 | 0.74276916 | 0.24006928 | 3.09397842 | 0.00197492 |
| 18430 | oxtr | 7.46962396 | -2.3236965 | 0.75136689 | -3.0926256 | 0.00198394 |
| 210762 | PPP1R36 | 5.37639872 | -3.6651874 | 1.18527982 | -3.092255 | 0.00198642 |
| 18107 | NMT1 | 3958.32367 | -0.3239411 | 0.10477362 | -3.0918192 | 0.00198934 |
| 272322 | ARNTL2 | 62.536724 | -4.0530066 | 1.31130869 | -3.0908104 | 0.00199611 |
| 225028 | MAP4K3 | 1548.73087 | -0.5454106 | 0.17647509 | -3.0905813 | 0.00199765 |
| 68525 | EVC2 | 1514.60103 | 0.68626204 | 0.22210037 | 3.08987343 | 0.00200242 |
| 20437 | Siah1a | 428.765115 | -0.3892338 | 0.12599996 | -3.0891579 | 0.00200725 |
| 107746 | RAPGEF1 | 6722.51684 | 0.23669627 | 0.07664622 | 3.0881662 | 0.00201396 |
| 16164 | IL13RA1 | 854.667293 | -0.2196115 | 0.07112685 | -3.0876033 | 0.00201778 |
| 66588 | CMPK1 | 1977.37064 | 0.85893096 | 0.27824776 | 3.08692856 | 0.00202236 |
| 107376 | E330013P04I | 4.43641718 | -5.582325 | 1.80849005 | -3.0867325 | 0.0020237 |
| 101197 | Zfp956 | 160.543816 | -0.4599428 | 0.14900652 | -3.0867297 | 0.00202372 |
| 110350 | DYNC2H1 | 640.495067 | 0.4443313 | 0.14395637 | 3.08656919 | 0.00202481 |
| 30853 | MLF2 | 6888.94429 | 0.24176072 | 0.07836072 | 3.08522827 | 0.00203396 |
| 58988 | RPS6KB2 | 392.257082 | -0.3893817 | 0.12625613 | -3.0840615 | 0.00204195 |
| 104349 | Zfp119a | 105.187094 | -0.4699747 | 0.15240263 | -3.0837704 | 0.00204395 |
| 67389 | FAM132A | 78.3920199 | -0.7471055 | 0.24250825 | -3.0807425 | 0.00206485 |
| 69440 | DENND6B | 168.078895 | -0.3949973 | 0.1282487 | -3.0799322 | 0.00207048 |
| 52013 | R3HCC1L | 603.876161 | -0.4274205 | 0.13878608 | -3.0797076 | 0.00207204 |
| 231915 | USPL1 | 1135.88044 | 0.29120436 | 0.0945823 | 3.07884625 | 0.00207804 |
| 69820 | 1810059H22 | 33.8717819 | 1.07136541 | 0.34802255 | 3.07843671 | 0.0020809 |
| 211482 | EFHB | 4.56936538 | -3.0111207 | 0.97837159 | -3.0776862 | 0.00208614 |
| 229613 | 6330549D23 | 142.295401 | 0.43851435 | 0.14250858 | 3.07710835 | 0.00209019 |
| 100041286 | Snhg15 | 388.320337 | -0.3618054 | 0.11761804 | -3.0761043 | 0.00209724 |
| 74521 | PPP4R4 | 24.3974269 | 1.03335103 | 0.33593964 | 3.07600207 | 0.00209796 |
| 11600 | ANGPT1 | 657.224693 | 4.06331426 | 1.32103808 | 3.07584946 | 0.00209904 |
| 228858 | GDAP1L1 | 5.91625537 | -3.0962585 | 1.00681551 | -3.0752987 | 0.00210292 |
| 76498 | PAQR4 | 297.194319 | -0.5928357 | 0.19278362 | -3.0751354 | 0.00210407 |
| 70354 | SECISBP2L | 4948.73876 | -0.6097896 | 0.19829944 | -3.0750947 | 0.00210436 |
| 15288 | HMBS | 785.321282 | 0.2558647 | 0.08324639 | 3.0735833 | 0.00211505 |
| 72151 | RFC5 | 1409.3692 | 0.35532891 | 0.1156134 | 3.07342325 | 0.00211618 |
| 64453 | Zfp280b | 850.17023 | 0.27548933 | 0.08964086 | 3.07325611 | 0.00211737 |
| 243842 | GLTSCR1 | 587.009901 | 0.4753935 | 0.15471729 | 3.07265914 | 0.00212161 |

| | | | | | | |
|---|---|---|---|---|---|---|
| 74318 | HOPX | 15.6976927 | 1.7291894 | 0.56290348 | 3.07191103 | 0.00212693 |
| 320019 | Ptgs2os | 74.4423771 | 1.45972076 | 0.47535133 | 3.07082502 | 0.00213468 |
| 66610 | ABI3 | 4.16950483 | -3.9338553 | 1.28105895 | -3.070784 | 0.00213498 |
| 12913 | CREB3 | 1203.22223 | -0.5893281 | 0.19191833 | -3.0707236 | 0.00213541 |
| 433940 | FAM222A | 19.436393 | -2.1512165 | 0.70069774 | -3.0701062 | 0.00213983 |
| 56524 | MPP6 | 2899.77631 | 0.58817077 | 0.19162472 | 3.06938879 | 0.00214497 |
| 218333 | ICE1 | 2264.42509 | 0.35178936 | 0.11462306 | 3.06909759 | 0.00214706 |
| 107173 | GPR137 | 436.997299 | 0.36645173 | 0.11946525 | 3.06743373 | 0.00215905 |
| 18777 | LYPLA1 | 2291.93879 | -0.4110803 | 0.1340673 | -3.0662238 | 0.00216781 |
| 107448 | UNC5A | 3.83562263 | 3.87714689 | 1.26464197 | 3.06580597 | 0.00217084 |
| 100306945 | Snora73b | 126.310011 | 0.90099767 | 0.29396259 | 3.06500792 | 0.00217664 |
| 80883 | NTNG1 | 6.30391339 | 4.02123322 | 1.31254893 | 3.06368254 | 0.00218631 |
| 216154 | MED16 | 1335.93579 | 0.21310447 | 0.06955841 | 3.06367636 | 0.00218635 |
| 380664 | lemd3 | 1083.88423 | -0.3313592 | 0.1082003 | -3.062461 | 0.00219525 |
| 70885 | INTS10 | 1113.28864 | 0.2084884 | 0.06810689 | 3.06119397 | 0.00220456 |
| 100038882 | ISG15 | 155.887451 | -1.0062272 | 0.32877458 | -3.0605384 | 0.00220939 |
| 69870 | POLR3GL | 99.2048068 | -0.5805489 | 0.18975235 | -3.0595083 | 0.00221701 |
| 53330 | VAMP4 | 326.31627 | 0.28703984 | 0.09382629 | 3.0592686 | 0.00221878 |
| 68240 | RPA3 | 271.218016 | 0.40367691 | 0.13198371 | 3.0585359 | 0.00222421 |
| 50496 | E2F6 | 1502.42288 | 0.63347422 | 0.20713842 | 3.05821691 | 0.00222658 |
| 103724 | TBC1D10A | 845.201059 | -0.3240164 | 0.1059569 | -3.058002 | 0.00222818 |
| 70701 | NIPAL1 | 1707.5651 | -0.6497881 | 0.21250322 | -3.0577798 | 0.00222983 |
| 384061 | FNDC5 | 5.8051315 | -3.0214159 | 0.98814764 | -3.0576563 | 0.00223075 |
| 67049 | PUS3 | 307.464142 | 0.6416652 | 0.20988251 | 3.05725906 | 0.00223371 |
| 66427 | CYB5B | 8807.26359 | 0.25228472 | 0.08252484 | 3.05707615 | 0.00223507 |
| 100503041 | PDZD7 | 8.26840458 | -1.7900397 | 0.58559746 | -3.056775 | 0.00223732 |
| 68194 | NDUFB4 | 83.18356 | -0.522704 | 0.17101494 | -3.0564813 | 0.00223951 |
| 66880 | RSRC1 | 1151.84217 | -0.2105366 | 0.06889653 | -3.0558375 | 0.00224433 |
| 218952 | FERMT2 | 2651.66029 | 0.53091686 | 0.17377346 | 3.05522411 | 0.00224892 |
| 108797 | MEX3B | 183.776411 | 0.97947577 | 0.32059326 | 3.05519766 | 0.00224912 |
| 71764 | C2CD2L | 834.369134 | 0.5680362 | 0.18592983 | 3.05511062 | 0.00224978 |
| 17346 | MKNK1 | 494.942302 | 0.45831845 | 0.15002537 | 3.05493958 | 0.00225106 |
| 103844 | INCA1 | 24.8266162 | -1.7449952 | 0.57131962 | -3.0543239 | 0.00225568 |
| 108767 | PNRC1 | 1035.36253 | -1.1859912 | 0.38842719 | -3.0533167 | 0.00226327 |
| 218639 | ARL15 | 517.349161 | 0.69831624 | 0.22871187 | 3.0532575 | 0.00226372 |
| 230936 | PHF13 | 905.95358 | 0.55036295 | 0.18026161 | 3.05313447 | 0.00226464 |
| 622127 | Cyp3a57 | 11.2701274 | -6.9233172 | 2.26820349 | -3.0523352 | 0.00227068 |
| 241447 | CERS6 | 1213.86599 | 0.66375072 | 0.21752966 | 3.0513113 | 0.00227844 |
| 13394 | DLX4 | 24.7083715 | -2.4770757 | 0.81210887 | -3.0501768 | 0.00228707 |
| 320473 | HEATR5B | 1540.92995 | 0.2867889 | 0.09402941 | 3.04999156 | 0.00228848 |
| 100043726 | Gm4610 | 107.064525 | -3.0868691 | 1.01218517 | -3.0497079 | 0.00229064 |
| 208638 | SLC25A38 | 308.102603 | 0.44173794 | 0.14484687 | 3.0496893 | 0.00229078 |
| 353310 | Zfp703 | 2116.75083 | 0.52840748 | 0.17329014 | 3.04926448 | 0.00229402 |
| 215160 | rhbdd2 | 667.023211 | -0.4600277 | 0.15092711 | -3.0480125 | 0.0023036 |
| 434218 | Trim34b | 10.1817912 | -2.0804501 | 0.68307653 | -3.0457057 | 0.00232135 |

| | | | | | | |
|---|---|---|---|---|---|---|
| 105988 | ESPL1 | 2271.67056 | -0.3353407 | 0.11011536 | -3.0453577 | 0.00232404 |
| 66069 | SNUPN | 215.189144 | 0.36733921 | 0.12064112 | 3.04489233 | 0.00232764 |
| 12943 | PCDHA10 | 4.05091761 | -5.4521605 | 1.79126227 | -3.0437534 | 0.00233647 |
| 320206 | A730028G07 | 82.480041 | -0.6372959 | 0.20939104 | -3.0435682 | 0.0023379 |
| 100683 | TRRAP | 5842.29695 | 0.29519392 | 0.09700612 | 3.04304412 | 0.00234198 |
| 70252 | Jpx | 27.7244325 | -1.1219383 | 0.36873437 | -3.0426735 | 0.00234487 |
| 27632 | NELFE | 1214.7136 | -0.2089868 | 0.0686903 | -3.0424508 | 0.0023466 |
| 224640 | LEMD2 | 1498.38936 | 0.60166187 | 0.19777502 | 3.04215291 | 0.00234893 |
| 210503 | Zfp677 | 143.753177 | -0.512243 | 0.16838237 | -3.0421414 | 0.00234902 |
| 20200 | S100A6 | 29006.2657 | -0.5537582 | 0.18209925 | -3.0409694 | 0.00235818 |
| 69806 | SLC39A11 | 939.633033 | -0.3316588 | 0.10913439 | -3.0389942 | 0.00237369 |
| 22763 | ZFR | 5166.31181 | 0.25682572 | 0.08451092 | 3.03896476 | 0.00237393 |
| 93889 | PCDHB18 | 52.0475827 | 1.21053442 | 0.39840773 | 3.03843105 | 0.00237814 |
| 381695 | N4BP2L2 | 1729.64422 | -0.2474025 | 0.0814317 | -3.0381591 | 0.00238028 |
| 237615 | ANKRD52 | 5205.25835 | -0.1854884 | 0.06107674 | -3.0369723 | 0.00238967 |
| 229709 | AHCYL1 | 8158.37288 | 0.42825183 | 0.14108998 | 3.03531002 | 0.00240289 |
| 378702 | SERF2 | 666.998401 | -0.3967926 | 0.13083409 | -3.032792 | 0.00242303 |
| 22036 | TRAIP | 500.833284 | 0.33949511 | 0.11194269 | 3.03275823 | 0.0024233 |
| 30951 | CBX8 | 182.852233 | -0.4379628 | 0.14462163 | -3.0283355 | 0.00245905 |
| 100040297 | Gm2695 | 745.91837 | 0.2170115 | 0.07167218 | 3.0278347 | 0.00246313 |
| 218506 | MRPS27 | 723.808596 | 0.43193042 | 0.14271719 | 3.02647784 | 0.00247421 |
| 338350 | Acad12 | 17.810192 | -1.3268873 | 0.43854155 | -3.0256821 | 0.00248073 |
| 74978 | LRRIQ1 | 6.38925422 | -4.0487984 | 1.33935864 | -3.0229382 | 0.00250333 |
| 170711 | OTUD7A | 2.9320011 | -4.2041 | 1.39160971 | -3.0210338 | 0.00251913 |
| 73061 | Cldn34c1 | 3.69587354 | -5.3114795 | 1.7582253 | -3.0209322 | 0.00251998 |
| 56452 | ORC6 | 1444.09305 | 0.31098224 | 0.10295583 | 3.02054052 | 0.00252324 |
| 243906 | ZFP14 | 27.7088778 | -1.204072 | 0.39864133 | -3.0204396 | 0.00252408 |
| 14287 | FPGS | 1007.68462 | 0.26272044 | 0.08699138 | 3.0200743 | 0.00252713 |
| 270035 | Ietm2 | 24.807081 | -1.1972882 | 0.39645101 | -3.0200154 | 0.00252762 |
| 545389 | CEP170 | 5236.77469 | 0.22239649 | 0.07365346 | 3.01949827 | 0.00253194 |
| 270624 | SPIN4 | 178.189011 | 0.42157572 | 0.13963946 | 3.01902996 | 0.00253585 |
| 20318 | SDF4 | 2401.62119 | 0.21983952 | 0.07283185 | 3.018453 | 0.00254069 |
| 26407 | MAP3K4 | 1197.05531 | 0.70948872 | 0.23507261 | 3.01816838 | 0.00254308 |
| 65964 | ZAK | 4683.27882 | -0.3089397 | 0.10236496 | -3.0180216 | 0.00254431 |
| 12704 | CIT | 1882.52211 | 0.3393244 | 0.112443 | 3.01774593 | 0.00254662 |
| 67705 | 1810058I24R | 984.390714 | 0.37472221 | 0.12422097 | 3.01657778 | 0.00255646 |
| 16451 | JAK1 | 4644.42359 | 0.57346295 | 0.19022541 | 3.01464955 | 0.00257276 |
| 235435 | LCTL | 4.49911067 | -2.9905511 | 0.99240024 | -3.0134526 | 0.00258293 |
| 100504278 | 2810402E24I | 53.7045926 | -0.7714112 | 0.25599146 | -3.0134257 | 0.00258316 |
| 208084 | PIF1 | 587.74719 | 0.40695913 | 0.13505073 | 3.01337975 | 0.00258355 |
| 22371 | VWF | 92.1579714 | -2.9108036 | 0.9662145 | -3.0125853 | 0.00259033 |
| 654795 | SDR39U1 | 634.538585 | -0.7926116 | 0.26312383 | -3.0123141 | 0.00259264 |
| 242466 | Zfp462 | 2684.7723 | -0.4060243 | 0.13480673 | -3.0118994 | 0.00259619 |
| 67220 | PLEKHO1 | 833.874825 | 0.79731911 | 0.26477227 | 3.01133913 | 0.00260098 |
| 24131 | LDB3 | 43.2040131 | 2.0538834 | 0.68226158 | 3.01040461 | 0.002609 |

| | | | | | | |
|---|---|---|---|---|---|---|
| 100689 | SPON2 | 10.3136269 | 2.15595322 | 0.7161716 | 3.01038636 | 0.00260916 |
| 12558 | CDH2 | 5410.58359 | -1.2495145 | 0.41506904 | -3.0103774 | 0.00260923 |
| 67707 | MRPL24 | 863.054282 | -0.3252748 | 0.10809194 | -3.0092421 | 0.002619 |
| 14085 | FAH | 366.206052 | -1.2370845 | 0.41110973 | -3.0091347 | 0.00261993 |
| 627626 | PTCHD4 | 102.700649 | -3.4756453 | 1.15505613 | -3.0090705 | 0.00262048 |
| 16651 | SSPN | 235.430692 | -1.6538108 | 0.54964308 | -3.0088813 | 0.00262211 |
| 18637 | PFDN2 | 1053.60039 | -0.3252949 | 0.10814561 | -3.0079348 | 0.0026303 |
| 76238 | GRHPR | 472.812251 | -0.7595575 | 0.25251934 | -3.007918 | 0.00263044 |
| 94065 | MRPL34 | 585.570457 | 0.33464842 | 0.11128293 | 3.0071855 | 0.00263679 |
| 66845 | MRPL33 | 1292.49559 | -0.2608246 | 0.08674779 | -3.0067006 | 0.002641 |
| 14004 | CHCHD2 | 961.880891 | -0.2125551 | 0.07069439 | -3.0066755 | 0.00264122 |
| 110385 | PDE4C | 2.96040626 | -5.0041582 | 1.66445071 | -3.0064923 | 0.00264281 |
| 68735 | MRPS18C | 517.802244 | -0.3069094 | 0.10209669 | -3.006066 | 0.00264652 |
| 54153 | rasa4 | 921.488727 | -0.7207469 | 0.23977554 | -3.0059232 | 0.00264776 |
| 56280 | MRPL37 | 1194.94663 | 0.53024533 | 0.17643407 | 3.00534551 | 0.00265279 |
| 66101 | PPIH | 133.360442 | 0.67039914 | 0.22309261 | 3.00502618 | 0.00265558 |
| 18205 | NTF3 | 4.06937832 | 4.02429401 | 1.33923551 | 3.00491884 | 0.00265652 |
| 73689 | BLOC1S2 | 192.11038 | -0.3624414 | 0.12062809 | -3.0046187 | 0.00265914 |
| 210135 | Zfp180 | 976.092047 | 0.61568254 | 0.20491966 | 3.00450685 | 0.00266012 |
| 54384 | MTMR7 | 67.3297764 | -2.8179976 | 0.93824173 | -3.0034878 | 0.00266904 |
| 73078 | PMPCB | 1677.60522 | -0.2367225 | 0.07885578 | -3.0019673 | 0.00268241 |
| 230075 | NDUFB6 | 918.168806 | -0.4143691 | 0.13803779 | -3.0018528 | 0.00268342 |
| 28077 | MED10 | 929.207768 | 0.31097017 | 0.10360109 | 3.00161083 | 0.00268555 |
| 74525 | FAM234B | 530.02686 | 0.28277117 | 0.09421708 | 3.00127299 | 0.00268853 |
| 72289 | MALAT1 | 23752.5952 | 0.45343696 | 0.15112712 | 3.00036787 | 0.00269654 |
| 16815 | LBX2 | 5.01996271 | -3.6091746 | 1.20298708 | -3.0001774 | 0.00269822 |
| 13169 | DBNL | 3232.69404 | -0.112059 | 0.03736271 | -2.9992193 | 0.00270672 |
| 218311 | Zfp455 | 22.4976731 | -1.5661081 | 0.52231785 | -2.9983813 | 0.00271418 |
| 64833 | Acot10 | 484.385871 | -0.2539201 | 0.08468587 | -2.9983761 | 0.00271422 |
| 18438 | P2RX4 | 875.725473 | -0.7794364 | 0.25995759 | -2.9983213 | 0.00271471 |
| 140630 | UBE4A | 2281.69656 | -0.1404849 | 0.04686157 | -2.9978708 | 0.00271873 |
| 327766 | TMEM26 | 15.65858 | -4.0600133 | 1.35454494 | -2.9973264 | 0.00272359 |
| 269881 | MAP3K10 | 695.482479 | 0.53926627 | 0.18007277 | 2.99471295 | 0.00274703 |
| 68953 | CHMP2A | 1204.54131 | -0.8209397 | 0.27415642 | -2.9944208 | 0.00274966 |
| 241230 | ST8SIA6 | 4.84750506 | -2.7521502 | 0.91922335 | -2.9939951 | 0.0027535 |
| 11636 | AK1 | 2634.99115 | -0.5810065 | 0.19407899 | -2.9936598 | 0.00275653 |
| 70383 | COX10 | 866.548168 | 0.2449855 | 0.08184229 | 2.99338528 | 0.00275901 |
| 76303 | OSBP | 2382.43957 | 0.16893385 | 0.05644722 | 2.99277546 | 0.00276453 |
| 74198 | DTX2 | 714.425733 | 0.55031091 | 0.1838918 | 2.99257989 | 0.0027663 |
| 50791 | MAGI2 | 121.493686 | 1.50331811 | 0.50266512 | 2.99069512 | 0.00278343 |
| 69923 | AGK | 93.4583008 | 0.48822605 | 0.1632899 | 2.98993414 | 0.00279038 |
| 233789 | SMG1 | 7322.94893 | 0.48258532 | 0.1614127 | 2.98976046 | 0.00279196 |
| 239510 | PHF20L1 | 3119.6322 | 0.18960485 | 0.06343585 | 2.98892294 | 0.00279963 |
| 68982 | 1500015A07 | 62.3117789 | -0.8072553 | 0.27017251 | -2.9879255 | 0.00280878 |
| 71911 | BDH1 | 45.2373574 | -2.8269633 | 0.94634639 | -2.9872395 | 0.00281509 |

| | | | | | | |
|---|---|---|---|---|---|---|
| 67966 | ZCCHC10 | 391.961101 | 0.53539383 | 0.17922925 | 2.98720129 | 0.00281544 |
| 101497 | PLEKHG2 | 1330.3375 | 0.62970533 | 0.21080891 | 2.98709069 | 0.00281646 |
| 276846 | PIGS | 1698.91353 | -0.490406 | 0.16418303 | -2.9869468 | 0.00281779 |
| 665137 | Gm7511 | 118.903887 | 0.50970635 | 0.17065352 | 2.9867907 | 0.00281923 |
| 15024 | H2-T10 | 500.962383 | -0.5978982 | 0.20019361 | -2.9865998 | 0.00282099 |
| 212111 | INPP5A | 984.011617 | -0.3109845 | 0.10413005 | -2.9865013 | 0.0028219 |
| 16510 | KCNH1 | 15.3234813 | -2.5388658 | 0.85098733 | -2.9834356 | 0.00285032 |
| 107568 | WWP1 | 1312.18788 | -0.7541396 | 0.25286787 | -2.9823464 | 0.00286048 |
| 12607 | CEBPZ | 2688.95738 | 0.29022157 | 0.09731641 | 2.98224684 | 0.00286141 |
| 13669 | EIF3A | 19640.8387 | 0.13650652 | 0.04577315 | 2.98224028 | 0.00286147 |
| 210529 | METTL14 | 823.309568 | -0.2955847 | 0.09911543 | -2.9822265 | 0.0028616 |
| 68626 | ELAC2 | 1377.71903 | 0.39111278 | 0.13115081 | 2.9821605 | 0.00286222 |
| 67149 | NKAIN1 | 907.279804 | -0.3843145 | 0.12890517 | -2.9813737 | 0.00286958 |
| 620419 | Zfp963 | 78.630697 | -0.6337094 | 0.21259429 | -2.9808393 | 0.0028746 |
| 72392 | tmem175 | 306.672268 | 0.43082018 | 0.14461272 | 2.97913061 | 0.00289068 |
| 100861953 | #N/A | 11.2379109 | -6.1826717 | 2.07574541 | -2.9785308 | 0.00289634 |
| 76959 | CHMP5 | 1618.6338 | -0.50959 | 0.17110981 | -2.9781461 | 0.00289998 |
| 74125 | ARMC8 | 1096.06905 | 0.38046687 | 0.12775505 | 2.97809654 | 0.00290045 |
| 17722 | ND6 | 3725.94804 | -0.3798766 | 0.12758559 | -2.9774254 | 0.0029068 |
| 56292 | NAA10 | 696.086512 | 0.4682999 | 0.15728733 | 2.97735301 | 0.00290749 |
| 53333 | TOMM40 | 3761.80177 | 0.46099501 | 0.15487839 | 2.97649657 | 0.00291562 |
| 70911 | PHYHIPL | 3.37523696 | -4.4084969 | 1.48159358 | -2.9755103 | 0.00292502 |
| 21422 | tfcp2 | 322.190281 | -0.4632004 | 0.15575391 | -2.9739247 | 0.00294017 |
| 387314 | TMTC1 | 5.26719862 | 3.27267357 | 1.10076815 | 2.97308164 | 0.00294826 |
| 217232 | CDC27 | 4836.08104 | 0.13041959 | 0.04386698 | 2.97306952 | 0.00294838 |
| 214292 | Syna | 4.76196496 | 3.14536878 | 1.05877274 | 2.97076857 | 0.00297056 |
| 208194 | EXOG | 400.240117 | 0.36019432 | 0.12128328 | 2.9698596 | 0.00297936 |
| 68889 | UBAC2 | 761.751916 | -0.795456 | 0.26786609 | -2.9696033 | 0.00298185 |
| 408062 | Zfp873 | 128.079365 | -0.4908355 | 0.16529351 | -2.9694781 | 0.00298306 |
| 20392 | SGCE | 166.167995 | 1.25968861 | 0.42438509 | 2.96826786 | 0.00299483 |
| 13681 | EIF4A1 | 2324.52973 | 0.15267477 | 0.05146272 | 2.96670628 | 0.00301008 |
| 230577 | PARS2 | 191.917625 | 0.45399874 | 0.15312261 | 2.96493602 | 0.00302746 |
| 434179 | Zfp975 | 38.5614699 | -1.009601 | 0.34052547 | -2.9648325 | 0.00302848 |
| 20598 | SMPD2 | 330.098389 | -0.463623 | 0.15638943 | -2.9645418 | 0.00303134 |
| 637776 | Zfp977 | 2.84007788 | -4.9396442 | 1.66625278 | -2.9645227 | 0.00303153 |
| 19699 | RELN | 3.98077695 | -3.9119934 | 1.31996551 | -2.9637088 | 0.00303956 |
| 238393 | Serpina3f | 147.279127 | -1.1206551 | 0.37817449 | -2.9633282 | 0.00304332 |
| 230757 | 5730409E04I | 591.662156 | 0.48997086 | 0.16534694 | 2.96328954 | 0.0030437 |
| 110052 | DEK | 5827.78655 | 0.54129757 | 0.18280948 | 2.96099285 | 0.00306649 |
| 66314 | TPD52L2 | 1732.12261 | 0.24434456 | 0.0825275 | 2.96076532 | 0.00306876 |
| 12557 | CDH17 | 49.4593737 | -1.9287123 | 0.6514457 | -2.9606648 | 0.00306976 |
| 77891 | UBE2S | 194.548919 | 0.73841171 | 0.2494256 | 2.96044874 | 0.00307191 |
| 26404 | MAP3K12 | 367.708753 | -1.0796645 | 0.36478088 | -2.9597617 | 0.00307877 |
| 100503842 | 2410022M11 | 36.9727705 | 1.19139127 | 0.40255774 | 2.95955373 | 0.00308085 |
| 100503631 | Gm19805 | 5.46599713 | 2.76172151 | 0.93325017 | 2.959251 | 0.00308388 |

| | | | | | | |
|---|---|---|---|---|---|---|
| 544963 | IQGAP2 | 32.1492095 | -2.6222511 | 0.88618105 | -2.9590466 | 0.00308592 |
| 76846 | RPS9 | 6538.49351 | 0.35578151 | 0.12023804 | 2.95897624 | 0.00308663 |
| 17721 | ND5 | 49298.0045 | -0.6190876 | 0.20922535 | -2.958951 | 0.00308688 |
| 214444 | CDK5RAP2 | 1900.51262 | 0.31545389 | 0.10661762 | 2.95874052 | 0.00308899 |
| 15945 | CXCL10 | 70.57165 | 1.02887168 | 0.34776052 | 2.95856382 | 0.00309076 |
| 16825 | LDB1 | 2329.47097 | -0.1324302 | 0.0447689 | -2.9580835 | 0.00309558 |
| 66455 | CNPY4 | 817.466086 | 0.35835121 | 0.12114705 | 2.9579853 | 0.00309657 |
| 381218 | 4430402I18R | 5.63794826 | -2.400296 | 0.81161308 | -2.9574388 | 0.00310206 |
| 399626 | C330024C12I | 3.20449623 | -4.317472 | 1.45996203 | -2.9572495 | 0.00310397 |
| 71902 | CAND1 | 8224.24298 | 0.14150769 | 0.04786118 | 2.95662759 | 0.00311024 |
| 16904 | GZMM | 27.3789997 | -0.886097 | 0.29970992 | -2.9565155 | 0.00311137 |
| 244281 | MYO16 | 4.46904668 | -3.4289472 | 1.15994397 | -2.9561318 | 0.00311524 |
| 208968 | Zfp280c | 1238.05332 | 0.34531296 | 0.11683685 | 2.95551425 | 0.00312148 |
| 20715 | Serpina3g | 349.321805 | -1.2022115 | 0.40677784 | -2.9554498 | 0.00312213 |
| 381284 | CROCC2 | 19.8522645 | 1.34953601 | 0.45678866 | 2.95439907 | 0.00313279 |
| 59031 | CHST12 | 630.63307 | 0.41265377 | 0.1396757 | 2.95437047 | 0.00313308 |
| 68304 | KDELC2 | 1147.49707 | 0.42839249 | 0.14501152 | 2.95419637 | 0.00313484 |
| 69441 | 1700023F06I | 9.41720352 | -2.888353 | 0.97774108 | -2.9541082 | 0.00313574 |
| 108900 | FAM72A | 150.397979 | 0.59048681 | 0.19988886 | 2.95407569 | 0.00313607 |
| 224904 | 2410015M2C | 793.98312 | -0.3905687 | 0.1322821 | -2.9525439 | 0.00315167 |
| 18412 | SQSTM1 | 23286.3556 | -0.2521699 | 0.08541363 | -2.9523379 | 0.00315378 |
| 14184 | FGFR3 | 29.7720358 | -1.7071398 | 0.57845364 | -2.9512129 | 0.00316529 |
| 67246 | 2810474O19 | 2079.53381 | -0.6512998 | 0.22075251 | -2.9503619 | 0.00317402 |
| 68559 | PDRG1 | 831.892825 | -0.5253164 | 0.17806977 | -2.9500597 | 0.00317713 |
| 99100 | CEP152 | 609.533962 | 0.26207974 | 0.08885187 | 2.94962561 | 0.00318159 |
| 68298 | NCAPD2 | 4963.08923 | 0.54477573 | 0.1847249 | 2.94911917 | 0.00318681 |
| 384214 | EPHX4 | 6.39796661 | -3.5626289 | 1.20880603 | -2.9472296 | 0.00320635 |
| 52064 | COQ5 | 814.740742 | -0.2855298 | 0.09688423 | -2.9471237 | 0.00320745 |
| 100502925 | #N/A | 73.5454466 | 0.63252856 | 0.2146669 | 2.94655841 | 0.00321332 |
| 194952 | JMJD4 | 703.5443 | -0.323153 | 0.10968174 | -2.946279 | 0.00321622 |
| 24074 | taf7 | 160.92893 | -0.4987604 | 0.16939167 | -2.9444214 | 0.00323559 |
| 72931 | SWI5 | 3082.66841 | -0.2400786 | 0.08154442 | -2.9441458 | 0.00323847 |
| 12330 | CANX | 28647.7753 | 0.07751849 | 0.02633244 | 2.94384065 | 0.00324167 |
| 24068 | SRA1 | 961.197856 | -0.3788771 | 0.12871545 | -2.9435247 | 0.00324498 |
| 54382 | Tcstv1 | 3.8754054 | -5.3795115 | 1.82828793 | -2.9423765 | 0.00325704 |
| 20778 | SCARB1 | 3380.46311 | 0.32318795 | 0.10984607 | 2.94218945 | 0.003259 |
| 72603 | 2700033N17 | 12.549025 | 1.74852551 | 0.59451034 | 2.94111876 | 0.00327029 |
| 11669 | ALDH2 | 8119.11287 | -0.5862594 | 0.19933605 | -2.9410603 | 0.00327091 |
| 102607 | SNX19 | 1794.22704 | 0.18956507 | 0.06449149 | 2.93938134 | 0.00328868 |
| 18984 | POR | 3868.40349 | -0.5441395 | 0.18513131 | -2.9392084 | 0.00329052 |
| 233726 | IPO7 | 9676.97038 | 0.7056077 | 0.24007098 | 2.93916284 | 0.003291 |
| 100504267 | Trp53cor1 | 24.7064263 | -1.7214569 | 0.58572506 | -2.9390186 | 0.00329253 |
| 224648 | UHRF1BP1 | 1570.57456 | 0.64114051 | 0.21814859 | 2.93900827 | 0.00329264 |
| 626275 | A930012L18I | 24.654653 | 0.91630464 | 0.31182701 | 2.93850316 | 0.00329801 |
| 101543 | WTIP | 472.109265 | 2.29863471 | 0.7827407 | 2.93664902 | 0.00331779 |

| | | | | | | |
|---|---|---|---|---|---|---|
| 50997 | MPP2 | 169.57405 | 2.10368649 | 0.71637993 | 2.9365514 | 0.00331884 |
| 67150 | RNF141 | 179.633922 | -0.5427195 | 0.18483788 | -2.936192 | 0.00332269 |
| 66197 | CKS2 | 63.6976114 | 0.83725166 | 0.28515317 | 2.93614717 | 0.00332317 |
| 14105 | SRSF10 | 2016.49353 | 0.30783693 | 0.10484669 | 2.93606725 | 0.00332402 |
| 170625 | SNX18 | 3677.65648 | -0.406066 | 0.13830364 | -2.9360472 | 0.00332424 |
| 15139 | Hc | 7.31843174 | -2.0724923 | 0.70602809 | -2.9354247 | 0.00333091 |
| 93679 | TRIM8 | 2704.47646 | -0.2129955 | 0.07256594 | -2.9351994 | 0.00333333 |
| 74504 | FAM53A | 884.606516 | 0.31038414 | 0.10574863 | 2.93511277 | 0.00333426 |
| 208666 | DIRAS1 | 8.78117336 | -4.0311672 | 1.37343002 | -2.9351093 | 0.0033343 |
| 66599 | RDM1 | 137.859954 | -1.01031 | 0.34429895 | -2.9343976 | 0.00334196 |
| 237754 | BTNL9 | 3.51563988 | -4.4592699 | 1.51989436 | -2.9339341 | 0.00334695 |
| 22690 | ZFP28 | 104.36728 | -0.5302767 | 0.18082602 | -2.9325241 | 0.00336219 |
| 66277 | KLF15 | 9.03979758 | 2.08466635 | 0.71155884 | 2.92971747 | 0.0033927 |
| 17218 | MCM5 | 6297.52797 | 0.32968287 | 0.11253334 | 2.92964627 | 0.00339348 |
| 79464 | LIAS | 740.216882 | -0.2530581 | 0.08639285 | -2.929156 | 0.00339884 |
| 69020 | Zfp707 | 185.020344 | -0.4180597 | 0.14272847 | -2.9290562 | 0.00339993 |
| 241919 | SLC7A14 | 4.01082056 | -3.3035929 | 1.12790474 | -2.9289644 | 0.00340093 |
| 66234 | MSMO1 | 7188.07974 | -0.3822128 | 0.13049961 | -2.9288424 | 0.00340227 |
| 77782 | POLQ | 1002.40474 | 0.29238662 | 0.0998446 | 2.92841701 | 0.00340693 |
| 67881 | MDP1 | 539.106158 | 0.61722651 | 0.21078147 | 2.92827692 | 0.00340846 |
| 17120 | MAD1L1 | 1613.26372 | 0.22241269 | 0.07597952 | 2.92727157 | 0.0034195 |
| 217265 | ABCA5 | 1522.10835 | 0.42407459 | 0.14487294 | 2.92721733 | 0.0034201 |
| 225651 | MPPE1 | 286.402387 | 0.39884128 | 0.13628818 | 2.92645539 | 0.00342849 |
| 116873 | STIM2 | 1877.41705 | -0.2140882 | 0.07315701 | -2.9264213 | 0.00342886 |
| 112419 | Ifit1bl2 | 3.59283634 | -4.5091355 | 1.5408877 | -2.9263233 | 0.00342994 |
| 213988 | TNRC6B | 2638.79539 | 0.34593529 | 0.11821609 | 2.92629606 | 0.00343024 |
| 75785 | KLHL24 | 1316.82714 | -0.6377943 | 0.21801109 | -2.9255132 | 0.00343889 |
| 17449 | MDH1 | 3432.27262 | -0.1323578 | 0.04524426 | -2.9254059 | 0.00344007 |
| 74691 | TDRD9 | 37.729934 | 2.28131061 | 0.77983007 | 2.92539454 | 0.0034402 |
| 74754 | DHCR24 | 8012.00117 | 0.63156429 | 0.21598766 | 2.92407573 | 0.00345481 |
| 100041677 | Zfp984 | 218.166005 | 0.37648914 | 0.12876649 | 2.92381304 | 0.00345772 |
| 13713 | ELK3 | 5064.79299 | 0.16527117 | 0.05653435 | 2.92337606 | 0.00346258 |
| 227700 | SH3GLB2 | 965.338613 | -0.3707061 | 0.1268329 | -2.9227911 | 0.00346909 |
| 80915 | DUSP12 | 643.82497 | 0.19316838 | 0.06609638 | 2.92252611 | 0.00347205 |
| 215384 | FCGBP | 3.62078537 | -3.7279645 | 1.27571658 | -2.9222514 | 0.00347511 |
| 30838 | FBXW4 | 431.499168 | -0.3729536 | 0.12763747 | -2.9219755 | 0.00347819 |
| 77106 | Tmem181a | 253.336201 | 0.45079983 | 0.15433007 | 2.92101094 | 0.00348898 |
| 14211 | SMC2 | 7193.07824 | 0.235825 | 0.08074842 | 2.92049044 | 0.00349481 |
| 70681 | FAM175A | 399.685494 | -0.3095328 | 0.10600523 | -2.9199771 | 0.00350057 |
| 224139 | GOLGB1 | 3575.14093 | -0.3726914 | 0.12765489 | -2.9195235 | 0.00350567 |
| 100504501 | Gm20257 | 35.5533928 | 0.96518648 | 0.33061003 | 2.91941075 | 0.00350694 |
| 76987 | HDHD2 | 571.454705 | -0.2641792 | 0.09055233 | -2.9174202 | 0.0035294 |
| 68552 | SMIM14 | 1045.88409 | -0.6063681 | 0.20786502 | -2.9171244 | 0.00353275 |
| 170643 | KIRREL | 7211.63478 | 0.36587784 | 0.12542965 | 2.91699652 | 0.0035342 |
| 53619 | BLCAP | 1617.95477 | -0.766103 | 0.26264622 | -2.9168626 | 0.00353571 |

| | | | | | | |
|---|---|---|---|---|---|---|
| 17951 | Naip5 | 9.71067642 | -1.7289218 | 0.59284009 | -2.9163376 | 0.00354167 |
| 67065 | POLR3D | 938.103186 | 0.78507374 | 0.26920238 | 2.91629567 | 0.00354215 |
| 320203 | C130071C03I | 4.22285165 | 4.0547992 | 1.39073655 | 2.91557677 | 0.00355032 |
| 16365 | ACOD1 | 57.5049441 | -1.4557985 | 0.49933673 | -2.9154644 | 0.0035516 |
| 71835 | LANCL2 | 971.678132 | 0.32025139 | 0.10986213 | 2.91502982 | 0.00355655 |
| 56632 | SPHK2 | 804.988254 | 0.3941071 | 0.13522274 | 2.91450312 | 0.00356255 |
| 83429 | CTNS | 803.697441 | -0.6580446 | 0.22580539 | -2.9142113 | 0.00356588 |
| 22221 | UBP1 | 1936.38314 | 0.25910126 | 0.08892282 | 2.91377693 | 0.00357085 |
| 106795 | TCF19 | 1698.65073 | 0.21712156 | 0.07453704 | 2.91293519 | 0.00358049 |
| 14756 | GPLD1 | 20.2478181 | -1.5434029 | 0.52989785 | -2.9126423 | 0.00358385 |
| 631584 | Gm7068 | 2.9475163 | -4.2095772 | 1.44637497 | -2.9104328 | 0.00360929 |
| 68634 | TM2D3 | 298.391727 | -0.4903928 | 0.16852442 | -2.9099215 | 0.0036152 |
| 68209 | rnaseh2c | 526.067844 | 0.33147515 | 0.1139126 | 2.90990759 | 0.00361536 |
| 67784 | PLXND1 | 1261.97497 | 1.21473233 | 0.41750938 | 2.90947312 | 0.00362039 |
| 71099 | TSSK4 | 13.1780463 | 1.26908933 | 0.43620111 | 2.90941336 | 0.00362108 |
| 100038725 | CEP85L | 92.8924265 | 0.64331102 | 0.22111424 | 2.90940558 | 0.00362117 |
| 403187 | OPA3 | 1378.8301 | 0.50824045 | 0.17469346 | 2.90932735 | 0.00362207 |
| 13846 | EPHB4 | 3111.27272 | -0.4847204 | 0.16663898 | -2.9088056 | 0.00362812 |
| 14356 | TIMM10B | 567.185334 | 0.67460351 | 0.23197941 | 2.90803182 | 0.00363711 |
| 19181 | PSMC2 | 5108.32078 | -0.1232343 | 0.04238958 | -2.9071847 | 0.00364698 |
| 100090 | ZBTB48 | 199.920643 | 0.4657613 | 0.16029368 | 2.9056748 | 0.00366462 |
| 18419 | OTOG | 7.51729747 | -6.3385996 | 2.18152799 | -2.905578 | 0.00366576 |
| 227522 | RPP38 | 221.167465 | -0.3855623 | 0.132709 | -2.9053217 | 0.00366876 |
| 13199 | DDN | 6.99824893 | 2.4731133 | 0.8515449 | 2.90426646 | 0.00368115 |
| 108934 | SMIM13 | 699.25032 | 0.61047008 | 0.21025093 | 2.90353095 | 0.0036898 |
| 67038 | 2010109I03R | 3.35034795 | -5.1682307 | 1.78043973 | -2.9027833 | 0.00369862 |
| 64652 | NISCH | 6884.07978 | 0.51752282 | 0.17828618 | 2.9027646 | 0.00369885 |
| 70458 | 2610318N02 | 3.83891388 | -3.81988 | 1.31617568 | -2.902257 | 0.00370484 |
| 12850 | COQ7 | 757.722559 | -0.3252393 | 0.11206804 | -2.902159 | 0.003706 |
| 407823 | BAZ2B | 1145.02429 | 0.38837711 | 0.13386652 | 2.90122666 | 0.00371705 |
| 67095 | trak1 | 1470.51496 | 0.26617002 | 0.09176239 | 2.90064393 | 0.00372397 |
| 239719 | MKL2 | 1926.14341 | -0.4049172 | 0.13962096 | -2.9001177 | 0.00373023 |
| 100503021 | Gm19510 | 16.4609736 | -2.0384733 | 0.70294625 | -2.8998992 | 0.00373283 |
| 21754 | TESK1 | 1320.89657 | -0.4900185 | 0.16908127 | -2.8981239 | 0.00375402 |
| 15277 | HK2 | 2631.40636 | 0.69380568 | 0.23940961 | 2.897986 | 0.00375567 |
| 73545 | 1700094D03 | 89.8810534 | -0.6029898 | 0.20810161 | -2.8975739 | 0.00376061 |
| 100504448 | Gm20234 | 3.59058016 | -5.2695986 | 1.81868298 | -2.8974806 | 0.00376173 |
| 13207 | DDX5 | 11831.9484 | 0.25155226 | 0.08682192 | 2.89733588 | 0.00376347 |
| 100043147 | Gm4262 | 96.1946367 | 1.05449138 | 0.36396711 | 2.89721612 | 0.0037649 |
| 68865 | ARV1 | 202.857148 | -0.4204419 | 0.14513605 | -2.8968813 | 0.00376892 |
| 621976 | tmem170b | 627.093552 | -0.3706045 | 0.12793816 | -2.8967475 | 0.00377053 |
| 14659 | Glrp1 | 21.8836503 | -2.4305228 | 0.83933373 | -2.8957764 | 0.00378222 |
| 225896 | UBXN1 | 2359.58709 | -0.2775383 | 0.09584296 | -2.8957605 | 0.00378241 |
| 208846 | DAAM1 | 2462.85599 | 0.15836851 | 0.05470198 | 2.89511469 | 0.0037902 |
| 12018 | BAK1 | 3515.66036 | 0.78634194 | 0.27162208 | 2.89498538 | 0.00379176 |

| | | | | | | |
|---|---|---|---|---|---|---|
| 11647 | ALPL | 5.98505696 | -2.3121745 | 0.79873896 | -2.8947812 | 0.00379423 |
| 19113 | Prl7a1 | 3.59795491 | -3.7417639 | 1.29265465 | -2.8946354 | 0.00379599 |
| 319748 | Zfp865 | 790.698424 | 0.38739491 | 0.13384349 | 2.89438741 | 0.00379899 |
| 28030 | GFM1 | 2406.31578 | 0.13848078 | 0.04785061 | 2.8940233 | 0.0038034 |
| 15042 | H2-T24 | 14.9414632 | -1.5213905 | 0.52570283 | -2.8940123 | 0.00380353 |
| 19045 | PPP1CA | 10283.1445 | -0.1681623 | 0.05810753 | -2.8939841 | 0.00380387 |
| 73046 | GLRX5 | 2294.40709 | 0.65440577 | 0.22618543 | 2.89322684 | 0.00381306 |
| 241627 | WDR76 | 735.447826 | -0.4348307 | 0.15030914 | -2.8929092 | 0.00381692 |
| 72727 | B3GAT3 | 869.362308 | -0.2262295 | 0.07821324 | -2.8924699 | 0.00382226 |
| 100504040 | Gm17586 | 153.892005 | -0.4611919 | 0.15947716 | -2.8918994 | 0.00382921 |
| 432730 | Gm5446 | 237.713308 | 0.36984212 | 0.12789951 | 2.89166184 | 0.0038321 |
| 73490 | MIPOL1 | 185.047913 | 0.44292295 | 0.15318785 | 2.89137135 | 0.00383565 |
| 67471 | GPATCH1 | 420.02384 | 0.37464161 | 0.12960059 | 2.89074009 | 0.00384336 |
| 66362 | EXOSC3 | 884.21503 | 0.56735014 | 0.19631281 | 2.89003121 | 0.00385204 |
| 76899 | GOLGA1 | 1086.61581 | -0.1831787 | 0.06338486 | -2.8899447 | 0.0038531 |
| 170826 | PPARGC1B | 91.4796212 | 1.32571953 | 0.4588575 | 2.88917479 | 0.00386254 |
| 232196 | C87436 | 458.881741 | 0.4740353 | 0.16408016 | 2.88904702 | 0.00386411 |
| 242574 | C130073F10I | 3.91601384 | -4.6264919 | 1.60169733 | -2.8884932 | 0.00387092 |
| 56332 | AMOTL2 | 6674.29994 | 0.77673654 | 0.2689306 | 2.88824159 | 0.00387402 |
| 18505 | PAX3 | 14.7806395 | 2.8666656 | 0.99270415 | 2.88773406 | 0.00388028 |
| 13180 | PCBD1 | 9.71850501 | -6.7120104 | 2.32546481 | -2.8863092 | 0.00389789 |
| 67579 | CPEB4 | 530.649627 | -0.3396538 | 0.11768853 | -2.8860401 | 0.00390122 |
| 212898 | DSE | 2436.32927 | 0.81106398 | 0.28103159 | 2.88602421 | 0.00390142 |
| 108645 | MAT2B | 1606.42051 | 0.17201005 | 0.05961837 | 2.88518532 | 0.00391183 |
| 75786 | CKAP5 | 7013.86128 | 0.22427138 | 0.07773232 | 2.88517537 | 0.00391196 |
| 18242 | OAT | 2704.39011 | 0.67343904 | 0.23342267 | 2.88506272 | 0.00391336 |
| 71296 | CRNDE | 90.521299 | -0.6861499 | 0.23786902 | -2.8845703 | 0.00391948 |
| 72355 | CDPF1 | 130.37602 | -0.4760388 | 0.16510227 | -2.8832967 | 0.00393537 |
| 320271 | SCAI | 721.433232 | 0.47916258 | 0.1662043 | 2.88297339 | 0.00393941 |
| 620235 | SIGLEC15 | 11.2460582 | -2.3125385 | 0.80223889 | -2.8826058 | 0.00394401 |
| 56508 | RAPGEF4 | 15.4937371 | 2.18827445 | 0.75916353 | 2.88248099 | 0.00394557 |
| 432467 | HNRNPH3 | 518.022257 | 0.49929175 | 0.17325762 | 2.88178803 | 0.00395426 |
| 57277 | SLURP1 | 4.38114578 | 3.48039034 | 1.20783139 | 2.88152003 | 0.00395762 |
| 14453 | GAS2 | 100.091031 | 0.85619571 | 0.29718418 | 2.88102717 | 0.00396381 |
| 23897 | HAX1 | 230.383386 | -0.3383361 | 0.11746442 | -2.8803282 | 0.00397261 |
| 18572 | PDCD11 | 3853.67497 | 0.23239756 | 0.0807113 | 2.87936828 | 0.00398473 |
| 67298 | GPRASP1 | 361.44175 | -0.4240692 | 0.14731745 | -2.8786083 | 0.00399434 |
| 66067 | GTPBP8 | 134.914259 | -0.4648448 | 0.16148443 | -2.8785732 | 0.00399479 |
| 17156 | MAN1A2 | 5004.77561 | 0.28582234 | 0.09930629 | 2.87818986 | 0.00399964 |
| 68910 | Zfp467 | 19.2098121 | -3.715229 | 1.29099864 | -2.8777947 | 0.00400466 |
| 54197 | RNF5 | 420.322516 | -0.3783593 | 0.13149557 | -2.8773536 | 0.00401026 |
| 666170 | Gm7964 | 6200.03891 | 0.37753724 | 0.13128425 | 2.87572382 | 0.00403102 |
| 17160 | man2b2 | 1562.86626 | -0.8211268 | 0.28554282 | -2.8756694 | 0.00403172 |
| 235041 | KANK2 | 2969.96982 | 0.76727931 | 0.26685659 | 2.87524966 | 0.00403708 |
| 97387 | STRN4 | 2935.8131 | 0.43182284 | 0.15021528 | 2.87469322 | 0.0040442 |

| | | | | | | |
|---|---|---|---|---|---|---|
| 66799 | UBE2W | 984.065748 | -0.2720998 | 0.09465585 | -2.8746226 | 0.00404511 |
| 269152 | KIF26B | 156.967716 | 0.88547912 | 0.3080517 | 2.8744497 | 0.00404732 |
| 71673 | RNF215 | 377.537581 | -0.3612849 | 0.12570213 | -2.8741355 | 0.00405135 |
| 19017 | PPARGC1A | 82.3849655 | -3.8357252 | 1.33457591 | -2.8741154 | 0.00405161 |
| 67015 | CCDC91 | 619.271579 | 0.2955261 | 0.10289595 | 2.87208692 | 0.00407771 |
| 11972 | ATP6V0D1 | 2726.69132 | -0.4503283 | 0.15685808 | -2.8709283 | 0.00409268 |
| 78772 | HHIPL2 | 12.1377899 | -3.287134 | 1.14518003 | -2.870408 | 0.00409942 |
| 18655 | PGK1 | 692.882081 | 0.29458466 | 0.10262825 | 2.87040514 | 0.00409946 |
| 54392 | NCAPG | 3017.21138 | 0.33790761 | 0.11772562 | 2.87029807 | 0.00410085 |
| 68277 | 2310057M21 | 423.652696 | 0.45752877 | 0.15949519 | 2.86860544 | 0.00412286 |
| 55991 | PANX1 | 328.411971 | 2.03336907 | 0.70892964 | 2.86822409 | 0.00412783 |
| 209966 | PGBD5 | 8.2152666 | -3.3204783 | 1.15787593 | -2.8677324 | 0.00413425 |
| 72937 | #N/A | 3.5755849 | 3.79333902 | 1.3233719 | 2.86641949 | 0.00415144 |
| 100504343 | #N/A | 7.83290007 | 2.03237923 | 0.70926879 | 2.86545702 | 0.00416408 |
| 27359 | SYTL4 | 3.86136024 | -3.8335139 | 1.33802827 | -2.865047 | 0.00416947 |
| 107182 | BTAF1 | 3462.65393 | 0.22416046 | 0.078253 | 2.86456066 | 0.00417588 |
| 14672 | GNA11 | 4706.71674 | 0.14961751 | 0.05223526 | 2.86430103 | 0.00417931 |
| 57266 | CXCL14 | 7.63965135 | -5.6136706 | 1.96010982 | -2.8639572 | 0.00418385 |
| 19240 | TMSB10 | 3147.59996 | 0.48920555 | 0.17081571 | 2.86393771 | 0.0041841 |
| 101142 | ITFG2 | 686.483881 | 0.33979085 | 0.11865306 | 2.86373447 | 0.00418679 |
| 233878 | SEZ6L2 | 8.99591694 | -1.9456365 | 0.67941901 | -2.8636768 | 0.00418755 |
| 105504 | EXOC5 | 3507.82537 | 0.51161808 | 0.17877072 | 2.86186731 | 0.00421153 |
| 20431 | PMEL | 16.9876945 | -2.0014907 | 0.69959658 | -2.8609212 | 0.00422412 |
| 71817 | TMEM50A | 1405.92491 | 0.26997384 | 0.09437557 | 2.8606328 | 0.00422796 |
| 235599 | 6430571L13F | 6.9689474 | -2.3404648 | 0.81819436 | -2.8605242 | 0.00422941 |
| 14084 | FAF1 | 2414.73119 | 0.38029508 | 0.13295127 | 2.86040965 | 0.00423094 |
| 239760 | Gm4943 | 32.4380271 | -0.7357999 | 0.25728275 | -2.8598884 | 0.0042379 |
| 11847 | ARG2 | 36.051845 | -1.5916172 | 0.55660579 | -2.8595054 | 0.00424302 |
| 20753 | SPRR1A | 6.80437485 | -2.7745017 | 0.97044803 | -2.8589905 | 0.00424991 |
| 69814 | Prss32 | 3.56045241 | -5.2584432 | 1.83954309 | -2.8585594 | 0.00425569 |
| 66190 | ACER3 | 1812.80292 | -0.3543707 | 0.12396891 | -2.8585452 | 0.00425588 |
| 234734 | AARS | 11017.8203 | 0.11197688 | 0.03917384 | 2.85846038 | 0.00425702 |
| 11765 | AP1G1 | 4472.60041 | -0.1803695 | 0.06310148 | -2.8584033 | 0.00425779 |
| 140546 | ERI3 | 991.106207 | 0.54447013 | 0.19049536 | 2.85818055 | 0.00426078 |
| 192970 | DHRS11 | 187.280833 | -0.6521963 | 0.22822187 | -2.857729 | 0.00426685 |
| 217127 | KAT7 | 3965.79289 | 0.16784081 | 0.05873554 | 2.85756808 | 0.00426901 |
| 70361 | LMAN1 | 4748.16424 | 0.34029735 | 0.11909455 | 2.85737126 | 0.00427166 |
| 27412 | Peg12 | 4.26846359 | -5.5199587 | 1.93246192 | -2.8564385 | 0.00428423 |
| 109032 | SP110 | 6.27429946 | -2.3885236 | 0.83624383 | -2.8562526 | 0.00428674 |
| 226695 | Ifi205 | 52.6009297 | -1.1304345 | 0.39589616 | -2.8553812 | 0.00429852 |
| 59030 | MKKS | 678.863726 | 0.27990895 | 0.09806042 | 2.85445397 | 0.00431109 |
| 14545 | GDAP1 | 3.44774767 | -5.2114208 | 1.82597426 | -2.8540494 | 0.00431658 |
| 266781 | SNX17 | 2230.4614 | 0.24925423 | 0.08733386 | 2.85403877 | 0.00431673 |
| 668724 | Gm9320 | 13.0820296 | -1.6127029 | 0.56517775 | -2.8534437 | 0.00432482 |
| 54204 | 41882 | 17.484649 | -1.2326863 | 0.43201432 | -2.8533458 | 0.00432615 |

| | | | | | | |
|---|---|---|---|---|---|---|
| 14113 | FBL | 476.814808 | 0.57234612 | 0.20074313 | 2.85113683 | 0.00435632 |
| 70314 | RABEP2 | 515.681498 | 0.24527908 | 0.0860288 | 2.85112736 | 0.00435645 |
| 245622 | FAM199X | 899.995926 | 0.29075619 | 0.10202222 | 2.84992987 | 0.00437289 |
| 80898 | ERAP1 | 1168.95628 | -0.3649113 | 0.12804409 | -2.8498879 | 0.00437346 |
| 75686 | NUDT16 | 61.7167684 | 1.3439785 | 0.4716567 | 2.84948459 | 0.00437901 |
| 19365 | RAD52 | 243.035916 | 0.39223765 | 0.13766009 | 2.84932012 | 0.00438128 |
| 230996 | 9430015G10 | 555.421729 | 0.52043704 | 0.18270692 | 2.84848014 | 0.00439286 |
| 54343 | ATF7IP | 2091.47807 | -0.4506958 | 0.15824622 | -2.8480665 | 0.00439857 |
| 72900 | NDUFV2 | 1651.73451 | -0.2020428 | 0.07096504 | -2.8470756 | 0.00441229 |
| 229285 | SPG20 | 1838.84002 | 0.30720084 | 0.10791267 | 2.8467541 | 0.00441675 |
| 621699 | Gm6252 | 270.848222 | 0.4789897 | 0.16834652 | 2.84526048 | 0.00443751 |
| 11606 | AGT | 7.32951123 | 2.68320411 | 0.9432572 | 2.84461558 | 0.00444651 |
| 109245 | LRRC39 | 11.296836 | -3.5203322 | 1.23758225 | -2.8445238 | 0.00444779 |
| 66262 | ING5 | 1212.08801 | 0.23401063 | 0.08227069 | 2.84439868 | 0.00444953 |
| 109676 | ANK2 | 214.495272 | 1.13463187 | 0.39902847 | 2.84348598 | 0.0044623 |
| 227731 | SLC25A25 | 720.543517 | -0.6080822 | 0.21386979 | -2.8432355 | 0.00446581 |
| 215707 | CCDC92 | 424.698969 | -0.6803899 | 0.23932472 | -2.8429572 | 0.00446971 |
| 117599 | HELB | 645.284208 | -0.2353019 | 0.08276694 | -2.8429452 | 0.00446988 |
| 50770 | ATP11A | 5580.08692 | -0.5574015 | 0.19606917 | -2.8428818 | 0.00447076 |
| 18087 | NKTR | 2830.08586 | 0.2846089 | 0.10011355 | 2.84286097 | 0.00447106 |
| 269470 | WDR3 | 2431.37875 | 0.29483052 | 0.10372641 | 2.84238635 | 0.00447772 |
| 68833 | PDCL3 | 1620.33196 | -0.5610047 | 0.19737931 | -2.8422669 | 0.0044794 |
| 245469 | PDZD4 | 316.652358 | 0.41239404 | 0.14512483 | 2.84165039 | 0.00448807 |
| 20511 | SLC1A2 | 70.7398882 | -2.1947498 | 0.77252733 | -2.8409995 | 0.00449724 |
| 12479 | Cd1d1 | 117.93056 | 0.66007032 | 0.23236013 | 2.84072105 | 0.00450117 |
| 54371 | CHST2 | 1080.19146 | -0.6731688 | 0.23704551 | -2.8398295 | 0.00451377 |
| 56520 | NME4 | 130.159976 | 0.66531329 | 0.2343045 | 2.83952415 | 0.00451809 |
| 67959 | PUF60 | 3561.41623 | -0.1514311 | 0.05334356 | -2.8387896 | 0.0045285 |
| 19085 | PRKAR1B | 19.6480662 | 0.99929754 | 0.35202366 | 2.8387227 | 0.00452945 |
| 319660 | AGMO | 25.9245957 | -1.7566616 | 0.61884726 | -2.8386028 | 0.00453115 |
| 268933 | WDR24 | 633.985864 | 0.45055285 | 0.1587498 | 2.83813167 | 0.00453785 |
| 16194 | il6ra | 353.305201 | -0.8293282 | 0.29228612 | -2.837385 | 0.00454847 |
| 69754 | FBXO7 | 537.047419 | -0.2184596 | 0.07700729 | -2.8368697 | 0.00455582 |
| 11727 | ANG | 8.34491718 | 1.69445609 | 0.59733657 | 2.83668565 | 0.00455845 |
| 14674 | GNA13 | 4152.78163 | 0.20175739 | 0.07113413 | 2.83629522 | 0.00456402 |
| 394435 | Ugt1a6b | 72.2141794 | 1.05598577 | 0.3723559 | 2.83595816 | 0.00456884 |
| 14777 | Gpx2-ps1 | 4.1438757 | -5.4770206 | 1.93127967 | -2.8359541 | 0.0045689 |
| 18007 | NEO1 | 8344.23147 | 0.36914188 | 0.13018702 | 2.83547381 | 0.00457578 |
| 192166 | SARDH | 206.864364 | -1.0960812 | 0.38658746 | -2.8352736 | 0.00457865 |
| 67834 | IDH3A | 3885.83476 | 0.30000985 | 0.10582867 | 2.83486369 | 0.00458452 |
| 69068 | 1810011O10 | 336.059707 | -2.0119721 | 0.70972582 | -2.8348582 | 0.0045846 |
| 329908 | USP24 | 5126.07701 | 0.52112622 | 0.18384862 | 2.8345397 | 0.00458918 |
| 244058 | RGMA | 622.795424 | 0.96152791 | 0.33927922 | 2.83403126 | 0.00459648 |
| 72016 | 1600002H07 | 372.781794 | 0.74418149 | 0.26264315 | 2.83343196 | 0.00460511 |
| 108155 | OGT | 4490.68724 | -0.3919053 | 0.13835745 | -2.8325566 | 0.00461774 |

| | | | | | | |
|---|---|---|---|---|---|---|
| 26914 | h2afy | 3337.32403 | 0.38192607 | 0.13495659 | 2.8299919 | 0.00465492 |
| 100043489 | 1300002E11I | 143.86089 | -0.7304878 | 0.2581731 | -2.8294499 | 0.00466281 |
| 246257 | OVCA2 | 237.311157 | 0.39134537 | 0.13834274 | 2.82881039 | 0.00467214 |
| 94094 | Trim34a | 9.20184382 | -2.0426122 | 0.72210461 | -2.8286929 | 0.00467385 |
| 329877 | DENND4C | 1697.57338 | 0.33853711 | 0.11969118 | 2.82842155 | 0.00467782 |
| 67345 | HERC4 | 1988.09864 | -0.2184334 | 0.07726049 | -2.8272326 | 0.00469522 |
| 71904 | PAQR7 | 92.4879281 | -1.2555172 | 0.44428108 | -2.8259523 | 0.00471403 |
| 12034 | PHB2 | 6459.1125 | 0.34084083 | 0.12061194 | 2.82592939 | 0.00471437 |
| 18294 | OGG1 | 535.051744 | -0.2895642 | 0.10247934 | -2.8255862 | 0.00471942 |
| 654799 | 4930594M22 | 23.004481 | -1.3783415 | 0.48797994 | -2.8245863 | 0.00473417 |
| 78294 | RPS27A | 115.701579 | 0.45260948 | 0.16028038 | 2.82386076 | 0.0047449 |
| 52808 | TSPYL2 | 690.358378 | 0.22084723 | 0.07822548 | 2.8232136 | 0.00475449 |
| 235293 | SC5D | 3471.64103 | -0.2456087 | 0.08699745 | -2.8231714 | 0.00475511 |
| 17105 | lyz2 | 4.70300321 | -4.9124165 | 1.7401651 | -2.82296 | 0.00475825 |
| 70370 | fbln7 | 11.3007809 | -1.6146957 | 0.57206765 | -2.8225608 | 0.00476418 |
| 269952 | GDPGP1 | 278.443131 | 0.36182888 | 0.1282057 | 2.82225277 | 0.00476876 |
| 17259 | MEF2B | 5.72390923 | -3.0049135 | 1.06524534 | -2.8208652 | 0.00478943 |
| 30050 | FBXW2 | 1910.94075 | -0.4717731 | 0.16728182 | -2.8202296 | 0.00479893 |
| 20274 | SCN9A | 3.41749716 | -5.2028484 | 1.84485529 | -2.8201933 | 0.00479947 |
| 54409 | RAMP2 | 10.0719311 | 2.84144475 | 1.00768215 | 2.81978275 | 0.00480562 |
| 320127 | DGKI | 3.41291884 | -4.4255223 | 1.56949823 | -2.8197052 | 0.00480678 |
| 56554 | Raet1d | 97.9933042 | -3.28182 | 1.16411558 | -2.8191531 | 0.00481505 |
| 629557 | Gm6981 | 8.91537393 | 1.50282855 | 0.53311928 | 2.8189349 | 0.00481833 |
| 12514 | CD68 | 361.829215 | -1.3399642 | 0.47543555 | -2.8183929 | 0.00482647 |
| 104086 | CYP27A1 | 18.3429918 | -2.5774616 | 0.91502851 | -2.8168102 | 0.00485032 |
| 14857 | GSTA1 | 128.726571 | 0.72419301 | 0.25714649 | 2.81626631 | 0.00485854 |
| 100503636 | 1700123M08 | 14.3932374 | -1.6209196 | 0.57556545 | -2.8162211 | 0.00485922 |
| 227357 | ESPNL | 6.34554992 | -2.2361817 | 0.79425129 | -2.8154587 | 0.00487077 |
| 23789 | CORO1B | 4565.66597 | -0.4101818 | 0.14570525 | -2.8151481 | 0.00487548 |
| 15976 | IFNAR2 | 813.077641 | -0.3708302 | 0.13178575 | -2.8138871 | 0.00489464 |
| 225207 | Zfp521 | 9.47704205 | -4.1118426 | 1.46134616 | -2.8137362 | 0.00489694 |
| 234663 | DYNC1LI2 | 3970.98578 | -0.1982135 | 0.07045125 | -2.8134846 | 0.00490077 |
| 319934 | SBF2 | 2323.87719 | 0.59268178 | 0.21067637 | 2.8132333 | 0.00490461 |
| 193286 | BC049762 | 3.83356342 | -2.7176394 | 0.96617243 | -2.8127892 | 0.00491138 |
| 69482 | NUP35 | 240.86561 | 0.38422299 | 0.13660008 | 2.81275809 | 0.00491186 |
| 74407 | TTC25 | 3.95849639 | -3.2991807 | 1.17322076 | -2.8120715 | 0.00492236 |
| 102866 | PLS3 | 10715.8989 | 0.25984477 | 0.09240674 | 2.81196767 | 0.00492395 |
| 20133 | RRM1 | 11289.3547 | 0.54647082 | 0.19436491 | 2.8115714 | 0.00493001 |
| 22404 | WIZ | 2087.50039 | 0.15024609 | 0.05345658 | 2.81061909 | 0.00494463 |
| 66084 | RMND1 | 116.54315 | -0.6041437 | 0.21499116 | -2.8100861 | 0.00495283 |
| 72749 | TONSL | 888.5071 | 0.23318041 | 0.0829805 | 2.81006281 | 0.00495318 |
| 791299 | Gm10069 | 52.9821055 | -0.7521014 | 0.26769548 | -2.8095409 | 0.00496122 |
| 211147 | 41708 | 7.39000017 | 3.92042 | 1.39552216 | 2.80928538 | 0.00496516 |
| 67872 | NSMCE4A | 1433.95083 | 0.47658891 | 0.16968703 | 2.80863491 | 0.0049752 |
| 78330 | NDUFV3 | 659.430074 | -0.6008246 | 0.2139237 | -2.8085929 | 0.00497585 |

| | | | | | | |
|---|---|---|---|---|---|---|
| 245866 | IFT52 | 732.61028 | -0.3228534 | 0.11499522 | -2.8075376 | 0.00499218 |
| 241490 | RBM45 | 1281.8764 | 0.21814666 | 0.07771546 | 2.80699184 | 0.00500065 |
| 216848 | CHD3 | 2587.49904 | -0.4931786 | 0.17574469 | -2.806222 | 0.00501261 |
| 99237 | TM9SF4 | 2473.38991 | -0.4253604 | 0.15158411 | -2.8061014 | 0.00501449 |
| 14163 | FGD1 | 1019.85235 | 0.25220194 | 0.08987762 | 2.80605933 | 0.00501515 |
| 105446 | GMPR2 | 609.091978 | 0.64174966 | 0.22874582 | 2.80551429 | 0.00502364 |
| 192287 | SLC25A36 | 562.590007 | 0.42825666 | 0.15265386 | 2.8054099 | 0.00502526 |
| 56210 | REV1 | 1314.11945 | -0.1420708 | 0.05066084 | -2.804352 | 0.00504178 |
| 231014 | 9330182L06F | 92.7912446 | -1.5496839 | 0.55263344 | -2.8041805 | 0.00504447 |
| 236790 | INTS6L | 245.051826 | 0.63406554 | 0.22614689 | 2.80377736 | 0.00505078 |
| 23853 | DEF6 | 664.693267 | 1.62271399 | 0.57877255 | 2.80371622 | 0.00505173 |
| 246316 | LGI2 | 15.6597195 | -1.9899764 | 0.70982772 | -2.8034639 | 0.00505569 |
| 56224 | TSPAN5 | 2068.40935 | 0.46039295 | 0.16433195 | 2.80160348 | 0.00508493 |
| 68083 | PAK1IP1 | 2364.32191 | 0.16086389 | 0.05743855 | 2.80062577 | 0.00510036 |
| 15200 | HBEGF | 1834.66035 | -0.7667575 | 0.2738102 | -2.800325 | 0.00510512 |
| 224814 | ABCC10 | 255.809674 | -0.5410155 | 0.19323805 | -2.7997357 | 0.00511445 |
| 320229 | 9530052C20I | 16.8150092 | -1.2510476 | 0.44709007 | -2.7982005 | 0.00513882 |
| 66989 | KCTD20 | 1381.84778 | 0.69647523 | 0.24894533 | 2.7977036 | 0.00514673 |
| 320938 | TNPO3 | 4569.80557 | 0.14490752 | 0.05180997 | 2.79690379 | 0.00515949 |
| 14187 | Akr1b8 | 786.173958 | 2.04379443 | 0.73081266 | 2.79660513 | 0.00516426 |
| 67979 | ATAD1 | 3517.69047 | -0.2471038 | 0.08836562 | -2.7963794 | 0.00516787 |
| 74498 | GORASP1 | 596.313321 | 0.42968964 | 0.15368924 | 2.79583422 | 0.00517659 |
| 67501 | CCDC50 | 4186.80729 | 0.42563989 | 0.15236946 | 2.79347242 | 0.00521455 |
| 68215 | FAM98B | 1548.79352 | 0.22695814 | 0.08124607 | 2.79346605 | 0.00521465 |
| 266459 | Gm5039 | 3.77004754 | -3.7734652 | 1.35142019 | -2.792222 | 0.00523474 |
| 75695 | RILPL1 | 444.691319 | -0.4887911 | 0.17507089 | -2.7919611 | 0.00523896 |
| 230822 | NCMAP | 39.3652466 | -2.2246109 | 0.7968095 | -2.791898 | 0.00523999 |
| 16175 | IL1A | 5.19180179 | -4.2568319 | 1.52486934 | -2.7916043 | 0.00524474 |
| 14585 | GFRA1 | 53.7595502 | 2.39165579 | 0.85700212 | 2.7907233 | 0.00525904 |
| 108927 | LHFP | 4331.8247 | 0.4422867 | 0.15849156 | 2.79060087 | 0.00526103 |
| 225929 | patl1 | 2025.60213 | 0.18603304 | 0.06666487 | 2.79057065 | 0.00526152 |

| padj |
|---|
| 0 |
| 0 |
| 0 |
| 0 |
| 0 |
| 0 |
| 0 |
| 0 |
| 0 |
| 0 |
| 0 |
| 0 |
| 0 |
| 0 |
| 0 |
| 0 |
| 0 |
| 0 |
| 0 |
| 0 |
| 0 |
| 1.21E-277 |
| 1.94E-276 |
| 3.11E-270 |
| 1.50E-264 |
| 2.44E-252 |
| 5.99E-251 |
| 2.00E-250 |
| 3.40E-239 |
| 2.57E-236 |
| 6.45E-236 |
| 1.21E-229 |
| 8.42E-228 |
| 6.18E-220 |
| 1.21E-218 |
| 5.18E-215 |
| 1.66E-208 |
| 1.95E-208 |
| 9.80E-202 |
| 3.82E-198 |
| 1.36E-197 |
| 1.07E-195 |
| 3.05E-194 |

| |
|---|
| 1.86E-193 |
| 3.75E-186 |
| 9.11E-184 |
| 4.32E-180 |
| 6.30E-179 |
| 3.30E-178 |
| 5.13E-176 |
| 8.84E-176 |
| 9.08E-175 |
| 5.05E-172 |
| 2.59E-170 |
| 1.20E-169 |
| 1.50E-167 |
| 4.54E-167 |
| 8.24E-167 |
| 2.86E-166 |
| 1.71E-165 |
| 4.51E-163 |
| 1.53E-162 |
| 1.70E-162 |
| 3.62E-162 |
| 2.24E-161 |
| 2.56E-158 |
| 1.86E-157 |
| 5.25E-157 |
| 2.74E-156 |
| 6.61E-155 |
| 6.64E-155 |
| 7.90E-155 |
| 1.35E-154 |
| 1.90E-154 |
| 1.72E-153 |
| 3.32E-153 |
| 1.64E-150 |
| 8.40E-150 |
| 2.17E-148 |
| 4.39E-148 |
| 1.12E-146 |
| 4.67E-146 |
| 1.98E-145 |
| 7.00E-145 |
| 1.00E-144 |
| 1.25E-144 |
| 4.39E-143 |
| 5.55E-142 |

2.30E-141
1.77E-137
5.96E-135
1.12E-134
6.96E-134
4.54E-133
4.67E-133
2.00E-132
3.28E-132
9.48E-132
1.32E-125
4.00E-125
5.01E-124
1.23E-123
2.29E-123
2.89E-123
1.28E-122
2.33E-122
2.89E-122
3.79E-122
4.38E-121
3.59E-120
1.18E-119
3.82E-119
5.87E-119
1.45E-117
5.18E-117
1.46E-115
6.89E-115
1.01E-113
5.40E-113
5.67E-113
1.56E-112
5.65E-112
9.21E-112
9.94E-112
3.65E-111
5.02E-111
5.78E-111
5.98E-111
1.10E-110
2.34E-110
5.49E-110
1.84E-109
4.07E-109

| |
|---|
| 1.08E-108 |
| 2.32E-108 |
| 9.03E-108 |
| 1.86E-107 |
| 2.37E-107 |
| 1.18E-106 |
| 1.22E-106 |
| 8.51E-106 |
| 1.21E-105 |
| 1.25E-105 |
| 2.77E-105 |
| 7.65E-105 |
| 2.45E-104 |
| 7.98E-104 |
| 1.25E-103 |
| 3.20E-103 |
| 6.87E-103 |
| 1.44E-102 |
| 2.31E-102 |
| 4.04E-102 |
| 1.28E-101 |
| 2.60E-100 |
| 2.34E-99 |
| 8.65E-99 |
| 3.48E-98 |
| 1.27E-97 |
| 2.09E-97 |
| 4.50E-97 |
| 2.37E-96 |
| 1.06E-95 |
| 3.35E-95 |
| 4.78E-95 |
| 5.46E-95 |
| 1.01E-94 |
| 1.15E-94 |
| 5.53E-94 |
| 1.19E-93 |
| 1.29E-93 |
| 2.14E-93 |
| 6.67E-93 |
| 7.90E-93 |
| 5.11E-92 |
| 9.93E-92 |
| 2.13E-91 |
| 2.23E-91 |

2.45E-91
5.71E-91
1.25E-90
1.42E-89
1.93E-89
3.13E-89
3.68E-89
5.89E-88
7.54E-88
7.72E-88
5.93E-87
9.12E-87
9.12E-87
2.56E-86
4.77E-86
1.80E-85
3.98E-85
1.86E-84
5.24E-84
6.60E-84
7.65E-84
1.87E-83
2.99E-83
4.73E-83
7.23E-83
1.07E-82
1.19E-82
3.24E-82
4.05E-82
5.15E-82
1.10E-81
3.59E-81
1.21E-80
1.50E-80
3.41E-80
3.86E-80
4.27E-80
2.02E-79
2.42E-79
3.16E-79
5.73E-79
1.12E-78
1.33E-78
1.38E-78
1.97E-78

3.58E-78
6.88E-78
1.00E-77
4.99E-77
1.86E-76
3.74E-76
4.75E-76
1.55E-75
1.84E-75
4.44E-75
4.48E-75
5.03E-75
1.81E-74
3.07E-74
6.77E-74
7.19E-74
1.00E-73
1.96E-73
3.29E-73
7.53E-73
1.17E-72
2.77E-72
3.46E-72
8.20E-72
1.01E-71
1.27E-71
2.44E-71
2.84E-71
7.14E-71
1.53E-70
4.15E-70
4.92E-70
1.82E-69
2.48E-69
3.42E-69
3.60E-69
4.44E-69
7.50E-69
1.00E-68
1.22E-68
2.45E-68
4.20E-68
5.25E-68
5.68E-68
7.64E-68

2.84E-67
4.19E-67
8.21E-67
1.40E-66
2.63E-66
8.09E-66
8.56E-66
1.36E-65
1.51E-65
3.80E-65
1.70E-64
1.89E-64
2.09E-64
2.30E-64
2.39E-64
2.81E-64
4.42E-64
4.59E-64
7.13E-64
7.26E-64
7.46E-64
8.36E-64
1.05E-63
1.28E-63
1.43E-63
2.11E-63
2.22E-63
2.38E-63
2.57E-63
2.71E-63
5.78E-63
5.99E-63
1.15E-62
1.63E-62
1.74E-62
1.91E-62
1.97E-62
2.10E-62
2.56E-62
4.07E-62
4.37E-62
4.79E-62
5.63E-62
6.35E-62
1.84E-61

| |
|---|
| 2.09E-61 |
| 2.78E-61 |
| 3.97E-61 |
| 4.45E-61 |
| 5.39E-61 |
| 7.03E-61 |
| 7.42E-61 |
| 8.01E-61 |
| 8.18E-61 |
| 1.79E-60 |
| 2.59E-60 |
| 2.66E-60 |
| 7.96E-60 |
| 1.01E-59 |
| 1.87E-59 |
| 2.24E-59 |
| 2.29E-59 |
| 2.86E-59 |
| 3.69E-59 |
| 6.27E-59 |
| 7.33E-59 |
| 8.98E-59 |
| 1.04E-58 |
| 1.77E-58 |
| 3.47E-58 |
| 3.80E-58 |
| 5.17E-58 |
| 1.04E-57 |
| 1.31E-57 |
| 2.13E-57 |
| 2.75E-57 |
| 2.79E-57 |
| 2.88E-57 |
| 4.27E-57 |
| 4.73E-57 |
| 5.87E-57 |
| 6.43E-57 |
| 9.41E-57 |
| 1.08E-56 |
| 1.08E-56 |
| 1.84E-56 |
| 3.99E-56 |
| 5.30E-56 |
| 6.43E-56 |
| 7.31E-56 |

| |
|---|
| 7.97E-56 |
| 9.99E-56 |
| 1.02E-55 |
| 1.31E-55 |
| 2.04E-55 |
| 2.80E-55 |
| 3.57E-55 |
| 4.00E-55 |
| 4.85E-55 |
| 4.89E-55 |
| 6.62E-55 |
| 6.76E-55 |
| 7.72E-55 |
| 1.16E-54 |
| 1.67E-54 |
| 3.37E-54 |
| 4.66E-54 |
| 6.83E-54 |
| 7.46E-54 |
| 7.68E-54 |
| 1.05E-53 |
| 1.36E-53 |
| 1.75E-53 |
| 2.08E-53 |
| 2.12E-53 |
| 2.19E-53 |
| 2.68E-53 |
| 2.70E-53 |
| 2.98E-53 |
| 4.29E-53 |
| 6.40E-53 |
| 7.10E-53 |
| 1.01E-52 |
| 2.13E-52 |
| 3.40E-52 |
| 3.42E-52 |
| 3.61E-52 |
| 4.98E-52 |
| 5.17E-52 |
| 8.70E-52 |
| 1.01E-51 |
| 1.22E-51 |
| 1.40E-51 |
| 1.40E-51 |
| 1.54E-51 |

1.55E-51
1.68E-51
1.91E-51
1.96E-51
2.26E-51
2.47E-51
3.12E-51
4.33E-51
4.48E-51
6.53E-51
7.98E-51
1.04E-50
1.91E-50
2.39E-50
2.73E-50
2.82E-50
3.32E-50
4.26E-50
4.79E-50
9.65E-50
9.85E-50
9.96E-50
1.21E-49
1.22E-49
1.28E-49
1.33E-49
1.76E-49
2.96E-49
3.30E-49
3.40E-49
6.49E-49
8.82E-49
9.02E-49
1.22E-48
1.34E-48
1.61E-48
1.67E-48
2.49E-48
2.65E-48
4.76E-48
1.06E-47
1.20E-47
1.32E-47
1.63E-47
2.15E-47

2.83E-47
3.12E-47
3.38E-47
3.79E-47
5.24E-47
6.01E-47
7.06E-47
7.18E-47
8.07E-47
1.16E-46
1.27E-46
1.62E-46
1.79E-46
2.34E-46
2.68E-46
3.09E-46
3.17E-46
3.62E-46
4.27E-46
4.32E-46
5.66E-46
7.44E-46
1.56E-45
1.99E-45
2.21E-45
2.57E-45
3.27E-45
3.76E-45
5.62E-45
6.80E-45
6.83E-45
7.02E-45
1.01E-44
1.03E-44
1.07E-44
1.27E-44
1.88E-44
1.94E-44
2.71E-44
3.02E-44
3.06E-44
3.32E-44
3.37E-44
3.93E-44
3.99E-44

4.00E-44
5.81E-44
8.30E-44
8.96E-44
9.87E-44
1.08E-43
1.55E-43
1.79E-43
1.91E-43
2.97E-43
4.05E-43
4.70E-43
4.86E-43
5.22E-43
6.78E-43
7.80E-43
8.78E-43
9.93E-43
1.24E-42
1.56E-42
1.61E-42
1.75E-42
1.83E-42
2.29E-42
2.34E-42
2.46E-42
3.38E-42
3.38E-42
3.70E-42
3.81E-42
4.37E-42
5.02E-42
5.14E-42
5.35E-42
6.45E-42
7.53E-42
8.01E-42
1.03E-41
1.19E-41
1.23E-41
1.25E-41
1.38E-41
1.61E-41
1.66E-41
2.03E-41

2.81E-41
3.08E-41
3.13E-41
3.86E-41
5.02E-41
5.85E-41
7.02E-41
8.56E-41
9.70E-41
1.11E-40
1.14E-40
1.25E-40
1.61E-40
1.63E-40
1.66E-40
2.15E-40
2.55E-40
2.64E-40
2.64E-40
3.07E-40
3.18E-40
4.02E-40
4.32E-40
4.41E-40
4.78E-40
5.95E-40
6.88E-40
6.94E-40
8.02E-40
8.66E-40
8.76E-40
9.20E-40
1.28E-39
1.55E-39
1.61E-39
1.66E-39
2.07E-39
2.29E-39
2.43E-39
3.10E-39
3.79E-39
3.91E-39
5.40E-39
6.14E-39
6.82E-39

7.32E-39
7.37E-39
7.38E-39
7.68E-39
9.81E-39
1.11E-38
1.28E-38
1.52E-38
1.55E-38
1.69E-38
1.76E-38
1.84E-38
2.23E-38
2.53E-38
2.55E-38
2.82E-38
3.02E-38
3.07E-38
4.18E-38
5.09E-38
6.14E-38
6.88E-38
7.36E-38
1.03E-37
1.04E-37
1.06E-37
1.15E-37
1.26E-37
1.40E-37
1.52E-37
1.61E-37
1.75E-37
1.77E-37
1.87E-37
2.43E-37
2.46E-37
2.55E-37
2.64E-37
3.30E-37
4.39E-37
5.08E-37
5.08E-37
6.08E-37
6.08E-37
6.43E-37

7.28E-37
8.09E-37
9.04E-37
1.29E-36
1.34E-36
1.59E-36
1.66E-36
1.85E-36
2.01E-36
2.42E-36
2.58E-36
2.63E-36
2.94E-36
3.44E-36
3.55E-36
3.88E-36
3.90E-36
4.64E-36
5.15E-36
6.34E-36
6.98E-36
7.52E-36
7.60E-36
7.74E-36
8.21E-36
1.09E-35
1.12E-35
1.18E-35
1.20E-35
1.24E-35
1.30E-35
1.40E-35
1.58E-35
2.00E-35
2.29E-35
2.50E-35
2.58E-35
2.59E-35
2.68E-35
3.16E-35
3.24E-35
3.31E-35
3.43E-35
3.59E-35
4.12E-35

4.12E-35
4.47E-35
4.73E-35
4.93E-35
5.07E-35
5.08E-35
5.46E-35
5.75E-35
6.38E-35
6.93E-35
7.86E-35
7.87E-35
8.09E-35
9.19E-35
9.39E-35
1.11E-34
1.19E-34
1.31E-34
1.32E-34
1.43E-34
1.55E-34
1.89E-34
1.99E-34
2.13E-34
2.17E-34
2.20E-34
2.20E-34
2.33E-34
2.59E-34
2.63E-34
2.72E-34
2.82E-34
3.06E-34
3.06E-34
3.06E-34
3.12E-34
3.32E-34
3.48E-34
3.50E-34
3.93E-34
4.08E-34
4.53E-34
4.70E-34
4.85E-34
5.04E-34

7.11E-34
7.80E-34
8.31E-34
9.16E-34
9.49E-34
1.07E-33
1.37E-33
1.38E-33
1.48E-33
1.75E-33
1.93E-33
2.46E-33
2.55E-33
2.91E-33
3.01E-33
3.27E-33
3.42E-33
3.59E-33
3.91E-33
4.18E-33
4.20E-33
4.29E-33
5.21E-33
5.60E-33
5.70E-33
5.79E-33
6.65E-33
7.08E-33
8.18E-33
9.07E-33
9.66E-33
1.35E-32
1.52E-32
1.64E-32
1.74E-32
1.83E-32
1.96E-32
2.11E-32
2.16E-32
2.29E-32
2.37E-32
2.51E-32
2.54E-32
2.55E-32
2.76E-32

3.02E-32
3.30E-32
3.41E-32
3.94E-32
4.02E-32
4.47E-32
5.93E-32
6.03E-32
6.11E-32
6.31E-32
6.43E-32
6.71E-32
6.96E-32
6.98E-32
8.29E-32
8.37E-32
8.60E-32
9.04E-32
9.69E-32
1.08E-31
1.61E-31
1.61E-31
2.17E-31
2.20E-31
3.19E-31
3.27E-31
4.03E-31
4.11E-31
4.68E-31
4.95E-31
5.02E-31
5.34E-31
5.90E-31
6.51E-31
8.09E-31
8.11E-31
8.81E-31
9.36E-31
9.68E-31
1.04E-30
1.09E-30
1.20E-30
1.41E-30
1.49E-30
1.98E-30

| |
|---|
| 2.19E-30 |
| 2.67E-30 |
| 3.02E-30 |
| 3.05E-30 |
| 3.48E-30 |
| 3.63E-30 |
| 3.83E-30 |
| 3.84E-30 |
| 4.02E-30 |
| 4.11E-30 |
| 4.59E-30 |
| 4.60E-30 |
| 5.00E-30 |
| 5.04E-30 |
| 5.55E-30 |
| 6.00E-30 |
| 6.07E-30 |
| 6.11E-30 |
| 6.29E-30 |
| 6.64E-30 |
| 6.91E-30 |
| 8.21E-30 |
| 8.50E-30 |
| 8.85E-30 |
| 9.56E-30 |
| 1.02E-29 |
| 1.06E-29 |
| 1.07E-29 |
| 1.07E-29 |
| 1.17E-29 |
| 1.33E-29 |
| 1.35E-29 |
| 1.41E-29 |
| 1.50E-29 |
| 1.52E-29 |
| 1.53E-29 |
| 1.70E-29 |
| 2.08E-29 |
| 2.18E-29 |
| 2.29E-29 |
| 2.48E-29 |
| 2.93E-29 |
| 2.97E-29 |
| 3.18E-29 |
| 3.22E-29 |

3.33E-29
3.58E-29
3.94E-29
3.99E-29
5.89E-29
6.27E-29
6.71E-29
7.14E-29
7.44E-29
8.28E-29
9.09E-29
9.24E-29
9.73E-29
1.01E-28
1.11E-28
1.11E-28
1.12E-28
1.20E-28
1.38E-28
1.56E-28
1.62E-28
1.90E-28
2.31E-28
2.34E-28
2.63E-28
2.96E-28
3.23E-28
3.89E-28
4.13E-28
4.43E-28
4.63E-28
4.94E-28
5.42E-28
5.75E-28
5.81E-28
6.18E-28
6.43E-28
6.62E-28
7.46E-28
8.09E-28
9.90E-28
9.97E-28
1.00E-27
1.05E-27
1.27E-27

1.28E-27
1.60E-27
1.63E-27
1.65E-27
1.90E-27
1.92E-27
1.93E-27
2.53E-27
2.62E-27
2.77E-27
2.96E-27
3.05E-27
3.10E-27
3.13E-27
3.16E-27
3.22E-27
3.42E-27
3.53E-27
3.83E-27
3.87E-27
3.93E-27
4.04E-27
4.11E-27
4.21E-27
4.21E-27
4.47E-27
4.74E-27
5.01E-27
6.15E-27
6.45E-27
6.62E-27
7.08E-27
7.16E-27
7.25E-27
7.34E-27
7.47E-27
7.58E-27
7.75E-27
7.83E-27
9.33E-27
9.34E-27
9.51E-27
9.66E-27
1.03E-26
1.16E-26

1.16E-26
1.18E-26
1.21E-26
1.28E-26
1.53E-26
1.59E-26
1.62E-26
1.76E-26
1.79E-26
1.86E-26
2.12E-26
2.12E-26
2.15E-26
2.30E-26
2.45E-26
2.53E-26
2.70E-26
2.95E-26
3.55E-26
3.84E-26
4.76E-26
5.15E-26
5.20E-26
5.68E-26
5.74E-26
6.04E-26
6.12E-26
6.70E-26
6.84E-26
6.88E-26
7.21E-26
7.44E-26
7.82E-26
8.37E-26
8.37E-26
9.08E-26
9.97E-26
1.04E-25
1.06E-25
1.10E-25
1.22E-25
1.28E-25
1.32E-25
1.37E-25
1.41E-25

1.42E-25
1.44E-25
1.50E-25
1.82E-25
1.91E-25
1.91E-25
1.98E-25
2.01E-25
2.03E-25
2.05E-25
2.47E-25
2.56E-25
2.57E-25
3.01E-25
3.13E-25
3.16E-25
3.19E-25
3.41E-25
3.58E-25
3.79E-25
3.99E-25
4.23E-25
4.56E-25
5.43E-25
5.59E-25
5.97E-25
6.21E-25
6.62E-25
7.26E-25
7.27E-25
7.89E-25
8.30E-25
8.94E-25
9.75E-25
9.85E-25
1.01E-24
1.13E-24
1.16E-24
1.23E-24
1.30E-24
1.37E-24
1.42E-24
1.44E-24
1.44E-24
1.57E-24

1.62E-24
1.65E-24
1.92E-24
1.98E-24
2.06E-24
2.17E-24
2.19E-24
2.22E-24
2.33E-24
2.38E-24
2.38E-24
2.42E-24
2.42E-24
2.46E-24
2.66E-24
2.76E-24
2.77E-24
2.88E-24
3.02E-24
3.18E-24
3.18E-24
3.24E-24
3.38E-24
3.41E-24
3.41E-24
3.50E-24
3.55E-24
3.56E-24
3.63E-24
3.67E-24
3.89E-24
4.09E-24
4.10E-24
4.22E-24
4.60E-24
4.64E-24
4.90E-24
4.99E-24
5.08E-24
5.50E-24
5.64E-24
5.95E-24
6.01E-24
6.18E-24
6.25E-24

6.47E-24
6.85E-24
7.15E-24
7.28E-24
7.82E-24
7.91E-24
8.19E-24
8.53E-24
8.82E-24
9.11E-24
9.63E-24
9.66E-24
1.03E-23
1.03E-23
1.05E-23
1.09E-23
1.11E-23
1.19E-23
1.24E-23
1.27E-23
1.27E-23
1.31E-23
1.33E-23
1.45E-23
1.49E-23
1.51E-23
1.72E-23
1.78E-23
1.80E-23
1.88E-23
1.93E-23
1.97E-23
1.99E-23
2.08E-23
2.20E-23
2.33E-23
2.53E-23
2.70E-23
2.73E-23
2.98E-23
3.11E-23
3.12E-23
3.27E-23
3.27E-23
3.33E-23

3.37E-23
3.43E-23
3.56E-23
3.59E-23
3.73E-23
3.80E-23
3.84E-23
3.85E-23
3.90E-23
4.01E-23
4.24E-23
4.29E-23
4.46E-23
4.68E-23
4.76E-23
4.82E-23
4.88E-23
5.12E-23
5.18E-23
5.30E-23
5.35E-23
5.54E-23
5.57E-23
5.78E-23
5.93E-23
6.27E-23
6.30E-23
6.35E-23
6.37E-23
6.69E-23
6.96E-23
7.04E-23
7.66E-23
7.76E-23
8.91E-23
9.13E-23
9.31E-23
9.46E-23
9.54E-23
9.76E-23
1.04E-22
1.07E-22
1.08E-22
1.12E-22
1.16E-22

1.16E-22
1.21E-22
1.22E-22
1.28E-22
1.34E-22
1.42E-22
1.42E-22
1.54E-22
1.55E-22
1.61E-22
1.71E-22
1.90E-22
2.15E-22
2.24E-22
2.28E-22
2.33E-22
2.43E-22
2.44E-22
2.49E-22
2.60E-22
2.67E-22
3.01E-22
3.15E-22
3.17E-22
3.26E-22
3.52E-22
3.62E-22
3.65E-22
3.84E-22
3.90E-22
3.94E-22
4.45E-22
5.07E-22
5.17E-22
6.00E-22
6.20E-22
6.22E-22
6.40E-22
6.47E-22
6.73E-22
6.77E-22
7.03E-22
8.10E-22
8.15E-22
8.15E-22

8.27E-22
8.33E-22
8.47E-22
9.31E-22
1.07E-21
1.09E-21
1.16E-21
1.18E-21
1.18E-21
1.18E-21
1.23E-21
1.24E-21
1.26E-21
1.28E-21
1.30E-21
1.30E-21
1.30E-21
1.34E-21
1.34E-21
1.35E-21
1.41E-21
1.43E-21
1.44E-21
1.48E-21
1.50E-21
1.53E-21
1.55E-21
1.58E-21
1.60E-21
1.60E-21
1.83E-21
1.95E-21
2.05E-21
2.07E-21
2.17E-21
2.26E-21
2.37E-21
2.45E-21
2.50E-21
2.57E-21
2.63E-21
2.65E-21
2.69E-21
2.84E-21
2.89E-21

2.92E-21
2.94E-21
2.97E-21
3.08E-21
3.32E-21
3.37E-21
3.39E-21
3.63E-21
3.65E-21
3.69E-21
3.72E-21
3.79E-21
4.14E-21
4.33E-21
4.37E-21
4.43E-21
4.57E-21
4.91E-21
4.97E-21
4.98E-21
4.99E-21
5.00E-21
5.26E-21
5.26E-21
5.65E-21
5.79E-21
5.79E-21
6.42E-21
6.94E-21
7.65E-21
8.05E-21
8.35E-21
8.51E-21
8.95E-21
9.11E-21
9.25E-21
9.99E-21
1.01E-20
1.02E-20
1.03E-20
1.06E-20
1.09E-20
1.10E-20
1.15E-20
1.17E-20

1.18E-20
1.24E-20
1.25E-20
1.32E-20
1.38E-20
1.41E-20
1.43E-20
1.47E-20
1.50E-20
1.50E-20
1.51E-20
1.94E-20
2.02E-20
2.03E-20
2.08E-20
2.12E-20
2.13E-20
2.14E-20
2.18E-20
2.19E-20
2.25E-20
2.25E-20
2.32E-20
2.34E-20
2.46E-20
2.55E-20
2.56E-20
2.82E-20
2.93E-20
2.94E-20
3.07E-20
3.19E-20
3.34E-20
3.53E-20
3.54E-20
3.59E-20
3.86E-20
4.22E-20
4.27E-20
4.31E-20
4.51E-20
4.56E-20
4.62E-20
4.83E-20
5.12E-20

5.15E-20
5.32E-20
5.39E-20
5.45E-20
5.56E-20
5.70E-20
5.79E-20
5.82E-20
5.88E-20
5.94E-20
6.00E-20
6.19E-20
6.23E-20
6.75E-20
6.91E-20
7.03E-20
7.26E-20
7.34E-20
7.44E-20
8.07E-20
8.28E-20
8.60E-20
9.00E-20
9.67E-20
9.93E-20
9.94E-20
1.07E-19
1.10E-19
1.12E-19
1.12E-19
1.15E-19
1.16E-19
1.26E-19
1.28E-19
1.29E-19
1.33E-19
1.34E-19
1.35E-19
1.36E-19
1.37E-19
1.42E-19
1.43E-19
1.47E-19
1.48E-19
1.52E-19

1.53E-19
1.53E-19
1.55E-19
1.55E-19
1.56E-19
1.56E-19
1.63E-19
1.68E-19
1.70E-19
1.75E-19
1.80E-19
1.90E-19
1.91E-19
1.92E-19
2.01E-19
2.11E-19
2.20E-19
2.23E-19
2.32E-19
2.36E-19
2.43E-19
2.55E-19
2.62E-19
2.64E-19
2.65E-19
2.79E-19
2.83E-19
2.88E-19
2.96E-19
3.00E-19
3.00E-19
3.15E-19
3.15E-19
3.36E-19
3.52E-19
3.53E-19
3.74E-19
3.75E-19
3.84E-19
3.94E-19
3.96E-19
3.98E-19
4.02E-19
4.11E-19
4.12E-19

4.12E-19
4.14E-19
4.31E-19
4.60E-19
4.66E-19
4.71E-19
4.78E-19
5.03E-19
5.25E-19
6.33E-19
6.53E-19
6.63E-19
6.64E-19
7.24E-19
7.35E-19
7.49E-19
7.86E-19
7.91E-19
8.87E-19
9.27E-19
9.55E-19
9.60E-19
9.64E-19
9.71E-19
1.10E-18
1.12E-18
1.18E-18
1.19E-18
1.19E-18
1.25E-18
1.27E-18
1.42E-18
1.44E-18
1.45E-18
1.46E-18
1.48E-18
1.50E-18
1.61E-18
1.66E-18
1.74E-18
1.78E-18
1.89E-18
1.98E-18
1.98E-18
2.01E-18

2.05E-18
2.15E-18
2.21E-18
2.21E-18
2.23E-18
2.50E-18
2.79E-18
2.87E-18
2.88E-18
2.97E-18
3.10E-18
3.28E-18
3.28E-18
3.43E-18
3.52E-18
3.60E-18
3.60E-18
3.63E-18
3.82E-18
3.94E-18
4.06E-18
4.19E-18
4.25E-18
4.47E-18
4.51E-18
4.53E-18
4.53E-18
4.67E-18
4.67E-18
4.72E-18
4.79E-18
4.87E-18
4.90E-18
5.21E-18
5.33E-18
5.67E-18
5.68E-18
5.82E-18
5.86E-18
5.96E-18
6.56E-18
6.74E-18
6.90E-18
6.94E-18
7.13E-18

7.18E-18
7.20E-18
7.75E-18
7.75E-18
7.85E-18
7.97E-18
8.44E-18
8.67E-18
8.70E-18
9.44E-18
9.59E-18
9.59E-18
9.89E-18
9.98E-18
1.07E-17
1.11E-17
1.16E-17
1.16E-17
1.18E-17
1.20E-17
1.25E-17
1.26E-17
1.33E-17
1.42E-17
1.44E-17
1.46E-17
1.46E-17
1.50E-17
1.52E-17
1.53E-17
1.56E-17
1.57E-17
1.62E-17
1.66E-17
1.68E-17
1.68E-17
1.76E-17
1.79E-17
1.81E-17
1.83E-17
1.85E-17
1.89E-17
2.03E-17
2.10E-17
2.11E-17

| |
|---|
| 2.23E-17 |
| 2.45E-17 |
| 2.51E-17 |
| 2.51E-17 |
| 2.62E-17 |
| 2.68E-17 |
| 2.70E-17 |
| 2.77E-17 |
| 2.80E-17 |
| 3.05E-17 |
| 3.10E-17 |
| 3.23E-17 |
| 3.38E-17 |
| 3.41E-17 |
| 3.81E-17 |
| 3.85E-17 |
| 3.88E-17 |
| 4.15E-17 |
| 4.33E-17 |
| 4.55E-17 |
| 4.56E-17 |
| 4.76E-17 |
| 4.87E-17 |
| 4.90E-17 |
| 4.94E-17 |
| 5.19E-17 |
| 5.26E-17 |
| 5.41E-17 |
| 5.43E-17 |
| 5.57E-17 |
| 5.75E-17 |
| 5.75E-17 |
| 5.76E-17 |
| 5.83E-17 |
| 6.06E-17 |
| 6.17E-17 |
| 6.33E-17 |
| 6.38E-17 |
| 6.52E-17 |
| 6.52E-17 |
| 6.57E-17 |
| 6.59E-17 |
| 7.35E-17 |
| 7.37E-17 |
| 7.39E-17 |

7.65E-17
7.69E-17
7.95E-17
8.07E-17
8.12E-17
9.14E-17
1.01E-16
1.01E-16
1.05E-16
1.06E-16
1.10E-16
1.10E-16
1.10E-16
1.12E-16
1.14E-16
1.17E-16
1.19E-16
1.20E-16
1.20E-16
1.25E-16
1.27E-16
1.31E-16
1.33E-16
1.35E-16
1.40E-16
1.44E-16
1.48E-16
1.49E-16
1.54E-16
1.54E-16
1.60E-16
1.74E-16
1.75E-16
1.75E-16
1.77E-16
1.84E-16
1.98E-16
1.98E-16
1.98E-16
2.06E-16
2.07E-16
2.18E-16
2.19E-16
2.20E-16
2.20E-16

2.22E-16
2.24E-16
2.27E-16
2.28E-16
2.29E-16
2.39E-16
2.39E-16
2.41E-16
2.41E-16
2.43E-16
2.43E-16
2.48E-16
2.51E-16
2.51E-16
2.56E-16
2.60E-16
2.60E-16
2.66E-16
2.67E-16
2.74E-16
2.86E-16
2.92E-16
2.94E-16
3.07E-16
3.16E-16
3.22E-16
3.31E-16
3.53E-16
3.58E-16
3.78E-16
3.88E-16
3.93E-16
4.18E-16
4.19E-16
4.32E-16
4.37E-16
4.48E-16
4.58E-16
4.62E-16
4.69E-16
4.72E-16
4.80E-16
4.85E-16
5.25E-16
5.32E-16

| |
|---|
| 5.39E-16 |
| 5.51E-16 |
| 5.52E-16 |
| 5.54E-16 |
| 5.76E-16 |
| 5.77E-16 |
| 5.85E-16 |
| 5.91E-16 |
| 5.96E-16 |
| 5.99E-16 |
| 6.96E-16 |
| 7.14E-16 |
| 7.22E-16 |
| 7.39E-16 |
| 7.71E-16 |
| 7.91E-16 |
| 8.09E-16 |
| 8.80E-16 |
| 8.93E-16 |
| 8.93E-16 |
| 8.95E-16 |
| 9.09E-16 |
| 9.26E-16 |
| 9.26E-16 |
| 9.34E-16 |
| 9.51E-16 |
| 9.58E-16 |
| 9.58E-16 |
| 1.02E-15 |
| 1.04E-15 |
| 1.09E-15 |
| 1.14E-15 |
| 1.14E-15 |
| 1.15E-15 |
| 1.21E-15 |
| 1.22E-15 |
| 1.26E-15 |
| 1.29E-15 |
| 1.29E-15 |
| 1.45E-15 |
| 1.61E-15 |
| 1.77E-15 |
| 1.78E-15 |
| 1.78E-15 |
| 1.84E-15 |

1.88E-15
2.13E-15
2.15E-15
2.25E-15
2.28E-15
2.34E-15
2.36E-15
2.36E-15
2.41E-15
2.43E-15
2.47E-15
2.56E-15
2.68E-15
2.70E-15
2.91E-15
2.94E-15
2.95E-15
2.99E-15
3.01E-15
3.06E-15
3.17E-15
3.18E-15
3.20E-15
3.22E-15
3.24E-15
3.27E-15
3.40E-15
3.48E-15
3.63E-15
3.72E-15
3.82E-15
3.82E-15
3.83E-15
3.83E-15
3.87E-15
3.93E-15
3.99E-15
4.02E-15
4.05E-15
4.12E-15
4.28E-15
4.35E-15
4.36E-15
4.39E-15
4.39E-15

4.76E-15
4.83E-15
4.85E-15
4.89E-15
4.91E-15
5.04E-15
5.14E-15
5.15E-15
5.34E-15
5.40E-15
5.42E-15
5.69E-15
5.72E-15
5.77E-15
5.81E-15
5.81E-15
5.89E-15
6.03E-15
6.14E-15
6.91E-15
7.43E-15
7.75E-15
7.83E-15
8.02E-15
8.03E-15
8.06E-15
8.36E-15
8.72E-15
8.73E-15
8.74E-15
8.98E-15
9.01E-15
9.11E-15
9.13E-15
9.21E-15
9.41E-15
9.75E-15
1.01E-14
1.02E-14
1.02E-14
1.02E-14
1.09E-14
1.09E-14
1.12E-14
1.22E-14

1.22E-14
1.25E-14
1.27E-14
1.30E-14
1.35E-14
1.35E-14
1.36E-14
1.39E-14
1.41E-14
1.43E-14
1.55E-14
1.55E-14
1.58E-14
1.60E-14
1.60E-14
1.60E-14
1.61E-14
1.62E-14
1.64E-14
1.65E-14
1.69E-14
1.76E-14
1.81E-14
1.82E-14
1.90E-14
1.90E-14
1.90E-14
1.91E-14
1.92E-14
1.93E-14
1.94E-14
2.05E-14
2.05E-14
2.09E-14
2.16E-14
2.23E-14
2.26E-14
2.26E-14
2.26E-14
2.26E-14
2.27E-14
2.28E-14
2.40E-14
2.43E-14
2.44E-14

2.48E-14
2.49E-14
2.58E-14
2.73E-14
2.78E-14
2.86E-14
2.86E-14
2.91E-14
3.01E-14
3.03E-14
3.04E-14
3.06E-14
3.08E-14
3.23E-14
3.25E-14
3.32E-14
3.34E-14
3.37E-14
3.45E-14
3.51E-14
3.55E-14
3.55E-14
3.61E-14
3.63E-14
3.66E-14
3.68E-14
3.72E-14
3.90E-14
3.90E-14
4.01E-14
4.04E-14
4.07E-14
4.16E-14
4.21E-14
4.22E-14
4.28E-14
4.38E-14
4.39E-14
4.39E-14
4.39E-14
4.45E-14
4.55E-14
4.56E-14
4.78E-14
4.94E-14

| |
|---|
| 4.96E-14 |
| 5.00E-14 |
| 5.05E-14 |
| 5.26E-14 |
| 5.42E-14 |
| 5.47E-14 |
| 5.49E-14 |
| 5.68E-14 |
| 5.70E-14 |
| 5.72E-14 |
| 5.74E-14 |
| 5.79E-14 |
| 6.05E-14 |
| 6.15E-14 |
| 6.26E-14 |
| 6.29E-14 |
| 6.32E-14 |
| 6.40E-14 |
| 6.46E-14 |
| 6.48E-14 |
| 6.72E-14 |
| 6.75E-14 |
| 6.77E-14 |
| 6.81E-14 |
| 6.85E-14 |
| 7.09E-14 |
| 7.17E-14 |
| 7.27E-14 |
| 7.37E-14 |
| 7.46E-14 |
| 7.76E-14 |
| 7.80E-14 |
| 7.94E-14 |
| 7.98E-14 |
| 8.01E-14 |
| 8.03E-14 |
| 8.44E-14 |
| 8.53E-14 |
| 8.85E-14 |
| 8.87E-14 |
| 8.88E-14 |
| 9.10E-14 |
| 9.21E-14 |
| 9.25E-14 |
| 9.34E-14 |

9.43E-14
9.48E-14
9.56E-14
9.62E-14
9.99E-14
1.01E-13
1.02E-13
1.02E-13
1.03E-13
1.12E-13
1.12E-13
1.14E-13
1.15E-13
1.21E-13
1.22E-13
1.24E-13
1.26E-13
1.30E-13
1.31E-13
1.32E-13
1.35E-13
1.39E-13
1.44E-13
1.46E-13
1.46E-13
1.47E-13
1.50E-13
1.52E-13
1.55E-13
1.57E-13
1.58E-13
1.69E-13
1.71E-13
1.72E-13
1.74E-13
1.74E-13
1.74E-13
1.74E-13
1.76E-13
1.77E-13
1.80E-13
1.85E-13
1.85E-13
1.85E-13
1.93E-13

2.05E-13
2.06E-13
2.12E-13
2.12E-13
2.15E-13
2.22E-13
2.32E-13
2.34E-13
2.34E-13
2.39E-13
2.41E-13
2.42E-13
2.48E-13
2.48E-13
2.54E-13
2.56E-13
2.56E-13
2.63E-13
2.63E-13
2.66E-13
2.67E-13
2.73E-13
2.74E-13
2.78E-13
2.79E-13
2.81E-13
2.81E-13
2.93E-13
2.95E-13
2.95E-13
3.01E-13
3.14E-13
3.25E-13
3.31E-13
3.34E-13
3.35E-13
3.36E-13
3.41E-13
3.42E-13
3.46E-13
3.60E-13
3.73E-13
3.74E-13
3.74E-13
3.84E-13

4.01E-13
4.06E-13
4.11E-13
4.13E-13
4.14E-13
4.21E-13
4.36E-13
4.36E-13
4.38E-13
4.62E-13
4.65E-13
4.65E-13
4.70E-13
4.74E-13
4.76E-13
4.83E-13
4.97E-13
5.21E-13
5.28E-13
5.40E-13
5.40E-13
5.43E-13
5.43E-13
5.44E-13
5.47E-13
5.95E-13
5.98E-13
6.02E-13
6.02E-13
6.04E-13
6.12E-13
6.33E-13
6.34E-13
6.36E-13
6.49E-13
6.61E-13
6.67E-13
6.72E-13
6.87E-13
6.93E-13
6.93E-13
6.94E-13
6.99E-13
7.05E-13
7.21E-13

7.43E-13
7.45E-13
7.47E-13
7.47E-13
8.17E-13
8.52E-13
8.59E-13
8.66E-13
8.78E-13
8.83E-13
8.95E-13
9.22E-13
9.45E-13
9.48E-13
9.66E-13
9.76E-13
9.78E-13
9.81E-13
9.85E-13
9.87E-13
1.01E-12
1.06E-12
1.06E-12
1.10E-12
1.10E-12
1.13E-12
1.13E-12
1.16E-12
1.16E-12
1.17E-12
1.18E-12
1.19E-12
1.24E-12
1.24E-12
1.28E-12
1.28E-12
1.28E-12
1.28E-12
1.30E-12
1.30E-12
1.30E-12
1.33E-12
1.34E-12
1.35E-12
1.38E-12

1.38E-12
1.39E-12
1.43E-12
1.43E-12
1.46E-12
1.47E-12
1.47E-12
1.53E-12
1.54E-12
1.55E-12
1.56E-12
1.58E-12
1.58E-12
1.62E-12
1.70E-12
1.71E-12
1.72E-12
1.73E-12
1.75E-12
1.83E-12
1.84E-12
1.88E-12
1.90E-12
1.94E-12
1.96E-12
1.97E-12
1.97E-12
2.03E-12
2.07E-12
2.07E-12
2.10E-12
2.11E-12
2.12E-12
2.22E-12
2.23E-12
2.25E-12
2.27E-12
2.34E-12
2.35E-12
2.37E-12
2.37E-12
2.37E-12
2.41E-12
2.43E-12
2.58E-12

2.59E-12
2.61E-12
2.61E-12
2.66E-12
2.70E-12
2.76E-12
2.78E-12
2.78E-12
2.79E-12
2.89E-12
2.91E-12
2.92E-12
2.96E-12
2.96E-12
3.01E-12
3.04E-12
3.11E-12
3.11E-12
3.15E-12
3.15E-12
3.23E-12
3.31E-12
3.35E-12
3.37E-12
3.38E-12
3.49E-12
3.60E-12
3.60E-12
3.63E-12
3.78E-12
3.81E-12
3.84E-12
3.88E-12
3.92E-12
3.93E-12
3.98E-12
4.06E-12
4.14E-12
4.34E-12
4.34E-12
4.36E-12
4.49E-12
4.49E-12
4.49E-12
4.63E-12

4.63E-12
4.68E-12
4.72E-12
4.88E-12
4.88E-12
4.88E-12
4.93E-12
4.96E-12
5.02E-12
5.14E-12
5.31E-12
5.34E-12
5.34E-12
5.38E-12
5.51E-12
5.53E-12
5.56E-12
5.65E-12
5.66E-12
5.68E-12
5.70E-12
5.81E-12
5.89E-12
6.06E-12
6.08E-12
6.13E-12
6.22E-12
6.27E-12
6.38E-12
6.60E-12
6.60E-12
6.64E-12
6.74E-12
6.81E-12
6.82E-12
7.13E-12
7.31E-12
7.33E-12
7.48E-12
7.55E-12
7.55E-12
7.62E-12
7.66E-12
7.75E-12
7.75E-12

7.85E-12
7.87E-12
7.88E-12
7.92E-12
7.95E-12
7.98E-12
8.07E-12
8.10E-12
8.14E-12
8.18E-12
8.34E-12
8.37E-12
8.54E-12
8.61E-12
8.80E-12
8.89E-12
8.95E-12
8.95E-12
9.05E-12
9.26E-12
9.38E-12
9.93E-12
1.00E-11
1.01E-11
1.03E-11
1.04E-11
1.05E-11
1.05E-11
1.08E-11
1.13E-11
1.13E-11
1.14E-11
1.16E-11
1.19E-11
1.21E-11
1.21E-11
1.23E-11
1.23E-11
1.28E-11
1.30E-11
1.30E-11
1.31E-11
1.31E-11
1.31E-11
1.34E-11

1.35E-11
1.42E-11
1.47E-11
1.48E-11
1.49E-11
1.50E-11
1.50E-11
1.50E-11
1.50E-11
1.52E-11
1.53E-11
1.53E-11
1.53E-11
1.55E-11
1.58E-11
1.59E-11
1.59E-11
1.60E-11
1.60E-11
1.60E-11
1.61E-11
1.62E-11
1.63E-11
1.63E-11
1.64E-11
1.64E-11
1.67E-11
1.68E-11
1.77E-11
1.78E-11
1.84E-11
1.84E-11
1.87E-11
1.87E-11
1.88E-11
1.89E-11
1.93E-11
1.95E-11
2.01E-11
2.02E-11
2.02E-11
2.03E-11
2.04E-11
2.05E-11
2.12E-11

2.19E-11
2.20E-11
2.20E-11
2.22E-11
2.26E-11
2.26E-11
2.37E-11
2.42E-11
2.57E-11
2.58E-11
2.61E-11
2.61E-11
2.61E-11
2.68E-11
2.71E-11
2.72E-11
2.79E-11
2.85E-11
2.93E-11
3.00E-11
3.01E-11
3.17E-11
3.18E-11
3.18E-11
3.26E-11
3.28E-11
3.32E-11
3.37E-11
3.39E-11
3.42E-11
3.52E-11
3.54E-11
3.58E-11
3.58E-11
3.59E-11
3.59E-11
3.78E-11
3.78E-11
3.89E-11
3.93E-11
3.96E-11
3.97E-11
3.98E-11
4.02E-11
4.17E-11

4.18E-11
4.21E-11
4.23E-11
4.28E-11
4.31E-11
4.34E-11
4.49E-11
4.61E-11
4.64E-11
4.68E-11
4.70E-11
4.81E-11
4.89E-11
4.98E-11
5.02E-11
5.10E-11
5.10E-11
5.13E-11
5.19E-11
5.22E-11
5.27E-11
5.28E-11
5.48E-11
5.66E-11
5.74E-11
5.82E-11
5.91E-11
5.91E-11
5.92E-11
6.07E-11
6.19E-11
6.21E-11
6.26E-11
6.39E-11
6.43E-11
6.45E-11
6.45E-11
6.48E-11
6.54E-11
6.55E-11
6.62E-11
6.62E-11
6.62E-11
6.62E-11
6.65E-11

6.69E-11
6.89E-11
6.95E-11
7.10E-11
7.17E-11
7.34E-11
7.43E-11
7.53E-11
7.62E-11
7.72E-11
7.78E-11
7.85E-11
7.90E-11
8.11E-11
8.12E-11
8.42E-11
8.78E-11
8.84E-11
8.88E-11
8.93E-11
8.98E-11
9.35E-11
9.35E-11
9.46E-11
9.52E-11
9.54E-11
9.55E-11
9.55E-11
9.74E-11
9.94E-11
1.00E-10
1.01E-10
1.03E-10
1.03E-10
1.03E-10
1.05E-10
1.06E-10
1.06E-10
1.07E-10
1.07E-10
1.09E-10
1.09E-10
1.09E-10
1.12E-10
1.18E-10

1.18E-10
1.19E-10
1.22E-10
1.23E-10
1.23E-10
1.24E-10
1.24E-10
1.25E-10
1.25E-10
1.25E-10
1.25E-10
1.30E-10
1.30E-10
1.33E-10
1.34E-10
1.37E-10
1.38E-10
1.41E-10
1.43E-10
1.45E-10
1.45E-10
1.47E-10
1.47E-10
1.47E-10
1.49E-10
1.55E-10
1.57E-10
1.57E-10
1.57E-10
1.61E-10
1.64E-10
1.64E-10
1.64E-10
1.65E-10
1.66E-10
1.68E-10
1.70E-10
1.71E-10
1.71E-10
1.73E-10
1.75E-10
1.75E-10
1.78E-10
1.79E-10
1.81E-10

1.81E-10
1.83E-10
1.84E-10
1.85E-10
1.85E-10
1.86E-10
1.87E-10
1.88E-10
1.88E-10
1.92E-10
1.96E-10
1.96E-10
1.96E-10
1.96E-10
2.00E-10
2.05E-10
2.08E-10
2.11E-10
2.13E-10
2.25E-10
2.28E-10
2.29E-10
2.31E-10
2.33E-10
2.35E-10
2.36E-10
2.37E-10
2.37E-10
2.39E-10
2.39E-10
2.40E-10
2.40E-10
2.45E-10
2.45E-10
2.46E-10
2.50E-10
2.51E-10
2.52E-10
2.52E-10
2.56E-10
2.58E-10
2.60E-10
2.60E-10
2.61E-10
2.62E-10

2.62E-10
2.62E-10
2.66E-10
2.69E-10
2.72E-10
2.75E-10
2.79E-10
2.79E-10
2.80E-10
2.90E-10
2.91E-10
3.06E-10
3.09E-10
3.10E-10
3.15E-10
3.16E-10
3.17E-10
3.17E-10
3.18E-10
3.22E-10
3.23E-10
3.26E-10
3.26E-10
3.28E-10
3.30E-10
3.30E-10
3.34E-10
3.54E-10
3.58E-10
3.60E-10
3.68E-10
3.69E-10
3.70E-10
3.78E-10
3.78E-10
3.81E-10
3.85E-10
3.89E-10
3.98E-10
4.06E-10
4.14E-10
4.17E-10
4.27E-10
4.35E-10
4.45E-10

4.76E-10
4.79E-10
4.82E-10
4.84E-10
4.84E-10
4.84E-10
4.95E-10
4.98E-10
4.99E-10
5.02E-10
5.03E-10
5.07E-10
5.17E-10
5.24E-10
5.26E-10
5.33E-10
5.34E-10
5.38E-10
5.44E-10
5.47E-10
5.57E-10
5.59E-10
5.67E-10
5.76E-10
5.82E-10
5.83E-10
5.90E-10
5.98E-10
6.05E-10
6.17E-10
6.18E-10
6.22E-10
6.27E-10
6.32E-10
6.36E-10
6.36E-10
6.37E-10
6.38E-10
6.68E-10
6.89E-10
6.93E-10
7.00E-10
7.01E-10
7.03E-10
7.05E-10

7.25E-10
7.44E-10
7.53E-10
7.55E-10
7.55E-10
7.58E-10
7.58E-10
7.60E-10
7.63E-10
7.64E-10
7.68E-10
7.86E-10
8.09E-10
8.20E-10
8.29E-10
8.30E-10
8.37E-10
8.40E-10
8.42E-10
8.44E-10
8.48E-10
8.75E-10
8.81E-10
8.82E-10
8.88E-10
8.91E-10
8.94E-10
9.00E-10
9.08E-10
9.08E-10
9.13E-10
9.16E-10
9.25E-10
9.27E-10
9.41E-10
9.44E-10
9.47E-10
9.54E-10
9.55E-10
9.73E-10
9.81E-10
9.92E-10
9.92E-10
9.96E-10
1.00E-09

| |
|---|
| 1.01E-09 |
| 1.02E-09 |
| 1.02E-09 |
| 1.03E-09 |
| 1.03E-09 |
| 1.03E-09 |
| 1.04E-09 |
| 1.05E-09 |
| 1.07E-09 |
| 1.09E-09 |
| 1.09E-09 |
| 1.10E-09 |
| 1.14E-09 |
| 1.16E-09 |
| 1.19E-09 |
| 1.19E-09 |
| 1.20E-09 |
| 1.23E-09 |
| 1.24E-09 |
| 1.25E-09 |
| 1.25E-09 |
| 1.25E-09 |
| 1.26E-09 |
| 1.27E-09 |
| 1.27E-09 |
| 1.27E-09 |
| 1.30E-09 |
| 1.31E-09 |
| 1.31E-09 |
| 1.32E-09 |
| 1.33E-09 |
| 1.34E-09 |
| 1.35E-09 |
| 1.36E-09 |
| 1.36E-09 |
| 1.38E-09 |
| 1.38E-09 |
| 1.38E-09 |
| 1.40E-09 |
| 1.41E-09 |
| 1.44E-09 |
| 1.46E-09 |
| 1.46E-09 |
| 1.49E-09 |
| 1.52E-09 |

1.52E-09
1.52E-09
1.53E-09
1.57E-09
1.57E-09
1.57E-09
1.58E-09
1.63E-09
1.65E-09
1.67E-09
1.71E-09
1.72E-09
1.73E-09
1.76E-09
1.78E-09
1.82E-09
1.82E-09
1.82E-09
1.82E-09
1.84E-09
1.88E-09
1.94E-09
1.97E-09
1.98E-09
1.98E-09
2.01E-09
2.03E-09
2.07E-09
2.11E-09
2.12E-09
2.14E-09
2.16E-09
2.16E-09
2.17E-09
2.22E-09
2.23E-09
2.27E-09
2.27E-09
2.28E-09
2.28E-09
2.30E-09
2.31E-09
2.35E-09
2.37E-09
2.37E-09

2.38E-09
2.41E-09
2.43E-09
2.45E-09
2.50E-09
2.52E-09
2.52E-09
2.54E-09
2.54E-09
2.56E-09
2.58E-09
2.59E-09
2.59E-09
2.60E-09
2.66E-09
2.75E-09
2.75E-09
2.76E-09
2.77E-09
2.79E-09
2.79E-09
2.81E-09
2.83E-09
2.83E-09
2.84E-09
2.89E-09
2.90E-09
2.97E-09
2.98E-09
3.01E-09
3.03E-09
3.03E-09
3.06E-09
3.07E-09
3.08E-09
3.09E-09
3.09E-09
3.09E-09
3.09E-09
3.11E-09
3.14E-09
3.14E-09
3.17E-09
3.22E-09
3.25E-09

3.25E-09
3.27E-09
3.35E-09
3.37E-09
3.38E-09
3.40E-09
3.51E-09
3.53E-09
3.55E-09
3.59E-09
3.60E-09
3.60E-09
3.60E-09
3.61E-09
3.64E-09
3.66E-09
3.66E-09
3.66E-09
3.72E-09
3.77E-09
3.79E-09
3.79E-09
3.82E-09
3.86E-09
3.86E-09
3.86E-09
3.87E-09
3.88E-09
3.93E-09
3.96E-09
4.06E-09
4.10E-09
4.19E-09
4.21E-09
4.22E-09
4.37E-09
4.43E-09
4.46E-09
4.53E-09
4.60E-09
4.60E-09
4.73E-09
4.76E-09
4.82E-09
4.85E-09

4.85E-09
4.91E-09
4.91E-09
4.99E-09
5.02E-09
5.04E-09
5.04E-09
5.05E-09
5.05E-09
5.08E-09
5.15E-09
5.36E-09
5.54E-09
5.54E-09
5.60E-09
5.63E-09
5.68E-09
5.68E-09
5.71E-09
5.72E-09
5.85E-09
5.85E-09
5.87E-09
5.87E-09
5.89E-09
5.95E-09
6.01E-09
6.05E-09
6.19E-09
6.21E-09
6.21E-09
6.26E-09
6.27E-09
6.46E-09
6.49E-09
6.53E-09
6.60E-09
6.63E-09
6.75E-09
6.80E-09
6.81E-09
6.82E-09
6.87E-09
6.89E-09
6.89E-09

6.97E-09
7.07E-09
7.21E-09
7.22E-09
7.39E-09
7.44E-09
7.52E-09
7.53E-09
7.72E-09
7.84E-09
7.88E-09
7.93E-09
8.29E-09
8.37E-09
8.37E-09
8.40E-09
8.42E-09
8.58E-09
8.58E-09
8.59E-09
8.60E-09
8.68E-09
8.87E-09
8.91E-09
9.13E-09
9.15E-09
9.35E-09
9.37E-09
9.40E-09
9.48E-09
9.54E-09
9.57E-09
9.62E-09
9.72E-09
9.93E-09
1.00E-08
1.01E-08
1.01E-08
1.02E-08
1.02E-08
1.03E-08
1.05E-08
1.05E-08
1.05E-08
1.07E-08

1.07E-08
1.08E-08
1.09E-08
1.09E-08
1.09E-08
1.11E-08
1.12E-08
1.12E-08
1.12E-08
1.13E-08
1.14E-08
1.14E-08
1.14E-08
1.14E-08
1.14E-08
1.15E-08
1.15E-08
1.15E-08
1.15E-08
1.16E-08
1.17E-08
1.18E-08
1.18E-08
1.18E-08
1.19E-08
1.22E-08
1.22E-08
1.23E-08
1.26E-08
1.26E-08
1.27E-08
1.27E-08
1.28E-08
1.29E-08
1.29E-08
1.30E-08
1.31E-08
1.32E-08
1.33E-08
1.34E-08
1.34E-08
1.35E-08
1.35E-08
1.36E-08
1.38E-08

| |
|---|
| 1.38E-08 |
| 1.40E-08 |
| 1.41E-08 |
| 1.41E-08 |
| 1.42E-08 |
| 1.45E-08 |
| 1.45E-08 |
| 1.45E-08 |
| 1.47E-08 |
| 1.48E-08 |
| 1.48E-08 |
| 1.49E-08 |
| 1.50E-08 |
| 1.51E-08 |
| 1.56E-08 |
| 1.58E-08 |
| 1.59E-08 |
| 1.60E-08 |
| 1.60E-08 |
| 1.61E-08 |
| 1.63E-08 |
| 1.63E-08 |
| 1.64E-08 |
| 1.64E-08 |
| 1.64E-08 |
| 1.67E-08 |
| 1.68E-08 |
| 1.70E-08 |
| 1.70E-08 |
| 1.72E-08 |
| 1.74E-08 |
| 1.74E-08 |
| 1.75E-08 |
| 1.78E-08 |
| 1.78E-08 |
| 1.80E-08 |
| 1.81E-08 |
| 1.81E-08 |
| 1.82E-08 |
| 1.82E-08 |
| 1.85E-08 |
| 1.87E-08 |
| 1.87E-08 |
| 1.89E-08 |
| 1.92E-08 |

1.93E-08
1.97E-08
1.99E-08
2.01E-08
2.04E-08
2.06E-08
2.07E-08
2.07E-08
2.07E-08
2.08E-08
2.12E-08
2.13E-08
2.15E-08
2.15E-08
2.16E-08
2.18E-08
2.21E-08
2.23E-08
2.30E-08
2.30E-08
2.35E-08
2.35E-08
2.35E-08
2.35E-08
2.37E-08
2.37E-08
2.38E-08
2.38E-08
2.39E-08
2.41E-08
2.43E-08
2.47E-08
2.53E-08
2.56E-08
2.57E-08
2.60E-08
2.62E-08
2.62E-08
2.64E-08
2.71E-08
2.72E-08
2.72E-08
2.73E-08
2.76E-08
2.79E-08

2.88E-08
2.88E-08
2.90E-08
2.90E-08
2.90E-08
2.92E-08
2.98E-08
3.03E-08
3.05E-08
3.05E-08
3.09E-08
3.10E-08
3.10E-08
3.13E-08
3.15E-08
3.17E-08
3.17E-08
3.20E-08
3.21E-08
3.21E-08
3.22E-08
3.25E-08
3.26E-08
3.27E-08
3.42E-08
3.44E-08
3.46E-08
3.48E-08
3.48E-08
3.49E-08
3.50E-08
3.53E-08
3.55E-08
3.60E-08
3.60E-08
3.66E-08
3.67E-08
3.73E-08
3.78E-08
3.81E-08
3.82E-08
3.83E-08
3.87E-08
3.87E-08
3.96E-08

3.98E-08
4.02E-08
4.05E-08
4.07E-08
4.17E-08
4.17E-08
4.19E-08
4.19E-08
4.21E-08
4.25E-08
4.27E-08
4.32E-08
4.32E-08
4.32E-08
4.34E-08
4.36E-08
4.36E-08
4.40E-08
4.40E-08
4.41E-08
4.55E-08
4.56E-08
4.60E-08
4.60E-08
4.62E-08
4.63E-08
4.64E-08
4.67E-08
4.74E-08
4.80E-08
4.93E-08
4.93E-08
4.96E-08
4.97E-08
4.98E-08
5.04E-08
5.11E-08
5.11E-08
5.14E-08
5.17E-08
5.18E-08
5.19E-08
5.24E-08
5.27E-08
5.32E-08

5.36E-08
5.49E-08
5.51E-08
5.57E-08
5.60E-08
5.67E-08
5.72E-08
5.85E-08
5.87E-08
5.97E-08
5.97E-08
6.02E-08
6.03E-08
6.06E-08
6.08E-08
6.09E-08
6.12E-08
6.13E-08
6.14E-08
6.17E-08
6.20E-08
6.29E-08
6.31E-08
6.37E-08
6.40E-08
6.41E-08
6.50E-08
6.55E-08
6.62E-08
6.79E-08
6.80E-08
6.91E-08
6.92E-08
6.94E-08
7.03E-08
7.04E-08
7.05E-08
7.07E-08
7.15E-08
7.20E-08
7.24E-08
7.32E-08
7.47E-08
7.49E-08
7.54E-08

7.55E-08
7.55E-08
7.58E-08
7.66E-08
7.75E-08
7.83E-08
8.11E-08
8.11E-08
8.14E-08
8.20E-08
8.39E-08
8.41E-08
8.51E-08
8.58E-08
8.58E-08
8.58E-08
8.59E-08
8.61E-08
8.63E-08
8.67E-08
8.80E-08
8.90E-08
9.10E-08
9.13E-08
9.28E-08
9.28E-08
9.29E-08
9.30E-08
9.41E-08
9.45E-08
9.55E-08
9.72E-08
9.83E-08
9.92E-08
9.95E-08
1.01E-07
1.01E-07
1.03E-07
1.04E-07
1.05E-07
1.05E-07
1.05E-07
1.05E-07
1.06E-07
1.06E-07

1.07E-07
1.07E-07
1.09E-07
1.10E-07
1.10E-07
1.10E-07
1.11E-07
1.11E-07
1.11E-07
1.11E-07
1.13E-07
1.13E-07
1.16E-07
1.16E-07
1.17E-07
1.18E-07
1.18E-07
1.18E-07
1.19E-07
1.20E-07
1.21E-07
1.22E-07
1.22E-07
1.25E-07
1.25E-07
1.26E-07
1.26E-07
1.26E-07
1.29E-07
1.29E-07
1.29E-07
1.32E-07
1.33E-07
1.33E-07
1.33E-07
1.35E-07
1.36E-07
1.36E-07
1.37E-07
1.38E-07
1.38E-07
1.39E-07
1.39E-07
1.39E-07
1.41E-07

1.42E-07
1.43E-07
1.45E-07
1.45E-07
1.45E-07
1.47E-07
1.49E-07
1.50E-07
1.51E-07
1.51E-07
1.51E-07
1.51E-07
1.51E-07
1.52E-07
1.52E-07
1.53E-07
1.54E-07
1.54E-07
1.54E-07
1.55E-07
1.55E-07
1.55E-07
1.55E-07
1.56E-07
1.58E-07
1.58E-07
1.64E-07
1.64E-07
1.64E-07
1.65E-07
1.65E-07
1.66E-07
1.68E-07
1.70E-07
1.71E-07
1.71E-07
1.74E-07
1.74E-07
1.74E-07
1.75E-07
1.75E-07
1.79E-07
1.81E-07
1.81E-07
1.81E-07

1.83E-07
1.87E-07
1.87E-07
1.90E-07
1.92E-07
1.93E-07
1.93E-07
1.96E-07
1.96E-07
1.98E-07
2.01E-07
2.02E-07
2.04E-07
2.06E-07
2.06E-07
2.07E-07
2.08E-07
2.08E-07
2.13E-07
2.13E-07
2.17E-07
2.19E-07
2.19E-07
2.19E-07
2.20E-07
2.20E-07
2.26E-07
2.27E-07
2.27E-07
2.27E-07
2.28E-07
2.28E-07
2.29E-07
2.29E-07
2.29E-07
2.29E-07
2.30E-07
2.31E-07
2.33E-07
2.42E-07
2.45E-07
2.45E-07
2.45E-07
2.45E-07
2.48E-07

2.50E-07
2.53E-07
2.53E-07
2.55E-07
2.59E-07
2.60E-07
2.61E-07
2.61E-07
2.61E-07
2.61E-07
2.61E-07
2.61E-07
2.65E-07
2.67E-07
2.67E-07
2.67E-07
2.69E-07
2.70E-07
2.71E-07
2.73E-07
2.73E-07
2.76E-07
2.78E-07
2.79E-07
2.79E-07
2.80E-07
2.81E-07
2.81E-07
2.83E-07
2.84E-07
2.85E-07
2.85E-07
2.86E-07
2.87E-07
2.88E-07
2.94E-07
3.00E-07
3.03E-07
3.03E-07
3.10E-07
3.12E-07
3.13E-07
3.16E-07
3.17E-07
3.17E-07

3.19E-07
3.22E-07
3.25E-07
3.30E-07
3.31E-07
3.32E-07
3.35E-07
3.37E-07
3.37E-07
3.39E-07
3.44E-07
3.44E-07
3.46E-07
3.48E-07
3.48E-07
3.52E-07
3.54E-07
3.55E-07
3.56E-07
3.58E-07
3.59E-07
3.61E-07
3.66E-07
3.69E-07
3.70E-07
3.73E-07
3.74E-07
3.75E-07
3.77E-07
3.77E-07
3.82E-07
3.82E-07
3.83E-07
3.83E-07
3.87E-07
3.89E-07
3.90E-07
3.92E-07
3.96E-07
3.98E-07
3.98E-07
4.00E-07
4.01E-07
4.02E-07
4.07E-07

4.09E-07
4.09E-07
4.10E-07
4.18E-07
4.18E-07
4.21E-07
4.29E-07
4.36E-07
4.36E-07
4.40E-07
4.42E-07
4.45E-07
4.49E-07
4.49E-07
4.50E-07
4.50E-07
4.51E-07
4.52E-07
4.65E-07
4.66E-07
4.71E-07
4.77E-07
4.78E-07
4.84E-07
4.84E-07
4.90E-07
4.91E-07
4.91E-07
4.94E-07
5.04E-07
5.05E-07
5.07E-07
5.08E-07
5.11E-07
5.11E-07
5.12E-07
5.22E-07
5.23E-07
5.24E-07
5.27E-07
5.30E-07
5.58E-07
5.59E-07
5.60E-07
5.61E-07

5.62E-07
5.66E-07
5.75E-07
5.86E-07
5.86E-07
5.87E-07
5.90E-07
5.92E-07
6.05E-07
6.06E-07
6.07E-07
6.08E-07
6.08E-07
6.10E-07
6.14E-07
6.15E-07
6.16E-07
6.22E-07
6.23E-07
6.29E-07
6.32E-07
6.32E-07
6.35E-07
6.41E-07
6.45E-07
6.48E-07
6.52E-07
6.53E-07
6.61E-07
6.62E-07
6.64E-07
6.64E-07
6.68E-07
6.69E-07
6.79E-07
6.80E-07
6.80E-07
6.83E-07
6.83E-07
6.84E-07
6.96E-07
7.02E-07
7.23E-07
7.30E-07
7.31E-07

7.31E-07
7.40E-07
7.43E-07
7.46E-07
7.57E-07
7.59E-07
7.64E-07
7.65E-07
7.67E-07
7.75E-07
7.82E-07
7.85E-07
7.91E-07
7.92E-07
7.94E-07
7.96E-07
8.05E-07
8.08E-07
8.24E-07
8.33E-07
8.40E-07
8.41E-07
8.42E-07
8.43E-07
8.44E-07
8.46E-07
8.47E-07
8.47E-07
8.50E-07
8.57E-07
8.64E-07
8.65E-07
8.69E-07
8.72E-07
8.73E-07
8.75E-07
8.93E-07
9.00E-07
9.11E-07
9.35E-07
9.35E-07
9.40E-07
9.49E-07
9.52E-07
9.61E-07

9.65E-07
9.82E-07
9.86E-07
9.86E-07
9.95E-07
1.00E-06
1.01E-06
1.02E-06
1.03E-06
1.04E-06
1.05E-06
1.06E-06
1.07E-06
1.08E-06
1.08E-06
1.08E-06
1.08E-06
1.09E-06
1.09E-06
1.11E-06
1.11E-06
1.12E-06
1.12E-06
1.12E-06
1.14E-06
1.14E-06
1.15E-06
1.15E-06
1.16E-06
1.16E-06
1.16E-06
1.16E-06
1.17E-06
1.17E-06
1.20E-06
1.20E-06
1.21E-06
1.21E-06
1.22E-06
1.23E-06
1.23E-06
1.23E-06
1.24E-06
1.25E-06
1.25E-06

1.28E-06
1.29E-06
1.30E-06
1.31E-06
1.31E-06
1.32E-06
1.32E-06
1.34E-06
1.34E-06
1.34E-06
1.34E-06
1.34E-06
1.35E-06
1.35E-06
1.36E-06
1.36E-06
1.36E-06
1.37E-06
1.38E-06
1.38E-06
1.39E-06
1.40E-06
1.40E-06
1.41E-06
1.44E-06
1.46E-06
1.47E-06
1.47E-06
1.47E-06
1.50E-06
1.50E-06
1.51E-06
1.51E-06
1.52E-06
1.53E-06
1.54E-06
1.54E-06
1.55E-06
1.56E-06
1.58E-06
1.58E-06
1.59E-06
1.60E-06
1.61E-06
1.62E-06

1.62E-06
1.62E-06
1.62E-06
1.63E-06
1.63E-06
1.63E-06
1.63E-06
1.70E-06
1.70E-06
1.70E-06
1.71E-06
1.71E-06
1.74E-06
1.75E-06
1.75E-06
1.75E-06
1.75E-06
1.76E-06
1.77E-06
1.77E-06
1.77E-06
1.78E-06
1.80E-06
1.81E-06
1.81E-06
1.82E-06
1.83E-06
1.84E-06
1.85E-06
1.87E-06
1.87E-06
1.87E-06
1.87E-06
1.89E-06
1.91E-06
1.94E-06
1.99E-06
2.00E-06
2.01E-06
2.03E-06
2.07E-06
2.07E-06
2.09E-06
2.09E-06
2.10E-06

| |
|---|
| 2.11E-06 |
| 2.14E-06 |
| 2.16E-06 |
| 2.16E-06 |
| 2.16E-06 |
| 2.19E-06 |
| 2.19E-06 |
| 2.19E-06 |
| 2.20E-06 |
| 2.20E-06 |
| 2.20E-06 |
| 2.21E-06 |
| 2.23E-06 |
| 2.24E-06 |
| 2.25E-06 |
| 2.26E-06 |
| 2.29E-06 |
| 2.30E-06 |
| 2.31E-06 |
| 2.31E-06 |
| 2.31E-06 |
| 2.31E-06 |
| 2.33E-06 |
| 2.34E-06 |
| 2.36E-06 |
| 2.36E-06 |
| 2.36E-06 |
| 2.39E-06 |
| 2.41E-06 |
| 2.41E-06 |
| 2.42E-06 |
| 2.43E-06 |
| 2.44E-06 |
| 2.44E-06 |
| 2.46E-06 |
| 2.47E-06 |
| 2.48E-06 |
| 2.49E-06 |
| 2.49E-06 |
| 2.50E-06 |
| 2.50E-06 |
| 2.50E-06 |
| 2.50E-06 |
| 2.51E-06 |
| 2.52E-06 |

2.52E-06
2.52E-06
2.54E-06
2.55E-06
2.56E-06
2.57E-06
2.58E-06
2.59E-06
2.60E-06
2.62E-06
2.62E-06
2.63E-06
2.64E-06
2.64E-06
2.64E-06
2.66E-06
2.66E-06
2.67E-06
2.69E-06
2.70E-06
2.70E-06
2.72E-06
2.75E-06
2.75E-06
2.77E-06
2.78E-06
2.78E-06
2.82E-06
2.83E-06
2.83E-06
2.83E-06
2.83E-06
2.85E-06
2.86E-06
2.88E-06
2.91E-06
2.93E-06
2.94E-06
2.95E-06
2.99E-06
3.00E-06
3.00E-06
3.00E-06
3.01E-06
3.01E-06

3.06E-06
3.06E-06
3.07E-06
3.09E-06
3.09E-06
3.11E-06
3.12E-06
3.12E-06
3.13E-06
3.14E-06
3.14E-06
3.15E-06
3.15E-06
3.16E-06
3.17E-06
3.17E-06
3.21E-06
3.24E-06
3.24E-06
3.26E-06
3.28E-06
3.28E-06
3.30E-06
3.30E-06
3.32E-06
3.38E-06
3.38E-06
3.43E-06
3.43E-06
3.45E-06
3.50E-06
3.51E-06
3.53E-06
3.55E-06
3.56E-06
3.58E-06
3.61E-06
3.61E-06
3.62E-06
3.65E-06
3.65E-06
3.68E-06
3.69E-06
3.71E-06
3.75E-06

3.75E-06
3.76E-06
3.77E-06
3.80E-06
3.80E-06
3.82E-06
3.82E-06
3.83E-06
3.83E-06
3.84E-06
3.87E-06
3.87E-06
3.89E-06
3.90E-06
3.91E-06
3.93E-06
3.94E-06
3.95E-06
3.95E-06
3.97E-06
3.98E-06
3.99E-06
3.99E-06
4.02E-06
4.03E-06
4.03E-06
4.04E-06
4.06E-06
4.08E-06
4.10E-06
4.12E-06
4.12E-06
4.15E-06
4.15E-06
4.17E-06
4.20E-06
4.21E-06
4.21E-06
4.23E-06
4.25E-06
4.31E-06
4.32E-06
4.32E-06
4.38E-06
4.40E-06

4.40E-06
4.43E-06
4.46E-06
4.46E-06
4.55E-06
4.59E-06
4.61E-06
4.64E-06
4.65E-06
4.69E-06
4.71E-06
4.73E-06
4.73E-06
4.74E-06
4.75E-06
4.81E-06
4.84E-06
4.84E-06
4.86E-06
4.88E-06
4.89E-06
4.89E-06
4.92E-06
4.94E-06
4.94E-06
4.95E-06
4.95E-06
4.98E-06
5.01E-06
5.07E-06
5.07E-06
5.08E-06
5.09E-06
5.11E-06
5.13E-06
5.14E-06
5.17E-06
5.20E-06
5.23E-06
5.23E-06
5.24E-06
5.25E-06
5.28E-06
5.30E-06
5.30E-06

5.38E-06
5.40E-06
5.41E-06
5.41E-06
5.42E-06
5.46E-06
5.50E-06
5.54E-06
5.54E-06
5.55E-06
5.55E-06
5.59E-06
5.59E-06
5.63E-06
5.64E-06
5.64E-06
5.67E-06
5.69E-06
5.69E-06
5.74E-06
5.76E-06
5.80E-06
5.80E-06
5.82E-06
5.85E-06
5.85E-06
5.90E-06
5.93E-06
5.95E-06
5.96E-06
5.98E-06
5.98E-06
6.01E-06
6.03E-06
6.03E-06
6.08E-06
6.12E-06
6.13E-06
6.13E-06
6.14E-06
6.17E-06
6.24E-06
6.24E-06
6.25E-06
6.31E-06

6.47E-06
6.51E-06
6.51E-06
6.52E-06
6.52E-06
6.52E-06
6.53E-06
6.55E-06
6.56E-06
6.58E-06
6.58E-06
6.59E-06
6.63E-06
6.66E-06
6.70E-06
6.73E-06
6.79E-06
6.85E-06
6.91E-06
6.93E-06
6.98E-06
7.11E-06
7.12E-06
7.16E-06
7.19E-06
7.19E-06
7.19E-06
7.20E-06
7.25E-06
7.28E-06
7.28E-06
7.30E-06
7.32E-06
7.34E-06
7.44E-06
7.46E-06
7.49E-06
7.51E-06
7.62E-06
7.62E-06
7.65E-06
7.67E-06
7.68E-06
7.72E-06
7.73E-06

7.76E-06
7.76E-06
7.84E-06
7.89E-06
7.96E-06
7.98E-06
7.98E-06
7.99E-06
8.05E-06
8.08E-06
8.08E-06
8.08E-06
8.13E-06
8.13E-06
8.14E-06
8.14E-06
8.16E-06
8.24E-06
8.36E-06
8.38E-06
8.39E-06
8.40E-06
8.40E-06
8.44E-06
8.46E-06
8.53E-06
8.59E-06
8.60E-06
8.60E-06
8.66E-06
8.73E-06
8.75E-06
8.78E-06
8.78E-06
8.85E-06
8.96E-06
9.00E-06
9.02E-06
9.03E-06
9.07E-06
9.08E-06
9.12E-06
9.12E-06
9.21E-06
9.32E-06

9.46E-06
9.47E-06
9.56E-06
9.61E-06
9.63E-06
9.63E-06
9.78E-06
9.78E-06
9.85E-06
9.93E-06
9.95E-06
9.96E-06
9.97E-06
1.00E-05
1.01E-05
1.01E-05
1.01E-05
1.01E-05
1.01E-05
1.01E-05
1.01E-05
1.02E-05
1.03E-05
1.03E-05
1.05E-05
1.05E-05
1.06E-05
1.06E-05
1.06E-05
1.06E-05
1.07E-05
1.07E-05
1.07E-05
1.07E-05
1.08E-05
1.08E-05
1.09E-05
1.10E-05
1.10E-05
1.10E-05
1.10E-05
1.11E-05
1.11E-05
1.11E-05
1.11E-05

1.11E-05
1.12E-05
1.12E-05
1.12E-05
1.13E-05
1.13E-05
1.14E-05
1.14E-05
1.15E-05
1.16E-05
1.17E-05
1.17E-05
1.17E-05
1.17E-05
1.18E-05
1.18E-05
1.19E-05
1.19E-05
1.19E-05
1.19E-05
1.20E-05
1.20E-05
1.20E-05
1.20E-05
1.20E-05
1.20E-05
1.22E-05
1.22E-05
1.23E-05
1.25E-05
1.27E-05
1.27E-05
1.28E-05
1.28E-05
1.28E-05
1.30E-05
1.31E-05
1.32E-05
1.33E-05
1.33E-05
1.34E-05
1.34E-05
1.34E-05
1.37E-05
1.37E-05

1.37E-05
1.38E-05
1.38E-05
1.39E-05
1.39E-05
1.39E-05
1.39E-05
1.39E-05
1.40E-05
1.40E-05
1.40E-05
1.41E-05
1.42E-05
1.43E-05
1.43E-05
1.44E-05
1.45E-05
1.46E-05
1.47E-05
1.48E-05
1.48E-05
1.48E-05
1.49E-05
1.49E-05
1.50E-05
1.50E-05
1.50E-05
1.51E-05
1.51E-05
1.51E-05
1.52E-05
1.53E-05
1.54E-05
1.54E-05
1.54E-05
1.55E-05
1.56E-05
1.57E-05
1.57E-05
1.57E-05
1.57E-05
1.59E-05
1.59E-05
1.59E-05
1.60E-05

1.61E-05
1.61E-05
1.61E-05
1.61E-05
1.61E-05
1.62E-05
1.62E-05
1.63E-05
1.64E-05
1.64E-05
1.64E-05
1.64E-05
1.65E-05
1.66E-05
1.66E-05
1.66E-05
1.66E-05
1.67E-05
1.68E-05
1.69E-05
1.69E-05
1.69E-05
1.73E-05
1.74E-05
1.74E-05
1.74E-05
1.75E-05
1.75E-05
1.76E-05
1.76E-05
1.78E-05
1.78E-05
1.78E-05
1.78E-05
1.78E-05
1.80E-05
1.80E-05
1.81E-05
1.81E-05
1.81E-05
1.81E-05
1.82E-05
1.82E-05
1.82E-05
1.83E-05

1.83E-05
1.85E-05
1.85E-05
1.85E-05
1.86E-05
1.87E-05
1.87E-05
1.88E-05
1.88E-05
1.89E-05
1.89E-05
1.89E-05
1.91E-05
1.92E-05
1.93E-05
1.95E-05
1.95E-05
1.96E-05
1.97E-05
1.97E-05
1.98E-05
1.99E-05
1.99E-05
1.99E-05
2.00E-05
2.01E-05
2.03E-05
2.04E-05
2.04E-05
2.05E-05
2.06E-05
2.09E-05
2.09E-05
2.09E-05
2.10E-05
2.10E-05
2.10E-05
2.11E-05
2.12E-05
2.13E-05
2.13E-05
2.14E-05
2.14E-05
2.14E-05
2.15E-05

|  |
| --- |
| 2.15E-05 |
| 2.15E-05 |
| 2.17E-05 |
| 2.20E-05 |
| 2.21E-05 |
| 2.21E-05 |
| 2.22E-05 |
| 2.22E-05 |
| 2.22E-05 |
| 2.23E-05 |
| 2.23E-05 |
| 2.24E-05 |
| 2.27E-05 |
| 2.28E-05 |
| 2.28E-05 |
| 2.28E-05 |
| 2.29E-05 |
| 2.32E-05 |
| 2.35E-05 |
| 2.37E-05 |
| 2.39E-05 |
| 2.40E-05 |
| 2.40E-05 |
| 2.42E-05 |
| 2.42E-05 |
| 2.43E-05 |
| 2.45E-05 |
| 2.46E-05 |
| 2.47E-05 |
| 2.50E-05 |
| 2.51E-05 |
| 2.51E-05 |
| 2.53E-05 |
| 2.54E-05 |
| 2.54E-05 |
| 2.55E-05 |
| 2.58E-05 |
| 2.60E-05 |
| 2.61E-05 |
| 2.61E-05 |
| 2.62E-05 |
| 2.62E-05 |
| 2.63E-05 |
| 2.63E-05 |
| 2.64E-05 |

2.65E-05
2.65E-05
2.71E-05
2.71E-05
2.72E-05
2.72E-05
2.73E-05
2.74E-05
2.74E-05
2.74E-05
2.75E-05
2.75E-05
2.75E-05
2.77E-05
2.77E-05
2.81E-05
2.82E-05
2.83E-05
2.85E-05
2.88E-05
2.89E-05
2.89E-05
2.91E-05
2.91E-05
2.92E-05
2.95E-05
2.97E-05
2.98E-05
2.99E-05
3.00E-05
3.01E-05
3.01E-05
3.01E-05
3.02E-05
3.02E-05
3.03E-05
3.03E-05
3.04E-05
3.04E-05
3.04E-05
3.05E-05
3.05E-05
3.06E-05
3.06E-05
3.07E-05

3.08E-05
3.08E-05
3.08E-05
3.10E-05
3.11E-05
3.12E-05
3.14E-05
3.15E-05
3.15E-05
3.16E-05
3.17E-05
3.19E-05
3.20E-05
3.23E-05
3.25E-05
3.26E-05
3.29E-05
3.30E-05
3.30E-05
3.30E-05
3.31E-05
3.33E-05
3.33E-05
3.33E-05
3.35E-05
3.37E-05
3.37E-05
3.38E-05
3.39E-05
3.41E-05
3.42E-05
3.45E-05
3.46E-05
3.47E-05
3.48E-05
3.48E-05
3.49E-05
3.49E-05
3.51E-05
3.51E-05
3.53E-05
3.56E-05
3.59E-05
3.60E-05
3.61E-05

3.66E-05
3.66E-05
3.72E-05
3.73E-05
3.73E-05
3.74E-05
3.75E-05
3.75E-05
3.76E-05
3.77E-05
3.77E-05
3.77E-05
3.78E-05
3.78E-05
3.81E-05
3.81E-05
3.82E-05
3.82E-05
3.85E-05
3.88E-05
3.90E-05
3.90E-05
3.90E-05
3.91E-05
3.91E-05
3.94E-05
3.97E-05
3.98E-05
4.01E-05
4.02E-05
4.03E-05
4.03E-05
4.03E-05
4.04E-05
4.05E-05
4.05E-05
4.07E-05
4.08E-05
4.12E-05
4.12E-05
4.14E-05
4.14E-05
4.15E-05
4.17E-05
4.17E-05

4.19E-05
4.19E-05
4.21E-05
4.25E-05
4.27E-05
4.29E-05
4.30E-05
4.31E-05
4.33E-05
4.35E-05
4.36E-05
4.38E-05
4.41E-05
4.41E-05
4.43E-05
4.43E-05
4.44E-05
4.45E-05
4.46E-05
4.46E-05
4.47E-05
4.49E-05
4.49E-05
4.50E-05
4.54E-05
4.58E-05
4.60E-05
4.63E-05
4.65E-05
4.65E-05
4.70E-05
4.71E-05
4.76E-05
4.76E-05
4.77E-05
4.80E-05
4.81E-05
4.81E-05
4.82E-05
4.85E-05
4.86E-05
4.87E-05
4.89E-05
4.89E-05
4.92E-05

| |
|---|
| 4.93E-05 |
| 4.93E-05 |
| 4.96E-05 |
| 4.97E-05 |
| 4.98E-05 |
| 4.98E-05 |
| 4.99E-05 |
| 5.04E-05 |
| 5.06E-05 |
| 5.07E-05 |
| 5.07E-05 |
| 5.07E-05 |
| 5.08E-05 |
| 5.08E-05 |
| 5.08E-05 |
| 5.13E-05 |
| 5.13E-05 |
| 5.15E-05 |
| 5.16E-05 |
| 5.18E-05 |
| 5.19E-05 |
| 5.20E-05 |
| 5.20E-05 |
| 5.23E-05 |
| 5.24E-05 |
| 5.25E-05 |
| 5.26E-05 |
| 5.28E-05 |
| 5.30E-05 |
| 5.30E-05 |
| 5.32E-05 |
| 5.32E-05 |
| 5.33E-05 |
| 5.35E-05 |
| 5.38E-05 |
| 5.40E-05 |
| 5.41E-05 |
| 5.41E-05 |
| 5.41E-05 |
| 5.44E-05 |
| 5.46E-05 |
| 5.49E-05 |
| 5.49E-05 |
| 5.50E-05 |
| 5.50E-05 |

5.50E-05
5.53E-05
5.53E-05
5.54E-05
5.54E-05
5.54E-05
5.55E-05
5.57E-05
5.57E-05
5.57E-05
5.58E-05
5.59E-05
5.61E-05
5.63E-05
5.65E-05
5.66E-05
5.67E-05
5.70E-05
5.74E-05
5.74E-05
5.75E-05
5.75E-05
5.77E-05
5.78E-05
5.79E-05
5.79E-05
5.80E-05
5.80E-05
5.80E-05
5.81E-05
5.82E-05
5.84E-05
5.84E-05
5.86E-05
5.87E-05
5.88E-05
5.89E-05
5.90E-05
5.92E-05
5.93E-05
5.93E-05
5.94E-05
5.96E-05
5.97E-05
5.98E-05

5.99E-05
6.06E-05
6.08E-05
6.08E-05
6.10E-05
6.13E-05
6.15E-05
6.15E-05
6.17E-05
6.17E-05
6.21E-05
6.24E-05
6.24E-05
6.25E-05
6.31E-05
6.34E-05
6.36E-05
6.36E-05
6.37E-05
6.38E-05
6.40E-05
6.40E-05
6.45E-05
6.45E-05
6.50E-05
6.51E-05
6.51E-05
6.51E-05
6.52E-05
6.56E-05
6.59E-05
6.61E-05
6.66E-05
6.73E-05
6.80E-05
6.80E-05
6.81E-05
6.84E-05
6.84E-05
6.84E-05
6.91E-05
6.95E-05
6.96E-05
7.05E-05
7.05E-05

7.13E-05
7.15E-05
7.18E-05
7.19E-05
7.20E-05
7.25E-05
7.26E-05
7.35E-05
7.37E-05
7.43E-05
7.45E-05
7.45E-05
7.51E-05
7.54E-05
7.55E-05
7.61E-05
7.64E-05
7.65E-05
7.71E-05
7.72E-05
7.72E-05
7.73E-05
7.75E-05
7.80E-05
7.81E-05
7.82E-05
7.84E-05
7.85E-05
7.86E-05
7.86E-05
7.87E-05
8.00E-05
8.00E-05
8.02E-05
8.04E-05
8.04E-05
8.04E-05
8.16E-05
8.20E-05
8.20E-05
8.23E-05
8.25E-05
8.37E-05
8.38E-05
8.38E-05

8.38E-05
8.44E-05
8.45E-05
8.46E-05
8.47E-05
8.48E-05
8.53E-05
8.53E-05
8.62E-05
8.64E-05
8.64E-05
8.65E-05
8.66E-05
8.67E-05
8.67E-05
8.70E-05
8.77E-05
8.78E-05
8.78E-05
8.82E-05
8.83E-05
8.85E-05
8.95E-05
8.96E-05
8.96E-05
8.96E-05
8.97E-05
8.98E-05
9.00E-05
9.04E-05
9.11E-05
9.11E-05
9.14E-05
9.15E-05
9.19E-05
9.20E-05
9.20E-05
9.24E-05
9.28E-05
9.35E-05
9.44E-05
9.49E-05
9.53E-05
9.60E-05
9.66E-05

| |
|---|
| 9.67E-05 |
| 9.73E-05 |
| 9.76E-05 |
| 9.80E-05 |
| 9.85E-05 |
| 9.85E-05 |
| 9.94E-05 |
| 9.99E-05 |
| 9.99E-05 |
| 0.00010009 |
| 0.00010057 |
| 0.00010085 |
| 0.00010086 |
| 0.00010086 |
| 0.0001011 |
| 0.00010111 |
| 0.00010122 |
| 0.00010123 |
| 0.0001018 |
| 0.0001018 |
| 0.00010202 |
| 0.00010203 |
| 0.00010206 |
| 0.00010246 |
| 0.00010319 |
| 0.00010325 |
| 0.00010325 |
| 0.00010346 |
| 0.00010354 |
| 0.00010466 |
| 0.00010544 |
| 0.00010575 |
| 0.00010598 |
| 0.00010604 |
| 0.00010648 |
| 0.00010699 |
| 0.00010699 |
| 0.00010714 |
| 0.00010747 |
| 0.00010776 |
| 0.00010785 |
| 0.00010797 |
| 0.00010834 |
| 0.00010917 |
| 0.00010932 |

0.00010932
0.00010939
0.00010987
0.00010987
0.0001102
0.00011024
0.00011024
0.0001104
0.00011134
0.00011136
0.00011238
0.00011257
0.00011319
0.00011342
0.00011342
0.00011347
0.00011481
0.00011491
0.00011517
0.00011601
0.00011601
0.00011605
0.00011622
0.00011636
0.00011664
0.00011687
0.00011703
0.0001176
0.00011775
0.00011783
0.00011849
0.00011863
0.00011878
0.00011901
0.00011956
0.00011966
0.00011968
0.0001203
0.00012046
0.00012068
0.00012071
0.00012139
0.0001214
0.0001214
0.00012142

| |
|---|
| 0.00012147 |
| 0.00012159 |
| 0.00012277 |
| 0.0001229 |
| 0.00012307 |
| 0.00012333 |
| 0.00012357 |
| 0.00012374 |
| 0.00012426 |
| 0.00012463 |
| 0.00012468 |
| 0.00012472 |
| 0.00012482 |
| 0.00012523 |
| 0.00012603 |
| 0.00012614 |
| 0.00012646 |
| 0.00012649 |
| 0.00012713 |
| 0.00012713 |
| 0.00012721 |
| 0.00012762 |
| 0.00012813 |
| 0.00012833 |
| 0.00012833 |
| 0.00012887 |
| 0.00012926 |
| 0.00012929 |
| 0.00012936 |
| 0.00012963 |
| 0.00012979 |
| 0.00012981 |
| 0.00013012 |
| 0.00013105 |
| 0.00013111 |
| 0.00013159 |
| 0.00013168 |
| 0.00013175 |
| 0.00013204 |
| 0.00013204 |
| 0.00013213 |
| 0.00013261 |
| 0.00013267 |
| 0.00013273 |
| 0.00013282 |

| |
|---|
| 0.0001329 |
| 0.00013294 |
| 0.00013355 |
| 0.00013355 |
| 0.00013376 |
| 0.0001348 |
| 0.00013619 |
| 0.00013634 |
| 0.0001373 |
| 0.00013739 |
| 0.00013739 |
| 0.00013792 |
| 0.00013844 |
| 0.00013857 |
| 0.00013857 |
| 0.00013859 |
| 0.00013939 |
| 0.00013964 |
| 0.00013996 |
| 0.00014012 |
| 0.0001405 |
| 0.00014119 |
| 0.00014193 |
| 0.00014241 |
| 0.00014242 |
| 0.00014268 |
| 0.00014268 |
| 0.00014308 |
| 0.00014352 |
| 0.00014482 |
| 0.00014546 |
| 0.00014599 |
| 0.00014623 |
| 0.0001464 |
| 0.00014655 |
| 0.00014682 |
| 0.00014971 |
| 0.00015 |
| 0.00015012 |
| 0.0001502 |
| 0.0001504 |
| 0.00015068 |
| 0.00015074 |
| 0.00015155 |
| 0.00015222 |

0.00015222
0.00015283
0.00015335
0.00015382
0.00015408
0.00015413
0.00015447
0.00015473
0.00015505
0.0001565
0.00015686
0.00015737
0.00015795
0.00015796
0.00015807
0.00015808
0.00015817
0.00015836
0.00015845
0.00015863
0.00016015
0.00016038
0.00016062
0.00016163
0.00016217
0.00016218
0.00016308
0.00016308
0.00016329
0.00016329
0.0001634
0.00016393
0.00016398
0.00016435
0.00016461
0.00016518
0.00016621
0.00016728
0.00016775
0.0001683
0.00016842
0.00016896
0.00016979
0.00017054
0.00017063

| |
|---|
| 0.00017081 |
| 0.00017081 |
| 0.00017081 |
| 0.00017138 |
| 0.00017162 |
| 0.00017172 |
| 0.00017174 |
| 0.00017217 |
| 0.00017246 |
| 0.00017273 |
| 0.00017319 |
| 0.0001745 |
| 0.00017514 |
| 0.00017516 |
| 0.00017603 |
| 0.00017625 |
| 0.00017626 |
| 0.00017637 |
| 0.00017688 |
| 0.00017809 |
| 0.00017829 |
| 0.00017977 |
| 0.00018018 |
| 0.00018036 |
| 0.00018128 |
| 0.0001813 |
| 0.00018245 |
| 0.00018248 |
| 0.00018287 |
| 0.00018305 |
| 0.0001835 |
| 0.00018367 |
| 0.00018627 |
| 0.00018689 |
| 0.00018801 |
| 0.00018892 |
| 0.00018909 |
| 0.00018924 |
| 0.00018977 |
| 0.00018995 |
| 0.00019057 |
| 0.00019096 |
| 0.00019098 |
| 0.00019104 |
| 0.0001913 |

| |
|---|
| 0.00019144 |
| 0.00019158 |
| 0.00019208 |
| 0.00019246 |
| 0.00019255 |
| 0.00019315 |
| 0.00019384 |
| 0.00019401 |
| 0.00019437 |
| 0.00019449 |
| 0.00019463 |
| 0.00019512 |
| 0.00019547 |
| 0.00019635 |
| 0.00019653 |
| 0.00019759 |
| 0.00019764 |
| 0.00019799 |
| 0.00020053 |
| 0.00020082 |
| 0.00020121 |
| 0.00020165 |
| 0.00020175 |
| 0.00020262 |
| 0.00020295 |
| 0.0002031 |
| 0.00020364 |
| 0.00020465 |
| 0.00020501 |
| 0.00020605 |
| 0.00020605 |
| 0.00020606 |
| 0.00020637 |
| 0.00020651 |
| 0.00020652 |
| 0.00020654 |
| 0.00020957 |
| 0.00020957 |
| 0.00020961 |
| 0.00020999 |
| 0.00021008 |
| 0.00021008 |
| 0.00021008 |
| 0.00021166 |
| 0.00021215 |

| |
|---|
| 0.00021217 |
| 0.00021286 |
| 0.00021286 |
| 0.00021399 |
| 0.00021482 |
| 0.00021495 |
| 0.00021497 |
| 0.00021628 |
| 0.0002168 |
| 0.00021682 |
| 0.00021811 |
| 0.00022064 |
| 0.00022236 |
| 0.00022268 |
| 0.00022319 |
| 0.00022351 |
| 0.00022443 |
| 0.00022597 |
| 0.00022611 |
| 0.00022752 |
| 0.00022753 |
| 0.00022868 |
| 0.00022885 |
| 0.00022985 |
| 0.00022994 |
| 0.00023391 |
| 0.00023404 |
| 0.0002342 |
| 0.00023464 |
| 0.00023474 |
| 0.00023476 |
| 0.00023504 |
| 0.00023551 |
| 0.00023557 |
| 0.0002367 |
| 0.0002367 |
| 0.0002374 |
| 0.00023777 |
| 0.00023777 |
| 0.00023777 |
| 0.00023784 |
| 0.00023875 |
| 0.00024031 |
| 0.00024139 |
| 0.00024186 |

| |
|---|
| 0.00024214 |
| 0.00024214 |
| 0.00024219 |
| 0.0002424 |
| 0.00024263 |
| 0.00024427 |
| 0.00024521 |
| 0.00024522 |
| 0.00024646 |
| 0.00024706 |
| 0.00024725 |
| 0.00024804 |
| 0.00024804 |
| 0.00025052 |
| 0.00025114 |
| 0.00025277 |
| 0.00025407 |
| 0.00025447 |
| 0.00025495 |
| 0.00025533 |
| 0.00025559 |
| 0.00025673 |
| 0.00025701 |
| 0.00025732 |
| 0.0002594 |
| 0.00025982 |
| 0.00026022 |
| 0.00026058 |
| 0.0002645 |
| 0.00026475 |
| 0.00026582 |
| 0.00026655 |
| 0.00026715 |
| 0.00026795 |
| 0.00026857 |
| 0.00027028 |
| 0.00027041 |
| 0.00027041 |
| 0.00027068 |
| 0.00027068 |
| 0.00027346 |
| 0.00027386 |
| 0.000275 |
| 0.00027506 |
| 0.00027506 |

| |
|---|
| 0.0002753 |
| 0.00027576 |
| 0.00027643 |
| 0.0002791 |
| 0.00027921 |
| 0.00028093 |
| 0.00028357 |
| 0.00028561 |
| 0.0002862 |
| 0.0002862 |
| 0.00029189 |
| 0.00029339 |
| 0.00029339 |
| 0.00029442 |
| 0.00029561 |
| 0.0002958 |
| 0.00029596 |
| 0.00029653 |
| 0.00029678 |
| 0.00029684 |
| 0.0002989 |
| 0.00029993 |
| 0.00030144 |
| 0.00030144 |
| 0.00030144 |
| 0.00030171 |
| 0.00030218 |
| 0.00030481 |
| 0.00030583 |
| 0.00030626 |
| 0.00030659 |
| 0.00030894 |
| 0.00030929 |
| 0.00030964 |
| 0.00031049 |
| 0.00031068 |
| 0.00031094 |
| 0.00031405 |
| 0.00031482 |
| 0.00031529 |
| 0.00031703 |
| 0.00031715 |
| 0.00031774 |
| 0.00031896 |
| 0.00032101 |

0.00032222
0.00032237
0.00032237
0.00032284
0.00032448
0.00032521
0.00032526
0.0003267
0.00032709
0.00032719
0.00032721
0.00032721
0.00032732
0.00032766
0.00032875
0.0003289
0.00032971
0.00033035
0.00033103
0.00033187
0.00033187
0.00033266
0.00033278
0.00033426
0.00033512
0.00033645
0.00033739
0.00033755
0.00033762
0.0003379
0.00033852
0.00034008
0.0003402
0.00034101
0.00034106
0.00034106
0.00034106
0.00034304
0.00034308
0.00034537
0.00034538
0.00034624
0.00034642
0.00034666
0.00034666

| |
|---|
| 0.00034727 |
| 0.0003476 |
| 0.00034994 |
| 0.00035033 |
| 0.00035079 |
| 0.00035221 |
| 0.0003529 |
| 0.0003533 |
| 0.00035333 |
| 0.00035367 |
| 0.0003558 |
| 0.00035868 |
| 0.00036125 |
| 0.00036189 |
| 0.00036241 |
| 0.00036357 |
| 0.00036452 |
| 0.000365 |
| 0.00036532 |
| 0.00036599 |
| 0.00036619 |
| 0.00036651 |
| 0.00036753 |
| 0.00036753 |
| 0.00036759 |
| 0.00036835 |
| 0.00036898 |
| 0.00036899 |
| 0.00036919 |
| 0.00036919 |
| 0.00036919 |
| 0.00036926 |
| 0.00036966 |
| 0.00036973 |
| 0.00037216 |
| 0.00037477 |
| 0.00037547 |
| 0.00037582 |
| 0.00037582 |
| 0.00037585 |
| 0.00037755 |
| 0.00037808 |
| 0.00037852 |
| 0.00037861 |
| 0.000379 |

| |
|---|
| 0.00037905 |
| 0.00037905 |
| 0.00037907 |
| 0.00037937 |
| 0.00037947 |
| 0.00037947 |
| 0.0003797 |
| 0.00037992 |
| 0.00038254 |
| 0.00038595 |
| 0.00038602 |
| 0.00038652 |
| 0.00038696 |
| 0.00038879 |
| 0.00038885 |
| 0.00038915 |
| 0.00038979 |
| 0.00038979 |
| 0.00039045 |
| 0.00039095 |
| 0.00039232 |
| 0.00039373 |
| 0.00039491 |
| 0.00039522 |
| 0.00039526 |
| 0.00039738 |
| 0.0003983 |
| 0.00039981 |
| 0.00040146 |
| 0.00040196 |
| 0.00040236 |
| 0.00040245 |
| 0.00040247 |
| 0.00040284 |
| 0.00040312 |
| 0.00040703 |
| 0.00040824 |
| 0.00040859 |
| 0.00040878 |
| 0.00041082 |
| 0.0004121 |
| 0.00041246 |
| 0.00041262 |
| 0.00041308 |
| 0.00041373 |

0.00041458
0.00041565
0.00041594
0.00041661
0.00041792
0.00042005
0.00042095
0.00042179
0.0004232
0.00042558
0.00042603
0.00042722
0.00042922
0.00042944
0.0004305
0.00043244
0.00043301
0.0004331
0.00043366
0.00043551
0.00043657
0.00043679
0.00043713
0.00044195
0.00044287
0.00044479
0.00044498
0.00044506
0.00044641
0.00044697
0.00044711
0.00044788
0.0004502
0.00045039
0.00045063
0.00045193
0.00045201
0.00045201
0.0004538
0.00045609
0.00045641
0.0004566
0.00045747
0.00045865
0.00045995

0.00046069
0.00046138
0.00046179
0.00046294
0.00046297
0.00046313
0.00046684
0.00046688
0.00046773
0.00046783
0.00046866
0.000471
0.00047101
0.00047474
0.00047502
0.00047666
0.0004783
0.00047854
0.00047893
0.00047978
0.00048019
0.00048345
0.00048444
0.0004868
0.00048753
0.00048794
0.00048948
0.00049369
0.00049462
0.00050071
0.0005014
0.00050232
0.00050305
0.0005039
0.0005039
0.00050441
0.00050466
0.0005049
0.00050552
0.00050571
0.00050616
0.00050665
0.00050994
0.00051105
0.00051409

| |
|---|
| 0.00051561 |
| 0.00051855 |
| 0.00051878 |
| 0.00052145 |
| 0.00052399 |
| 0.00052525 |
| 0.00052549 |
| 0.00052694 |
| 0.00052986 |
| 0.00053298 |
| 0.00053368 |
| 0.00053465 |
| 0.00053856 |
| 0.00053864 |
| 0.00053997 |
| 0.00054005 |
| 0.00054046 |
| 0.00054171 |
| 0.00054335 |
| 0.0005439 |
| 0.00054772 |
| 0.00054792 |
| 0.00054812 |
| 0.00055053 |
| 0.00055205 |
| 0.00055432 |
| 0.0005547 |
| 0.00055624 |
| 0.00055741 |
| 0.00056925 |
| 0.00056954 |
| 0.00057065 |
| 0.00057109 |
| 0.00057211 |
| 0.00057239 |
| 0.00057388 |
| 0.00057651 |
| 0.00057728 |
| 0.00057874 |
| 0.00057959 |
| 0.00058706 |
| 0.00058709 |
| 0.00058814 |
| 0.00059089 |
| 0.00059202 |

0.00059231
0.00059286
0.00059333
0.00059693
0.0005976
0.00059808
0.00059816
0.00059882
0.00060222
0.00060331
0.00060379
0.00060379
0.00060446
0.00060538
0.00060749
0.00060837
0.00060962
0.00061097
0.00061148
0.0006118
0.00061274
0.00061795
0.00061977
0.00062001
0.0006219
0.00062249
0.00062424
0.00062449
0.00062484
0.00062535
0.00062572
0.00062625
0.00062735
0.00062834
0.000629
0.00063301
0.0006342
0.00063635
0.0006382
0.00064027
0.00064027
0.00064109
0.00064218
0.00064304
0.00064409

0.00064497
0.00064599
0.00064622
0.00064679
0.00064865
0.00064897
0.0006507
0.0006526
0.00065307
0.0006533
0.00065529
0.00065603
0.00065668
0.00065746
0.00065908
0.00066201
0.00066317
0.00066515
0.00066553
0.00066575
0.00066942
0.00067186
0.00067308
0.00067374
0.00067551
0.00067817
0.00068112
0.00068112
0.00068492
0.00068653
0.00068698
0.00068698
0.000689
0.00068994
0.00069978
0.00070086
0.00070172
0.00070409
0.00070426
0.00070426
0.00070516
0.0007052
0.0007052
0.00070611
0.00070634

0.00070888
0.00070989
0.00071128
0.00071128
0.00071155
0.00071267
0.00071301
0.00071465
0.00071605
0.00072078
0.00072239
0.00072432
0.00072466
0.00072494
0.00072534
0.00072694
0.00072736
0.00072851
0.00072888
0.00072906
0.00072989
0.00073085
0.00073085
0.0007321
0.00073352
0.00073409
0.00073476
0.00073489
0.00073548
0.0007369
0.000737
0.00073847
0.00073881
0.00073956
0.00074173
0.00074242
0.00074534
0.0007459
0.00075006
0.00075835
0.00075897
0.00076072
0.00076129
0.00076173
0.00076402

| |
|---|
| 0.0007659 |
| 0.00076806 |
| 0.00077274 |
| 0.00077402 |
| 0.00077666 |
| 0.00077725 |
| 0.00078425 |
| 0.000786 |
| 0.00078601 |
| 0.00078625 |
| 0.00078724 |
| 0.00078834 |
| 0.00078834 |
| 0.00078837 |
| 0.00079074 |
| 0.00079171 |
| 0.00079361 |
| 0.00079393 |
| 0.00079472 |
| 0.00079524 |
| 0.00079524 |
| 0.00079801 |
| 0.00080095 |
| 0.00080161 |
| 0.00080161 |
| 0.00080297 |
| 0.00080332 |
| 0.00080811 |
| 0.00081223 |
| 0.00081343 |
| 0.00081391 |
| 0.00081536 |
| 0.00081577 |
| 0.00081577 |
| 0.00081723 |
| 0.00081869 |
| 0.00081988 |
| 0.0008199 |
| 0.00082075 |
| 0.00082169 |
| 0.00082307 |
| 0.00082529 |
| 0.00082742 |
| 0.00083253 |
| 0.00083377 |

| |
|---|
| 0.00083377 |
| 0.00083567 |
| 0.00083567 |
| 0.00083824 |
| 0.0008384 |
| 0.00083908 |
| 0.00083948 |
| 0.00083951 |
| 0.00083964 |
| 0.00084308 |
| 0.00084955 |
| 0.00084992 |
| 0.00085008 |
| 0.00085029 |
| 0.00085238 |
| 0.00085246 |
| 0.00085343 |
| 0.00085533 |
| 0.00086398 |
| 0.00087137 |
| 0.00087205 |
| 0.00087268 |
| 0.00087429 |
| 0.00087713 |
| 0.00087922 |
| 0.00088098 |
| 0.00088107 |
| 0.00088402 |
| 0.00088533 |
| 0.00088546 |
| 0.00088768 |
| 0.0008886 |
| 0.00089226 |
| 0.00089258 |
| 0.00089338 |
| 0.00089446 |
| 0.00089457 |
| 0.00089565 |
| 0.00089565 |
| 0.00089565 |
| 0.00089725 |
| 0.00089745 |
| 0.00089828 |
| 0.00090447 |
| 0.0009069 |

| |
|---|
| 0.0009069 |
| 0.00090859 |
| 0.00091067 |
| 0.00091285 |
| 0.00091357 |
| 0.00091357 |
| 0.00091683 |
| 0.00091742 |
| 0.00091779 |
| 0.00091819 |
| 0.00091973 |
| 0.00092088 |
| 0.0009229 |
| 0.00092815 |
| 0.00093089 |
| 0.00093441 |
| 0.00093479 |
| 0.00093848 |
| 0.00094103 |
| 0.00094103 |
| 0.00094127 |
| 0.00094131 |
| 0.00094336 |
| 0.00094654 |
| 0.0009489 |
| 0.00094918 |
| 0.00095124 |
| 0.00095129 |
| 0.00095372 |
| 0.00095395 |
| 0.00095399 |
| 0.00095484 |
| 0.00095526 |
| 0.00095526 |
| 0.00095845 |
| 0.00095845 |
| 0.00096138 |
| 0.00096295 |
| 0.00096391 |
| 0.00096759 |
| 0.00096891 |
| 0.0009733 |
| 0.00097921 |
| 0.00097987 |
| 0.00098097 |

| |
|---|
| 0.00098621 |
| 0.00098855 |
| 0.00098875 |
| 0.00098906 |
| 0.00099311 |
| 0.00099311 |
| 0.00099421 |
| 0.00099478 |
| 0.00099478 |
| 0.00099697 |
| 0.00099728 |
| 0.00099739 |
| 0.00099872 |
| 0.00100083 |
| 0.00100418 |
| 0.00100435 |
| 0.00100482 |
| 0.00100509 |
| 0.00100712 |
| 0.00100799 |
| 0.00101286 |
| 0.00101386 |
| 0.00101387 |
| 0.00101563 |
| 0.001017 |
| 0.00102078 |
| 0.00102166 |
| 0.00102523 |
| 0.00102535 |
| 0.0010262 |
| 0.00102706 |
| 0.00102966 |
| 0.00103121 |
| 0.00103591 |
| 0.00103775 |
| 0.00104239 |
| 0.00104328 |
| 0.00104418 |
| 0.00104643 |
| 0.00105016 |
| 0.00105203 |
| 0.00105404 |
| 0.00105449 |
| 0.00105488 |
| 0.00105574 |

| |
|---|
| 0.00106467 |
| 0.00106583 |
| 0.00106837 |
| 0.0010694 |
| 0.00107032 |
| 0.00107116 |
| 0.00107222 |
| 0.0010807 |
| 0.00108081 |
| 0.00108722 |
| 0.00108856 |
| 0.00109334 |
| 0.00109573 |
| 0.00110085 |
| 0.00110265 |
| 0.00110378 |
| 0.00110954 |
| 0.00110954 |
| 0.00111753 |
| 0.00112058 |
| 0.00112098 |
| 0.00112104 |
| 0.00112176 |
| 0.001125 |
| 0.00112621 |
| 0.00112855 |
| 0.00113224 |
| 0.00113284 |
| 0.00113309 |
| 0.00113505 |
| 0.00113662 |
| 0.00113961 |
| 0.00113961 |
| 0.00114089 |
| 0.00115068 |
| 0.00115212 |
| 0.00115523 |
| 0.00115859 |
| 0.00115861 |
| 0.00115889 |
| 0.00116027 |
| 0.0011603 |
| 0.0011603 |
| 0.00116043 |
| 0.00116154 |

| |
|---|
| 0.0011626 |
| 0.00116373 |
| 0.00116478 |
| 0.00116478 |
| 0.00116653 |
| 0.00116975 |
| 0.00117032 |
| 0.00117513 |
| 0.00117796 |
| 0.00117976 |
| 0.001182 |
| 0.00118735 |
| 0.00118923 |
| 0.00118931 |
| 0.00119019 |
| 0.00119122 |
| 0.00119147 |
| 0.00119147 |
| 0.00119195 |
| 0.00119384 |
| 0.00119574 |
| 0.001198 |
| 0.001198 |
| 0.00120056 |
| 0.00120085 |
| 0.00120253 |
| 0.00121868 |
| 0.00121881 |
| 0.00122442 |
| 0.00122949 |
| 0.00122962 |
| 0.00123211 |
| 0.00123601 |
| 0.00123655 |
| 0.00123776 |
| 0.00123832 |
| 0.0012431 |
| 0.00124657 |
| 0.00125497 |
| 0.00125694 |
| 0.0012587 |
| 0.00126169 |
| 0.00126175 |
| 0.00126175 |
| 0.00126723 |

| |
|---|
| 0.00127167 |
| 0.00127218 |
| 0.00127332 |
| 0.00127617 |
| 0.0012775 |
| 0.00128125 |
| 0.00128874 |
| 0.00129114 |
| 0.00129338 |
| 0.0012934 |
| 0.00129516 |
| 0.00129754 |
| 0.00130356 |
| 0.00130902 |
| 0.00130992 |
| 0.00132077 |
| 0.00132435 |
| 0.00132552 |
| 0.00132552 |
| 0.00132772 |
| 0.00132866 |
| 0.00133027 |
| 0.00133321 |
| 0.00133404 |
| 0.00133513 |
| 0.00134815 |
| 0.00134819 |
| 0.00134869 |
| 0.00135134 |
| 0.00135592 |
| 0.00136244 |
| 0.00136244 |
| 0.00136244 |
| 0.00136998 |
| 0.0013707 |
| 0.00137515 |
| 0.00138035 |
| 0.00138035 |
| 0.00138281 |
| 0.00138478 |
| 0.00138626 |
| 0.00139355 |
| 0.00139721 |
| 0.00140059 |
| 0.00140085 |

| |
|---|
| 0.00140174 |
| 0.00141078 |
| 0.00141464 |
| 0.00141606 |
| 0.00141915 |
| 0.00142345 |
| 0.00142453 |
| 0.00142614 |
| 0.00142716 |
| 0.00143161 |
| 0.00143306 |
| 0.00143492 |
| 0.00143835 |
| 0.0014391 |
| 0.00143967 |
| 0.00144047 |
| 0.001442 |
| 0.00144265 |
| 0.00144391 |
| 0.00145001 |
| 0.00145043 |
| 0.00145246 |
| 0.00145678 |
| 0.00145761 |
| 0.00145934 |
| 0.00146363 |
| 0.00146473 |
| 0.00146485 |
| 0.00146607 |
| 0.00146861 |
| 0.00146875 |
| 0.00147597 |
| 0.00147783 |
| 0.00147974 |
| 0.00148036 |
| 0.00148681 |
| 0.00149921 |
| 0.00150494 |
| 0.00150787 |
| 0.00151306 |
| 0.00151405 |
| 0.00152011 |
| 0.00152098 |
| 0.00152225 |
| 0.00152305 |

| |
|---|
| 0.0015268 |
| 0.00153052 |
| 0.00154788 |
| 0.00154862 |
| 0.00155727 |
| 0.00155743 |
| 0.00155912 |
| 0.00156129 |
| 0.00156252 |
| 0.00156278 |
| 0.00156428 |
| 0.00156711 |
| 0.00156997 |
| 0.00157749 |
| 0.00158162 |
| 0.00158196 |
| 0.00158242 |
| 0.00158361 |
| 0.00158563 |
| 0.00158667 |
| 0.00158795 |
| 0.00158795 |
| 0.00158874 |
| 0.00158874 |
| 0.00158938 |
| 0.00158938 |
| 0.00159163 |
| 0.0015942 |
| 0.00160016 |
| 0.00160134 |
| 0.00160273 |
| 0.00160479 |
| 0.00160522 |
| 0.00160615 |
| 0.00161397 |
| 0.00161708 |
| 0.001619 |
| 0.00161902 |
| 0.00161942 |
| 0.00162 |
| 0.00162015 |
| 0.00162015 |
| 0.00162577 |
| 0.00162703 |
| 0.00162854 |

0.00163102
0.00163208
0.0016353
0.00164343
0.00164343
0.00164424
0.00164613
0.00164771
0.00164875
0.00165282
0.00165574
0.00165606
0.0016618
0.00166429
0.00166429
0.00166556
0.00167167
0.00167359
0.00167497
0.00167527
0.00168777
0.00168798
0.0016935
0.00169653
0.00169949
0.00170569
0.00170682
0.00171462
0.00171684
0.00173011
0.00173376
0.00173798
0.0017423
0.00174768
0.00174863
0.00175035
0.00175176
0.00175218
0.00175309
0.00175464
0.00175716
0.00175776
0.00176028
0.0017685
0.00177033

| |
|---|
| 0.00177154 |
| 0.00177154 |
| 0.00177691 |
| 0.00178877 |
| 0.00179141 |
| 0.00179819 |
| 0.00179974 |
| 0.00180238 |
| 0.00180406 |
| 0.00181626 |
| 0.0018166 |
| 0.00181731 |
| 0.00181817 |
| 0.00182783 |
| 0.00183147 |
| 0.00183387 |
| 0.0018349 |
| 0.00183736 |
| 0.00184214 |
| 0.00184996 |
| 0.00185206 |
| 0.00185335 |
| 0.00185378 |
| 0.0018543 |
| 0.00185735 |
| 0.00186132 |
| 0.00186189 |
| 0.00186189 |
| 0.00186378 |
| 0.00187189 |
| 0.00187224 |
| 0.00187575 |
| 0.00187641 |
| 0.00187641 |
| 0.00188417 |
| 0.00189076 |
| 0.00189818 |
| 0.0019026 |
| 0.00190597 |
| 0.00190597 |
| 0.00190787 |
| 0.0019124 |
| 0.00191242 |
| 0.00191823 |
| 0.00191932 |

| |
|---|
| 0.00192311 |
| 0.001925 |
| 0.00192527 |
| 0.00193201 |
| 0.00194143 |
| 0.0019428 |
| 0.00194589 |
| 0.00195062 |
| 0.00195224 |
| 0.00195516 |
| 0.00196205 |
| 0.00196654 |
| 0.0019691 |
| 0.00196974 |
| 0.00197735 |
| 0.00199104 |
| 0.0019916 |
| 0.0019986 |
| 0.00200023 |
| 0.00200023 |
| 0.00200061 |
| 0.00200061 |
| 0.00200263 |
| 0.00200616 |
| 0.00200863 |
| 0.00200994 |
| 0.00200994 |
| 0.00201055 |
| 0.00202121 |
| 0.00202121 |
| 0.00202138 |
| 0.00202302 |
| 0.00202999 |
| 0.00203047 |
| 0.00203464 |
| 0.00204497 |
| 0.00204535 |
| 0.00205242 |
| 0.00205324 |
| 0.00206143 |
| 0.00206433 |
| 0.00207131 |
| 0.00207428 |
| 0.0020747 |
| 0.00208154 |

0.00208452
0.00208705
0.00208741
0.0020985
0.00210349
0.00210989
0.00211013
0.00211462
0.00212329
0.00213526
0.00213938
0.00213938
0.00213938
0.00214065
0.00214982
0.00216054
0.00216238
0.00216927
0.00216981
0.00217074
0.00217074
0.00217574
0.00218164
0.00218164
0.00218734
0.00218971
0.00219
0.0021987
0.00220584
0.00221276
0.00221527
0.00221867
0.00223354
0.00223424
0.00224771
0.00225064
0.00225064
0.00225704
0.00225853
0.00226002
0.00226113
0.00226113
0.00226588
0.00227247
0.00227293

0.00227429
0.00227429
0.0022757
0.002277
0.00228433
0.0022852
0.00228965
0.00228965
0.00229133
0.00229153
0.00229714
0.00229797
0.00230082
0.00230148
0.0023016
0.00230593
0.00230997
0.0023132
0.00233128
0.00233213
0.00235653
0.00235671
0.00235762
0.00235993
0.00236406
0.00237229
0.00237833
0.00238108
0.00238184
0.00238538
0.00239067
0.00239067
0.00239075
0.00239075
0.00239207
0.00239266
0.00239279
0.00239301
0.00239929
0.0024063
0.00241546
0.00241637
0.0024196
0.0024198
0.0024198

0.00242492
0.00243026
0.00243955
0.00244148
0.00244169
0.00244249
0.00244661
0.00244692
0.00244742
0.00244742
0.00244749
0.00245141
0.00245141
0.00246189
0.00246267
0.00246721
0.00246987
0.00247228
0.00247527
0.00247543
0.0024762
0.00248451
0.00248517
0.00248517
0.00249088
0.00249229
0.00250161
0.00250288
0.00250592
0.00250736
0.00251178
0.00251178
0.00251249
0.00251331
0.00252764
0.00253539
0.00254062
0.00254663
0.00254822
0.00255518
0.00256091
0.00256541
0.00256899
0.00257615
0.00258399

0.00258834
0.00259305
0.00259518
0.00259686
0.00260387
0.00260974
0.00261432
0.00261666
0.00262146
0.00262343
0.00262509
0.00262556
0.00262854
0.00263126
0.00264509
0.00265654
0.00266264
0.00266264
0.00266271
0.00266401
0.00266681
0.00266791
0.00267028
0.00267486
0.00267486
0.00268024
0.00268293
0.00268431
0.0026899
0.00269823
0.00270413
0.00270633
0.00270793
0.00270979
0.00270979
0.00271078
0.00271239
0.00271781
0.00272823
0.00272963
0.00273188
0.0027358
0.0027364
0.00273956
0.00273982

| |
|---|
| 0.00274599 |
| 0.00275545 |
| 0.00276347 |
| 0.00276503 |
| 0.00276646 |
| 0.00277377 |
| 0.00277377 |
| 0.00277779 |
| 0.00277977 |
| 0.00278639 |
| 0.00279153 |
| 0.0027951 |
| 0.00279554 |
| 0.00279906 |
| 0.00279948 |
| 0.00280417 |
| 0.00280417 |
| 0.00281672 |
| 0.00281741 |
| 0.00284085 |
| 0.00284483 |
| 0.00284624 |
| 0.00284873 |
| 0.00285295 |
| 0.00285307 |
| 0.00285395 |
| 0.00287359 |
| 0.00288981 |
| 0.00289202 |
| 0.00290198 |
| 0.00290312 |
| 0.00290312 |
| 0.00290633 |
| 0.00291035 |
| 0.00291601 |
| 0.0029217 |
| 0.00292468 |
| 0.00292508 |
| 0.00292757 |
| 0.0029443 |
| 0.00295044 |
| 0.00295044 |
| 0.00297994 |
| 0.00297999 |
| 0.00298044 |

0.00298579
0.00299888
0.00300637
0.00300646
0.0030076
0.00302164
0.00302356
0.00302548
0.00302898
0.00303125
0.00304048
0.00305276
0.00305723
0.00307133
0.0030752
0.00307704
0.00308581
0.00309017
0.00309208
0.00310479
0.00311176
0.00311176
0.00311217
0.00311217
0.00311635
0.00312082
0.00312249
0.00312512
0.0031291
0.0031426
0.00314464
0.00314959
0.00315478
0.00317288
0.00318831
0.00320489
0.00321064
0.00321211
0.00321593
0.00321686
0.00322004
0.00322708
0.00322708
0.00323341
0.00323736

0.00323998
0.0032401
0.00324748
0.00326692
0.00327038
0.00327175
0.00327503
0.00328249
0.00328249
0.00328831
0.00328849
0.00328998
0.00329065
0.00329241
0.00330461
0.00330752
0.00331519
0.00331701
0.00332054
0.00332054
0.0033289
0.00335057
0.00336071
0.00336522
0.00336717
0.00337893
0.00339286
0.00340902
0.00342085
0.00342718
0.00343902
0.00344099
0.00344446
0.00344919
0.00345511
0.00345643
0.00345643
0.00346285
0.00346625
0.00350725
0.00352454
0.00352864
0.00353447
0.00353447
0.00354686

0.00355035
0.00355297
0.00356189
0.00356218
0.00356352
0.00358843
0.0035961
0.00360359
0.00361169
0.00361977
0.00362398
0.00362436
0.00362556
0.00362875
0.00367028
0.00367232
0.00367232
0.00367631
0.00368129
0.00368224
0.00368244
0.00369289
0.00371674
0.00371694
0.0037182
0.00372069
0.00372327
0.00372387
0.00372387
0.00373303
0.00373303
0.00373462
0.00374127
0.0037475
0.00375885
0.00376088
0.00376184
0.00376283
0.0037669
0.00376697
0.00377555
0.00378444
0.00378504
0.00378676
0.00378676

| |
|---|
| 0.00378874 |
| 0.00379948 |
| 0.00380304 |
| 0.00380583 |
| 0.00382085 |
| 0.00382855 |
| 0.00383197 |
| 0.00383197 |
| 0.00384612 |
| 0.00386284 |
| 0.00386449 |
| 0.00386715 |
| 0.00387645 |
| 0.00388906 |
| 0.00392104 |
| 0.00393065 |
| 0.00393277 |
| 0.0039683 |
| 0.00397199 |
| 0.00399033 |
| 0.00399333 |
| 0.00400782 |
| 0.00401091 |
| 0.0040126 |
| 0.00403138 |
| 0.00403964 |
| 0.00404198 |
| 0.00404289 |
| 0.00405842 |
| 0.00406857 |
| 0.00407869 |
| 0.00407869 |
| 0.00407869 |
| 0.00408235 |
| 0.00409664 |
| 0.0041026 |
| 0.0041124 |
| 0.00411952 |
| 0.00412481 |
| 0.00412829 |
| 0.00413021 |
| 0.00413595 |
| 0.00415716 |
| 0.00415716 |
| 0.00415842 |

0.00416381
0.00417041
0.00417175
0.00417614
0.0041961
0.00420644
0.00422029
0.00422029
0.00422066
0.00422508
0.00422943
0.004248
0.00426677
0.00427146
0.0042771
0.00429101
0.00429368
0.00430328
0.00431301
0.00432678
0.00433434
0.00434354
0.00434514
0.00434514
0.00434684
0.00436583
0.00438234
0.00438597
0.00443015
0.00444156
0.00444424
0.00445645
0.00446191
0.00447249
0.0044805
0.00449495
0.00449582
0.00449745
0.00450509
0.00450682
0.00450682
0.00452904
0.00453079
0.00453266
0.00454106

0.00455177
0.00456763
0.00456763
0.00456787
0.00457664
0.00458696
0.00459075
0.0046157
0.00463373
0.00463951
0.00465122
0.00467058
0.00467058
0.00468889
0.00470808
0.00471769
0.00473324
0.00473633
0.00474722
0.00475157
0.00475427
0.00475709
0.00475834
0.00476394
0.00476614
0.00477023
0.00477419
0.00478374
0.00479254
0.00479254
0.00479322
0.00479524
0.00480438
0.00481982
0.00482006
0.00482132
0.00483346
0.00484925
0.00486689
0.00486917
0.00487262
0.00487262
0.0048788
0.00489844
0.00493545

| |
|---|
| 0.00494043 |
| 0.00494735 |
| 0.00496538 |
| 0.0049677 |
| 0.00497562 |
| 0.00498102 |
| 0.00498397 |
| 0.00498762 |
| 0.00498762 |
| 0.00500633 |
| 0.00503827 |
| 0.00503827 |
| 0.00504646 |
| 0.00505027 |
| 0.00506944 |
| 0.00509672 |
| 0.00513849 |
| 0.00513849 |
| 0.00521353 |
| 0.00522141 |
| 0.00524413 |
| 0.00525717 |
| 0.00530429 |
| 0.00533697 |
| 0.00533798 |
| 0.0053441 |
| 0.00534509 |
| 0.00535075 |
| 0.005351 |
| 0.00535935 |
| 0.00536685 |
| 0.00537629 |
| 0.00538055 |
| 0.00538236 |
| 0.00538647 |
| 0.00540647 |
| 0.00544015 |
| 0.00546053 |
| 0.00546053 |
| 0.00546056 |
| 0.00547406 |
| 0.00547815 |
| 0.00548483 |
| 0.00549415 |
| 0.00550915 |

| |
|---|
| 0.00550915 |
| 0.00550915 |
| 0.00552896 |
| 0.00553011 |
| 0.00553046 |
| 0.00553309 |
| 0.00554903 |
| 0.00554903 |
| 0.0055616 |
| 0.0055693 |
| 0.0055693 |
| 0.00557184 |
| 0.00557884 |
| 0.00558064 |
| 0.00559043 |
| 0.00559548 |
| 0.00559663 |
| 0.00560133 |
| 0.00560257 |
| 0.00562054 |
| 0.00564786 |
| 0.00564916 |
| 0.00565282 |
| 0.00565827 |
| 0.00567428 |
| 0.00567699 |
| 0.00569404 |
| 0.00570815 |
| 0.00570815 |
| 0.00570834 |
| 0.00571594 |
| 0.00572532 |
| 0.00577376 |
| 0.00577844 |
| 0.00578566 |
| 0.00579118 |
| 0.00579554 |
| 0.00580628 |
| 0.00580915 |
| 0.00584427 |
| 0.00585799 |
| 0.00586047 |
| 0.0058757 |
| 0.00589404 |
| 0.0059063 |

| |
|---|
| 0.0059063 |
| 0.00590757 |
| 0.00590949 |
| 0.00591164 |
| 0.00591447 |
| 0.00591551 |
| 0.00597422 |
| 0.00599436 |
| 0.00599436 |
| 0.00599436 |
| 0.00599436 |
| 0.00599478 |
| 0.00600933 |
| 0.00601895 |
| 0.00605173 |
| 0.0060627 |
| 0.00606943 |
| 0.00606953 |
| 0.00608195 |
| 0.0060825 |
| 0.00609862 |
| 0.00611738 |
| 0.00614818 |
| 0.00616354 |
| 0.00616354 |
| 0.006209 |
| 0.00622649 |
| 0.00623078 |
| 0.00623241 |
| 0.00625609 |
| 0.00628704 |
| 0.00632241 |
| 0.00632362 |
| 0.00632815 |
| 0.00632815 |
| 0.00634398 |
| 0.00635078 |
| 0.00635078 |
| 0.0063974 |
| 0.0064012 |
| 0.00640236 |
| 0.00640592 |
| 0.00641929 |
| 0.00642269 |
| 0.00642807 |

| |
|---|
| 0.0064314 |
| 0.00643153 |
| 0.00643153 |
| 0.00643499 |
| 0.00643774 |
| 0.00644685 |
| 0.00644796 |
| 0.00645847 |
| 0.0064615 |
| 0.0064736 |
| 0.00647502 |
| 0.00648214 |
| 0.00649419 |
| 0.0064946 |
| 0.00651547 |
| 0.00651547 |
| 0.00651821 |
| 0.00651887 |
| 0.00651887 |
| 0.00655035 |
| 0.00655377 |
| 0.00657674 |
| 0.00659393 |
| 0.00659942 |
| 0.00660775 |
| 0.00661763 |
| 0.00665724 |
| 0.00665855 |
| 0.00666977 |
| 0.00667483 |
| 0.00671406 |
| 0.00671907 |
| 0.00672473 |
| 0.00673062 |
| 0.00675465 |
| 0.00675776 |
| 0.00678018 |
| 0.00678048 |
| 0.00681634 |
| 0.00681916 |
| 0.00681918 |
| 0.00682061 |
| 0.00682061 |
| 0.00683075 |
| 0.00687073 |

| |
|---|
| 0.0068719 |
| 0.00687887 |
| 0.00687887 |
| 0.0068791 |
| 0.0068791 |
| 0.00689193 |
| 0.00689594 |
| 0.00689595 |
| 0.00689595 |
| 0.00691079 |
| 0.00692012 |
| 0.00695062 |
| 0.00701269 |
| 0.00701329 |
| 0.00702335 |
| 0.00702459 |
| 0.00702566 |
| 0.0070274 |
| 0.00703601 |
| 0.00703817 |
| 0.00705995 |
| 0.00706016 |
| 0.00707622 |
| 0.00707622 |
| 0.00707703 |
| 0.00707703 |
| 0.00709384 |
| 0.00709451 |
| 0.00709451 |
| 0.00712361 |
| 0.0071286 |
| 0.00713759 |
| 0.00714999 |
| 0.00715505 |
| 0.00716033 |
| 0.00716565 |
| 0.00718684 |
| 0.00719782 |
| 0.00720865 |
| 0.00721812 |
| 0.00721969 |
| 0.00726489 |
| 0.00727074 |
| 0.00727268 |
| 0.00727476 |

| |
|---|
| 0.0072859 |
| 0.0072859 |
| 0.00730166 |
| 0.00730325 |
| 0.00731238 |
| 0.00732368 |
| 0.00732948 |
| 0.00733863 |
| 0.00735739 |
| 0.00736324 |
| 0.00741444 |
| 0.00742478 |
| 0.00742478 |
| 0.00743352 |
| 0.00743352 |
| 0.00743352 |
| 0.00743432 |
| 0.00744567 |
| 0.00746305 |
| 0.00748222 |
| 0.00751735 |
| 0.0075186 |
| 0.00752368 |
| 0.00754801 |
| 0.00756468 |
| 0.00758105 |
| 0.00758105 |
| 0.00759226 |
| 0.00759355 |
| 0.00761509 |
| 0.00762818 |
| 0.00763991 |
| 0.00764414 |
| 0.00768645 |
| 0.00768873 |
| 0.00769775 |
| 0.00769894 |
| 0.00770139 |
| 0.00770323 |
| 0.00771036 |
| 0.00771255 |
| 0.00773464 |
| 0.00773464 |
| 0.00774947 |
| 0.00775155 |

0.00775549
0.00775799
0.00776301
0.00776967
0.00776967
0.00776967
0.00778732
0.00779409
0.00780389
0.00781696
0.00782176
0.00782788
0.00784251
0.0078591
0.00786014
0.00787829
0.00788037
0.00789314
0.00789834
0.00790997
0.00794474
0.00794968
0.00794968
0.00796888
0.00796888
0.0079706
0.00798195
0.00801316
0.00802025
0.00802847
0.00803051
0.00804705
0.00805276
0.00806422
0.00808097
0.00810446
0.00812262
0.00812262
0.00813134
0.00814038
0.00815061
0.00819165
0.00819191
0.00820164
0.00821495

| |
|---|
| 0.00821562 |
| 0.00821896 |
| 0.00822533 |
| 0.00822533 |
| 0.00827715 |
| 0.00830637 |
| 0.00831777 |
| 0.00831777 |
| 0.00831941 |
| 0.00836288 |
| 0.00837178 |
| 0.00838362 |
| 0.00841728 |
| 0.00844172 |
| 0.00845146 |
| 0.00846325 |
| 0.008469 |
| 0.00847632 |
| 0.00847632 |
| 0.00848057 |
| 0.00848091 |
| 0.00852828 |
| 0.00855256 |
| 0.00855914 |
| 0.00856086 |
| 0.00856275 |
| 0.00857562 |
| 0.00858478 |
| 0.00859751 |
| 0.00860716 |
| 0.00860716 |
| 0.00860825 |
| 0.00860858 |
| 0.00861341 |
| 0.00862446 |
| 0.00862762 |
| 0.00863176 |
| 0.00865594 |
| 0.00865979 |
| 0.00868237 |
| 0.00870653 |
| 0.00871546 |
| 0.00871546 |
| 0.00873057 |
| 0.00873203 |

| |
|---|
| 0.00879072 |
| 0.00879072 |
| 0.00882257 |
| 0.00882257 |
| 0.00883252 |
| 0.00883584 |
| 0.00885796 |
| 0.00886823 |
| 0.00889463 |
| 0.00890237 |
| 0.00894297 |
| 0.00895983 |
| 0.00896116 |
| 0.00896342 |
| 0.00898787 |
| 0.00899367 |
| 0.00899919 |
| 0.00899919 |
| 0.00899919 |
| 0.00899919 |
| 0.00901134 |
| 0.00901345 |
| 0.00902963 |
| 0.00904681 |
| 0.00905345 |
| 0.00907752 |
| 0.00908493 |
| 0.00910459 |
| 0.00910523 |
| 0.00910737 |
| 0.00911955 |
| 0.00913963 |
| 0.00915311 |
| 0.0091571 |
| 0.00916702 |
| 0.00917425 |
| 0.00917425 |
| 0.00918677 |
| 0.00919124 |
| 0.00920063 |
| 0.00920063 |
| 0.00920852 |
| 0.00922189 |
| 0.00923791 |
| 0.00926195 |

0.00933521
0.00934973
0.00936713
0.00936926
0.00937589
0.00940946
0.00944519
0.00944519
0.00945399
0.00948222
0.00950238
0.00952018
0.00952018
0.00952513
0.00953567
0.0095435
0.00958354
0.00960095
0.00960095
0.0096119
0.00961289
0.0096281
0.0096333
0.00964823
0.00969456
0.00970963
0.00970964
0.00973136
0.00973941
0.00977634
0.00977956
0.00978586
0.00979215
0.0098039
0.0098039
0.00982348
0.00982529
0.00983603
0.00986382
0.00987814
0.00987814
0.00989279
0.00989927
0.00991783
0.00991783

| |
|---|
| 0.009949 |
| 0.00996449 |
| 0.00998694 |
| 0.00998921 |
| 0.00998921 |
| 0.01000473 |
| 0.01000659 |
| 0.01003809 |
| 0.01004204 |
| 0.0100532 |
| 0.01005372 |
| 0.01006019 |
| 0.01011698 |
| 0.01014627 |
| 0.01015432 |
| 0.01017147 |
| 0.01021853 |
| 0.01023284 |
| 0.01025678 |
| 0.01026484 |
| 0.0102706 |
| 0.01028651 |
| 0.01035926 |
| 0.01035926 |
| 0.01039774 |
| 0.01040468 |
| 0.01040527 |
| 0.01041328 |
| 0.01044022 |
| 0.01044225 |
| 0.01044225 |
