## Supplementary material for "CCL5 promotes breast cancer recurrence through macrophage recruitment in residual tumors": Figure 6 Source Data 2

| Entrez | Symbol | baseMean | log2FoldChai | lfcSE | stat | pvalue |
|---|---|---|---|---|---|---|
| 80797 | Clca3a2 | 2368.78067 | 9.16617641 | 0.75326057 | 12.1686662 | 4.56E-34 |
| 209195 | CLIC6 | 747.270404 | 8.65505976 | 0.83800005 | 10.328233 | 5.25E-25 |
| 100038347 | FAM174B | 262.759229 | 5.93800875 | 0.60562766 | 9.8047185 | 1.07E-22 |
| 12990 | CSN1S1 | 944.388978 | 8.53567108 | 0.91221661 | 9.35706607 | 8.20E-21 |
| 58860 | ADAMDEC1 | 182.557155 | 4.71281119 | 0.5575432 | 8.45281797 | 2.84E-17 |
| 15213 | HEY1 | 261.350239 | 5.23167918 | 0.64308832 | 8.13524205 | 4.11E-16 |
| 232345 | A2M | 178.832212 | 9.44403925 | 1.23713643 | 7.63378961 | 2.28E-14 |
| 229933 | CLCA2 | 89.5712528 | 5.93478831 | 0.78484652 | 7.56171829 | 3.98E-14 |
| 16005 | IGFALS | 37.0293544 | 5.11994004 | 0.69177777 | 7.40113408 | 1.35E-13 |
| 320007 | SIDT1 | 189.492385 | 4.44239726 | 0.60095159 | 7.39227139 | 1.44E-13 |
| 26366 | Ceacam10 | 109.30804 | 8.06758392 | 1.09877706 | 7.34233014 | 2.10E-13 |
| 70574 | CPM | 490.169865 | 5.33236163 | 0.73089117 | 7.29569855 | 2.97E-13 |
| 20496 | SLC12A2 | 1706.57295 | 2.59906512 | 0.36014541 | 7.21671042 | 5.33E-13 |
| 12319 | car8 | 97.0366342 | 8.94683505 | 1.24696784 | 7.17487234 | 7.24E-13 |
| 12671 | CHRM3 | 195.583395 | 6.84561516 | 0.96630977 | 7.0842864 | 1.40E-12 |
| 99571 | FGG | 111.654877 | 5.37759654 | 0.7691062 | 6.99200782 | 2.71E-12 |
| 12353 | Car6 | 130.841721 | 6.25502329 | 0.91024015 | 6.87183846 | 6.34E-12 |
| 16529 | KCNK5 | 428.314362 | 4.1424042 | 0.60906014 | 6.8013057 | 1.04E-11 |
| 57816 | TESC | 100.6796 | 9.78767519 | 1.44059625 | 6.79418346 | 1.09E-11 |
| 233099 | #N/A | 57.22855 | 9.48959628 | 1.40460355 | 6.75606743 | 1.42E-11 |
| 23945 | MGLL | 329.003615 | 2.0528727 | 0.30510746 | 6.72835948 | 1.72E-11 |
| 17304 | MFGE8 | 3632.69534 | 3.50717201 | 0.53840006 | 6.5140632 | 7.31E-11 |
| 58187 | CLDN10 | 57.6234542 | 4.91420782 | 0.7655862 | 6.41888241 | 1.37E-10 |
| 78896 | 1500015O10 | 192.263927 | 5.14430139 | 0.80244718 | 6.41076633 | 1.45E-10 |
| 100294583 | #N/A | 80.1042564 | 8.10937091 | 1.27339601 | 6.36830242 | 1.91E-10 |
| 216643 | GABRP | 445.798945 | 6.13759281 | 0.9662003 | 6.3522986 | 2.12E-10 |
| 50500 | TTPA | 66.8980295 | 6.00492124 | 0.96670651 | 6.21173145 | 5.24E-10 |
| 17436 | ME1 | 298.197044 | 3.36365539 | 0.54324192 | 6.19181849 | 5.95E-10 |
| 20203 | S100B | 97.3908138 | 4.49442379 | 0.74067455 | 6.06801432 | 1.30E-09 |
| 232413 | CLEC12A | 585.854091 | -2.4403832 | 0.40652357 | -6.0030547 | 1.94E-09 |
| 723792 | #N/A | 21.3812076 | 8.39596743 | 1.41026396 | 5.95347232 | 2.63E-09 |
| 22359 | VLDLR | 289.374943 | 4.07891972 | 0.69006299 | 5.91093827 | 3.40E-09 |
| 224116 | MUC20 | 19.7963379 | 7.73317229 | 1.32996515 | 5.81456761 | 6.08E-09 |
| 108079 | PRKAA2 | 84.4409816 | 4.08455306 | 0.70529231 | 5.79129105 | 6.98E-09 |
| 56175 | BACE2 | 49.0337856 | 4.49159631 | 0.77977797 | 5.76009645 | 8.41E-09 |
| 72432 | SPINK5 | 28.0142393 | 7.6964523 | 1.33978462 | 5.74454444 | 9.22E-09 |
| 12831 | COL5A1 | 994.825288 | -1.8313239 | 0.3203661 | -5.7163473 | 1.09E-08 |
| 12993 | #N/A | 739.48523 | 8.25386038 | 1.44803425 | 5.70004498 | 1.20E-08 |
| 26381 | ESRRG | 63.6470635 | 7.8605703 | 1.38166544 | 5.68919949 | 1.28E-08 |
| 328287 | Gm20554 | 118.519834 | 3.55370207 | 0.62711237 | 5.66677077 | 1.46E-08 |
| 330409 | CECR2 | 51.1990112 | 5.21914083 | 0.92197571 | 5.66082251 | 1.51E-08 |
| 12991 | CSN2 | 672.626471 | 8.6861747 | 1.53675518 | 5.6522827 | 1.58E-08 |
| 12992 | #N/A | 14.2587682 | 8.14940952 | 1.45537166 | 5.59953842 | 2.15E-08 |

| | | | | | | |
|---|---|---|---|---|---|---|
| 320782 | TMEM154 | 142.387526 | 2.89530516 | 0.51899311 | 5.57869669 | 2.42E-08 |
| 93882 | PCDHB11 | 18.7561816 | 4.05315194 | 0.73284722 | 5.53069156 | 3.19E-08 |
| 76683 | #N/A | 57.7843873 | 6.74589039 | 1.22817835 | 5.49259838 | 3.96E-08 |
| 269799 | Clec4a1 | 526.1571 | -1.922855 | 0.35342105 | -5.4406917 | 5.31E-08 |
| 329735 | 4933431E20I | 67.4076243 | 2.51535855 | 0.4625882 | 5.43757614 | 5.40E-08 |
| 26422 | NBEA | 591.539125 | 2.86033823 | 0.5295867 | 5.40107639 | 6.62E-08 |
| 16367 | IRS1 | 683.923243 | 2.33794629 | 0.43325759 | 5.39620386 | 6.81E-08 |
| 20277 | SCNN1B | 17.7014591 | 7.82360369 | 1.45120983 | 5.3910906 | 7.00E-08 |
| 16770 | LALBA | 16.5057924 | 6.30748577 | 1.17227702 | 5.38054202 | 7.43E-08 |
| 22239 | Ugt8a | 18.6722178 | 6.39467437 | 1.18917619 | 5.37739861 | 7.56E-08 |
| 171284 | #N/A | 29.0195946 | 6.64402539 | 1.2355534 | 5.37736806 | 7.56E-08 |
| 16564 | KIF21A | 286.334011 | 3.56488481 | 0.66645511 | 5.34902469 | 8.84E-08 |
| 12722 | Clca3a1 | 59.1651059 | 4.84907509 | 0.91243463 | 5.31443563 | 1.07E-07 |
| 70377 | DERL3 | 23.6811222 | 3.46363558 | 0.65190368 | 5.3131094 | 1.08E-07 |
| 12023 | barx2 | 94.4451678 | 4.55854434 | 0.85892562 | 5.30726321 | 1.11E-07 |
| 209558 | ENPP3 | 350.668848 | 4.44697656 | 0.84084037 | 5.28872866 | 1.23E-07 |
| 66695 | ASPN | 402.682241 | -4.6066199 | 0.87417838 | -5.2696566 | 1.37E-07 |
| 54140 | AVPR1A | 16.1786655 | 3.38730068 | 0.64617332 | 5.2420931 | 1.59E-07 |
| 217305 | CD300LD | 1046.88957 | -1.250716 | 0.23930304 | -5.2264945 | 1.73E-07 |
| 108096 | Slco1a5 | 114.783279 | 3.96530309 | 0.75973733 | 5.21930793 | 1.80E-07 |
| 14066 | F3 | 402.076859 | 3.3367218 | 0.63992481 | 5.21424042 | 1.85E-07 |
| 107527 | IL1RL2 | 192.309831 | 1.82352558 | 0.35189835 | 5.18196679 | 2.20E-07 |
| 16371 | IRX1 | 288.145274 | 3.21912089 | 0.62311463 | 5.16617771 | 2.39E-07 |
| 12669 | CHRM1 | 74.3630479 | 4.69789481 | 0.91983911 | 5.10730058 | 3.27E-07 |
| 216551 | LGALSL | 211.197385 | 2.06308061 | 0.40405899 | 5.10588966 | 3.29E-07 |
| 68075 | LURAP1 | 22.6607265 | 3.6093 | 0.7079624 | 5.09815208 | 3.43E-07 |
| 11537 | CFD | 80.7060125 | 6.81006182 | 1.33724005 | 5.09262477 | 3.53E-07 |
| 100503289 | #N/A | 19.9959998 | 7.57244709 | 1.49017947 | 5.08156719 | 3.74E-07 |
| 55990 | FMO2 | 32.4563784 | 3.62728566 | 0.71868042 | 5.04714691 | 4.48E-07 |
| 17341 | BHLHA15 | 48.9213386 | 5.05920275 | 1.00379934 | 5.04005386 | 4.65E-07 |
| 68588 | CTHRC1 | 83.4504447 | -4.7609566 | 0.9579793 | -4.9697907 | 6.70E-07 |
| 546144 | WDR72 | 41.9132146 | 4.42892259 | 0.89920559 | 4.92537262 | 8.42E-07 |
| 20296 | CCL2 | 1995.85544 | -1.2583054 | 0.26025844 | -4.8348305 | 1.33E-06 |
| 100740 | AI839979 | 94.7219078 | -2.5898877 | 0.53613843 | -4.8306326 | 1.36E-06 |
| 229722 | 5330417C22I | 95.0859216 | 4.5797209 | 0.94849622 | 4.82840184 | 1.38E-06 |
| 93961 | B3GALT5 | 338.044981 | 3.35219811 | 0.69601474 | 4.81627456 | 1.46E-06 |
| 209743 | AF529169 | 234.249464 | 2.97641378 | 0.61896716 | 4.80867802 | 1.52E-06 |
| 22373 | #N/A | 9.82263879 | 7.55312135 | 1.57530239 | 4.79471203 | 1.63E-06 |
| 13106 | CYP2E1 | 17.6814672 | 8.43088048 | 1.76598868 | 4.7740286 | 1.81E-06 |
| 14233 | FOXI1 | 132.938657 | 5.22492332 | 1.09889735 | 4.75469642 | 1.99E-06 |
| 12772 | CCR2 | 2404.51303 | -1.8830917 | 0.39711439 | -4.7419377 | 2.12E-06 |
| 13992 | KHDRBS3 | 78.3625197 | 2.79735049 | 0.59162262 | 4.72826834 | 2.26E-06 |
| 71520 | GRAP | 105.216227 | -1.4638729 | 0.31312478 | -4.6750465 | 2.94E-06 |
| 67393 | CXXC5 | 268.425277 | 1.71530855 | 0.37006816 | 4.63511516 | 3.57E-06 |
| 14807 | GRIK3 | 186.585146 | 4.99806967 | 1.07913106 | 4.63156872 | 3.63E-06 |

| | | | | | | |
|---|---|---|---|---|---|---|
| 107321 | LPXN | 776.035699 | -1.0873264 | 0.23495887 | -4.6277306 | 3.70E-06 |
| 67719 | #N/A | 284.047172 | 8.31997992 | 1.79816919 | 4.62691718 | 3.71E-06 |
| 71355 | COL24A1 | 15.7653167 | -3.80168 | 0.82238604 | -4.6227439 | 3.79E-06 |
| 100047282 | Gm20568 | 185.016398 | -0.8747139 | 0.18942674 | -4.6176898 | 3.88E-06 |
| 77018 | COL25A1 | 35.6470073 | 4.58660247 | 0.9940555 | 4.61403057 | 3.95E-06 |
| 215085 | #N/A | 23.572073 | 7.18223137 | 1.56678908 | 4.58404483 | 4.56E-06 |
| 98256 | KMO | 106.700271 | -1.5812628 | 0.34642356 | -4.5645361 | 5.01E-06 |
| 16494 | KCNA6 | 22.5738043 | 5.16086816 | 1.14560666 | 4.50492156 | 6.64E-06 |
| 78354 | 2210407C18I | 10.395509 | 4.80611179 | 1.07030253 | 4.49042366 | 7.11E-06 |
| 217082 | HLF | 121.834224 | 3.36354553 | 0.75241904 | 4.47030892 | 7.81E-06 |
| 20284 | SCRG1 | 14.4800167 | 5.848012 | 1.30799761 | 4.47096536 | 7.79E-06 |
| 21677 | TEAD2 | 115.180219 | 2.02990021 | 0.45481575 | 4.46312645 | 8.08E-06 |
| 67017 | FAM210B | 253.315157 | 1.29666134 | 0.2908843 | 4.45765322 | 8.29E-06 |
| 233274 | Siglech | 57.1250035 | 3.50390968 | 0.78648858 | 4.45513106 | 8.38E-06 |
| 360213 | TRIM46 | 30.7363155 | 3.38129639 | 0.76025691 | 4.44757074 | 8.68E-06 |
| 77569 | LIMCH1 | 143.578121 | 3.84970018 | 0.86750134 | 4.43768788 | 9.09E-06 |
| 330914 | ARHGAP32 | 280.592166 | 2.25205164 | 0.509948 | 4.41623783 | 1.00E-05 |
| 50706 | POSTN | 3638.86321 | -2.9268598 | 0.66393504 | -4.4083527 | 1.04E-05 |
| 68891 | CD177 | 157.954228 | 4.77965178 | 1.08483481 | 4.40587979 | 1.05E-05 |
| 26365 | CEACAM1 | 543.356992 | 2.40659468 | 0.54773952 | 4.39368459 | 1.11E-05 |
| 11554 | ADRB1 | 82.1955209 | 3.91973256 | 0.89247652 | 4.39197276 | 1.12E-05 |
| 58233 | DNAJA4 | 342.935513 | 2.25872233 | 0.51551557 | 4.38148231 | 1.18E-05 |
| 100043636 | AI662270 | 698.76105 | -0.7996475 | 0.18396926 | -4.3466364 | 1.38E-05 |
| 16858 | LGALS7 | 21.3719053 | 2.75721382 | 0.63833443 | 4.31938763 | 1.56E-05 |
| 22354 | VIPR1 | 548.619521 | 2.08657515 | 0.48557891 | 4.29708767 | 1.73E-05 |
| 80718 | RAB27B | 56.3560295 | 2.48294852 | 0.57810322 | 4.29499168 | 1.75E-05 |
| 11979 | #N/A | 10.806848 | 5.82892872 | 1.36080993 | 4.28342606 | 1.84E-05 |
| 18595 | PDGFRA | 828.789022 | -3.7997586 | 0.8875787 | -4.2810385 | 1.86E-05 |
| 70984 | 4931406C07I | 577.646208 | 1.49064797 | 0.34824654 | 4.28043869 | 1.87E-05 |
| 382253 | CDKL5 | 90.6794929 | 1.86910659 | 0.43925524 | 4.25517196 | 2.09E-05 |
| 217306 | CD300E | 76.6629366 | -2.12909 | 0.50129522 | -4.2471779 | 2.16E-05 |
| 12845 | COMP | 464.54086 | 3.39355715 | 0.80047979 | 4.23940389 | 2.24E-05 |
| 237387 | lrrc3 | 367.78001 | 4.25591746 | 1.00496586 | 4.23488761 | 2.29E-05 |
| 22433 | XBP1 | 3370.07023 | 1.04702364 | 0.24812961 | 4.21966423 | 2.45E-05 |
| 12350 | Car3 | 78.8084927 | 5.56235116 | 1.32123493 | 4.20996374 | 2.55E-05 |
| 381680 | Nxpe5 | 640.71147 | -1.4240219 | 0.33878405 | -4.2033323 | 2.63E-05 |
| 13081 | cyp24a1 | 71.1261629 | 6.72387214 | 1.60260169 | 4.1955978 | 2.72E-05 |
| 64214 | RGS18 | 109.795216 | -2.085592 | 0.49871367 | -4.1819428 | 2.89E-05 |
| 78754 | GALNT15 | 66.1341145 | 3.28134428 | 0.78501487 | 4.17997722 | 2.92E-05 |
| 319259 | BRICD5 | 25.0336974 | 4.74008232 | 1.13391459 | 4.1802816 | 2.91E-05 |
| 18542 | PCOLCE | 162.192245 | -2.2837637 | 0.54891871 | -4.160477 | 3.18E-05 |
| 74053 | #N/A | 21.7948475 | 5.76711501 | 1.38951744 | 4.1504445 | 3.32E-05 |
| 77669 | #N/A | 32.1917289 | 4.26851725 | 1.03118674 | 4.13942216 | 3.48E-05 |
| 71884 | CHIT1 | 28.2953027 | 5.96296979 | 1.44556635 | 4.12500595 | 3.71E-05 |
| 242594 | 1700024P16I | 13.193547 | 3.62769956 | 0.87966099 | 4.12397457 | 3.72E-05 |

| | | | | | | |
|---|---|---|---|---|---|---|
| 71069 | STOX2 | 254.455906 | 2.12514424 | 0.51705792 | 4.11006998 | 3.96E-05 |
| 66255 | Hsbp1l1 | 12.0132068 | 5.5123299 | 1.34371577 | 4.10230349 | 4.09E-05 |
| 74511 | LRRC17 | 50.3179334 | -3.785413 | 0.92324617 | -4.1001123 | 4.13E-05 |
| 14245 | LPIN1 | 659.352265 | 1.78986112 | 0.43661792 | 4.09937623 | 4.14E-05 |
| 66198 | THEM5 | 5.45884195 | 6.61199118 | 1.62181416 | 4.07691049 | 4.56E-05 |
| 12578 | CDKN2A | 28.4854427 | 3.61964904 | 0.88964595 | 4.06863994 | 4.73E-05 |
| 77781 | EPM2AIP1 | 740.333681 | 0.85123225 | 0.20931023 | 4.06684497 | 4.77E-05 |
| 72169 | TRIM29 | 27.419801 | 3.22519218 | 0.79362919 | 4.06385279 | 4.83E-05 |
| 99439 | DUOX1 | 154.038144 | 4.64055369 | 1.15221802 | 4.02749619 | 5.64E-05 |
| 76491 | ABHD14B | 52.511876 | 2.02627319 | 0.50348035 | 4.02453281 | 5.71E-05 |
| 230903 | FBXO44 | 70.9563203 | 1.20285543 | 0.30012885 | 4.00779674 | 6.13E-05 |
| 67856 | ECHDC3 | 86.086158 | 2.96220517 | 0.74062161 | 3.99962025 | 6.34E-05 |
| 100041546 | Ly6c2 | 57.5171471 | -2.6448033 | 0.66222234 | -3.9938298 | 6.50E-05 |
| 14804 | #N/A | 5.58288357 | 5.84149547 | 1.46543738 | 3.98617884 | 6.71E-05 |
| 76905 | LRG1 | 140.923048 | 2.24196391 | 0.562744 | 3.98398542 | 6.78E-05 |
| 210146 | IRGQ | 637.8019 | 1.40228205 | 0.3526831 | 3.97603985 | 7.01E-05 |
| 68867 | RNF122 | 236.261884 | 1.56554366 | 0.39430903 | 3.97034692 | 7.18E-05 |
| 20352 | SEMA4B | 516.902233 | 1.82461323 | 0.46072608 | 3.96029941 | 7.49E-05 |
| 100038573 | #N/A | 143.210751 | 4.24013588 | 1.07648449 | 3.93887317 | 8.19E-05 |
| 13869 | #N/A | 24.8331633 | 4.88483917 | 1.24360918 | 3.92795362 | 8.57E-05 |
| 16160 | IL12B | 274.547591 | 3.31783826 | 0.84555874 | 3.92384124 | 8.71E-05 |
| 217379 | UBXN2A | 365.591743 | 0.88226835 | 0.22511336 | 3.91921807 | 8.88E-05 |
| 100503073 | #N/A | 30.2454703 | 3.2374958 | 0.82964585 | 3.90226239 | 9.53E-05 |
| 330723 | HTRA4 | 219.398118 | 1.58539343 | 0.4087488 | 3.87864971 | 0.00010504 |
| 330119 | ADAMTS3 | 67.951456 | 4.32030828 | 1.1170916 | 3.86746107 | 0.00010997 |
| 277154 | NYNRIN | 135.561335 | -1.4488553 | 0.37554556 | -3.8580015 | 0.00011432 |
| 99633 | ADGRL2 | 580.770625 | 1.22278062 | 0.31819476 | 3.84286851 | 0.0001216 |
| 215798 | ADGRG6 | 424.161937 | 1.84895267 | 0.48384378 | 3.82138357 | 0.00013271 |
| 320910 | ITGB8 | 266.204173 | 2.42891576 | 0.63692282 | 3.8135166 | 0.000137 |
| 11450 | ADIPOQ | 14.7207604 | 5.80654437 | 1.5255876 | 3.80610355 | 0.00014117 |
| 381485 | TRIM55 | 35.1568762 | 3.2832876 | 0.86508384 | 3.79534034 | 0.00014744 |
| 18968 | POLA1 | 845.701641 | -0.6019434 | 0.15876502 | -3.7914106 | 0.00014979 |
| 18636 | CFP | 2182.93983 | -1.980512 | 0.52284125 | -3.7879796 | 0.00015188 |
| 72029 | CNPY3 | 1137.1161 | -0.6221614 | 0.16438939 | -3.7846814 | 0.00015391 |
| 100862305 | #N/A | 16.6237514 | -2.9145229 | 0.77123824 | -3.7790177 | 0.00015745 |
| 73652 | 2210408F21I | 39.5453265 | 1.82087017 | 0.48243055 | 3.77436745 | 0.00016041 |
| 67412 | #N/A | 11.1532626 | 3.32649543 | 0.88184843 | 3.77218503 | 0.00016182 |
| 66957 | SERPINB11 | 10.7454574 | 6.56358664 | 1.74308313 | 3.76550407 | 0.00016621 |
| 13449 | dok2 | 231.802077 | -1.623857 | 0.43168863 | -3.7616395 | 0.0001688 |
| 450219 | #N/A | 4.69104947 | 6.05781643 | 1.6125361 | 3.75670128 | 0.00017217 |
| 56177 | OLFM1 | 319.828862 | -1.2671353 | 0.3374298 | -3.7552561 | 0.00017316 |
| 319973 | A630077J23I | 12.4786383 | 4.5634428 | 1.21566593 | 3.75386255 | 0.00017413 |
| 13800 | ENAH | 170.557277 | 1.97415829 | 0.52721292 | 3.74451801 | 0.00018074 |
| 245404 | DCAF12L1 | 13.6299584 | 4.96774164 | 1.32791193 | 3.74101739 | 0.00018328 |
| 12738 | CLDN2 | 19.5514418 | 4.52369762 | 1.21472141 | 3.72406182 | 0.00019604 |

| | | | | | | |
|---|---|---|---|---|---|---|
| 12509 | Cd59a | 84.1857315 | 1.44422687 | 0.38791527 | 3.7230472 | 0.00019683 |
| 20965 | SYN2 | 6.7110815 | 5.50802957 | 1.47935854 | 3.72325533 | 0.00019667 |
| 13858 | EPS15 | 2529.89045 | -0.9620751 | 0.25880897 | -3.7173174 | 0.00020135 |
| 93689 | LMOD1 | 56.8144228 | 3.27323791 | 0.88190731 | 3.71154416 | 0.000206 |
| 241452 | DHRS9 | 84.1479988 | -2.0303482 | 0.54717873 | -3.7105759 | 0.00020679 |
| 239652 | #N/A | 15.9603034 | 4.1427529 | 1.11685254 | 3.70931054 | 0.00020782 |
| 76157 | #N/A | 10.7692742 | 5.41050667 | 1.45970935 | 3.70656437 | 0.00021009 |
| 14734 | GPC3 | 42.6526352 | 2.46494031 | 0.66537174 | 3.70460628 | 0.00021172 |
| 20312 | CX3CL1 | 207.765633 | 1.76006492 | 0.47540131 | 3.70227189 | 0.00021368 |
| 27027 | TSPAN32 | 187.844018 | -1.1976496 | 0.32399932 | -3.696457 | 0.00021863 |
| 260299 | CADM4 | 93.6713431 | 2.49701509 | 0.67556265 | 3.6962006 | 0.00021885 |
| 50781 | DKK3 | 94.4660588 | -2.9165517 | 0.79291817 | -3.6782506 | 0.00023484 |
| 233332 | ADAMTS17 | 10.6461231 | 3.50660346 | 0.95388647 | 3.67612246 | 0.00023681 |
| 80891 | Fcrls | 6219.86778 | -1.8011409 | 0.49143076 | -3.6650959 | 0.00024725 |
| 75320 | ETNK1 | 3750.70114 | 1.3420301 | 0.3661926 | 3.6648204 | 0.00024751 |
| 66857 | PLBD1 | 1288.41669 | -1.2887429 | 0.35202406 | -3.6609512 | 0.00025128 |
| 14133 | Fcna | 261.994713 | 2.16172003 | 0.59107702 | 3.65725609 | 0.00025493 |
| 636104 | #N/A | 7.57022619 | 5.58826099 | 1.52996307 | 3.65254632 | 0.00025965 |
| 327900 | UBTD2 | 309.53547 | 0.93750339 | 0.25860695 | 3.62520575 | 0.00028873 |
| 20304 | CCL5 | 381.275483 | 1.97503036 | 0.54539066 | 3.62131315 | 0.00029311 |
| 21834 | THRB | 72.0175399 | 2.7832411 | 0.76982458 | 3.6154225 | 0.00029986 |
| 320440 | #N/A | 69.9880188 | 2.26714721 | 0.62792292 | 3.6105502 | 0.00030555 |
| 229898 | GBP5 | 637.864995 | 1.37564916 | 0.3814933 | 3.60595892 | 0.000311 |
| 63954 | RBP7 | 120.503871 | 5.70611913 | 1.58577862 | 3.59830752 | 0.00032029 |
| 268807 | KLHL38 | 52.70968 | 2.18491313 | 0.60764765 | 3.59569093 | 0.00032353 |
| 16852 | LGALS1 | 4734.56173 | -2.2306104 | 0.6209147 | -3.5924586 | 0.00032757 |
| 226245 | PLEKHS1 | 21.7339638 | 3.14577307 | 0.87982165 | 3.57546677 | 0.0003496 |
| 21828 | THBS4 | 217.510051 | -2.4073829 | 0.67353566 | -3.5742471 | 0.00035124 |
| 63959 | SLC29A1 | 1841.96697 | 1.82215855 | 0.50982325 | 3.57409854 | 0.00035144 |
| 13602 | SPARCL1 | 1119.53298 | 2.07707288 | 0.58478642 | 3.55184869 | 0.00038253 |
| 26557 | HOMER2 | 156.084763 | 2.35926146 | 0.6660598 | 3.54211658 | 0.00039693 |

| padj |
| --- |
| 2.30E-30 |
| 1.98E-21 |
| 2.71E-19 |
| 1.61E-17 |
| 3.07E-14 |
| 3.45E-13 |
| 1.23E-11 |
| 1.94E-11 |
| 6.37E-11 |
| 6.61E-11 |
| 9.06E-11 |
| 1.25E-10 |
| 2.12E-10 |
| 2.70E-10 |
| 4.91E-10 |
| 8.73E-10 |
| 1.92E-09 |
| 3.01E-09 |
| 3.11E-09 |
| 3.89E-09 |
| 4.63E-09 |
| 1.63E-08 |
| 2.96E-08 |
| 3.08E-08 |
| 3.90E-08 |
| 4.22E-08 |
| 9.54E-08 |
| 1.07E-07 |
| 2.20E-07 |
| 3.11E-07 |
| 3.97E-07 |
| 5.09E-07 |
| 8.75E-07 |
| 9.86E-07 |
| 1.17E-06 |
| 1.27E-06 |
| 1.44E-06 |
| 1.56E-06 |
| 1.65E-06 |
| 1.86E-06 |
| 1.91E-06 |
| 1.99E-06 |
| 2.60E-06 |

| |
|---|
| 2.91E-06 |
| 3.74E-06 |
| 4.57E-06 |
| 5.90E-06 |
| 5.96E-06 |
| 7.10E-06 |
| 7.24E-06 |
| 7.40E-06 |
| 7.79E-06 |
| 7.82E-06 |
| 7.82E-06 |
| 9.03E-06 |
| 1.07E-05 |
| 1.07E-05 |
| 1.08E-05 |
| 1.18E-05 |
| 1.27E-05 |
| 1.47E-05 |
| 1.57E-05 |
| 1.62E-05 |
| 1.66E-05 |
| 1.94E-05 |
| 2.10E-05 |
| 2.82E-05 |
| 2.83E-05 |
| 2.93E-05 |
| 3.00E-05 |
| 3.13E-05 |
| 3.66E-05 |
| 3.78E-05 |
| 5.22E-05 |
| 6.46E-05 |
| 9.87E-05 |
| 0.0001003 |
| 0.00010094 |
| 0.00010573 |
| 0.00010867 |
| 0.00011448 |
| 0.00012457 |
| 0.00013648 |
| 0.00014471 |
| 0.00015138 |
| 0.00018578 |
| 0.00022089 |
| 0.00022379 |

0.00022702
0.00022702
0.00022977
0.00023356
0.00023677
0.00026915
0.00029201
0.00037686
0.00039922
0.00043225
0.00043225
0.00044537
0.00045358
0.00045729
0.0004658
0.00048435
0.00052531
0.00053892
0.0005414
0.00056631
0.00056757
0.00059362
0.0006781
0.00074569
0.0008145
0.00081714
0.00084769
0.00085395
0.00085395
0.00093925
0.0009705
0.00099294
0.0010072
0.00107143
0.00111203
0.00113532
0.0011678
0.00122315
0.0012269
0.0012269
0.00132859
0.00136975
0.00142944
0.00150968
0.00151238

| |
|---|
| 0.0015809 |
| 0.00163062 |
| 0.0016418 |
| 0.00164271 |
| 0.00178627 |
| 0.00184132 |
| 0.00185079 |
| 0.00186317 |
| 0.00210296 |
| 0.00212437 |
| 0.00225838 |
| 0.00232086 |
| 0.00236684 |
| 0.00243856 |
| 0.00245529 |
| 0.0025206 |
| 0.00256936 |
| 0.00264853 |
| 0.00286965 |
| 0.00297547 |
| 0.0030198 |
| 0.00306426 |
| 0.00327218 |
| 0.0035581 |
| 0.00370043 |
| 0.00383803 |
| 0.00405564 |
| 0.00436799 |
| 0.0044899 |
| 0.00461655 |
| 0.00479041 |
| 0.00485641 |
| 0.00491028 |
| 0.00492628 |
| 0.00502902 |
| 0.00510218 |
| 0.00512545 |
| 0.00520986 |
| 0.00526917 |
| 0.00535208 |
| 0.005361 |
| 0.0053689 |
| 0.00550529 |
| 0.00553789 |
| 0.0058883 |

0.00588861
0.00588861
0.00598816
0.00609047
0.00610187
0.00612049
0.00616492
0.00618695
0.00623212
0.00635844
0.00635844
0.00676666
0.00678894
0.00702897
0.00702897
0.0071226
0.0072125
0.0073324
0.00806313
0.00817035
0.00832768
0.00843916
0.00857413
0.00879821
0.00887099
0.00894931
0.00949966
0.00951524
0.00951524
0.01030185
0.01059509
